# Long-term stability of RNA nucleoside standards for accurate LC-MS quantification

**DOI:** 10.1101/2025.07.30.667630

**Authors:** Kira Kerkhoff, Hagen Wesseling, Yuyang Qi, Sofia Obersteiner, Kuangjie Liu, Maximilian Berg, Leona Rusling, Hendrik Zipse, Stefanie Kaiser

## Abstract

Accurate LC-MS (liquid chromatography coupled mass spectrometry) analysis of RNA modifications relies on synthetic nucleoside standards whose chemical integrity determines both qualitative identification and quantitative measurements. While the purity of these standards is typically verified prior to use, their long-term chemical stability during storage has not been systematically investigated.

Here, we evaluated the stability of 44 canonical and modified ribonucleosides in aqueous solution during storage at -80 °C and -20 °C. Initial quality control confirmed the identity of all tested standards and revealed purity issues in selected compounds, including contamination of 1-methyladenosine (m^1^A) with 6-methyladenosine (m^6^A) and the presence of S- and R-isomers of 5-(carboxyhydroxymethyl)-2′-O-methyluridine (mchm^5^Um).

Long-term LC-UV-MS monitoring over 12 months showed that 30 nucleosides remained stable, two were stable for at least six months, and 12 exhibited substantial quantitative changes. Seven nucleosides formed detectable degradation products, most of which could be structurally assigned. Quantum-chemical calculations of reaction free energies for deglycosylation, deamination, deacetylation and desulfurization correlated with the experimentally observed stability trends.

Based on these results, we propose a practical guideline for the preparation, storage and analytical quality control of nucleoside standards, including recommendations for purity verification by UV spectroscopy and quantitative NMR. These guidelines provide an experimental framework to improve the robustness and inter-laboratory comparability of LC-MS-based RNA modification analysis.

## Introduction

The four letter-code of RNA is expanded in all living organisms by chemical modifications of the major building blocks cytidine (C), uridine (U), guanosine (G) and adenosine (A). Roughly 170 chemical modifications have been reported and collected in the major database MODOMICS so far (1). Some of these modifications are available as synthetic standards from either commercial sources (Table S1), through chemical synthesis in academic laboratories (2–4) or purification from biological materials (5,6). These standards are commonly used by mass spectrometrists in their efforts to (i) unambiguously identify an RNA modification from a biological sample or (ii) quantify its absolute abundance within the sample. For this purpose, the standards are typically dissolved in water and diluted to an ideal working concentration of 10 mM.

To validate the chemical identity of the synthetic standards, NMR and high-resolution mass spectrometry (HRMS) are the methods of choice. Analytical data are collected in the Pub-Chem database (NIH,(7)). Both technologies give a first indication of contamination with other nucleosides. Regarding purity, HPLC coupled to UV and/or MS detection gives a valuable orientation regarding the presence of contaminants or isomers of a compound. Here, relative retention times for various RNA modifications are collected at MODOMICS (1). An even simpler possibility is the use of 2D-thin-layer chromatography using fluorescent cellulose plates followed by UV-detection or radioactivity (8).

If the extinction coefficient (ε) of the nucleoside is known, it is possible to confirm the concentration of the standard solution by using a simple (calibrated) photometer. Currently, the extinction coefficient of ∼40 modified nucleosides are known (9) and have been summarized in Table S2. However, this approach requires an analyte solution free of other compounds that absorb light at the specified wavelength.

Typically, aqueous standard solutions of nucleosides are prepared in batches and stored at - 20 °C or -80 °C for extended periods. According to the ICH guideline M10 on bioanalytical method validation, the shelf-life of reference standards must be established to ensure reliable quantification. However, this is not common practice in the field of modified nucleosides and very little is known about how these aqueous stocks change in quality and quantity over time and what their shelf-life is. The need to have this information is emphasized by studies that reported chemical instabilities of modified nucleosides in the past. For 7-10 modifications a shelf-life of aqueous stock solutions is reported and summarized in Table 1. While chemical instability causes the formation of impurities and thus favours mis-identifications, it also reduces the quantity of the target nucleoside in solution. This introduces a significant bias into both relative and absolute quantitative analysis. Relative quantification is affected if samples, not measured on the same day, are compared. Absolute quantification, namely with the goal to provide absolute modification number per purified transcript, relies on the use of external calibration using synthetic standards of known concentration. These standards are serially diluted and analyzed before or after analysis of the biological samples. If a standard is unstable, the stock concentration decreases and thus a lower signal arises. Yet, the analyst assumes that the signal corresponds to the stock concentration and thus the analyte in the sample suffers from over-quantification. This means that the experimentally determined value is higher than the true value in the sample. This bias is unknown to the analyst and thus inaccurate data may be received which might affect biological interpretation.

**Table 1.** Known chemical instabilities of ribonucleosides in aqueous solution.

|  |  |  |
| --- | --- | --- |
| **A** known shelf-lives of modified nucleoside chemical standards |  |  |
| (Modified) ribonucleoside | Shelf life at -20 °C | product |
| m<sup>1</sup>A(m) (10) | 3 months (11) | formation of m<sup>6</sup>A(m) (dimroth rearrangement) |
| m<sup>3</sup>C (11) | 3 months (11) | formation of m<sup>3</sup>U |
| **B** Known instabilities without defined shelf-life |  |  |
| A (12) | Chemical deamination of adenosine to inosine formation |  |
| s<sup>4</sup>U (13) | *In-vitro* oxidation using 1 mM iodine leads to s<sup>4</sup>U dimer formation |  |
| s<sup>4</sup>U (14) | Formation of cytidine in alkaline hydrolysis buffer |  |
| x<sup>5</sup>s<sup>2</sup>U (14) | In ammonium bicarbonate buffer, formation of (5-modified)-isocytidine |  |
| ct<sup>6</sup>A (15,16) | Ring opening and formation of t<sup>6</sup>A, epimerization between hydantoin form (natural) and oxazolone form (potential intermediate in ct<sup>6</sup>A biogenesis) |  |
| ct<sup>6</sup>A (17) | Esterification and glycine addition |  |

To address the critical but largely unexplored question of how long synthetic nucleoside standards remain analytically reliable, we evaluated the aqueous stability of 44 canonical and modified nucleosides. Using LC-UV-MS we monitored their abundance after storage for up to 12 months at -80 °C and -20 °C in aqueous solutions. In addition, we monitored the stability at elevated temperatures for 3 months to assess short-term thermal robustness and facilitate the identification of degradation products. 30 out of 44 nucleosides were stable in aqueous solution at either -20 °C or -80 °C for 12 months, two for 6-12 months while 12 showed a quantity affecting behaviour. Prominent examples include desulfurization and dimer formation of 4-thiouridine (s^4^U), deamination of 3-methylcytidine (m^3^C) to 3-methyluridine (m^3^U), and stepwise hydrolysis and deamination of N -acetylcytidine (ac^4^C) (only at elevated temperature, not at -20 or -80 °C). The wobble nucleoside 5-methoxycarbonylmethyl-2-thiouridine (mcm s²U) underwent partial desulfurization, and N6-isopentenyladenosine (i A) displayed low-level de-prenylation. Quantum-chemical calculations of solution-phase reaction free energies for deglycosylation, deamination, deacetylation and desulfurization correlated with the experimental trends, providing a predictive framework for untested modifications in aqueous conditions. Additionally, we monitored stability in dimethyl sulfoxide (DMSO) for 3 months to explore alternative storage conditions. While this timeframe did not exceed the aqueous shelf-life of all tested compounds, we successfully demonstrated a clear protective effect of DMSO for N2-methylguanosine (m^2^G), s^4^U, and mcm^5^s^2^U. Together with this mitigation strategy, we provide a guideline for the preparation, quality control and storage of nucleoside solutions. The resulting standard operating procedure and practical handling recommendations will facilitate robust, inter-laboratory-comparable quantification of RNA modifications in the hands of RNA biologists and analytical chemists confronted with this sample class.

## Materials and Methods

### List of chemicals and reagents

All chemicals and reagents were acquired from Merck Chemicals (Darmstadt, Germany) unless stated otherwise.

### Quantitative NMR spectroscopy of standards

Nucleosides and internal calibrants were weighed in a suitable sample tube using a Ultramicrobalance XPR2U (Mettler Toledo, Columbus, OH, USA). Both components were dissolved in deuterated solvent and vortexed for 1 min. If dissolution was incomplete, the mixture was additionally sonicated for 15 min. Subsequently, the solution was transferred to Boroeco-5-7 NMR tubes (Deutero GmbH, Kastellaun, Germany) and immediately analyzed using a Bruker AVANCE III spectrometer (800 MHz) equipped with a 5 mm TXO cryoprobe head with z-gradient. The instrument was controlled using TopSpin version 3.6.3. Shimming (TopShim tool ‘ts’) and probe head tuning and matching (‘atma’) was applied before each measurement. Key experimental processing and acquisition parameters are listed as follows: nucleus = ^1^H, number of scans (NS) = 4, receiver gain (RG) = determined before each measurement (‘rga’), relaxation delay (D1) = 63 s, pulse width (P1) = determined before each measurement (‘pulsecal’), acquisition time (AQ) = 7.7 s, spectral width (SW) = 17.0 ppm, time domain (TD) = 128000, spectrum size (SI) = 132 000, apodization function = [em], line broadening factor = 0.2 Hz, transmitter offset (O1P) = 4.5 ppm.

### Data analysis of quantitative NMR

NMR raw data were processed and analyzed using MestReLab-Software MNova (version 14.1.2). The phase of each spectrum was adjusted manually and baseline corrections were done using the software’s polynomial fit algorithm. Integration limits of each signal were set to 0.05 ppm from the ^13^C satellite signals. The purity of the analyte/nucleoside was calculated using the following equation:

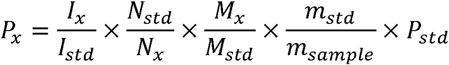

x = analyte

std = internal calibrant

P = purity (%)

I = integral value

N = number of spins

M = molecular weight (g/mol)

m = weighing amount (g)

For calculations and graph generations, Microsoft Excel and GraphPad Prism software (version 10.4.1) were used.

### Weighing nucleoside standards

Nucleoside aliquots between 0.5 mg and 3.0 mg (vendor information in Table S3, exact masses in Table S4) were dispensed into disposable polypropylene weigh boats (PP61.1, Carl Roth, Karlsruhe, Germany) on the same analytical balance as used for NMR analysis. Each aliquot was then transferred to a 1.5 mL polypropylene vial (cat. no. 525-0651, VWR International, Darmstadt, Germany) for subsequent dissolution. The weigh boat was reweighed after transfer, and the net difference was recorded as the precise mass of nucleoside in the vial. Every nucleoside was weighed in triplicate. The spatula was rinsed with ultrapure water and dried with lint-free tissue between samples to prevent cross-contamination.

### Dilution and preparation before analysis – aqueous solutions

Ultrapure water produced by a Milli-Q Integral system (Merck Millipore, Darmstadt, Germany) was added to each vial to obtain a final nucleoside concentration of 10 mM (Table S4). The contents were vortex-mixed until fully dissolved. For the subsequent stability assay, 20 µL of each 10 mM stock were used to prepare a serial dilution as follows.

1. **First dilution (1 : 10).** 20 µl of the 10 mM stock were pipetted into a 96-well polypropylene plate, followed by 180 µL ultrapure water. The solution was mixed by aspirating and dispensing ten times.
2. **Second dilution (1 : 100).** 20 µl of the 1 : 10 dilution were transferred to a new 96-well plate and combined with 180 µL water, using the same mixing procedure.
3. **Third dilution and master plate preparation (1 : 1000).** 150 µl of the 1 : 100 solution were added to 1350 µL water in a 96 deep-well plate (cat. no. AXYP2MLSQCS, Axygen/Corning, supplied by Sigma-Aldrich, Darmstadt, Germany). After thorough mixing, this deep-well plate served as the master plate for all storage conditions.
4. **Aliquoting for storage.** Using an eight-channel pipette set to 100 µL, 30 µL from each well of the master plate were dispensed into individual wells of a standard 96-well plate (part no. 5043-9311, Agilent InfinityLab, Denmark) allocated to the analyti-cal time point. Plates were sealed with aluminium foil (cat. no. Z721549, Excel Scientific, supplied by Sigma-Aldrich, Darmstadt, Germany), labelled with the designated storage temperature and sampling interval, and immediately placed under the specified conditions.
5. **Analysis plate preparation.** At the respective analysis time point, the plate was brought to room temperature and spun down. The aluminium foil was then removed and replaced with a silicone sealing mat (part no. 5042-1389, Agilent InfinityLab, Denmark).
6. **Start point analysis.** The first plate was immediately analyzed by LC-UV-MS.

### Dilution and preparation before analysis – DMSO solutions

DMSO (Art. No.: 00905-50mL, Deutero, Kastellaun, Germany) was added to each vial to obtain a final nucleoside concentration of 10 mM. The contents were vortexed until fully dissolved. The solutions were stored at -20 °C in glass vials with screw caps (Art. No. 702282 and 702287.1, Macherey-Nagel, Düren, Germany) and thawed immediately prior to the subsequent stability assay. 5 µL of each 10 mM thawed stock were used to prepare a serial dilution immediately as follows.

1. **First dilution (1 : 10).** The 5 µl aliquot was diluted with 45 µl of ultrapure water in 1.5 mL reaction tubes. The solution was mixed by vortexing.
2. **Second dilution (1 : 100).** 20 µL of the 1 : 10 dilution were diluted with 180 µL ultrapure water, using the same mixing procedure.
3. **Third dilution (1 : 1000).** 20 µL of the 1 : 100 dilution were diluted with 180 µL ultrapure water, using the same mixing procedure.
4. **Analysis plate preparation.** 30 µL of the third dilution were dispensed into individual wells of a standard 96-well plate, which was then sealed with a silicone sealing mat.
5. **Start point analysis.** Immediately after dissolving the nucleosides, the first serial dilution was performed. The resulting analysis plate was then analyzed by LC-UV-MS.

### Liquid chromatography coupled mass spectrometry

LC-UV-MS experiments were carried out using an Agilent 1290 Infinity II system (Agilent Technologies, Santa Clara, CA, USA), equipped with a diode array detector (DAD) for UV detection and coupled to an Agilent 6470 triple quadrupole mass spectrometer with an Agilent Jet Stream ESI ion source. The diode array detector (DAD) wavelengths were set to 230, 254, and 320 nm for the quantification and identification of potential degradation products. The specific wavelength used for the quantification of each nucleoside is provided in Table S7. 10 µL of each 10 µM nucleoside (100 pmol per inject) solution was injected into the system. Chromatographic separation was performed on a Phenomenex Synergy Fusion-RP column (2.5 µm, 100 Å, 100 × 2 mm; Phenomenex, Torrance, CA, USA) at 35 °C with a flow rate of 350 µL/min. Buffer A consisted of 5 mM ammonium acetate (adjusted to pH 5.3 with glacial acetic acid) and buffer B was Ultra LC-MS grade acetonitrile. The gradient began with 100% solvent A for 1 min, followed by a linear increase to 10% solvent B over 3 min. From 3 to 5 min, solvent B was increased to 40% and held for 1 min, then returned to 100% solvent A over 2 min, followed by a 1 min re-equilibration phase. The mass spectrometer was operated in MS2 scan mode over an *m/z* range of 100 to 600. Data acquisition was done in positive ion mode, with fragmentor voltage set to 90 V and cell acceleration voltage of 5 V.

### Analysis of LC-UV-MS data

UV peak areas were integrated, and MS2 scans were analyzed using MassHunter Qualitative Analysis Navigator (version B.08.00). Subsequently, analyte recovery was calculated in Microsoft Excel by normalizing the peak area of each nucleoside to its initial value at time point zero, expressed as a percentage. Data were then exported to GraphPad Prism (version 10.6.1) for linear regression analysis, with the y-intercept constrained to 100% at x = 0, and the 95% confidence interval (CI) was computed. The Estimated Shelf Life (Est. SL) was determined as the point of intersection between the 95% CI and the predefined acceptance criteria of 95–105% recovery.

### HRAMS analysis

To further investigate nucleosides with observed degradation products, high resolution accurate mass (HRAM) measurements were performed on a Q Exactive^TM^ Plus (Thermo Fisher Scientific, Dreieich, Germany). Thermally stressed nucleoside solutions of ms^2^i^6^A and mcm^5^s^2^U and non-stressed nucleoside solution of mcm^5^U for comparison, 10 mM each, were diluted to 5 µM in 500:500:1 water/methanol/formic acid and measured on the Q Exactive by direct infusion at 5 µL/min into the heated electrospray ionization (H-ESI) source in positive ion mode with a capillary temperature of 320 °C, sheath gas of 10 (nominal units), spray voltage of 4 kV, and an S-lens RF level of 55. Full MS1, MS2 and pseudo-MS3 scans were acquired for at least 1 min with an AGC target of 1e5 and a maximum injection time of 30 ms as manual acquisitions in the Thermo Tune software (version 2.9 Build 2926). Thermo Xcalibur Qual Browser (Thermo Xcalibur v.4.1.31.9) was used to obtain extracted ion chromatograms for each acquisition.

### Quantum chemical calculations

Conformational sampling for all systems was performed with CREST 2.12 (18,19) using the GFN2-xTB (20) method. The structures of all conformers (including those of the most stable tautomers) were subsequently reoptimized at the B3LYP-D3/def2-TZVPP (21–23) level of theory, followed by single point calculations at the SMD(H_2_O)/B3LYP-D3/def2-TZVPP (24) level using (25). Refined energies have subsequently been obtained using the DLPNO-CCSD(T)/CBS (26,27) scheme implemented in ORCA 6.0.0 in combination with the cc-pVTZ and cc-pVQZ basis sets. (28) The DLPNO-CCSD(T)/CBS energies were then combined with solvation free energies in water and thermochemical corrections obtained at B3LYP-D3/def2-TZVPP level to yield final free energies in water at 298.15 K (ΔG_298_) referenced to a standard state of 1 mol/L.

## Results

### Identity and purity assessment of commercial nucleoside standards

To verify the identity and purity of commercially available nucleoside standards, we purchased four canonical nucleosides and 41 modified nucleosides from multiple vendors and analyzed freshly prepared aqueous solutions using LC-UV-MS. For the majority of compounds, chromatographic analysis revealed a single dominant peak corresponding to the expected nucleoside. However, we observed notable exceptions. We found 1-methyladenosine (m^1^A) to be contaminated with 6-methyladenosine (m^6^A) (Figure S1), consistent with the thermodynamically favored Dimroth rearrangement in aqueous solution (Section S3.2.2 of Supplementary Information 2 “SI_2”). Furthermore, we found 5- (carboxyhydroxymethyl)-2’-*O*-methyluridine (mchm^5^Um) to be sold in a mixture of its S-and R-forms which complicates accurate quantification (Figure S2, (29,30)). Overall, these results indicate that commercially available nucleoside standards generally match their expected chemical identity but may contain impurities or stereochemical mixtures that require consideration during quantitative analysis. A comprehensive list of currently available commercial nucleosides is provided in Table S1.

### Considerations for quantitative use of nucleoside standards

For quantitative assessment of nucleoside abundance in native RNA samples, commercial standards are weighed (1), dissolved to reach a defined concentration (2) and stored at -20 °C or -80 °C (3) prior to their use. Due to the high price of many modified nucleosides (hundreds to thousands of euros per mg), it is common practice for laboratories to prepare these stocks once, store them at -20 or -80°C and pull aliquots upon need (4). Each of these steps may affect the identity and quantity of the nucleoside stocks and thereby introduce analytical bias. We therefore set out to systematically assess all steps of this common practice and provide practical guidelines and best practices for both analytical chemists and RNA biologists working with RNA modification data.

While weighing is often considered a minor technical step, it can introduce a substantial source of quantitative bias. Many modified nucleosides are only affordable in small quantities which requires specialized weighing capabilities to minimize errors. From Table S1 we selected 45 nucleosides (Table S3) and purchased them in quantities above 10 mg. This allowed us to weigh each nucleoside three times within a defined mass range of 0.5-3 mg (Table S4). This range was selected because it exceeds the balance partition value by a factor of 5000, resulting in a theoretical weighing uncertainty below 1%.

After dilution and LC-UV analysis, we calculated the relative standard deviation (RSD) of the resulting peak areas and we assessed the weighing and dilution accuracy. As shown in Figure S3, the RSD was below 5% for most nucleosides, with 8 nucleosides being between 5-10% and 2 being above 10%. These results indicate that weighing and dilution can introduce quantitative deviations of several percent. For the tested mass ranges, the relative error remained constant, with only a slight increase in the 0.5 - 1.0 mg range. Based on these results, we recommend weighing more than 1 mg of nucleoside standard whenever possible.

### Determination of nucleoside concentration using UV or quantitative NMR spectroscopy

Weighing and dilution can introduce a quantitative bias into nucleoside stock solutions at the stage of preparation. A second bias is the assumed purity of the utilized nucleoside powder. Most commercial standards display a purity value in their certificate of analysis. We want to point out that some vendors state on their websites that their standards are only for qualitative analysis and not for quantitative. This information is important as it tells the analyst that the given purity value is potentially inaccurate which may introduce bias in quantitative down-stream analyses. For assessment of the true purity, and the weighing/dilution bias, we recommend further quality assurance measures to ensure the quantity of the modified nucleoside solutions. For ∼ 40 nucleosides, molar extinction coefficients are known (Table S2) which allows, in theory, quantification by UV absorption. Using the Beer-Lambert law, the concentration of nucleoside solutions can be determined in a cuvette with a UV photometer (attention: pH must be adjusted correctly). A comparison of the theoretical concentration and the experimentally determined concentration allow assessment of the nucleosides’ purity. This may be especially useful if the storage container is not gas tight and water evaporates during the months and years of storage. Importantly, we noticed that polypropylene (PP) vials commonly used for containing nucleoside stocks leach UV-active compounds into the solution. The inset of Figure 1A shows the UV-spectrum of water stored in a standard PP vial for 2 months. An absorption maximum at λ = 230 nm is apparent with a remaining absorption at 254 nm, which is the standard absorption wavelength of nucleosides. The impact of this contaminant is also apparent in LC-UV analysis, where a distinct peak becomes visible after nucleoside storage in PP (Figure 1A). Our mitigation strategy is the use of glass vials. Here, we observe no occurrence of contaminants. Thus, we recommend the use of glass vials for long-term storage of nucleoside solutions in combination with regular quantification using UV absorption. An additional benefit of glass containers was uncovered when we gravimetrically assessed evaporation. While PP vials showed ∼ 4.5 % evaporation of water at 60 °C, glass vials had less than 1 % of water loss (Figure S4). To account for evaporation, we recommend the immediate analysis of the nucleoside solution by UV absorption and an in-house determination of extinction coefficients, especially for those nucleosides with unknown extinction coefficients.

**Figure 1.**
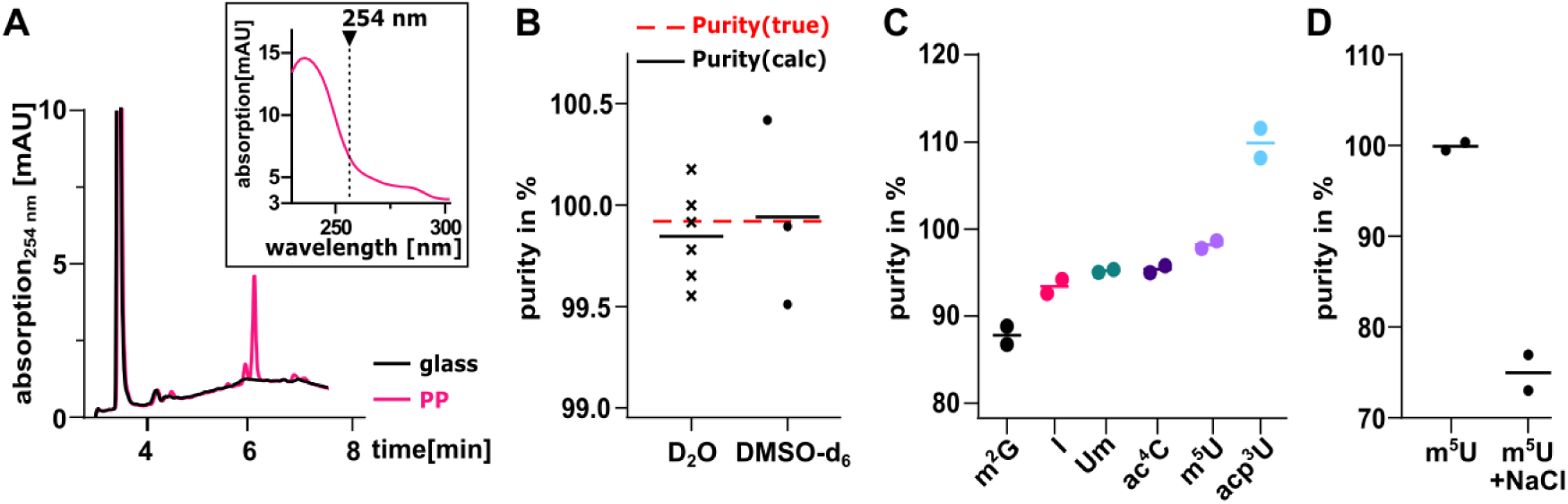
Quantitative assessment of nucleoside stock solution purity by UV and NMR spectroscopy. **(A)** LC-UV chromatogram at λ= 254 nm of inosine solution stored in polypropylene (PP) and glass vials for 2 months. Inset: UV spectrum of water stored in PP vials. **(B)** Purity determination by ^1^H qNMR of secondary standard DMS (dimethyl sulfone P = 99.96 %) using secondary standard MA (maleic acid P = 99.93 %) as internal calibrant in D_2_O (n = 6) and DMSO-d_6_ (n = 3), respectively. Red, dotted line indicates the certified purity of DMS and black lines indicate the calculated purity in D_2_O and DMSO-d_6_, respectively. **(C)** Purity values of six different nucleoside standards obtained by qNMR (n = 2) **(D)** Purity of m^5^U and deliberately contaminated m^5^U (with NaCl as contaminant).

For modified nucleosides with unknown extinction coefficient (> 90 nucleosides) and unclear purity of the used powder, quantitative ^1^H NMR (qNMR) can be useful. Extended considerations for qNMR of nucleoside quantification are given in supplementary information 1 (“SI_1”). Briefly, qNMR is considered a relative-primary method, as the integrated signal area is directly proportional to the number of nuclei contributing to the signal and, consequently, to the amount of substance (31). qNMR is considered to be the most reliable solution to assess the purity of powders. Typically, an internal calibration method is used, where both the analyte and a standard substance are measured within the same sample, which allows direct referencing of the analyte to the standard within a single spectrum. qNMR offers exceptional accuracy and an overall uncertainty as low as 0.15 % (32). As a downside, sample amounts of more than 5 mg are required and complete dissolution in the required 0.5 mL D_2_O is often challenging for nucleosides. An alternative is DMSO-d_6_ (for more detail we refer to SI_1 and Figure S5 and Table S5). Dimethylsulfone (DMS) was selected as the internal standard for nucleoside quantification, and, in a first step, we assessed the purity of DMS with maleic acid as a standard. The results – 99.89% purity of DMS in D_2_O with an RSD of 0.23 % and 99.98% purity in DMSO-d_6_ with an RSD of 0.46% – confirmed both the accuracy and high precision of the qNMR method (Figure 1B).

Next, we analyzed six different nucleoside standards commonly used for LC-MS-based quantification. The qNMR purity assessment revealed that most of these standards exhibit variations in purity (Figure 1C and Table S6). 2’-O-methyluridine (Um), 4-acetylcytidine (ac^4^C), and inosine (I) showed purity values ranging between 95% and 99%. As seen in the ^1^H NMR spectrum of ac^4^C (Figure S6), unassignable signals are present that cannot be attributed to any protons of ac^4^C. These signals likely originate from the synthesis process and may indicate the presence of byproducts or residual solvents detectable by NMR. However, no such unassignable signals were observed in the spectra of the other nucleosides. 5-methyluridine (m^5^U) exhibited a purity close to 100%, while 3-(3-amino-3-carboxypropyl)uridine (acp^3^U) showed a purity exceeding 110%, and m^2^G had a purity below 90%. The unusually high purity value of acp^3^U may be explained by the processing issues as described in Figure S5B. For m^2^G, the corresponding ^1^H NMR spectrum (Figure S6) indicates no detectable proton-based impurities, suggesting that residual salt from synthesis may contribute to the lower purity value. To investigate this hypothesis, we conducted a proof-of-principle experiment, where we deliberately contaminated the m^5^U stock powder with NaCl before sample preparation. The m^5^U:NaCl mixture was prepared in a 3:1 ratio, homogenized, and analyzed by qNMR under the same conditions. As shown in Figure 1D, the calculated purity of m^5^U dropped to approximately 75%, which supports our idea that residual salts may influence purity assessments. While this experiment does not directly confirm our hypothesis, it suggests that varying nucleoside purities may stem from residual non-^1^H-detectable contaminants.

In summary, we identified weighing, dissolution and powder purity as sources for incorrect quantities of nucleoside stock solutions. A potential mitigation strategy is the use of initial UV absorption measurements and subsequent storage in glass. If the purity of the powder is unclear and more than 10 mg (2 replicates à 5 mg) are available, we recommend the use of qNMR.

### Determination of nucleoside stability in aqueous solution over 12 months

Modified nucleosides display a vast variety of chemically reactive groups and their most common solvent, water, is a highly reactive chemical as well. We hypothesized that some nucleosides may react with water which impacts the qualitative and quantitative data recorded by LC-MS. Thus, we set out to define the stability of 40 modified and the four canonical nucleosides in aqueous solution following the guideline for stability testing of new drug substances and products (CPMP/ICH/2736/99) from the European Medicines Agency (EMA). An outline of the study design is given in Figure 2A. The analysis of each nucleoside should be conducted in three technical replicates. The long-term shelf life of the nucleosides was assessed under normal storage conditions (-20 °C/-80 °C). In this manuscript, we assess stability over a 12-month period by taking measurements at 1, 2, 3, 6, and 12 months for samples stored at -20 °C, and at 3, 6, and 12 months for those stored at -80 °C. Additionally, the substances were monitored at elevated temperatures of 8 °C (refrigerator) and room temperature (RT) over a period of three months, with timepoints at one, two and three months, to verify their sensitivity to short-term temperature increases. Such long-term experiments require substantial logistical planning, and we critically discuss the decisions that had to be made. We opted to store the nucleosides in 96-well plates sealed with aluminium foil, as previous studies have shown that evaporation is minimal (33). Furthermore, we decided to store the nucleosides in a diluted form (10 µM) rather than at their stock concentration (10 mM). This decision is backed by the fact that dilution of all replicates to 10 µM at the start avoids dilution errors at each measuring timepoint. Thus, all nucleoside stock solutions were dissolved at 10 mM, distributed onto two 96-well deep-well plates (1.5 ml per well; designated as ’master plates’) and serially diluted to 10 µM prior to distribution into the 40 standard 96-well plates (30 µl per well; designated as ’storage plates’). The first two plates were analyzed by LC-UV-MS immediately after dissolution to establish a reference point. The remaining plates were sealed with aluminium foil and stored at their respective temperatures. Evaporation was assessed by storing nucleoside solutions in cryo-vials (PP) and comparing LC-UV signals to those obtained from nucleosides stored in 96-well plates. Cryo-vials showed higher apparent recovery, consistent with concentration effects caused by stronger solvent evaporation during storage (Figure S7). These results indicate that solvent evaporation is lower in 96-well plates than in standard PP cryo-vials.

**Figure 2.**
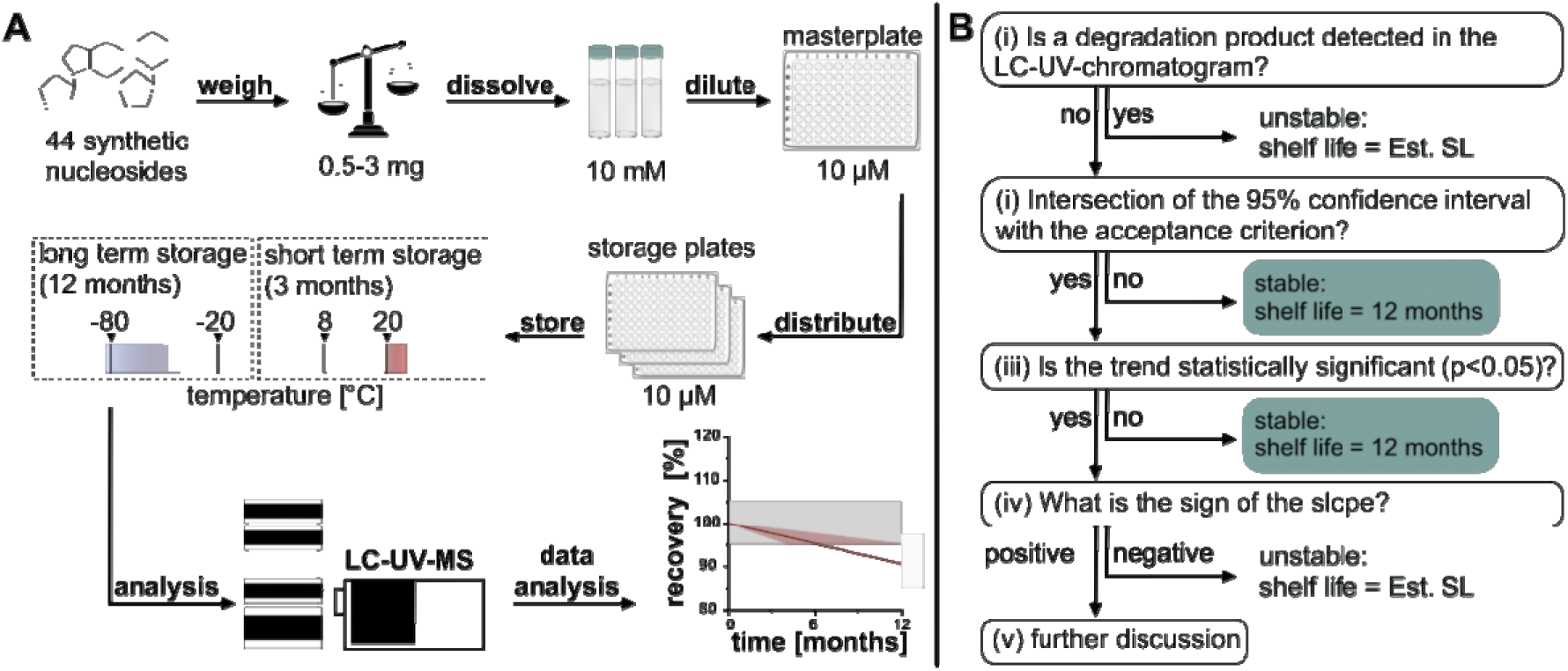
**(A)** Graphical overview of the study design and workflow which was used to determine the short-term and long-term stability of nucleoside solutions at different storage temperatures. Stability of the compound is assessed over time using LC-UV-MS (liquid chromatography–ultraviolet–mass spectrometry) **(B)** Decision tree that assesses all statistical and physicochemical parameters needed to judge nucleoside stability.

### LC-UV detection revealed degradation products in seven of the 44 nucleosides

Each nucleoside was injected into a prepared LC-UV-MS system with a 9-minute run time which resulted in a runtime of ∼ 20 hours per storage condition. Each plate stayed in storage until the morning of its use and was then thawed for 1 hour at room temperature followed by a spin-vortex-spin procedure to ensure homogeneity of the analyte solutions. The plate was then placed in the autosampler at 8°C. Analytes were separated by reverse-phase chromatography and the UV signal at a designated wavelength (see Table S7) was recorded prior to a mass scan ranging from *m/z* 100-600. The UV signal of the analyte and all emerging peaks were integrated and used to monitor the stability of the nucleosides. The MS scan data from the emerging peaks was analyzed to allow first hypotheses on the chemical structures of the decomposition products. After analysis, the plates were discarded. For each nucleoside replicate, the UV areas of the nucleoside peak were integrated and related to the replicate’s area at time point 0. Time point 0 is the reference point and equals 100 % recovery.

All nucleosides were screened by inspection of the resulting LC-UV and LC-MS chromatograms after -20 °C / -80°C storage. We followed the decision tree in Figure 2B to determine whether a nucleoside can be considered stable or unstable. The first decision is made by screening for degradation products. If a degradation product is found, the nucleoside is immediately considered unstable. The occurrence of new peaks (not those from PP vials, see Figure 1A) is interpretated as formation of degradation products. The chromatograms of the affected nucleosides m^1^A, 3-methylcytidine (m^3^C), mcm^5^s^2^U, i^6^A, 2-methylthio-N6-isopentenyladenosine (ms^2^i^6^A) and s^4^U are given in Figure 3 alongside the quantitative impact on the recovery. For m^1^A, m^3^C and s^4^U we find the previously described degradation products, namely m^6^A, m^3^U and di-s^4^U (11,13,34). For the bacterial tRNA modification s^4^U we identified uridine as a second degradation product. The relative distribution of the di-s^4^U and U products is temperature dependent. At 8 °C, RT and -20 °C, uridine formation dominates, which is consistent with a desulfurization reaction. In contrast, lower temperatures (-80 °C) favour the formation of di-s^4^U, suggesting a radical-mediated mechanism that proceeds via thiol oxidation and disulfide bond formation (Figure S8).

**Figure 3.**
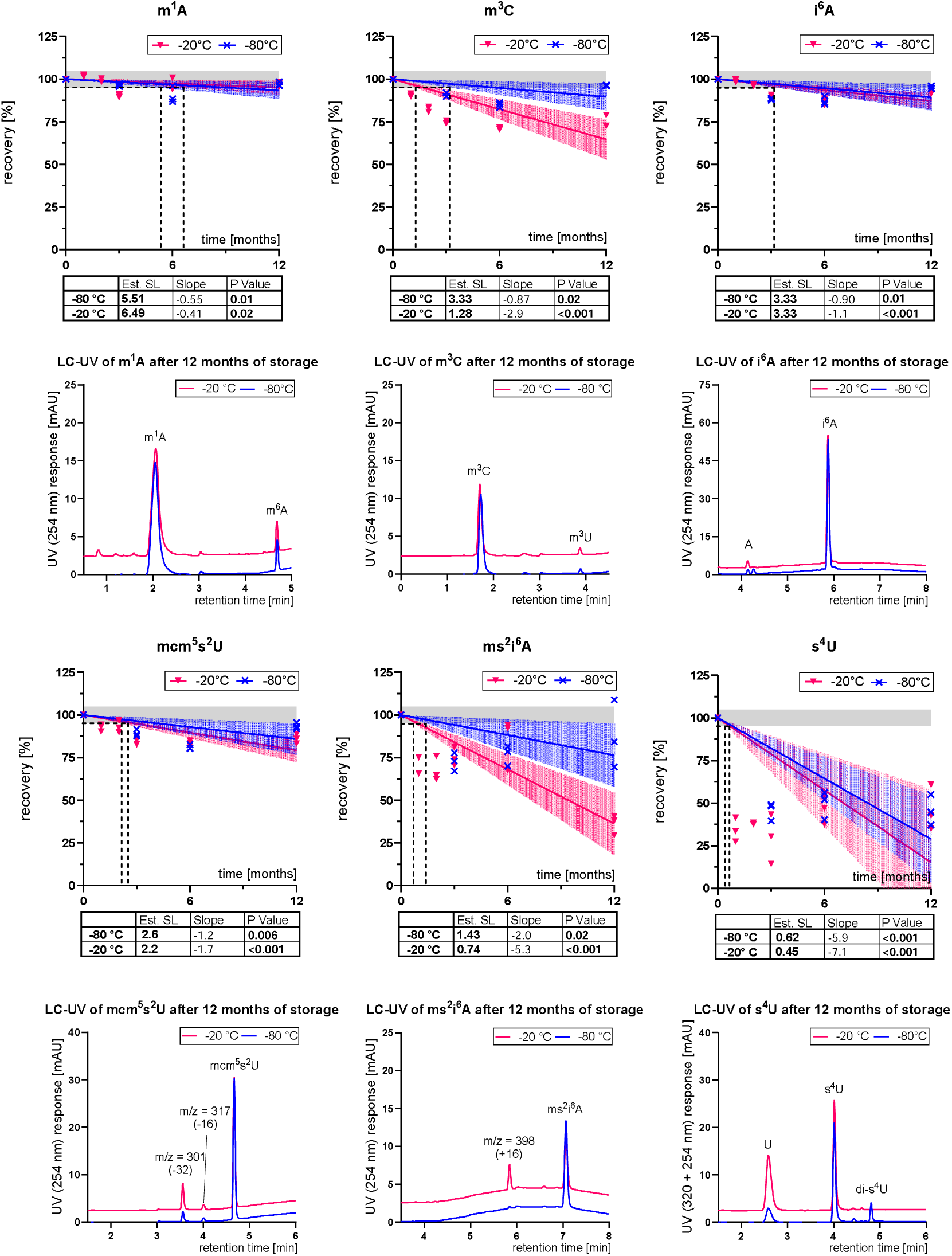
Unstable nucleosides under storage conditions (-20 °C/ -80 °C) with corresponding chromatograms of degradation products. Shelf life was determined by comparing the UV area after storage to the initial value and normalized to 100 %. The grey shaded region indicates the acceptance criterion of ±5 % deviation (95–105 %). Coloured shaded bands represent the 95 % confidence interval of the linear regression. Statistical parameters (slope, p-value, and Est. SL) are provided in the inset tables.

For 6-isopentenyladenosine (i^6^A), we identified a time-dependent conversion into adenosine. (Figure S8). These results indicate a small (∼5 %) but consistent chemical lability of i^6^A under prolonged storage. The identification of degradation products for mcm^5^s^2^U, ms^2^i^6^A required a more detailed analysis using HRAMS (high resolution accuracy mass spectrometry).

Hypermodified uridine derivatives show a particularly interesting behaviour in their degradation pathways. In Figure 3 the chromatogram of mcm^5^s^2^U shows the formation of two degradation products (DP). One with *m/z* 301 (DP1) and one with *m/z* 317 (DP2). HRAMS analysis revealed that the *m/z* 301 product results from a complete loss of sulfur (Figure 4A). This highly uncommon degradation product is further supported by a series of MS fragmentation products consistent with the sequential loss of the ribose and methyl group (Figure S9). An additional strong indication of the structure is the observed mass deviation of less than 3 ppm relative to the calculated theoretical value, supporting accurate structural assignment. The value of HRAMS for such analyses becomes apparent in Figure S11. A zoom into the *m/z* range of the M+2 range clearly indicates the presence of the natural sulfur-34 isotope in mcm^5^s^2^U and its absence in the *m/z* 301 product, while both show the M+2 peak expected for natural oxygen-18. Importantly, recent literature explicitly demonstrated that mcm^5^s^2^U undergoes oxidative desulfurization under cellular stress *in vivo*, forming the stable 4-pyrimidinone derivative mcm^5^H^2^U (35).The second degradation product (DP2) has a *m/z* 317 which matches the *m/z* of mcm^5^U and its retention time. Yet, HRAMS analysis reported an *m/z* of 317.2153 for the second degradation product and an *m/z* of 317.0984 for mcm^5^U (Figure 4A and Figure S11). These masses deviate by 368 ppm which is a clear sign that the second degradation product is not mcm^5^U but another, yet unknown, nucleoside. These findings intrigued us to investigate the stability of related hypermodified uridine derivatives in more detail. We therefore extended the short-term storage at elevated temperatures for hypermodified uridines (cm^5^U, mcm^5^U, ncm^5^U and mcm^5^Um). After 6 months of storage at 8 °C and 20 °C, all hypermodified uridines showed recoveries below 100%, indicating partial degradation, with mcm^5^Um being most affected (Figure S12). To investigate the degradation pathway of mcm^5^Um in more detail, we analyzed the corresponding degradation products. For mcm^5^Um we observed a cleavage of the ether leading to cm^5^Um as the product. Interestingly, this cleavage was not observed for the structurally related derivatives mcm^5^s^2^U and mcm^5^U under the same conditions (Figure S13). While this observation does not impact storage at -20 and -80 °C, it may impact the stability of these modified nucleotides within native tRNA.

**Figure 4.**
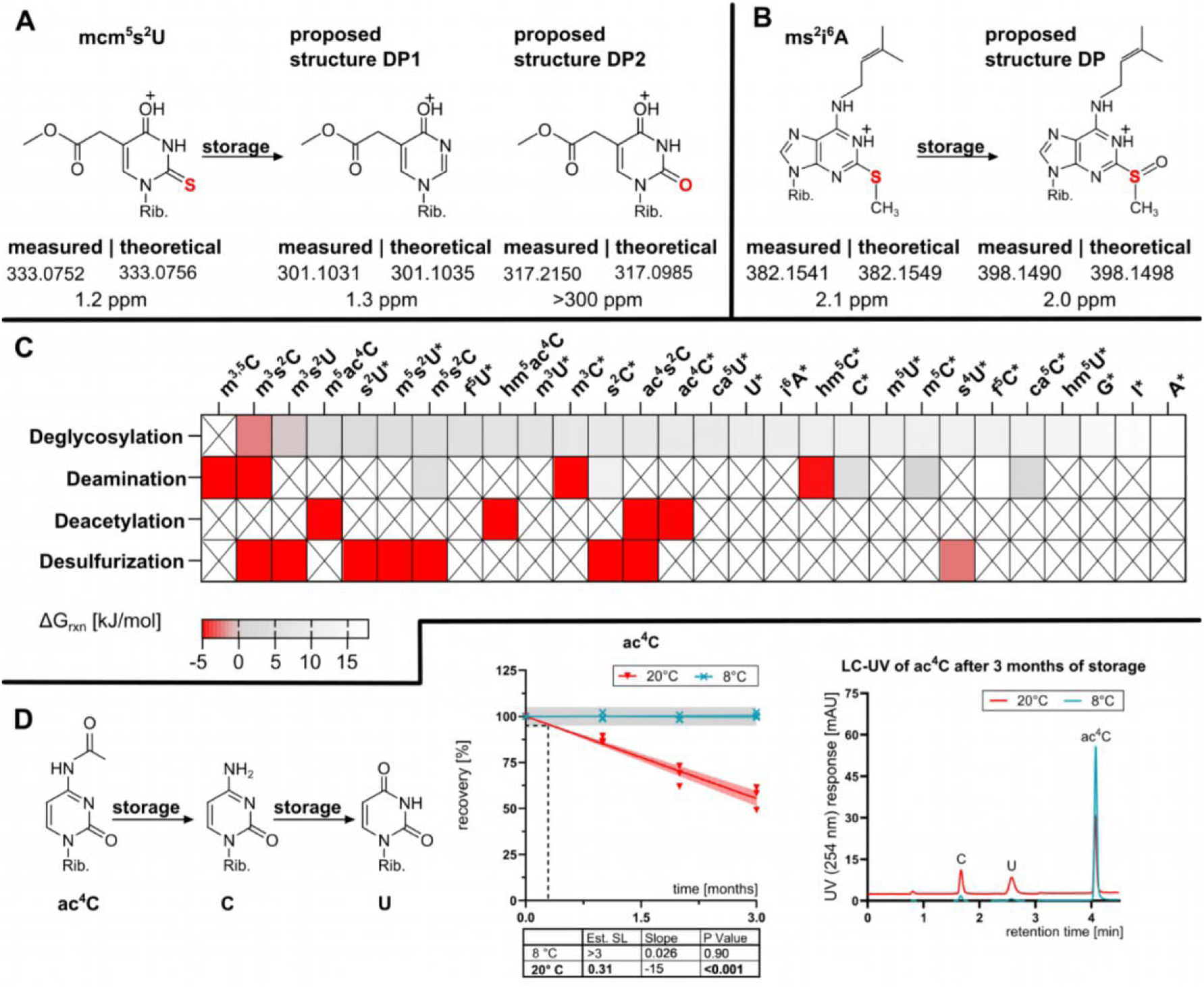
High-resolution mass spectrometry (HRMS) analysis of proposed degradation products. The figure shows the proposed structures of storage-induced degradation products alongside their corresponding MS1 HRMS data (measured mass, theoretical mass, and mass error in ppm). **(A)** For the degradation of mcm□s²U, HRMS data supports the proposed structure DP1 (1.3 ppm error), while the proposed structure DP2 is not supported by the data (>300 ppm error). **(B)** For the degradation of ms²i□A, the HRMS data is consistent with the proposed structure DP (2.0 ppm error)**. (C)** Reaction free energies for hydrolytic deglycosylation, hydrolytic deamination, hydrolytic deacetylation of C4-acetylated cytosine derivatives and hydrolytic de-sulfurization of nucleosides and nucleobases with thiocarbonyl groups. * indicates nucleosides found in native RNA. Red color indicates negative free reaction energies equaling exothermic reactions. X indicates that the reaction was not calculated. **(D)** Shelf life of ac^4^C at elevated temperature (8 °C and RT) with corresponding chromatograms of degradation products. Shelf life was determined by comparing the UV area after storage to the initial value and normalized to 100%. The grey shaded region indicates the acceptance criterion of ±5 % deviation (95–105 %). Coloured shaded bands represent the 95 % confidence interval of the linear regression. Statistical parameters (slope, p-value, and Est. SL) are provided in the inset tables.

HRAMS analysis of ms^2^i^6^A and its degradation product revealed a mass shift corresponding to the addition of one oxygen atom (Figure 4B). Chemically, the most probable degradation pathway is the oxidation of the reactive thioether group to a sulfoxide. Yet, MS/MS fragmentation of the base indicates the location of the oxygen on the nucleobase and ruling out an oxidation at the isopentenyl moiety (which would have yielded the natural nucleoside ms^2^io^6^A) (Figure S10). HRAMS and MS/MS fragmentation strongly indicate formation of a sulfoxide. According to our own guideline on nucleoside structure identification (36), we cannot confirm this hypothesis and truly identify the degradation product of ms^2^i^6^A without synthetic reference standards.

While the previously discussed nucleosides exhibited degradation products already at common storage temperatures of -20 °C and -80 °C, N4-acetylcytidine (ac^4^C) was the only nucleoside that degraded exclusively at room temperature (Figure 4D). LC-UV-MS analysis indicates that ac^4^C decays via a two-step hydrolytic pathway: initial deacetylation to cytidine (C) followed by deamination to uridine (U). This is confirmed by new peaks at the retention times of C and U, with corresponding [M+H]^+^ *m/z* signals of 244 and 245. These results confirm that ac^4^C lacks stability in aqueous solutions at ambient temperatures and is likely sensitive to freeze-thaw cycles. This impacts the current debate on the presence of ac^4^C in human mRNA, as discussed below.

### Theoretical analysis of degradation pathways

Intrigued by the identified degradation products, we asked the question which degradation products are chemically most likely. The degradation pathways for nucleosides analyzed here include (a) the deglycosylation reaction with water to yield the free nucleobase and ribofuranose; (b) the hydrolytic deamination of cytosine derivatives to yield the respective uracil bases and ammonia; (c) the hydrolytic deacetylation of 4-acetylaminocytosine derivatives to yield acetic acid and the respective cytosines; and (d) hydrolytic desulfurization of C2-thiocarbonyl derivatives of cytosine and uracil to yield hydrogen sulfide and the respective parent compounds. Reaction free energies (ΔG_rxn_, in kJ mol^-1^) for all four reaction types are obtained by subtracting the free energies ΔG_298_ of all reactants from those of the products (Table S8). Results for all reactions are summarized in the heatmap of Figure 4C and in a graphical manner in Figure S14 and S15 of SI_1. Introducing a positive charge into the nucleobase by N-methylation is expected to facilitate the initial heterolytic C1’−N bond cleavage in SN1-like hydrolysis, which eliminates the need to generate charge-separation. In line with this picture, deglycosylation becomes less ender-gonic upon methylation in our calculations (ΔΔGrxn ≈ -5 kJ mol-1, Section S3.2.1 of SI_2), and these nucleosides indeed displayed reduced stability in aqueous storage. For most of the nucleosides studied here the deglycosylation reaction is endergonic by 3 - 11 kJ mol^-1^ and these systems can be considered to be stable. The reaction free energies for the hydrolytic deamination of cytosine derivatives are collected for the full nucleoside (Figure 4C) as well as the respective free bases (Figure S14B). For cytidine (C) as the reference system of this group the deamination is moderately endergonic by 4.5 kJ mol^-1^. Among the natural nucleoside modifications, only m^3^C and hm^5^C show negative ΔG_rxn_ indicating their preference for spontaneous deamination (ΔG_rxn_ -29.2 and -4.6, respectively). For m^3^C, the detected deamination product m^3^U (Figure 3) confirms the overlap of theoretical calculation and wet lab experiment. In contrast, while hm^5^C demonstrated experimental instability during storage at -20 °C (with an estimated shelf life of 11.4 months), we could not detect any corresponding degradation product (Figure S18) Adenosine to inosine has a positive ΔG_rxn_ indicating the low tendency for deamination in buffer-free water.

Hydrolytic de-acetylation of C4 amino group-acetylated cytosine derivatives is exergonic for all nucleosides and nucleobases studied here (Figure S15A). The driving force for deacetylation amounts to -10.2 kJ mol^-1^ for cytidine itself, which is in accordance with our experimental data described in Figure 4D. It becomes notably larger for C5-substituted cytidine derivatives. The most favorable reaction energy is found for the 5-methyl substituted system (m^5^ac^4^C) with ΔG_rxn_ = -24.9 kJ mol^-1^. These results open room for speculation: Is 4-acetyl-5-methylcytidine a non-natural RNA modification, or is it natural but due to its instability undetectable in current analytical systems?

Reaction energies for hydrolytic de-sulfurization as the fourth degradation pathway analyzed here are shown in Figure 4C and in Figure S15B in a graphical manner. De-sulfurization is found to be energetically favorable for all systems studied here with reaction energies ranging from -2.2 kJ mol^-1^ for 4-thiouridine (s^4^U) to -16.5 mol^-1^ for 2-thiouridine (s^2^U) an intermediate of hypermodified uridines. This indicates that desulfurization of *e.g.* mcm^5^s^2^U to mcm^5^U is chemically favored and plausible. Interestingly, although our HRAMS analysis clearly detected a degradation product exhibiting a complete loss of sulfur (m/z 301), the specific formation of mcm^5^U could not be confirmed, highlighting the complexity of these degradation pathways in solution (Figure 4A).

Finally, reaction energies were also calculated for the two competing pathways for the decomposition of s^4^U shown in Figure 3 and Figure S8. For the free base of this latter system oxidative dimerization employing triplet oxygen as the reaction partner to yield the respective s^4^U dimer and water as the products is highly exothermic at ΔG_rxn_ = -101.6 kJ/mol, whereas hydrolytic loss of sulfur is only marginally exergonic (ΔG_rxn_ = –3.8 kJ/mol). However, in dilute aqueous solution there is an abundance of water available as a reactant, while the availability of dissolved oxygen may, together with the requirement of radical-chain initiation, limit the role of oxidative dimerization. That oxidative dimerization becomes less competitive at higher temperatures may also be due to the entropic requirements of a reaction turning three separate reactants into two products. This complex interplay of individual factors apparently generates a scenario, where both pathways coexist in parallel.

All taken together, the calculation of reaction free energies for the four general degradation reactions analyzed here point to hydrolytic de-sulfurization and hydrolytic de-acetylation are the more favorable degradation reactions, while hydrolytic deglycosylation is least favorable for all systems studied. As already implied in the definition of these processes, reactions (a) - (d) are hydrolytic in the sense of requiring water as the reactant. While the concentration of this reactant is high in aqueous solution, it may be quite low in DMSO solution. Among others, this may represent one of the reasons for the observed reactivity differences in these two solvents.

### Handling of physical artifacts and conservative shelf-life estimation

Following the decision tree in Figure 2B and the guidance of ICH Q1E, we performed a linear regression of the recovery calculated for each timepoint, followed by the calculation of the 95 % confidence interval. Whenever the confidence interval intersects the 5 % acceptance criterion (as recommended by ICH), it indicates potential chemical instability. The time point of intersection is defined as the estimated shelf life (Est. SL).

However, applying strict ICH guidelines, which are originally designed for highly optimized, single-substance assays, is not entirely suitable for the simultaneous analysis of so many compounds (∼2000 datapoints). In such broad analytical setups, normal technical variations can naturally inflate the confidence intervals. Evaluating the data solely by strict ICH criteria would thus falsely classify many nucleosides as unstable due to standard signal fluctuations rather than actual degradation. Therefore, we also considered the statistical significance of the linear regression slope (Figure 2B).

Although an increase in analyte amount is chemically impossible, we observed cases with a significant regression but a positive slope. While we do not yet understand why this phenomenon occurs, it is highly reproducible, occurs in 96-well plates, cryo-vials (both PP), at all temperatures and for inosine even in glass vials stored at room temperature (Figure S22 and S23). The most likely explanation for this is evaporation. As shown in Figure S7, evaporation is a factor in the cryo-vials and is thus also likely to occur in the 96-well plate format. For the short-term storage at elevated temperatures (8 °C and RT), this evaporation was in some cases even visually apparent through a loss of sample volume. Because this evaporation systematically increases the apparent analyte concentration, we decided to strictly exclude the intersection with the upper acceptance limit when evaluating stability at these elevated temperatures. Consequently, determining an accurate shelf life at 8 °C and RT is not feasible under these conditions. Instead, these conditions serve as a qualitative stress test, where any observed degradation clearly identifies highly temperature-sensitive nucleosides.

Interestingly, we also observed a statistically significant increase in concentration (a positive slope) for certain nucleosides during long-term storage at -20 °C and -80 °C (Figure 5A and S20). While this could generally be attributed to evaporation, two nucleosides (Im, m^2^G) exhibited a remarkably steep positive slope (Figure 5A). Alternative explanations include precipitation and solubility issues, or initial adsorption to the polypropylene of the 96-well plates followed by a gradual release back into solution. While we cannot explain the cause of this phenomenon, we had to decide on the stability of these nucleosides, and we adopted a strictly conservative approach: for these cases at -20 °C and -80 °C the estimated shelf life (Est. SL) resulting from the intersection with the upper limit was reported as the true shelf life. This ensures that we do not falsely guarantee a longer shelf life beyond what is analytically certain.

**Figure 5.**
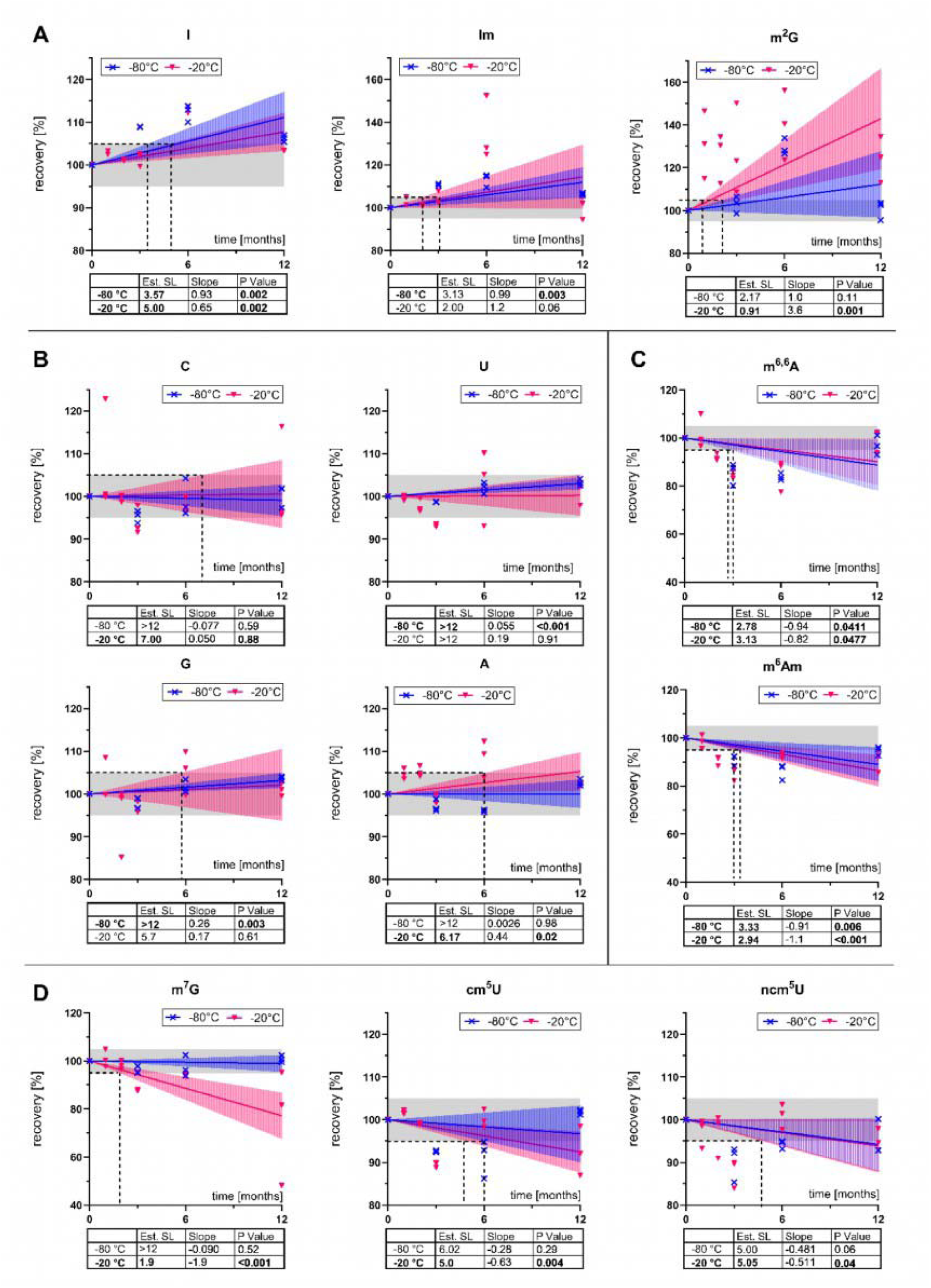
Stability assessment of aqueous nucleoside solutions using LC-UV-MS. Recovery was determined by comparing the UV area after storage to the initial value and normalized to 100 %. The grey shaded region indicates the acceptance criterion of ±5 % deviation (95–105 %). Coloured shaded bands represent the 95% confidence interval of the linear regression. Statistical parameters (slope, p-value, and estimated shelf life (Est. SL)) are provided in the inset tables. **(A)** The canonical nucleosides are stable under at least one long-term storage condition (-80 °C and -20 °C) **(B)** m^6,6^A and m^6,6^Am showing a statistically significant negative slope (degradation), but without detectable degradation products in LC-UV for -80 °C and -20 °C. **(C)** m^7^G, cm^5^U and ncm^5^U showing a statistically significant negative slope (degradation), but without detectable degradation products in LC-UV for -20 °C but not for -80 °C **(D)** I, Im and m^2^G and show a significant positive slope for -20 °C and -80 °C

### 30 out of 44 studied nucleosides are stable for 12 months in aqueous solution at -80 °C or -20 °C

In summary, the four canonicals and 26 modified nucleosides are considered to be stable for 12 months at either -20 or -80 °C (Figure 5B, S17, S18, S20 and summarized in Table 2). Notably, for 15 of these nucleosides, we could confirm stability over 12 months at both common storage temperatures. (Figure S16). Whenever a nucleoside showed a statistically significant regression without detectable degradation products, the intersection of the regression with the acceptance criterion was used to calculate its shelf life. m^6,6^A, m^6^Am are the only nucleosides exhibiting instability at both -20 °C and -80 °C, characterized by a negative regression without the formation of detectable degradation products (Figure 5C). Furthermore, nine nucleosides (Cm, m^5^C, m^4^C, hm^5^C, m^5^Um, mcm^5^U, m^7^G, ncm^5^U and cm^5^U) are stable at -80 °C, but show lower recoveries when stored at -20 °C (Figure 5D and S18). For m^7^G potential spontaneous depurination (37) likely explains the absence of detectable degradation products in our analysis. Finally, short-term storage at elevated temperatures revealed pronounced instability for all nucleosides that formed degradation products, as well as for ncm U, cm U, m, A, and m Am (Figures S19 and S21).

**Table 2:** Shelf life of nucleoside solutions. (^#^) indicates nucleosides that should be stored in DMSO. (*) indicates Intersection between the confidence interval (CI) and acceptance criteria, without a significant slope. (§) within 3 months of storage.

|  |  |  |  |  |  |  |
| --- | --- | --- | --- | --- | --- | --- |
|  |  | water |  |  | DMSO |  |
|  | Nucleoside | -80 °C | -20 °C | Instability at 8 °C or RT<sup>§</sup> | -20 °C | Degradation product |
| Shelf life >12 months at -20 °C and -80 °C | C | 12.0 | 12.0* |  | - |  |
|  | Ψ | 12.0 | 12.0* |  | - |  |
|  | U | 12.0 | 12.0 |  | - |  |
|  | Um | 12.0 | 12.0* |  | - |  |
|  | m<sup>5</sup>U | 12.0 | 12.0* |  | - |  |
|  | G | 12.0 | 12.0* |  | - |  |
|  | m<sup>2,2</sup>G | 12.0 | 12.0 |  | - |  |
|  | mcm<sup>5</sup>Um | 12.0* | 12.0* |  | - |  |
|  | t<sup>6</sup>A | 12.0 | 12.0 |  | - |  |
|  | m<sup>6</sup>t<sup>6</sup>A | 12.0* | 12.0* |  | - |  |
|  | Am | 12.0 | 12.0 |  | - |  |
|  | m<sup>6</sup>A | 12.0* | 12.0* | Yes (8°C) | - |  |
|  | m<sup>4</sup>Cm | 12.0 | 12.0 |  | - |  |
|  | m<sup>5</sup>Cm | 12.0 | 12.0* |  | - |  |
|  | m<sup>2</sup>A | 12.0 | 12.0* |  | - |  |
| Shelf life >12 months at -80 °C | Cm | 12.0 | 7.7 |  | - |  |
|  | m<sup>5</sup>C | 12.0 | 10.2 |  | - |  |
|  | hm<sup>5</sup>C | 12.0 | 11.4 |  | - |  |
|  | A | 12.0 | 6.2 |  | 3.0 |  |
|  | Gm | 12.0* | 5.1 |  | - |  |
|  | m<sup>7</sup>G | 12.0* | 1.9 |  | 3.0 |  |
|  | m<sup>3</sup>U | 12.0 | - |  | - |  |
|  | m<sup>4</sup>C | 12.0 | 10.4 |  | - |  |
|  | m<sup>5</sup>Um | 12.0* | 5.4 |  | - |  |
|  | ncm<sup>5</sup>U | 12.0* | 5.0 | Yes (RT; 8°C) | - |  |
|  | cm<sup>5</sup>U | 12.0* | 5.0 | Yes (RT; 8°C) | 3.0 |  |
|  | mcm<sup>5</sup>U | 12.0* | 5.4 |  | - |  |
| Shelf life >12 months at -20 °C | f<sup>5</sup>C | 5.6 | 12.0 |  | - |  |
|  | m<sup>1</sup>G | 10.9 | 12.0* |  | - |  |
|  | m<sup>1</sup>Ψ | 6.0 | 12.0 |  | - |  |
| Shelf life < 12 months | Im | 3.1 | 2.0 |  | 1.0 |  |
|  | m<sup>3</sup>C | 3.3 | 1.3 | Yes (RT; 8°C) | 3.0 | m<sup>3</sup>U |
|  | I | 3.6 | 5.0 |  | 2.3 |  |
|  | m<sup>1</sup>A | 5.5 | 6.0 | Yes (8°C) | 2.1 | m<sup>6</sup>A |
|  | ac<sup>4</sup>C<sup>#</sup> | 3.1 | 6.0 | Yes (RT) | 3.0 | C, U |
|  | m<sup>2</sup>G<sup>#</sup> | 2.2 | <1 |  | 3.0 |  |
|  | mcm<sup>5</sup>s<sup>2</sup>U<sup>#</sup> | 2.5 | 2.2 | Yes (RT; 8°C) | 3.0 | 301; 317 |
|  | i<sup>6</sup>A | 3.3 | 3.3 | Yes (RT; 8°C) | 1.9 | A |
|  | acp<sup>3</sup>U | 10.6 | - |  | - |  |
|  | ms<sup>2</sup>t<sup>6</sup>A | 3.9 | 4.6 |  | 3.0 |  |
|  | m<sup>6,6</sup>A | 2.8 | 3.1 | Yes (8°C) | 3.0 |  |
|  | m<sup>6</sup>Am | 3.3 | 2.9 | Yes (8°C) | 3.0 |  |
|  | s<sup>4</sup>U<sup>#</sup> | <1 | <1 | Yes (RT; 8°C) | 3.0 | U; di-s<sup>4</sup>U |
|  | ms<sup>2</sup>i<sup>6</sup>A | 1.4 | <1 | Yes (RT; 8°C) | - | 398 |

### m^2^G, s^4^U, and mcm^5^s^2^U show improved stability in DMSO

After 6 months of storage in aqueous solution more than a dozen nucleosides were found to change their quantities during storage. While this observation has no consequence for relative quantification where the abundance of a modification of sample A and sample B are compared within the same analytical run, it introduces a bias into analyses separated by months. A direct comparison of sample batches would not be possible. Further, absolute quantification that determines the stoichiometric number of a modification per canonical nucleoside or per RNA is not accurate. Therefore, we searched for mitigation strategies and overcome the instability of these nucleosides. One possibility is storage in a solvent that is chemically less aggressive, and we decided to use DMSO (dimethylsulfoxide) as nucleosides have a high solubility in DMSO which may mitigate the phenomenon observed for I, Im and m^2^G. Due to our studies in PP and the fact that DMSO liberates even more leachables than water, we decided that glass vials should be used. While glass containers are compatible with DMSO, the caps and/or seals cannot withstand DMSO easily and lose integrity, especially below -40 °C. Thus, storage of nucleosides in DMSO at -80 °C would cause the seals/caps to erode and evaporation is accelerated.

For this reason, we selected glass vials and caps/seals compatible with DMSO and a storage temperature of -20 °C. The 15 nucleosides with the highest quantitative change at 6 months storage at any temperature were selected, weighed fresh (n=3), dissolved in DMSO and an aliquot diluted with water for immediate LC-UV-MS analysis (start point) and UV spectroscopy. The remaining tubes were stored at -20 °C. Extinction coefficients for nucleoside stocks stored in DMSO were determined spectrophotometrically after 1:1000 dilution in water for additional quality control (Table S9). Freeze-thaw cycles were included in the experimental design, as we hypothesized that repeated temperature shifts could induce degradation, especially for nucleosides like ac C that are stable when frozen but highly temperature-sensitive.

7 nucleosides (m^3^C, mcm^5^s^2^U, s^4^U, cm^5^U, m^7^G, m^6,6^A, and m^6^Am) that had a negative slope or degradation products in aqueous storage can be considered stable over 3 months in DMSO at -80 °C (Figure 6A-C and Figure S24). Yet, m^1^A and i^6^A degrade significantly in DMSO within the 3 months, consistent with their behavior in water, with no improvement in the estimated shelf life (Figure 6D). Nucleosides showing a highly positive slope in water, I and Im, still show a positive slope in DMSO, indicating that the underlying cause is not solvent-dependent. Only m^2^G, which had a substantial positive slope in water, appears to show normal and stable behavior in DMSO at -20 °C (Figure 6E).

**Figure 6.**
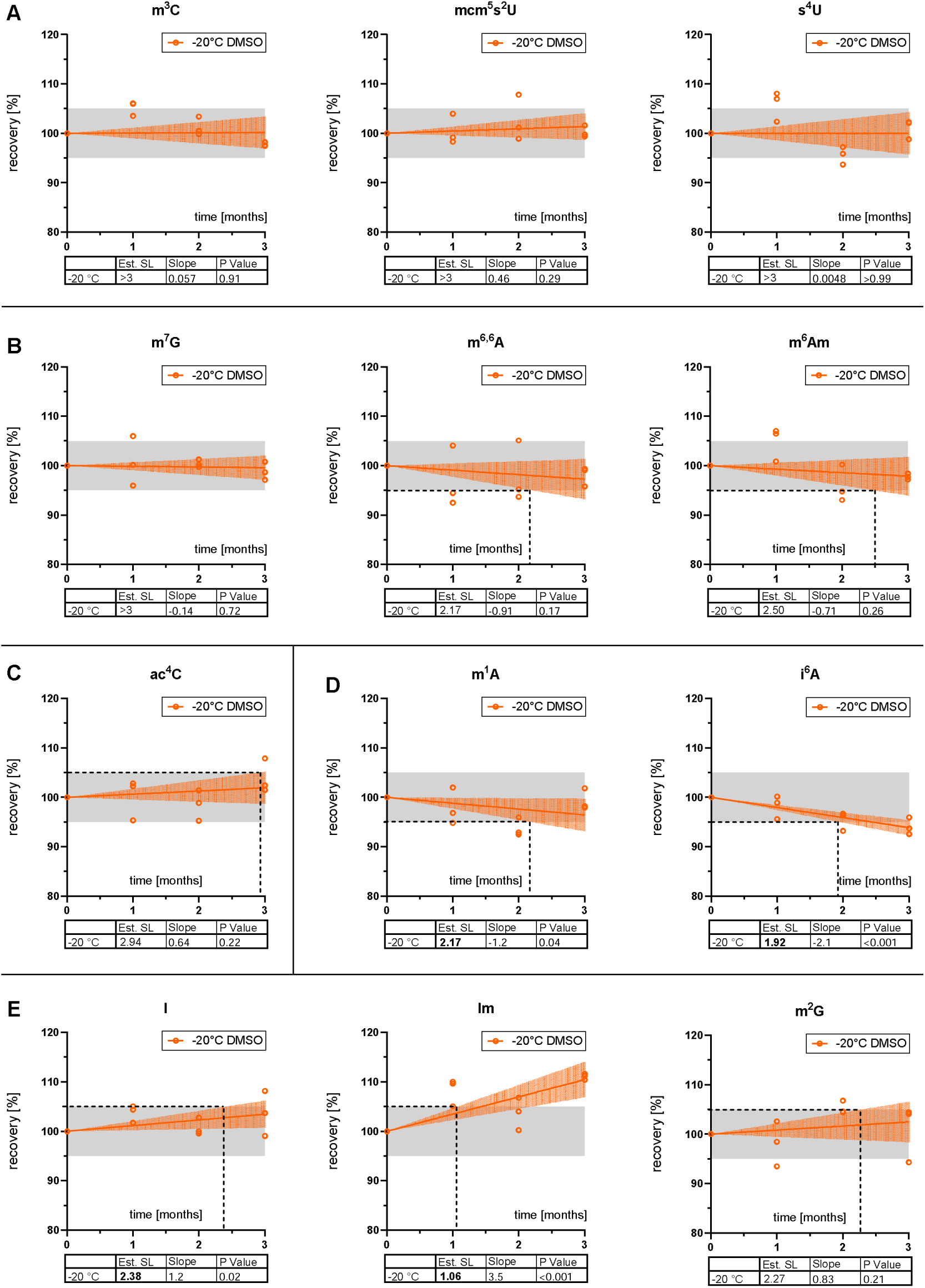
Stability of nucleoside stock solutions in DMSO at -20□°C over three months. Recovery was determined by comparing the UV area after storage to the initial value and normalized to 100%. The grey shaded region indicates the acceptance criterion of ±5% deviation (95–105%). Coloured shaded bands represent the 95% confidence interval of the linear regression. Statistical parameters (slope, p-value, and estimated shelf life (Est. SL)) are provided in the inset tables.

Notably, ac C remained stable over 3 months despite undergoing three freeze-thaw cycles. While its freeze-thaw stability in water remains untested in this setup, these results confidently validate DMSO as a secure storage solvent for ac C under fluctuating temperature conditions (Figure 6C).

### Standard operating procedure (SOP) for handling synthetic nucleoside standards

This SOP provides experimentally derived recommendations for the storage, handling and analytical assessment of modified nucleosides used as analytical standards. The goal is to ensure compound integrity, accurate quantification, and reproducible results. These recommendations apply to analytical standards analyzed under controlled laboratory conditions and are intended for LC-MS analytical workflows.

1. Purity check of synthetic nucleosides

Synthetic nucleosides are purchased in solid form with a declared purity. It is crucial to verify the exact purity and identity of these standards, as they are not always strictly intended for quantitative purposes by the manufacturer.

- Preferred Method (qNMR): If >10 mg of powder is available, purity should ideally be determined using quantitative NMR (qNMR, 2 replicates à 5 mg).
- Alternative 1 (UV Quantification): If qNMR is not feasible but the extinction coefficient (ε) is known, purity can be verified by dissolving/diluting the standard and measuring it via UV spectroscopy. Crucial: All dissolution and dilution steps must be performed exclusively in glass containers to prevent material leakage (leachables), which can distort UV readings. Note that the literature extinction coefficient typically applies only to water as the solvent and a specified pH (Table S2 and S9).
- Alternative 2 (LC-UV or LC-MS fallback): As a last resort, LC-UV can be used to check for the presence of impurities. However, this method does not allow for calculation of purity as ε from the impurity might be different from the analytes’ ε. The same applies to ionization efficiency and thus MS response.

3. Preparation of new nucleoside stock solutions

- Solvent and weighing: The weighed mass should be >1 mg (even when using an ultra-microbalance). Water or DMSO (to increase stability) should be used as solvents.
- Container selection: Glass containers are superior to plastic, based on our findings regarding material leakage and evaporation.
- Quality check: Freshly prepared stocks must be verified. If an extinction coefficient is known, the experimental extinction coefficient of the stock must not deviate by more than 5 % from the literature value. If no extinction coefficient is known (or UV analysis is unavailable): Stocks must be prepared in triplicates. The measurement values obtained via LC-UV or LC-MS must have a relative standard deviation (RSD) of <5 % between the triplicates.
- Evaporation: To monitor evaporation over time, the initial weight of the storage vial containing the stock solution must be measured and documented.

4. Storage temperature:

- For long-term storage, -80 °C or -20 °C should be selected depending on the specific compound.
- Handling precautions: All compounds should be protected from direct light during storage and handling. Prolonged exposure to ambient air should be minimized.
- Freeze-thaw cycles: Freeze-thaw cycles must be avoided, especially for the nucleosides highlighted in Table 2, which exhibit significant instability during short-term storage at room temperature (RT) and 8 °C.
- Evaporation monitoring: To track evaporation, the weight of the standard vial should be documented before and after each use.
- Nucleoside solutions must be freshly prepared once their designated shelf life (see Table 2) has expired. For nucleosides with identified degradation products, routine screening for these specific products must be performed.

5. Documentation and quality control:

Storage temperature, storage duration, solvent, container material, and preparation date must be documented for each compound. Freshly prepared solutions should be clearly labelled with the date and used within the defined timeframe.

6. Limitations

- Study Duration: The underlying stability study only covers up to 12 months for water and up to 3 months for DMSO. Therefore, no shelf life beyond these periods can be safely assumed.
- Freeze-thaw cycles: Stability in water was not tested over multiple freeze-thaw cycles. While it is reasonable to hypothesize that nucleosides insensitive to RT or 8 °C might also withstand freeze-thaw cycles, this has not been proven. Consequently, freeze-thaw cycles should be avoided for all nucleosides whenever possible.
- Conservative estimates: Shelf life estimates are based on worst-case calculations; therefore, some nucleosides may have longer stability than stated. Additionally, for certain nucleosides, storage at -20 °C proved to be more stable than at -80 °C. Always use the storage temperature that experimentally demonstrated the longer shelf life.
- Extrapolated Shelf Life: For nucleosides with an estimated shelf life <3 months, the exact duration cannot be considered precise due to a limited number of measurement points (in accordance with ICH guideline CPMP/ICH/2736/99). Therefore, rigorous quality control checks should be performed prior to each measurement, even when analyses are conducted within the estimated shelf life.

A graphical and table summary of this SOP is found in Figure 7 and Table 2.

**Figure 7.**
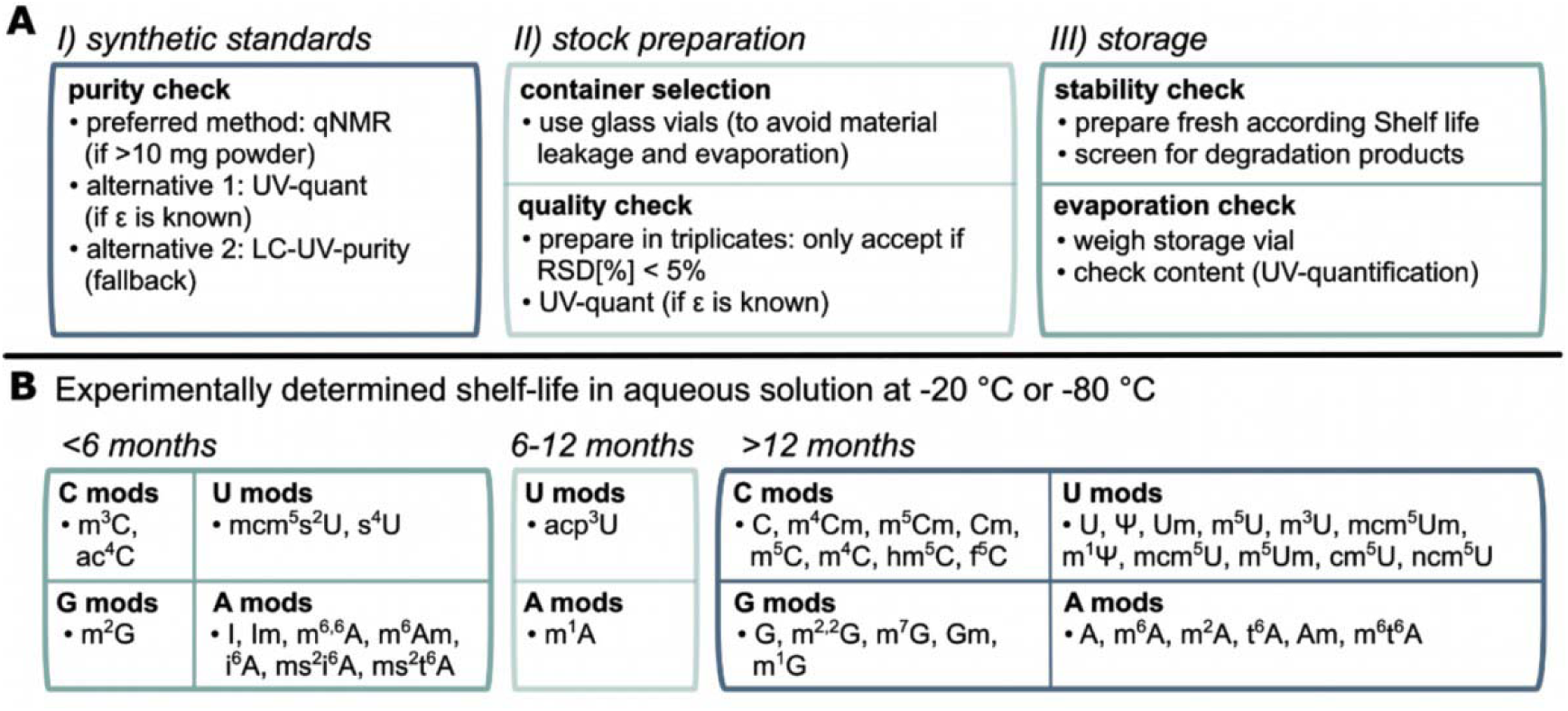
Recommended workflow and shelf-life categorization for synthetic nucleosides. **(A)** A three-step protocol detailing best practices for standard purity checks, stock solution preparation, and quality control during long-term storage. **(B)** Expected shelf-life in aqueous solution of different C, U, G, and A modifications, grouped into stability classes (<6 months, 6-12 months, and >12 months) to ensure standard integrity.

## Discussion

LC-MS analysis of modified nucleosides remains an important tool to study RNA modifications and define their elusive function and impact. This tool has substantially advanced since the first benchmark study by Crain, Pomerantz and McCloskey (38,39) and several guidelines and best practices were presented for both qualitative and quantitative analysis in the last years (11,36,40,41). In parallel, various studies appeared that study and exploit the chemical reactivity of modified nucleosides (reviewed in (42) and recent studies (43,44)) and even artefacts arising from unwanted chemical reactivity exist (13–15). In this manuscript we merge both lines of research, determine the stability of 44 ribonucleosides and present a practical guideline that summarizes the resulting best practices for synthetic standard handling. Yet, a few topics require further discussion and critical reflection.

We report the stability of nucleosides purchased from a single vendor using either pure water or DMSO as solvent. Several aspects of this statement require discussion. First, we want to discuss the fact that we used pure solvents without buffer addition. While it is possible, that some of the instabilities observed here can be avoided by using buffers, we decided against the use of buffers and salts. Our decision was guided by a recent report of deamination reactions of hypermodified uridine nucleosides in the presence of bicarbonate ions in the buffer (14). Based on this study and the fact that synthesis of nucleosides requires the use of various reagents, salts and solvents, it is possible that commercial nucleoside powders contain residual amounts of salts and reagents that can cause or accelerate the instabilities observed in this manuscript. Thus, stability could in principle be influenced by the source of the nucleoside powder. Due to the high price of nucleoside powders, we could not purchase nucleosides from various vendors and study whether the stability is impacted by the source of the nucleoside powder. As a practical consequence for the community, we designed Figure 7, and we strongly recommend assessing standard stability for each batch of prepared solution. Regarding instabilities, our quantum chemical calculations often predicted the experimentally-observed instability and thus we are confident that the instabilities reported in this manuscript are connected to the analyte’s chemistry and not necessarily to remaining contaminants in the powder. Potential examples are m^7^G, which most likely deglycosylates theoretically (Figure 4C) and degrades at -20 °C (but not -80 °C). Another important aspect that requires discussion is the fact that we studied nucleoside stability in diluted aqueous solution and not in the 1000-fold more concentrated stock solutions. In contrast, our DMSO stability studies were (due to its experimental need) conducted with the concentrated stocks. Therefore, we cannot experimentally prove that the better stability of some nucleosides in DMSO is caused by the solvent or the storage concentration. The requirement of water as a reactant (not solvent) in the degradation reactions (a) – (d) does, of course, provide one reason for different effective reactivities in water vs. DMSO solution. For acetal hydrolysis reactions such as the deglycosylation (a) it is well known that the reaction rate in water/DMSO mixtures decreases with increasing DMSO content, which may be seen as a general solvent effect on hydrolysis reactions (45,46). The combination of both effects may thus be responsible for the higher stability in DMSO solutions observed here.

Based on these considerations, we strongly recommend in-house assessment of nucleoside concentration by UV spectroscopy and degradation analysis by LC-UV. In this context, we want to emphasize that the storage vessel immediately impacts the quality and quantity of a nucleoside solution. Therefore, we recommend testing of vials towards evaporation, leaching of UV-active compounds and solvent/temperature stability. We want to emphasize that glass vials are superior to PP vials and that storage at -80 °C is not possible for any container if DMSO is the solvent.

Physical effects introduce a second layer of bias. Evaporation during storage, even in sealed cryo-vials, and currently unknown effects (I, Im, m^2^G), inflate apparent concentrations. Accurate quantification therefore requires a dual-control strategy: volumetric checks to correct for solvent loss and targeted monitoring of known degradation products. Another physical bias is observed for m^2^G, I and Im where we observe an increase in abundance that cannot be explained by evaporation alone. As no chemical process could generate these nucleosides in sealed storage, and the effect is less pronounced for I and Im, and even absent for m^2^G in DMSO, we hypothesize physical effects as the basis of this phenomenon. These findings highlight a broader analytical concern in nucleoside quantification. Namely, stability studies must consider not only degradation but also factors that may transiently influence apparent analyte abundance.

Further, we want to raise awareness to the instabilities observed at 20 °C. While these are of no practical consequence to the analytical chemist or core facility, they are a warning signal to the RNA scientist performing the biological study and potential other analytical tools. We found that ac^4^C quickly decomposes at room temperature. This observation may clarify conflicting reports in the epitranscriptomics literature surrounding ac^4^C in human mRNA. Several studies have reported widespread presence of ac^4^C across coding regions, attributing functional relevance to this modification (47,48). However, more recent investigations, such as (49), have not detected ac^4^C and attributed earlier findings to technical artefacts or misinterpretations. The pronounced chemical instability observed here for ac^4^C in aqueous solution, especially its rapid deacetylation to cytidine and subsequent deamination to uridine, offers a unifying hypothesis: ac^4^C may be genuinely present in mRNA but is lost prior to detection due to degradation during RNA preparation, handling, or storage. Thus, our results provide a compelling mechanistic explanation for the ongoing controversy surrounding the detection of ac^4^C in human mRNA. Similarly, studies of hypermodified uridines at position 34 of human tRNAs might suffer from an instability of the modification within its RNA environment. It is possible that this group of modifications degrades inside the cells, during RNA isolation or subsequent sample preparation. Chemical instability during RNA isolation or storage may therefore affect not only LC-MS detection but also sequencing-based technologies such as Illumina-based modification mapping or nanopore direct RNA sequencing, where modification detection likewise depends on the chemical integrity of the nucleoside.

With the SOP presented in this manuscript, together with recent guidelines on LC-MS analysis of ribonucleosides (11), key steps in the analytical workflow from nucleoside standard preparation to data acquisition can now be addressed systematically. This framework minimizes chemical degradation and physical concentration artefacts and thereby improves the robustness and inter-laboratory comparability of quantitative RNA modification measurements, providing a more reliable foundation for studying the biological roles of RNA modifications.

## Acknowledgements

The authors would like to thank Jens Wöhnert and his working group for providing the instrument (NMR). Special thanks to Elke Duchardt-Ferner for her time, instructions and advice.

## Data availability

The data supporting this article have been included as part of the Supplementary Information. Unless stated otherwise Supplementary Figures are found in Supplementary Information 1 (SI_1) or the Supplementary Tables (spread sheet). Reaction free energy considerations are detailed in Supplementary Information 2 (SI_2).

