## Supplement Information 1 for "Long-term stability of RNA nucleoside standards for accurate LC-MS quantification"

Extended Information on qNMR 2
Extended information on theoretical chemical analysis 2
Supplementary Figures 5
For Supplementary Tables S1-S9, please see the attached Excel sheet

**Extended information on qNMR of modified nucleosides**

To achieve results with high metrological quality and the required accuracy, reference standards with certified purity are used [33]. Examples of commonly used $^1$H NMR certified reference standards for internal calibration are listed in Table S5. These certified standards can be used for high-precision measurements with exceptional accuracy and an overall uncertainty as low as 0.15 % [34].

One potential issue when using aprotic and/or hygroscopic solvents, such as DMSO-$d_6$, is that acidic or alkaline protons may lead to broad signals in the spectra, potentially affecting spectral resolution and quantification accuracy. As shown in Figure S4A, the acidic internal calibrant maleic acid (MA) affects the water signal in DMSO-$d_6$, causing a shift toward higher ppm values and signal broadening. This distortion complicates spectral processing and may lead to biased peak integration and quantification, as seen in the case of 2′-*O*-methyluridine (Um), where signals nearby the broad water peak are directly affected. These issues are not limited to internal calibrants, as acidic or alkaline nucleosides can exhibit similar effects. For instance, as illustrated in Figure S4B, the spectrum of 3-(3-amino-3-carboxypropyl)uridine (acp$^3$U) also shows broad signals. Consequently, to ensure optimal spectral resolution and quantification accuracy, we selected dimethylsulfone (DMS) as the internal calibrant of choice for all subsequent experiments. All subsequent qNMR experiments followed the protocol outlined in the Materials and Methods section and Table S5, beginning with validation of the method by quantifying the purity of DMS using certified MA as an internal calibrant. The results – 99.89% purity in $D_2O$ with an RSD of 0.23 % and 99.98% purity in DMSO-$d_6$ with an RSD of 0.46% – confirmed both the accuracy and high precision of the qNMR method (Figure 1B).

**Extended theoretical analysis of degradation pathways**

The degradation pathways for nucleosides analyzed include (a) the deglycosylation reaction with water to yield the free nucleobase and ribofuranose; (b) the hydrolytic deamination of cytosine derivatives to yield the respective uracil bases and ammonia; (c) the hydrolytic deacetylation of 4-acetylaminocytosine derivatives to yield acetic acid and the respective cytosines; and (d) hydrolytic desulfurization of C2-thiocarbonyl derivatives of cytosine and uracil to yield hydrogen sulfide and the respective parent compounds. Detailed description of the used methods is summarized in Supplementary Information 2 (SI_2). Reaction free energies ($\Delta G_{rxn}$, in kJ mol$^{-1}$) for all four reaction types are obtained by subtracting the free energies $\Delta G_{298}$ of all reactants from those of the products (Table S8). For most of the nucleosides studied here the deglycosylation reaction is endergonic by 3 - 11 kJ mol$^{-1}$ The sole exceptions are the non-native 2-thiocarbonyl derivatives 2-thio-3-methylcytidine (m$^3$s$^2$C) and

2-thio-3-methyluridine ($m^3s^2U$) with small negative (-0.3 - -2.0 kJ $mol^{-1}$) reaction energies (Figure S13A). It is, from a general point of view, comforting to see that reaction energies are least favorable for canonical nucleosides such as uridine (U) and cytidine (C), which simply speaks for the stability of these systems.

The reaction free energies for the hydrolytic deamination of cytosine derivatives are collected for the full nucleoside as well as the respective free bases. For cytidine (C) and cytosine (c) as the reference systems of this group the deamination is moderately endergonic by 3.7 - 4.8 kJ $mol^{-1}$. While moderately exergonic reactions are predicted for some of the C5-substituted cytosine derivatives, the reaction becomes quite favorable for 3-methyl substituted cytosine derivatives. These may, depending on solution p*H* values, either be present in their neutral imino or protonated iminium ion form. For the neutral forms analyzed here the reaction energies amount to -26 - -31 kJ $mol^{-1}$. The impact of amino group protonation was studied in more detail for 3-methylcytosine ($m^3c$) as the most relevant parent system. While hydrolytic deamination is exergonic by -30.5 kJ $mol^{-1}$ for the neutral imino form shown in Figure S13B, the reaction energy is reduced to -19.5 kJ $mol^{-1}$ for the protonated iminium ion form (see SI for details). Deamination is found to be least favorable for adenosine (A) and its free base adenine (a). In more general terms, the deamination energetics of nucleosides and their free base equivalents are closely similar, which points to a rather limited influence of the N1 substituent on the deamination process.

Hydrolytic de-acetylation of cytosine derivatives carrying an acetyl group at the C4 amino group is exergonic for all nucleosides and nucleobases studied here (Figure S14A). The driving force for de-acetylation amounts to -10.2 kJ $mol^{-1}$ for cytidine itself, which is in accordance with our experimental data described in Figure 4D. It becomes notably larger for C5-substituted cytidine derivatives. The most favorable reaction energy is found for the 5-methyl substituted system ($m^5ac^4C$) with $\Delta G_{rxn}$ = -24.9 kJ $mol^{-1}$. These results open room for speculation: Is 4-acetyl-5-methylcytidine a non-natural RNA modification, or is it natural but due to its instability undetectable in current analytical systems?

Reaction energies for hydrolytic de-sulfurisation as the fourth degradation pathway analyzed here are shown in Figure 4D and in Figure S14B in a graphical manner. De-sulfurisation is found to be energetically favorable for all systems studied here with reaction energies ranging from -2.2 kJ $mol^{-1}$ for 4-thiouridine ($s^4U$) to -23.0 $mol^{-1}$ for the non-native 2-thio-3-methyluridine ($m^3s^2U$). De-sulfurisation energies differ, in selected cases, quite significantly between nucleosides and free bases. A point in case is 2-thiouridine ($s^2U$) the natural derivative of uridine with $\Delta G_{rxn}$ = -16.9 kJ $mol^{-1}$ as compared to 2-thiouracil ($s^2u$) with $\Delta G_{rxn}$ = -9.5 kJ $mol^{-1}$ (Table S8). Conformational analysis of the reactant and product nucleosides of the de-sulfurisation reaction indicates that this is most likely due to differences in hydrogen bonding interactions between the ribosyl hydroxy substituents and the C2 carbonyl resp. thiocarbonyl group.

Finally, reaction energies were also calculated for the two competing pathways for the decomposition of $s^4U$ shown in Figure 4D. For the free base of this latter system oxidative dimerization employing triplet oxygen as the reaction partner to yield the respective $s^4U$ dimer and water as the products is highly exothermic at $\Delta G_{rxn}$ = -101.6 kJ/mol, whereas hydrolytic loss of sulfur is only marginally exergonic ($\Delta G_{rxn}$ = –3.8 kJ/mol). However, in dilute aqueous solution there is an abundance of water available as a reactant, while the availability of dissolved oxygen may, together with the requirement of radical-chain initiation, limit the role of oxidative dimerization. That the oxidative dimerization becomes less competitive at higher temperatures may also be due to the entropic requirements of a reaction turning three separate reactants into two products. This complex interplay of individual factors apparently generates a scenario, where both pathways coexist in parallel. The hydrolytic deprenylation of $i^6A$ shown in Figure S7 is, in comparison, thermochemically quite unfavorable (with $\Delta G_{rxn}$ = +24.5 kJ/mol for the nucleoside and +31.6 kJ/mol for the free base), which makes this process at least in the hydrolytic mechanism considered here quite unlikely.

**Supplementary Figures**

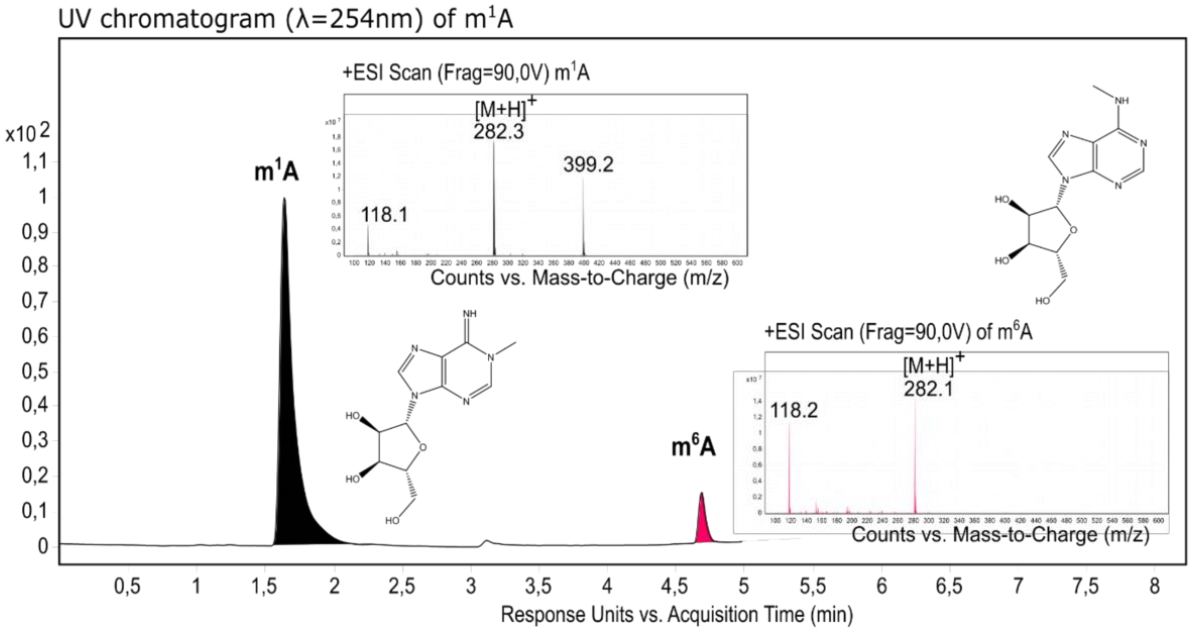


**Figure S1.** UV-chromatogram and low-resolution mass spectra of a fresh solution of 1-methyladenosine ($m^1A$). Fresh prepared $m^1A$ solution already shows impurity with $m^6A$.

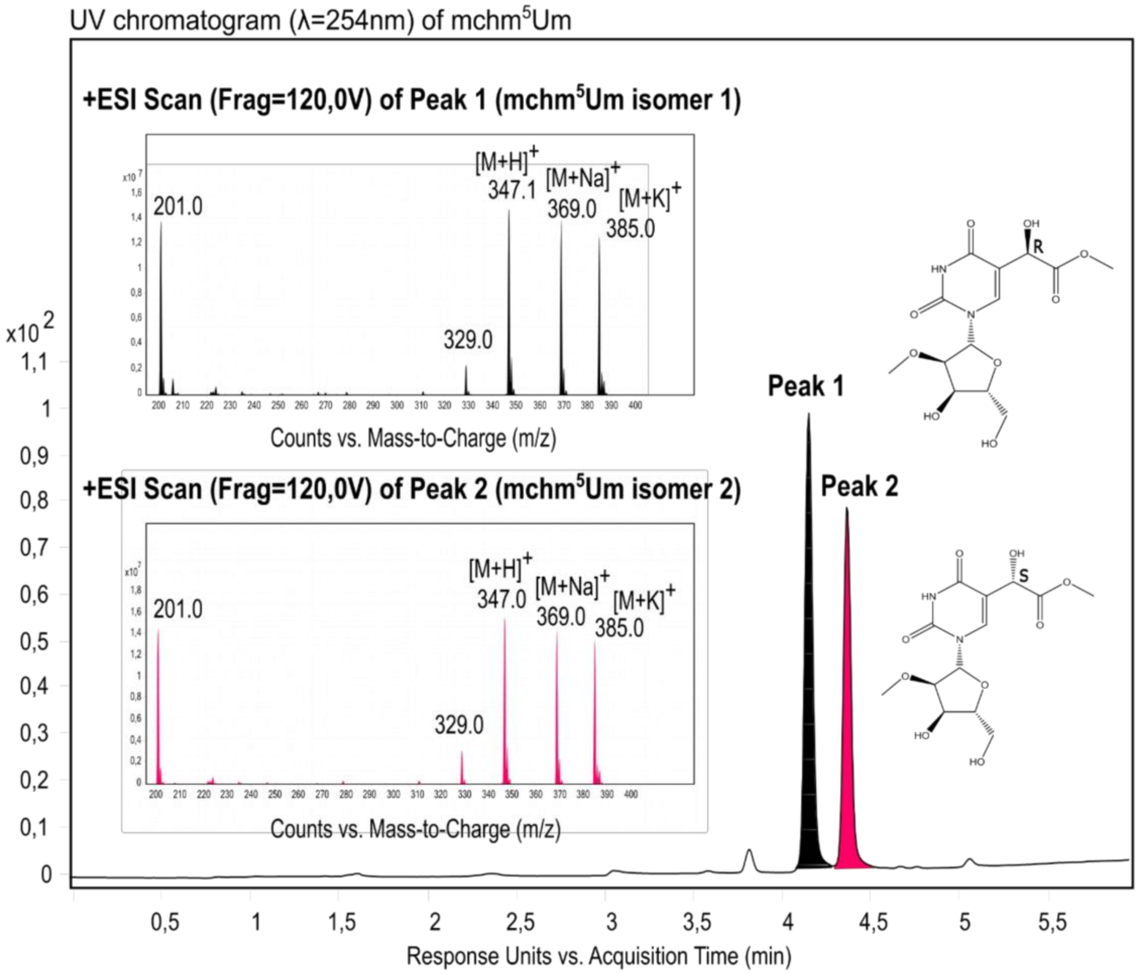


**Figure S2.** UV-chromatogram and low-resolution mass spectra of 5-(carboxyhydroxymethyl)-2'-O-methyluridine (mchm$^5$Um). This compound is sold as a mixture of the two diastereomers.

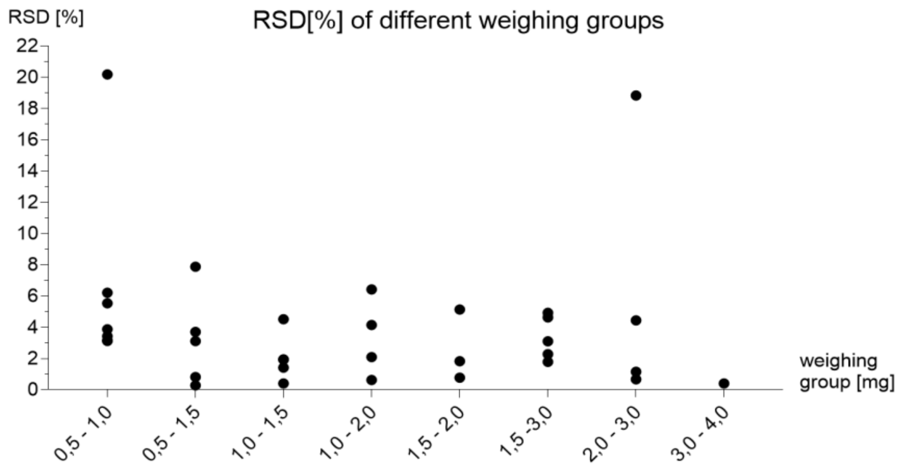


**Figure S3.** Relative standard deviation (RSD) of nucleosides binned into 8 different weighing groups. Except for 2 nucleosides, the RSD was below 10 %.

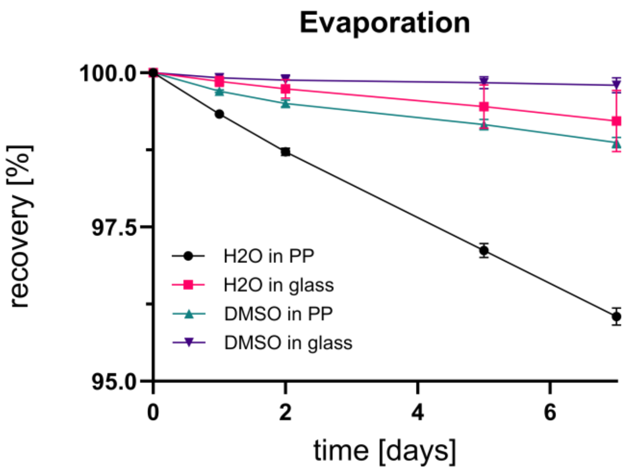


**Figure S4.** Gravimetrically determined recovery (%) of $H_2O$ and DMSO in polypropylene (PP) cryo-vials and glass containers at 60 °C over 7 days to assess evaporative losses. Values represent mean ± SD; recovery is compared to the initial value and normalized to 100 %. For $H_2O$, a pronounced decrease in recovery was observed in cryo-vials, whereas losses in glass were markedly lower (remaining at ~99 %). These data indicate substantially higher evaporation and/or permeation of water in PP cryo-vials compared to glass.

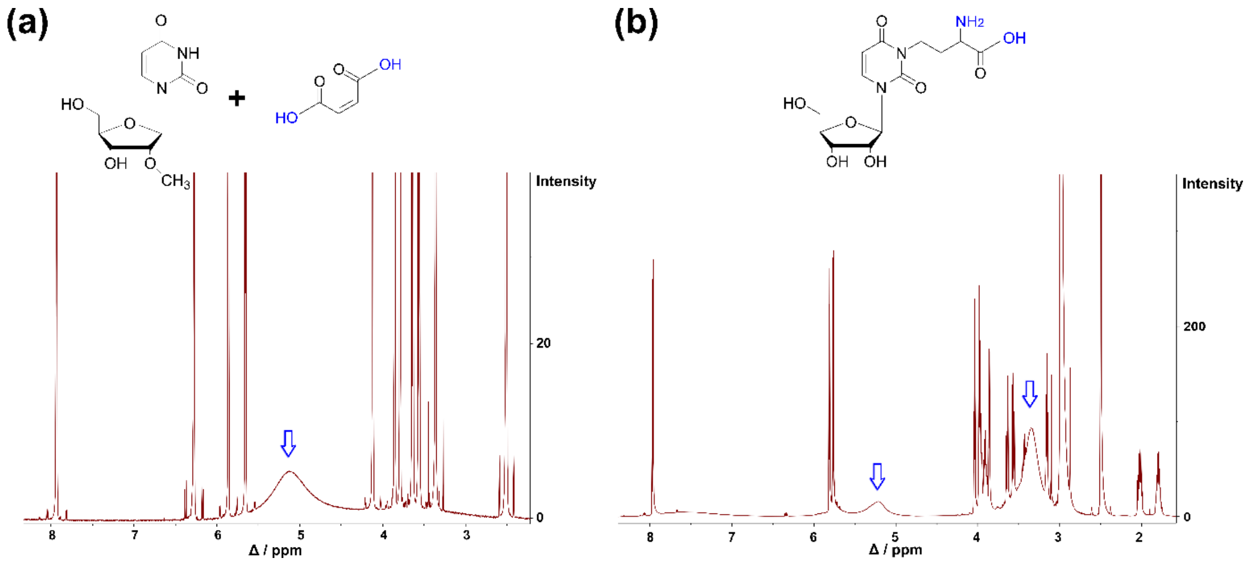


**Figure S5.** $^{1}H$ NMR spectra of nucleosides Um and acp$^{3}$U in DMSO-$d_6$. Broad and non-distinct water peaks are indicated by blue arrows. **(a)** $^{1}H$ NMR of Um and maleic acid (MA – internal calibrant) in DMSO-$d_6$. **(b)** $^{1}H$ NMR of acp$^{3}$U and dimethylsulfone (DMS – internal calibrant) in DMSO-$d_6$.

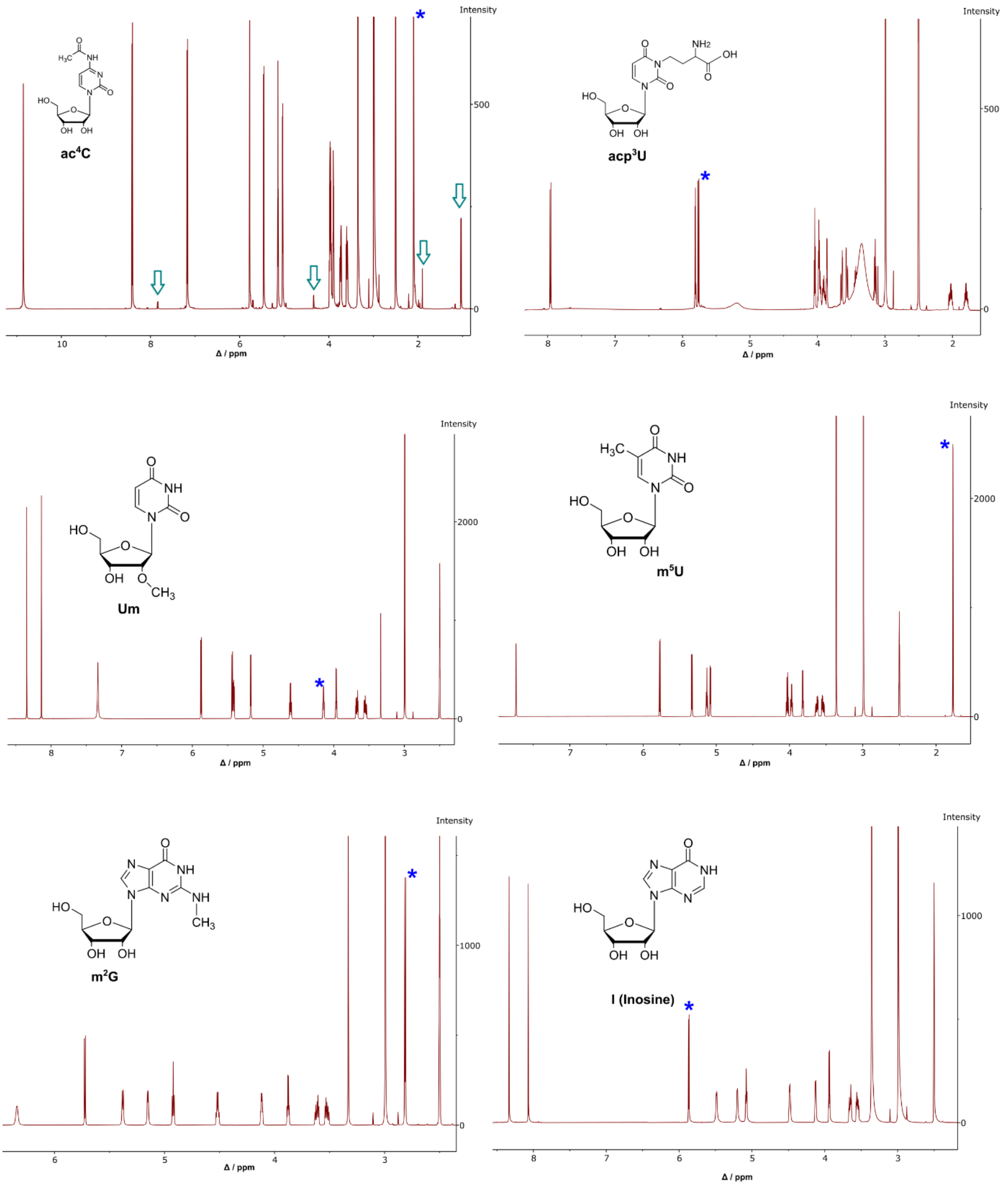


**Figure S6.** $^{1}H$ NMR of the six nucleosides (ac$^4$C, acp$^3$U, Um, m$^5$U, m$^2$G and I) being quantified in the study. Blue asterisks indicate signals which have been used for quantification. Green arrows highlight non-assignable protons in the spectrum of ac$^4$C.

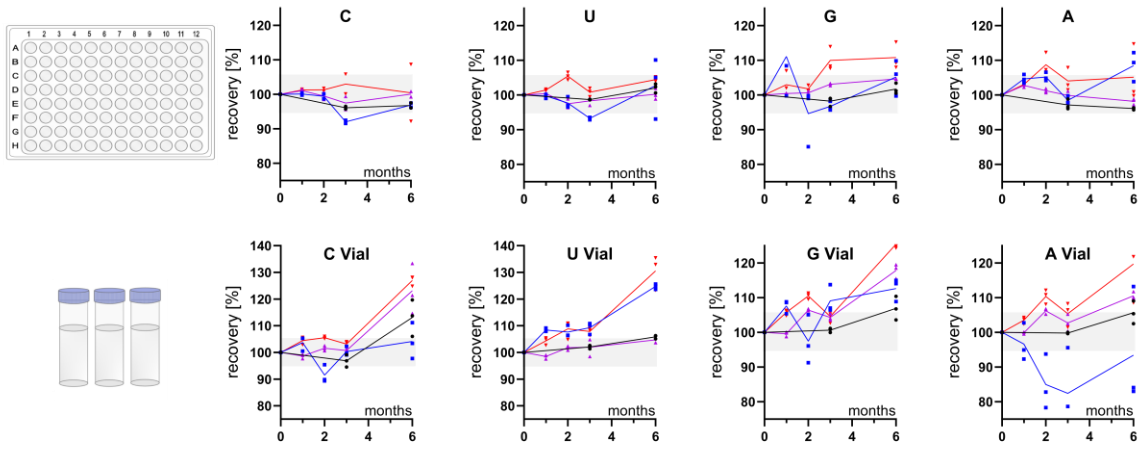


**Figure S7.** Stability of canonical nucleosides following storage in well-plates (top) and cryo-vials (bottom). Data points are color-coded by storage temperature: -80 °C (black), -20 °C (blue), 8 °C (purple), and 20 °C (red). The apparent increase in recovery observed for samples stored in cryo-vials indicates a higher rate of solvent evaporation compared to well-plates.

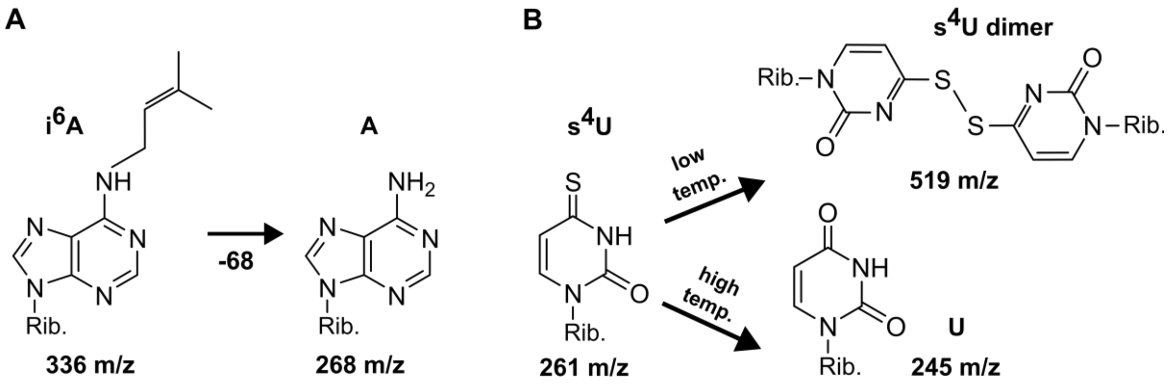


**Figure S8.** Temperature-dependent degradation/dimerization. **(A)** 6-isopentenyladenosine ($i^6A$) decomposes to adenosine. **(B)** 4-thiouridine ($s^4U$) decomposes to uridine (U) and forms dimers (di-$s^4U$)

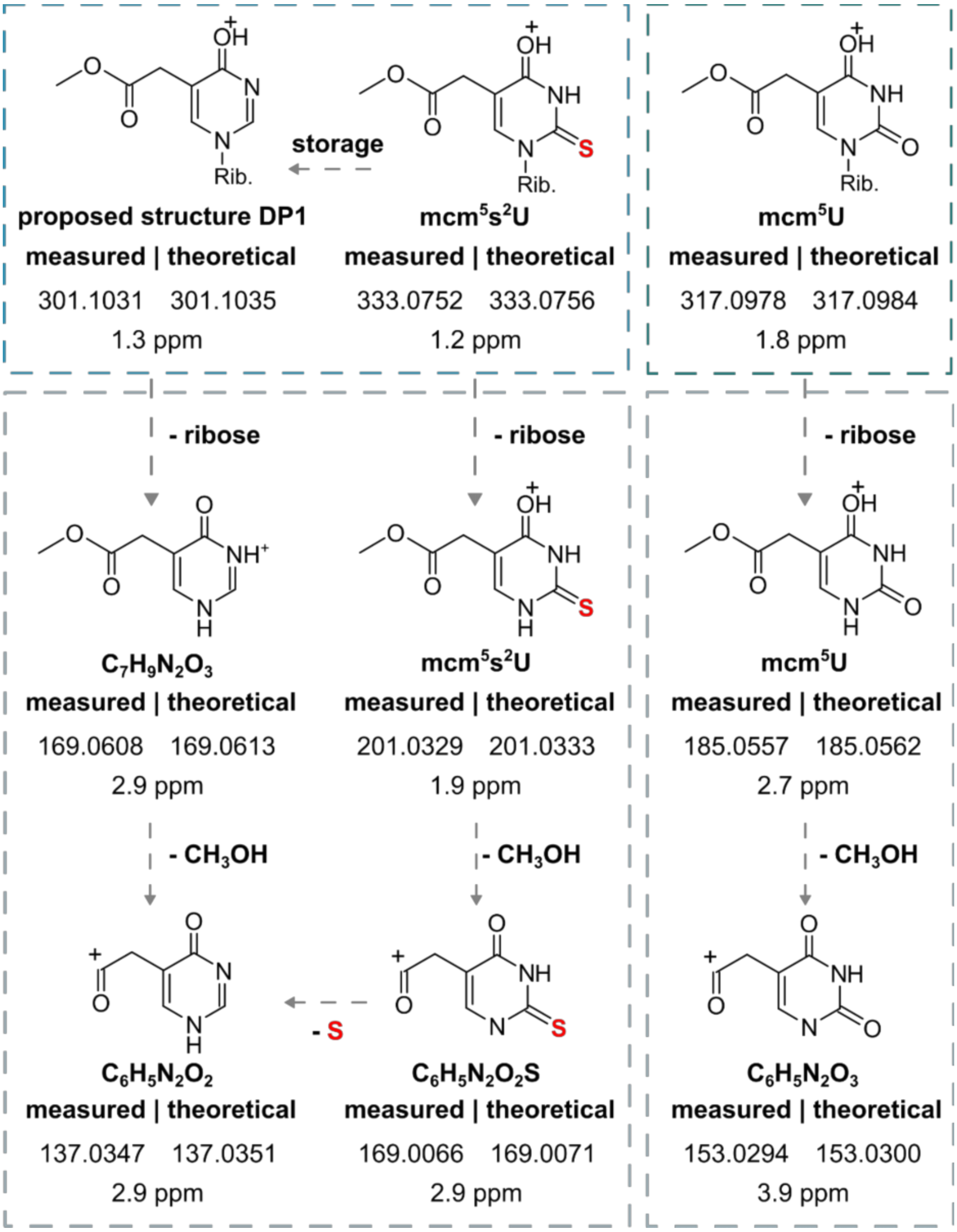


**Figure S9.** HRAMS fragmentation pattern (including MS2 and MS3 scans) of mcm$^5$s$^2$U, degradation product 1 (DP1) at m/z 301 and mcm$^5$U. Mass deviation of suggested structures is indicated in ppm.

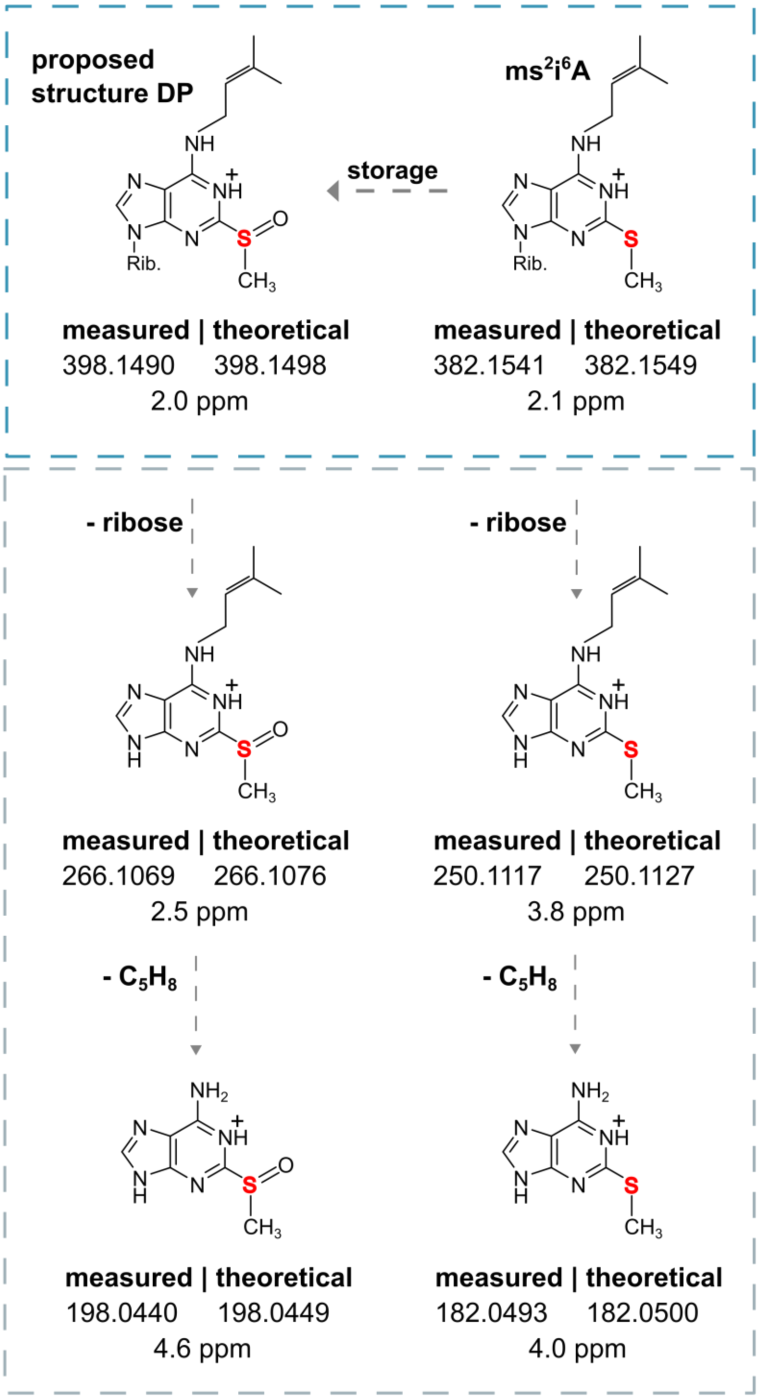


**Figure S10.** HRAMS fragmentation pattern (including MS2 and MS3 scans) of ms$^2$i$^6$A and degradation product (DP) at m/z 398. Mass deviation of suggested structures is indicated in ppm.

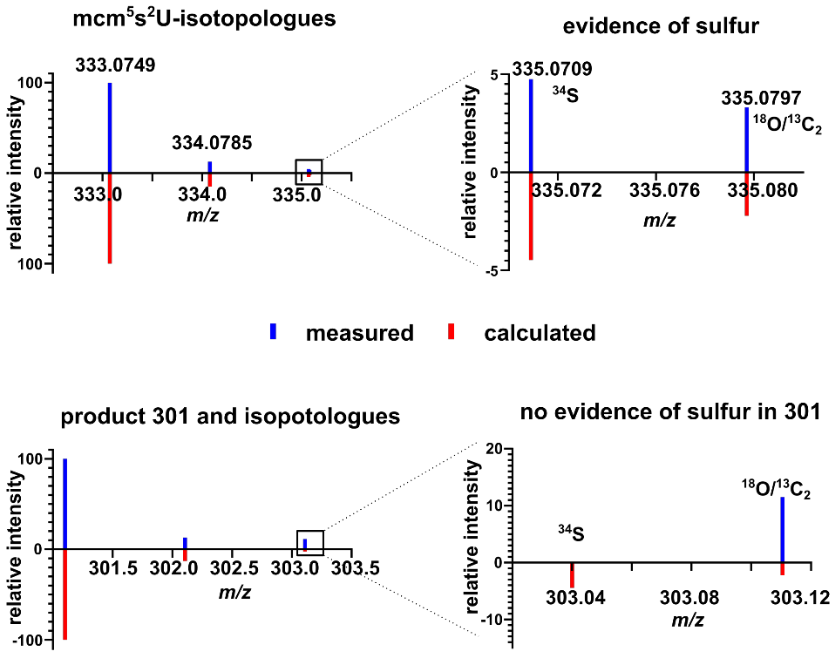


**Figure S11.** Isotopic distribution ($^{13}C$, $^{34}S$, $^{18}O$) of mcm$^5$s$^2$U and its degradation product 1 (DP1) at m/z 301. Experimental values were acquired via HRAMS and compared to theoretical values calculated from the natural abundance of isotopologues. While sulfur was clearly detected in mcm$^5$s$^2$U, DP1 shows no isotopic evidence of sulfur.

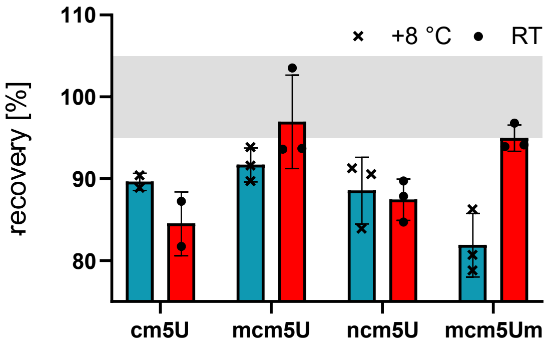


**Figure S12**. Stability of human U34 modifications in dependence of temperature after 6 months at 8 °C and room temperature.

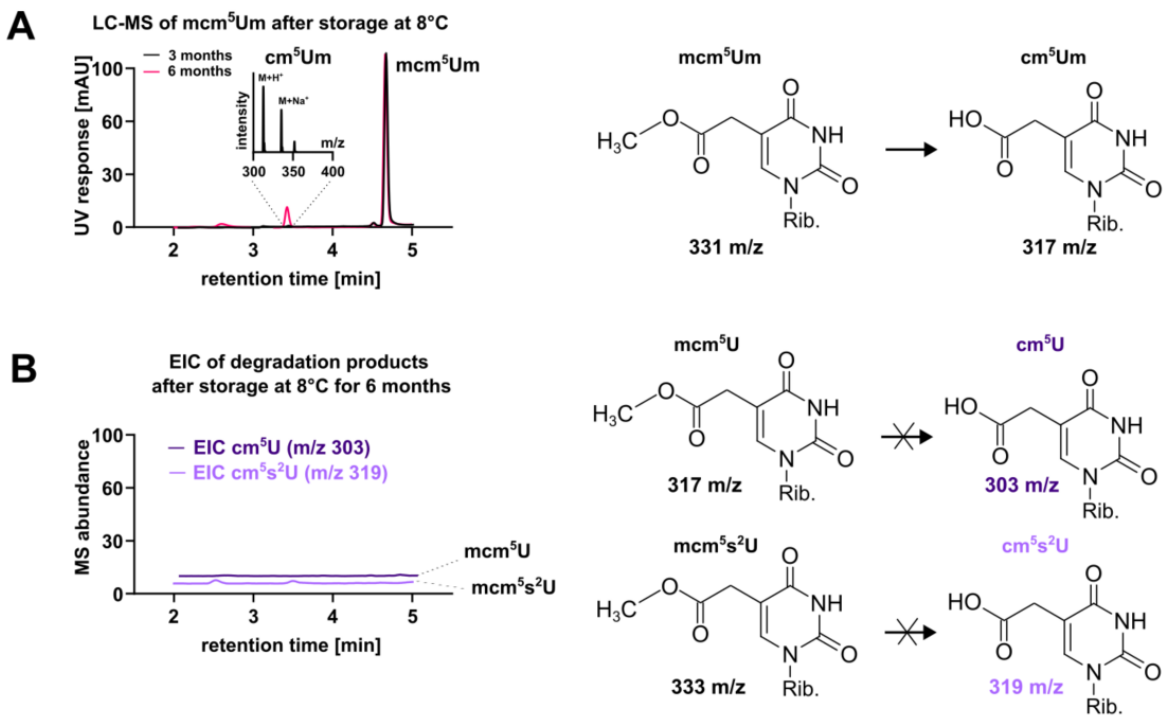


**Figure S13**. Degradation products from mcm5Um, mcm5U and mcm5s2U. **(A)** LC-UV-MS chromatograms of 5-methoxycarbonylmethyl-2'-O-methyluridine (mcm5Um) after storage at 8 °C for 3 months (black) and 6 months (pink). **(B)** EIC of the corresponding degradation products from mcm5U and mcm5s2U.

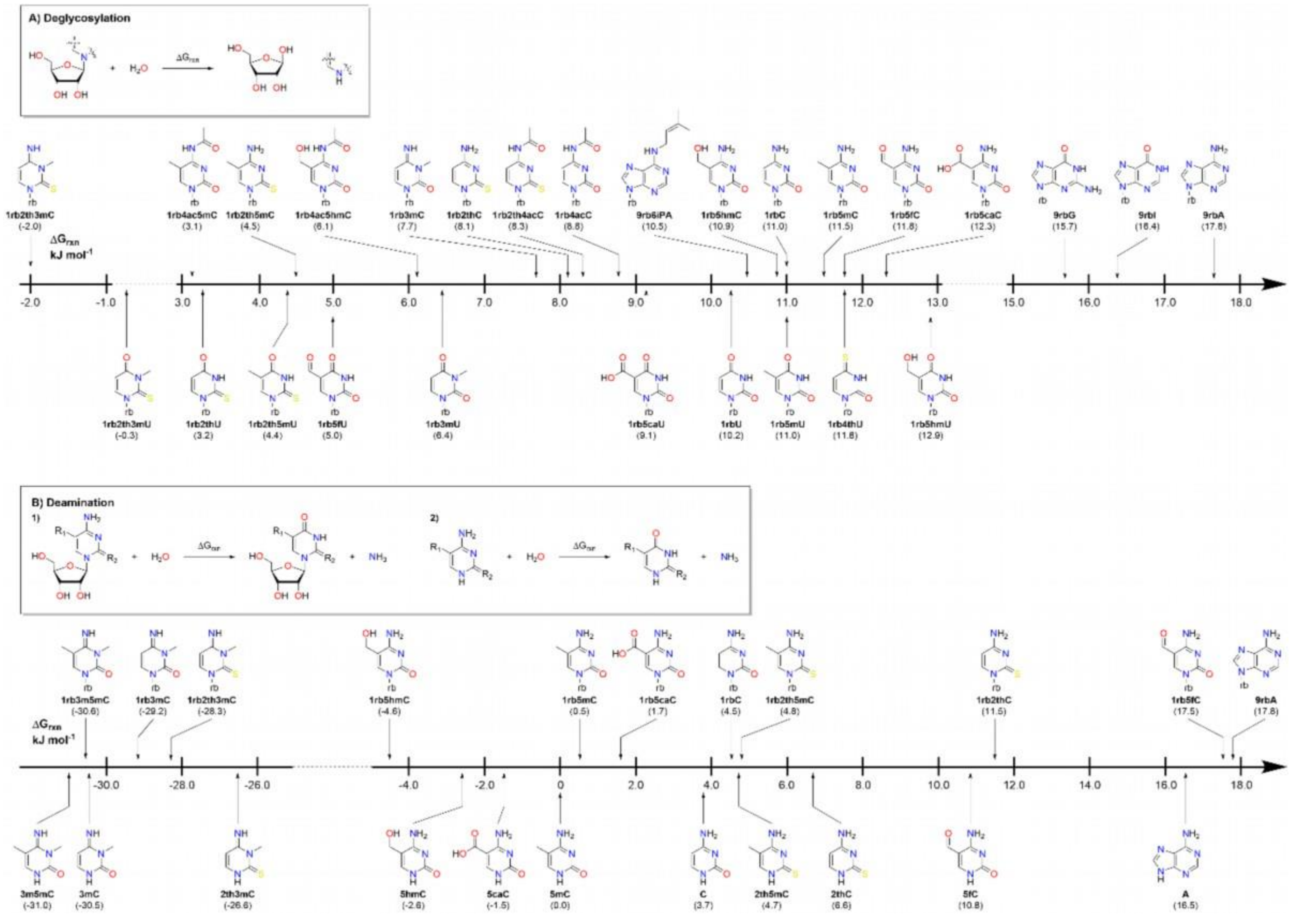


**Figure S14.** Reaction free energies for hydrolytic deglycosylation **(A)** and hydrolytic deamination **(B)** of different natural and non-natural nucleosides.

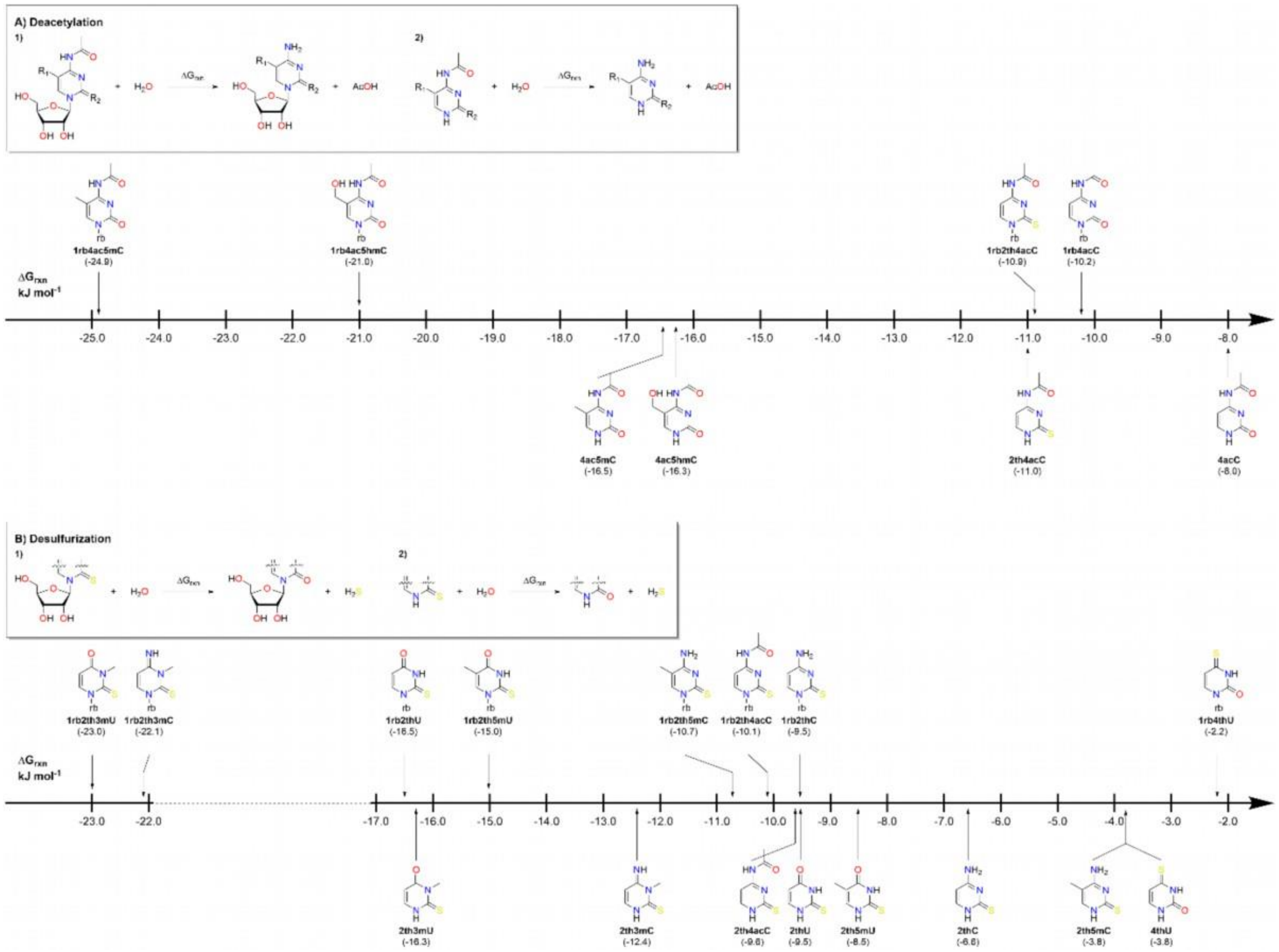


**Figure S15.** Reaction free energies **(A)** for hydrolytic deacetylation of C4-acetylated cytosine derivatives and **(B)** hydrolytic de-sulfurization of nucleosides and nucleobases with thiocarbonyl groups

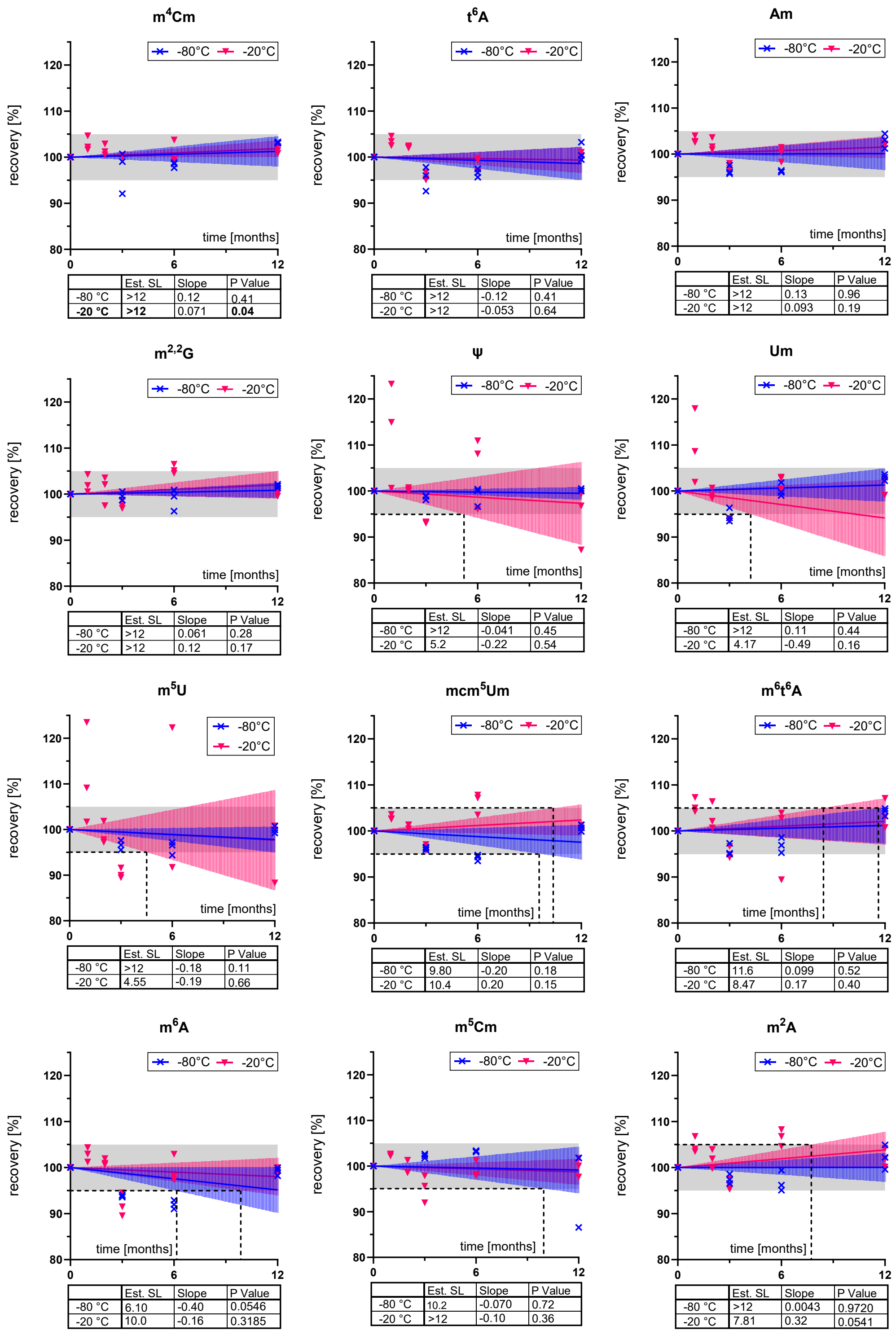


**Figure S16.** Nucleosides remain stable under both common storage conditions (-20 °C and - 80 °C). Shelf life was determined by comparing the UV peak area after storage to the initial value, normalized to 100 %. The grey shaded region indicates the acceptance criterion of ± 5 % deviation (95–105 %). Colored shaded bands represent the 95 % confidence interval of the linear regression. Statistical parameters (slope, p-value, and Est. SL) are provided in the inset tables.

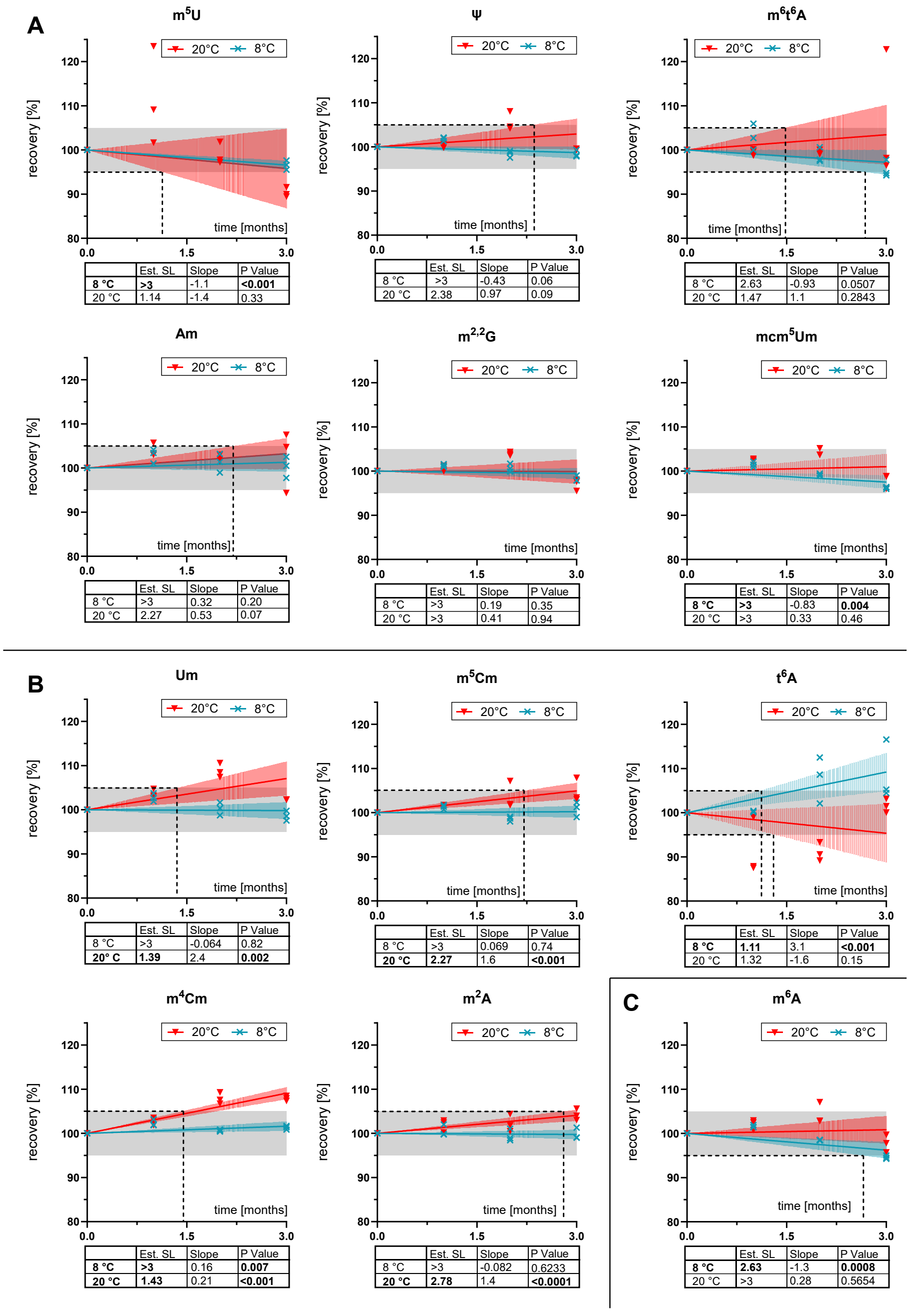


**Figure S17.** Short-term stability assessment (8 °C and RT) for nucleosides demonstrating stability at both common storage temperatures (-20 °C and -80 °C). Short-term shelf life was determined by comparing the UV peak area after storage to the initial value, normalized to 100 %. The grey shaded region indicates the acceptance criterion of ± 5 % deviation (95–105 %). Coloured shaded bands represent the 95 % confidence interval of the linear regression. Statistical parameters (slope, p-value, and Est. SL) are provided in the inset tables. **(A)** Nucleosides demonstrating stability at both RT and 8 °C. **(B)** Nucleosides exhibiting an apparent increase in recovery at RT or 8 °C. This effect is attributed to elevated solvent evaporation at these temperatures. Therefore, these compounds are still considered stable. **(C)** Stability profile of m$^6$A, which shows a slight but significant downward trend at 8 °C. However, with an estimated shelf life of > 2.5 months, it exhibits low sensitivity to elevated temperatures.

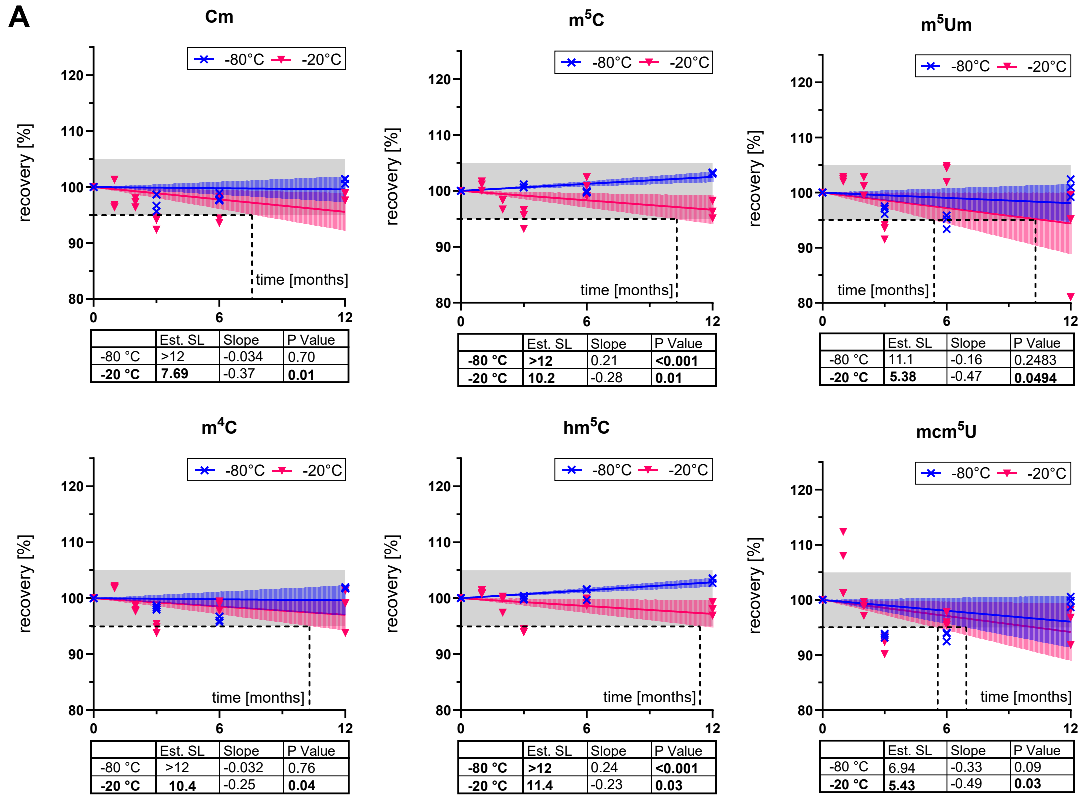


Cm

| | Est. SL | Slope | P Value |
|---|---|---|---|
| -80 °C | >12 | -0.034 | 0.70 |
| **-20 °C** | **7.69** | -0.37 | **0.01** |

m$^5$C

| | Est. SL | Slope | P Value |
|---|---|---|---|
| **-80 °C** | **>12** | 0.21 | **<0.001** |
| **-20 °C** | **10.2** | -0.28 | **0.01** |

m$^5$Um

| | Est. SL | Slope | P Value |
|---|---|---|---|
| -80 °C | 11.1 | -0.16 | 0.2483 |
| **-20 °C** | **5.38** | -0.47 | **0.0494** |

m$^4$C

| | Est. SL | Slope | P Value |
|---|---|---|---|
| -80 °C | >12 | -0.032 | 0.76 |
| **-20 °C** | **10.4** | -0.25 | **0.04** |

hm$^5$C

| | Est. SL | Slope | P Value |
|---|---|---|---|
| **-80 °C** | **>12** | 0.24 | **<0.001** |
| **-20 °C** | **11.4** | -0.23 | **0.03** |

mcm$^5$U

| | Est. SL | Slope | P Value |
|---|---|---|---|
| -80 °C | 6.94 | -0.33 | 0.09 |
| **-20 °C** | **5.43** | -0.49 | **0.03** |

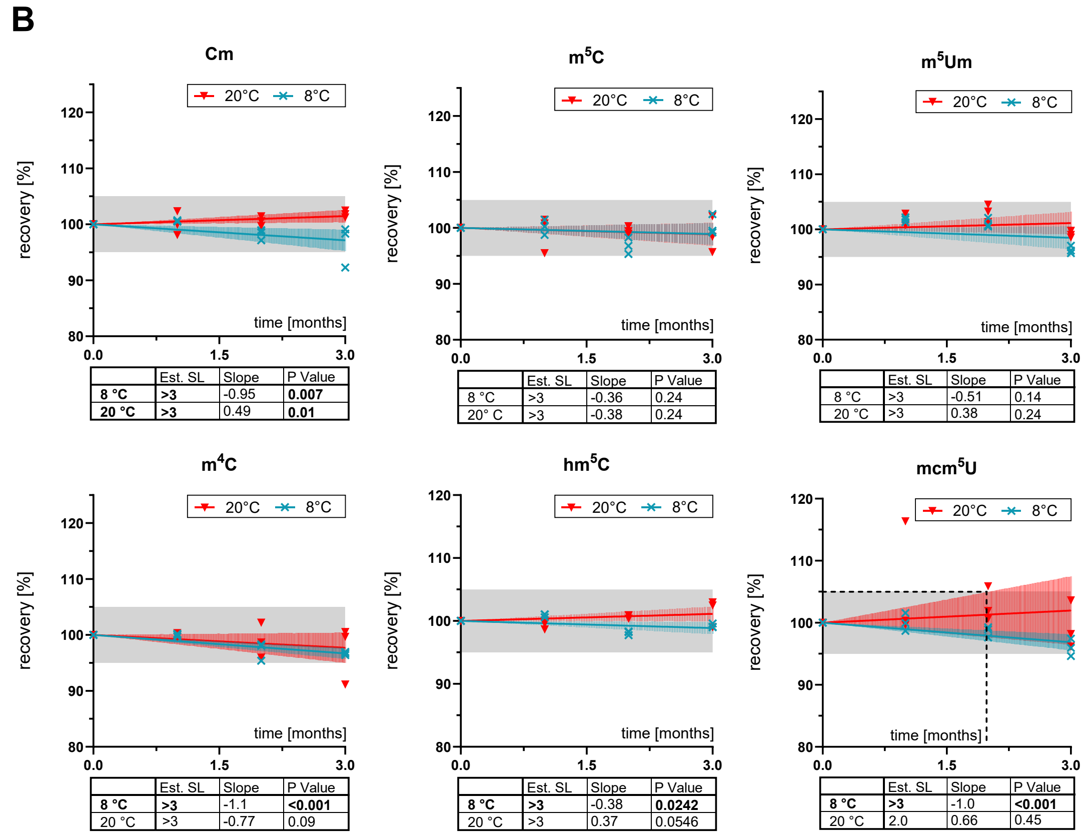


Cm

| | Est. SL | Slope | P Value |
|---|---|---|---|
| **8 °C** | **>3** | -0.95 | **0.007** |
| **20 °C** | **>3** | 0.49 | **0.01** |

m$^5$C

| | Est. SL | Slope | P Value |
|---|---|---|---|
| 8 °C | >3 | -0.36 | 0.24 |
| 20° C | >3 | -0.38 | 0.24 |

m$^5$Um

| | Est. SL | Slope | P Value |
|---|---|---|---|
| 8 °C | >3 | -0.51 | 0.14 |
| 20 °C | >3 | 0.38 | 0.24 |

m$^4$C

| | Est. SL | Slope | P Value |
|---|---|---|---|
| **8 °C** | **>3** | -1.1 | **<0.001** |
| 20 °C | >3 | -0.77 | 0.09 |

hm$^5$C

| | Est. SL | Slope | P Value |
|---|---|---|---|
| **8 °C** | **>3** | -0.38 | **0.0242** |
| 20 °C | >3 | 0.37 | 0.0546 |

mcm$^5$U

| | Est. SL | Slope | P Value |
|---|---|---|---|
| **8 °C** | **>3** | -1.0 | **<0.001** |
| 20 °C | 2.0 | 0.66 | 0.45 |

**Figure S18.** Long-term (-20 °C and -80 °C) and short-term (RT and 8 °C) shelf life for nucleosides demonstrating stability at -80 °C (First part). Shelf life was determined by comparing the UV area after storage to the initial value and normalized to 100 %. The grey shaded region indicates the acceptance criterion of ± 5 % deviation (95–105 %). Coloured shaded bands represent the 95 % confidence interval of the linear regression. Statistical parameters (Slope, P-value, and Est. SL) are provided in the inset tables. **(A)** Long-term stability at -80 °C and -20 °C, showing that these nucleosides are stable at -80 °C but unstable at -20 °C. **(B)** Short-term storage at RT and 8 °C, where all nucleosides remain stable.

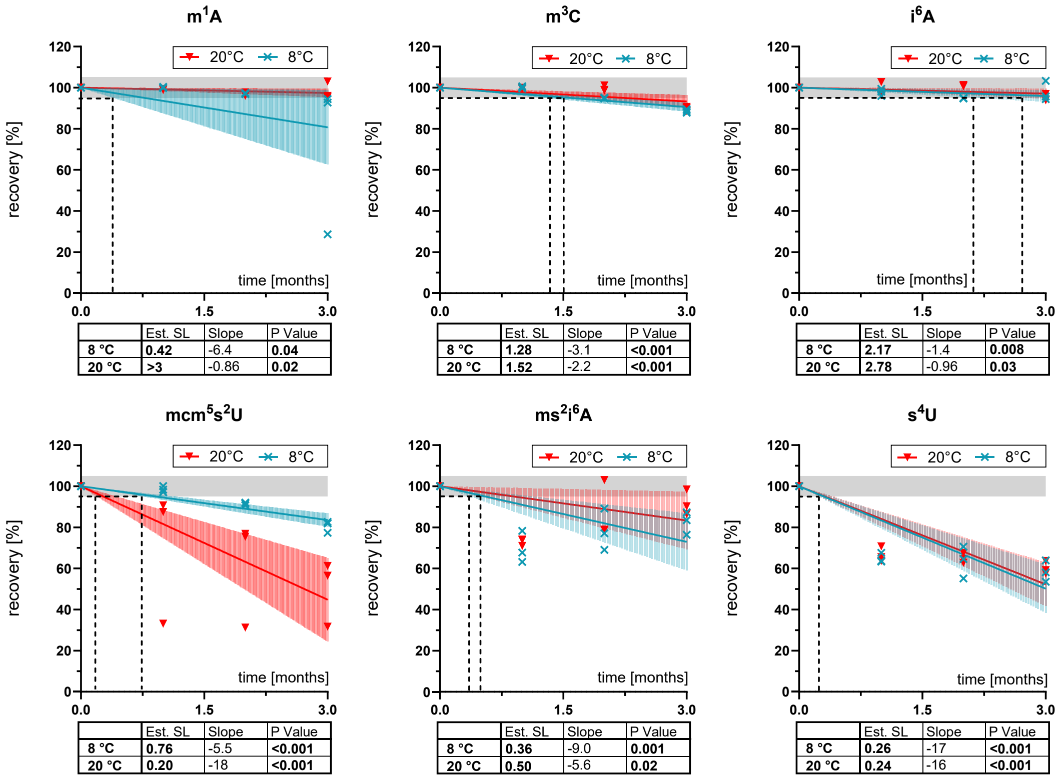


**Figure S19.** Short-term (RT and 8 °C) shelf life for nucleosides that were unstable at both common storage conditions (-20 °C and -80 °C) and showed degradation products in the UV chromatogram. Shelf life was determined by comparing the UV area after storage to the initial value and normalized to 100 %. The grey shaded region indicates the acceptance criterion of ± 5 % deviation (95–105 %). Coloured shaded bands represent the 95% confidence interval of the linear regression. Statistical parameters (Slope, P-value, and Est. SL) are provided in the inset tables. Long-term stability data (-20 °C and -80 °C) are presented in Figure 3 of the main manuscript.

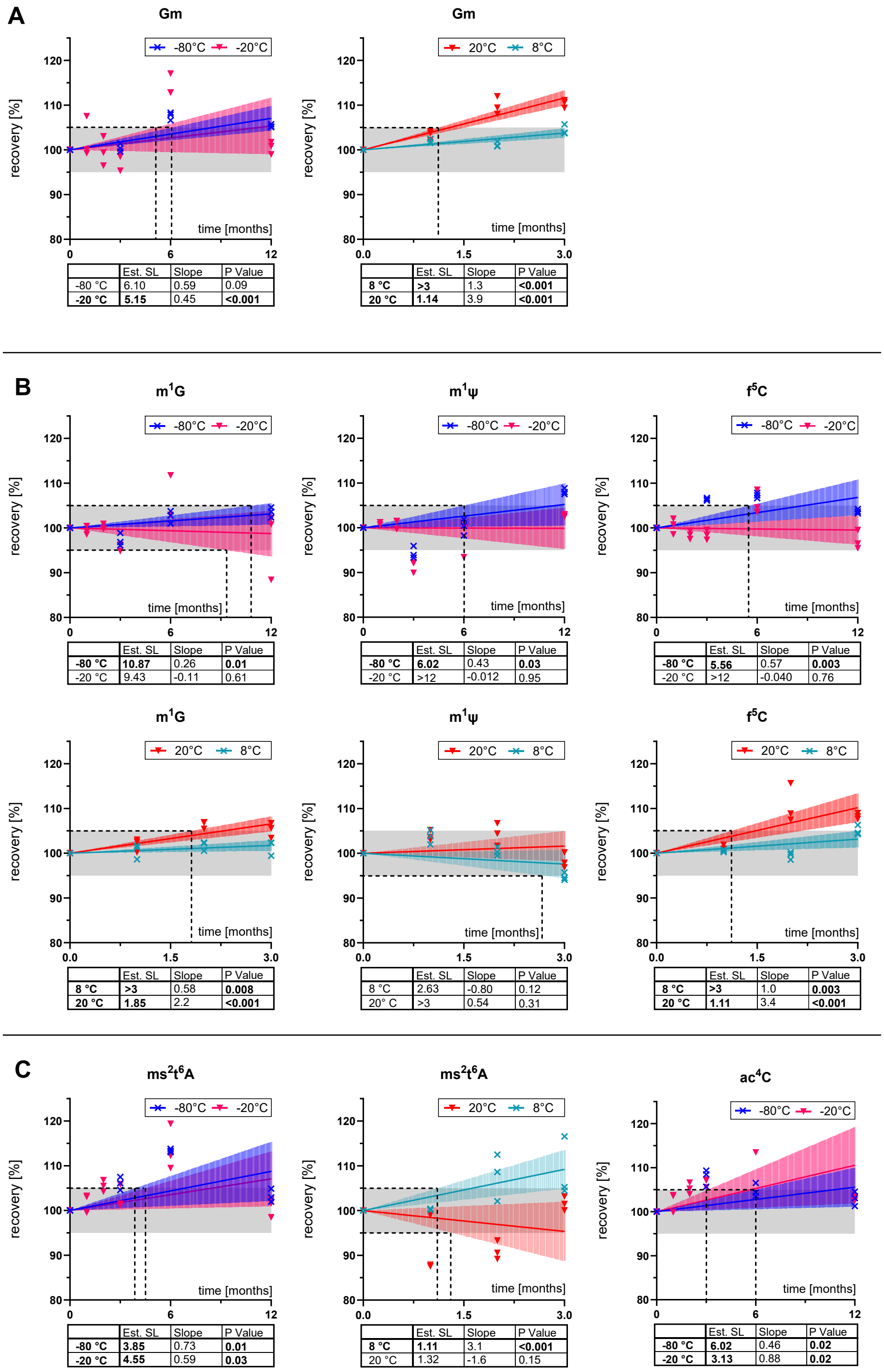


**A** Gm (-80 °C / -20 °C)

| | Est. SL | Slope | P Value |
|---|---|---|---|
| -80 °C | 6.10 | 0.59 | 0.09 |
| **-20 °C** | **5.15** | 0.45 | **<0.001** |

**A** Gm (8 °C / 20 °C)

| | Est. SL | Slope | P Value |
|---|---|---|---|
| **8 °C** | **>3** | 1.3 | **<0.001** |
| **20 °C** | **1.14** | 3.9 | **<0.001** |

**B** m1G (-80 °C / -20 °C)

| | Est. SL | Slope | P Value |
|---|---|---|---|
| **-80 °C** | **10.87** | 0.26 | **0.01** |
| -20 °C | 9.43 | -0.11 | 0.61 |

**B** m1ψ (-80 °C / -20 °C)

| | Est. SL | Slope | P Value |
|---|---|---|---|
| **-80 °C** | **6.02** | 0.43 | **0.03** |
| -20 °C | >12 | -0.012 | 0.95 |

**B** f5C (-80 °C / -20 °C)

| | Est. SL | Slope | P Value |
|---|---|---|---|
| **-80 °C** | **5.56** | 0.57 | **0.003** |
| -20 °C | >12 | -0.040 | 0.76 |

**B** m1G (8 °C / 20 °C)

| | Est. SL | Slope | P Value |
|---|---|---|---|
| **8 °C** | **>3** | 0.58 | **0.008** |
| **20 °C** | **1.85** | 2.2 | **<0.001** |

**B** m1ψ (8 °C / 20 °C)

| | Est. SL | Slope | P Value |
|---|---|---|---|
| 8 °C | 2.63 | -0.80 | 0.12 |
| 20° C | >3 | 0.54 | 0.31 |

**B** f5C (8 °C / 20 °C)

| | Est. SL | Slope | P Value |
|---|---|---|---|
| **8 °C** | **>3** | 1.0 | **0.003** |
| **20 °C** | **1.11** | 3.4 | **<0.001** |

**C** ms2t6A (-80 °C / -20 °C)

| | Est. SL | Slope | P Value |
|---|---|---|---|
| **-80 °C** | **3.85** | 0.73 | **0.01** |
| **-20 °C** | **4.55** | 0.59 | **0.03** |

**C** ms2t6A (8 °C / 20 °C)

| | Est. SL | Slope | P Value |
|---|---|---|---|
| **8 °C** | **1.11** | 3.1 | **<0.001** |
| 20 °C | 1.32 | -1.6 | 0.15 |

**C** ac4C (-80 °C / -20 °C)

| | Est. SL | Slope | P Value |
|---|---|---|---|
| **-80 °C** | **6.02** | 0.46 | **0.02** |
| **-20 °C** | **3.13** | 0.88 | **0.02** |

**Figure S20.** Long-term (-20 °C and -80 °C) and short-term (RT and 8 °C) shelf life for nucleosides categorized as unstable due to a statistically significant positive slope. Shelf life was evaluated by normalizing the UV peak area after storage to the initial baseline (100 %). The grey shaded region indicates the acceptance criterion of ± 5 % deviation (95–105 %). Coloured shaded bands represent the 95 % confidence interval of the linear regression. Inset tables summarize the statistical parameters (slope, p-value, and Est. SL). **(A)** Gm exhibits instability at -80 °C. **(B)** m$^1$G, m$^1$Y, and f$^5$C exhibit instability at -20 °C. **(C)** ms$^2$t$^6$A and ac$^4$C are unstable under both standard long-term storage conditions (-20 °C and -80 °C). Short-term stability data for ac$^4$C can be found in Figure 5 of the main manuscript.

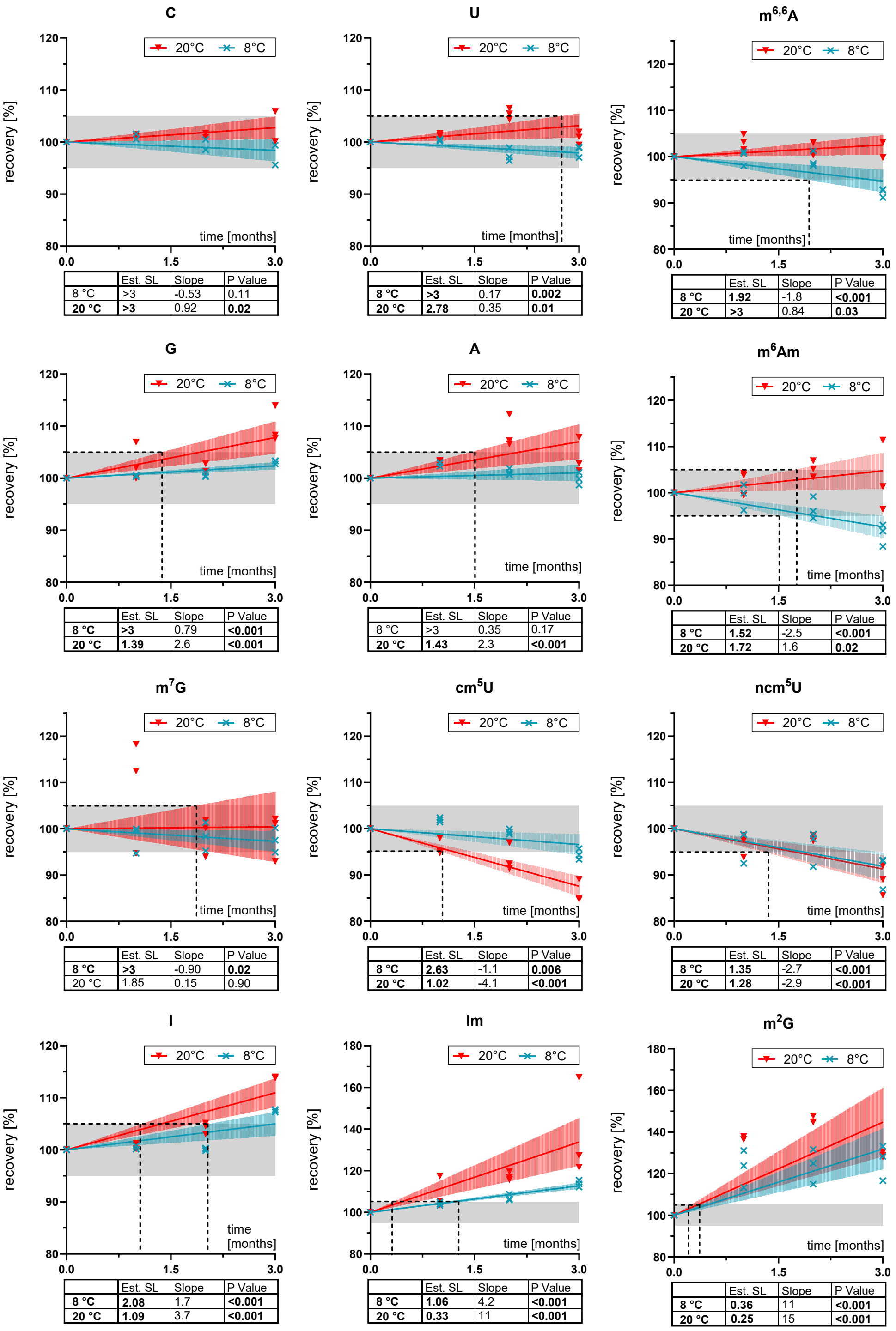


**Figure S21.** Short-term (RT and 8 °C) shelf life for nucleosides shown in Figure 5 of the main manuscript. Shelf life was evaluated by normalizing the UV peak area after storage to the initial baseline (100 %). The grey shaded region indicates the acceptance criterion of ± 5 % deviation (95–105 %). Coloured shaded bands represent the 95 % confidence interval of the linear regression. Inset tables summarize the statistical parameters (slope, p-value, and Est. SL).

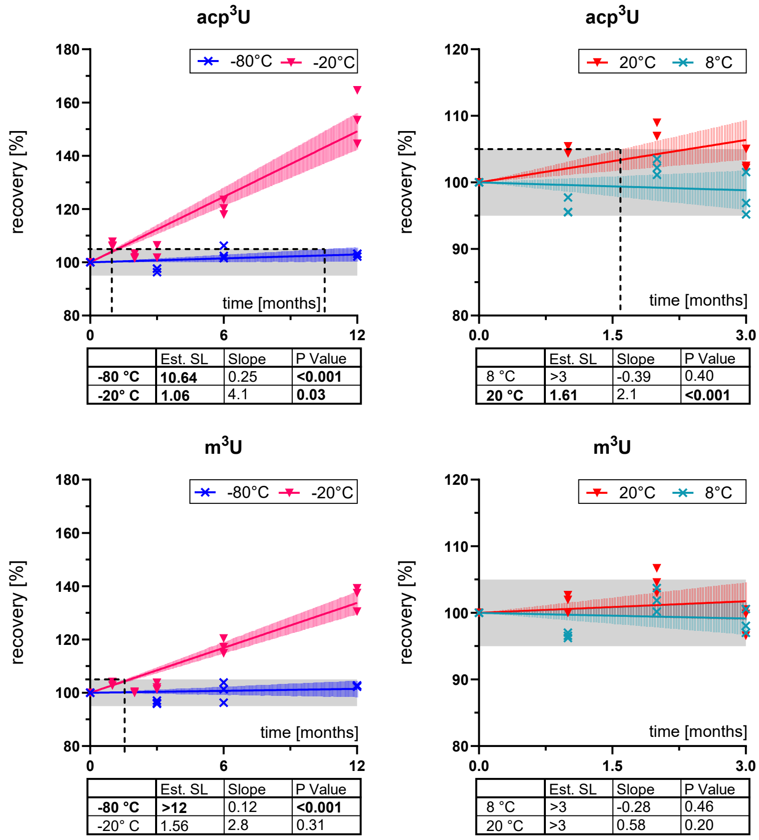


acp3U (-80 °C / -20 °C):

| | Est. SL | Slope | P Value |
|---|---|---|---|
| **-80 °C** | **10.64** | 0.25 | **<0.001** |
| **-20° C** | **1.06** | 4.1 | **0.03** |

acp3U (8 °C / 20 °C):

| | Est. SL | Slope | P Value |
|---|---|---|---|
| 8 °C | >3 | -0.39 | 0.40 |
| **20 °C** | **1.61** | 2.1 | **<0.001** |

m3U (-80 °C / -20 °C):

| | Est. SL | Slope | P Value |
|---|---|---|---|
| **-80 °C** | **>12** | 0.12 | **<0.001** |
| -20° C | 1.56 | 2.8 | 0.31 |

m3U (8 °C / 20 °C):

| | Est. SL | Slope | P Value |
|---|---|---|---|
| 8 °C | >3 | -0.28 | 0.46 |
| 20 °C | >3 | 0.58 | 0.20 |

**Figure S22.** Long-term (-20 °C and -80°C) and short-term (RT and 8 °C) shelf life for m$^3$U and acp$^3$U. Unlike the other samples, these nucleosides were stored in cryo-vials rather than 96-well plates. Due to the high rate of solvent evaporation observed in cryo-vials, an accurate assessment of stability at -20 °C is not possible; therefore, these specific data points are considered inconclusive. Shelf life was evaluated by normalizing the UV peak area after storage to the initial baseline (100 %). The grey shaded region indicates the acceptance criterion of ± 5 % deviation (95–105 %). Coloured shaded bands represent the 95 % confidence interval of the linear regression. Inset tables summarize the statistical parameters (slope, p-value, and Est. SL).

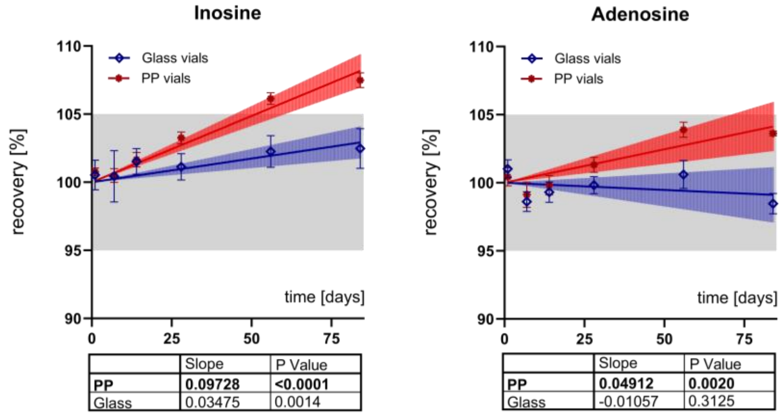


Inosine:

| | Slope | P Value |
|---|---|---|
| **PP** | **0.09728** | **<0.0001** |
| Glass | 0.03475 | 0.0014 |

Adenosine:

| | Slope | P Value |
|---|---|---|
| **PP** | **0.04912** | **0.0020** |
| Glass | -0.01057 | 0.3125 |

**Figure S23.** Observation at room temperature storage of inosine and adenosine in polypropylene and glass vials for 3 months. Significant regression with a positive slope for the PP-stored nucleosides and inosine in glass vials most likely due to evaporation.

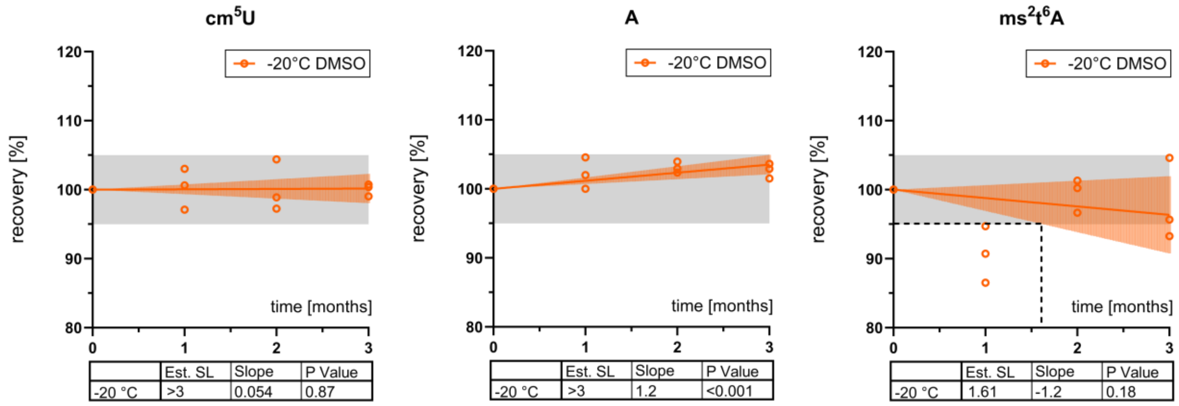


**Figure S24** Additional examples of stable nucleosides complementing the data presented in Figure 6 of the main text (storage at -20 °C in DMSO). cm$^5$U and A do not fall outside the acceptance criteria (95-105 %), while ms$^2$t$^6$A does not show a statistically significant slope. Shelf life was determined by comparing the UV area after storage to the initial value and normalized to 100 %. The grey shaded region indicates the acceptance criterion of ± 5 % deviation (95–105 %). Coloured shaded bands represent the 95 % confidence interval of the linear regression. Statistical parameters (Slope, P-value, and Est.SL) are provided in the inset tables.
