## Supplement Information 2 for "Long-term stability of RNA nucleoside standards for accurate LC-MS quantification"

### Table of Contents

**S1 Computational Details** ···· **3**

S1.1 Method Development for Conformational Search ···· 3

**S2 Theoretical Study** ···· **4**

S2.1 Calculation Method ···· 4

S2.1.1 Complete Basis Set ···· 4

S2.1.2 Reaction Energy ···· 5

S2.2 Conformer Analysis ···· 9

**S3 Results and Discussion** ···· **104**

S3.1 Four Reaction Categories ···· 104

S3.1.1 Deglycosylation ···· 105

S3.1.2 Deamination ···· 107

S3.1.3 Deacetylation ···· 109

S3.1.4 Desulfurization ···· 111

S3.2 Corresponding Computational Results to Experimental Findings ···· 113

S3.2.1 Degradation of 4-thiouridine ···· 113

S3.2.2 Degradation of 3-methylcytidine ···· 114

S3.2.3 Degradation of $N^6$-isoprenyladenosine ···· 115

S3.2.4 Degradation of $N^4$-acetylcytidine ···· 116

**S4 Reference** ···· **117**

### S1 Computational Details

#### S1.1 Method Development for Conformational Search

Appropriate conformational sampling is important to identify the conformer with the lowest energy. Cytidine was selected as the model compound to explore the conformational space using different computational techniques. The initial conformer search employed GOAT (Global Optimization Automated Toolkit) module in ORCA 6.0.0[1] and CREST 2.12[2-3] (Conformer-Rotamer Ensemble Sampling Tool) to have a robust assessment of the conformational space covered by different approaches. GOAT was applied at the GFN2-xTB[4] level to map the initial conformers of cytidine, while CREST was used here with both GFN2-xTB and GFNFF[5], yielding two additional sets of conformers for cytidine. Before conducting the conformational search, the input structure was pre-optimzed at the same level serving as a reference for topology check. The default energy window 6 kcal/mol was applied as a sorting threshold. The xTB method from GOAT and CREST yielded similar conformer sets, while GFNFF from CREST produced a distinct set. To evaluate each method's coverage of the conformational space, two key structural parameters were analyzed, the O1'-C1'-N1-C2 and C1'-C2'-C3'-C4' dihedral angle. The former determines whether the carbonyl group of cytosine is in an anti or gauche orientation, while the latter determines whether the ribose ring's C3' position is endo (positive) or exo (negative dihedral angle). After confirming the validity of this method, we chose the conformational search with CREST at GFN2-xTB level.

### S2 Theoretical Study

#### S2.1 Calculation Method

##### S2.1.1Complete Basis Set

Following the conformer sampling, all conformers were further optimized using Gaussian 09[6] at the (U)B3LYP[7]-D3[8]/def2-TZVPP[9] level of theory in the gas phase, followed by single-point calculation at the SMD($H_2O$)[10]/(U)B3LYP-D3/def2-TZVPP level of theory in the aqueous phase. Single point energies have subsequently been calculated with the DLPNO-CCSD(T)[11-12] method as implemented in ORCA 6.0.0 in combination with the cc-pVTZ and cc-pVQZ basis sets[13-15], followed by extrapolation to the complete basis set (CBS) limit to DLPNO-CCSD(T)/CBS total energies.[13] Each of the DLPNO is separated into $E_{HF}$ as the Hartree-Fock reference energy and $E_C$ as the final correlation energy. For the cc-pVTZ and cc-pVQZ basis sets, the 3/4 extrapolation scheme with $n = 3$ for triple- and $m = 4$ for quadruple-zeta (eq. (2.1)), and the constants with $\alpha = 5.46$ and $\beta = 3.05$ were used (eq. (2.2)).[11, 16] $E_{CBS,HF}$ was summed up with $E_{CBS,C}$ to get the correct extrapolated DLPNO energy $E_{CBS}$ (eq. (2.3)).

$$\mathrm{E}_{\mathrm{HF}}^{\mathrm{CBS}} = \frac{\mathrm{E}_{\mathrm{HF}}^{n} \cdot e^{-\alpha\sqrt{m}} - \mathrm{E}_{\mathrm{HF}}^{m} \cdot e^{-\alpha\sqrt{n}}}{e^{-\alpha\sqrt{m}} - e^{-\alpha\sqrt{n}}} \tag{2.1}$$

$$\mathrm{E}_{\mathrm{C}}^{\mathrm{CBS}} = \frac{\mathrm{E}_{\mathrm{C}}^{n} \cdot n^{\beta} - \mathrm{E}_{\mathrm{C}}^{m} \cdot m^{\beta}}{n^{\beta} - m^{\beta}} \tag{2.2}$$

$$\mathrm{E}_{\mathrm{CBS}} = \mathrm{E}_{\mathrm{HF}}^{\mathrm{CBS}} + \mathrm{E}_{\mathrm{C}}^{\mathrm{CBS}} \tag{2.3}$$

#### S2.1.2Reaction Energy

The Gibbs free energy of solvation in water $\Delta G_{Solv}$ was obtained as the difference of total energies without zero-point correction in the aqueous phase $E_{Tot,W}$ and in the gas phase $E_{Tot}$ (eq. (2.4)). The Gibbs free energy at standard state in the aqueous phase $G_{298,W}$ was obtained as the sum of $E_{Tot}$, $\Delta G_{Solv}$, thermal correction to Gibbs free energy $\delta G$, and standard state correction energy to 1 mol/L $\Delta G_{Corr}$ (+7.908 kJ/mol) (eq. (2.5)).

$$\Delta G_{Solv} = E_{Tot,W} - E_{Tot} \tag{2.4}$$

$$G_{298,W} = E_{Tot} + \Delta G_{Solv} + \delta G + \Delta G_{Corr} \tag{2.5}$$

The Gibbs free energy at standard state in the gas phase $G_{CBS}$ at DLPNO-CCSD(T)/CBS level of theory was obtained as the sum of $E_{CBS}$ and $\delta G$ (eq. (2.6)), while the aqueous phase $G_{CBS,W}$ was obtained as the sum of $G_{CBS}$, $\Delta G_{Solv}$, and $\Delta G_{Corr}$ (eq. (2.7)).

$$G_{CBS} = E_{CBS} + \delta G \tag{2.6}$$

$$G_{CBS,W} = G_{CBS} + \Delta G_{Solv} + \Delta G_{Corr} \tag{2.7}$$

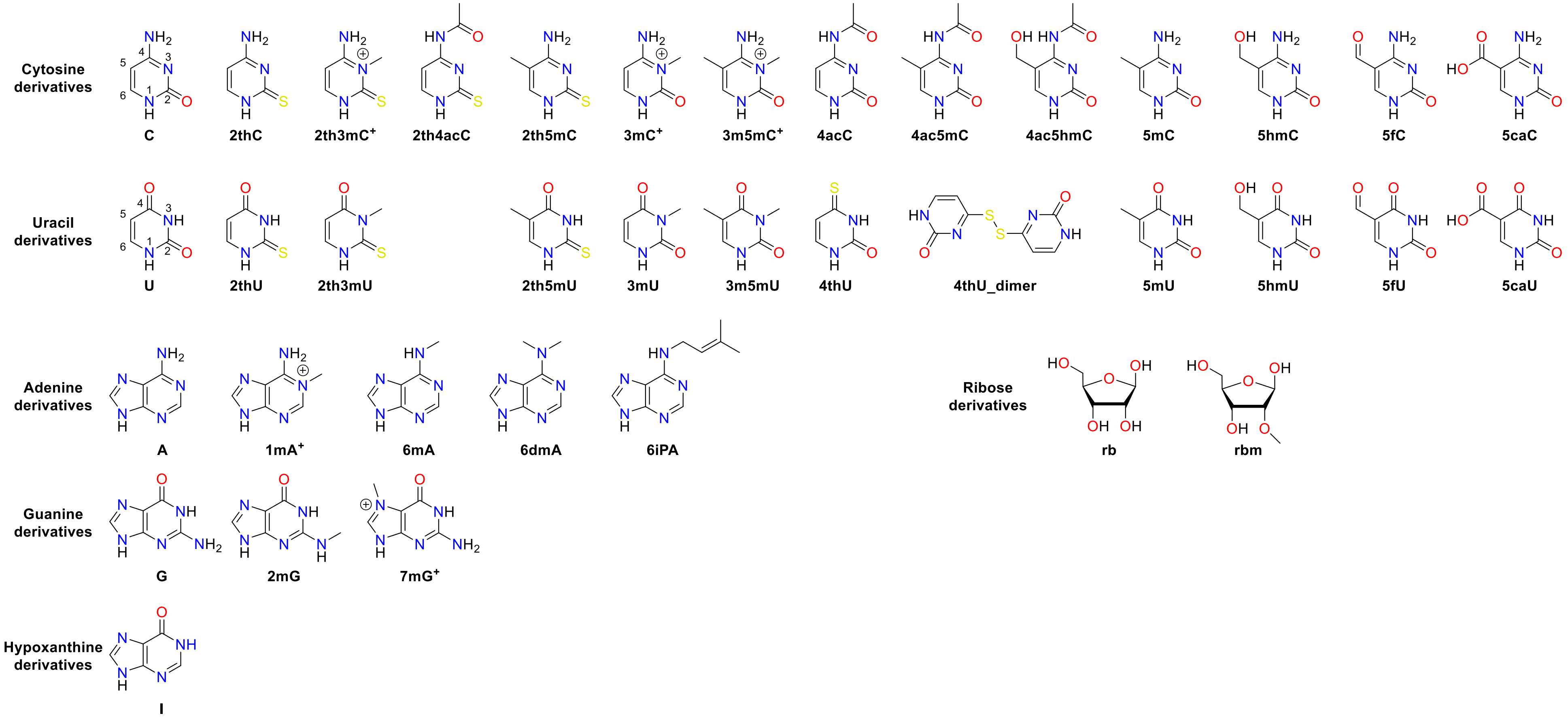

**Figure S1.** The 24 modified cytosine and uracil derivatives.

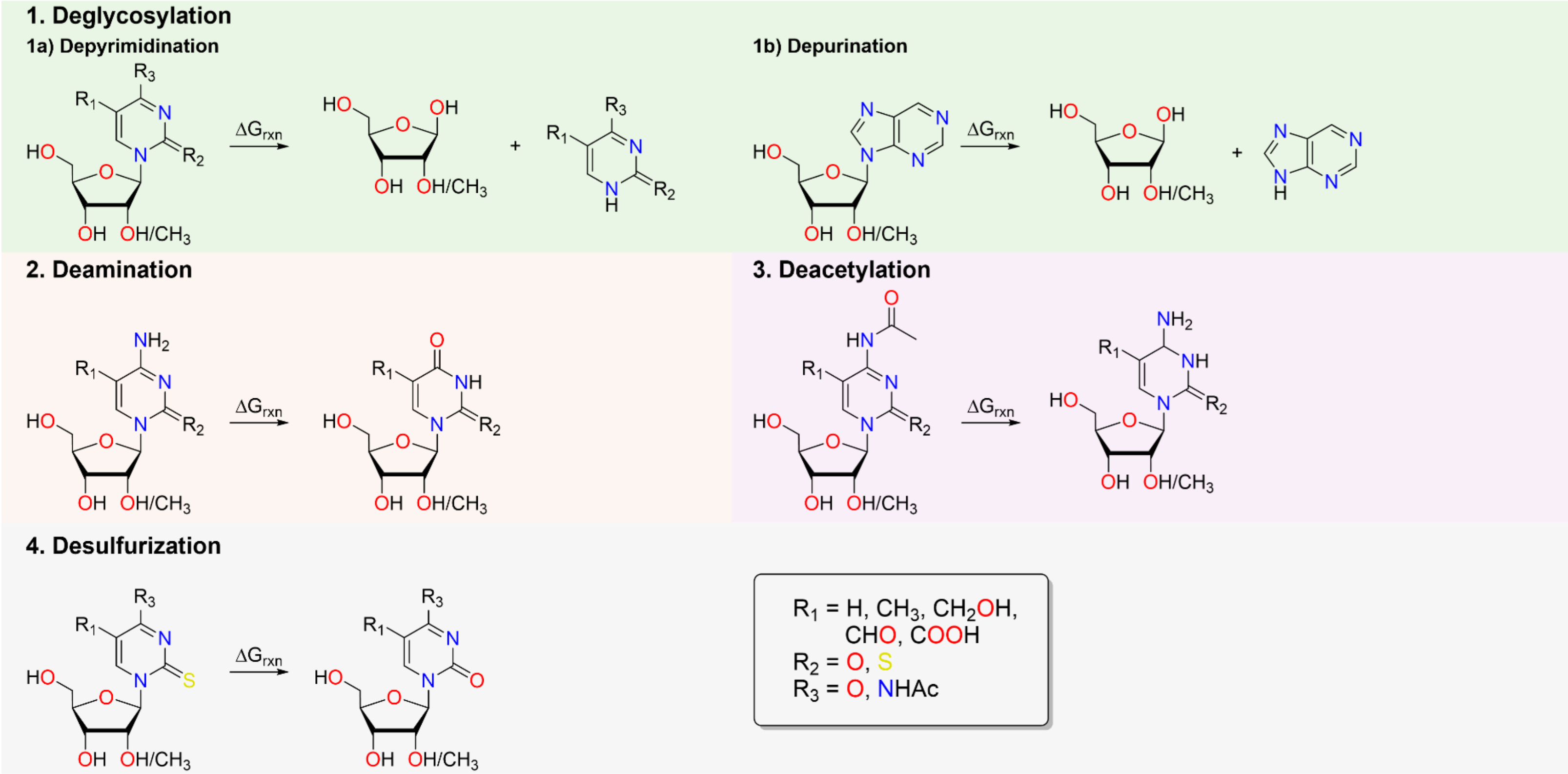

**Figure S2.** Overview of four potential decomposition reactions of modified cytidines and uridines: (1) Deglycosylation[17]; (2) Deamination[18]; (3) Deacetylation[19]; (4) Desulfurization[20].

The thermodynamic driving force for different reactions is reflected in the reaction free energies $\Delta E_{rxn,gas}$, $\Delta G_{rxn,gas}$, $\Delta G_{rxn}$ (eq. (2.8), eq. (2.9), eq. (2.10)).

$$\Delta E_{rxn,gas} = \sum E_{CBS}^{Product} - \sum E_{CBS}^{Reactant} \tag{2.8}$$

$$\Delta G_{rxn,gas} = \sum G_{CBS}^{Product} - \sum G_{CBS}^{Reactant} \tag{2.9}$$

$$\Delta G_{rxn} = \sum G_{CBS,W}^{Product} - \sum G_{CBS,W}^{Reactant} \tag{2.10}$$

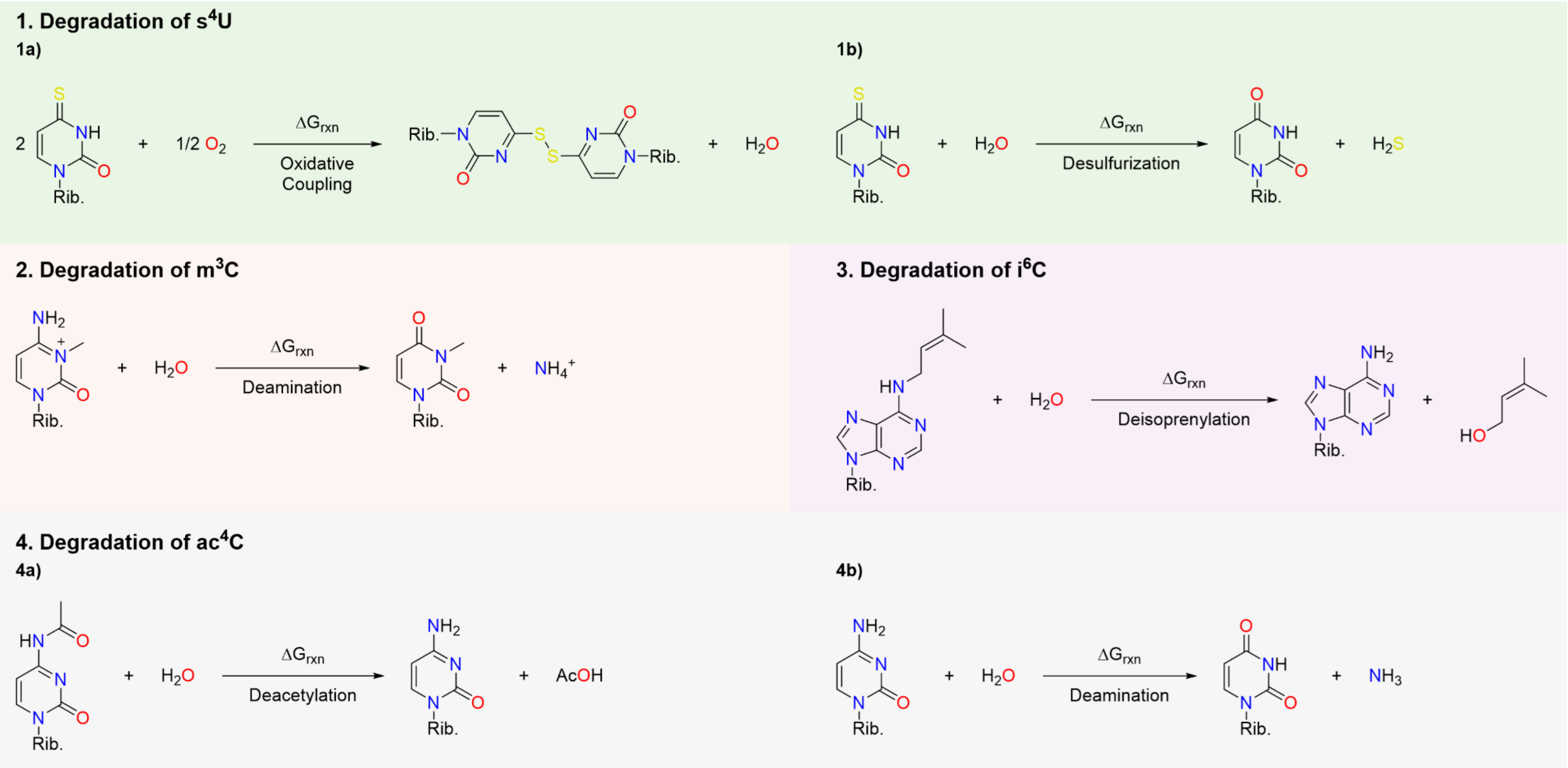

**Figure S3.** Corresponding four categories of reaction in **Figure 4**: (1) Degradation of 4-thiouridine; (2) Degradation of 3-methylcytidine; (3) Degradation of 6-isopentenyladenosine; (4) Degradation of $N^4$-acetylcytidine.

The thermodynamic driving force for different reactions is reflected in the reaction free energies $\Delta E_{rxn,gas}$, $\Delta G_{rxn,gas}$, $\Delta G_{rxn}$ (eq. (2.8), eq. (2.9), eq. (2.10)).

$$\Delta E_{rxn,gas} = \sum E_{CBS}^{Product} - \sum E_{CBS}^{Reactant} \tag{2.11}$$

$$\Delta G_{rxn,gas} = \sum G_{CBS}^{Product} - \sum G_{CBS}^{Reactant} \tag{2.12}$$

$$\Delta G_{rxn} = \sum G_{CBS,W}^{Product} - \sum G_{CBS,W}^{Reactant} \tag{2.13}$$

#### S2.2 Conformer Analysis

**Table S1.** Gas-phase optimized basic molecules at the (U)B3LYP-D3/def2-TZVPP level of theory followed by aqueous phase single-point calculation. The columns display total energy without zero-point correction ($E_{Tot}$), Gibbs free energy ($\delta G$), total energy without zero-point correction ($E_{Tot,W}$), Gibbs free energy ($G_{298,W}$) in water (W), total single-point energy ($E_{CBS}$) calculated at DLPNO-CCSD(T)/CBS level of theory, and their corresponding free energy $G_{CBS}$. $G_{298,W}$ and $G_{CBS,W}$ have been corrected to the standard state of 1 mol/L by addition of +7.908 kJ/mol. $\Delta G_{Solv}$ represents the Gibbs free energy of solvation.

| | (U)B3LYP-D3/def2-TZVPP | | | SMD($H_2O$)/(U)B3LYP-D3/def2-TZVPP[a] | | | | DLPNO-CCSD(T)/CBS | | |
|---|---|---|---|---|---|---|---|---|---|---|
| No. | $E_{Tot}$ (Hartree) | $\delta H$ (Hartree) | $\delta G$ (Hartree) | $E_{Tot,W}$ (Hartree) | $\Delta G_{Solv}$ (kJ/mol) | $H_{298,W}$ (Hartree) | $G_{298,W}$ (Hartree) | $E_{CBS,HF}$ (Hartree) | $E_{CBS,C}$ (Hartree) | $G_{CBS,W}$ (Hartree) |
| **$H_2O$** | -76.466681 | 0.025091 | 0.003672 | -76.478486 | -31.0 | -76.453395 | -76.471802 | -76.066879 | -0.309226 | -76.381226 |
| **$NH_3$** | -56.588660 | 0.038040 | 0.015162 | -56.593896 | -13.7 | -56.555856 | -56.575722 | -56.224610 | -0.280348 | -56.492019 |
| **$NH_4^+$** | -56.927233 | 0.053214 | 0.032115 | -57.058092 | -343.6 | -57.004878 | -57.022965 | -56.569082 | -0.273837 | -56.938651 |
| **$H_2S$** | -399.426820 | 0.018837 | -0.004508 | -399.428801 | -5.2 | -399.409964 | -399.430297 | -398.719353 | -0.248411 | -398.971242 |
| **$O_2$** | -150.389771 | 0.007037 | -0.016233 | -150.390443 | -1.8 | -150.383406 | -150.403664 | -149.691643 | -0.532628 | -150.238165 |
| **A** | -467.515091 | 0.120035 | 0.079489 | -467.540992 | -68.0 | -467.420957 | -467.458491 | -464.721333 | -2.041510 | -466.706244 |
| **1mA** | -506.811845 | 0.149407 | 0.106576 | -506.852798 | -107.5 | -506.703391 | -506.743210 | -503.729300 | -2.257015 | -505.917680 |
| **$1mA^+$** | -507.221283 | 0.163147 | 0.120183 | -507.320812 | -261.3 | -507.157665 | -507.197617 | -504.149115 | -2.241892 | -506.367342 |
| **6dmA** | -546.158371 | 0.179071 | 0.130722 | -546.175493 | -45.0 | -546.041759 | -546.172481 | -542.797280 | -2.461293 | -545.141962 |
| **I** | -487.391157 | 0.107798 | 0.068741 | -487.423645 | -85.3 | -487.315847 | -487.351892 | -484.559106 | -2.069309 | -486.589150 |
| **G** | -542.787393 | 0.125836 | 0.083717 | -542.826154 | -101.8 | -542.700318 | -542.739425 | -539.635285 | -2.309574 | -541.896892 |
| **7mG** | -582.116335 | 0.155488 | 0.108785 | -582.150821 | -90.5 | -581.995333 | -582.039024 | -578.681077 | -2.518395 | -581.122161 |
| **$7mG^+$** | -582.503931 | 0.168848 | 0.121988 | -582.606659 | -269.7 | -582.437811 | -582.481659 | -579.073730 | -2.507212 | -581.558669 |
| **C** | -395.107825 | 0.105973 | 0.067228 | -395.138152 | -79.6 | -395.032179 | -395.067912 | -392.797146 | -1.690307 | -394.447539 |
| **2thC** | -718.056461 | 0.104157 | 0.063605 | -718.085841 | -77.1 | -717.981684 | -718.019224 | -715.431408 | -1.640875 | -717.035045 |
| **2th5mC** | -757.392145 | 0.133521 | 0.089506 | -757.421758 | -77.8 | -757.288237 | -757.329240 | -754.483796 | -1.851279 | -756.272171 |
| **$3mC^+$** | -434.819005 | 0.149496 | 0.108303 | -434.918911 | -262.3 | -434.769415 | -434.807596 | -432.230823 | -1.890489 | -434.109903 |
| **5mC** | -434.442854 | 0.135362 | 0.093276 | -434.473215 | -79.7 | -434.337853 | -434.376927 | -431.848606 | -1.900906 | -433.683584 |
| **U** | -415.003094 | 0.094185 | 0.056709 | -415.024647 | -56.6 | -414.930462 | -414.964926 | -412.658470 | -1.715049 | -414.335351 |
| **2thU** | -737.951572 | 0.092349 | 0.053663 | -737.971990 | -53.6 | -737.879641 | -737.915315 | -735.291119 | -1.666870 | -736.921732 |
| **2th5mU** | -777.289446 | 0.121734 | 0.079397 | -777.308811 | -50.8 | -777.187077 | -777.226402 | -774.346473 | -1.876144 | -776.159573 |
| **3mU** | -454.328792 | 0.123500 | 0.081697 | -454.347123 | -48.1 | -454.223623 | -454.262414 | -451.700717 | -1.925575 | -453.559914 |
| **3m5mU** | -493.665904 | 0.152864 | 0.107839 | -493.682978 | -44.8 | -493.530114 | -493.572127 | -490.755138 | -2.135190 | -492.796552 |
| **5mU** | -454.340292 | 0.123509 | 0.082343 | -454.360920 | -54.2 | -454.237411 | -454.275565 | -451.713007 | -1.924502 | -453.572781 |
| **5ncmU** | -623.125333 | 0.154567 | 0.106331 | -623.155382 | -78.9 | -623.000815 | -623.046039 | -619.578777 | -2.602439 | -622.101922 |

[a]: Single-point calculation in aqueous phase with SMD model.

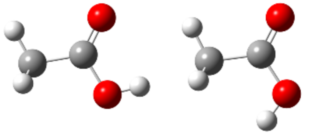

conformer_1 conformer_2

**Figure S4.** B3LYP-D3/def2-TZVPP optimized geometries of conformers for acetic acid (**AcOH**).

**Table S2.** Conformers of gas-phase optimized acetic acid (**AcOH**) at the B3LYP-D3/def2-TZVPP level of theory followed by aqueous phase single-point calculation. The columns display total energy without zero-point correction ($E_{Tot}$), thermal correction to enthalpy (δH), Gibbs free energy (δG), total energy without zero-point correction ($E_{Tot,W}$), Gibbs free energy ($G_{298,W}$) in water (W), total single-point energy ($E_{CBS}$) calculated at DLPNO-CCSD(T)/CBS level of theory, and their corresponding free energy $G_{CBS,W}$. $G_{298,W}$ and $G_{CBS,W}$ have been corrected to the standard state of 1 mol/L by addition of +7.908 kJ/mol. $\Delta G_{Solv}$ represents the Gibbs free energy of solvation. The data are arranged in the ascending numeric order of $E_{Tot,W}$. $\Delta G_{298,W}$ represents the respective energy difference to the lowest structure. Only conformers within the 24 kJ/mol (6 kcal/mol) energy window above the lowest in CREST are included in initial conformer sampling. Duplicates of the same structure are excluded. The overall optimum is marked bold.

| | **B3LYP-D3/def2-TZVPP** | | | **SMD($H_2O$)/B3LYP-D3/def2-TZVPP[a]** | | | | | **DLPNO-CCSD(T)/CBS** | | |
|---|---|---|---|---|---|---|---|---|---|---|---|
| **AcOH No.** | **$E_{Tot}$** (Hartree) | **δH** (Hartree) | **δG** (Hartree) | **$E_{Tot,W}$** (Hartree) | **$\Delta G_{Solv}$** (kJ/mol) | **$H_{298,W}$** (Hartree) | **$G_{298,W}$** (Hartree) | **$\Delta G_{298,W}$** (kJ/mol) | **$E_{CBS,HF}$** (Hartree) | **$E_{CBS,C}$** (Hartree) | **$G_{CBS,W}$** (Hartree) |
| **conformer_1** | **-229.195033** | **0.067093** | **0.034344** | **-229.204206** | **-24.1** | **-229.137113** | **-229.166850** | **0** | **-227.924951** | **-0.941485** | **-228.838252** |
| conformer_2 | -229.186670 | 0.066885 | 0.034463 | -229.202476 | -41.5 | -229.135591 | -229.165001 | 4.9 | -227.915807 | -0.942454 | -228.836592 |

[a]: Single-point calculation in aqueous phase with SMD model.

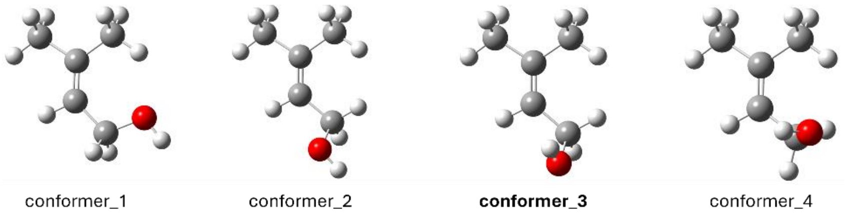

**Figure S5.** B3LYP-D3/def2-TZVPP optimized geometries of conformers for prenol (**iPOH**).

**Table S3.** Conformers of gas-phase optimized prenol (**iPOH**) at the B3LYP-D3/def2-TZVPP level of theory followed by aqueous phase single-point calculation. The columns display total energy without zero-point correction ($E_{Tot}$), thermal correction to enthalpy (δH), Gibbs free energy (δG), total energy without zero-point correction ($E_{Tot,W}$), Gibbs free energy ($G_{298,W}$) in water (W), total single-point energy ($E_{CBS}$) calculated at DLPNO-CCSD(T)/CBS level of theory, and their corresponding free energy $G_{CBS,W}$. $G_{298,W}$ and $G_{CBS,W}$ have been corrected to the standard state of 1 mol/L by addition of +7.908 kJ/mol. $\Delta G_{Solv}$ represents the Gibbs free energy of solvation. The data are arranged in the ascending numeric order of $E_{Tot,W}$. $\Delta G_{298,W}$ represents the respective energy difference to the lowest structure. Only conformers within the 24 kJ/mol (6 kcal/mol) energy window above the lowest in CREST are included in initial conformer sampling. Duplicates of the same structure are excluded. The overall optimum is marked bold.

| | B3LYP-D3/def2-TZVPP | | | SMD($H_2O$)/B3LYP-D3/def2-TZVPP[a] | | | | | DLPNO-CCSD(T)/CBS | | |
|---|---|---|---|---|---|---|---|---|---|---|---|
| **iPOH No.** | **$E_{Tot}$** (Hartree) | **δH** (Hartree) | **δG** (Hartree) | **$E_{Tot,W}$** (Hartree) | **$\Delta G_{Solv}$** (kJ/mol) | **$H_{298,W}$** (Hartree) | **$G_{298,W}$** (Hartree) | **$\Delta G_{298,W}$** (kJ/mol) | **$E_{CBS,HF}$** (Hartree) | **$E_{CBS,C}$** (Hartree) | **$G_{CBS,W}$** (Hartree) |
| **conformer_3** | **-271.880351** | **0.149829** | **0.109215** | **-271.886671** | **-16.6** | **-271.736842** | **-271.774444** | **0** | **-270.121870** | **-1.299424** | **-271.315388** |
| conformer_4 | -271.879258 | 0.149690 | 0.108902 | -271.884967 | -15.0 | -271.735277 | -271.773053 | 3.7 | -270.119787 | -1.300259 | -271.313842 |
| conformer_2 | -271.878153 | 0.149612 | 0.108563 | -271.885463 | -19.2 | -271.735851 | -271.773888 | 1.5 | -270.120382 | -1.299039 | -271.315156 |
| conformer_1 | -271.875227 | 0.149587 | 0.107986 | -271.880246 | -13.2 | -271.730659 | -271.769248 | 13.6 | -270.116702 | -1.299543 | -271.310266 |

[a]: Single-point calculation in aqueous phase with SMD model.

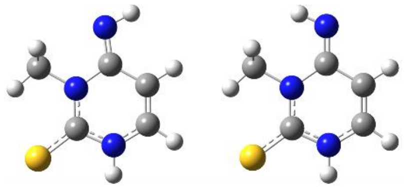

**Figure S6.** B3LYP-D3/def2-TZVPP optimized geometries of conformers for neutral 2-thio-3-methylcytosine (**2th3mC**).

**Table S4.** Conformers of gas-phase optimized neutral 2-thio-3-methylcytosine (**2th3mC**) at the B3LYP-D3/def2-TZVPP level of theory followed by aqueous phase single-point calculation. The columns display total energy without zero-point correction ($E_{Tot}$), thermal correction to enthalpy (δH), Gibbs free energy (δG), total energy without zero-point correction ($E_{Tot,W}$), Gibbs free energy ($G_{298,W}$) in water (W), total single-point energy ($E_{CBS}$) calculated at DLPNO-CCSD(T)/CBS level of theory, and their corresponding free energy $G_{CBS,W}$. $G_{298,W}$ and $G_{CBS,W}$ have been corrected to the standard state of 1 mol/L by addition of +7.908 kJ/mol. $\Delta G_{Solv}$ represents the Gibbs free energy of solvation. The data are arranged in the ascending numeric order of $E_{Tot,W}$. $\Delta G_{298,W}$ represents the respective energy difference to the lowest structure. Only conformers within the 24 kJ/mol (6 kcal/mol) energy window above the lowest in CREST are included in initial conformer sampling. Duplicates of the same structure are excluded. The overall optimum is marked bold.

| | **B3LYP-D3/def2-TZVPP** | | | **SMD(H2O)/B3LYP-D3/def2-TZVPP[a]** | | | | | **DLPNO-CCSD(T)/CBS** | | |
|---|---|---|---|---|---|---|---|---|---|---|---|
| **2th3mC (neutral) No.** | **$E_{Tot}$** (Hartree) | **δH** (Hartree) | **δG** (Hartree) | **$E_{Tot,W}$** (Hartree) | **$\Delta G_{Solv}$** (kJ/mol) | **$H_{298,W}$** (Hartree) | **$G_{298,W}$** (Hartree) | **$\Delta G_{298,W}$** (kJ/mol) | **$E_{CBS,HF}$** (Hartree) | **$E_{CBS,C}$** (Hartree) | **$G_{CBS,W}$** (Hartree) |
| **conformer_1** | **-757.376687** | **0.133987** | **0.091126** | **-757.3929946** | **-42.8** | **-757.259008** | **-757.298857** | **0** | **-754.465868** | **-1.856343** | **-756.244381** |
| conformer_2 | -757.374004 | 0.133911 | 0.091407 | -757.392241 | -47.9 | -757.258330 | -757.297822 | 2.7 | -754.462266 | -1.857289 | -756.243373 |

[a]: Single-point calculation in aqueous phase with SMD model.

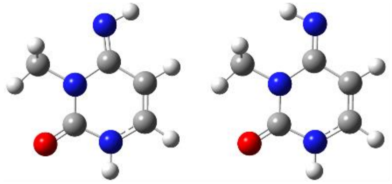

**Figure S7.** B3LYP-D3/def2-TZVPP optimized geometries of conformers for neutral 3-methylcytosine (**3mC**).

**Table S5.** Conformers of gas-phase optimized neutral 3-methylcytosine (**3mC**) at the B3LYP-D3/def2-TZVPP level of theory followed by aqueous phase single-point calculation. The columns display total energy without zero-point correction ($E_{Tot}$), thermal correction to enthalpy (δH), Gibbs free energy (δG), total energy without zero-point correction ($E_{Tot,W}$), Gibbs free energy ($G_{298,W}$) in water (W), total single-point energy ($E_{CBS}$) calculated at DLPNO-CCSD(T)/CBS level of theory, and their corresponding free energy $G_{CBS,W}$. $G_{298,W}$ and $G_{CBS,W}$ have been corrected to the standard state of 1 mol/L by addition of +7.908 kJ/mol. $\Delta G_{Solv}$ represents the Gibbs free energy of solvation. The data are arranged in the ascending numeric order of $E_{Tot,W}$. $\Delta G_{298,W}$ represents the respective energy difference to the lowest structure. Only conformers within the 24 kJ/mol (6 kcal/mol) energy window above the lowest in CREST are included in initial conformer sampling. Duplicates of the same structure are excluded. The overall optimum is marked bold.

| | **B3LYP-D3/def2-TZVPP** | | | **SMD($H_2O$)/B3LYP-D3/def2-TZVPP[a]** | | | | | **DLPNO-CCSD(T)/CBS** | | |
|---|---|---|---|---|---|---|---|---|---|---|---|
| **3mC No.** | **$E_{Tot}$** (Hartree) | **δH** (Hartree) | **δG** (Hartree) | **$E_{Tot,W}$** (Hartree) | **$\Delta G_{Solv}$** (kJ/mol) | **$H_{298,W}$** (Hartree) | **$G_{298,W}$** (Hartree) | **$\Delta G_{298,W}$** (kJ/mol) | **$E_{CBS,HF}$** (Hartree) | **$E_{CBS,C}$** (Hartree) | **$G_{CBS,W}$** (Hartree) |
| **conformer_1** | **-434.430332** | **0.135993** | **0.094736** | **-434.448131** | **-46.7** | **-434.312138** | **-434.350383** | **0** | **-431.836464** | **-1.902566** | **-433.659082** |
| conformer_2 | -434.427760 | 0.135855 | 0.094797 | -434.447343 | -51.4 | -434.311488 | -434.349534 | 2.2 | -431.833151 | -1.903354 | -433.658279 |

[a]: Single-point calculation in aqueous phase with SMD model.

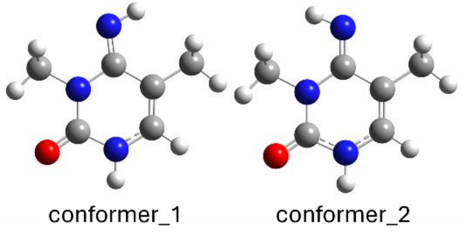

**Figure S8.** B3LYP-D3/def2-TZVPP optimized geometries of conformers for neutral 3,5-dimethylcytosine (**3m5mC**).

**Table S6.** Conformers of gas-phase optimized 3,5-dimethylcytosine (**3m5mC**) at the B3LYP-D3/def2-TZVPP level of theory followed by aqueous phase single-point calculation. The columns display total energy without zero-point correction ($E_{Tot}$), thermal correction to enthalpy (δH), Gibbs free energy (δG), total energy without zero-point correction ($E_{Tot,W}$), Gibbs free energy ($G_{298,W}$) in water (W), total single-point energy ($E_{CBS}$) calculated at DLPNO-CCSD(T)/CBS level of theory, and their corresponding free energy $G_{CBS,W}$. $G_{298,W}$ and $G_{CBS,W}$ have been corrected to the standard state of 1 mol/L by addition of +7.908 kJ/mol. $\Delta G_{Solv}$ represents the Gibbs free energy of solvation. The data are arranged in the ascending numeric order of $E_{Tot,W}$. $\Delta G_{298,W}$ represents the respective energy difference to the lowest structure. Only conformers within the 24 kJ/mol (6 kcal/mol) energy window above the lowest in CREST are included in initial conformer sampling. Duplicates of the same structure are excluded. The overall optimum is marked bold.

| 3m5mC No. | B3LYP-D3/def2-TZVPP | | | SMD($H_2O$)/B3LYP-D3/def2-TZVPP[a] | | | | | DLPNO-CCSD(T)/CBS | | |
|---|---|---|---|---|---|---|---|---|---|---|---|
| | **$E_{Tot}$** (Hartree) | **δH** (Hartree) | **δG** (Hartree) | **$E_{Tot,W}$** (Hartree) | **$\Delta G_{Solv}$** (kJ/mol) | **$H_{298,W}$** (Hartree) | **$G_{298,W}$** (Hartree) | **$\Delta G_{298,W}$** (kJ/mol) | **$E_{CBS,HF}$** (Hartree) | **$E_{CBS,C}$** (Hartree) | **$G_{CBS,W}$** (Hartree) |
| **conformer_1** | **-473.766469** | **0.165386** | **0.120951** | **-473.783531** | **-44.8** | **-473.618145** | **-473.659568** | **0** | **-470.889491** | **-2.112950** | **-472.895541** |
| conformer_2 | -473.764582 | 0.165271 | 0.120880 | -473.782060 | -45.9 | -473.616789 | -473.658168 | 3.7 | -470.887268 | -2.113253 | -472.894107 |

[a]: Single-point calculation in aqueous phase with SMD model.

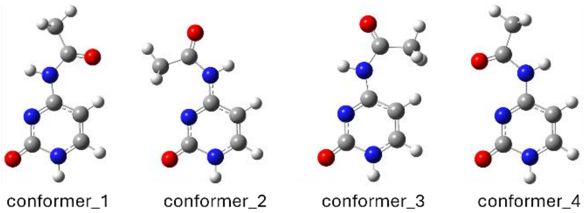

**Figure S9.** B3LYP-D3/def2-TZVPP optimized geometries of conformers for 4-acetylcytosine (**4acC**).

**Table S7.** Conformers of gas-phase optimized 4-acetylcytosine (**4acC**) at the B3LYP-D3/def2-TZVPP level of theory followed by aqueous phase single-point calculation. The columns display total energy without zero-point correction ($E_{Tot}$), thermal correction to enthalpy (δH), Gibbs free energy (δG), total energy without zero-point correction ($E_{Tot,W}$), Gibbs free energy ($G_{298,W}$) in water (W), total single-point energy ($E_{CBS}$) calculated at DLPNO-CCSD(T)/CBS level of theory, and their corresponding free energy $G_{CBS,W}$. $G_{298,W}$ and $G_{CBS,W}$ have been corrected to the standard state of 1 mol/L by addition of +7.908 kJ/mol. $\Delta G_{Solv}$ represents the Gibbs free energy of solvation. The data are arranged in the ascending numeric order of $E_{Tot,W}$. $\Delta G_{298,W}$ represents the respective energy difference to the lowest structure. Only conformers within the 24 kJ/mol (6 kcal/mol) energy window above the lowest in CREST are included in initial conformer sampling. Duplicates of the same structure are excluded. The overall optimum is marked bold.

| | B3LYP-D3/def2-TZVPP | | | SMD($H_2O$)/B3LYP-D3/def2-TZVPP[a] | | | | | DLPNO-CCSD(T)/CBS | | |
|---|---|---|---|---|---|---|---|---|---|---|---|
| **4acC No.** | **$E_{Tot}$** (Hartree) | **δH** (Hartree) | **δG** (Hartree) | **$E_{Tot,W}$** (Hartree) | **$\Delta G_{Solv}$** (kJ/mol) | **$H_{298,W}$** (Hartree) | **$G_{298,W}$** (Hartree) | **$\Delta G_{298,W}$** (kJ/mol) | **$E_{CBS,HF}$** (Hartree) | **$E_{CBS,C}$** (Hartree) | **$G_{CBS,W}$** (Hartree) |
| **conformer_1** | **-547.831137** | **0.147337** | **0.099975** | **-547.860219** | **-76.4** | **-547.712882** | **-547.757232** | **0** | **-544.644668** | **-2.330744** | **-546.901507** |
| conformer_2 | -547.827390 | 0.147197 | 0.100795 | -547.855859 | -74.7 | -547.708662 | -547.752052 | 13.6 | -544.640735 | -2.330880 | -546.896277 |
| conformer_3 | -547.822974 | 0.147389 | 0.099928 | -547.855310 | -84.9 | -547.707921 | -547.752370 | 12.8 | -544.634064 | -2.332871 | -546.896331 |
| conformer_4 | -547.815264 | 0.146738 | 0.097880 | -547.855236 | -104.9 | -547.708498 | -547.754344 | 7.6 | -544.628938 | -2.331160 | -546.899178 |

[a]: Single-point calculation in aqueous phase with SMD model.

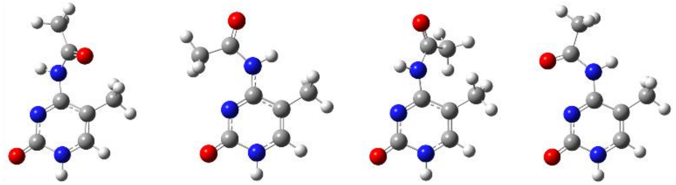

**Figure S10.** B3LYP-D3/def2-TZVPP optimized geometries of conformers for 4-acetylcytosine (**4ac5mC**).

**Table S8.** Conformers of gas-phase optimized 4-acetylcytosine (**4ac5mC**) at the B3LYP-D3/def2-TZVPP level of theory followed by aqueous phase single-point calculation. The columns display total energy without zero-point correction ($E_{Tot}$), thermal correction to enthalpy (δH), Gibbs free energy (δG), total energy without zero-point correction ($E_{Tot,W}$), Gibbs free energy ($G_{298,W}$) in water (W), total single-point energy ($E_{CBS}$) calculated at DLPNO-CCSD(T)/CBS level of theory, and their corresponding free energy $G_{CBS,W}$. $G_{298,W}$ and $G_{CBS,W}$ have been corrected to the standard state of 1 mol/L by addition of +7.908 kJ/mol. $\Delta G_{Solv}$ represents the Gibbs free energy of solvation. The data are arranged in the ascending numeric order of $E_{Tot,W}$. $\Delta G_{298,W}$ represents the respective energy difference to the lowest structure. Only conformers within the 24 kJ/mol (6 kcal/mol) energy window above the lowest in CREST are included in initial conformer sampling. Duplicates of the same structure are excluded. The overall optimum is marked bold.

| | **B3LYP-D3/def2-TZVPP** | | | **SMD(H2O)/B3LYP-D3/def2-TZVPP[a]** | | | | | **DLPNO-CCSD(T)/CBS** | | |
|---|---|---|---|---|---|---|---|---|---|---|---|
| **4ac5mC No.** | **$E_{Tot}$** (Hartree) | **δH** (Hartree) | **δG** (Hartree) | **$E_{Tot,W}$** (Hartree) | **$\Delta G_{Solv}$** (kJ/mol) | **$H_{298,W}$** (Hartree) | **$G_{298,W}$** (Hartree) | **$\Delta G_{298,W}$** (kJ/mol) | **$E_{CBS,HF}$** (Hartree) | **$E_{CBS,C}$** (Hartree) | **$G_{CBS,W}$** (Hartree) |
| conformer_2 | -587.162198 | 0.176592 | 0.126910 | -587.190234 | -73.6 | -587.013642 | -587.060312 | 5.3 | -583.691616 | -2.542006 | -586.131736 |
| **conformer_4** | **-587.149641** | **0.176070** | **0.123965** | **-587.189295** | **-104.1** | **-587.013225** | **-587.062318** | **0** | **-583.679316** | **-2.542341** | **-586.134333** |
| conformer_1 | -587.157924 | 0.176490 | 0.126200 | -587.187525 | -77.7 | -587.011035 | -587.058313 | 10.5 | -583.685786 | -2.543686 | -586.129860 |
| conformer_3 | -587.150942 | 0.176474 | 0.126892 | -587.183673 | -85.9 | -587.007199 | -587.053769 | 22.4 | -583.675770 | -2.546055 | -586.124653 |

[a]: Single-point calculation in aqueous phase with SMD model.

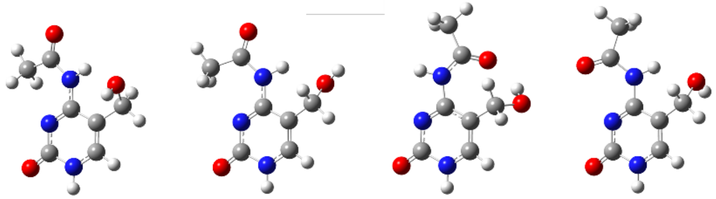

**Figure S11.** B3LYP-D3/def2-TZVPP optimized geometries of conformers for 4-acetylcytosine (**4ac5hmC**).

**Table S9.** Conformers of gas-phase optimized 4-acetylcytosine (**4ac5hmC**) at the B3LYP-D3/def2-TZVPP level of theory followed by aqueous phase single-point calculation. The columns display total energy without zero-point correction ($E_{Tot}$), thermal correction to enthalpy (δH), Gibbs free energy (δG), total energy without zero-point correction ($E_{Tot,W}$), Gibbs free energy ($G_{298,W}$) in water (W), total single-point energy ($E_{CBS}$) calculated at DLPNO-CCSD(T)/CBS level of theory, and their corresponding free energy $G_{CBS,W}$. $G_{298,W}$ and $G_{CBS,W}$ have been corrected to the standard state of 1 mol/L by addition of +7.908 kJ/mol. $\Delta G_{Solv}$ represents the Gibbs free energy of solvation. The data are arranged in the ascending numeric order of $E_{Tot,W}$. $\Delta G_{298,W}$ represents the respective energy difference to the lowest structure. Only conformers within the 24 kJ/mol (6 kcal/mol) energy window above the lowest in CREST are included in initial conformer sampling. Duplicates of the same structure are excluded. The overall optimum is marked bold.

| | **B3LYP-D3/def2-TZVPP** | | | **SMD(H2O)/B3LYP-D3/def2-TZVPP[a]** | | | | | **DLPNO-CCSD(T)/CBS** | | |
|---|---|---|---|---|---|---|---|---|---|---|---|
| **4ac5hmC No.** | **$E_{Tot}$** (Hartree) | **δH** (Hartree) | **δG** (Hartree) | **$E_{Tot,W}$** (Hartree) | **$\Delta G_{Solv}$** (kJ/mol) | **$H_{298,W}$** (Hartree) | **$G_{298,W}$** (Hartree) | **$\Delta G_{298,W}$** (kJ/mol) | **$E_{CBS,HF}$** (Hartree) | **$E_{CBS,C}$** (Hartree) | **$G_{CBS,W}$** (Hartree) |
| conformer_1 | -662.412066 | 0.182943 | 0.131049 | -662.445463 | -87.7 | -662.262520 | -662.311402 | 3.5 | -658.581907 | -2.813391 | -661.294634 |
| **conformer_15** | **-662.402703** | **0.182621** | **0.129526** | **-662.445270** | **-111.8** | **-662.262649** | **-662.312732** | **0** | **-658.572657** | **-2.813795** | **-661.296480** |
| conformer_3 | -662.410839 | 0.183186 | 0.131095 | -662.444975 | -89.6 | -662.261789 | -662.310868 | 4.9 | -658.579563 | -2.814871 | -661.294463 |
| conformer_2 | -662.411005 | 0.182831 | 0.130877 | -662.444484 | -87.9 | -662.261653 | -662.310595 | 5.6 | -658.581367 | -2.813052 | -661.294009 |
| conformer_16 | -662.400768 | 0.182439 | 0.129083 | -662.443816 | -113.0 | -662.261377 | -662.311721 | 2.7 | -658.571231 | -2.813508 | -661.295691 |
| conformer_5 | -662.406978 | 0.182763 | 0.129772 | -662.442723 | -93.8 | -662.259960 | -662.309939 | 7.3 | -658.577271 | -2.813000 | -661.293231 |
| conformer_7 | -662.406430 | 0.182845 | 0.130390 | -662.442361 | -94.3 | -662.259516 | -662.308959 | 9.9 | -658.575843 | -2.814425 | -661.292797 |
| conformer_4 | -662.407181 | 0.182847 | 0.130097 | -662.442123 | -91.7 | -662.259276 | -662.309014 | 9.8 | -658.578076 | -2.812661 | -661.292569 |
| conformer_10 | -662.404823 | 0.182702 | 0.129992 | -662.441740 | -96.9 | -662.259038 | -662.308736 | 10.5 | -658.574089 | -2.814485 | -661.292487 |
| conformer_6 | -662.407561 | 0.182672 | 0.129464 | -662.441242 | -88.4 | -662.258570 | -662.308766 | 10.4 | -658.579614 | -2.812061 | -661.292879 |
| conformer_12 | -662.395451 | 0.182244 | 0.127123 | -662.441115 | -119.9 | -662.258871 | -662.310980 | 4.6 | -658.565494 | -2.813909 | -661.294932 |
| conformer_8 | -662.406348 | 0.182778 | 0.129973 | -662.440633 | -90.0 | -662.257855 | -662.307648 | 13.3 | -658.576593 | -2.813945 | -661.291838 |
| conformer_14 | -662.394662 | 0.182109 | 0.127403 | -662.440034 | -119.1 | -662.257925 | -662.309619 | 8.2 | -658.566571 | -2.812655 | -661.294182 |

[a]: Single-point calculation in aqueous phase with SMD model.

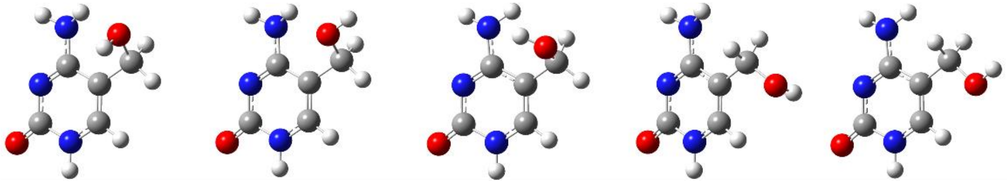

**Figure S12.** B3LYP-D3/def2-TZVPP optimized geometries of conformers for 5-hydroxymethylcytosine (**5hmC**).

**Table S10.** Conformers of gas-phase optimized 5-hydroxymethylcytosine (**5hmC**) at the B3LYP-D3/def2-TZVPP level of theory followed by aqueous phase single-point calculation. The columns display total energy without zero-point correction ($E_{Tot}$), thermal correction to enthalpy (δH), Gibbs free energy (δG), total energy without zero-point correction ($E_{Tot,W}$), Gibbs free energy ($G_{298,W}$) in water (W), total single-point energy ($E_{CBS}$) calculated at DLPNO-CCSD(T)/CBS level of theory, and their corresponding free energy $G_{CBS,W}$. $G_{298,W}$ and $G_{CBS,W}$ have been corrected to the standard state of 1 mol/L by addition of +7.908 kJ/mol. $\Delta G_{Solv}$ represents the Gibbs free energy of solvation. The data are arranged in the ascending numeric order of $E_{Tot,W}$. $\Delta G_{298,W}$ represents the respective energy difference to the lowest structure. Only conformers within the 24 kJ/mol (6 kcal/mol) energy window above the lowest in CREST are included in initial conformer sampling. Duplicates of the same structure are excluded. The overall optimum is marked bold.

| | **B3LYP-D3/def2-TZVPP** | | | **SMD($H_2O$)/B3LYP-D3/def2-TZVPP[a]** | | | | | **DLPNO-CCSD(T)/CBS** | | |
|---|---|---|---|---|---|---|---|---|---|---|---|
| **5hmC No.** | **$E_{Tot}$** (Hartree) | **δH** (Hartree) | **δG** (Hartree) | **$E_{Tot,W}$** (Hartree) | **$\Delta G_{Solv}$** (kJ/mol) | **$H_{298,W}$** (Hartree) | **$G_{298,W}$** (Hartree) | **$\Delta G_{298,W}$** (kJ/mol) | **$E_{CBS,HF}$** (Hartree) | **$E_{CBS,C}$** (Hartree) | **$G_{CBS,W}$** (Hartree) |
| **conformer_1** | **-509.694654** | **0.142043** | **0.098546** | **-509.728660** | **-89.3** | **-509.586617** | **-509.627102** | **0** | **-506.740818** | **-2.172386** | **-508.845652** |
| conformer_2 | -509.692566 | 0.141943 | 0.098366 | -509.727183 | -90.9 | -509.585240 | -509.625805 | 3.4 | -506.739355 | -2.172144 | -508.844739 |
| conformer_3 | -509.689268 | 0.141826 | 0.097581 | -509.725824 | -96.0 | -509.583998 | -509.625231 | 4.9 | -506.735798 | -2.172362 | -508.844123 |
| conformer_4 | -509.688441 | 0.141776 | 0.097365 | -509.725804 | -98.1 | -509.584028 | -509.625427 | 4.4 | -506.735622 | -2.171677 | -508.844285 |
| conformer_5 | -509.688129 | 0.141576 | 0.096764 | -509.724763 | -96.2 | -509.583187 | -509.624987 | 5.6 | -506.736446 | -2.171085 | -508.844389 |

[a]: Single-point calculation in aqueous phase with SMD model.

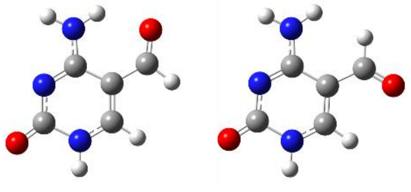

**Figure S13.** B3LYP-D3/def2-TZVPP optimized geometries of conformers for 5-formylcytosine (**5fC**).

**Table S11.** Conformers of gas-phase optimized 5-formylcytosine (**5fC**) at the B3LYP-D3/def2-TZVPP level of theory followed by aqueous phase single-point calculation. The columns display total energy without zero-point correction ($E_{Tot}$), thermal correction to enthalpy (δH), Gibbs free energy (δG), total energy without zero-point correction ($E_{Tot,W}$), Gibbs free energy ($G_{298,W}$) in water (W), total single-point energy ($E_{CBS}$) calculated at DLPNO-CCSD(T)/CBS level of theory, and their corresponding free energy $G_{CBS,W}$. $G_{298,W}$ and $G_{CBS,W}$ have been corrected to the standard state of 1 mol/L by addition of +7.908 kJ/mol. $\Delta G_{Solv}$ represents the Gibbs free energy of solvation. The data are arranged in the ascending numeric order of $E_{Tot,W}$. $\Delta G_{298,W}$ represents the respective energy difference to the lowest structure. Only conformers within the 24 kJ/mol (6 kcal/mol) energy window above the lowest in CREST are included in initial conformer sampling. Duplicates of the same structure are excluded. The overall optimum is marked bold.

| | B3LYP-D3/def2-TZVPP | | | SMD($H_2O$)/B3LYP-D3/def2-TZVPP[a] | | | | | DLPNO-CCSD(T)/CBS | | |
|---|---|---|---|---|---|---|---|---|---|---|---|
| **5fC No.** | **$E_{Tot}$** (Hartree) | **δH** (Hartree) | **δG** (Hartree) | **$E_{Tot,W}$** (Hartree) | **$\Delta G_{Solv}$** (kJ/mol) | **$H_{298,W}$** (Hartree) | **$G_{298,W}$** (Hartree) | **$\Delta G_{298,W}$** (kJ/mol) | **$E_{CBS,HF}$** (Hartree) | **$E_{CBS,C}$** (Hartree) | **$G_{CBS,W}$** (Hartree) |
| **conformer_1** | **-508.491666** | **0.117684** | **0.075559** | **-508.521492** | **-78.3** | **-508.403808** | **-508.442921** | **0** | **-505.582485** | **-2.125234** | **-507.658974** |
| conformer_2 | -508.481782 | 0.117328 | 0.073134 | -508.514541 | -86.0 | -508.397213 | -508.438395 | 11.9 | -505.574374 | -2.123434 | -507.654422 |

[a]: Single-point calculation in aqueous phase with SMD model.

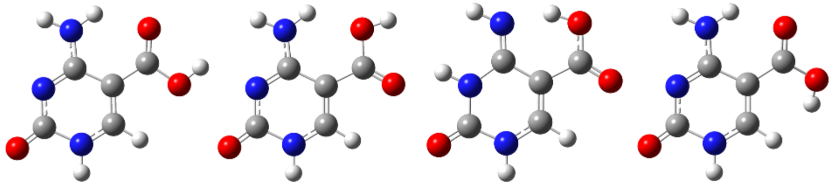

**Figure S14.** B3LYP-D3/def2-TZVPP optimized geometries of conformers for 5-carboxylcytosine (**5caC**).

**Table S12.** Conformers of gas-phase optimized 5-carboxylcytosine (**5caC**) at the B3LYP-D3/def2-TZVPP level of theory followed by aqueous phase single-point calculation. The columns display total energy without zero-point correction ($E_{Tot}$), thermal correction to enthalpy (δH), Gibbs free energy (δG), total energy without zero-point correction ($E_{Tot,W}$), Gibbs free energy ($G_{298,W}$) in water (W), total single-point energy ($E_{CBS}$) calculated at DLPNO-CCSD(T)/CBS level of theory, and their corresponding free energy $G_{CBS,W}$. $G_{298,W}$ and $G_{CBS,W}$ have been corrected to the standard state of 1 mol/L by addition of +7.908 kJ/mol. $\Delta G_{Solv}$ represents the Gibbs free energy of solvation. The data are arranged in the ascending numeric order of $E_{Tot,W}$. $\Delta G_{298,W}$ represents the respective energy difference to the lowest structure. Only conformers within the 24 kJ/mol (6 kcal/mol) energy window above the lowest in CREST are included in initial conformer sampling. Duplicates of the same structure are excluded. The overall optimum is marked bold.

| **5caC No.** | **B3LYP-D3/def2-TZVPP** $E_{Tot}$ (Hartree) | **δH** (Hartree) | **δG** (Hartree) | **SMD(H₂O)/B3LYP-D3/def2-TZVPP[a]** $E_{Tot,W}$ (Hartree) | $\Delta G_{Solv}$ (kJ/mol) | $H_{298,W}$ (Hartree) | $G_{298,W}$ (Hartree) | $\Delta G_{298,W}$ (kJ/mol) | **DLPNO-CCSD(T)/CBS** $E_{CBS,HF}$ (Hartree) | $E_{CBS,C}$ (Hartree) | $G_{CBS,W}$ (Hartree) |
|---|---|---|---|---|---|---|---|---|---|---|---|
| **conformer_1** | **-583.779532** | **0.124166** | **0.079604** | **-583.810296** | **-80.8** | **-583.686130** | **-583.727680** | **0** | **-580.514377** | **-2.394689** | **-582.857215** |
| conformer_2 | -583.775473 | 0.124122 | 0.079178 | -583.807575 | -84.3 | -583.683453 | -583.725385 | 6.0 | -580.510307 | -2.394526 | -582.854746 |
| conformer_4 | -583.765791 | 0.123745 | 0.078854 | -583.804643 | -102.0 | -583.680898 | -583.722777 | 12.9 | -580.497909 | -2.397918 | -582.852814 |
| conformer_3 | -583.773088 | 0.124301 | 0.080409 | -583.803086 | -78.8 | -583.678785 | -583.719665 | 21.0 | -580.505302 | -2.399492 | -582.851372 |

[a]: Single-point calculation in aqueous phase with SMD model.

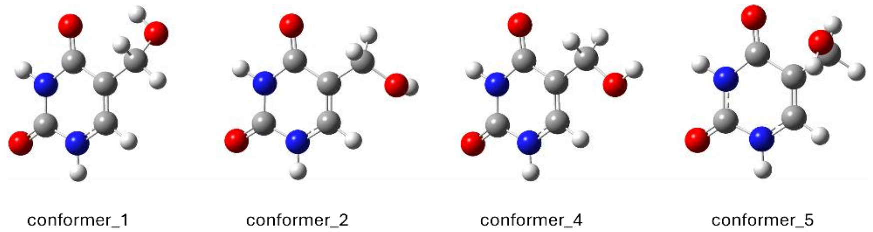

**Figure S15.** B3LYP-D3/def2-TZVPP optimized geometries of conformers for 5-hydroxymethyluracil (**5hmU**).

**Table S13.** Conformers of gas-phase optimized 5-hydroxymethyluracil (**5hmU**) at the B3LYP-D3/def2-TZVPP level of theory followed by aqueous phase single-point calculation. The columns display total energy without zero-point correction ($E_{Tot}$), thermal correction to enthalpy (δH), Gibbs free energy (δG), total energy without zero-point correction ($E_{Tot,W}$), Gibbs free energy ($G_{298,W}$) in water (W), total single-point energy ($E_{CBS}$) calculated at DLPNO-CCSD(T)/CBS level of theory, and their corresponding free energy $G_{CBS,W}$. $G_{298,W}$ and $G_{CBS,W}$ have been corrected to the standard state of 1 mol/L by addition of +7.908 kJ/mol. $\Delta G_{Solv}$ represents the Gibbs free energy of solvation. The data are arranged in the ascending numeric order of $E_{Tot,W}$. $\Delta G_{298,W}$ represents the respective energy difference to the lowest structure. Only conformers within the 24 kJ/mol (6 kcal/mol) energy window above the lowest in CREST are included in initial conformer sampling. Duplicates of the same structure are excluded. The overall optimum is marked bold.

| | **B3LYP-D3/def2-TZVPP** | | | **SMD($H_2O$)/B3LYP-D3/def2-TZVPP**[a] | | | | | **DLPNO-CCSD(T)/CBS** | | |
|---|---|---|---|---|---|---|---|---|---|---|---|
| **5hmU No.** | **$E_{Tot}$ (Hartree)** | **δH (Hartree)** | **δG (Hartree)** | **$E_{Tot,W}$ (Hartree)** | **$\Delta G_{Solv}$ (kJ/mol)** | **$H_{298,W}$ (Hartree)** | **$G_{298,W}$ (Hartree)** | **$\Delta G_{298,W}$ (kJ/mol)** | **$E_{CBS,HF}$ (Hartree)** | **$E_{CBS,C}$ (Hartree)** | **$G_{CBS,W}$ (Hartree)** |
| **conformer_1** | **-529.590156** | **0.130046** | **0.086799** | **-529.616085** | **-68.1** | **-529.486039** | **-529.526274** | **0** | **-526.604259** | **-2.195477** | **-528.735853** |
| conformer 5 | -529.584124 | 0.129789 | 0.085797 | -529.615046 | -81.2 | -529.485257 | -529.526237 | 0.1 | -526.598368 | -2.195413 | -528.735893 |
| conformer 2 | -529.587412 | 0.129812 | 0.085955 | -529.614210 | -70.4 | -529.484398 | -529.525243 | 2.7 | -526.601871 | -2.194935 | -528.734637 |
| conformer 4 | -529.588097 | 0.129726 | 0.085718 | -529.613519 | -66.7 | -529.483793 | -529.524789 | 3.9 | -526.603498 | -2.194382 | -528.734572 |

[a]: Single-point calculation in aqueous phase with SMD model.

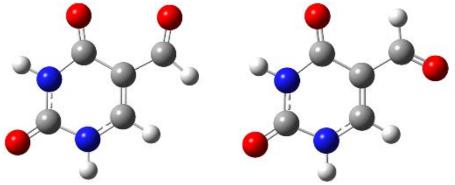

**Figure S16.** B3LYP-D3/def2-TZVPP optimized geometries of conformers for 5-formyluracil (**5fU**).

**Table S14.** Conformers of gas-phase optimized 5-formyluracil (**5fU**) at the B3LYP-D3/def2-TZVPP level of theory followed by aqueous phase single-point calculation. The columns display total energy without zero-point correction ($E_{Tot}$), thermal correction to enthalpy (δH), Gibbs free energy (δG), total energy without zero-point correction ($E_{Tot,W}$), Gibbs free energy ($G_{298,W}$) in water (W), total single-point energy ($E_{CBS}$) calculated at DLPNO-CCSD(T)/CBS level of theory, and their corresponding free energy $G_{CBS,W}$. $G_{298,W}$ and $G_{CBS,W}$ have been corrected to the standard state of 1 mol/L by addition of +7.908 kJ/mol. $\Delta G_{Solv}$ represents the Gibbs free energy of solvation. The data are arranged in the ascending numeric order of $E_{Tot,W}$. $\Delta G_{298,W}$ represents the respective energy difference to the lowest structure. Only conformers within the 24 kJ/mol (6 kcal/mol) energy window above the lowest in CREST are included in initial conformer sampling. Duplicates of the same structure are excluded. The overall optimum is marked bold.

| | **B3LYP-D3/def2-TZVPP** | | | **SMD($H_2O$)/B3LYP-D3/def2-TZVPP[a]** | | | | | **DLPNO-CCSD(T)/CBS** | | |
|---|---|---|---|---|---|---|---|---|---|---|---|
| **5fU No.** | **$E_{Tot}$** (Hartree) | **δH** (Hartree) | **δG** (Hartree) | **$E_{Tot,W}$** (Hartree) | **$\Delta G_{Solv}$** (kJ/mol) | **$H_{298,W}$** (Hartree) | **$G_{298,W}$** (Hartree) | **$\Delta G_{298,W}$** (kJ/mol) | **$E_{CBS,HF}$** (Hartree) | **$E_{CBS,C}$** (Hartree) | **$G_{CBS,W}$** (Hartree) |
| conformer_2 | -528.377221 | 0.105589 | 0.063191 | -528.401381 | -63.4 | -528.295792 | -528.335178 | 0.8 | -525.438085 | -2.147268 | -527.543310 |
| **conformer_1** | **-528.369075** | **0.105223** | **0.062527** | **-528.401035** | **-83.9** | **-528.295812** | **-528.335496** | **0** | **-525.428418** | **-2.149230** | **-527.544069** |

[a]: Single-point calculation in aqueous phase with SMD model.

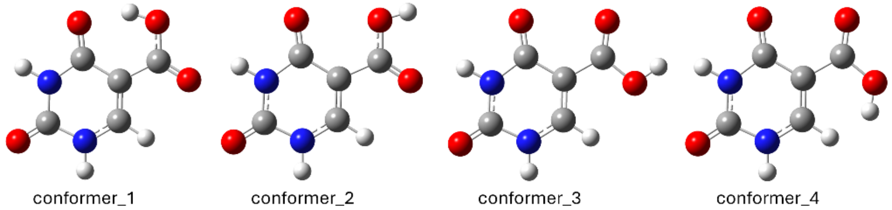

**Figure S17.** B3LYP-D3/def2-TZVPP optimized geometries of conformers for 5-carboxyluracil (**5caU**).

**Table S15.** Conformers of gas-phase optimized 5-carboxyluracil (**5caU**) at the B3LYP-D3/def2-TZVPP level of theory followed by aqueous phase single-point calculation. The columns display total energy without zero-point correction ($E_{Tot}$), thermal correction to enthalpy (δH), Gibbs free energy (δG), total energy without zero-point correction ($E_{Tot,W}$), Gibbs free energy ($G_{298,W}$) in water (W), total single-point energy ($E_{CBS}$) calculated at DLPNO-CCSD(T)/CBS level of theory, and their corresponding free energy $G_{CBS,W}$. $G_{298,W}$ and $G_{CBS,W}$ have been corrected to the standard state of 1 mol/L by addition of +7.908 kJ/mol. $\Delta G_{Solv}$ represents the Gibbs free energy of solvation. The data are arranged in the ascending numeric order of $E_{Tot,W}$. $\Delta G_{298,W}$ represents the respective energy difference to the lowest structure. Only conformers within the 24 kJ/mol (6 kcal/mol) energy window above the lowest in CREST are included in initial conformer sampling. Duplicates of the same structure are excluded. The overall optimum is marked bold.

| | B3LYP-D3/def2-TZVPP | | | SMD($H_2O$)/B3LYP-D3/def2-TZVPP[a] | | | | | DLPNO-CCSD(T)/CBS | | |
|---|---|---|---|---|---|---|---|---|---|---|---|
| **5caU No.** | **$E_{Tot}$** (Hartree) | **δH** (Hartree) | **δG** (Hartree) | **$E_{Tot,W}$** (Hartree) | **$\Delta G_{Solv}$** (kJ/mol) | **$H_{298,W}$** (Hartree) | **$G_{298,W}$** (Hartree) | **$\Delta G_{298,W}$** (kJ/mol) | **$E_{CBS,HF}$** (Hartree) | **$E_{CBS,C}$** (Hartree) | **$G_{CBS,W}$** (Hartree) |
| **conformer_1** | **-603.669696** | **0.112187** | **0.068539** | **-603.697177** | **-72.2** | **-603.584990** | **-603.625626** | **0** | **-600.369687** | **-2.421361** | **-602.746978** |
| conformer_2 | -603.661029 | 0.112020 | 0.066968 | -603.690814 | -78.2 | -603.578794 | -603.620834 | 12.6 | -600.363939 | -2.418454 | -602.742199 |
| conformer_3 | -603.658339 | 0.111849 | 0.066209 | -603.690700 | -85.0 | -603.578851 | -603.621479 | 10.9 | -600.360465 | -2.419575 | -602.743180 |
| conformer_4 | -603.646065 | 0.111622 | 0.066374 | -603.684591 | -101.1 | -603.572969 | -603.615205 | 27.4 | -600.345924 | -2.422098 | -602.737162 |

[a]: Single-point calculation in aqueous phase with SMD model.

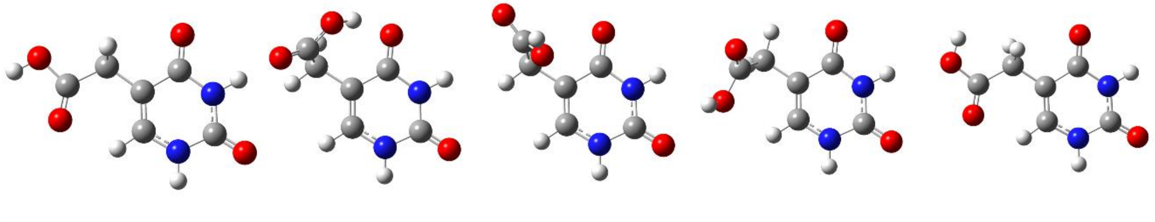

**Figure S18.** B3LYP-D3/def2-TZVPP optimized geometries of conformers for 5-carboxymethyluracil (**5cmU**).

**Table S16.** Conformers of gas-phase optimized 5-carboxymethyluracil (**5cmU**) at the B3LYP-D3/def2-TZVPP level of theory followed by aqueous phase single-point calculation. The columns display total energy without zero-point correction ($E_{Tot}$), thermal correction to enthalpy (δH), Gibbs free energy (δG), total energy without zero-point correction ($E_{Tot,W}$), Gibbs free energy ($G_{298,W}$) in water (W), total single-point energy ($E_{CBS}$) calculated at DLPNO-CCSD(T)/CBS level of theory, and their corresponding free energy $G_{CBS,W}$. $G_{298,W}$ and $G_{CBS,W}$ have been corrected to the standard state of 1 mol/L by addition of +7.908 kJ/mol. $\Delta G_{Solv}$ represents the Gibbs free energy of solvation. The data are arranged in the ascending numeric order of $E_{Tot,W}$. $\Delta G_{298,W}$ represents the respective energy difference to the lowest structure. Only conformers within the 24 kJ/mol (6 kcal/mol) energy window above the lowest in CREST are included in initial conformer sampling. Duplicates of the same structure are excluded. The overall optimum is marked bold.

| | **B3LYP-D3/def2-TZVPP** | | | **SMD($H_2O$)/B3LYP-D3/def2-TZVPP[a]** | | | | | **DLPNO-CCSD(T)/CBS** | | |
|---|---|---|---|---|---|---|---|---|---|---|---|
| **5cmU No.** | **$E_{Tot}$** (Hartree) | **δH** (Hartree) | **δG** (Hartree) | **$E_{Tot,W}$** (Hartree) | **$\Delta G_{Solv}$** (kJ/mol) | **$H_{298,W}$** (Hartree) | **$G_{298,W}$** (Hartree) | **$\Delta G_{298,W}$** (kJ/mol) | **$E_{CBS,HF}$** (Hartree) | **$E_{CBS,C}$** (Hartree) | **$G_{CBS,W}$** (Hartree) |
| conformer_2 | -642.998192 | 0.142057 | 0.095002 | -643.029047 | -81.0 | -642.886990 | -642.931033 | 0.1 | -639.415992 | -2.630713 | -641.979546 |
| **conformer_3** | **-642.995056** | **0.141599** | **0.092742** | **-643.026842** | **-83.5** | **-642.885243** | **-642.931088** | **0.0** | **-639.416842** | **-2.627645** | **-641.980518** |
| conformer_1 | -642.997261 | 0.141619 | 0.091287 | -643.023508 | -68.9 | -642.881889 | -642.929209 | 4.9 | -639.419199 | -2.627304 | -641.978451 |
| conformer_4 | -642.993550 | 0.141864 | 0.091957 | -643.022769 | -76.7 | -642.880905 | -642.927800 | 8.6 | -639.414254 | -2.628366 | -641.976870 |
| conformer_5 | -642.990022 | 0.141470 | 0.091590 | -643.021644 | -83.0 | -642.880174 | -642.927042 | 10.6 | -639.411282 | -2.628192 | -641.976494 |

[a]: Single-point calculation in aqueous phase with SMD model.

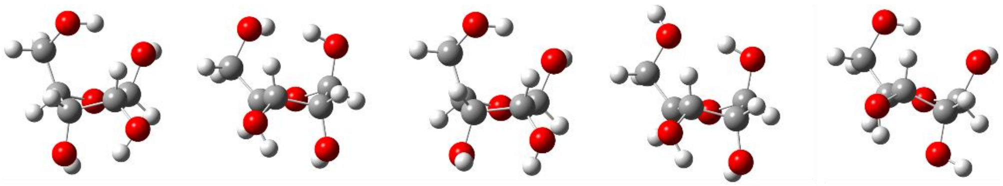

conformer_1 conformer_3 conformer_4 conformer_5 conformer_13

**Figure S19.** B3LYP-D3/def2-TZVPP optimized geometries of conformers for ribose (**rb**).

**Table S17.** Conformers of gas-phase optimized ribose (**rb**) at the B3LYP-D3/def2-TZVPP level of theory followed by aqueous phase single-point calculation. The columns display total energy without zero-point correction ($E_{Tot}$), thermal correction to enthalpy (δH), Gibbs free energy (δG), total energy without zero-point correction ($E_{Tot,W}$), Gibbs free energy ($G_{298,W}$) in water (W), total single-point energy ($E_{CBS}$) calculated at DLPNO-CCSD(T)/CBS level of theory, and their corresponding free energy $G_{CBS,W}$. $G_{298,W}$ and $G_{CBS,W}$ have been corrected to the standard state of 1 mol/L by addition of +7.908 kJ/mol. $\Delta G_{Solv}$ represents the Gibbs free energy of solvation. The data are arranged in the ascending numeric order of $E_{Tot,W}$. $\Delta G_{298,W}$ represents the respective energy difference to the lowest structure. Only conformers within the 24 kJ/mol (6 kcal/mol) energy window above the lowest in CREST are included in initial conformer sampling. Duplicates of the same structure are excluded. The overall optimum is marked bold.

| | B3LYP-D3/def2-TZVPP | | | SMD($H_2O$)/B3LYP-D3/def2-TZVPP[a] | | | | | DLPNO-CCSD(T)/CBS | | |
|---|---|---|---|---|---|---|---|---|---|---|---|
| **rb No.** | **$E_{Tot}$** (Hartree) | **δH** (Hartree) | **δG** (Hartree) | **$E_{Tot,W}$** (Hartree) | **$\Delta G_{Solv}$** (kJ/mol) | **$H_{298,W}$** (Hartree) | **$G_{298,W}$** (Hartree) | **$\Delta G_{298,W}$** (kJ/mol) | **$E_{CBS,HF}$** (Hartree) | **$E_{CBS,C}$** (Hartree) | **$G_{CBS,W}$** (Hartree) |
| conformer_6 | -572.914169 | 0.175027 | 0.129285 | -572.942252 | -73.7 | -572.767225 | -572.809955 | 2.9 | -569.719089 | -2.389356 | -572.004231 |
| conformer_66 | -572.915161 | 0.174926 | 0.128628 | -572.942244 | -71.1 | -572.767318 | -572.810604 | 1.2 | -569.722553 | -2.387387 | -572.005384 |
| **conformer_13** | **-572.913711** | **0.174756** | **0.127858** | **-572.941939** | **-74.1** | **-572.767183** | **-572.811069** | **0** | **-569.722387** | **-2.386200** | **-572.005945** |
| conformer_46 | -572.913320 | 0.175143 | 0.129354 | -572.941857 | -74.9 | -572.766714 | -572.809491 | 4.1 | -569.719682 | -2.388424 | -572.004277 |
| conformer_7 | -572.913779 | 0.175086 | 0.128953 | -572.941314 | -72.3 | -572.766228 | -572.809349 | 4.5 | -569.720795 | -2.387875 | -572.004239 |
| conformer_4 | -572.914421 | 0.175282 | 0.129553 | -572.941290 | -70.5 | -572.766008 | -572.808725 | 6.2 | -569.719054 | -2.389879 | -572.003236 |
| conformer_26 | -572.911705 | 0.174931 | 0.127248 | -572.941013 | -76.9 | -572.766082 | -572.810753 | 0.8 | -569.721909 | -2.385083 | -572.006040 |
| conformer_1 | -572.914745 | 0.175313 | 0.130317 | -572.940993 | -68.9 | -572.765680 | -572.807664 | 8.9 | -569.718074 | -2.391333 | -572.002326 |
| conformer_3 | -572.914494 | 0.175223 | 0.129700 | -572.940917 | -69.4 | -572.765694 | -572.808205 | 7.5 | -569.718603 | -2.390395 | -572.002709 |
| conformer_24 | -572.913413 | 0.174921 | 0.129324 | -572.940907 | -72.2 | -572.765986 | -572.808571 | 6.6 | -569.717582 | -2.390331 | -572.003072 |
| conformer_11 | -572.913026 | 0.174923 | 0.128617 | -572.940813 | -73.0 | -572.765890 | -572.809184 | 4.9 | -569.720956 | -2.387248 | -572.004362 |
| conformer_25 | -572.909467 | 0.175115 | 0.129866 | -572.940692 | -82.0 | -572.765577 | -572.807814 | 8.5 | -569.712574 | -2.391117 | -572.002038 |
| conformer_5 | -572.914557 | 0.175028 | 0.129666 | -572.940628 | -68.4 | -572.765600 | -572.807950 | 8.2 | -569.719445 | -2.389992 | -572.002831 |
| conformer_78 | -572.912321 | 0.174825 | 0.128746 | -572.940490 | -74.0 | -572.765665 | -572.808732 | 6.1 | -569.719661 | -2.388030 | -572.004102 |
| conformer_62 | -572.909504 | 0.174902 | 0.127432 | -572.940339 | -81.0 | -572.765437 | -572.809895 | 3.1 | -569.718795 | -2.385801 | -572.004987 |
| conformer_44 | -572.912235 | 0.174700 | 0.127409 | -572.940301 | -73.7 | -572.765601 | -572.809880 | 3.1 | -569.723221 | -2.384577 | -572.005443 |
| conformer_35 | -572.909060 | 0.174753 | 0.127925 | -572.940204 | -81.8 | -572.765451 | -572.809267 | 4.7 | -569.715867 | -2.387730 | -572.003805 |
| conformer_12 | -572.913218 | 0.174733 | 0.128323 | -572.940033 | -70.4 | -572.765300 | -572.808698 | 6.2 | -569.721289 | -2.387371 | -572.004140 |
| conformer_31 | -572.909719 | 0.174781 | 0.128136 | -572.939872 | -79.2 | -572.765091 | -572.808724 | 6.2 | -569.717891 | -2.387031 | -572.003927 |
| conformer_22 | -572.910769 | 0.175143 | 0.129796 | -572.939830 | -76.3 | -572.764687 | -572.807022 | 10.6 | -569.715178 | -2.390372 | -572.001803 |
| conformer_14 | -572.914090 | 0.174995 | 0.128615 | -572.939518 | -66.8 | -572.764523 | -572.807891 | 8.3 | -569.721023 | -2.388164 | -572.002989 |
| conformer_50 | -572.908797 | 0.174476 | 0.126821 | -572.939449 | -80.5 | -572.764973 | -572.809616 | 3.8 | -569.720011 | -2.384338 | -572.005168 |
| conformer_85 | -572.909534 | 0.174731 | 0.128028 | -572.939448 | -78.5 | -572.764717 | -572.808408 | 7.0 | -569.718357 | -2.386629 | -572.003859 |
| conformer_23 | -572.910686 | 0.175052 | 0.128971 | -572.939391 | -75.4 | -572.764339 | -572.807408 | 9.6 | -569.717058 | -2.388981 | -572.002762 |
| conformer_53 | -572.911157 | 0.174600 | 0.127001 | -572.939308 | -73.9 | -572.764708 | -572.809295 | 4.7 | -569.722102 | -2.384782 | -572.005021 |
| conformer_63 | -572.907046 | 0.174802 | 0.128159 | -572.939308 | -84.7 | -572.764506 | -572.808137 | 7.7 | -569.712567 | -2.389177 | -572.002835 |
| conformer_52 | -572.911235 | 0.174704 | 0.127979 | -572.939005 | -72.9 | -572.764301 | -572.808014 | 8.0 | -569.720535 | -2.385919 | -572.003233 |
| conformer_70 | -572.908511 | 0.174754 | 0.128168 | -572.938916 | -79.8 | -572.764162 | -572.807736 | 8.8 | -569.715922 | -2.387724 | -572.002872 |
| conformer_42 | -572.909719 | 0.174558 | 0.126928 | -572.938898 | -76.6 | -572.764340 | -572.808958 | 5.5 | -569.719999 | -2.384867 | -572.004105 |
| conformer_83 | -572.907449 | 0.174717 | 0.126151 | -572.938874 | -82.5 | -572.764157 | -572.809711 | 3.6 | -569.717792 | -2.384824 | -572.004878 |

| | | | | | | | | | | | |
|---|---|---|---|---|---|---|---|---|---|---|---|
| conformer_19 | -572.909783 | 0.174892 | 0.128346 | -572.938746 | -76.0 | -572.763854 | -572.807388 | 9.7 | -569.717291 | -2.387566 | -572.002463 |
| conformer_20 | -572.910234 | 0.174709 | 0.127316 | -572.938746 | -74.9 | -572.764037 | -572.808418 | 7.0 | -569.720284 | -2.385502 | -572.003970 |
| conformer_75 | -572.904562 | 0.174508 | 0.126383 | -572.938668 | -89.5 | -572.764160 | -572.809273 | 4.7 | -569.712836 | -2.386144 | -572.003692 |
| conformer_47 | -572.908863 | 0.174674 | 0.127142 | -572.938628 | -78.1 | -572.763954 | -572.808474 | 6.8 | -569.718449 | -2.386140 | -572.004200 |
| conformer_91 | -572.905779 | 0.174644 | 0.126283 | -572.938613 | -86.2 | -572.763969 | -572.809318 | 4.6 | -569.716250 | -2.384433 | -572.004221 |
| conformer_39 | -572.912880 | 0.174808 | 0.128096 | -572.938551 | -67.4 | -572.763743 | -572.807443 | 9.5 | -569.723299 | -2.385080 | -572.002943 |
| conformer_17 | -572.908222 | 0.174864 | 0.126862 | -572.938547 | -79.6 | -572.763683 | -572.808673 | 6.3 | -569.717442 | -2.385887 | -572.003780 |
| conformer_86 | -572.903503 | 0.174355 | 0.126037 | -572.938503 | -91.9 | -572.764148 | -572.809454 | 4.2 | -569.711862 | -2.386032 | -572.003846 |
| conformer_37 | -572.911295 | 0.174867 | 0.128196 | -572.938475 | -71.4 | -572.763608 | -572.807267 | 10.0 | -569.719153 | -2.386954 | -572.002079 |
| conformer_90 | -572.907223 | 0.174546 | 0.127208 | -572.938465 | -82.0 | -572.763919 | -572.808245 | 7.4 | -569.717515 | -2.385012 | -572.003549 |
| conformer_38 | -572.910571 | 0.175014 | 0.128804 | -572.938459 | -73.2 | -572.763445 | -572.806643 | 11.6 | -569.715599 | -2.389289 | -572.000960 |
| conformer_93 | -572.907853 | 0.174386 | 0.126499 | -572.938260 | -79.8 | -572.763874 | -572.808749 | 6.1 | -569.717978 | -2.385061 | -572.003935 |
| conformer_21 | -572.911210 | 0.175068 | 0.129831 | -572.938256 | -71.0 | -572.763188 | -572.805413 | 14.9 | -569.714899 | -2.390671 | -571.999773 |
| conformer_71 | -572.911410 | 0.174740 | 0.127758 | -572.938177 | -70.3 | -572.763437 | -572.807407 | 9.6 | -569.721638 | -2.384909 | -572.002545 |
| conformer_15 | -572.908803 | 0.174955 | 0.128540 | -572.937960 | -76.6 | -572.763005 | -572.806408 | 12.2 | -569.715634 | -2.388479 | -572.001718 |
| conformer_68 | -572.910368 | 0.174773 | 0.128050 | -572.937947 | -72.4 | -572.763174 | -572.806885 | 11.0 | -569.719436 | -2.385848 | -572.001802 |
| conformer_57 | -572.910502 | 0.174654 | 0.127445 | -572.937946 | -72.1 | -572.763292 | -572.807489 | 9.4 | -569.721853 | -2.384482 | -572.003323 |
| conformer_41 | -572.905443 | 0.174647 | 0.127354 | -572.937882 | -85.2 | -572.763235 | -572.807516 | 9.3 | -569.711638 | -2.388301 | -572.002013 |
| conformer_30 | -572.907921 | 0.174928 | 0.128347 | -572.937823 | -78.5 | -572.762895 | -572.806464 | 12.1 | -569.714636 | -2.388506 | -572.001685 |
| conformer_100 | -572.906032 | 0.174481 | 0.126663 | -572.937788 | -83.4 | -572.763307 | -572.808113 | 7.8 | -569.716872 | -2.384440 | -572.003393 |
| conformer_79 | -572.908241 | 0.174764 | 0.128091 | -572.937721 | -77.4 | -572.762957 | -572.806618 | 11.7 | -569.716614 | -2.387128 | -572.002119 |
| conformer_72 | -572.909765 | 0.174603 | 0.127800 | -572.937653 | -73.2 | -572.763050 | -572.806841 | 11.1 | -569.720177 | -2.385087 | -572.002339 |
| conformer_64 | -572.913360 | 0.174785 | 0.127995 | -572.937592 | -63.6 | -572.762807 | -572.806585 | 11.8 | -569.724659 | -2.384647 | -572.002531 |
| conformer_95 | -572.904329 | 0.174530 | 0.126308 | -572.937507 | -87.1 | -572.762977 | -572.808187 | 7.6 | -569.712884 | -2.386139 | -572.002880 |
| conformer_32 | -572.908053 | 0.175088 | 0.128960 | -572.937467 | -77.2 | -572.762379 | -572.805495 | 14.6 | -569.712254 | -2.390051 | -571.999747 |
| conformer_45 | -572.908572 | 0.174566 | 0.127089 | -572.937364 | -75.6 | -572.762798 | -572.807263 | 10.0 | -569.720304 | -2.383855 | -572.002850 |
| conformer_73 | -572.908136 | 0.174564 | 0.127491 | -572.937258 | -76.5 | -572.762694 | -572.806755 | 11.3 | -569.718901 | -2.384525 | -572.002044 |
| conformer_48 | -572.909858 | 0.174759 | 0.128350 | -572.937242 | -71.9 | -572.762483 | -572.805880 | 13.6 | -569.716023 | -2.388520 | -572.000564 |
| conformer_54 | -572.906434 | 0.174906 | 0.128356 | -572.937074 | -80.4 | -572.762168 | -572.805706 | 14.1 | -569.713037 | -2.388674 | -572.000983 |
| conformer_80 | -572.910623 | 0.174714 | 0.127720 | -572.936988 | -69.2 | -572.762274 | -572.806256 | 12.6 | -569.720636 | -2.385403 | -572.001672 |
| conformer_34 | -572.908829 | 0.174921 | 0.128357 | -572.936843 | -73.5 | -572.761922 | -572.805474 | 14.7 | -569.716397 | -2.388043 | -572.001085 |
| conformer_74 | -572.906245 | 0.174535 | 0.126109 | -572.936700 | -80.0 | -572.762165 | -572.807579 | 9.2 | -569.715953 | -2.385913 | -572.003200 |
| conformer_88 | -572.906497 | 0.174811 | 0.128150 | -572.936615 | -79.1 | -572.761804 | -572.805453 | 14.7 | -569.713875 | -2.388333 | -572.001163 |
| conformer_97 | -572.907706 | 0.174469 | 0.126694 | -572.936190 | -74.8 | -572.761721 | -572.806484 | 12.0 | -569.720033 | -2.383568 | -572.002379 |
| conformer_61 | -572.907744 | 0.174873 | 0.128234 | -572.936108 | -74.5 | -572.761235 | -572.804862 | 16.3 | -569.714050 | -2.388972 | -572.000140 |
| conformer_55 | -572.907734 | 0.174620 | 0.126803 | -572.936044 | -74.3 | -572.761424 | -572.806229 | 12.7 | -569.718095 | -2.385541 | -572.002131 |

[a]: Single-point calculation in aqueous phase with SMD model.

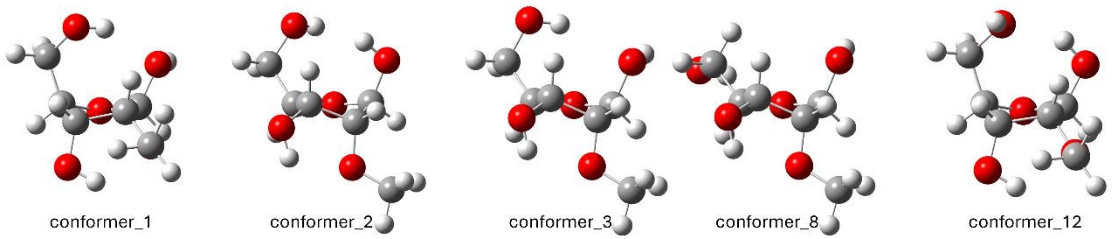

**Figure S20.** B3LYP-D3/def2-TZVPP optimized geometries of conformers for 2'-*O*-methylribose (**rbm**).

**Table S18.** Conformers of gas-phase optimized 2'-*O*-methylribose (**rbm**) at the B3LYP-D3/def2-TZVPP level of theory followed by aqueous phase single-point calculation. The columns display total energy without zero-point correction ($E_{Tot}$), thermal correction to enthalpy (δH), Gibbs free energy (δG), total energy without zero-point correction ($E_{Tot,W}$), Gibbs free energy ($G_{298,W}$) in water (W), total single-point energy ($E_{CBS}$) calculated at DLPNO-CCSD(T)/CBS level of theory, and their corresponding free energy $G_{CBS,W}$. $G_{298,W}$ and $G_{CBS,W}$ have been corrected to the standard state of 1 mol/L by addition of +7.908 kJ/mol. $\Delta G_{Solv}$ represents the Gibbs free energy of solvation. The data are arranged in the ascending numeric order of $E_{Tot,W}$. $\Delta G_{298,W}$ represents the respective energy difference to the lowest structure. Only conformers within the 24 kJ/mol (6 kcal/mol) energy window above the lowest in CREST are included in initial conformer sampling. Duplicates of the same structure are excluded. The overall optimum is marked bold.

| | B3LYP-D3/def2-TZVPP | | | SMD(H2O)/B3LYP-D3/def2-TZVPP[a] | | | | | DLPNO-CCSD(T)/CBS | | |
|---|---|---|---|---|---|---|---|---|---|---|---|
| **rbm No.** | $E_{Tot}$ (Hartree) | δH (Hartree) | δG (Hartree) | $E_{Tot,W}$ (Hartree) | $\Delta G_{Solv}$ (kJ/mol) | $H_{298,W}$ (Hartree) | $G_{298,W}$ (Hartree) | $\Delta G_{298,W}$ (kJ/mol) | $E_{CBS,HF}$ (Hartree) | $E_{CBS,C}$ (Hartree) | $G_{CBS,W}$ (Hartree) |
| conformer_3 | -612.232256 | 0.204230 | 0.153791 | -612.255416 | -60.8 | -612.051186 | -612.098613 | 0.0 | -608.759757 | -2.591365 | -611.217480 |
| conformer_2 | -612.232213 | 0.204604 | 0.155155 | -612.254679 | -59.0 | -612.050075 | -612.096512 | 5.5 | -608.758142 | -2.593059 | -611.215500 |
| conformer_1 | -612.233033 | 0.204629 | 0.155613 | -612.254433 | -56.2 | -612.049804 | -612.095808 | 7.4 | -608.756621 | -2.595471 | -611.214867 |
| conformer_12 | -612.227931 | 0.204349 | 0.155113 | -612.253850 | -68.0 | -612.049501 | -612.095725 | 7.6 | -608.751023 | -2.595572 | -611.214390 |
| conformer_8 | -612.230756 | 0.204199 | 0.153604 | -612.253699 | -60.2 | -612.049500 | -612.097083 | 4.0 | -608.760818 | -2.589617 | -611.216761 |
| conformer_9 | -612.230393 | 0.204259 | 0.153912 | -612.253458 | -60.6 | -612.049199 | -612.096534 | 5.5 | -608.756635 | -2.592836 | -611.215611 |
| conformer_5 | -612.231589 | 0.204364 | 0.154772 | -612.253353 | -57.1 | -612.048989 | -612.095569 | 8.0 | -608.758506 | -2.592546 | -611.215031 |
| conformer_4 | -612.232657 | 0.204487 | 0.155306 | -612.253325 | -54.3 | -612.048838 | -612.095007 | 9.5 | -608.758408 | -2.593578 | -611.214336 |
| conformer_27 | -612.227279 | 0.203977 | 0.152838 | -612.252959 | -67.4 | -612.048982 | -612.097109 | 3.9 | -608.757522 | -2.589360 | -611.216713 |
| conformer_14 | -612.227847 | 0.204226 | 0.153589 | -612.252585 | -64.9 | -612.048359 | -612.095984 | 6.9 | -608.754953 | -2.592696 | -611.215786 |
| conformer_18 | -612.228175 | 0.204019 | 0.152720 | -612.252401 | -63.6 | -612.048382 | -612.096669 | 5.1 | -608.757525 | -2.589929 | -611.215948 |
| conformer_57 | -612.226880 | 0.203918 | 0.152747 | -612.252320 | -66.8 | -612.048402 | -612.096561 | 5.4 | -608.757641 | -2.589094 | -611.216416 |
| conformer_36 | -612.225548 | 0.203985 | 0.153241 | -612.252032 | -69.5 | -612.048047 | -612.095779 | 7.4 | -608.754094 | -2.590868 | -611.215193 |
| conformer_50 | -612.226631 | 0.204418 | 0.155536 | -612.251919 | -66.4 | -612.047501 | -612.093371 | 13.8 | -608.752037 | -2.593606 | -611.212383 |
| conformer_24 | -612.228306 | 0.204166 | 0.153669 | -612.251584 | -61.1 | -612.047418 | -612.094903 | 9.7 | -608.756656 | -2.590851 | -611.214104 |
| conformer_6 | -612.229900 | 0.204202 | 0.153756 | -612.251561 | -56.9 | -612.047359 | -612.094793 | 10.0 | -608.759222 | -2.589946 | -611.214062 |
| conformer_25 | -612.228226 | 0.204165 | 0.153223 | -612.251532 | -61.2 | -612.047367 | -612.095297 | 8.7 | -608.757419 | -2.590670 | -611.215160 |
| conformer_52 | -612.223961 | 0.203840 | 0.152721 | -612.251427 | -72.1 | -612.047587 | -612.095694 | 7.7 | -608.753036 | -2.590245 | -611.215015 |
| conformer_44 | -612.223961 | 0.203840 | 0.152721 | -612.251427 | -72.1 | -612.047587 | -612.095694 | 7.7 | -608.753036 | -2.590245 | -611.215015 |
| conformer_55 | -612.227363 | 0.203962 | 0.152523 | -612.250751 | -61.4 | -612.046789 | -612.095216 | 8.9 | -608.755674 | -2.590677 | -611.214204 |
| conformer_54 | -612.227363 | 0.203962 | 0.152523 | -612.250751 | -61.4 | -612.046789 | -612.095216 | 8.9 | -608.755674 | -2.590677 | -611.214204 |
| conformer_59 | -612.225555 | 0.204160 | 0.153810 | -612.250731 | -66.1 | -612.046571 | -612.093909 | 12.4 | -608.752151 | -2.592198 | -611.212703 |
| conformer_10 | -612.226269 | 0.204237 | 0.153541 | -612.250658 | -64.0 | -612.046421 | -612.094105 | 11.8 | -608.753225 | -2.592648 | -611.213709 |
| conformer_17 | -612.227291 | 0.204509 | 0.155347 | -612.250579 | -61.1 | -612.046070 | -612.092220 | 16.8 | -608.748563 | -2.596798 | -611.210290 |
| conformer_51 | -612.227291 | 0.204509 | 0.155347 | -612.250579 | -61.1 | -612.046070 | -612.092220 | 16.8 | -608.748563 | -2.596798 | -611.210290 |
| conformer_19 | -612.226254 | 0.204099 | 0.152425 | -612.250351 | -63.3 | -612.046252 | -612.094914 | 9.7 | -608.756049 | -2.589990 | -611.214699 |
| conformer_42 | -612.226865 | 0.204049 | 0.153417 | -612.250270 | -61.4 | -612.046221 | -612.093841 | 12.5 | -608.756146 | -2.590146 | -611.213268 |
| conformer_43 | -612.226865 | 0.204049 | 0.153417 | -612.250270 | -61.4 | -612.046221 | -612.093841 | 12.5 | -608.756146 | -2.590146 | -611.213268 |
| conformer_33 | -612.227604 | 0.204051 | 0.152640 | -612.249984 | -58.8 | -612.045933 | -612.094332 | 11.2 | -608.756046 | -2.590599 | -611.213372 |

| | | | | | | | | | | | |
|---|---|---|---|---|---|---|---|---|---|---|---|
| conformer_30 | -612.225877 | 0.204381 | 0.155359 | -612.249920 | -63.1 | -612.045539 | -612.091549 | 18.5 | -608.746770 | -2.597398 | -611.209840 |
| conformer_15 | -612.227063 | 0.204765 | 0.155922 | -612.249860 | -59.9 | -612.045095 | -612.090926 | 20.2 | -608.747935 | -2.597354 | -611.209151 |
| conformer_38 | -612.225113 | 0.203891 | 0.152792 | -612.249694 | -64.5 | -612.045803 | -612.093890 | 12.4 | -608.754882 | -2.589578 | -611.213237 |
| conformer_40 | -612.225113 | 0.203891 | 0.152792 | -612.249694 | -64.5 | -612.045803 | -612.093890 | 12.4 | -608.754882 | -2.589578 | -611.213237 |
| conformer_35 | -612.225476 | 0.204140 | 0.153090 | -612.249573 | -63.3 | -612.045433 | -612.093471 | 13.5 | -608.753167 | -2.592229 | -611.213391 |
| conformer_20 | -612.227048 | 0.204472 | 0.155706 | -612.249477 | -58.9 | -612.045005 | -612.090759 | 20.6 | -608.748791 | -2.596911 | -611.209413 |
| conformer_46 | -612.225493 | 0.204008 | 0.152310 | -612.249198 | -62.2 | -612.045190 | -612.093876 | 12.4 | -608.755872 | -2.589727 | -611.213982 |
| conformer_45 | -612.225493 | 0.204008 | 0.152310 | -612.249198 | -62.2 | -612.045190 | -612.093876 | 12.4 | -608.755872 | -2.589727 | -611.213982 |
| conformer_32 | -612.225593 | 0.204304 | 0.154434 | -612.249053 | -61.6 | -612.044749 | -612.091607 | 18.4 | -608.750300 | -2.594408 | -611.210721 |
| conformer_48 | -612.221904 | 0.204212 | 0.153980 | -612.248810 | -70.6 | -612.044598 | -612.091818 | 17.8 | -608.745309 | -2.594978 | -611.210201 |
| conformer_61 | -612.223188 | 0.204004 | 0.152523 | -612.248738 | -67.1 | -612.044734 | -612.093203 | 14.2 | -608.750420 | -2.592632 | -611.213067 |
| conformer_49 | -612.226525 | 0.204201 | 0.154027 | -612.248662 | -58.1 | -612.044461 | -612.091623 | 18.4 | -608.755912 | -2.589893 | -611.210903 |
| conformer_23 | -612.224245 | 0.204171 | 0.153107 | -612.248226 | -63.0 | -612.044055 | -612.092107 | 17.1 | -608.750535 | -2.593110 | -611.211506 |
| conformer_28 | -612.224245 | 0.204171 | 0.153106 | -612.248225 | -63.0 | -612.044054 | -612.092107 | 17.1 | -608.750535 | -2.593110 | -611.211507 |
| conformer_47 | -612.222221 | 0.204132 | 0.153929 | -612.248029 | -67.8 | -612.043897 | -612.091088 | 19.8 | -608.747220 | -2.594164 | -611.210251 |
| conformer_60 | -612.226515 | 0.204113 | 0.153689 | -612.247500 | -55.1 | -612.043387 | -612.090799 | 20.5 | -608.756934 | -2.589396 | -611.210614 |
| conformer_37 | -612.222963 | 0.204316 | 0.154460 | -612.247476 | -64.4 | -612.043160 | -612.090004 | 22.6 | -608.747413 | -2.594462 | -611.208916 |
| conformer_53 | -612.223600 | 0.204237 | 0.153532 | -612.247338 | -62.3 | -612.043101 | -612.090794 | 20.5 | -608.748309 | -2.594081 | -611.209584 |
| conformer_29 | -612.223600 | 0.204237 | 0.153532 | -612.247338 | -62.3 | -612.043101 | -612.090794 | 20.5 | -608.748309 | -2.594081 | -611.209584 |
| conformer_56 | -612.218636 | 0.204106 | 0.153580 | -612.247259 | -75.1 | -612.043153 | -612.090667 | 20.9 | -608.741829 | -2.595133 | -611.208993 |

[a]: Single-point calculation in aqueous phase with SMD model.

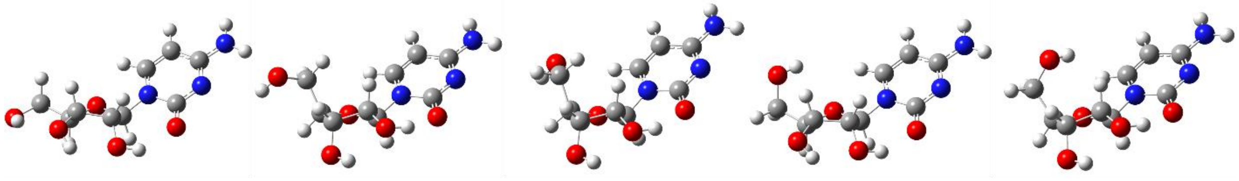

conformer_1 conformer_2 conformer_3 conformer_4 conformer_7

**Figure S21.** B3LYP-D3/def2-TZVPP optimized geometries of conformers for cytidine (**1rbC**).

**Table S19.** Conformers of gas-phase optimized cytidine (**1rbC**) at the B3LYP-D3/def2-TZVPP level of theory followed by aqueous phase single-point calculation. The columns display total energy without zero-point correction ($E_{Tot}$), thermal correction to enthalpy (δH), Gibbs free energy (δG), total energy without zero-point correction ($E_{Tot,W}$), Gibbs free energy ($G_{298,W}$) in water (W), total single-point energy ($E_{CBS}$) calculated at DLPNO-CCSD(T)/CBS level of theory, and their corresponding free energy $G_{CBS,W}$. $G_{298,W}$ and $G_{CBS,W}$ have been corrected to the standard state of 1 mol/L by addition of +7.908 kJ/mol. $\Delta G_{Solv}$ represents the Gibbs free energy of solvation. The data are arranged in the ascending numeric order of $E_{Tot,W}$. $\Delta G_{298,W}$ represents the respective energy difference to the lowest structure. Only conformers within the 24 kJ/mol (6 kcal/mol) energy window above the lowest in CREST are included in initial conformer sampling. Duplicates of the same structure are excluded. The overall optimum is marked bold.

| | B3LYP-D3/def2-TZVPP | | | SMD($H_2O$)/B3LYP-D3/def2-TZVPP[a] | | | | | DLPNO-CCSD(T)/CBS | | |
|---|---|---|---|---|---|---|---|---|---|---|---|
| **1rbC No.** | **$E_{Tot}$** (Hartree) | **δH** (Hartree) | **δG** (Hartree) | **$E_{Tot,W}$** (Hartree) | **$\Delta G_{Solv}$** (kJ/mol) | **$H_{298,W}$** (Hartree) | **$G_{298,W}$** (Hartree) | **$\Delta G_{298,W}$** (kJ/mol) | **$E_{CBS,HF}$** (Hartree) | **$E_{CBS,C}$** (Hartree) | **$G_{CBS,W}$** (Hartree) |
| **conformer_7** | **-891.565221** | **0.254695** | **0.194103** | **-891.604799** | **-103.9** | **-891.350104** | **-891.407684** | **0** | **-886.451845** | **-3.782145** | **-890.076454** |
| conformer_25 | -891.563627 | 0.254916 | 0.195898 | -891.605731 | -110.5 | -891.350815 | -891.406821 | 2.3 | -886.451786 | -3.780856 | -890.075836 |
| conformer_15 | -891.562518 | 0.254768 | 0.195391 | -891.605471 | -112.8 | -891.350703 | -891.407068 | 1.6 | -886.450539 | -3.780712 | -890.075802 |
| conformer_6 | -891.564129 | 0.254640 | 0.194280 | -891.603856 | -104.3 | -891.349216 | -891.406564 | 2.9 | -886.451808 | -3.781278 | -890.075520 |
| conformer_19 | -891.560777 | 0.254783 | 0.195480 | -891.605474 | -117.3 | -891.350691 | -891.406982 | 1.8 | -886.446882 | -3.782128 | -890.075214 |
| conformer_2 | -891.563042 | 0.254626 | 0.193970 | -891.602942 | -104.8 | -891.348316 | -891.405960 | 4.5 | -886.452157 | -3.780070 | -890.075145 |
| conformer_10 | -891.562510 | 0.254511 | 0.193903 | -891.602865 | -106.0 | -891.348354 | -891.405950 | 4.6 | -886.451199 | -3.780450 | -890.075090 |
| conformer_8 | -891.565219 | 0.254648 | 0.194880 | -891.604165 | -102.3 | -891.349517 | -891.406273 | 3.7 | -886.452608 | -3.781391 | -890.075052 |
| conformer_3 | -891.564468 | 0.254761 | 0.195507 | -891.603840 | -103.4 | -891.349079 | -891.405321 | 6.2 | -886.452584 | -3.781361 | -890.074799 |
| conformer_23 | -891.563068 | 0.254663 | 0.195473 | -891.603220 | -105.4 | -891.348557 | -891.404735 | 7.7 | -886.451868 | -3.781256 | -890.074791 |
| conformer_27 | -891.560288 | 0.254794 | 0.195200 | -891.604398 | -115.8 | -891.349604 | -891.406186 | 3.9 | -886.448300 | -3.780572 | -890.074770 |
| conformer_4 | -891.562886 | 0.254741 | 0.193160 | -891.600626 | -99.1 | -891.345885 | -891.404454 | 8.5 | -886.453586 | -3.779437 | -890.074591 |
| conformer_22 | -891.561537 | 0.254892 | 0.194936 | -891.603173 | -109.3 | -891.348281 | -891.405225 | 6.5 | -886.452502 | -3.778189 | -890.074379 |
| conformer_18 | -891.562491 | 0.254772 | 0.195367 | -891.603569 | -107.9 | -891.348797 | -891.405190 | 6.5 | -886.450566 | -3.780976 | -890.074241 |
| conformer_21 | -891.562174 | 0.254581 | 0.194119 | -891.601787 | -104.0 | -891.347206 | -891.404656 | 8.0 | -886.451848 | -3.779867 | -890.074197 |
| conformer_12 | -891.565774 | 0.254731 | 0.195169 | -891.602963 | -97.6 | -891.348232 | -891.404782 | 7.6 | -886.454194 | -3.780945 | -890.074147 |
| conformer_29 | -891.556819 | 0.254811 | 0.193660 | -891.601252 | -116.7 | -891.346441 | -891.404580 | 8.2 | -886.447329 | -3.778958 | -890.074048 |
| conformer_31 | -891.557991 | 0.254722 | 0.193771 | -891.600721 | -112.2 | -891.345999 | -891.403938 | 9.8 | -886.448614 | -3.779305 | -890.073866 |
| conformer_16 | -891.561358 | 0.254802 | 0.193480 | -891.600210 | -102.0 | -891.345408 | -891.403718 | 10.4 | -886.453484 | -3.778010 | -890.073855 |
| conformer_5 | -891.562654 | 0.254796 | 0.193869 | -891.600275 | -98.8 | -891.345479 | -891.403394 | 11.3 | -886.454527 | -3.778501 | -890.073768 |
| conformer_14 | -891.560458 | 0.254607 | 0.193590 | -891.599728 | -103.1 | -891.345121 | -891.403126 | 12.0 | -886.452730 | -3.778297 | -890.073695 |
| conformer_24 | -891.560039 | 0.254593 | 0.193870 | -891.599090 | -102.5 | -891.344497 | -891.402208 | 14.4 | -886.452784 | -3.778078 | -890.073031 |
| conformer_37 | -891.556241 | 0.254715 | 0.194681 | -891.601380 | -118.5 | -891.346665 | -891.403687 | 10.5 | -886.443629 | -3.781716 | -890.072791 |
| conformer_32 | -891.555475 | 0.254829 | 0.194120 | -891.600200 | -117.4 | -891.345371 | -891.403068 | 12.1 | -886.446102 | -3.778623 | -890.072318 |
| conformer_39 | -891.556779 | 0.254641 | 0.193434 | -891.598162 | -108.7 | -891.343521 | -891.401716 | 15.7 | -886.448210 | -3.778440 | -890.071588 |
| conformer_34 | -891.555395 | 0.254752 | 0.194254 | -891.598692 | -113.7 | -891.343940 | -891.401426 | 16.4 | -886.445807 | -3.779141 | -890.070978 |
| conformer_1 | -891.562372 | 0.254923 | 0.194558 | -891.598283 | -94.3 | -891.343360 | -891.400713 | 18.3 | -886.453828 | -3.778664 | -890.070833 |
| conformer_28 | -891.557771 | 0.254722 | 0.193273 | -891.596922 | -102.8 | -891.342200 | -891.400637 | 18.5 | -886.449667 | -3.778257 | -890.070789 |

[a]: Single-point calculation in aqueous phase with SMD model.

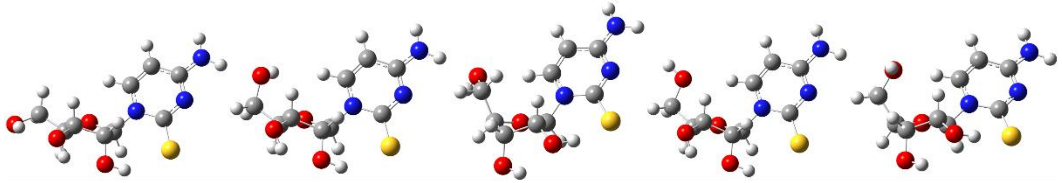

**Figure S18.** B3LYP-D3/def2-TZVPP optimized geometries of conformers for 2-thiocytidine (**1rb2thC**).

**Table S20.** Conformers of gas-phase optimized 2-thiocytidine (**1rb2thC**) at the B3LYP-D3/def2-TZVPP level of theory followed by aqueous phase single-point calculation. The columns display total energy without zero-point correction ($E_{Tot}$), thermal correction to enthalpy (δH), Gibbs free energy (δG), total energy without zero-point correction ($E_{Tot,W}$), Gibbs free energy ($G_{298,W}$) in water (W), total single-point energy ($E_{CBS}$) calculated at DLPNO-CCSD(T)/CBS level of theory, and their corresponding free energy $G_{CBS,W}$. $G_{298,W}$ and $G_{CBS,W}$ have been corrected to the standard state of 1 mol/L by addition of +7.908 kJ/mol. $\Delta G_{Solv}$ represents the Gibbs free energy of solvation. The data are arranged in the ascending numeric order of $E_{Tot,W}$. $\Delta G_{298,W}$ represents the respective energy difference to the lowest structure. Only conformers within the 24 kJ/mol (6 kcal/mol) energy window above the lowest in CREST are included in initial conformer sampling. Duplicates of the same structure are excluded. The overall optimum is marked bold.

| | B3LYP-D3/def2-TZVPP | | | SMD($H_2O$)/B3LYP-D3/def2-TZVPP[a] | | | | | DLPNO-CCSD(T)/CBS | | |
|---|---|---|---|---|---|---|---|---|---|---|---|
| **1rb2thC No.** | **$E_{Tot}$** (Hartree) | **δH** (Hartree) | **δG** (Hartree) | **$E_{Tot,W}$** (Hartree) | **$\Delta G_{Solv}$** (kJ/mol) | **$H_{298,W}$** (Hartree) | **$G_{298,W}$** (Hartree) | **$\Delta G_{298,W}$** (kJ/mol) | **$E_{CBS,HF}$** (Hartree) | **$E_{CBS,C}$** (Hartree) | **$G_{CBS,W}$** (Hartree) |
| conformer_9 | -1214.510975 | 0.252236 | 0.191626 | -1214.549342 | -100.7 | -1214.297106 | -1214.354704 | 3.6 | -1209.079501 | -3.736723 | -1212.659952 |
| conformer_24 | -1214.510682 | 0.252984 | 0.192007 | -1214.548791 | -100.1 | -1214.295807 | -1214.353772 | 6.0 | -1209.084028 | -3.733153 | -1212.660269 |
| conformer_14 | -1214.508972 | 0.252245 | 0.191168 | -1214.548783 | -104.5 | -1214.296538 | -1214.354603 | 3.8 | -1209.076947 | -3.737707 | -1212.660285 |
| conformer_4 | -1214.509693 | 0.252343 | 0.191912 | -1214.548646 | -102.3 | -1214.296303 | -1214.353722 | 6.1 | -1209.078221 | -3.737742 | -1212.659992 |
| **conformer_3** | **-1214.509607** | **0.252677** | **0.189552** | **-1214.548622** | **-102.4** | **-1214.295945** | **-1214.356058** | **0** | **-1209.082595** | **-3.733794** | **-1212.662840** |
| conformer_10 | -1214.507229 | 0.252207 | 0.190580 | -1214.548325 | -107.9 | -1214.296118 | -1214.354733 | 3.5 | -1209.077281 | -3.735762 | -1212.660549 |
| conformer_13 | -1214.511326 | 0.252249 | 0.191322 | -1214.548210 | -96.8 | -1214.295961 | -1214.353876 | 5.7 | -1209.081053 | -3.736160 | -1212.659763 |
| conformer_16 | -1214.507703 | 0.252128 | 0.190570 | -1214.547897 | -105.5 | -1214.295769 | -1214.354315 | 4.6 | -1209.073689 | -3.739706 | -1212.660007 |
| conformer_18 | -1214.508085 | 0.252333 | 0.192311 | -1214.547876 | -104.5 | -1214.295543 | -1214.352553 | 9.2 | -1209.077065 | -3.737860 | -1212.659392 |
| conformer_8 | -1214.510845 | 0.252958 | 0.191910 | -1214.547657 | -96.6 | -1214.294699 | -1214.352735 | 8.7 | -1209.085010 | -3.732749 | -1212.659649 |
| conformer_28 | -1214.509468 | 0.252698 | 0.191076 | -1214.547409 | -99.6 | -1214.294711 | -1214.353321 | 7.2 | -1209.083437 | -3.733350 | -1212.660640 |
| conformer_20 | -1214.506164 | 0.252104 | 0.190374 | -1214.547192 | -107.7 | -1214.295088 | -1214.353806 | 5.9 | -1209.076726 | -3.735591 | -1212.659959 |
| conformer_12 | -1214.507076 | 0.252496 | 0.190661 | -1214.546732 | -104.1 | -1214.294236 | -1214.353059 | 7.9 | -1209.081543 | -3.733135 | -1212.660661 |
| conformer_17 | -1214.506570 | 0.252414 | 0.190536 | -1214.546008 | -103.5 | -1214.293594 | -1214.352460 | 9.4 | -1209.081403 | -3.733038 | -1212.660330 |
| conformer_30 | -1214.496908 | 0.252422 | 0.189306 | -1214.545832 | -128.5 | -1214.293410 | -1214.353514 | 6.7 | -1209.068711 | -3.733176 | -1212.658493 |
| conformer_1 | -1214.509534 | 0.252780 | 0.191549 | -1214.545549 | -94.6 | -1214.292769 | -1214.350988 | 13.3 | -1209.082690 | -3.734015 | -1212.658158 |
| conformer_27 | -1214.500700 | 0.252775 | 0.191981 | -1214.545105 | -116.6 | -1214.292330 | -1214.350112 | 15.6 | -1209.070416 | -3.736136 | -1212.655965 |
| conformer_19 | -1214.502268 | 0.252646 | 0.191667 | -1214.543891 | -109.3 | -1214.291245 | -1214.349212 | 18.0 | -1209.071966 | -3.736600 | -1212.655511 |
| conformer_21 | -1214.502860 | 0.252357 | 0.189413 | -1214.543673 | -107.2 | -1214.291316 | -1214.351248 | 12.6 | -1209.077998 | -3.732389 | -1212.658775 |
| conformer_29 | -1214.497583 | 0.252745 | 0.190194 | -1214.542020 | -116.7 | -1214.289275 | -1214.348814 | 19.0 | -1209.069880 | -3.734283 | -1212.655394 |
| conformer_26 | -1214.498063 | 0.252624 | 0.190094 | -1214.540866 | -112.4 | -1214.288242 | -1214.347760 | 21.8 | -1209.070084 | -3.734985 | -1212.654766 |
| conformer_25 | -1214.497880 | 0.252548 | 0.189949 | -1214.536902 | -102.5 | -1214.284354 | -1214.343941 | 31.8 | -1209.071174 | -3.733985 | -1212.651219 |

[a]: Single-point calculation in aqueous phase with SMD model.

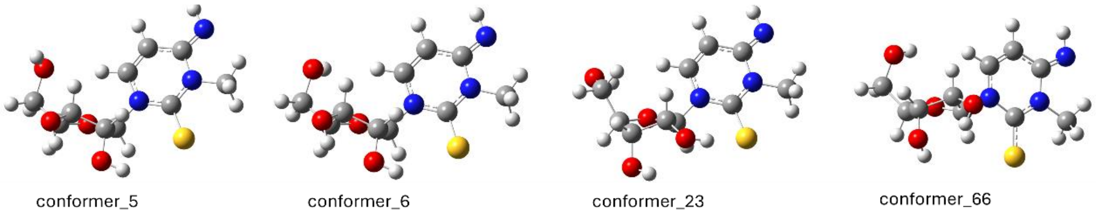

**Figure S23.** B3LYP-D3/def2-TZVPP optimized geometries of conformers for neutral 2-thio-3-methylcytidine (**1rb2th3mC**).

**Table S21.** Conformers of gas-phase optimized neutral 2-thio-3-methylcytidine (**1rb2th3mC**) at the B3LYP-D3/def2-TZVPP level of theory followed by aqueous phase single-point calculation. The columns display total energy without zero-point correction ($E_{Tot}$), thermal correction to enthalpy (δH), Gibbs free energy (δG), total energy without zero-point correction ($E_{Tot,W}$), Gibbs free energy ($G_{298,W}$) in water (W), total single-point energy ($E_{CBS}$) calculated at DLPNO-CCSD(T)/CBS level of theory, and their corresponding free energy $G_{CBS,W}$. $G_{298,W}$ and $G_{CBS,W}$ have been corrected to the standard state of 1 mol/L by addition of +7.908 kJ/mol. $\Delta G_{Solv}$ represents the Gibbs free energy of solvation. The data are arranged in the ascending numeric order of $E_{Tot,W}$. $\Delta G_{298,W}$ represents the respective energy difference to the lowest structure. Only conformers within the 24 kJ/mol (6 kcal/mol) energy window above the lowest in CREST are included in initial conformer sampling. Duplicates of the same structure are excluded. The overall optimum is marked bold.

| 1rb2th3mC No. | B3LYP-D3/def2-TZVPP $E_{Tot}$ (Hartree) | δH (Hartree) | δG (Hartree) | SMD(H₂O)/B3LYP-D3/def2-TZVPP[a] $E_{Tot,W}$ (Hartree) | $\Delta G_{Solv}$ (kJ/mol) | $H_{298,W}$ (Hartree) | $G_{298,W}$ (Hartree) | $\Delta G_{298,W}$ (kJ/mol) | DLPNO-CCSD(T)/CBS $E_{CBS,HF}$ (Hartree) | $E_{CBS,C}$ (Hartree) | $G_{CBS,W}$ (Hartree) |
|---|---|---|---|---|---|---|---|---|---|---|---|
| conformer_5 | -1253.826108 | 0.282751 | 0.219086 | -1253.853401 | -71.7 | -1253.570650 | -1253.631303 | 2.6 | -1248.112991 | -3.949351 | -1251.867537 |
| conformer_6 | -1253.825184 | 0.282673 | 0.218367 | -1253.853286 | -73.8 | -1253.570613 | -1253.631907 | 1.1 | -1248.111949 | -3.949490 | -1251.868162 |
| conformer_23 | -1253.823516 | 0.282230 | 0.218817 | -1253.852742 | -76.7 | -1253.570512 | -1253.630913 | 3.7 | -1248.106931 | -3.952554 | -1251.866882 |
| conformer_15 | -1253.825468 | 0.282272 | 0.219363 | -1253.852667 | -71.4 | -1253.570395 | -1253.630292 | 5.3 | -1248.108843 | -3.952243 | -1251.865911 |
| conformer_7 | -1253.826381 | 0.282709 | 0.219040 | -1253.852273 | -68.0 | -1253.569564 | -1253.630221 | 5.5 | -1248.114135 | -3.948886 | -1251.866862 |
| conformer_36 | -1253.823119 | 0.282084 | 0.217889 | -1253.852231 | -76.4 | -1253.570147 | -1253.631330 | 2.6 | -1248.106624 | -3.952548 | -1251.867382 |
| conformer_87 | -1253.818823 | 0.282508 | 0.218319 | -1253.852150 | -87.5 | -1253.569642 | -1253.630819 | 3.9 | -1248.104289 | -3.950717 | -1251.867001 |
| conformer_18 | -1253.821847 | 0.282606 | 0.218431 | -1253.852128 | -79.5 | -1253.569522 | -1253.630685 | 4.3 | -1248.107671 | -3.950575 | -1251.867084 |
| conformer_17 | -1253.821125 | 0.282500 | 0.217364 | -1253.852075 | -81.3 | -1253.569575 | -1253.631699 | 1.6 | -1248.106970 | -3.950608 | -1251.868152 |
| **conformer_66** | **-1253.815549** | **0.282359** | **0.216734** | **-1253.852054** | **-95.8** | **-1253.569695** | **-1253.632308** | **0** | **-1248.101619** | **-3.949974** | **-1251.868353** |
| conformer_3 | -1253.824518 | 0.282653 | 0.218674 | -1253.851863 | -71.8 | -1253.569210 | -1253.630177 | 5.6 | -1248.112526 | -3.948977 | -1251.867162 |
| conformer_47 | -1253.815634 | 0.282383 | 0.217981 | -1253.851750 | -94.8 | -1253.569367 | -1253.630757 | 4.1 | -1248.100830 | -3.949970 | -1251.865923 |
| conformer_14 | -1253.826346 | 0.282354 | 0.219463 | -1253.851694 | -66.6 | -1253.569340 | -1253.629219 | 8.1 | -1248.110807 | -3.951592 | -1251.865273 |
| conformer_60 | -1253.814734 | 0.282376 | 0.217234 | -1253.851627 | -96.9 | -1253.569251 | -1253.631381 | 2.4 | -1248.100702 | -3.949778 | -1251.867127 |
| conformer_9 | -1253.823654 | 0.282238 | 0.219220 | -1253.851600 | -73.4 | -1253.569362 | -1253.629368 | 7.7 | -1248.106431 | -3.953458 | -1251.865602 |
| conformer_59 | -1253.816703 | 0.282343 | 0.217637 | -1253.851595 | -91.6 | -1253.569252 | -1253.630946 | 3.6 | -1248.101555 | -3.951285 | -1251.867083 |
| conformer_45 | -1253.819306 | 0.282123 | 0.218505 | -1253.851467 | -84.4 | -1253.569344 | -1253.629950 | 6.2 | -1248.102207 | -3.953195 | -1251.866046 |
| conformer_12 | -1253.821900 | 0.282142 | 0.217645 | -1253.851444 | -77.6 | -1253.569302 | -1253.630787 | 4.0 | -1248.106933 | -3.950983 | -1251.866803 |
| conformer_13 | -1253.821900 | 0.282142 | 0.217645 | -1253.851444 | -77.6 | -1253.569302 | -1253.630787 | 4.0 | -1248.106933 | -3.950983 | -1251.866803 |
| conformer_102 | -1253.815714 | 0.282254 | 0.217568 | -1253.851364 | -93.6 | -1253.569110 | -1253.630784 | 4.0 | -1248.102768 | -3.948620 | -1251.866458 |
| conformer_26 | -1253.821104 | 0.281998 | 0.217338 | -1253.851353 | -79.4 | -1253.569355 | -1253.631003 | 3.4 | -1248.105648 | -3.951398 | -1251.866946 |
| conformer_34 | -1253.821083 | 0.282145 | 0.218773 | -1253.851353 | -79.5 | -1253.569208 | -1253.629568 | 7.2 | -1248.103723 | -3.953044 | -1251.865252 |
| conformer_43 | -1253.816999 | 0.282440 | 0.218146 | -1253.851157 | -89.7 | -1253.568717 | -1253.629999 | 6.1 | -1248.103204 | -3.949484 | -1251.865689 |
| conformer_76 | -1253.815311 | 0.282332 | 0.216992 | -1253.851050 | -93.8 | -1253.568718 | -1253.631046 | 3.3 | -1248.103245 | -3.948345 | -1251.867325 |
| conformer_21 | -1253.822208 | 0.282573 | 0.218545 | -1253.850977 | -75.5 | -1253.568404 | -1253.629420 | 7.6 | -1248.108900 | -3.950103 | -1251.866215 |
| conformer_16 | -1253.821900 | 0.282416 | 0.217912 | -1253.850927 | -76.2 | -1253.568511 | -1253.630003 | 6.1 | -1248.110331 | -3.948858 | -1251.867292 |
| conformer_112 | -1253.821900 | 0.282416 | 0.217912 | -1253.850927 | -76.2 | -1253.568511 | -1253.630003 | 6.1 | -1248.110331 | -3.948858 | -1251.867292 |
| conformer_71 | -1253.819025 | 0.282011 | 0.217365 | -1253.850912 | -83.7 | -1253.568901 | -1253.630535 | 4.7 | -1248.101967 | -3.953162 | -1251.866639 |
| conformer_116 | -1253.819025 | 0.282011 | 0.217365 | -1253.850912 | -83.7 | -1253.568901 | -1253.630535 | 4.7 | -1248.101967 | -3.953162 | -1251.866639 |
| conformer_61 | -1253.822341 | 0.282104 | 0.218825 | -1253.850843 | -74.8 | -1253.568739 | -1253.629006 | 8.7 | -1248.105169 | -3.953961 | -1251.865795 |

| | | | | | | | | | | | |
|---|---|---|---|---|---|---|---|---|---|---|---|
| conformer_110 | -1253.811797 | 0.282274 | 0.216593 | -1253.850796 | -102.4 | -1253.568522 | -1253.631191 | 2.9 | -1248.097265 | -3.950683 | -1251.867341 |
| conformer_11 | -1253.820372 | 0.282484 | 0.217906 | -1253.850774 | -79.8 | -1253.568290 | -1253.629856 | 6.4 | -1248.107537 | -3.949891 | -1251.866912 |
| conformer_83 | -1253.817277 | 0.282288 | 0.217493 | -1253.850682 | -87.7 | -1253.568394 | -1253.630177 | 5.6 | -1248.105449 | -3.948034 | -1251.866383 |
| conformer_55 | -1253.812871 | 0.282260 | 0.216614 | -1253.850681 | -99.3 | -1253.568421 | -1253.631055 | 3.3 | -1248.100031 | -3.948637 | -1251.866853 |
| conformer_94 | -1253.811902 | 0.282163 | 0.216230 | -1253.850524 | -101.4 | -1253.568361 | -1253.631282 | 2.7 | -1248.098927 | -3.948685 | -1251.866992 |
| conformer_29 | -1253.822215 | 0.282250 | 0.218956 | -1253.850436 | -74.1 | -1253.568186 | -1253.628468 | 10.1 | -1248.105896 | -3.952539 | -1251.864688 |
| conformer_81 | -1253.818147 | 0.282292 | 0.218059 | -1253.850419 | -84.7 | -1253.568127 | -1253.629348 | 7.8 | -1248.104470 | -3.950357 | -1251.866028 |
| conformer_96 | -1253.812911 | 0.282272 | 0.217166 | -1253.850396 | -98.4 | -1253.568124 | -1253.630218 | 5.5 | -1248.096813 | -3.952420 | -1251.866540 |
| conformer_104 | -1253.811023 | 0.282293 | 0.216396 | -1253.850369 | -103.3 | -1253.568076 | -1253.630961 | 3.5 | -1248.096322 | -3.950631 | -1251.866891 |
| conformer_65 | -1253.820984 | 0.282061 | 0.217634 | -1253.850320 | -77.0 | -1253.568259 | -1253.629674 | 6.9 | -1248.106408 | -3.950872 | -1251.865970 |
| conformer_40 | -1253.821637 | 0.282366 | 0.217887 | -1253.850273 | -75.2 | -1253.567907 | -1253.629374 | 7.7 | -1248.110449 | -3.948694 | -1251.866881 |
| conformer_27 | -1253.822930 | 0.282125 | 0.219026 | -1253.850183 | -71.6 | -1253.568058 | -1253.628145 | 10.9 | -1248.102519 | -3.956382 | -1251.864116 |
| conformer_32 | -1253.817715 | 0.282027 | 0.217030 | -1253.850108 | -85.0 | -1253.568081 | -1253.630066 | 5.9 | -1248.102130 | -3.951810 | -1251.866292 |
| conformer_28 | -1253.819007 | 0.282125 | 0.218717 | -1253.850062 | -81.5 | -1253.567937 | -1253.628333 | 10.4 | -1248.100957 | -3.954322 | -1251.864604 |
| conformer_1 | -1253.824421 | 0.282645 | 0.218760 | -1253.849870 | -66.8 | -1253.567225 | -1253.628098 | 11.1 | -1248.111623 | -3.949608 | -1251.864907 |
| conformer_75 | -1253.814192 | 0.282152 | 0.215879 | -1253.849816 | -93.5 | -1253.567664 | -1253.630925 | 3.6 | -1248.103565 | -3.947087 | -1251.867386 |
| conformer_33 | -1253.817593 | 0.282260 | 0.217285 | -1253.849804 | -84.6 | -1253.567544 | -1253.629507 | 7.4 | -1248.105139 | -3.949780 | -1251.866833 |
| conformer_117 | -1253.811661 | 0.282234 | 0.216211 | -1253.849753 | -100.0 | -1253.567519 | -1253.630530 | 4.7 | -1248.098868 | -3.949183 | -1251.866920 |
| conformer_79 | -1253.813261 | 0.282340 | 0.217810 | -1253.849724 | -95.7 | -1253.567384 | -1253.628902 | 8.9 | -1248.098670 | -3.950391 | -1251.864702 |
| conformer_44 | -1253.817740 | 0.282606 | 0.219033 | -1253.849554 | -83.5 | -1253.566948 | -1253.627509 | 12.6 | -1248.101555 | -3.952528 | -1251.863852 |
| conformer_115 | -1253.812553 | 0.282182 | 0.216301 | -1253.849553 | -97.1 | -1253.567371 | -1253.630240 | 5.4 | -1248.100302 | -3.948333 | -1251.866323 |
| conformer_113 | -1253.817916 | 0.281995 | 0.218310 | -1253.849406 | -82.7 | -1253.567411 | -1253.628084 | 11.1 | -1248.099786 | -3.955040 | -1251.864994 |
| conformer_98 | -1253.809115 | 0.282157 | 0.215797 | -1253.849353 | -105.6 | -1253.567196 | -1253.630544 | 4.6 | -1248.095559 | -3.949483 | -1251.866472 |
| conformer_91 | -1253.818395 | 0.282262 | 0.217958 | -1253.849317 | -81.2 | -1253.567055 | -1253.628347 | 10.4 | -1248.105497 | -3.950038 | -1251.865486 |
| conformer_111 | -1253.817451 | 0.282226 | 0.217331 | -1253.849198 | -83.4 | -1253.566972 | -1253.628855 | 9.1 | -1248.105397 | -3.949602 | -1251.866404 |
| conformer_122 | -1253.814578 | 0.282176 | 0.217161 | -1253.849167 | -90.8 | -1253.566991 | -1253.628994 | 8.7 | -1248.099984 | -3.951349 | -1251.865750 |
| conformer_114 | -1253.816889 | 0.281938 | 0.216976 | -1253.849026 | -84.4 | -1253.567088 | -1253.629038 | 8.6 | -1248.101603 | -3.951812 | -1251.865565 |
| conformer_99 | -1253.814497 | 0.282182 | 0.216108 | -1253.848883 | -90.3 | -1253.566701 | -1253.629763 | 6.7 | -1248.104466 | -3.946674 | -1251.866405 |
| conformer_4 | -1253.820236 | 0.282522 | 0.218106 | -1253.848754 | -74.9 | -1253.566232 | -1253.627636 | 12.3 | -1248.106530 | -3.950522 | -1251.864452 |
| conformer_52 | -1253.818953 | 0.282029 | 0.218782 | -1253.848697 | -78.1 | -1253.566668 | -1253.626903 | 14.2 | -1248.097717 | -3.957394 | -1251.863061 |
| conformer_64 | -1253.818218 | 0.282231 | 0.216738 | -1253.848462 | -79.4 | -1253.566231 | -1253.628712 | 9.4 | -1248.107290 | -3.947978 | -1251.865762 |
| conformer_118 | -1253.810513 | 0.282005 | 0.215103 | -1253.848459 | -99.6 | -1253.566454 | -1253.630344 | 5.2 | -1248.099101 | -3.948037 | -1251.866969 |
| conformer_30 | -1253.819044 | 0.282506 | 0.218958 | -1253.848268 | -76.7 | -1253.565762 | -1253.626298 | 15.8 | -1248.102735 | -3.953092 | -1251.863081 |
| conformer_85 | -1253.813848 | 0.282473 | 0.218686 | -1253.848235 | -90.3 | -1253.565762 | -1253.626537 | 15.2 | -1248.097001 | -3.953395 | -1251.863085 |
| conformer_48 | -1253.815211 | 0.282682 | 0.218304 | -1253.847624 | -85.1 | -1253.564942 | -1253.626308 | 15.8 | -1248.101242 | -3.950750 | -1251.863088 |
| conformer_105 | -1253.814201 | 0.282100 | 0.216173 | -1253.847481 | -87.4 | -1253.565381 | -1253.628296 | 10.5 | -1248.102282 | -3.949110 | -1251.865486 |
| conformer_62 | -1253.815003 | 0.282314 | 0.218490 | -1253.846989 | -84.0 | -1253.564675 | -1253.625487 | 17.9 | -1248.097961 | -3.954015 | -1251.862460 |
| conformer_42 | -1253.815339 | 0.282494 | 0.218093 | -1253.846418 | -81.6 | -1253.563924 | -1253.625313 | 18.4 | -1248.101035 | -3.951530 | -1251.862539 |
| conformer_93 | -1253.811283 | 0.282591 | 0.218110 | -1253.846280 | -91.9 | -1253.563689 | -1253.625158 | 18.8 | -1248.096612 | -3.951553 | -1251.862041 |
| conformer_54 | -1253.813704 | 0.282651 | 0.218254 | -1253.845817 | -84.3 | -1253.563166 | -1253.624551 | 20.4 | -1248.099263 | -3.950988 | -1251.861098 |
| conformer_82 | -1253.811282 | 0.282411 | 0.217978 | -1253.845062 | -88.7 | -1253.562651 | -1253.624072 | 21.6 | -1248.096233 | -3.952379 | -1251.861402 |
| conformer_100 | -1253.809800 | 0.282593 | 0.218270 | -1253.844495 | -91.1 | -1253.561902 | -1253.623213 | 23.9 | -1248.094630 | -3.951838 | -1251.859882 |
| conformer_73 | -1253.812542 | 0.282502 | 0.218069 | -1253.844119 | -82.9 | -1253.561617 | -1253.623038 | 24.3 | -1248.097646 | -3.951690 | -1251.859831 |
| conformer_50 | -1253.814769 | 0.282478 | 0.218015 | -1253.843477 | -75.4 | -1253.560999 | -1253.622450 | 25.9 | -1248.101055 | -3.950886 | -1251.859623 |
| conformer_119 | -1253.808503 | 0.282477 | 0.218188 | -1253.842907 | -90.3 | -1253.560430 | -1253.621707 | 27.8 | -1248.092869 | -3.952549 | -1251.858621 |
| conformer_37 | -1253.815635 | 0.282513 | 0.218157 | -1253.842452 | -70.4 | -1253.559939 | -1253.621283 | 28.9 | -1248.102587 | -3.950591 | -1251.858826 |
| conformer_95 | -1253.810816 | 0.282405 | 0.217938 | -1253.842245 | -82.5 | -1253.559840 | -1253.621295 | 28.9 | -1248.096394 | -3.951770 | -1251.858643 |
| conformer_101 | -1253.811610 | 0.282420 | 0.218037 | -1253.841191 | -77.7 | -1253.558771 | -1253.620142 | 31.9 | -1248.097838 | -3.951513 | -1251.857882 |

[a]: Single-point calculation in aqueous phase with SMD model.

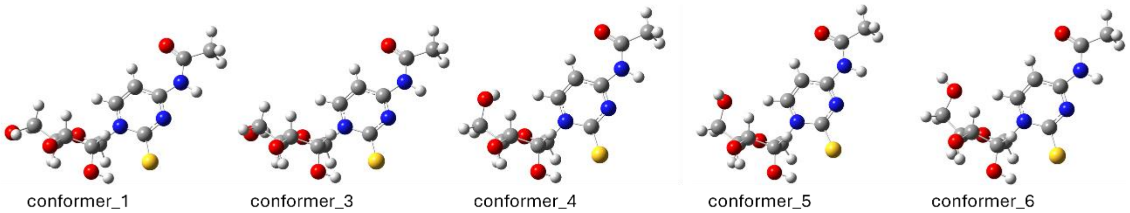

**Figure S24.** B3LYP-D3/def2-TZVPP optimized geometries of conformers for 2-thio-4-acetylcytidine (**1rb2th4acC**).

**Table S22.** Conformers of gas-phase optimized 2-thio-4-acetylcytidine (**1rb2th4acC**) at the B3LYP-D3/def2-TZVPP level of theory followed by aqueous phase single-point calculation. The columns display total energy without zero-point correction ($E_{Tot}$), thermal correction to enthalpy (δH), Gibbs free energy (δG), total energy without zero-point correction ($E_{Tot,W}$), Gibbs free energy ($G_{298,W}$) in water (W), total single-point energy ($E_{CBS}$) calculated at DLPNO-CCSD(T)/CBS level of theory, and their corresponding free energy $G_{CBS,W}$. $G_{298,W}$ and $G_{CBS,W}$ have been corrected to the standard state of 1 mol/L by addition of +7.908 kJ/mol. $\Delta G_{Solv}$ represents the Gibbs free energy of solvation. The data are arranged in the ascending numeric order of $E_{Tot,W}$. $\Delta G_{298,W}$ represents the respective energy difference to the lowest structure. Only conformers within the 24 kJ/mol (6 kcal/mol) energy window above the lowest in CREST are included in initial conformer sampling. Duplicates of the same structure are excluded. The overall optimum is marked bold.

| | B3LYP-D3/def2-TZVPP | | | SMD($H_2O$)/B3LYP-D3/def2-TZVPP[a] | | | | | DLPNO-CCSD(T)/CBS | | |
|---|---|---|---|---|---|---|---|---|---|---|---|
| 1rb2th4acC No. | $E_{Tot}$ (Hartree) | δH (Hartree) | δG (Hartree) | $E_{Tot,W}$ (Hartree) | $\Delta G_{Solv}$ (kJ/mol) | $H_{298,W}$ (Hartree) | $G_{298,W}$ (Hartree) | $\Delta G_{298,W}$ (kJ/mol) | $E_{CBS,HF}$ (Hartree) | $E_{CBS,C}$ (Hartree) | $G_{CBS,W}$ (Hartree) |
| conformer_10 | -1367.233762 | 0.293511 | 0.224834 | -1367.270901 | -97.5 | -1366.977390 | -1367.043055 | 3.7 | -1360.926460 | -4.377346 | -1365.113098 |
| conformer_5 | -1367.233622 | 0.294078 | 0.224399 | -1367.270859 | -97.8 | -1366.976781 | -1367.043448 | 2.6 | -1360.931148 | -4.373674 | -1365.114649 |
| **conformer_4** | **-1367.232625** | **0.293980** | **0.223236** | **-1367.270699** | **-100.0** | **-1366.976719** | **-1367.044451** | **0.0** | **-1360.929917** | **-4.373991** | **-1365.115733** |
| conformer_15 | -1367.231425 | 0.293350 | 0.223641 | -1367.270644 | -103.0 | -1366.977294 | -1367.043991 | 1.2 | -1360.924197 | -4.377620 | -1365.114383 |
| conformer_7 | -1367.232286 | 0.293536 | 0.224645 | -1367.270224 | -99.6 | -1366.976688 | -1367.042567 | 4.9 | -1360.925135 | -4.378085 | -1365.113501 |
| conformer_8 | -1367.234720 | 0.293576 | 0.224874 | -1367.269906 | -92.4 | -1366.976330 | -1367.042020 | 6.4 | -1360.928483 | -4.376791 | -1365.112575 |
| conformer_11 | -1367.230066 | 0.293404 | 0.223628 | -1367.269884 | -104.5 | -1366.976480 | -1367.043244 | 3.2 | -1360.924431 | -4.376200 | -1365.113809 |
| conformer_6 | -1367.234106 | 0.294035 | 0.224218 | -1367.269878 | -93.9 | -1366.975843 | -1367.042648 | 4.7 | -1360.932380 | -4.373323 | -1365.114245 |
| conformer_16 | -1367.229353 | 0.293272 | 0.223450 | -1367.269841 | -106.3 | -1366.976569 | -1367.043379 | 2.8 | -1360.923178 | -4.376614 | -1365.113818 |
| conformer_19 | -1367.231238 | 0.293527 | 0.224881 | -1367.269741 | -101.1 | -1366.976214 | -1367.041848 | 6.8 | -1360.924438 | -4.378229 | -1365.113277 |
| conformer_3 | -1367.232360 | 0.293951 | 0.224020 | -1367.269448 | -97.4 | -1366.975497 | -1367.042416 | 5.3 | -1360.930581 | -4.373658 | -1365.114295 |
| conformer_21 | -1367.229237 | 0.293317 | 0.223311 | -1367.268764 | -103.8 | -1366.975447 | -1367.042441 | 5.3 | -1360.924063 | -4.376044 | -1365.113311 |
| conformer_14 | -1367.229902 | 0.293749 | 0.223486 | -1367.268742 | -102.0 | -1366.974993 | -1367.042244 | 5.8 | -1360.928567 | -4.373474 | -1365.114384 |
| conformer_18 | -1367.229674 | 0.293706 | 0.223473 | -1367.268122 | -100.9 | -1366.974416 | -1367.041637 | 7.4 | -1360.928704 | -4.373304 | -1365.113972 |
| conformer_1 | -1367.232279 | 0.293929 | 0.224239 | -1367.267503 | -92.5 | -1366.973574 | -1367.040252 | 11.0 | -1360.929793 | -4.374221 | -1365.111988 |
| conformer_43 | -1367.219868 | 0.293708 | 0.221889 | -1367.267408 | -124.8 | -1366.973700 | -1367.042507 | 5.1 | -1360.916799 | -4.373512 | -1365.112949 |
| conformer_32 | -1367.220183 | 0.293632 | 0.222495 | -1367.267347 | -123.8 | -1366.973715 | -1367.041840 | 6.9 | -1360.916103 | -4.373770 | -1365.111529 |
| conformer_41 | -1367.219037 | 0.293609 | 0.221573 | -1367.266986 | -125.9 | -1366.973377 | -1367.042401 | 5.4 | -1360.915943 | -4.373284 | -1365.112591 |
| conformer_26 | -1367.221682 | 0.293703 | 0.222807 | -1367.266803 | -118.5 | -1366.973100 | -1367.040984 | 9.1 | -1360.918558 | -4.373339 | -1365.111199 |
| conformer_30 | -1367.223641 | 0.293966 | 0.224697 | -1367.266514 | -112.6 | -1366.972548 | -1367.038805 | 14.8 | -1360.917861 | -4.376347 | -1365.109371 |
| conformer_38 | -1367.226172 | 0.294126 | 0.224522 | -1367.266142 | -104.9 | -1366.972016 | -1367.038608 | 15.3 | -1360.921023 | -4.375767 | -1365.109227 |
| conformer_36 | -1367.217392 | 0.293550 | 0.221575 | -1367.266026 | -127.7 | -1366.972476 | -1367.041439 | 7.9 | -1360.915195 | -4.372448 | -1365.111690 |
| conformer_23 | -1367.225971 | 0.293523 | 0.222107 | -1367.265792 | -104.6 | -1366.972269 | -1367.040673 | 9.9 | -1360.925446 | -4.372635 | -1365.112784 |
| conformer_44 | -1367.224667 | 0.294101 | 0.223981 | -1367.265783 | -108.0 | -1366.971682 | -1367.038790 | 14.9 | -1360.918963 | -4.376280 | -1365.109367 |
| conformer_40 | -1367.224933 | 0.293677 | 0.225374 | -1367.265528 | -106.6 | -1366.971851 | -1367.037142 | 19.2 | -1360.915125 | -4.380299 | -1365.107634 |
| conformer_20 | -1367.225131 | 0.293841 | 0.224530 | -1367.265263 | -105.4 | -1366.971422 | -1367.037721 | 17.7 | -1360.919341 | -4.376800 | -1365.108731 |
| conformer_34 | -1367.220844 | 0.294046 | 0.223868 | -1367.263801 | -112.8 | -1366.969755 | -1367.036921 | 19.8 | -1360.917400 | -4.374519 | -1365.107996 |
| conformer_28 | -1367.221253 | 0.293871 | 0.223621 | -1367.262732 | -108.9 | -1366.968861 | -1367.036099 | 21.9 | -1360.917583 | -4.375124 | -1365.107553 |

[a]: Single-point calculation in aqueous phase with SMD model.

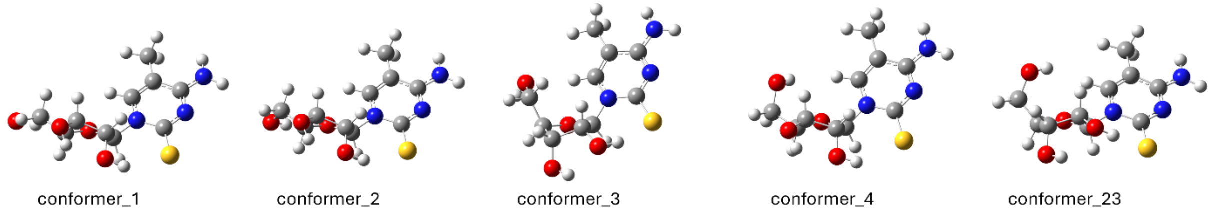

**Figure S25.** B3LYP-D3/def2-TZVPP optimized geometries of conformers for 2-thio-5-methylcytidine (**1rb2th5mC**).

**Table S23.** Conformers of gas-phase optimized 2-thio-5-methylcytidine (**1rb2th5mC**) at the B3LYP-D3/def2-TZVPP level of theory followed by aqueous phase single-point calculation. The columns display total energy without zero-point correction ($E_{Tot}$), thermal correction to enthalpy (δH), Gibbs free energy (δG), total energy without zero-point correction ($E_{Tot,W}$), Gibbs free energy ($G_{298,W}$) in water (W), total single-point energy ($E_{CBS}$) calculated at DLPNO-CCSD(T)/CBS level of theory, and their corresponding free energy $G_{CBS,W}$. $G_{298,W}$ and $G_{CBS,W}$ have been corrected to the standard state of 1 mol/L by addition of +7.908 kJ/mol. $\Delta G_{Solv}$ represents the Gibbs free energy of solvation. The data are arranged in the ascending numeric order of $E_{Tot,W}$. $\Delta G_{298,W}$ represents the respective energy difference to the lowest structure. Only conformers within the 24 kJ/mol (6 kcal/mol) energy window above the lowest in CREST are included in initial conformer sampling. Duplicates of the same structure are excluded. The overall optimum is marked bold.

| 1rb2th5mC No. | B3LYP-D3/def2-TZVPP $E_{Tot}$ (Hartree) | δH (Hartree) | δG (Hartree) | SMD(H2O)/B3LYP-D3/def2-TZVPP[a] $E_{Tot,W}$ (Hartree) | $\Delta G_{Solv}$ (kJ/mol) | $H_{298,W}$ (Hartree) | $G_{298,W}$ (Hartree) | $\Delta G_{298,W}$ (kJ/mol) | DLPNO-CCSD(T)/CBS $E_{CBS,HF}$ (Hartree) | $E_{CBS,C}$ (Hartree) | $G_{CBS,W}$ (Hartree) |
|---|---|---|---|---|---|---|---|---|---|---|---|
| conformer_7 | -1253.847325 | 0.281596 | 0.217315 | -1253.885714 | -100.8 | -1253.604118 | -1253.665387 | 1.3 | -1248.132147 | -3.947727 | -1251.897936 |
| **conformer_23** | **-1253.844576** | **0.281477** | **0.216475** | **-1253.885356** | **-107.1** | **-1253.603879** | **-1253.665869** | **0** | **-1248.129485** | **-3.947838** | **-1251.898615** |
| conformer_3 | -1253.846379 | 0.281824 | 0.218143 | -1253.885086 | -101.6 | -1253.603262 | -1253.663931 | 5.1 | -1248.130948 | -3.948812 | -1251.897312 |
| conformer_4 | -1253.845920 | 0.282221 | 0.216711 | -1253.885051 | -102.7 | -1253.602830 | -1253.665328 | 1.4 | -1248.135179 | -3.944613 | -1251.899200 |
| conformer_15 | -1253.845603 | 0.281718 | 0.217481 | -1253.885043 | -103.5 | -1253.603325 | -1253.664550 | 3.5 | -1248.129635 | -3.948944 | -1251.897525 |
| conformer_5 | -1253.846940 | 0.282277 | 0.217983 | -1253.884995 | -99.9 | -1253.602718 | -1253.664000 | 4.9 | -1248.136531 | -3.944193 | -1251.897783 |
| conformer_9 | -1253.843419 | 0.281458 | 0.216111 | -1253.884486 | -107.8 | -1253.603028 | -1253.665363 | 1.3 | -1248.129966 | -3.946482 | -1251.898393 |
| conformer_19 | -1253.847447 | 0.281637 | 0.217356 | -1253.884482 | -97.2 | -1253.602845 | -1253.664114 | 4.6 | -1248.133474 | -3.947103 | -1251.897245 |
| conformer_18 | -1253.842909 | 0.281367 | 0.216200 | -1253.884400 | -108.9 | -1253.603033 | -1253.665188 | 1.8 | -1248.128895 | -3.946942 | -1251.898117 |
| conformer_25 | -1253.844533 | 0.281720 | 0.218230 | -1253.884380 | -104.6 | -1253.602660 | -1253.663138 | 7.2 | -1248.129497 | -3.948958 | -1251.897060 |
| conformer_10 | -1253.846946 | 0.282221 | 0.217624 | -1253.883833 | -96.8 | -1253.601612 | -1253.663197 | 7.0 | -1248.137426 | -3.943748 | -1251.897426 |
| conformer_2 | -1253.845666 | 0.281983 | 0.216319 | -1253.883600 | -99.6 | -1253.601617 | -1253.664269 | 4.2 | -1248.136011 | -3.944102 | -1251.898717 |
| conformer_27 | -1253.842270 | 0.281345 | 0.215819 | -1253.883322 | -107.8 | -1253.601977 | -1253.664491 | 3.6 | -1248.129275 | -3.946382 | -1251.897877 |
| conformer_14 | -1253.843323 | 0.281843 | 0.216436 | -1253.882908 | -103.9 | -1253.601065 | -1253.663460 | 6.3 | -1248.134054 | -3.944030 | -1251.898222 |
| conformer_39 | -1253.833072 | 0.281940 | 0.216356 | -1253.882278 | -129.2 | -1253.600338 | -1253.662910 | 7.8 | -1248.121283 | -3.943909 | -1251.895031 |
| conformer_24 | -1253.842745 | 0.281810 | 0.216572 | -1253.882245 | -103.7 | -1253.600435 | -1253.662661 | 8.4 | -1248.133816 | -3.943926 | -1251.897658 |
| conformer_1 | -1253.845755 | 0.282143 | 0.217171 | -1253.881757 | -94.5 | -1253.599614 | -1253.661574 | 11.3 | -1248.135370 | -3.944666 | -1251.895854 |
| conformer_35 | -1253.836812 | 0.282028 | 0.217516 | -1253.881288 | -116.8 | -1253.599260 | -1253.660760 | 13.4 | -1248.123142 | -3.946766 | -1251.893856 |
| conformer_30 | -1253.838968 | 0.281700 | 0.214893 | -1253.880048 | -107.9 | -1253.598348 | -1253.662143 | 9.8 | -1248.130513 | -3.943115 | -1251.896802 |
| conformer_26 | -1253.838440 | 0.282051 | 0.217575 | -1253.879961 | -109.0 | -1253.597910 | -1253.659374 | 17.1 | -1248.124805 | -3.947113 | -1251.892852 |
| conformer_38 | -1253.833615 | 0.282071 | 0.216187 | -1253.877915 | -116.3 | -1253.595844 | -1253.658716 | 18.8 | -1248.122433 | -3.944852 | -1251.892385 |
| conformer_33 | -1253.834110 | 0.281953 | 0.215829 | -1253.876881 | -112.3 | -1253.594928 | -1253.658040 | 20.6 | -1248.122784 | -3.945436 | -1251.892149 |
| conformer_34 | -1253.833907 | 0.281975 | 0.215700 | -1253.872864 | -102.3 | -1253.590889 | -1253.654152 | 30.8 | -1248.123784 | -3.944391 | -1251.888421 |

[a]: Single-point calculation in aqueous phase with SMD model.

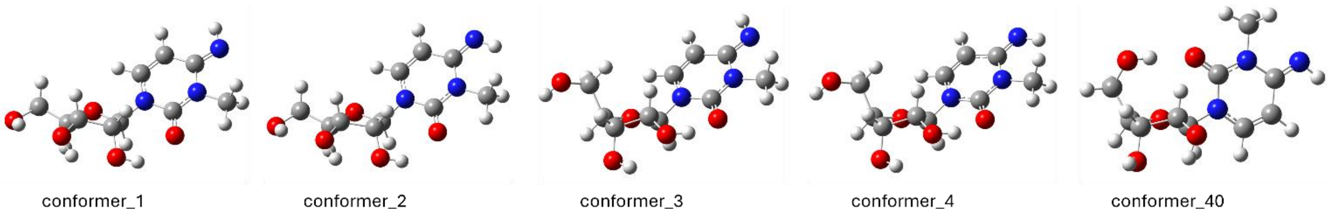

**Figure S26.** B3LYP-D3/def2-TZVPP optimized geometries of conformers for neutral 3-methylcytidine (**1rb3mC**).

**Table S24.** Conformers of gas-phase optimized 3-methylcytidine (**1rb3mC**) at the B3LYP-D3/def2-TZVPP level of theory followed by aqueous phase single-point calculation. The columns display total energy without zero-point correction ($E_{Tot}$), thermal correction to enthalpy (δH), Gibbs free energy (δG), total energy without zero-point correction ($E_{Tot,W}$), Gibbs free energy ($G_{298,W}$) in water (W), total single-point energy ($E_{CBS}$) calculated at DLPNO-CCSD(T)/CBS level of theory, and their corresponding free energy $G_{CBS,W}$. $G_{298,W}$ and $G_{CBS,W}$ have been corrected to the standard state of 1 mol/L by addition of +7.908 kJ/mol. $\Delta G_{Solv}$ represents the Gibbs free energy of solvation. The data are arranged in the ascending numeric order of $E_{Tot,W}$. $\Delta G_{298,W}$ represents the respective energy difference to the lowest structure. Only conformers within the 24 kJ/mol (6 kcal/mol) energy window above the lowest in CREST are included in initial conformer sampling. Duplicates of the same structure are excluded. The overall optimum is marked bold.

| | B3LYP-D3/def2-TZVPP | | | SMD(H2O)/B3LYP-D3/def2-TZVPP[a] | | | | | DLPNO-CCSD(T)/CBS | | |
|---|---|---|---|---|---|---|---|---|---|---|---|
| 1rb3mC No. | $E_{Tot}$ (Hartree) | δH (Hartree) | δG (Hartree) | $E_{Tot,W}$ (Hartree) | $\Delta G_{Solv}$ (kJ/mol) | $H_{298,W}$ (Hartree) | $G_{298,W}$ (Hartree) | $\Delta G_{298,W}$ (kJ/mol) | $E_{CBS,HF}$ (Hartree) | $E_{CBS,C}$ (Hartree) | $G_{CBS,W}$ (Hartree) |
| **conformer_40** | **-930.881573** | **0.284888** | **0.222273** | **-930.913892** | **-84.9** | **-930.629004** | **-930.688607** | **0** | **-925.486069** | **-3.993637** | **-929.286740** |
| conformer_58 | -930.882733 | 0.284854 | 0.223028 | -930.913458 | -80.7 | -930.628604 | -930.687418 | 3.1 | -925.487858 | -3.993498 | -929.286042 |
| conformer_25 | -930.879962 | 0.284787 | 0.222822 | -930.913423 | -87.9 | -930.628636 | -930.687589 | 2.7 | -925.483129 | -3.994674 | -929.285430 |
| conformer_14 | -930.881893 | 0.284773 | 0.222710 | -930.913315 | -82.5 | -930.628542 | -930.687593 | 2.7 | -925.486934 | -3.993335 | -929.285969 |
| conformer_63 | -930.882375 | 0.284783 | 0.222606 | -930.913296 | -81.2 | -930.628513 | -930.687678 | 2.4 | -925.487419 | -3.993450 | -929.286172 |
| conformer_34 | -930.879642 | 0.284713 | 0.222445 | -930.913250 | -88.2 | -930.628537 | -930.687793 | 2.1 | -925.482721 | -3.994643 | -929.285515 |
| conformer_20 | -930.881572 | 0.284707 | 0.222386 | -930.913148 | -82.9 | -930.628441 | -930.687750 | 2.2 | -925.486526 | -3.993303 | -929.286007 |
| conformer_29 | -930.877010 | 0.284666 | 0.222789 | -930.912780 | -93.9 | -930.628114 | -930.686979 | 4.3 | -925.479425 | -3.995446 | -929.284840 |
| conformer_59 | -930.879643 | 0.284728 | 0.222959 | -930.912754 | -86.9 | -930.628026 | -930.686783 | 4.8 | -925.483966 | -3.994365 | -929.285471 |
| conformer_68 | -930.879737 | 0.284895 | 0.222643 | -930.912670 | -86.5 | -930.627775 | -930.687015 | 4.2 | -925.484688 | -3.993140 | -929.285106 |
| conformer_19 | -930.878905 | 0.284654 | 0.222681 | -930.912635 | -88.6 | -930.627981 | -930.686942 | 4.4 | -925.483175 | -3.994151 | -929.285363 |
| conformer_175 | -930.879160 | 0.284825 | 0.221847 | -930.912282 | -87.0 | -930.627457 | -930.687423 | 3.1 | -925.484038 | -3.993055 | -929.285356 |
| conformer_96 | -930.876771 | 0.284760 | 0.222549 | -930.912009 | -92.5 | -930.627249 | -930.686448 | 5.7 | -925.480994 | -3.993922 | -929.284593 |
| conformer_26 | -930.881653 | 0.284714 | 0.222303 | -930.911775 | -79.1 | -930.627061 | -930.686460 | 5.6 | -925.486619 | -3.993569 | -929.284995 |
| conformer_170 | -930.873809 | 0.284590 | 0.220239 | -930.911710 | -99.5 | -930.627120 | -930.688459 | 0.4 | -925.480208 | -3.992148 | -929.287006 |
| conformer_173 | -930.880329 | 0.284832 | 0.222066 | -930.911403 | -81.6 | -930.626571 | -930.686325 | 6.0 | -925.487842 | -3.990865 | -929.284702 |
| conformer_48 | -930.879352 | 0.284847 | 0.220375 | -930.911197 | -83.6 | -930.626350 | -930.687810 | 2.1 | -925.486776 | -3.990994 | -929.286228 |
| conformer_133 | -930.878538 | 0.284572 | 0.222269 | -930.911082 | -85.4 | -930.626510 | -930.685801 | 7.4 | -925.482731 | -3.994400 | -929.284394 |
| conformer_9 | -930.882961 | 0.284608 | 0.221920 | -930.911063 | -73.8 | -930.626455 | -930.686131 | 6.5 | -925.486265 | -3.995143 | -929.284578 |
| conformer_145 | -930.871133 | 0.284509 | 0.220396 | -930.911062 | -104.8 | -930.626553 | -930.687654 | 2.5 | -925.476822 | -3.992853 | -929.286195 |
| conformer_140 | -930.873334 | 0.284566 | 0.221351 | -930.910958 | -98.8 | -930.626392 | -930.686595 | 5.3 | -925.478755 | -3.992223 | -929.284239 |
| conformer_185 | -930.876403 | 0.284674 | 0.221099 | -930.910826 | -90.4 | -930.626152 | -930.686715 | 5.0 | -925.482001 | -3.992814 | -929.285127 |
| conformer_24 | -930.881354 | 0.284719 | 0.221892 | -930.910784 | -77.3 | -930.626065 | -930.685880 | 7.2 | -925.486095 | -3.993670 | -929.284291 |
| conformer_49 | -930.876885 | 0.284721 | 0.222135 | -930.910745 | -88.9 | -930.626024 | -930.685598 | 7.9 | -925.483440 | -3.991795 | -929.283949 |
| conformer_30 | -930.882390 | 0.284642 | 0.221992 | -930.910739 | -74.4 | -930.626097 | -930.685735 | 7.5 | -925.486945 | -3.993573 | -929.283864 |
| conformer_5 | -930.881671 | 0.284739 | 0.222627 | -930.910692 | -76.2 | -930.625953 | -930.685053 | 9.3 | -925.486731 | -3.993759 | -929.283872 |
| conformer_17 | -930.882428 | 0.284551 | 0.221261 | -930.910675 | -74.2 | -930.626124 | -930.686402 | 5.8 | -925.485808 | -3.994964 | -929.284746 |
| conformer_31 | -930.882428 | 0.284551 | 0.221261 | -930.910675 | -74.2 | -930.626124 | -930.686402 | 5.8 | -925.485808 | -3.994964 | -929.284746 |
| conformer_196 | -930.876904 | 0.284843 | 0.221069 | -930.910599 | -88.5 | -930.625756 | -930.686518 | 5.5 | -925.484852 | -3.990953 | -929.285419 |
| conformer_103 | -930.876150 | 0.284746 | 0.220708 | -930.910532 | -90.3 | -930.625786 | -930.686812 | 4.7 | -925.482696 | -3.991839 | -929.285197 |

| | | | | | | | | | | | |
|---|---|---|---|---|---|---|---|---|---|---|---|
| conformer_129 | -930.870156 | 0.284384 | 0.219623 | -930.910446 | -105.8 | -930.626062 | -930.687811 | 2.1 | -925.476030 | -3.992488 | -929.286173 |
| conformer_179 | -930.873457 | 0.284514 | 0.220346 | -930.910423 | -97.1 | -930.625909 | -930.687065 | 4.0 | -925.482213 | -3.990158 | -929.285979 |
| conformer_178 | -930.873256 | 0.284619 | 0.219652 | -930.910417 | -97.6 | -930.625798 | -930.687753 | 2.2 | -925.481938 | -3.990012 | -929.286448 |
| conformer_128 | -930.874766 | 0.284603 | 0.221494 | -930.910325 | -93.4 | -930.625722 | -930.685819 | 7.3 | -925.481177 | -3.991695 | -929.283924 |
| conformer_33 | -930.880779 | 0.284640 | 0.220962 | -930.910221 | -77.3 | -930.625581 | -930.686247 | 6.2 | -925.485553 | -3.993493 | -929.284514 |
| conformer_205 | -930.880541 | 0.284814 | 0.221946 | -930.910215 | -77.9 | -930.625401 | -930.685257 | 8.8 | -925.488982 | -3.990346 | -929.284044 |
| conformer_166 | -930.873893 | 0.284596 | 0.221260 | -930.910209 | -95.3 | -930.625613 | -930.685937 | 7.0 | -925.478837 | -3.993486 | -929.284367 |
| conformer_142 | -930.873346 | 0.284584 | 0.220949 | -930.910185 | -96.7 | -930.625601 | -930.686224 | 6.3 | -925.478702 | -3.993058 | -929.284637 |
| conformer_67 | -930.880617 | 0.284657 | 0.222562 | -930.910161 | -77.6 | -930.625504 | -930.684587 | 10.6 | -925.486248 | -3.993640 | -929.283859 |
| conformer_27 | -930.877966 | 0.284548 | 0.221672 | -930.910148 | -84.5 | -930.625600 | -930.685464 | 8.3 | -925.481978 | -3.994374 | -929.283851 |
| conformer_10 | -930.879885 | 0.284463 | 0.222120 | -930.910146 | -79.4 | -930.625683 | -930.685014 | 9.4 | -925.482244 | -3.996078 | -929.283451 |
| conformer_200 | -930.872898 | 0.284451 | 0.219242 | -930.910141 | -97.8 | -930.625690 | -930.687887 | 1.9 | -925.481651 | -3.990050 | -929.286691 |
| conformer_143 | -930.870871 | 0.284443 | 0.219910 | -930.910060 | -102.9 | -930.625617 | -930.687138 | 3.9 | -925.478326 | -3.990830 | -929.285423 |
| conformer_207 | -930.872814 | 0.284530 | 0.220175 | -930.910045 | -97.8 | -930.625515 | -930.686858 | 4.6 | -925.480969 | -3.990711 | -929.285725 |
| conformer_216 | -930.875120 | 0.284469 | 0.220888 | -930.910019 | -91.6 | -930.625550 | -930.686119 | 6.5 | -925.483534 | -3.990185 | -929.284717 |
| conformer_192 | -930.873959 | 0.284734 | 0.221074 | -930.909972 | -94.6 | -930.625238 | -930.685886 | 7.1 | -925.481206 | -3.991727 | -929.284861 |
| conformer_12 | -930.877992 | 0.284589 | 0.222615 | -930.909949 | -83.9 | -930.625360 | -930.684322 | 11.3 | -925.482135 | -3.994644 | -929.283109 |
| conformer_37 | -930.878785 | 0.284500 | 0.221919 | -930.909932 | -81.8 | -930.625432 | -930.685001 | 9.5 | -925.482492 | -3.994432 | -929.283140 |
| conformer_106 | -930.875569 | 0.284817 | 0.222534 | -930.909910 | -90.2 | -930.625093 | -930.684364 | 11.1 | -925.479782 | -3.994262 | -929.282839 |
| conformer_13 | -930.883495 | 0.284667 | 0.222006 | -930.909893 | -69.3 | -930.625226 | -930.684875 | 9.8 | -925.488940 | -3.993269 | -929.283589 |
| conformer_99 | -930.876336 | 0.284741 | 0.222279 | -930.909888 | -88.1 | -930.625147 | -930.684597 | 10.5 | -925.482687 | -3.992544 | -929.283493 |
| conformer_88 | -930.874841 | 0.284599 | 0.220553 | -930.909872 | -92.0 | -930.625273 | -930.686307 | 6.0 | -925.479259 | -3.993477 | -929.284201 |
| conformer_169 | -930.870890 | 0.284416 | 0.220437 | -930.909850 | -102.3 | -930.625434 | -930.686401 | 5.8 | -925.478918 | -3.990900 | -929.285328 |
| conformer_164 | -930.870719 | 0.284540 | 0.219870 | -930.909806 | -102.6 | -930.625266 | -930.686924 | 4.4 | -925.478742 | -3.990766 | -929.285713 |
| conformer_210 | -930.869836 | 0.284272 | 0.218835 | -930.909762 | -104.8 | -930.625490 | -930.687915 | 1.8 | -925.477447 | -3.990709 | -929.286236 |
| conformer_3 | -930.880364 | 0.284592 | 0.221255 | -930.909753 | -77.2 | -930.625161 | -930.685486 | 8.2 | -925.486624 | -3.992482 | -929.284228 |
| conformer_77 | -930.877853 | 0.284730 | 0.221262 | -930.909659 | -83.5 | -930.624929 | -930.685385 | 8.5 | -925.485617 | -3.991487 | -929.284636 |
| conformer_180 | -930.871702 | 0.284447 | 0.221084 | -930.909636 | -99.6 | -930.625189 | -930.685540 | 8.1 | -925.475739 | -3.993892 | -929.283469 |
| conformer_131 | -930.875854 | 0.284672 | 0.221452 | -930.909612 | -88.6 | -930.624940 | -930.685148 | 9.1 | -925.482165 | -3.992471 | -929.283930 |
| conformer_32 | -930.879590 | 0.284431 | 0.220657 | -930.909610 | -78.8 | -930.625179 | -930.685941 | 7.0 | -925.485448 | -3.992826 | -929.284625 |
| conformer_126 | -930.875090 | 0.284750 | 0.221829 | -930.909609 | -90.6 | -930.624859 | -930.684768 | 10.1 | -925.479254 | -3.994200 | -929.283132 |
| conformer_193 | -930.877186 | 0.284712 | 0.222045 | -930.909547 | -85.0 | -930.624835 | -930.684490 | 10.8 | -925.484661 | -3.991242 | -929.283207 |
| conformer_137 | -930.870437 | 0.284465 | 0.220961 | -930.909513 | -102.6 | -930.625048 | -930.685540 | 8.1 | -925.475027 | -3.993821 | -929.283951 |
| conformer_73 | -930.877182 | 0.284519 | 0.222597 | -930.909473 | -84.8 | -930.624954 | -930.683864 | 12.5 | -925.481834 | -3.994572 | -929.283089 |
| conformer_213 | -930.869874 | 0.284425 | 0.220358 | -930.909392 | -103.8 | -930.624967 | -930.686022 | 6.8 | -925.477296 | -3.991501 | -929.284946 |
| conformer_90 | -930.871980 | 0.284501 | 0.220834 | -930.909391 | -98.2 | -930.624890 | -930.685545 | 8.0 | -925.475669 | -3.994227 | -929.283460 |
| conformer_204 | -930.872519 | 0.284390 | 0.221058 | -930.909337 | -96.7 | -930.624947 | -930.685267 | 8.8 | -925.480097 | -3.991026 | -929.283870 |
| conformer_69 | -930.877331 | 0.284645 | 0.220658 | -930.909301 | -83.9 | -930.624656 | -930.685631 | 7.8 | -925.485088 | -3.991417 | -929.284806 |
| conformer_6 | -930.879800 | 0.284494 | 0.219939 | -930.909284 | -77.4 | -930.624790 | -930.686333 | 6.0 | -925.486090 | -3.992361 | -929.284983 |
| conformer_100 | -930.872555 | 0.284708 | 0.222574 | -930.909239 | -96.3 | -930.624531 | -930.683653 | 13.0 | -925.476001 | -3.995038 | -929.282138 |
| conformer_104 | -930.873309 | 0.284626 | 0.222316 | -930.909195 | -94.2 | -930.624569 | -930.683867 | 12.4 | -925.478878 | -3.993351 | -929.282787 |
| conformer_87 | -930.875822 | 0.284927 | 0.221811 | -930.909133 | -87.5 | -930.624206 | -930.684310 | 11.3 | -925.483248 | -3.991212 | -929.282949 |
| conformer_42 | -930.879031 | 0.284333 | 0.219314 | -930.909129 | -79.0 | -930.624796 | -930.686803 | 4.7 | -925.484918 | -3.992677 | -929.285368 |
| conformer_4 | -930.877015 | 0.284483 | 0.221333 | -930.909107 | -84.3 | -930.624624 | -930.684762 | 10.1 | -925.482414 | -3.993367 | -929.283528 |
| conformer_194 | -930.872311 | 0.284328 | 0.219160 | -930.909090 | -96.6 | -930.624762 | -930.686918 | 4.4 | -925.482221 | -3.989123 | -929.285952 |
| conformer_199 | -930.874041 | 0.284464 | 0.220542 | -930.908979 | -91.7 | -930.624515 | -930.685425 | 8.4 | -925.482028 | -3.991182 | -929.284594 |
| conformer_161 | -930.874757 | 0.284622 | 0.221321 | -930.908975 | -89.8 | -930.624353 | -930.684642 | 10.4 | -925.481747 | -3.992328 | -929.283959 |
| conformer_41 | -930.876153 | 0.284296 | 0.220271 | -930.908936 | -86.1 | -930.624640 | -930.685653 | 7.8 | -925.481171 | -3.993662 | -929.284333 |
| conformer_50 | -930.879659 | 0.284540 | 0.221262 | -930.908753 | -76.4 | -930.624213 | -930.684479 | 10.8 | -925.486431 | -3.992282 | -929.283531 |
| conformer_168 | -930.875271 | 0.284832 | 0.221025 | -930.908717 | -87.8 | -930.623885 | -930.684680 | 10.3 | -925.482680 | -3.991113 | -929.283203 |
| conformer_187 | -930.869712 | 0.284243 | 0.219300 | -930.908470 | -101.8 | -930.624227 | -930.686158 | 6.4 | -925.478865 | -3.989903 | -929.285214 |
| conformer_156 | -930.869512 | 0.284195 | 0.218745 | -930.908464 | -102.3 | -930.624269 | -930.686707 | 5.0 | -925.476205 | -3.991934 | -929.285334 |
| conformer_83 | -930.872880 | 0.284826 | 0.221937 | -930.908449 | -93.4 | -930.623623 | -930.683500 | 13.4 | -925.479541 | -3.992059 | -929.282220 |
| conformer_113 | -930.872878 | 0.284612 | 0.221131 | -930.908392 | -93.2 | -930.623780 | -930.684249 | 11.4 | -925.478017 | -3.992976 | -929.282365 |
| conformer_8 | -930.880519 | 0.284769 | 0.221344 | -930.908313 | -73.0 | -930.623544 | -930.683957 | 12.2 | -925.488467 | -3.991531 | -929.283437 |
| conformer_195 | -930.870936 | 0.284352 | 0.220613 | -930.908307 | -98.1 | -930.623955 | -930.684682 | 10.3 | -925.478133 | -3.992039 | -929.283918 |

| | | | | | | | | | | | |
|---|---|---|---|---|---|---|---|---|---|---|---|
| conformer_190 | -930.869649 | 0.284536 | 0.219761 | -930.908192 | -101.2 | -930.623656 | -930.685419 | 8.4 | -925.476718 | -3.991362 | -929.283849 |
| conformer_53 | -930.876388 | 0.284446 | 0.221405 | -930.908125 | -83.3 | -930.623679 | -930.683708 | 12.9 | -925.482277 | -3.993173 | -929.282770 |
| conformer_209 | -930.875431 | 0.284498 | 0.220965 | -930.908011 | -85.5 | -930.623513 | -930.684034 | 12.0 | -925.483463 | -3.991253 | -929.283318 |
| conformer_135 | -930.871781 | 0.284564 | 0.221032 | -930.907970 | -95.0 | -930.623406 | -930.683926 | 12.3 | -925.477506 | -3.992727 | -929.282378 |
| conformer_115 | -930.869932 | 0.284539 | 0.221321 | -930.907811 | -99.5 | -930.623272 | -930.683478 | 13.5 | -925.474260 | -3.993819 | -929.281624 |
| conformer_189 | -930.874415 | 0.284514 | 0.220092 | -930.907740 | -87.5 | -930.623226 | -930.684636 | 10.4 | -925.482056 | -3.991151 | -929.283427 |
| conformer_162 | -930.873194 | 0.284399 | 0.219684 | -930.907720 | -90.6 | -930.623321 | -930.685024 | 9.4 | -925.481223 | -3.990919 | -929.283972 |
| conformer_163 | -930.880203 | 0.284769 | 0.221312 | -930.907710 | -72.2 | -930.622941 | -930.683386 | 13.7 | -925.489353 | -3.990528 | -929.283064 |
| conformer_147 | -930.871331 | 0.284506 | 0.220148 | -930.907684 | -95.4 | -930.623178 | -930.684524 | 10.7 | -925.477024 | -3.992652 | -929.282868 |
| conformer_18 | -930.879880 | 0.284703 | 0.220233 | -930.907638 | -72.9 | -930.622935 | -930.684393 | 11.1 | -925.487753 | -3.991545 | -929.283812 |
| conformer_15 | -930.877208 | 0.284615 | 0.221278 | -930.907635 | -79.9 | -930.623020 | -930.683345 | 13.8 | -925.484341 | -3.992318 | -929.282796 |
| conformer_157 | -930.872532 | 0.284756 | 0.221170 | -930.907600 | -92.1 | -930.622844 | -930.683418 | 13.6 | -925.479036 | -3.991862 | -929.281784 |
| conformer_92 | -930.875438 | 0.284723 | 0.221617 | -930.907480 | -84.1 | -930.622757 | -930.682851 | 15.1 | -925.482598 | -3.991764 | -929.281775 |
| conformer_153 | -930.872164 | 0.284641 | 0.221644 | -930.907373 | -92.4 | -930.622732 | -930.682717 | 15.5 | -925.478148 | -3.992477 | -929.281178 |
| conformer_152 | -930.870737 | 0.284366 | 0.220732 | -930.907339 | -96.1 | -930.622973 | -930.683595 | 13.2 | -925.478027 | -3.991759 | -929.282643 |
| conformer_74 | -930.877437 | 0.284709 | 0.221450 | -930.907309 | -78.4 | -930.622600 | -930.682847 | 15.1 | -925.485590 | -3.991078 | -929.282079 |
| conformer_22 | -930.879528 | 0.284678 | 0.218944 | -930.907139 | -72.5 | -930.622461 | -930.685183 | 9.0 | -925.488633 | -3.990505 | -929.284794 |
| conformer_98 | -930.874945 | 0.284646 | 0.220943 | -930.907122 | -84.5 | -930.622476 | -930.683167 | 14.3 | -925.482073 | -3.991645 | -929.281940 |
| conformer_174 | -930.871916 | 0.284425 | 0.220227 | -930.907088 | -92.3 | -930.622663 | -930.683849 | 12.5 | -925.478903 | -3.991883 | -929.282720 |
| conformer_16 | -930.876871 | 0.284652 | 0.221327 | -930.907049 | -79.2 | -930.622397 | -930.682710 | 15.5 | -925.485194 | -3.991310 | -929.282343 |
| conformer_101 | -930.876918 | 0.284632 | 0.220624 | -930.906994 | -79.0 | -930.622362 | -930.683358 | 13.8 | -925.485036 | -3.990985 | -929.282461 |
| conformer_116 | -930.874512 | 0.284540 | 0.220733 | -930.906964 | -85.2 | -930.622424 | -930.683219 | 14.1 | -925.483329 | -3.990389 | -929.282425 |
| conformer_45 | -930.877879 | 0.284622 | 0.221090 | -930.906890 | -76.2 | -930.622268 | -930.682788 | 15.3 | -925.487356 | -3.990457 | -929.282722 |
| conformer_134 | -930.874085 | 0.284482 | 0.219907 | -930.906693 | -85.6 | -930.622211 | -930.683774 | 12.7 | -925.482881 | -3.990304 | -929.282874 |
| conformer_144 | -930.869356 | 0.284553 | 0.221787 | -930.906693 | -98.0 | -930.622140 | -930.681894 | 17.6 | -925.474596 | -3.993347 | -929.280480 |
| conformer_201 | -930.874715 | 0.284473 | 0.219954 | -930.906677 | -83.9 | -930.622204 | -930.683711 | 12.9 | -925.483215 | -3.990751 | -929.282962 |
| conformer_78 | -930.874495 | 0.284602 | 0.221545 | -930.906619 | -84.3 | -930.622017 | -930.682062 | 17.2 | -925.481852 | -3.991950 | -929.281370 |
| conformer_172 | -930.872216 | 0.284493 | 0.220428 | -930.906606 | -90.3 | -930.622113 | -930.683166 | 14.3 | -925.478838 | -3.992144 | -929.281932 |
| conformer_177 | -930.868266 | 0.284278 | 0.219436 | -930.906493 | -100.4 | -930.622215 | -930.684045 | 12.0 | -925.473515 | -3.993230 | -929.282524 |
| conformer_80 | -930.877690 | 0.284574 | 0.220993 | -930.906353 | -75.3 | -930.621779 | -930.682348 | 16.4 | -925.487622 | -3.990148 | -929.282427 |
| conformer_208 | -930.878132 | 0.284689 | 0.221258 | -930.906346 | -74.1 | -930.621657 | -930.682076 | 17.1 | -925.487037 | -3.990718 | -929.281700 |
| conformer_182 | -930.871790 | 0.284436 | 0.219596 | -930.906319 | -90.7 | -930.621883 | -930.683711 | 12.9 | -925.478380 | -3.992047 | -929.282348 |
| conformer_202 | -930.871490 | 0.284441 | 0.220912 | -930.906311 | -91.4 | -930.621870 | -930.682387 | 16.3 | -925.479538 | -3.991166 | -929.281601 |
| conformer_186 | -930.872293 | 0.284397 | 0.220118 | -930.906027 | -88.6 | -930.621630 | -930.682897 | 15.0 | -925.480117 | -3.991486 | -929.282207 |
| conformer_1 | -930.879960 | 0.284909 | 0.222021 | -930.906023 | -68.4 | -930.621114 | -930.680990 | 20.0 | -925.488460 | -3.990830 | -929.280320 |
| conformer_181 | -930.869229 | 0.284404 | 0.220657 | -930.905976 | -96.5 | -930.621572 | -930.682307 | 16.5 | -925.475081 | -3.992938 | -929.281097 |
| conformer_89 | -930.874281 | 0.284437 | 0.220930 | -930.905671 | -82.4 | -930.621234 | -930.681729 | 18.1 | -925.483342 | -3.990979 | -929.281769 |
| conformer_71 | -930.875090 | 0.284577 | 0.221360 | -930.905648 | -80.2 | -930.621071 | -930.681276 | 19.2 | -925.483214 | -3.991566 | -929.280966 |
| conformer_2 | -930.876555 | 0.284740 | 0.221867 | -930.905353 | -75.6 | -930.620613 | -930.680474 | 21.4 | -925.484199 | -3.991660 | -929.279777 |

[a]: Single-point calculation in aqueous phase with SMD model.

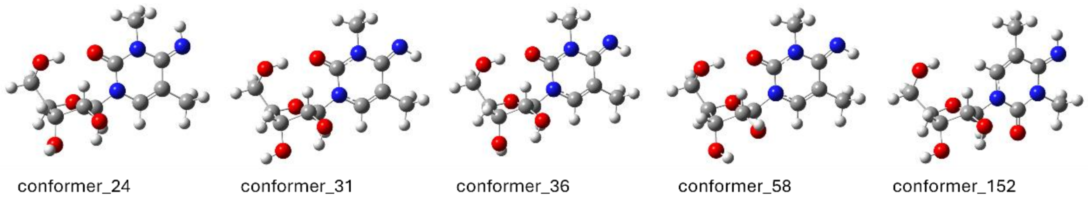

**Figure S27.** B3LYP-D3/def2-TZVPP optimized geometries of conformers for neutral 3,5-dimethylcytidine (**1rb3m5mC**).

**Table S25.** Conformers of gas-phase optimized 3,5-dimethylcytidine (**1rb3m5mC**) at the B3LYP-D3/def2-TZVPP level of theory followed by aqueous phase single-point calculation. The columns display total energy without zero-point correction ($E_{Tot}$), thermal correction to enthalpy (δH), Gibbs free energy (δG), total energy without zero-point correction ($E_{Tot,W}$), Gibbs free energy ($G_{298,W}$) in water (W), total single-point energy ($E_{CBS}$) calculated at DLPNO-CCSD(T)/CBS level of theory, and their corresponding free energy $G_{CBS,W}$. $G_{298,W}$ and $G_{CBS,W}$ have been corrected to the standard state of 1 mol/L by addition of +7.908 kJ/mol. $\Delta G_{Solv}$ represents the Gibbs free energy of solvation. The data are arranged in the ascending numeric order of $E_{Tot,W}$. $\Delta G_{298,W}$ represents the respective energy difference to the lowest structure. Only conformers within the 24 kJ/mol (6 kcal/mol) energy window above the lowest in CREST are included in initial conformer sampling. Duplicates of the same structure are excluded. The overall optimum is marked bold.

| | B3LYP-D3/def2-TZVPP | | | SMD(H2O)/B3LYP-D3/def2-TZVPP[a] | | | | | DLPNO-CCSD(T)/CBS | | |
|---|---|---|---|---|---|---|---|---|---|---|---|
| 1rb3m5mC No. | $E_{Tot}$ (Hartree) | δH (Hartree) | δG (Hartree) | $E_{Tot,W}$ (Hartree) | $\Delta G_{Solv}$ (kJ/mol) | $H_{298,W}$ (Hartree) | $G_{298,W}$ (Hartree) | $\Delta G_{298,W}$ (kJ/mol) | $E_{CBS,HF}$ (Hartree) | $E_{CBS,C}$ (Hartree) | $G_{CBS,W}$ (Hartree) |
| conformer_58 | -970.219369 | 0.314433 | 0.249772 | -970.249598 | -79.4 | -969.935165 | -969.996814 | 5.0 | -964.541091 | -4.204130 | -968.522666 |
| conformer_36 | -970.216587 | 0.314333 | 0.249458 | -970.249576 | -86.6 | -969.935243 | -969.997106 | 4.2 | -964.536406 | -4.205269 | -968.522194 |
| conformer_31 | -970.218466 | 0.314301 | 0.249363 | -970.249505 | -81.5 | -969.935204 | -969.997130 | 4.2 | -964.540150 | -4.203924 | -968.522738 |
| conformer_24 | -970.215818 | 0.314249 | 0.248898 | -970.248884 | -86.8 | -969.934635 | -969.996974 | 4.6 | -964.536610 | -4.204624 | -968.522389 |
| conformer_106 | -970.216237 | 0.314301 | 0.248900 | -970.248581 | -84.9 | -969.934280 | -969.996669 | 5.4 | -964.537845 | -4.203777 | -968.522053 |
| conformer_20 | -970.214280 | 0.314195 | 0.249314 | -970.248154 | -88.9 | -969.933959 | -969.995828 | 7.6 | -964.533771 | -4.205631 | -968.520950 |
| conformer_50 | -970.216895 | 0.314280 | 0.249587 | -970.248122 | -82.0 | -969.933842 | -969.995523 | 8.4 | -964.538234 | -4.204527 | -968.521390 |
| conformer_11 | -970.216104 | 0.314158 | 0.249194 | -970.248045 | -83.9 | -969.933887 | -969.995839 | 7.6 | -964.537447 | -4.204302 | -968.521483 |
| conformer_37 | -970.218199 | 0.314185 | 0.248865 | -970.247643 | -77.3 | -969.933458 | -969.995766 | 7.7 | -964.539748 | -4.204247 | -968.521562 |
| conformer_18 | -970.219543 | 0.314029 | 0.248309 | -970.247283 | -72.8 | -969.933254 | -969.995962 | 7.2 | -964.539132 | -4.206124 | -968.521674 |
| conformer_47 | -970.216728 | 0.314197 | 0.248114 | -970.247236 | -80.1 | -969.933039 | -969.996110 | 6.8 | -964.540537 | -4.201934 | -968.521853 |
| conformer_48 | -970.215869 | 0.314234 | 0.247433 | -970.247160 | -82.2 | -969.932926 | -969.996715 | 5.3 | -964.539638 | -4.201931 | -968.522415 |
| conformer_124 | -970.209370 | 0.313907 | 0.246329 | -970.247065 | -99.0 | -969.933158 | -969.997724 | 2.6 | -964.532154 | -4.202905 | -968.523413 |
| conformer_33 | -970.213893 | 0.314123 | 0.248661 | -970.247057 | -87.1 | -969.932934 | -969.995384 | 8.8 | -964.535191 | -4.204171 | -968.520852 |
| **conformer_152** | **-970.210896** | **0.313880** | **0.245269** | **-970.246998** | **-94.8** | **-969.933118** | **-969.998717** | **0** | **-964.535453** | **-4.201630** | **-968.524904** |
| conformer_34 | -970.218858 | 0.314108 | 0.248449 | -970.246732 | -73.2 | -969.932624 | -969.995271 | 9.0 | -964.539844 | -4.204454 | -968.520712 |
| conformer_30 | -970.217973 | 0.314024 | 0.247947 | -970.246666 | -75.3 | -969.932642 | -969.995707 | 7.9 | -964.539248 | -4.204514 | -968.521497 |
| conformer_10 | -970.218302 | 0.314142 | 0.248887 | -970.246652 | -74.4 | -969.932510 | -969.994753 | 10.4 | -964.539684 | -4.204784 | -968.520919 |
| conformer_155 | -970.209737 | 0.314001 | 0.245886 | -970.246589 | -96.8 | -969.932588 | -969.997691 | 2.7 | -964.535085 | -4.200747 | -968.523786 |
| conformer_69 | -970.213358 | 0.314330 | 0.247457 | -970.246473 | -86.9 | -969.932143 | -969.996004 | 7.1 | -964.537981 | -4.201553 | -968.522181 |
| conformer_156 | -970.209879 | 0.313853 | 0.246263 | -970.246435 | -96.0 | -969.932582 | -969.997160 | 4.1 | -964.535131 | -4.201013 | -968.523424 |
| conformer_103 | -970.208296 | 0.313878 | 0.246363 | -970.246385 | -100.0 | -969.932507 | -969.997010 | 4.5 | -964.530954 | -4.203199 | -968.522868 |
| conformer_133 | -970.207322 | 0.313872 | 0.246151 | -970.246314 | -102.4 | -969.932442 | -969.997151 | 4.1 | -964.531424 | -4.201452 | -968.522705 |
| conformer_65 | -970.217095 | 0.314018 | 0.248633 | -970.246239 | -76.5 | -969.932221 | -969.994594 | 10.8 | -964.538923 | -4.204808 | -968.521231 |
| conformer_21 | -970.215697 | 0.314002 | 0.248655 | -970.246122 | -79.9 | -969.932120 | -969.994455 | 11.2 | -964.536892 | -4.204665 | -968.520316 |
| conformer_66 | -970.216788 | 0.314145 | 0.247874 | -970.245963 | -76.6 | -969.931818 | -969.995077 | 9.6 | -964.541595 | -4.201312 | -968.521197 |
| conformer_43 | -970.213765 | 0.313982 | 0.247791 | -970.245755 | -84.0 | -969.931773 | -969.994952 | 9.9 | -964.537041 | -4.202471 | -968.520699 |
| conformer_42 | -970.213269 | 0.314045 | 0.247152 | -970.245754 | -85.3 | -969.931709 | -969.995590 | 8.2 | -964.536607 | -4.202384 | -968.521312 |
| conformer_107 | -970.212044 | 0.314254 | 0.249030 | -970.245675 | -88.3 | -969.931421 | -969.993633 | 13.3 | -964.532938 | -4.204954 | -968.519481 |
| conformer_4 | -970.217137 | 0.313940 | 0.248447 | -970.245644 | -74.8 | -969.931704 | -969.994185 | 11.9 | -964.536054 | -4.206838 | -968.519941 |

| | | | | | | | | | | | |
|---|---|---|---|---|---|---|---|---|---|---|---|
| conformer_87 | -970.207418 | 0.313755 | 0.246332 | -970.245639 | -100.3 | -969.931884 | -969.996295 | 6.4 | -964.529949 | -4.203169 | -968.521994 |
| conformer_108 | -970.212744 | 0.314167 | 0.248692 | -970.245626 | -86.3 | -969.931459 | -969.993922 | 12.6 | -964.535764 | -4.203251 | -968.520192 |
| conformer_62 | -970.214252 | 0.314100 | 0.247064 | -970.245619 | -82.4 | -969.931519 | -969.995543 | 8.3 | -964.538689 | -4.202115 | -968.522094 |
| conformer_7 | -970.216863 | 0.314123 | 0.247976 | -970.245561 | -75.3 | -969.931438 | -969.994573 | 10.9 | -964.539640 | -4.203197 | -968.520547 |
| conformer_116 | -970.208976 | 0.313699 | 0.244964 | -970.245547 | -96.0 | -969.931848 | -969.997571 | 3.0 | -964.533295 | -4.201910 | -968.523800 |
| conformer_38 | -970.216138 | 0.314003 | 0.247750 | -970.245487 | -77.1 | -969.931484 | -969.994725 | 10.5 | -964.538484 | -4.203581 | -968.520650 |
| conformer_101 | -970.207423 | 0.313888 | 0.247620 | -970.245387 | -99.7 | -969.931499 | -969.994755 | 10.4 | -964.529037 | -4.203429 | -968.519797 |
| conformer_13 | -970.215214 | 0.313898 | 0.247901 | -970.245200 | -78.7 | -969.931302 | -969.994287 | 11.6 | -964.536188 | -4.204818 | -968.520080 |
| conformer_119 | -970.207893 | 0.313946 | 0.246132 | -970.245172 | -97.9 | -969.931226 | -969.996028 | 7.1 | -964.533003 | -4.201039 | -968.522177 |
| conformer_27 | -970.215783 | 0.313912 | 0.248111 | -970.245052 | -76.8 | -969.931140 | -969.993929 | 12.6 | -964.536321 | -4.204921 | -968.519387 |
| conformer_5 | -970.215115 | 0.313947 | 0.248583 | -970.245052 | -78.6 | -969.931105 | -969.993457 | 13.8 | -964.536002 | -4.205250 | -968.519595 |
| conformer_78 | -970.212271 | 0.314369 | 0.248206 | -970.245031 | -86.0 | -969.930662 | -969.993813 | 12.9 | -964.536376 | -4.201812 | -968.519731 |
| conformer_121 | -970.207974 | 0.313700 | 0.246175 | -970.245020 | -97.3 | -969.931320 | -969.995833 | 7.6 | -964.532951 | -4.201318 | -968.522128 |
| conformer_54 | -970.211044 | 0.314189 | 0.247399 | -970.245000 | -89.2 | -969.930811 | -969.994589 | 10.8 | -964.535381 | -4.201879 | -968.520806 |
| conformer_19 | -970.216148 | 0.314035 | 0.246662 | -970.244989 | -75.7 | -969.930954 | -969.995315 | 8.9 | -964.538868 | -4.203152 | -968.521187 |
| conformer_97 | -970.205332 | 0.313733 | 0.246082 | -970.244917 | -103.9 | -969.931184 | -969.995823 | 7.6 | -964.529053 | -4.201918 | -968.521461 |
| conformer_95 | -970.208970 | 0.313945 | 0.247780 | -970.244790 | -94.0 | -969.930845 | -969.993998 | 12.4 | -964.531574 | -4.202889 | -968.519490 |
| conformer_144 | -970.204333 | 0.313643 | 0.245668 | -970.244784 | -106.2 | -969.931141 | -969.996104 | 6.9 | -964.527965 | -4.201933 | -968.521669 |
| conformer_59 | -970.209290 | 0.313930 | 0.247012 | -970.244781 | -93.2 | -969.930851 | -969.994757 | 10.4 | -964.529838 | -4.204715 | -968.520020 |
| conformer_55 | -970.214214 | 0.313851 | 0.248441 | -970.244747 | -80.2 | -969.930896 | -969.993294 | 14.2 | -964.535413 | -4.205424 | -968.519917 |
| conformer_51 | -970.216076 | 0.314027 | 0.247661 | -970.244514 | -74.7 | -969.930487 | -969.993841 | 12.8 | -964.539396 | -4.202983 | -968.520144 |
| conformer_159 | -970.209324 | 0.313799 | 0.247233 | -970.244462 | -92.3 | -969.930663 | -969.994217 | 11.8 | -964.533931 | -4.201337 | -968.520161 |
| conformer_56 | -970.213941 | 0.313938 | 0.247593 | -970.244449 | -80.1 | -969.930511 | -969.993844 | 12.8 | -964.538214 | -4.201868 | -968.519986 |
| conformer_148 | -970.206930 | 0.313912 | 0.246847 | -970.244403 | -98.4 | -969.930491 | -969.994544 | 11.0 | -964.531411 | -4.201686 | -968.520711 |
| conformer_14 | -970.217069 | 0.313961 | 0.247969 | -970.244259 | -71.4 | -969.930298 | -969.993278 | 14.3 | -964.538413 | -4.204803 | -968.519425 |
| conformer_73 | -970.209656 | 0.314107 | 0.248833 | -970.244212 | -90.7 | -969.930105 | -969.992367 | 16.7 | -964.530212 | -4.205290 | -968.518213 |
| conformer_88 | -970.210331 | 0.314022 | 0.248547 | -970.244123 | -88.7 | -969.930101 | -969.992564 | 16.2 | -964.533010 | -4.203617 | -968.518861 |
| conformer_52 | -970.211768 | 0.313937 | 0.246942 | -970.244108 | -84.9 | -969.930171 | -969.994154 | 12.0 | -964.535898 | -4.202487 | -968.520771 |
| conformer_2 | -970.214134 | 0.313974 | 0.247789 | -970.244063 | -78.6 | -969.930089 | -969.993262 | 14.3 | -964.536456 | -4.203679 | -968.519263 |
| conformer_117 | -970.209259 | 0.314036 | 0.247277 | -970.244025 | -91.3 | -969.929989 | -969.993736 | 13.1 | -964.530810 | -4.203921 | -968.519208 |
| conformer_26 | -970.213317 | 0.313847 | 0.247529 | -970.244015 | -80.6 | -969.930168 | -969.993474 | 13.8 | -964.535196 | -4.204038 | -968.519392 |
| conformer_22 | -970.216957 | 0.314186 | 0.247681 | -970.243756 | -70.4 | -969.929570 | -969.993063 | 14.8 | -964.541414 | -4.202366 | -968.519886 |
| conformer_93 | -970.206813 | 0.313684 | 0.245771 | -970.243720 | -96.9 | -969.930036 | -969.994937 | 9.9 | -964.530754 | -4.202099 | -968.520977 |
| conformer_143 | -970.209963 | 0.313939 | 0.247137 | -970.243660 | -88.5 | -969.929721 | -969.993511 | 13.7 | -964.534700 | -4.201580 | -968.519828 |
| conformer_129 | -970.208148 | 0.314022 | 0.247503 | -970.243562 | -93.0 | -969.929540 | -969.993047 | 14.9 | -964.530570 | -4.203342 | -968.518811 |
| conformer_25 | -970.216592 | 0.314199 | 0.247751 | -970.243530 | -70.7 | -969.929331 | -969.992767 | 15.6 | -964.542318 | -4.201314 | -968.519807 |
| conformer_63 | -970.209951 | 0.314202 | 0.247987 | -970.243446 | -87.9 | -969.929244 | -969.992447 | 16.5 | -964.533727 | -4.202176 | -968.518398 |
| conformer_131 | -970.207920 | 0.313675 | 0.246388 | -970.243431 | -93.2 | -969.929756 | -969.994031 | 12.3 | -964.532265 | -4.202209 | -968.520585 |
| conformer_94 | -970.211875 | 0.314180 | 0.247969 | -970.243414 | -82.8 | -969.929234 | -969.992433 | 16.5 | -964.535731 | -4.202352 | -968.518641 |
| conformer_145 | -970.208887 | 0.314183 | 0.247464 | -970.243388 | -90.6 | -969.929205 | -969.992912 | 15.2 | -964.532119 | -4.202508 | -968.518651 |
| conformer_130 | -970.206824 | 0.313897 | 0.246045 | -970.243384 | -96.0 | -969.929487 | -969.994327 | 11.5 | -964.530918 | -4.201702 | -968.520122 |
| conformer_120 | -970.208619 | 0.314012 | 0.247786 | -970.243218 | -90.8 | -969.929206 | -969.992420 | 16.5 | -964.531368 | -4.203095 | -968.518264 |
| conformer_75 | -970.213798 | 0.314153 | 0.247787 | -970.243133 | -77.0 | -969.928980 | -969.992334 | 16.8 | -964.538675 | -4.201640 | -968.518851 |
| conformer_44 | -970.213428 | 0.313882 | 0.247483 | -970.243045 | -77.8 | -969.929163 | -969.992550 | 16.2 | -964.536274 | -4.203461 | -968.518857 |
| conformer_158 | -970.207077 | 0.313666 | 0.245518 | -970.243030 | -94.4 | -969.929364 | -969.994500 | 11.1 | -964.533846 | -4.199774 | -968.521043 |
| conformer_83 | -970.206955 | 0.313922 | 0.247265 | -970.242649 | -93.7 | -969.928727 | -969.992372 | 16.7 | -964.528089 | -4.204330 | -968.517835 |
| conformer_126 | -970.209005 | 0.313849 | 0.246860 | -970.242340 | -87.5 | -969.928491 | -969.992468 | 16.4 | -964.533019 | -4.202232 | -968.518715 |
| conformer_6 | -970.214285 | 0.314058 | 0.247548 | -970.242287 | -73.5 | -969.928229 | -969.991727 | 18.4 | -964.538335 | -4.202735 | -968.518513 |
| conformer_112 | -970.207685 | 0.313840 | 0.247205 | -970.242250 | -90.8 | -969.928410 | -969.992033 | 17.5 | -964.532095 | -4.201944 | -968.518386 |
| conformer_149 | -970.208591 | 0.313948 | 0.246939 | -970.242237 | -88.3 | -969.928289 | -969.992286 | 16.9 | -964.531876 | -4.202780 | -968.518350 |
| conformer_96 | -970.205945 | 0.313937 | 0.247620 | -970.242154 | -95.1 | -969.928217 | -969.991522 | 18.9 | -964.528044 | -4.203655 | -968.517276 |
| conformer_85 | -970.213971 | 0.313916 | 0.247119 | -970.242120 | -73.9 | -969.928204 | -969.991989 | 17.7 | -964.540510 | -4.200957 | -968.519485 |
| conformer_8 | -970.213877 | 0.314055 | 0.247597 | -970.242044 | -74.0 | -969.927989 | -969.991435 | 19.1 | -964.539183 | -4.201705 | -968.518446 |
| conformer_68 | -970.214489 | 0.314125 | 0.247496 | -970.242028 | -72.3 | -969.927903 | -969.991520 | 18.9 | -964.540058 | -4.201317 | -968.518406 |
| conformer_71 | -970.209413 | 0.314035 | 0.247832 | -970.241839 | -85.1 | -969.927804 | -969.990995 | 20.3 | -964.532888 | -4.202774 | -968.517244 |
| conformer_113 | -970.206573 | 0.313995 | 0.247157 | -970.241827 | -92.6 | -969.927832 | -969.991658 | 18.5 | -964.529495 | -4.202856 | -968.517435 |

| | | | | | | | | | | | |
|---|---|---|---|---|---|---|---|---|---|---|---|
| conformer_86 | -970.206456 | 0.313882 | 0.247668 | -970.241756 | -92.7 | -969.927874 | -969.991076 | 20.1 | -964.528894 | -4.203468 | -968.516982 |
| conformer_60 | -970.211471 | 0.313991 | 0.247617 | -970.241562 | -79.0 | -969.927571 | -969.990933 | 20.4 | -964.535992 | -4.202060 | -968.517514 |
| conformer_3 | -970.216358 | 0.314329 | 0.248280 | -970.241465 | -65.9 | -969.927136 | -969.990173 | 22.4 | -964.541428 | -4.201628 | -968.516871 |
| conformer_122 | -970.205314 | 0.313793 | 0.246331 | -970.241315 | -94.5 | -969.927522 | -969.991972 | 17.7 | -964.527626 | -4.203445 | -968.517728 |
| conformer_99 | -970.208439 | 0.313880 | 0.247216 | -970.241239 | -86.1 | -969.927359 | -969.991011 | 20.2 | -964.533587 | -4.201365 | -968.517525 |
| conformer_134 | -970.209348 | 0.313833 | 0.246774 | -970.241203 | -83.6 | -969.927370 | -969.991417 | 19.2 | -964.534246 | -4.201799 | -968.518114 |
| conformer_41 | -970.211301 | 0.313819 | 0.247036 | -970.241197 | -78.5 | -969.927378 | -969.991149 | 19.9 | -964.536896 | -4.201708 | -968.518452 |
| conformer_123 | -970.206242 | 0.313857 | 0.247055 | -970.240842 | -90.8 | -969.926985 | -969.990775 | 20.9 | -964.529189 | -4.203131 | -968.516852 |
| conformer_67 | -970.211151 | 0.313779 | 0.247005 | -970.240655 | -77.5 | -969.926876 | -969.990638 | 21.2 | -964.537217 | -4.201401 | -968.518105 |
| conformer_57 | -970.212054 | 0.313953 | 0.247342 | -970.240462 | -74.6 | -969.926509 | -969.990108 | 22.6 | -964.537287 | -4.201721 | -968.517061 |
| conformer_1 | -970.213561 | 0.314202 | 0.248225 | -970.240035 | -69.5 | -969.925833 | -969.988798 | 26.0 | -964.538176 | -4.202033 | -968.515447 |
| conformer_104 | -970.204908 | 0.313988 | 0.247048 | -970.238990 | -89.5 | -969.925002 | -969.988930 | 25.7 | -964.523530 | -4.207586 | -968.515138 |

[a]: Single-point calculation in aqueous phase with SMD model.

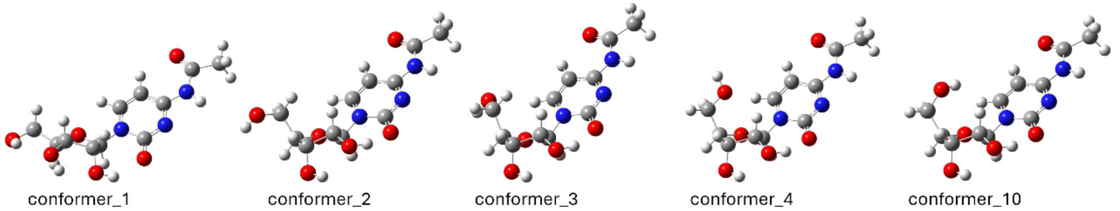

**Figure S28.** B3LYP-D3/def2-TZVPP optimized geometries of conformers for 4-acetylcytidine (**1rb4acC**).

**Table S26.** Conformers of gas-phase optimized 4-acetylcytidine (**1rb4acC**) at the B3LYP-D3/def2-TZVPP level of theory followed by aqueous phase single-point calculation. The columns display total energy without zero-point correction ($E_{Tot}$), thermal correction to enthalpy (δH), Gibbs free energy (δG), total energy without zero-point correction ($E_{Tot,W}$), Gibbs free energy ($G_{298,W}$) in water (W), total single-point energy ($E_{CBS}$) calculated at DLPNO-CCSD(T)/CBS level of theory, and their corresponding free energy $G_{CBS,W}$. $G_{298,W}$ and $G_{CBS,W}$ have been corrected to the standard state of 1 mol/L by addition of +7.908 kJ/mol. $\Delta G_{Solv}$ represents the Gibbs free energy of solvation. The data are arranged in the ascending numeric order of $E_{Tot,W}$. $\Delta G_{298,W}$ represents the respective energy difference to the lowest structure. Only conformers within the 24 kJ/mol (6 kcal/mol) energy window above the lowest in CREST are included in initial conformer sampling. Duplicates of the same structure are excluded. The overall optimum is marked bold.

| | B3LYP-D3/def2-TZVPP | | | SMD(H2O)/B3LYP-D3/def2-TZVPP[a] | | | | | DLPNO-CCSD(T)/CBS | | |
|---|---|---|---|---|---|---|---|---|---|---|---|
| 1rb4acC No. | $E_{Tot}$ (Hartree) | δH (Hartree) | δG (Hartree) | $E_{Tot,W}$ (Hartree) | $\Delta G_{Solv}$ (kJ/mol) | $H_{298,W}$ (Hartree) | $G_{298,W}$ (Hartree) | $\Delta G_{298,W}$ (kJ/mol) | $E_{CBS,HF}$ (Hartree) | $E_{CBS,C}$ (Hartree) | $G_{CBS,W}$ (Hartree) |
| **conformer_10** | **-1044.287910** | **0.295905** | **0.227591** | **-1044.326571** | **-101.5** | **-1044.030666** | **-1044.095968** | **0** | **-1038.298891** | **-4.422627** | **-1042.529577** |
| conformer_20 | -1044.285396 | 0.296110 | 0.228307 | -1044.326934 | -109.1 | -1044.030824 | -1044.095615 | 0.9 | -1038.297802 | -4.421114 | -1042.529135 |
| conformer_47 | -1044.286261 | 0.296198 | 0.228596 | -1044.327138 | -107.3 | -1044.030940 | -1044.095530 | 1.2 | -1038.298818 | -4.421241 | -1042.529329 |
| conformer_38 | -1044.284475 | 0.296038 | 0.227039 | -1044.325576 | -107.9 | -1044.029538 | -1044.095525 | 1.2 | -1038.299448 | -4.418858 | -1042.529357 |
| conformer_26 | -1044.283498 | 0.296110 | 0.228404 | -1044.326904 | -114.0 | -1044.030794 | -1044.095488 | 1.3 | -1038.294077 | -4.422396 | -1042.528464 |
| conformer_11 | -1044.286517 | 0.295853 | 0.227110 | -1044.325503 | -102.4 | -1044.029650 | -1044.095381 | 1.5 | -1038.298782 | -4.421450 | -1042.529095 |
| conformer_8 | -1044.288024 | 0.295898 | 0.228080 | -1044.326099 | -100.0 | -1044.030201 | -1044.095007 | 2.5 | -1038.299435 | -4.422028 | -1042.528445 |
| conformer_3 | -1044.286983 | 0.295902 | 0.228147 | -1044.325895 | -102.2 | -1044.029993 | -1044.094736 | 3.2 | -1038.299180 | -4.421984 | -1042.528917 |
| conformer_50 | -1044.282861 | 0.296094 | 0.228224 | -1044.325932 | -113.1 | -1044.029838 | -1044.094696 | 3.3 | -1038.295336 | -4.420972 | -1042.528143 |
| conformer_2 | -1044.285791 | 0.295889 | 0.227401 | -1044.324652 | -102.0 | -1044.028763 | -1044.094239 | 4.5 | -1038.299238 | -4.420423 | -1042.528109 |
| conformer_13 | -1044.285101 | 0.295803 | 0.227413 | -1044.324599 | -103.7 | -1044.028796 | -1044.094174 | 4.7 | -1038.298091 | -4.420800 | -1042.527964 |
| conformer_39 | -1044.286053 | 0.295855 | 0.228147 | -1044.325305 | -103.1 | -1044.029450 | -1044.094146 | 4.8 | -1038.298792 | -4.421898 | -1042.528783 |
| conformer_4 | -1044.289178 | 0.295949 | 0.228119 | -1044.325097 | -94.3 | -1044.029148 | -1044.093966 | 5.3 | -1038.301549 | -4.421633 | -1042.527970 |
| conformer_32 | -1044.285039 | 0.295762 | 0.226830 | -1044.323658 | -101.4 | -1044.027896 | -1044.093816 | 5.6 | -1038.299038 | -4.420248 | -1042.528063 |
| conformer_27 | -1044.285027 | 0.296074 | 0.228521 | -1044.325052 | -105.1 | -1044.028978 | -1044.093519 | 6.4 | -1038.297539 | -4.421358 | -1042.527389 |
| conformer_30 | -1044.284010 | 0.296029 | 0.226267 | -1044.322702 | -101.6 | -1044.026673 | -1044.093423 | 6.7 | -1038.300277 | -4.418449 | -1042.528140 |
| conformer_52 | -1044.279615 | 0.296157 | 0.227057 | -1044.323191 | -114.4 | -1044.027034 | -1044.093122 | 7.5 | -1038.294545 | -4.419263 | -1042.527314 |
| conformer_7 | -1044.285502 | 0.296055 | 0.227075 | -1044.323069 | -98.6 | -1044.027014 | -1044.092982 | 7.8 | -1038.300608 | -4.419760 | -1042.527848 |
| conformer_69 | -1044.276033 | 0.295769 | 0.226291 | -1044.322265 | -121.4 | -1044.026496 | -1044.092962 | 7.9 | -1038.288550 | -4.421004 | -1042.526483 |
| conformer_48 | -1044.280776 | 0.296031 | 0.226940 | -1044.322559 | -109.7 | -1044.026528 | -1044.092607 | 8.8 | -1038.295687 | -4.419698 | -1042.527217 |
| conformer_5 | -1044.285191 | 0.295996 | 0.226965 | -1044.322434 | -97.8 | -1044.026438 | -1044.092457 | 9.2 | -1038.301504 | -4.418753 | -1042.527523 |
| conformer_67 | -1044.279616 | 0.295879 | 0.227723 | -1044.323045 | -114.0 | -1044.027166 | -1044.092310 | 9.6 | -1038.293521 | -4.420499 | -1042.526714 |
| conformer_25 | -1044.282933 | 0.295833 | 0.226517 | -1044.321768 | -102.0 | -1044.025935 | -1044.092239 | 9.8 | -1038.299507 | -4.418696 | -1042.527510 |
| conformer_43 | -1044.282736 | 0.295771 | 0.226344 | -1044.321248 | -101.1 | -1044.025477 | -1044.091892 | 10.7 | -1038.299796 | -4.418424 | -1042.527376 |
| conformer_64 | -1044.278946 | 0.295997 | 0.228219 | -1044.323097 | -115.9 | -1044.027100 | -1044.091866 | 10.8 | -1038.290800 | -4.422095 | -1042.525815 |
| conformer_56 | -1044.278311 | 0.296204 | 0.227288 | -1044.321998 | -114.7 | -1044.025794 | -1044.091698 | 11.2 | -1038.293212 | -4.419005 | -1042.525604 |
| conformer_23 | -1044.283790 | 0.295727 | 0.227808 | -1044.322087 | -100.5 | -1044.026360 | -1044.091267 | 12.3 | -1038.294620 | -4.422684 | -1042.524781 |
| conformer_31 | -1044.281717 | 0.295910 | 0.228740 | -1044.322522 | -107.1 | -1044.026612 | -1044.090770 | 13.6 | -1038.293874 | -4.421270 | -1042.524197 |
| conformer_75 | -1044.279961 | 0.295983 | 0.228019 | -1044.321719 | -109.6 | -1044.025736 | -1044.090688 | 13.9 | -1038.288639 | -4.424518 | -1042.523884 |
| conformer_55 | -1044.280926 | 0.295979 | 0.227692 | -1044.321349 | -106.1 | -1044.025370 | -1044.090645 | 14.0 | -1038.290173 | -4.423749 | -1042.523641 |

| | | | | | | | | | | | |
|---|---|---|---|---|---|---|---|---|---|---|---|
| conformer_36 | -1044.279625 | 0.295875 | 0.228687 | -1044.322329 | -112.1 | -1044.026454 | -1044.090630 | 14.0 | -1038.289953 | -4.422599 | -1042.523557 |
| conformer_44 | -1044.282512 | 0.296014 | 0.229081 | -1044.322622 | -105.3 | -1044.026608 | -1044.090529 | 14.3 | -1038.294799 | -4.421356 | -1042.524171 |
| conformer_16 | -1044.284586 | 0.295784 | 0.228230 | -1044.321705 | -97.5 | -1044.025921 | -1044.090463 | 14.5 | -1038.295924 | -4.421937 | -1042.523739 |
| conformer_57 | -1044.279830 | 0.296000 | 0.226609 | -1044.320036 | -105.6 | -1044.024036 | -1044.090415 | 14.6 | -1038.295579 | -4.418803 | -1042.524967 |
| conformer_74 | -1044.277380 | 0.295844 | 0.226514 | -1044.319937 | -111.7 | -1044.024093 | -1044.090411 | 14.6 | -1038.293567 | -4.418277 | -1042.524875 |
| conformer_63 | -1044.278117 | 0.296023 | 0.227148 | -1044.320561 | -111.4 | -1044.024538 | -1044.090401 | 14.6 | -1038.292839 | -4.419506 | -1042.524629 |
| conformer_41 | -1044.281395 | 0.296011 | 0.227981 | -1044.321143 | -104.4 | -1044.025132 | -1044.090150 | 15.3 | -1038.296140 | -4.418874 | -1042.523769 |
| conformer_83 | -1044.274384 | 0.295879 | 0.226920 | -1044.320065 | -119.9 | -1044.024186 | -1044.090133 | 15.3 | -1038.287995 | -4.420166 | -1042.523910 |
| conformer_21 | -1044.282661 | 0.295855 | 0.228317 | -1044.321249 | -101.3 | -1044.025394 | -1044.089920 | 15.9 | -1038.294298 | -4.421810 | -1042.523367 |
| conformer_28 | -1044.281177 | 0.295632 | 0.227490 | -1044.320346 | -102.8 | -1044.024714 | -1044.089844 | 16.1 | -1038.293947 | -4.420943 | -1042.523556 |
| conformer_1 | -1044.284929 | 0.296188 | 0.227700 | -1044.320517 | -93.4 | -1044.024329 | -1044.089805 | 16.2 | -1038.300654 | -4.419054 | -1042.524584 |
| conformer_12 | -1044.281871 | 0.295741 | 0.227670 | -1044.320353 | -101.0 | -1044.024612 | -1044.089671 | 16.5 | -1038.295070 | -4.420585 | -1042.523455 |
| conformer_9 | -1044.283707 | 0.295908 | 0.228902 | -1044.321510 | -99.3 | -1044.025602 | -1044.089596 | 16.7 | -1038.295784 | -4.421910 | -1042.523582 |
| conformer_17 | -1044.285449 | 0.295855 | 0.228305 | -1044.320788 | -92.8 | -1044.024933 | -1044.089471 | 17.1 | -1038.297726 | -4.421584 | -1042.523331 |
| conformer_49 | -1044.280727 | 0.296024 | 0.226446 | -1044.318855 | -100.1 | -1044.022831 | -1044.089397 | 17.3 | -1038.297021 | -4.418605 | -1042.524296 |
| conformer_46 | -1044.282391 | 0.295777 | 0.228521 | -1044.320854 | -101.0 | -1044.025077 | -1044.089321 | 17.5 | -1038.295167 | -4.421733 | -1042.523830 |
| conformer_45 | -1044.281723 | 0.295984 | 0.227863 | -1044.320075 | -100.7 | -1044.024091 | -1044.089200 | 17.8 | -1038.297330 | -4.418416 | -1042.523223 |
| conformer_62 | -1044.281620 | 0.295991 | 0.228018 | -1044.320190 | -101.3 | -1044.024199 | -1044.089160 | 17.9 | -1038.291847 | -4.423332 | -1042.522720 |
| conformer_34 | -1044.281190 | 0.295903 | 0.228681 | -1044.320695 | -103.7 | -1044.024792 | -1044.089002 | 18.3 | -1038.293441 | -4.421494 | -1042.522746 |
| conformer_71 | -1044.275855 | 0.295945 | 0.226818 | -1044.318809 | -112.8 | -1044.022864 | -1044.088979 | 18.3 | -1038.290582 | -4.419329 | -1042.523036 |
| conformer_40 | -1044.281101 | 0.295661 | 0.227412 | -1044.319288 | -100.3 | -1044.023627 | -1044.088864 | 18.7 | -1038.294872 | -4.420342 | -1042.522977 |
| conformer_35 | -1044.280609 | 0.296014 | 0.227025 | -1044.318423 | -99.3 | -1044.022409 | -1044.088386 | 19.9 | -1038.296692 | -4.418229 | -1042.522698 |
| conformer_59 | -1044.277118 | 0.295859 | 0.226821 | -1044.318203 | -107.9 | -1044.022344 | -1044.088370 | 19.9 | -1038.291880 | -4.419745 | -1042.522878 |
| conformer_68 | -1044.277671 | 0.296150 | 0.227260 | -1044.318271 | -106.6 | -1044.022121 | -1044.087999 | 20.9 | -1038.290491 | -4.421604 | -1042.522422 |
| conformer_19 | -1044.281503 | 0.295911 | 0.227434 | -1044.318378 | -96.8 | -1044.022467 | -1044.087932 | 21.1 | -1038.296390 | -4.419825 | -1042.522644 |
| conformer_18 | -1044.281269 | 0.295910 | 0.227507 | -1044.318039 | -96.5 | -1044.022129 | -1044.087520 | 22.2 | -1038.297279 | -4.418911 | -1042.522441 |
| conformer_65 | -1044.277385 | 0.296111 | 0.227330 | -1044.317605 | -105.6 | -1044.021494 | -1044.087263 | 22.9 | -1038.291398 | -4.420595 | -1042.521871 |
| conformer_33 | -1044.279292 | 0.295762 | 0.227247 | -1044.317378 | -100.0 | -1044.021616 | -1044.087119 | 23.2 | -1038.295694 | -4.418730 | -1042.522251 |
| conformer_76 | -1044.274624 | 0.296004 | 0.227561 | -1044.317599 | -112.8 | -1044.021595 | -1044.087026 | 23.5 | -1038.289321 | -4.419124 | -1042.520847 |
| conformer_51 | -1044.279062 | 0.295734 | 0.227221 | -1044.316818 | -99.1 | -1044.021084 | -1044.086585 | 24.6 | -1038.295909 | -4.418455 | -1042.521886 |
| conformer_73 | -1044.276293 | 0.295808 | 0.226978 | -1044.315736 | -103.6 | -1044.019928 | -1044.085746 | 26.8 | -1038.291848 | -4.418916 | -1042.520217 |
| conformer_81 | -1044.274525 | 0.295877 | 0.227565 | -1044.316082 | -109.1 | -1044.020205 | -1044.085505 | 27.5 | -1038.288994 | -4.419604 | -1042.519578 |
| conformer_6 | -1044.281063 | 0.296059 | 0.228184 | -1044.316137 | -92.1 | -1044.020078 | -1044.084941 | 29.0 | -1038.296553 | -4.419132 | -1042.519563 |
| conformer_58 | -1044.277236 | 0.295843 | 0.226908 | -1044.314691 | -98.3 | -1044.018848 | -1044.084771 | 29.4 | -1038.293337 | -4.418721 | -1042.519593 |

[a]: Single-point calculation in aqueous phase with SMD model.

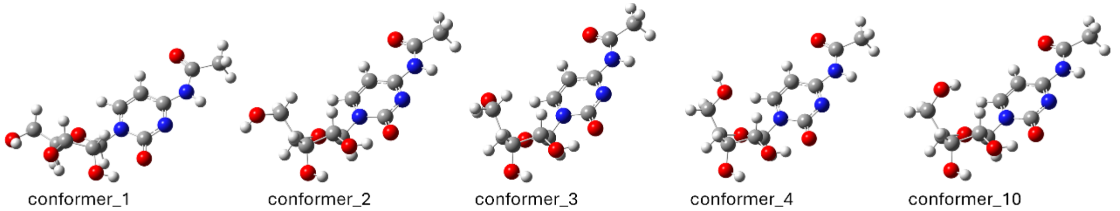

**Figure S29.** B3LYP-D3/def2-TZVPP optimized geometries of conformers for 4-acetyl-5-methylcytidine (**1rb4ac5mC**).

**Table S27.** Conformers of gas-phase optimized 4-acetyl-5-methylcytidine (**1rb4ac5mC**) at the B3LYP-D3/def2-TZVPP level of theory followed by aqueous phase single-point calculation. The columns display total energy without zero-point correction ($E_{Tot}$), thermal correction to enthalpy (δH), Gibbs free energy (δG), total energy without zero-point correction ($E_{Tot,W}$), Gibbs free energy ($G_{298,W}$) in water (W), total single-point energy ($E_{CBS}$) calculated at DLPNO-CCSD(T)/CBS level of theory, and their corresponding free energy $G_{CBS,W}$. $G_{298,W}$ and $G_{CBS,W}$ have been corrected to the standard state of 1 mol/L by addition of +7.908 kJ/mol. $\Delta G_{Solv}$ represents the Gibbs free energy of solvation. The data are arranged in the ascending numeric order of $E_{Tot,W}$. $\Delta G_{298,W}$ represents the respective energy difference to the lowest structure. Only conformers within the 24 kJ/mol (6 kcal/mol) energy window above the lowest in CREST are included in initial conformer sampling. Duplicates of the same structure are excluded. The overall optimum is marked bold.

| 1rb4ac5mC No. | B3LYP-D3/def2-TZVPP $E_{Tot}$ (Hartree) | δH (Hartree) | δG (Hartree) | SMD(H2O)/B3LYP-D3/def2-TZVPP[a] $E_{Tot,W}$ (Hartree) | $\Delta G_{Solv}$ (kJ/mol) | $H_{298,W}$ (Hartree) | $G_{298,W}$ (Hartree) | $\Delta G_{298,W}$ (kJ/mol) | DLPNO-CCSD(T)/CBS $E_{CBS,HF}$ (Hartree) | $E_{CBS,C}$ (Hartree) | $G_{CBS,W}$ (Hartree) |
|---|---|---|---|---|---|---|---|---|---|---|---|
| conformer_53 | -1083.617803 | 0.325507 | 0.255524 | -1083.657379 | -103.9 | -1083.331872 | -1083.398843 | 1.7 | -1077.345945 | -4.632736 | -1081.759720 |
| conformer_25 | -1083.614946 | 0.325331 | 0.254986 | -1083.657186 | -110.9 | -1083.331855 | -1083.399188 | 0.8 | -1077.341136 | -4.633917 | -1081.759295 |
| conformer_21 | -1083.616949 | 0.325339 | 0.254918 | -1083.657146 | -105.5 | -1083.331807 | -1083.399216 | 0.7 | -1077.344983 | -4.632600 | -1081.759851 |
| **conformer_14** | **-1083.619047** | **0.325202** | **0.254281** | **-1083.656771** | **-99.0** | **-1083.331569** | **-1083.399478** | **0.0** | **-1077.345273** | **-4.634519** | **-1081.760223** |
| conformer_5 | -1083.619737 | 0.325188 | 0.254263 | -1083.656453 | -96.4 | -1083.331265 | -1083.399178 | 0.8 | -1077.346664 | -4.633595 | -1081.759700 |
| conformer_1 | -1083.619047 | 0.325264 | 0.254834 | -1083.656228 | -97.6 | -1083.330964 | -1083.398382 | 2.9 | -1077.346626 | -4.633716 | -1081.759677 |
| conformer_58 | -1083.614213 | 0.325434 | 0.255023 | -1083.656189 | -110.2 | -1083.330755 | -1083.398154 | 3.5 | -1077.342220 | -4.632491 | -1081.758653 |
| conformer_13 | -1083.617995 | 0.325152 | 0.254064 | -1083.655953 | -99.7 | -1083.330801 | -1083.398877 | 1.6 | -1077.345182 | -4.633668 | -1081.759732 |
| conformer_27 | -1083.616489 | 0.325423 | 0.254148 | -1083.65579 | -103.2 | -1083.330367 | -1083.398630 | 2.2 | -1077.346750 | -4.630649 | -1081.759540 |
| conformer_35 | -1083.617554 | 0.325163 | 0.254422 | -1083.655607 | -99.9 | -1083.330444 | -1083.398173 | 3.4 | -1077.345788 | -4.633584 | -1081.759991 |
| conformer_8 | -1083.620377 | 0.325269 | 0.254438 | -1083.655428 | -92.0 | -1083.330159 | -1083.397978 | 3.9 | -1077.348300 | -4.633155 | -1081.759056 |
| conformer_23 | -1083.616473 | 0.325351 | 0.255016 | -1083.65537 | -102.1 | -1083.330019 | -1083.397342 | 5.6 | -1077.344549 | -4.632869 | -1081.758288 |
| conformer_3 | -1083.617021 | 0.325187 | 0.253934 | -1083.654968 | -99.6 | -1083.329781 | -1083.398022 | 3.8 | -1077.345905 | -4.632155 | -1081.759060 |
| conformer_18 | -1083.616387 | 0.325081 | 0.253775 | -1083.654934 | -101.2 | -1083.329853 | -1083.398147 | 3.5 | -1077.344839 | -4.632551 | -1081.759151 |
| conformer_28 | -1083.616636 | 0.32538 | 0.253926 | -1083.654649 | -99.8 | -1083.329269 | -1083.397711 | 4.6 | -1077.347815 | -4.630130 | -1081.759020 |
| conformer_71 | -1083.612513 | 0.325183 | 0.254106 | -1083.654538 | -110.3 | -1083.329355 | -1083.397420 | 5.4 | -1077.339001 | -4.634320 | -1081.758229 |
| conformer_54 | -1083.614897 | 0.325009 | 0.253361 | -1083.654536 | -104.1 | -1083.329527 | -1083.398163 | 3.5 | -1077.339211 | -4.636432 | -1081.758908 |
| conformer_42 | -1083.615405 | 0.324971 | 0.252864 | -1083.65438 | -102.3 | -1083.329409 | -1083.398504 | 2.6 | -1077.340184 | -4.636260 | -1081.759542 |
| conformer_44 | -1083.615656 | 0.325054 | 0.253693 | -1083.654352 | -101.6 | -1083.329298 | -1083.397647 | 4.8 | -1077.340490 | -4.635716 | -1081.758196 |
| conformer_49 | -1083.614111 | 0.325037 | 0.253453 | -1083.654175 | -105.2 | -1083.329138 | -1083.397710 | 4.6 | -1077.339225 | -4.635768 | -1081.758591 |
| conformer_34 | -1083.614665 | 0.325128 | 0.25428 | -1083.653995 | -103.3 | -1083.328867 | -1083.396703 | 7.3 | -1077.340207 | -4.635810 | -1081.758054 |
| conformer_37 | -1083.614419 | 0.325131 | 0.254111 | -1083.653971 | -103.8 | -1083.328840 | -1083.396848 | 6.9 | -1077.340321 | -4.635657 | -1081.758407 |
| conformer_31 | -1083.616177 | 0.325094 | 0.253612 | -1083.65388 | -99.0 | -1083.328786 | -1083.397256 | 5.8 | -1077.345645 | -4.631895 | -1081.758619 |
| conformer_48 | -1083.615062 | 0.32497 | 0.253503 | -1083.653869 | -101.9 | -1083.328899 | -1083.397354 | 5.6 | -1077.340317 | -4.635572 | -1081.758180 |
| conformer_87 | -1083.606847 | 0.325094 | 0.253339 | -1083.65384 | -123.4 | -1083.328746 | -1083.397489 | 5.2 | -1077.335282 | -4.631553 | -1081.757478 |
| conformer_51 | -1083.613750 | 0.324973 | 0.25249 | -1083.653511 | -104.4 | -1083.328538 | -1083.398009 | 3.9 | -1077.339673 | -4.635151 | -1081.759083 |
| conformer_91 | -1083.613744 | 0.325082 | 0.25411 | -1083.653385 | -104.1 | -1083.328303 | -1083.396263 | 8.4 | -1077.340144 | -4.635663 | -1081.758326 |
| conformer_63 | -1083.610894 | 0.325368 | 0.253003 | -1083.653275 | -111.3 | -1083.327907 | -1083.397260 | 5.8 | -1077.341584 | -4.630642 | -1081.758592 |
| conformer_50 | -1083.616434 | 0.325094 | 0.253573 | -1083.653269 | -96.7 | -1083.328175 | -1083.396684 | 7.3 | -1077.342253 | -4.635261 | -1081.757765 |
| conformer_78 | -1083.61115 | 0.325263 | 0.254639 | -1083.653242 | -110.5 | -1083.327979 | -1083.395591 | 10.2 | -1077.340617 | -4.631956 | -1081.757014 |

| | | | | | | | | | | | |
|---|---|---|---|---|---|---|---|---|---|---|---|
| conformer_76 | -1083.610366 | 0.325272 | 0.254771 | -1083.653214 | -112.5 | -1083.327942 | -1083.395431 | 10.6 | -1077.337778 | -4.633614 | -1081.756457 |
| conformer_89 | -1083.607966 | 0.325119 | 0.253304 | -1083.653177 | -118.7 | -1083.328058 | -1083.396861 | 6.9 | -1077.337483 | -4.631055 | -1081.757433 |
| conformer_40 | -1083.616365 | 0.325069 | 0.253969 | -1083.653028 | -96.3 | -1083.327959 | -1083.396047 | 9.0 | -1077.342335 | -4.635664 | -1081.757680 |
| conformer_77 | -1083.612367 | 0.325205 | 0.25404 | -1083.652968 | -106.6 | -1083.327763 | -1083.395916 | 9.4 | -1077.338871 | -4.634661 | -1081.757081 |
| conformer_39 | -1083.612949 | 0.324974 | 0.2529 | -1083.65281 | -104.7 | -1083.327836 | -1083.396898 | 6.8 | -1077.339948 | -4.634062 | -1081.757959 |
| conformer_61 | -1083.612226 | 0.324862 | 0.252799 | -1083.652727 | -106.3 | -1083.327865 | -1083.396916 | 6.7 | -1077.338756 | -4.634470 | -1081.757916 |
| conformer_11 | -1083.616639 | 0.325298 | 0.253593 | -1083.652719 | -94.7 | -1083.327421 | -1083.396114 | 8.8 | -1077.347199 | -4.631485 | -1081.758159 |
| conformer_56 | -1083.612154 | 0.325252 | 0.253024 | -1083.65268 | -106.4 | -1083.327428 | -1083.396644 | 7.4 | -1077.342768 | -4.631154 | -1081.758413 |
| conformer_24 | -1083.615635 | 0.325337 | 0.252832 | -1083.65266 | -97.2 | -1083.327323 | -1083.396816 | 7.0 | -1077.347508 | -4.629871 | -1081.758560 |
| conformer_12 | -1083.616346 | 0.325318 | 0.253867 | -1083.652513 | -95.0 | -1083.327195 | -1083.395634 | 10.1 | -1077.348111 | -4.630479 | -1081.757877 |
| conformer_55 | -1083.612375 | 0.324867 | 0.252338 | -1083.652493 | -105.3 | -1083.327626 | -1083.397143 | 6.1 | -1077.339316 | -4.634281 | -1081.758364 |
| conformer_38 | -1083.612967 | 0.324972 | 0.252779 | -1083.652471 | -103.7 | -1083.327499 | -1083.396680 | 7.3 | -1077.340315 | -4.634008 | -1081.758036 |
| conformer_86 | -1083.607405 | 0.325039 | 0.252515 | -1083.652208 | -117.6 | -1083.327169 | -1083.396681 | 7.3 | -1077.335346 | -4.632676 | -1081.757298 |
| conformer_66 | -1083.609656 | 0.325385 | 0.253489 | -1083.652077 | -111.4 | -1083.326692 | -1083.395576 | 10.2 | -1077.340330 | -4.630416 | -1081.756665 |
| conformer_22 | -1083.61438 | 0.325099 | 0.253241 | -1083.651817 | -98.3 | -1083.326718 | -1083.395564 | 10.3 | -1077.346562 | -4.630293 | -1081.758039 |
| conformer_84 | -1083.612371 | 0.324917 | 0.252667 | -1083.651703 | -103.3 | -1083.326786 | -1083.396024 | 9.1 | -1077.340241 | -4.633897 | -1081.757791 |
| conformer_43 | -1083.614055 | 0.325069 | 0.253269 | -1083.651249 | -97.7 | -1083.326180 | -1083.394968 | 11.8 | -1077.346691 | -4.630032 | -1081.757636 |
| conformer_47 | -1083.612672 | 0.325236 | 0.253238 | -1083.650796 | -100.1 | -1083.325560 | -1083.394546 | 12.9 | -1077.341459 | -4.633270 | -1081.756603 |
| conformer_70 | -1083.609586 | 0.325271 | 0.253585 | -1083.650547 | -107.5 | -1083.325276 | -1083.393950 | 14.5 | -1077.340045 | -4.630929 | -1081.755338 |
| conformer_2 | -1083.616166 | 0.325452 | 0.254292 | -1083.650376 | -89.8 | -1083.324924 | -1083.393072 | 16.8 | -1077.347413 | -4.630711 | -1081.755030 |
| conformer_65 | -1083.611263 | 0.325214 | 0.253248 | -1083.650308 | -102.5 | -1083.325094 | -1083.394048 | 14.3 | -1077.342768 | -4.630252 | -1081.755805 |
| conformer_85 | -1083.608971 | 0.325147 | 0.253159 | -1083.650267 | -108.4 | -1083.325120 | -1083.394096 | 14.1 | -1077.340827 | -4.629734 | -1081.755687 |
| conformer_46 | -1083.612372 | 0.325246 | 0.253316 | -1083.650229 | -99.4 | -1083.324983 | -1083.393901 | 14.6 | -1077.342576 | -4.632183 | -1081.756287 |
| conformer_75 | -1083.611326 | 0.325182 | 0.252351 | -1083.650208 | -102.1 | -1083.325026 | -1083.394845 | 12.2 | -1077.341245 | -4.632028 | -1081.756792 |
| conformer_45 | -1083.612371 | 0.32509 | 0.252585 | -1083.65014 | -99.2 | -1083.325050 | -1083.394543 | 13.0 | -1077.342299 | -4.632344 | -1081.756815 |
| conformer_72 | -1083.610262 | 0.325014 | 0.252609 | -1083.64955 | -103.1 | -1083.324536 | -1083.393929 | 14.6 | -1077.340816 | -4.632074 | -1081.756556 |
| conformer_73 | -1083.610057 | 0.324988 | 0.252278 | -1083.649406 | -103.3 | -1083.324418 | -1083.394116 | 14.1 | -1077.340212 | -4.632296 | -1081.756567 |
| conformer_57 | -1083.612184 | 0.325234 | 0.252864 | -1083.649192 | -97.2 | -1083.323958 | -1083.393316 | 16.2 | -1077.344240 | -4.630025 | -1081.755397 |
| conformer_32 | -1083.612192 | 0.325328 | 0.253456 | -1083.64817 | -94.5 | -1083.322842 | -1083.391702 | 20.4 | -1077.341817 | -4.632498 | -1081.753826 |
| conformer_33 | -1083.612032 | 0.325324 | 0.253753 | -1083.648074 | -94.6 | -1083.322750 | -1083.391309 | 21.4 | -1077.341437 | -4.632673 | -1081.753387 |

[a]: Single-point calculation in aqueous phase with SMD model.

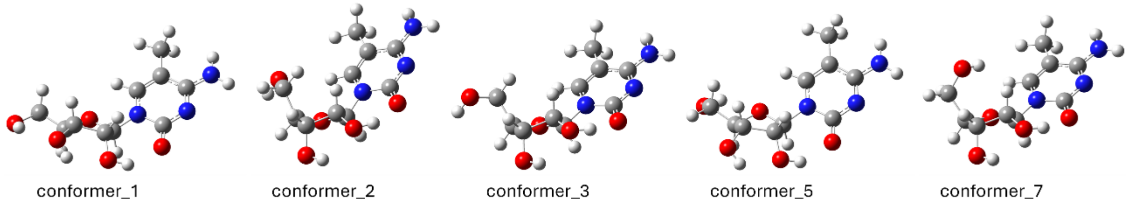

**Figure S30.** B3LYP-D3/def2-TZVPP optimized geometries of conformers for 5-methylcytidine (**1rb5mC**).

**Table S28.** Conformers of gas-phase optimized 5-methylcytidine (**1rb5mC**) at the B3LYP-D3/def2-TZVPP level of theory followed by aqueous phase single-point calculation. The columns display total energy without zero-point correction ($E_{Tot}$), Gibbs free energy ($\delta G$), total energy without zero-point correction ($E_{Tot,W}$), Gibbs free energy ($G_{298,W}$) in water (W), total single-point energy ($E_{CBS}$) calculated at DLPNO-CCSD(T)/CBS level of theory, and their corresponding free energy $G_{CBS}$. $G_{298,W}$ and $G_{CBS}$ have been corrected to the standard state of 1 mol/L by addition of +7.908 kJ/mol. $\Delta G_{Solv}$ represents the Gibbs free energy of solvation. The data are arranged in the ascending numeric order of $E_{Tot,W}$. $\Delta G_{298,W}$ represents the respective energy difference to the lowest structure. Only conformers within the 24 kJ/mol (6 kcal/mol) energy window above the lowest in CREST are included in initial conformer sampling. Duplicates of the same structure are excluded.

| 1rb5mC No. | B3LYP-D3/def2-TZVPP $E_{Tot}$ (Hartree) | $\delta H$ (Hartree) | $\delta G$ (Hartree) | SMD($H_2O$)/B3LYP-D3/def2-TZVPP[a] $E_{Tot,W}$ (Hartree) | $\Delta G_{Solv}$ (kJ/mol) | $H_{298,W}$ (Hartree) | $G_{298,W}$ (Hartree) | $\Delta G_{298,W}$ (kJ/mol) | DLPNO-CCSD(T)/CBS $E_{CBS,HF}$ (Hartree) | $E_{CBS,C}$ (Hartree) | $G_{CBS,W}$ (Hartree) |
|---|---|---|---|---|---|---|---|---|---|---|---|
| conformer_25 | -930.899142 | 0.284342 | 0.222093 | -930.940987 | -109.9 | -930.656645 | -930.715882 | 2.4 | -925.503549 | -3.991700 | -929.311988 |
| conformer_19 | -930.896257 | 0.284151 | 0.221356 | -930.940726 | -116.8 | -930.656575 | -930.716358 | 1.1 | -925.498659 | -3.992882 | -929.311642 |
| conformer_16 | -930.897969 | 0.284165 | 0.221397 | -930.940705 | -112.2 | -930.656540 | -930.716296 | 1.3 | -925.502261 | -3.991533 | -929.312122 |
| **conformer_7** | **-930.900704** | **0.284058** | **0.220222** | **-930.940029** | **-103.2** | **-930.655971** | **-930.716795** | **0** | **-925.503321** | **-3.993280** | **-929.312691** |
| conformer_26 | -930.895709 | 0.284302 | 0.221667 | -930.939653 | -115.4 | -930.655351 | -930.714974 | 4.8 | -925.499965 | -3.991451 | -929.310681 |
| conformer_9 | -930.900665 | 0.284050 | 0.221014 | -930.939451 | -101.8 | -930.655401 | -930.715425 | 3.6 | -925.503962 | -3.992635 | -929.311357 |
| conformer_6 | -930.899716 | 0.284063 | 0.219654 | -930.939327 | -104.0 | -930.655264 | -930.716661 | 0.4 | -925.503283 | -3.992563 | -929.312792 |
| conformer_12 | -930.899716 | 0.284064 | 0.219653 | -930.939327 | -104.0 | -930.655263 | -930.716662 | 0.3 | -925.503283 | -3.992563 | -929.312793 |
| conformer_2 | -930.900051 | 0.284158 | 0.221629 | -930.939046 | -102.4 | -930.654888 | -930.714405 | 6.3 | -925.503986 | -3.992759 | -929.311100 |
| conformer_18 | -930.897849 | 0.284149 | 0.221430 | -930.938762 | -107.4 | -930.654613 | -930.714320 | 6.5 | -925.502191 | -3.991929 | -929.310591 |
| conformer_23 | -930.898516 | 0.284113 | 0.221796 | -930.938392 | -104.7 | -930.654279 | -930.713584 | 8.4 | -925.503094 | -3.992593 | -929.310754 |
| conformer_22 | -930.896801 | 0.284234 | 0.220851 | -930.938325 | -109.0 | -930.654091 | -930.714462 | 6.1 | -925.503766 | -3.989462 | -929.310889 |
| conformer_14 | -930.901004 | 0.284074 | 0.221179 | -930.938157 | -97.5 | -930.654083 | -930.713966 | 7.4 | -925.505389 | -3.992128 | -929.310479 |
| conformer_3 | -930.898445 | 0.284101 | 0.220236 | -930.938120 | -104.2 | -930.654019 | -930.714872 | 5.0 | -925.503633 | -3.991070 | -929.311130 |
| conformer_10 | -930.897966 | 0.283963 | 0.219866 | -930.938033 | -105.2 | -930.654070 | -930.715155 | 4.3 | -925.502707 | -3.991488 | -929.311384 |
| conformer_20 | -930.897475 | 0.283979 | 0.220169 | -930.936961 | -103.7 | -930.652982 | -930.713780 | 7.9 | -925.503256 | -3.990873 | -929.310433 |
| conformer_37 | -930.891700 | 0.284227 | 0.221067 | -930.936579 | -117.8 | -930.652352 | -930.712500 | 11.3 | -925.495350 | -3.992602 | -929.308751 |
| conformer_40 | -930.892134 | 0.284099 | 0.220668 | -930.936497 | -116.5 | -930.652398 | -930.712817 | 10.4 | -925.497907 | -3.991022 | -929.309613 |
| conformer_28 | -930.892092 | 0.284297 | 0.220142 | -930.936163 | -115.7 | -930.651866 | -930.713009 | 9.9 | -925.498907 | -3.989802 | -929.309625 |
| conformer_33 | -930.898264 | 0.284223 | 0.219894 | -930.935967 | -99.0 | -930.651744 | -930.713061 | 9.8 | -925.505007 | -3.990580 | -929.310384 |
| conformer_31 | -930.893267 | 0.284133 | 0.219783 | -930.935741 | -111.5 | -930.651608 | -930.712946 | 10.1 | -925.500212 | -3.990120 | -929.310011 |
| conformer_5 | -930.898021 | 0.284193 | 0.220229 | -930.935579 | -98.6 | -930.651386 | -930.712338 | 11.7 | -925.505966 | -3.989587 | -929.309870 |
| conformer_32 | -930.890803 | 0.284350 | 0.220533 | -930.935272 | -116.8 | -930.650922 | -930.711727 | 13.3 | -925.497698 | -3.989443 | -929.308065 |
| conformer_17 | -930.896689 | 0.284194 | 0.219632 | -930.935217 | -101.2 | -930.651023 | -930.712573 | 11.1 | -925.504838 | -3.989221 | -929.309943 |
| conformer_15 | -930.895877 | 0.284031 | 0.219829 | -930.934989 | -102.7 | -930.650958 | -930.712148 | 12.2 | -925.504178 | -3.989388 | -929.309836 |
| conformer_24 | -930.895365 | 0.283994 | 0.219995 | -930.934402 | -102.5 | -930.650408 | -930.711395 | 14.2 | -925.504174 | -3.989138 | -929.309341 |
| conformer_36 | -930.891978 | 0.284082 | 0.219705 | -930.933419 | -108.8 | -930.649337 | -930.710702 | 16.0 | -925.499799 | -3.989161 | -929.307684 |
| conformer_1 | -930.897735 | 0.284352 | 0.220592 | -930.933259 | -93.3 | -930.648907 | -930.709655 | 18.7 | -925.505334 | -3.989710 | -929.306964 |

[a]: Single-point calculation in aqueous phase with SMD model.

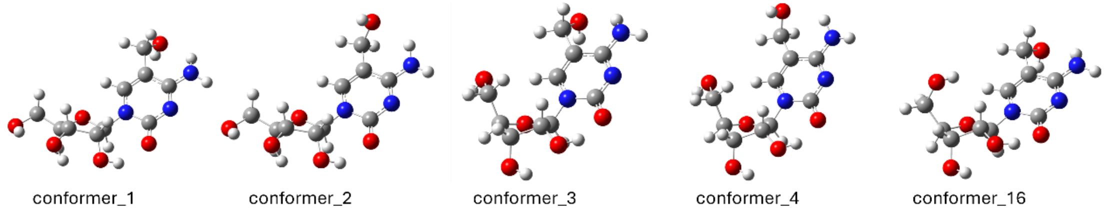

**Figure S31.** B3LYP-D3/def2-TZVPP optimized geometries of conformers for 5-hydroxymethylcytidine (**1rb5hmC**).

**Table S29.** Conformers of gas-phase optimized 5-hydroxymethylcytidine (**1rb5hmC**) at the B3LYP-D3/def2-TZVPP level of theory followed by aqueous phase single-point calculation. The columns display total energy without zero-point correction ($E_{Tot}$), thermal correction to enthalpy (δH), Gibbs free energy (δG), total energy without zero-point correction ($E_{Tot,W}$), Gibbs free energy ($G_{298,W}$) in water (W), total single-point energy ($E_{CBS}$) calculated at DLPNO-CCSD(T)/CBS level of theory, and their corresponding free energy $G_{CBS,W}$. $G_{298,W}$ and $G_{CBS,W}$ have been corrected to the standard state of 1 mol/L by addition of +7.908 kJ/mol. $\Delta G_{Solv}$ represents the Gibbs free energy of solvation. The data are arranged in the ascending numeric order of $E_{Tot,W}$. $\Delta G_{298,W}$ represents the respective energy difference to the lowest structure. Only conformers within the 24 kJ/mol (6 kcal/mol) energy window above the lowest in CREST are included in initial conformer sampling. Duplicates of the same structure are excluded. The overall optimum is marked bold.

| 1rb5hmC | B3LYP-D3/def2-TZVPP | | | SMD(H2O)/B3LYP-D3/def2-TZVPP[a] | | | | | DLPNO-CCSD(T)/CBS | | |
|---|---|---|---|---|---|---|---|---|---|---|---|
| No. | $E_{Tot}$ (Hartree) | δH (Hartree) | δG (Hartree) | $E_{Tot,W}$ (Hartree) | $\Delta G_{Solv}$ (kJ/mol) | $H_{298,W}$ (Hartree) | $G_{298,W}$ (Hartree) | $\Delta G_{298,W}$ (kJ/mol) | $E_{CBS,HF}$ (Hartree) | $E_{CBS,C}$ (Hartree) | $G_{CBS,W}$ (Hartree) |
| conformer_66 | -1006.150750 | 0.290989 | 0.227242 | -1006.196126 | -119.1 | -1005.905137 | -1005.965872 | 1.9 | -1000.395757 | -4.263029 | -1004.473907 |
| conformer_32 | -1006.149729 | 0.290785 | 0.226664 | -1006.195832 | -121.0 | -1005.905047 | -1005.966156 | 1.1 | -1000.394668 | -4.262943 | -1004.474039 |
| conformer_35 | -1006.149627 | 0.290857 | 0.226935 | -1006.195810 | -121.3 | -1005.904953 | -1005.965863 | 1.9 | -1000.394465 | -4.262883 | -1004.473584 |
| conformer_41 | -1006.147733 | 0.290809 | 0.226815 | -1006.195792 | -126.2 | -1005.904983 | -1005.965965 | 1.6 | -1000.390686 | -4.264211 | -1004.473129 |
| conformer_15 | -1006.152108 | 0.29068 | 0.225621 | -1006.195185 | -113.1 | -1005.904505 | -1005.966552 | 0.1 | -1000.395307 | -4.264741 | -1004.474492 |
| **conformer_16** | **-1006.152146** | **0.290599** | **0.225488** | **-1006.19509** | **-112.7** | **-1005.904491** | **-1005.966590** | **0** | **-1000.395800** | **-4.264276** | **-1004.474520** |
| conformer_6 | -1006.152729 | 0.29073 | 0.226479 | -1006.194924 | -110.8 | -1005.904194 | -1005.965433 | 3.0 | -1000.396734 | -4.263954 | -1004.473391 |
| conformer_8 | -1006.152830 | 0.290706 | 0.226309 | -1006.194889 | -110.4 | -1005.904183 | -1005.965568 | 2.7 | -1000.396744 | -4.263911 | -1004.473392 |
| conformer_120 | -1006.147245 | 0.290965 | 0.226801 | -1006.194846 | -125.0 | -1005.903881 | -1005.965033 | 4.1 | -1000.392090 | -4.262744 | -1004.472622 |
| conformer_129 | -1006.147131 | 0.290917 | 0.226554 | -1006.194759 | -125.0 | -1005.903842 | -1005.965193 | 3.7 | -1000.391997 | -4.262770 | -1004.472830 |
| conformer_3 | -1006.152228 | 0.290904 | 0.227094 | -1006.194739 | -111.6 | -1005.903835 | -1005.964633 | 5.1 | -1000.396750 | -4.263974 | -1004.473129 |
| conformer_12 | -1006.151418 | 0.290756 | 0.226071 | -1006.19473 | -113.7 | -1005.903974 | -1005.965647 | 2.5 | -1000.395364 | -4.264022 | -1004.473615 |
| conformer_150 | -1006.148678 | 0.290776 | 0.226803 | -1006.194614 | -120.6 | -1005.903838 | -1005.964799 | 4.7 | -1000.394310 | -4.262868 | -1004.473299 |
| conformer_4 | -1006.152013 | 0.290822 | 0.22675 | -1006.194532 | -111.6 | -1005.903710 | -1005.964770 | 4.8 | -1000.396793 | -4.263937 | -1004.473486 |
| conformer_119 | -1006.145585 | 0.290655 | 0.226480 | -1006.194335 | -128.0 | -1005.903680 | -1005.964843 | 4.6 | -1000.389110 | -4.264206 | -1004.472575 |
| conformer_14 | -1006.151043 | 0.290683 | 0.225666 | -1006.194287 | -113.5 | -1005.903604 | -1005.965609 | 2.6 | -1000.395342 | -4.263811 | -1004.473719 |
| conformer_110 | -1006.147355 | 0.290645 | 0.226418 | -1006.194285 | -123.2 | -1005.903640 | -1005.964855 | 4.6 | -1000.392826 | -4.262831 | -1004.473157 |
| conformer_116 | -1006.145722 | 0.290664 | 0.226522 | -1006.194234 | -127.4 | -1005.903570 | -1005.964700 | 5.0 | -1000.389262 | -4.264030 | -1004.472271 |
| conformer_106 | -1006.147477 | 0.290706 | 0.226633 | -1006.194226 | -122.7 | -1005.903520 | -1005.964581 | 5.3 | -1000.392906 | -4.262661 | -1004.472672 |
| conformer_65 | -1006.150746 | 0.290781 | 0.226792 | -1006.194025 | -113.6 | -1005.903244 | -1005.964221 | 6.2 | -1000.395932 | -4.263816 | -1004.473223 |
| conformer_38 | -1006.149387 | 0.290791 | 0.226522 | -1006.193962 | -117.0 | -1005.903171 | -1005.964428 | 5.7 | -1000.394375 | -4.263158 | -1004.472574 |
| conformer_60 | -1006.149229 | 0.290920 | 0.225951 | -1006.193954 | -117.4 | -1005.903034 | -1005.964991 | 4.2 | -1000.396645 | -4.260846 | -1004.473253 |
| conformer_37 | -1006.149434 | 0.290813 | 0.226527 | -1006.193939 | -116.8 | -1005.903126 | -1005.964400 | 5.7 | -1000.394428 | -4.263173 | -1004.472568 |
| conformer_83 | -1006.150142 | 0.290665 | 0.226384 | -1006.193824 | -114.7 | -1005.903159 | -1005.964428 | 5.7 | -1000.395556 | -4.263708 | -1004.473550 |
| conformer_73 | -1006.148862 | 0.290927 | 0.225796 | -1006.193816 | -118.0 | -1005.902889 | -1005.965008 | 4.2 | -1000.396443 | -4.260738 | -1004.473327 |
| conformer_48 | -1006.150457 | 0.290628 | 0.225809 | -1006.193757 | -113.7 | -1005.903129 | -1005.964936 | 4.3 | -1000.394223 | -4.264597 | -1004.473299 |
| conformer_20 | -1006.153253 | 0.290723 | 0.226268 | -1006.193684 | -106.2 | -1005.902961 | -1005.964404 | 5.7 | -1000.398203 | -4.263435 | -1004.472789 |
| conformer_52 | -1006.150112 | 0.290470 | 0.225343 | -1006.193599 | -114.2 | -1005.903129 | -1005.965244 | 3.5 | -1000.394245 | -4.264185 | -1004.473561 |
| conformer_23 | -1006.152906 | 0.290744 | 0.226395 | -1006.193572 | -106.8 | -1005.902828 | -1005.964165 | 6.4 | -1000.398003 | -4.263359 | -1004.472621 |
| conformer_154 | -1006.141637 | 0.290611 | 0.225783 | -1006.193451 | -136.0 | -1005.902840 | -1005.964656 | 5.1 | -1000.385421 | -4.263586 | -1004.472025 |

| | | | | | | | | | | | |
|---|---|---|---|---|---|---|---|---|---|---|---|
| conformer_136 | -1006.141902 | 0.290499 | 0.225406 | -1006.193423 | -135.3 | -1005.902924 | -1005.965005 | 4.2 | -1000.385722 | -4.263567 | -1004.472392 |
| conformer_5 | -1006.150140 | 0.290602 | 0.225213 | -1006.193397 | -113.6 | -1005.902795 | -1005.965172 | 3.7 | -1000.396023 | -4.262356 | -1004.473410 |
| conformer_17 | -1006.14964 | 0.290512 | 0.225221 | -1006.193384 | -114.9 | -1005.902872 | -1005.965151 | 3.8 | -1000.395078 | -4.262733 | -1004.473323 |
| conformer_138 | -1006.143359 | 0.290577 | 0.225656 | -1006.193382 | -131.3 | -1005.902805 | -1005.964714 | 4.9 | -1000.389069 | -4.262239 | -1004.472663 |
| conformer_50 | -1006.150668 | 0.290602 | 0.226217 | -1006.193350 | -112.1 | -1005.902748 | -1005.964121 | 6.5 | -1000.395069 | -4.263816 | -1004.472338 |
| conformer_128 | -1006.143504 | 0.290457 | 0.225263 | -1006.193267 | -130.7 | -1005.902810 | -1005.964992 | 4.2 | -1000.389258 | -4.262215 | -1004.472960 |
| conformer_54 | -1006.150190 | 0.290586 | 0.226099 | -1006.193211 | -113.0 | -1005.902625 | -1005.964100 | 6.5 | -1000.394734 | -4.263744 | -1004.472388 |
| conformer_42 | -1006.149577 | 0.290679 | 0.226000 | -1006.193206 | -114.5 | -1005.902527 | -1005.964194 | 6.3 | -1000.393988 | -4.263979 | -1004.472585 |
| conformer_167 | -1006.142037 | 0.290621 | 0.225846 | -1006.193152 | -134.2 | -1005.902531 | -1005.964294 | 6.0 | -1000.385494 | -4.263989 | -1004.471741 |
| conformer_72 | -1006.146500 | 0.290420 | 0.224806 | -1006.193133 | -122.4 | -1005.902713 | -1005.965315 | 3.3 | -1000.390556 | -4.264000 | -1004.473371 |
| conformer_28 | -1006.149858 | 0.290713 | 0.226555 | -1006.193116 | -113.6 | -1005.902403 | -1005.963549 | 8.0 | -1000.395132 | -4.263804 | -1004.472627 |
| conformer_27 | -1006.150095 | 0.290780 | 0.226904 | -1006.193108 | -112.9 | -1005.902328 | -1005.963192 | 8.9 | -1000.395094 | -4.263848 | -1004.472039 |
| conformer_7 | -1006.149945 | 0.290571 | 0.225165 | -1006.193107 | -113.3 | -1005.902536 | -1005.964930 | 4.4 | -1000.395935 | -4.262311 | -1004.473232 |
| conformer_145 | -1006.143865 | 0.290651 | 0.225901 | -1006.193106 | -129.3 | -1005.902455 | -1005.964193 | 6.3 | -1000.389192 | -4.262684 | -1004.472204 |
| conformer_177 | -1006.141754 | 0.290469 | 0.225035 | -1006.193103 | -134.8 | -1005.902634 | -1005.965056 | 4.0 | -1000.385241 | -4.264023 | -1004.472567 |
| conformer_22 | -1006.149390 | 0.290470 | 0.225257 | -1006.192956 | -114.4 | -1005.902486 | -1005.964687 | 5.0 | -1000.394955 | -4.262674 | -1004.472926 |
| conformer_82 | -1006.145802 | 0.290327 | 0.223927 | -1006.192941 | -123.8 | -1005.902614 | -1005.966002 | 1.5 | -1000.389246 | -4.264528 | -1004.473973 |
| conformer_44 | -1006.149199 | 0.290579 | 0.225500 | -1006.192896 | -114.7 | -1005.902317 | -1005.964384 | 5.8 | -1000.394138 | -4.263583 | -1004.472905 |
| conformer_89 | -1006.146664 | 0.290437 | 0.224705 | -1006.192796 | -121.1 | -1005.902359 | -1005.965079 | 4.0 | -1000.390139 | -4.264696 | -1004.473250 |
| conformer_163 | -1006.149268 | 0.290852 | 0.225577 | -1006.192741 | -114.1 | -1005.901889 | -1005.964152 | 6.4 | -1000.397626 | -4.260349 | -1004.472859 |
| conformer_140 | -1006.147005 | 0.290790 | 0.225601 | -1006.192700 | -120.0 | -1005.901910 | -1005.964087 | 6.6 | -1000.394990 | -4.260716 | -1004.472788 |
| conformer_144 | -1006.148628 | 0.290573 | 0.226205 | -1006.192546 | -115.3 | -1005.901973 | -1005.963329 | 8.6 | -1000.394524 | -4.263750 | -1004.472975 |
| conformer_26 | -1006.147843 | 0.290702 | 0.227132 | -1006.192538 | -117.3 | -1005.901836 | -1005.962394 | 11.0 | -1000.390878 | -4.265602 | -1004.471031 |
| conformer_105 | -1006.146198 | 0.290454 | 0.224922 | -1006.192500 | -121.6 | -1005.902046 | -1005.964566 | 5.3 | -1000.389942 | -4.264422 | -1004.472732 |
| conformer_112 | -1006.147413 | 0.290666 | 0.226331 | -1006.192425 | -118.2 | -1005.901759 | -1005.963082 | 9.2 | -1000.392969 | -4.263054 | -1004.471692 |
| conformer_162 | -1006.148233 | 0.290613 | 0.226436 | -1006.192391 | -115.9 | -1005.901778 | -1005.962943 | 9.6 | -1000.393974 | -4.263652 | -1004.472337 |
| conformer_146 | -1006.146822 | 0.290774 | 0.225684 | -1006.192378 | -119.6 | -1005.901604 | -1005.963682 | 7.6 | -1000.394849 | -4.260662 | -1004.472371 |
| conformer_53 | -1006.149260 | 0.290535 | 0.224883 | -1006.192369 | -113.2 | -1005.901834 | -1005.964474 | 5.6 | -1000.395643 | -4.262226 | -1004.473083 |
| conformer_39 | -1006.147729 | 0.290721 | 0.226575 | -1006.192165 | -116.7 | -1005.901444 | -1005.962578 | 10.5 | -1000.390205 | -4.265195 | -1004.470250 |
| conformer_76 | -1006.150871 | 0.290593 | 0.226038 | -1006.192121 | -108.3 | -1005.901528 | -1005.963071 | 9.2 | -1000.396358 | -4.263263 | -1004.471821 |
| conformer_71 | -1006.150769 | 0.290600 | 0.225891 | -1006.192104 | -108.5 | -1005.901504 | -1005.963201 | 8.9 | -1000.396265 | -4.263464 | -1004.472161 |
| conformer_103 | -1006.145187 | 0.290501 | 0.224822 | -1006.192079 | -123.1 | -1005.901578 | -1005.964245 | 6.2 | -1000.389762 | -4.263592 | -1004.472411 |
| conformer_156 | -1006.145329 | 0.290581 | 0.226540 | -1006.192051 | -122.7 | -1005.901470 | -1005.962499 | 10.7 | -1000.388012 | -4.265991 | -1004.471172 |
| conformer_92 | -1006.145521 | 0.290407 | 0.224821 | -1006.192036 | -122.1 | -1005.901629 | -1005.964203 | 6.3 | -1000.390108 | -4.263616 | -1004.472405 |
| conformer_67 | -1006.148927 | 0.290445 | 0.224677 | -1006.192011 | -113.1 | -1005.901566 | -1005.964322 | 6.0 | -1000.395485 | -4.262091 | -1004.472972 |
| conformer_181 | -1006.143827 | 0.290689 | 0.225297 | -1006.191986 | -126.4 | -1005.901297 | -1005.963677 | 7.6 | -1000.389603 | -4.262239 | -1004.471692 |
| conformer_95 | -1006.144872 | 0.290258 | 0.223963 | -1006.191976 | -123.7 | -1005.901718 | -1005.965001 | 4.2 | -1000.390366 | -4.262753 | -1004.473248 |
| conformer_109 | -1006.146494 | 0.290481 | 0.225060 | -1006.191923 | -119.3 | -1005.901442 | -1005.963851 | 7.2 | -1000.390768 | -4.263801 | -1004.471927 |
| conformer_88 | -1006.146629 | 0.290463 | 0.224721 | -1006.191901 | -118.9 | -1005.901438 | -1005.964168 | 6.4 | -1000.390992 | -4.263775 | -1004.472307 |
| conformer_36 | -1006.148046 | 0.290475 | 0.225130 | -1006.191855 | -115.0 | -1005.901380 | -1005.963713 | 7.6 | -1000.394442 | -4.262244 | -1004.472354 |
| conformer_59 | -1006.147541 | 0.290374 | 0.225031 | -1006.191803 | -116.2 | -1005.901429 | -1005.963760 | 7.4 | -1000.393480 | -4.262643 | -1004.472341 |
| conformer_69 | -1006.145962 | 0.290621 | 0.225734 | -1006.191779 | -120.3 | -1005.901158 | -1005.963033 | 9.3 | -1000.390896 | -4.263877 | -1004.471844 |
| conformer_179 | -1006.143345 | 0.290820 | 0.226611 | -1006.191759 | -127.1 | -1005.900939 | -1005.962136 | 11.7 | -1000.387584 | -4.264028 | -1004.470403 |
| conformer_193 | -1006.143247 | 0.290836 | 0.226839 | -1006.191747 | -127.3 | -1005.900911 | -1005.961896 | 12.3 | -1000.387459 | -4.263974 | -1004.470082 |
| conformer_34 | -1006.148033 | 0.290459 | 0.224773 | -1006.191715 | -114.7 | -1005.901256 | -1005.963930 | 7.0 | -1000.394595 | -4.262203 | -1004.472694 |
| conformer_182 | -1006.143936 | 0.290746 | 0.226295 | -1006.191701 | -125.4 | -1005.900955 | -1005.962394 | 11.0 | -1000.390298 | -4.262372 | -1004.471128 |
| conformer_137 | -1006.143724 | 0.290913 | 0.225054 | -1006.191680 | -125.9 | -1005.900767 | -1005.963614 | 7.8 | -1000.391252 | -4.261063 | -1004.472204 |
| conformer_141 | -1006.143750 | 0.290955 | 0.225408 | -1006.191618 | -125.7 | -1005.900663 | -1005.963198 | 8.9 | -1000.391075 | -4.261161 | -1004.471683 |
| conformer_57 | -1006.147607 | 0.290366 | 0.224926 | -1006.191604 | -115.5 | -1005.901238 | -1005.963666 | 7.7 | -1000.393769 | -4.262527 | -1004.472355 |
| conformer_107 | -1006.145149 | 0.290404 | 0.224523 | -1006.191590 | -121.9 | -1005.901186 | -1005.964055 | 6.7 | -1000.390173 | -4.263286 | -1004.472364 |
| conformer_131 | -1006.144983 | 0.290156 | 0.224055 | -1006.191565 | -122.3 | -1005.901409 | -1005.964498 | 5.5 | -1000.391212 | -4.262421 | -1004.473148 |
| conformer_155 | -1006.143121 | 0.290426 | 0.224883 | -1006.191533 | -127.1 | -1005.901107 | -1005.963638 | 7.7 | -1000.388964 | -4.262589 | -1004.472071 |
| conformer_148 | -1006.143129 | 0.290552 | 0.225528 | -1006.191449 | -126.9 | -1005.900897 | -1005.962909 | 9.7 | -1000.388879 | -4.262609 | -1004.471269 |
| conformer_9 | -1006.149823 | 0.290845 | 0.225548 | -1006.191391 | -109.1 | -1005.900546 | -1005.962831 | 9.9 | -1000.397220 | -4.261802 | -1004.472030 |
| conformer_94 | -1006.146942 | 0.290488 | 0.225347 | -1006.191296 | -116.4 | -1005.900808 | -1005.962937 | 9.6 | -1000.391722 | -4.263672 | -1004.471388 |
| conformer_78 | -1006.147042 | 0.290497 | 0.225255 | -1006.191279 | -116.1 | -1005.900782 | -1005.963012 | 9.4 | -1000.392328 | -4.263316 | -1004.471614 |

| | | | | | | | | | | | |
|---|---|---|---|---|---|---|---|---|---|---|---|
| conformer_11 | -1006.149797 | 0.290857 | 0.225623 | -1006.191253 | -108.8 | -1005.900396 | -1005.962618 | 10.4 | -1000.397352 | -4.261732 | -1004.471905 |
| conformer_97 | -1006.143267 | 0.290149 | 0.223457 | -1006.191197 | -125.8 | -1005.901048 | -1005.964728 | 4.9 | -1000.389457 | -4.262241 | -1004.473159 |
| conformer_125 | -1006.145026 | 0.290836 | 0.225243 | -1006.191173 | -121.2 | -1005.900337 | -1005.962918 | 9.6 | -1000.392485 | -4.261438 | -1004.471815 |
| conformer_172 | -1006.143381 | 0.290533 | 0.225343 | -1006.191170 | -125.5 | -1005.900637 | -1005.962815 | 9.9 | -1000.388860 | -4.262937 | -1004.471231 |
| conformer_79 | -1006.147911 | 0.290514 | 0.225194 | -1006.191159 | -113.5 | -1005.900645 | -1005.962953 | 9.5 | -1000.392950 | -4.263726 | -1004.471718 |
| conformer_170 | -1006.143440 | 0.290536 | 0.225316 | -1006.191142 | -125.2 | -1005.900606 | -1005.962814 | 9.9 | -1000.388906 | -4.262969 | -1004.471248 |
| conformer_123 | -1006.144978 | 0.290773 | 0.224867 | -1006.191079 | -121.0 | -1005.900306 | -1005.963200 | 8.9 | -1000.392568 | -4.261459 | -1004.472249 |
| conformer_68 | -1006.143600 | 0.290260 | 0.223601 | -1006.191022 | -124.5 | -1005.900762 | -1005.964409 | 5.7 | -1000.390010 | -4.262007 | -1004.472825 |
| conformer_185 | -1006.144495 | 0.290187 | 0.224243 | -1006.191018 | -122.1 | -1005.900831 | -1005.963763 | 7.4 | -1000.390178 | -4.263042 | -1004.472488 |
| conformer_126 | -1006.147222 | 0.290445 | 0.224860 | -1006.190891 | -114.7 | -1005.900446 | -1005.963019 | 9.4 | -1000.394265 | -4.262111 | -1004.472173 |
| conformer_13 | -1006.149433 | 0.290793 | 0.225474 | -1006.190885 | -108.8 | -1005.900092 | -1005.962399 | 11.0 | -1000.398215 | -4.260745 | -1004.471927 |
| conformer_74 | -1006.144277 | 0.290416 | 0.224513 | -1006.190880 | -122.4 | -1005.900464 | -1005.963355 | 8.5 | -1000.390618 | -4.262203 | -1004.471900 |
| conformer_111 | -1006.142927 | 0.290161 | 0.223879 | -1006.190864 | -125.9 | -1005.900703 | -1005.963973 | 6.9 | -1000.388914 | -4.262309 | -1004.472269 |
| conformer_114 | -1006.143700 | 0.290297 | 0.224478 | -1006.190843 | -123.8 | -1005.900546 | -1005.963353 | 8.5 | -1000.389640 | -4.262536 | -1004.471830 |
| conformer_159 | -1006.142451 | 0.290976 | 0.225637 | -1006.190808 | -127.0 | -1005.899832 | -1005.962159 | 11.6 | -1000.389852 | -4.260801 | -1004.470360 |
| conformer_31 | -1006.148671 | 0.290807 | 0.224787 | -1006.190803 | -110.6 | -1005.899996 | -1005.963004 | 9.4 | -1000.397548 | -4.260324 | -1004.472204 |
| conformer_135 | -1006.147021 | 0.290372 | 0.224570 | -1006.190742 | -114.8 | -1005.900370 | -1005.963160 | 9.0 | -1000.393937 | -4.262084 | -1004.472159 |
| conformer_10 | -1006.149575 | 0.290769 | 0.225311 | -1006.190739 | -108.1 | -1005.899970 | -1005.962416 | 11.0 | -1000.398249 | -4.260788 | -1004.471877 |
| conformer_33 | -1006.148514 | 0.290881 | 0.225158 | -1006.190704 | -110.8 | -1005.899823 | -1005.962534 | 10.6 | -1000.397499 | -4.260256 | -1004.471775 |
| conformer_77 | -1006.144036 | 0.290353 | 0.223966 | -1006.190664 | -122.4 | -1005.900311 | -1005.963686 | 7.6 | -1000.390311 | -4.262310 | -1004.472271 |
| conformer_149 | -1006.142550 | 0.290981 | 0.225621 | -1006.190648 | -126.3 | -1005.899667 | -1005.962015 | 12.0 | -1000.389976 | -4.260804 | -1004.470244 |
| conformer_108 | -1006.147252 | 0.290458 | 0.223151 | -1006.190639 | -113.9 | -1005.900181 | -1005.964476 | 5.6 | -1000.392647 | -4.263233 | -1004.473104 |
| conformer_122 | -1006.143344 | 0.290201 | 0.223244 | -1006.190583 | -124.0 | -1005.900382 | -1005.964327 | 5.9 | -1000.389234 | -4.262623 | -1004.472840 |
| conformer_157 | -1006.143156 | 0.290227 | 0.223801 | -1006.190281 | -123.7 | -1005.900054 | -1005.963468 | 8.2 | -1000.390242 | -4.261803 | -1004.472357 |
| conformer_30 | -1006.147456 | 0.290590 | 0.224863 | -1006.190242 | -112.3 | -1005.899652 | -1005.962367 | 11.1 | -1000.396641 | -4.260559 | -1004.472111 |
| conformer_29 | -1006.147643 | 0.290632 | 0.224977 | -1006.190126 | -111.5 | -1005.899494 | -1005.962137 | 11.7 | -1000.396671 | -4.260580 | -1004.471746 |
| conformer_191 | -1006.142866 | 0.290119 | 0.223478 | -1006.190099 | -124.0 | -1005.899980 | -1005.963609 | 7.8 | -1000.389122 | -4.262981 | -1004.472846 |
| conformer_124 | -1006.143201 | 0.290150 | 0.223542 | -1006.190071 | -123.1 | -1005.899921 | -1005.963517 | 8.1 | -1000.390727 | -4.261598 | -1004.472641 |
| conformer_180 | -1006.143645 | 0.290388 | 0.224226 | -1006.190042 | -121.8 | -1005.899654 | -1005.962804 | 9.9 | -1000.390443 | -4.262123 | -1004.471725 |
| conformer_175 | -1006.142652 | 0.289999 | 0.223012 | -1006.190002 | -124.3 | -1005.900003 | -1005.963978 | 6.9 | -1000.389779 | -4.261855 | -1004.472959 |
| conformer_45 | -1006.147873 | 0.290751 | 0.225354 | -1006.189924 | -110.4 | -1005.899173 | -1005.961558 | 13.2 | -1000.395798 | -4.261710 | -1004.471193 |
| conformer_43 | -1006.147902 | 0.290741 | 0.225330 | -1006.189731 | -109.8 | -1005.898990 | -1005.961389 | 13.7 | -1000.395978 | -4.261657 | -1004.471122 |
| conformer_102 | -1006.147014 | 0.290561 | 0.224892 | -1006.189704 | -112.1 | -1005.899143 | -1005.961800 | 12.6 | -1000.396669 | -4.260306 | -1004.471762 |
| conformer_121 | -1006.147055 | 0.290604 | 0.226247 | -1006.189558 | -111.6 | -1005.898954 | -1005.960299 | 16.5 | -1000.389094 | -4.266193 | -1004.468530 |
| conformer_87 | -1006.147221 | 0.290572 | 0.224883 | -1006.189541 | -111.1 | -1005.898969 | -1005.961646 | 13.0 | -1000.396763 | -4.260333 | -1004.471521 |
| conformer_192 | -1006.139465 | 0.290632 | 0.224292 | -1006.189506 | -131.4 | -1005.898874 | -1005.962202 | 11.5 | -1000.384140 | -4.263222 | -1004.470099 |
| conformer_47 | -1006.147585 | 0.290706 | 0.225305 | -1006.189438 | -109.9 | -1005.898732 | -1005.961121 | 14.4 | -1000.396953 | -4.260594 | -1004.471084 |
| conformer_169 | -1006.142391 | 0.290809 | 0.225394 | -1006.189347 | -123.3 | -1005.898538 | -1005.960941 | 14.8 | -1000.389672 | -4.261294 | -1004.469517 |
| conformer_101 | -1006.146432 | 0.290754 | 0.224883 | -1006.189309 | -112.6 | -1005.898555 | -1005.961414 | 13.6 | -1000.395989 | -4.260092 | -1004.471063 |
| conformer_96 | -1006.146529 | 0.290682 | 0.224790 | -1006.189222 | -112.1 | -1005.898540 | -1005.961420 | 13.6 | -1000.395911 | -4.260286 | -1004.471088 |
| conformer_46 | -1006.147644 | 0.290649 | 0.225145 | -1006.189219 | -109.2 | -1005.898570 | -1005.961062 | 14.5 | -1000.396829 | -4.260698 | -1004.470945 |
| conformer_80 | -1006.143615 | 0.290527 | 0.223997 | -1006.189120 | -119.5 | -1005.898593 | -1005.962111 | 11.8 | -1000.392052 | -4.260910 | -1004.471457 |
| conformer_161 | -1006.142458 | 0.290770 | 0.225287 | -1006.189074 | -122.4 | -1005.898304 | -1005.960775 | 15.3 | -1000.389761 | -4.261319 | -1004.469398 |
| conformer_75 | -1006.143795 | 0.290577 | 0.224403 | -1006.189015 | -118.7 | -1005.898438 | -1005.961600 | 13.1 | -1000.392085 | -4.261099 | -1004.470990 |
| conformer_171 | -1006.143819 | 0.290699 | 0.224850 | -1006.188979 | -118.6 | -1005.898280 | -1005.961117 | 14.4 | -1000.392187 | -4.260588 | -1004.470073 |
| conformer_176 | -1006.143890 | 0.290738 | 0.225031 | -1006.188931 | -118.3 | -1005.898193 | -1005.960888 | 15.0 | -1000.392197 | -4.260588 | -1004.469783 |
| conformer_189 | -1006.143890 | 0.290738 | 0.225031 | -1006.188931 | -118.3 | -1005.898193 | -1005.960888 | 15.0 | -1000.392196 | -4.260589 | -1004.469783 |
| conformer_2 | -1006.149277 | 0.290946 | 0.226081 | -1006.188834 | -103.9 | -1005.897888 | -1005.959741 | 18.0 | -1000.397556 | -4.261004 | -1004.469024 |
| conformer_84 | -1006.145590 | 0.290510 | 0.224771 | -1006.188825 | -113.5 | -1005.898315 | -1005.961042 | 14.6 | -1000.395344 | -4.260416 | -1004.471212 |
| conformer_98 | -1006.143931 | 0.290671 | 0.224677 | -1006.188794 | -117.8 | -1005.898123 | -1005.961105 | 14.4 | -1000.391873 | -4.261515 | -1004.470562 |
| conformer_1 | -1006.149418 | 0.290962 | 0.226157 | -1006.188671 | -103.1 | -1005.897709 | -1005.959502 | 18.6 | -1000.397626 | -4.261000 | -1004.468711 |
| conformer_90 | -1006.143924 | 0.290536 | 0.224148 | -1006.188645 | -117.4 | -1005.898109 | -1005.961485 | 13.4 | -1000.391955 | -4.261402 | -1004.470918 |
| conformer_86 | -1006.145485 | 0.290486 | 0.224732 | -1006.188555 | -113.1 | -1005.898069 | -1005.960811 | 15.2 | -1000.394989 | -4.260527 | -1004.470842 |
| conformer_91 | -1006.143704 | 0.290642 | 0.224747 | -1006.188466 | -117.5 | -1005.897824 | -1005.960707 | 15.4 | -1000.392907 | -4.260487 | -1004.470397 |
| conformer_153 | -1006.141766 | 0.290459 | 0.223717 | -1006.188308 | -122.2 | -1005.897849 | -1005.961579 | 13.2 | -1000.391254 | -4.259968 | -1004.471035 |
| conformer_165 | -1006.141425 | 0.290466 | 0.223748 | -1006.188286 | -123.0 | -1005.897820 | -1005.961526 | 13.3 | -1000.390901 | -4.259989 | -1004.470990 |

| | | | | | | | | | | | |
|---|---|---|---|---|---|---|---|---|---|---|---|
| conformer_187 | -1006.145110 | 0.290478 | 0.224776 | -1006.188280 | -113.3 | -1005.897802 | -1005.960492 | 16.0 | -1000.395341 | -4.260159 | -1004.470882 |
| conformer_85 | -1006.143079 | 0.290398 | 0.223873 | -1006.188221 | -118.5 | -1005.897823 | -1005.961336 | 13.8 | -1000.392543 | -4.260241 | -1004.471041 |
| conformer_81 | -1006.143083 | 0.290425 | 0.223813 | -1006.188191 | -118.4 | -1005.897766 | -1005.961366 | 13.7 | -1000.392618 | -4.260180 | -1004.471081 |
| conformer_99 | -1006.143586 | 0.290504 | 0.224175 | -1006.188154 | -117.0 | -1005.897650 | -1005.960967 | 14.8 | -1000.392783 | -4.260527 | -1004.470691 |
| conformer_127 | -1006.143472 | 0.290340 | 0.223540 | -1006.188008 | -116.9 | -1005.897668 | -1005.961456 | 13.5 | -1000.392866 | -4.260572 | -1004.471423 |
| conformer_132 | -1006.140993 | 0.290267 | 0.223541 | -1006.187822 | -122.9 | -1005.897555 | -1005.961269 | 14.0 | -1000.390791 | -4.259999 | -1004.471066 |
| conformer_152 | -1006.141585 | 0.290459 | 0.224287 | -1006.187754 | -121.2 | -1005.897295 | -1005.960455 | 16.1 | -1000.391181 | -4.260304 | -1004.470355 |
| conformer_133 | -1006.144794 | 0.290764 | 0.224702 | -1006.187464 | -112.0 | -1005.896700 | -1005.959750 | 18.0 | -1000.393620 | -4.260464 | -1004.469040 |
| conformer_134 | -1006.144869 | 0.290779 | 0.224743 | -1006.187457 | -111.8 | -1005.896678 | -1005.959702 | 18.1 | -1000.393607 | -4.260494 | -1004.468934 |
| conformer_24 | -1006.147434 | 0.290848 | 0.225896 | -1006.187392 | -104.9 | -1005.896544 | -1005.958484 | 21.3 | -1000.396304 | -4.260836 | -1004.468191 |
| conformer_164 | -1006.141336 | 0.290359 | 0.223824 | -1006.187355 | -120.8 | -1005.896996 | -1005.960519 | 15.9 | -1000.390834 | -4.260401 | -1004.470418 |
| conformer_25 | -1006.147355 | 0.290829 | 0.225884 | -1006.187127 | -104.4 | -1005.896298 | -1005.958231 | 21.9 | -1000.396106 | -4.260904 | -1004.467886 |
| conformer_142 | -1006.142767 | 0.290270 | 0.223471 | -1006.187017 | -116.2 | -1005.896747 | -1005.960534 | 15.9 | -1000.393349 | -4.259691 | -1004.470807 |
| conformer_55 | -1006.143474 | 0.290734 | 0.225000 | -1006.186354 | -112.6 | -1005.895620 | -1005.958342 | 21.7 | -1000.392128 | -4.260807 | -1004.467803 |
| conformer_49 | -1006.142885 | 0.290640 | 0.224746 | -1006.186294 | -114.0 | -1005.895654 | -1005.958536 | 21.1 | -1000.391926 | -4.260388 | -1004.467965 |
| conformer_56 | -1006.142621 | 0.290572 | 0.224637 | -1006.186265 | -114.6 | -1005.895693 | -1005.958616 | 20.9 | -1000.391590 | -4.260488 | -1004.468073 |
| conformer_61 | -1006.143317 | 0.290749 | 0.225085 | -1006.186046 | -112.2 | -1005.895297 | -1005.957949 | 22.7 | -1000.391965 | -4.260799 | -1004.467396 |
| conformer_115 | -1006.142215 | 0.290383 | 0.223981 | -1006.185056 | -112.5 | -1005.894673 | -1005.958063 | 22.4 | -1000.392329 | -4.259930 | -1004.468106 |
| conformer_183 | -1006.136770 | 0.291004 | 0.227066 | -1006.183038 | -121.5 | -1005.892034 | -1005.952960 | 35.8 | -1000.372909 | -4.272514 | -1004.461613 |

[a]: Single-point calculation in aqueous phase with SMD model.

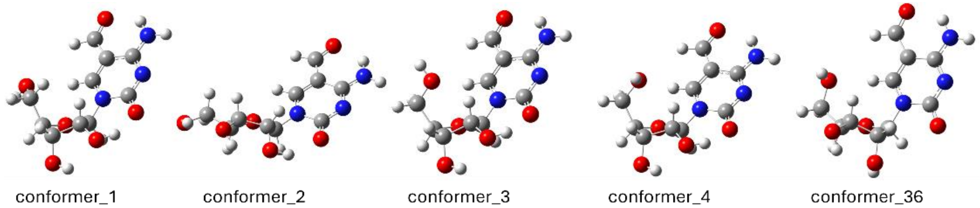

**Figure S32.** B3LYP-D3/def2-TZVPP optimized geometries of conformers for 5-formylcytidine (**1rb5fC**).

**Table S30.** Conformers of gas-phase optimized 5-formylcytidine (**1rb5fC**) at the B3LYP-D3/def2-TZVPP level of theory followed by aqueous phase single-point calculation. The columns display total energy without zero-point correction ($E_{Tot}$), thermal correction to enthalpy (δH), Gibbs free energy (δG), total energy without zero-point correction ($E_{Tot,W}$), Gibbs free energy ($G_{298,W}$) in water (W), total single-point energy ($E_{CBS}$) calculated at DLPNO-CCSD(T)/CBS level of theory, and their corresponding free energy $G_{CBS,W}$. $G_{298,W}$ and $G_{CBS,W}$ have been corrected to the standard state of 1 mol/L by addition of +7.908 kJ/mol. $\Delta G_{Solv}$ represents the Gibbs free energy of solvation. The data are arranged in the ascending numeric order of $E_{Tot,W}$. $\Delta G_{298,W}$ represents the respective energy difference to the lowest structure. Only conformers within the 24 kJ/mol (6 kcal/mol) energy window above the lowest in CREST are included in initial conformer sampling. Duplicates of the same structure are excluded. The overall optimum is marked bold.

| 1rb5fC No. | B3LYP-D3/def2-TZVPP $E_{Tot}$ (Hartree) | δH (Hartree) | δG (Hartree) | SMD(H2O)/B3LYP-D3/def2-TZVPP[a] $E_{Tot,W}$ (Hartree) | $\Delta G_{Solv}$ (kJ/mol) | $H_{298,W}$ (Hartree) | $G_{298,W}$ (Hartree) | $\Delta G_{298,W}$ (kJ/mol) | DLPNO-CCSD(T)/CBS $E_{CBS,HF}$ (Hartree) | $E_{CBS,C}$ (Hartree) | $G_{CBS,W}$ (Hartree) |
|---|---|---|---|---|---|---|---|---|---|---|---|
| conformer_23 | -1004.946847 | 0.266590 | 0.204108 | -1004.987772 | -107.4 | -1004.721182 | -1004.780652 | 5.7 | -999.237133 | -4.215404 | -1003.286341 |
| conformer_9 | -1004.948600 | 0.266272 | 0.202751 | -1004.987700 | -102.7 | -1004.721428 | -1004.781937 | 2.3 | -999.237276 | -4.217015 | -1003.287628 |
| conformer_15 | -1004.946002 | 0.266458 | 0.203686 | -1004.987558 | -109.1 | -1004.721100 | -1004.780860 | 5.1 | -999.236090 | -4.215249 | -1003.286197 |
| conformer_18 | -1004.943823 | 0.266405 | 0.203539 | -1004.987487 | -114.6 | -1004.721082 | -1004.780936 | 4.9 | -999.232036 | -4.216622 | -1003.285771 |
| conformer_4 | -1004.950083 | 0.266369 | 0.203525 | -1004.987484 | -98.2 | -1004.721115 | -1004.780947 | 4.9 | -999.239613 | -4.216167 | -1003.286643 |
| conformer_1 | -1004.949232 | 0.266508 | 0.204041 | -1004.987387 | -100.2 | -1004.720879 | -1004.780334 | 6.5 | -999.239531 | -4.216145 | -1003.286778 |
| conformer_14 | -1004.947025 | 0.266472 | 0.202628 | -1004.987309 | -105.8 | -1004.720837 | -1004.781669 | 3.0 | -999.239785 | -4.213164 | -1003.287594 |
| conformer_8 | -1004.947513 | 0.266328 | 0.202668 | -1004.987144 | -104.1 | -1004.720816 | -1004.781464 | 3.5 | -999.237021 | -4.216373 | -1003.287345 |
| conformer_32 | -1004.943202 | 0.266535 | 0.203497 | -1004.986725 | -114.3 | -1004.720190 | -1004.780216 | 6.8 | -999.233333 | -4.215271 | -1003.285618 |
| conformer_21 | -1004.948016 | 0.266339 | 0.203668 | -1004.986718 | -101.6 | -1004.720379 | -1004.780038 | 7.3 | -999.238824 | -4.216049 | -1003.286896 |
| conformer_3 | -1004.951051 | 0.266417 | 0.203500 | -1004.986546 | -93.2 | -1004.720129 | -1004.780034 | 7.3 | -999.241336 | -4.215850 | -1003.286170 |
| conformer_19 | -1004.947454 | 0.266454 | 0.202663 | -1004.986328 | -102.1 | -1004.719874 | -1004.780653 | 5.7 | -999.240959 | -4.212694 | -1003.286852 |
| conformer_5 | -1004.946893 | 0.266260 | 0.202295 | -1004.985930 | -102.5 | -1004.719670 | -1004.780623 | 5.7 | -999.238212 | -4.214770 | -1003.286711 |
| **conformer_36** | **-1004.940798** | **0.266158** | **0.200088** | **-1004.985912** | **-118.4** | **-1004.719754** | **-1004.782812** | **0** | **-999.230738** | **-4.215435** | **-1003.288186** |
| conformer_10 | -1004.946187 | 0.266168 | 0.202321 | -1004.985903 | -104.3 | -1004.719735 | -1004.780570 | 5.9 | -999.237118 | -4.215114 | -1003.286616 |
| conformer_17 | -1004.945530 | 0.266366 | 0.203273 | -1004.985891 | -106.0 | -1004.719525 | -1004.779606 | 8.4 | -999.235834 | -4.215585 | -1003.285495 |
| conformer_20 | -1004.946188 | 0.266193 | 0.202121 | -1004.985001 | -101.9 | -1004.718808 | -1004.779868 | 7.7 | -999.237991 | -4.214598 | -1003.286269 |
| conformer_7 | -1004.946370 | 0.266478 | 0.202499 | -1004.984342 | -99.7 | -1004.717864 | -1004.778831 | 10.5 | -999.239288 | -4.213965 | -1003.285714 |
| conformer_42 | -1004.938210 | 0.266229 | 0.202069 | -1004.984123 | -120.5 | -1004.717894 | -1004.779042 | 9.9 | -999.229053 | -4.214149 | -1003.284034 |
| conformer_30 | -1004.940315 | 0.266581 | 0.202313 | -1004.984067 | -114.9 | -1004.717486 | -1004.778742 | 10.7 | -999.232739 | -4.213675 | -1003.284842 |
| conformer_44 | -1004.940410 | 0.266323 | 0.203069 | -1004.983838 | -114.0 | -1004.717515 | -1004.777757 | 13.3 | -999.231954 | -4.214596 | -1003.283896 |
| conformer_46 | -1004.939456 | 0.266365 | 0.203248 | -1004.983721 | -116.2 | -1004.717356 | -1004.777461 | 14.0 | -999.228916 | -4.216273 | -1003.283194 |
| conformer_41 | -1004.937203 | 0.266146 | 0.201745 | -1004.983695 | -122.1 | -1004.717549 | -1004.778938 | 10.2 | -999.226938 | -4.215715 | -1003.284388 |
| conformer_6 | -1004.946134 | 0.266443 | 0.202522 | -1004.983641 | -98.5 | -1004.717198 | -1004.778107 | 12.4 | -999.240239 | -4.213028 | -1003.285240 |
| conformer_26 | -1004.941643 | 0.266433 | 0.202160 | -1004.983564 | -110.1 | -1004.717131 | -1004.778392 | 11.6 | -999.234172 | -4.214032 | -1003.284953 |
| conformer_34 | -1004.939033 | 0.266580 | 0.202537 | -1004.982977 | -115.4 | -1004.716397 | -1004.777428 | 14.1 | -999.231394 | -4.213527 | -1003.283316 |
| conformer_13 | -1004.944277 | 0.266276 | 0.201980 | -1004.982959 | -101.6 | -1004.716683 | -1004.777967 | 12.7 | -999.238962 | -4.212768 | -1003.285420 |
| conformer_24 | -1004.944098 | 0.266240 | 0.201979 | -1004.982452 | -100.7 | -1004.716212 | -1004.777461 | 14.0 | -999.239183 | -4.212505 | -1003.285052 |
| conformer_2 | -1004.946003 | 0.266574 | 0.202961 | -1004.981818 | -94.0 | -1004.715244 | -1004.775845 | 18.3 | -999.239590 | -4.213262 | -1003.282693 |
| conformer_40 | -1004.938936 | 0.266391 | 0.202183 | -1004.981550 | -111.9 | -1004.715159 | -1004.776355 | 17.0 | -999.231217 | -4.213995 | -1003.282631 |

| | | | | | | | | | | | |
|---|---|---|---|---|---|---|---|---|---|---|---|
| conformer_33 | -1004.940756 | 0.266350 | 0.201877 | -1004.981187 | -106.2 | -1004.714837 | -1004.776298 | 17.1 | -999.234063 | -4.213228 | -1003.282832 |
| conformer_48 | -1004.938562 | 0.266185 | 0.201763 | -1004.980946 | -111.3 | -1004.714761 | -1004.776171 | 17.4 | -999.232235 | -4.212637 | -1003.282481 |
| conformer_50 | -1004.938463 | 0.265951 | 0.201326 | -1004.980805 | -111.2 | -1004.714854 | -1004.776467 | 16.7 | -999.229287 | -4.214876 | -1003.282167 |
| conformer_29 | -1004.941182 | 0.266096 | 0.201781 | -1004.980126 | -102.2 | -1004.714030 | -1004.775333 | 19.6 | -999.232208 | -4.214589 | -1003.280949 |
| conformer_25 | -1004.941772 | 0.266368 | 0.201673 | -1004.979966 | -100.3 | -1004.713598 | -1004.775281 | 19.8 | -999.235627 | -4.212984 | -1003.282119 |
| conformer_38 | -1004.936256 | 0.265909 | 0.200518 | -1004.979772 | -114.3 | -1004.713863 | -1004.776242 | 17.2 | -999.229035 | -4.213175 | -1003.282196 |
| conformer_37 | -1004.934895 | 0.266121 | 0.200648 | -1004.975167 | -105.7 | -1004.709046 | -1004.771507 | 29.7 | -999.230094 | -4.211395 | -1003.278100 |

[a]: Single-point calculation in aqueous phase with SMD model.

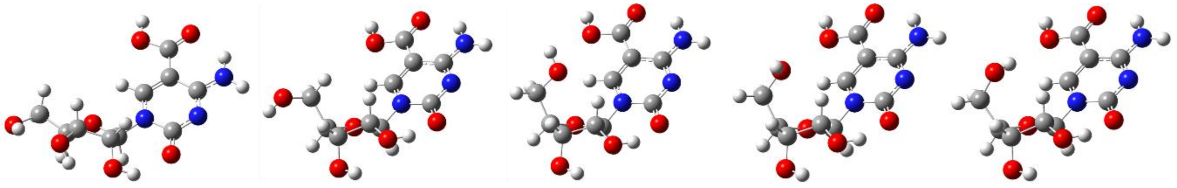

**Figure S33.** B3LYP-D3/def2-TZVPP optimized geometries of conformers for 5-carboxylcytidine (**1rb5caC**).

**Table S31.** Conformers of gas-phase optimized 5-carboxylcytidine (**1rb5caC**) at the B3LYP-D3/def2-TZVPP level of theory followed by aqueous phase single-point calculation. The columns display total energy without zero-point correction ($E_{Tot}$), thermal correction to enthalpy (δH), Gibbs free energy (δG), total energy without zero-point correction ($E_{Tot,W}$), Gibbs free energy ($G_{298,W}$) in water (W), total single-point energy ($E_{CBS}$) calculated at DLPNO-CCSD(T)/CBS level of theory, and their corresponding free energy $G_{CBS,W}$. $G_{298,W}$ and $G_{CBS,W}$ have been corrected to the standard state of 1 mol/L by addition of +7.908 kJ/mol. $\Delta G_{Solv}$ represents the Gibbs free energy of solvation. The data are arranged in the ascending numeric order of $E_{Tot,W}$. $\Delta G_{298,W}$ represents the respective energy difference to the lowest structure. Only conformers within the 24 kJ/mol (6 kcal/mol) energy window above the lowest in CREST are included in initial conformer sampling. Duplicates of the same structure are excluded. The overall optimum is marked bold.

| | B3LYP-D3/def2-TZVPP | | | SMD($H_2O$)/B3LYP-D3/def2-TZVPP[a] | | | | | DLPNO-CCSD(T)/CBS | | |
|---|---|---|---|---|---|---|---|---|---|---|---|
| 1rb5caC No. | $E_{Tot}$ (Hartree) | δH (Hartree) | δG (Hartree) | $E_{Tot,W}$ (Hartree) | $\Delta G_{Solv}$ (kJ/mol) | $H_{298,W}$ (Hartree) | $G_{298,W}$ (Hartree) | $\Delta G_{298,W}$ (kJ/mol) | $E_{CBS,HF}$ (Hartree) | $E_{CBS,C}$ (Hartree) | $G_{CBS,W}$ (Hartree) |
| **conformer_9** | **-1080.236967** | **0.272731** | **0.206577** | **-1080.277181** | **-105.6** | **-1080.004450** | **-1080.067592** | **0** | **-1074.169345** | **-4.486648** | **-1078.486617** |
| conformer_25 | -1080.234697 | 0.273021 | 0.208071 | -1080.276977 | -111.0 | -1080.003956 | -1080.065894 | 4.5 | -1074.168779 | -4.485109 | -1078.485085 |
| conformer_4 | -1080.237163 | 0.272751 | 0.207350 | -1080.276971 | -104.5 | -1080.004220 | -1080.066609 | 2.6 | -1074.170259 | -4.486232 | -1078.485937 |
| conformer_17 | -1080.231859 | 0.272940 | 0.207901 | -1080.276813 | -118.0 | -1080.003873 | -1080.065900 | 4.4 | -1074.163817 | -4.486336 | -1078.484193 |
| conformer_13 | -1080.233811 | 0.272881 | 0.207645 | -1080.276772 | -112.8 | -1080.003891 | -1080.066115 | 3.9 | -1074.167743 | -4.484891 | -1078.484938 |
| conformer_7 | -1080.235815 | 0.272782 | 0.207216 | -1080.276698 | -107.3 | -1080.003916 | -1080.066470 | 2.9 | -1074.169279 | -4.485737 | -1078.485671 |
| conformer_5 | -1080.235417 | 0.272787 | 0.207565 | -1080.276487 | -107.8 | -1080.003700 | -1080.065910 | 4.4 | -1074.169272 | -4.486073 | -1078.485838 |
| conformer_26 | -1080.234927 | 0.272763 | 0.207588 | -1080.276382 | -108.8 | -1080.003619 | -1080.065782 | 4.8 | -1074.168338 | -4.486707 | -1078.485900 |
| conformer_3 | -1080.238632 | 0.272811 | 0.207565 | -1080.276106 | -98.4 | -1080.003295 | -1080.065529 | 5.4 | -1074.172262 | -4.485991 | -1078.485150 |
| conformer_27 | -1080.232978 | 0.272855 | 0.206359 | -1080.275996 | -112.9 | -1080.003141 | -1080.066625 | 2.5 | -1074.169452 | -4.482987 | -1078.486086 |
| conformer_40 | -1080.231255 | 0.273010 | 0.207743 | -1080.275917 | -117.3 | -1080.002907 | -1080.065162 | 6.4 | -1074.165102 | -4.484952 | -1078.483961 |
| conformer_2 | -1080.234697 | 0.272782 | 0.206780 | -1080.275342 | -106.7 | -1080.002560 | -1080.065550 | 5.4 | -1074.169600 | -4.484657 | -1078.485111 |
| conformer_10 | -1080.234002 | 0.272666 | 0.206493 | -1080.275286 | -108.4 | -1080.002620 | -1080.065781 | 4.8 | -1074.168485 | -4.485012 | -1078.485275 |
| conformer_62 | -1080.225927 | 0.272570 | 0.205750 | -1080.275172 | -129.3 | -1080.002602 | -1080.066410 | 3.1 | -1074.159285 | -4.485578 | -1078.485346 |
| conformer_16 | -1080.233487 | 0.272882 | 0.207494 | -1080.275009 | -109.0 | -1080.002127 | -1080.064503 | 8.1 | -1074.167524 | -4.485272 | -1078.483812 |
| conformer_47 | -1080.229319 | 0.272952 | 0.206899 | -1080.274913 | -119.7 | -1080.001961 | -1080.065002 | 6.8 | -1074.162617 | -4.485075 | -1078.483374 |
| conformer_39 | -1080.230911 | 0.273006 | 0.209483 | -1080.274529 | -114.5 | -1080.001523 | -1080.062034 | 14.6 | -1074.159588 | -4.491455 | -1078.482165 |
| conformer_42 | -1080.227814 | 0.272916 | 0.207610 | -1080.274347 | -122.2 | -1080.001431 | -1080.063725 | 10.2 | -1074.159785 | -4.486132 | -1078.481827 |
| conformer_19 | -1080.233969 | 0.272685 | 0.206371 | -1080.274315 | -105.9 | -1080.001630 | -1080.064932 | 7.0 | -1074.169407 | -4.484450 | -1078.484820 |
| conformer_38 | -1080.229624 | 0.272837 | 0.207275 | -1080.274241 | -117.1 | -1080.001404 | -1080.063954 | 9.6 | -1074.163551 | -4.484772 | -1078.482653 |
| conformer_28 | -1080.232449 | 0.272696 | 0.207091 | -1080.274219 | -109.7 | -1080.001523 | -1080.064116 | 9.1 | -1074.165306 | -4.486291 | -1078.483264 |
| conformer_30 | -1080.231395 | 0.272703 | 0.206780 | -1080.274069 | -112.0 | -1080.001366 | -1080.064277 | 8.7 | -1074.165055 | -4.485286 | -1078.483223 |
| conformer_6 | -1080.234569 | 0.272924 | 0.206639 | -1080.273685 | -102.7 | -1080.000761 | -1080.064034 | 9.3 | -1074.171254 | -4.483554 | -1078.484274 |
| conformer_64 | -1080.222884 | 0.272453 | 0.204287 | -1080.273636 | -133.3 | -1080.001183 | -1080.066337 | 3.3 | -1074.157567 | -4.483975 | -1078.484995 |
| conformer_60 | -1080.231705 | 0.272886 | 0.208485 | -1080.273611 | -110.0 | -1080.000725 | -1080.062114 | 14.4 | -1074.162575 | -4.488280 | -1078.481265 |
| conformer_20 | -1080.234969 | 0.272846 | 0.207138 | -1080.273533 | -101.2 | -1080.000687 | -1080.063383 | 11.1 | -1074.167975 | -4.486028 | -1078.482416 |
| conformer_33 | -1080.229935 | 0.272687 | 0.206950 | -1080.273253 | -113.7 | -1080.000566 | -1080.063291 | 11.3 | -1074.163488 | -4.486119 | -1078.482963 |
| conformer_56 | -1080.227505 | 0.272957 | 0.207761 | -1080.273157 | -119.9 | -1080.000200 | -1080.062384 | 13.7 | -1074.160702 | -4.485916 | -1078.481497 |
| conformer_59 | -1080.228274 | 0.272847 | 0.207315 | -1080.273139 | -117.8 | -1080.000292 | -1080.062812 | 12.5 | -1074.163593 | -4.484229 | -1078.482359 |
| conformer_41 | -1080.228201 | 0.273056 | 0.206432 | -1080.273105 | -117.9 | -1080.000049 | -1080.063661 | 10.3 | -1074.164302 | -4.483412 | -1078.483174 |

| | | | | | | | | | | | |
|---|---|---|---|---|---|---|---|---|---|---|---|
| conformer_51 | -1080.225255 | 0.272651 | 0.205843 | -1080.273056 | -125.5 | -1080.000405 | -1080.064201 | 8.9 | -1074.158489 | -4.485532 | -1078.482966 |
| conformer_52 | -1080.225255 | 0.272651 | 0.205843 | -1080.273056 | -125.5 | -1080.000405 | -1080.064201 | 8.9 | -1074.158489 | -4.485531 | -1078.482966 |
| conformer_15 | -1080.232711 | 0.272932 | 0.205993 | -1080.272933 | -105.6 | -1080.000001 | -1080.063928 | 9.6 | -1074.170708 | -4.482302 | -1078.484227 |
| conformer_8 | -1080.234011 | 0.272902 | 0.206517 | -1080.272821 | -101.9 | -1079.999919 | -1080.063292 | 11.3 | -1074.171838 | -4.482777 | -1078.483896 |
| conformer_22 | -1080.230326 | 0.272709 | 0.206264 | -1080.272736 | -111.3 | -1080.000027 | -1080.063460 | 10.8 | -1074.165201 | -4.484496 | -1078.482830 |
| conformer_36 | -1080.229487 | 0.272964 | 0.206416 | -1080.272648 | -113.3 | -1079.999684 | -1080.063220 | 11.5 | -1074.165693 | -4.483752 | -1078.483178 |
| conformer_32 | -1080.229425 | 0.272576 | 0.205894 | -1080.272587 | -113.3 | -1080.000011 | -1080.063681 | 10.3 | -1074.163833 | -4.484899 | -1078.482989 |
| conformer_45 | -1080.229113 | 0.272822 | 0.207041 | -1080.272410 | -113.7 | -1079.999588 | -1080.062357 | 13.7 | -1074.163114 | -4.485114 | -1078.481472 |
| conformer_14 | -1080.231697 | 0.272704 | 0.206052 | -1080.272172 | -106.3 | -1079.999468 | -1080.063108 | 11.8 | -1074.170002 | -4.482519 | -1078.483933 |
| conformer_49 | -1080.226886 | 0.273069 | 0.206586 | -1080.271998 | -118.4 | -1079.998929 | -1080.062400 | 13.6 | -1074.163117 | -4.483031 | -1078.481662 |
| conformer_46 | -1080.229755 | 0.272621 | 0.205879 | -1080.271800 | -110.4 | -1079.999179 | -1080.062909 | 12.3 | -1074.165096 | -4.484348 | -1078.482597 |
| conformer_31 | -1080.231497 | 0.272679 | 0.206098 | -1080.271613 | -105.3 | -1079.998934 | -1080.062503 | 13.4 | -1074.170248 | -4.482231 | -1078.483484 |
| conformer_24 | -1080.230334 | 0.272893 | 0.206292 | -1080.271052 | -106.9 | -1079.998159 | -1080.061748 | 15.3 | -1074.167115 | -4.483211 | -1078.481741 |
| conformer_1 | -1080.233658 | 0.272961 | 0.206852 | -1080.270946 | -97.9 | -1079.997985 | -1080.061082 | 17.1 | -1074.170966 | -4.483051 | -1078.481442 |
| conformer_54 | -1080.226771 | 0.272865 | 0.206445 | -1080.270569 | -115.0 | -1079.997704 | -1080.061112 | 17.0 | -1074.162871 | -4.483551 | -1078.480764 |
| conformer_29 | -1080.229552 | 0.272841 | 0.206011 | -1080.270147 | -106.6 | -1079.997306 | -1080.061124 | 17.0 | -1074.167371 | -4.482563 | -1078.481506 |
| conformer_50 | -1080.228440 | 0.272800 | 0.205896 | -1080.270025 | -109.2 | -1079.997225 | -1080.061117 | 17.0 | -1074.165642 | -4.482819 | -1078.481138 |
| conformer_61 | -1080.226204 | 0.272711 | 0.206045 | -1080.269910 | -114.8 | -1079.997199 | -1080.060853 | 17.7 | -1074.163771 | -4.482159 | -1078.480579 |
| conformer_55 | -1080.226712 | 0.272639 | 0.205552 | -1080.269381 | -112.0 | -1079.996742 | -1080.060817 | 17.8 | -1074.164809 | -4.482526 | -1078.481441 |
| conformer_37 | -1080.229420 | 0.272774 | 0.205462 | -1080.268732 | -103.2 | -1079.995958 | -1080.060258 | 19.3 | -1074.167147 | -4.482604 | -1078.480589 |
| conformer_21 | -1080.229002 | 0.272908 | 0.206396 | -1080.268243 | -103.0 | -1079.995335 | -1080.058835 | 23.0 | -1074.166211 | -4.482952 | -1078.478996 |

[a]: Single-point calculation in aqueous phase with SMD model.

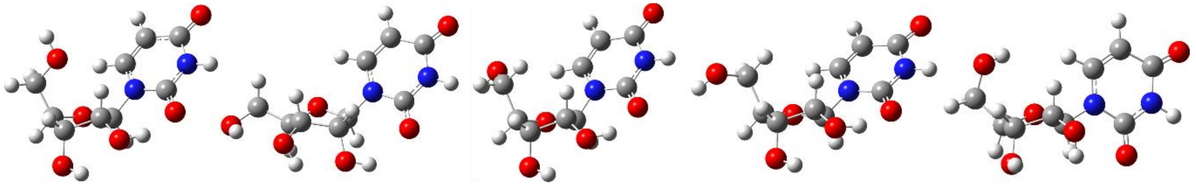

**Figure S34.** B3LYP-D3/def2-TZVPP optimized geometries of conformers for uridine (**1rbU**).

**Table S32.** Conformers of gas-phase optimized uridine (**1rbU**) at the B3LYP-D3/def2-TZVPP level of theory followed by aqueous phase single-point calculation. The columns display total energy without zero-point correction ($E_{Tot}$), thermal correction to enthalpy ($\delta H$), Gibbs free energy ($\delta G$), total energy without zero-point correction ($E_{Tot,W}$), Gibbs free energy ($G_{298,W}$) in water (W), total single-point energy ($E_{CBS}$) calculated at DLPNO-CCSD(T)/CBS level of theory, and their corresponding free energy $G_{CBS,W}$. $G_{298,W}$ and $G_{CBS,W}$ have been corrected to the standard state of 1 mol/L by addition of +7.908 kJ/mol. $\Delta G_{Solv}$ represents the Gibbs free energy of solvation. The data are arranged in the ascending numeric order of $E_{Tot,W}$. $\Delta G_{298,W}$ represents the respective energy difference to the lowest structure. Only conformers within the 24 kJ/mol (6 kcal/mol) energy window above the lowest in CREST are included in initial conformer sampling. Duplicates of the same structure are excluded. The overall optimum is marked bold.

| | B3LYP-D3/def2-TZVPP | | | SMD(H2O)/B3LYP-D3/def2-TZVPP[a] | | | | | DLPNO-CCSD(T)/CBS | | |
|---|---|---|---|---|---|---|---|---|---|---|---|
| 1rbU No. | $E_{Tot}$ (Hartree) | $\delta H$ (Hartree) | $\delta G$ (Hartree) | $E_{Tot,W}$ (Hartree) | $\Delta G_{Solv}$ (kJ/mol) | $H_{298,W}$ (Hartree) | $G_{298,W}$ (Hartree) | $\Delta G_{298,W}$ (kJ/mol) | $E_{CBS,HF}$ (Hartree) | $E_{CBS,C}$ (Hartree) | $G_{CBS,W}$ (Hartree) |
| conformer_26 | -911.455617 | 0.243099 | 0.185156 | -911.490299 | -91.1 | -911.247200 | -911.302131 | 4.2 | -906.310639 | -3.805105 | -909.962258 |
| conformer_5 | -911.454938 | 0.243014 | 0.184899 | -911.490193 | -92.6 | -911.247179 | -911.302282 | 3.8 | -906.309889 | -3.804903 | -909.962135 |
| conformer_10 | -911.452744 | 0.243000 | 0.184896 | -911.490152 | -98.2 | -911.247152 | -911.302244 | 3.9 | -906.305864 | -3.806212 | -909.961576 |
| conformer_32 | -911.451884 | 0.242992 | 0.184237 | -911.489004 | -97.5 | -911.246012 | -911.301755 | 5.2 | -906.306904 | -3.804647 | -909.961422 |
| **conformer_33** | **-911.449178** | **0.243039** | **0.182266** | **-911.488998** | **-104.5** | **-911.245959** | **-911.303720** | **0** | **-906.307035** | **-3.802388** | **-909.963965** |
| conformer_19 | -911.453419 | 0.242897 | 0.183629 | -911.488595 | -92.4 | -911.245698 | -911.301954 | 4.6 | -906.310414 | -3.802802 | -909.961751 |
| conformer_52 | -911.446833 | 0.242791 | 0.182355 | -911.488425 | -109.2 | -911.245634 | -911.303058 | 1.7 | -906.303116 | -3.803782 | -909.963123 |
| conformer_20 | -911.452153 | 0.242904 | 0.182425 | -911.488335 | -95.0 | -911.245431 | -911.302898 | 2.2 | -906.309089 | -3.802877 | -909.962711 |
| conformer_12 | -911.454067 | 0.242899 | 0.184298 | -911.488153 | -89.5 | -911.245254 | -911.300843 | 7.6 | -906.309031 | -3.805074 | -909.960881 |
| conformer_7 | -911.455413 | 0.242791 | 0.183912 | -911.487979 | -85.5 | -911.245188 | -911.301055 | 7.0 | -906.308729 | -3.806549 | -909.960920 |
| conformer_41 | -911.446889 | 0.242729 | 0.183019 | -911.487973 | -107.9 | -911.245244 | -911.301942 | 4.7 | -906.302148 | -3.803999 | -909.961200 |
| conformer_49 | -911.445703 | 0.242719 | 0.182352 | -911.487923 | -110.8 | -911.245204 | -911.302559 | 3.1 | -906.301873 | -3.803664 | -909.962393 |
| conformer_77 | -911.447275 | 0.242681 | 0.182849 | -911.487808 | -106.4 | -911.245127 | -911.301947 | 4.7 | -906.304457 | -3.802524 | -909.961653 |
| conformer_70 | -911.449695 | 0.242851 | 0.182826 | -911.487786 | -100.0 | -911.244935 | -911.301948 | 4.7 | -906.305190 | -3.804430 | -909.961873 |
| conformer_3 | -911.454347 | 0.242852 | 0.184546 | -911.487702 | -87.6 | -911.244850 | -911.300144 | 9.4 | -906.309102 | -3.805463 | -909.960362 |
| conformer_6 | -911.455244 | 0.242784 | 0.183806 | -911.487587 | -84.9 | -911.244803 | -911.300769 | 7.8 | -906.309488 | -3.805466 | -909.960480 |
| conformer_22 | -911.453859 | 0.242894 | 0.183607 | -911.487554 | -88.5 | -911.244660 | -911.300935 | 7.3 | -906.311729 | -3.802290 | -909.961095 |
| conformer_54 | -911.448020 | 0.242732 | 0.183105 | -911.487525 | -103.7 | -911.244793 | -911.301408 | 6.1 | -906.302466 | -3.804908 | -909.960762 |
| conformer_31 | -911.448534 | 0.242826 | 0.183410 | -911.487517 | -102.3 | -911.244691 | -911.301095 | 6.9 | -906.304684 | -3.803491 | -909.960735 |
| conformer_71 | -911.446289 | 0.242793 | 0.181766 | -911.487431 | -108.0 | -911.244638 | -911.302653 | 2.8 | -906.304669 | -3.801865 | -909.962897 |
| conformer_83 | -911.449577 | 0.242998 | 0.183147 | -911.487397 | -99.3 | -911.244399 | -911.301238 | 6.5 | -906.307433 | -3.802565 | -909.961659 |
| conformer_63 | -911.446664 | 0.242656 | 0.182060 | -911.487384 | -106.9 | -911.244728 | -911.302312 | 3.7 | -906.305005 | -3.802048 | -909.962701 |
| conformer_13 | -911.453456 | 0.242766 | 0.183602 | -911.487283 | -88.8 | -911.244517 | -911.300669 | 8.0 | -906.308164 | -3.805201 | -909.960578 |
| conformer_25 | -911.453479 | 0.242721 | 0.184236 | -911.487198 | -88.5 | -911.244477 | -911.299950 | 9.9 | -906.308752 | -3.805445 | -909.960668 |
| conformer_55 | -911.449205 | 0.242744 | 0.183001 | -911.487197 | -99.7 | -911.244453 | -911.301184 | 6.7 | -906.307341 | -3.801970 | -909.961291 |
| conformer_48 | -911.444037 | 0.242628 | 0.181993 | -911.487023 | -112.9 | -911.244395 | -911.302018 | 4.5 | -906.301332 | -3.802569 | -909.961883 |
| conformer_1 | -911.456743 | 0.242872 | 0.184196 | -911.486969 | -79.4 | -911.244097 | -911.299761 | 10.4 | -906.311736 | -3.805272 | -909.960027 |
| conformer_40 | -911.447629 | 0.242849 | 0.182864 | -911.486960 | -103.3 | -911.244111 | -911.301084 | 6.9 | -906.301735 | -3.805279 | -909.960468 |
| conformer_85 | -911.442958 | 0.242449 | 0.181253 | -911.486875 | -115.3 | -911.244426 | -911.302610 | 2.9 | -906.300203 | -3.802603 | -909.962458 |
| conformer_86 | -911.445697 | 0.242801 | 0.182703 | -911.486812 | -107.9 | -911.244011 | -911.301097 | 6.9 | -906.303631 | -3.802445 | -909.961476 |

| | | | | | | | | | | | |
|---|---|---|---|---|---|---|---|---|---|---|---|
| conformer_38 | -911.449125 | 0.242721 | 0.183741 | -911.486701 | -98.7 | -911.243980 | -911.299948 | 9.9 | -906.305609 | -3.804006 | -909.960439 |
| conformer_42 | -911.448087 | 0.242801 | 0.184033 | -911.486518 | -100.9 | -911.243717 | -911.299473 | 11.2 | -906.302505 | -3.805632 | -909.959522 |
| conformer_16 | -911.452099 | 0.242685 | 0.183213 | -911.486458 | -90.2 | -911.243773 | -911.300233 | 9.2 | -906.307642 | -3.804531 | -909.960308 |
| conformer_24 | -911.450681 | 0.242854 | 0.182926 | -911.486449 | -93.9 | -911.243595 | -911.300511 | 8.4 | -906.308295 | -3.803167 | -909.961292 |
| conformer_4 | -911.452986 | 0.242804 | 0.183402 | -911.486442 | -87.8 | -911.243638 | -911.300028 | 9.7 | -906.308968 | -3.804212 | -909.960222 |
| conformer_57 | -911.448189 | 0.242641 | 0.182479 | -911.486423 | -100.4 | -911.243782 | -911.300932 | 7.3 | -906.303715 | -3.804243 | -909.960701 |
| conformer_64 | -911.445848 | 0.242524 | 0.181252 | -911.486227 | -106.0 | -911.243703 | -911.301963 | 4.6 | -906.305381 | -3.800954 | -909.962450 |
| conformer_65 | -911.444861 | 0.242512 | 0.181192 | -911.485947 | -107.9 | -911.243435 | -911.301743 | 5.2 | -906.302039 | -3.802995 | -909.961916 |
| conformer_73 | -911.447066 | 0.242617 | 0.182403 | -911.485874 | -101.9 | -911.243257 | -911.300459 | 8.6 | -906.304863 | -3.802902 | -909.961158 |
| conformer_30 | -911.448532 | 0.243021 | 0.183469 | -911.485851 | -98.0 | -911.242830 | -911.299370 | 11.4 | -906.305735 | -3.802960 | -909.959533 |
| conformer_90 | -911.443899 | 0.242515 | 0.181540 | -911.485815 | -110.0 | -911.243300 | -911.301263 | 6.5 | -906.301829 | -3.802216 | -909.961408 |
| conformer_89 | -911.443305 | 0.242600 | 0.182163 | -911.485667 | -111.2 | -911.243067 | -911.300492 | 8.5 | -906.298344 | -3.804871 | -909.960403 |
| conformer_50 | -911.445454 | 0.242790 | 0.182973 | -911.485505 | -105.2 | -911.242715 | -911.299520 | 11.0 | -906.300083 | -3.804901 | -909.959050 |
| conformer_21 | -911.452403 | 0.242724 | 0.183143 | -911.485358 | -86.5 | -911.242634 | -911.299203 | 11.9 | -906.308940 | -3.803973 | -909.959712 |
| conformer_80 | -911.446253 | 0.242510 | 0.181115 | -911.485270 | -102.4 | -911.242760 | -911.301143 | 6.8 | -906.306446 | -3.800501 | -909.961836 |
| conformer_47 | -911.447080 | 0.242739 | 0.183018 | -911.485198 | -100.1 | -911.242459 | -911.299168 | 12.0 | -906.304966 | -3.802583 | -909.959637 |
| conformer_8 | -911.452832 | 0.242923 | 0.183175 | -911.485051 | -84.6 | -911.242128 | -911.298864 | 12.8 | -906.310401 | -3.803240 | -909.959673 |
| conformer_9 | -911.452476 | 0.242924 | 0.183396 | -911.484604 | -84.4 | -911.241680 | -911.298196 | 14.5 | -906.311183 | -3.802274 | -909.959177 |
| conformer_69 | -911.447760 | 0.242718 | 0.182272 | -911.484550 | -96.6 | -911.241832 | -911.299266 | 11.7 | -906.305174 | -3.802971 | -909.959650 |
| conformer_37 | -911.448310 | 0.242908 | 0.183545 | -911.484378 | -94.7 | -911.241470 | -911.297821 | 15.5 | -906.305209 | -3.803442 | -909.958162 |
| conformer_35 | -911.445187 | 0.242959 | 0.183142 | -911.484238 | -102.5 | -911.241279 | -911.298084 | 14.8 | -906.301502 | -3.803557 | -909.957956 |
| conformer_39 | -911.448021 | 0.242814 | 0.183154 | -911.484197 | -95.0 | -911.241383 | -911.298031 | 14.9 | -906.306668 | -3.801963 | -909.958641 |
| conformer_58 | -911.444660 | 0.242769 | 0.183402 | -911.484165 | -103.7 | -911.241396 | -911.297751 | 15.7 | -906.300389 | -3.804238 | -909.957718 |
| conformer_27 | -911.450469 | 0.242823 | 0.183182 | -911.484030 | -88.1 | -911.241207 | -911.297836 | 15.5 | -906.308367 | -3.802858 | -909.958591 |
| conformer_17 | -911.450286 | 0.242732 | 0.182858 | -911.483837 | -88.1 | -911.241105 | -911.297967 | 15.1 | -906.309398 | -3.802141 | -909.959220 |
| conformer_56 | -911.445437 | 0.242715 | 0.182571 | -911.483695 | -100.4 | -911.240980 | -911.298112 | 14.7 | -906.301989 | -3.803668 | -909.958333 |
| conformer_72 | -911.448176 | 0.242696 | 0.182161 | -911.483513 | -92.8 | -911.240817 | -911.298340 | 14.1 | -906.306470 | -3.802519 | -909.959153 |
| conformer_36 | -911.450263 | 0.242717 | 0.182905 | -911.483285 | -86.7 | -911.240568 | -911.297368 | 16.7 | -906.309800 | -3.801856 | -909.958761 |
| conformer_23 | -911.451279 | 0.242887 | 0.183170 | -911.483101 | -83.5 | -911.240214 | -911.296919 | 17.9 | -906.309871 | -3.802518 | -909.958028 |
| conformer_2 | -911.452255 | 0.243051 | 0.183891 | -911.482918 | -80.5 | -911.239867 | -911.296015 | 20.2 | -906.310261 | -3.802623 | -909.956644 |
| conformer_92 | -911.444352 | 0.242904 | 0.184266 | -911.482755 | -100.8 | -911.239851 | -911.295477 | 21.6 | -906.294879 | -3.809252 | -909.955256 |
| conformer_45 | -911.444673 | 0.242899 | 0.184571 | -911.482467 | -99.2 | -911.239568 | -911.294884 | 23.2 | -906.295687 | -3.808378 | -909.954276 |
| conformer_78 | -911.445803 | 0.242716 | 0.183122 | -911.481269 | -93.1 | -911.238553 | -911.295135 | 22.5 | -906.300994 | -3.805724 | -909.956050 |
| conformer_68 | -911.443943 | 0.242821 | 0.184235 | -911.480040 | -94.8 | -911.237219 | -911.292793 | 28.7 | -906.295929 | -3.808281 | -909.953060 |

[a]: Single-point calculation in aqueous phase with SMD model.

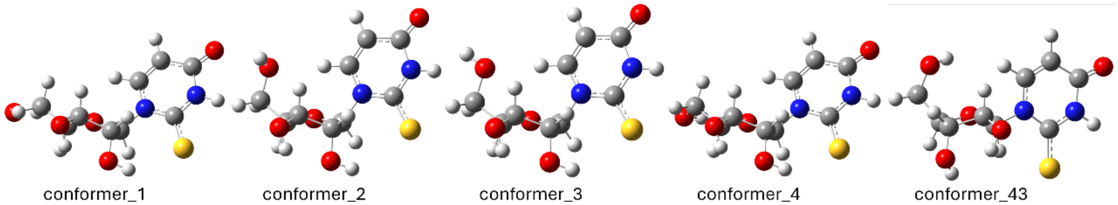

**Figure S35.** B3LYP-D3/def2-TZVPP optimized geometries of conformers for 2-thiouridine (**1rb2thU**).

**Table S33.** Conformers of gas-phase optimized 2-thiouridine (**1rb2thU**) at the B3LYP-D3/def2-TZVPP level of theory followed by aqueous phase single-point calculation. The columns display total energy without zero-point correction ($E_{Tot}$), thermal correction to enthalpy (δH), Gibbs free energy (δG), total energy without zero-point correction ($E_{Tot,W}$), Gibbs free energy ($G_{298,W}$) in water (W), total single-point energy ($E_{CBS}$) calculated at DLPNO-CCSD(T)/CBS level of theory, and their corresponding free energy $G_{CBS,W}$. $G_{298,W}$ and $G_{CBS,W}$ have been corrected to the standard state of 1 mol/L by addition of +7.908 kJ/mol. $\Delta G_{Solv}$ represents the Gibbs free energy of solvation. The data are arranged in the ascending numeric order of $E_{Tot,W}$. $\Delta G_{298,W}$ represents the respective energy difference to the lowest structure. Only conformers within the 24 kJ/mol (6 kcal/mol) energy window above the lowest in CREST are included in initial conformer sampling. Duplicates of the same structure are excluded. The overall optimum is marked bold.

| | B3LYP-D3/def2-TZVPP | | | SMD($H_2O$)/B3LYP-D3/def2-TZVPP[a] | | | | | DLPNO-CCSD(T)/CBS | | |
|---|---|---|---|---|---|---|---|---|---|---|---|
| 1rb2thU No. | $E_{Tot}$ (Hartree) | δH (Hartree) | δG (Hartree) | $E_{Tot,W}$ (Hartree) | $\Delta G_{Solv}$ (kJ/mol) | $H_{298,W}$ (Hartree) | $G_{298,W}$ (Hartree) | $\Delta G_{298,W}$ (kJ/mol) | $E_{CBS,HF}$ (Hartree) | $E_{CBS,C}$ (Hartree) | $G_{CBS,W}$ (Hartree) |
| conformer_2 | -1234.402284 | 0.241108 | 0.181132 | -1234.433897 | -83.0 | -1234.192789 | -1234.249753 | 2.7 | -1228.940961 | -3.758207 | -1232.546637 |
| conformer_3 | -1234.402795 | 0.241136 | 0.181324 | -1234.432967 | -79.2 | -1234.191831 | -1234.248631 | 5.7 | -1228.942261 | -3.757781 | -1232.545878 |
| conformer_59 | -1234.395512 | 0.240776 | 0.179573 | -1234.432817 | -97.9 | -1234.192041 | -1234.250232 | 1.5 | -1228.933165 | -3.759227 | -1232.547112 |
| conformer_68 | -1234.396743 | 0.240699 | 0.180481 | -1234.432477 | -93.8 | -1234.191778 | -1234.248984 | 4.7 | -1228.935246 | -3.759061 | -1232.546548 |
| conformer_20 | -1234.392587 | 0.240710 | 0.179910 | -1234.432377 | -104.5 | -1234.191667 | -1234.249455 | 3.5 | -1228.930086 | -3.758571 | -1232.545526 |
| conformer_47 | -1234.392495 | 0.240676 | 0.179914 | -1234.432372 | -104.7 | -1234.191696 | -1234.249446 | 3.5 | -1228.929182 | -3.760177 | -1232.546310 |
| **conformer_43** | **-1234.391895** | **0.240652** | **0.178559** | **-1234.432357** | **-106.2** | **-1234.191705** | **-1234.250786** | **0** | **-1228.930364** | **-3.758424** | **-1232.547678** |
| conformer_8 | -1234.401319 | 0.240535 | 0.181024 | -1234.432273 | -81.3 | -1234.191738 | -1234.248237 | 6.7 | -1228.936807 | -3.760818 | -1232.544543 |
| conformer_11 | -1234.401319 | 0.240535 | 0.181025 | -1234.432273 | -81.3 | -1234.191738 | -1234.248236 | 6.7 | -1228.936807 | -3.760818 | -1232.544542 |
| conformer_41 | -1234.390937 | 0.240681 | 0.179144 | -1234.432154 | -108.2 | -1234.191473 | -1234.249998 | 2.1 | -1228.929379 | -3.758183 | -1232.546623 |
| conformer_17 | -1234.398750 | 0.240421 | 0.179922 | -1234.432143 | -87.7 | -1234.191722 | -1234.249209 | 4.1 | -1228.934017 | -3.761493 | -1232.545968 |
| conformer_4 | -1234.400056 | 0.240853 | 0.180182 | -1234.432139 | -84.2 | -1234.191286 | -1234.248945 | 4.8 | -1228.939932 | -3.757853 | -1232.546674 |
| conformer_52 | -1234.392988 | 0.240622 | 0.179550 | -1234.432135 | -102.8 | -1234.191513 | -1234.249573 | 3.2 | -1228.932341 | -3.757170 | -1232.546096 |
| conformer_6 | -1234.399917 | 0.240586 | 0.181302 | -1234.431978 | -84.2 | -1234.191392 | -1234.247664 | 8.2 | -1228.934616 | -3.762273 | -1232.544637 |
| conformer_15 | -1234.394190 | 0.240797 | 0.180237 | -1234.431949 | -99.1 | -1234.191152 | -1234.248700 | 5.5 | -1228.932587 | -3.758143 | -1232.545240 |
| conformer_19 | -1234.398793 | 0.240375 | 0.179636 | -1234.431815 | -86.7 | -1234.191440 | -1234.249167 | 4.2 | -1228.934471 | -3.761043 | -1232.545889 |
| conformer_48 | -1234.391904 | 0.240642 | 0.178967 | -1234.431712 | -104.5 | -1234.191070 | -1234.249733 | 2.8 | -1228.932184 | -3.756858 | -1232.546871 |
| conformer_74 | -1234.392069 | 0.240398 | 0.178883 | -1234.431698 | -104.0 | -1234.191300 | -1234.249803 | 2.6 | -1228.930993 | -3.758638 | -1232.547365 |
| conformer_36 | -1234.394845 | 0.240690 | 0.179690 | -1234.431576 | -96.4 | -1234.190886 | -1234.248874 | 5.0 | -1228.935184 | -3.756612 | -1232.545826 |
| conformer_5 | -1234.402529 | 0.240583 | 0.181058 | -1234.431514 | -76.1 | -1234.190931 | -1234.247444 | 8.8 | -1228.939018 | -3.760323 | -1232.544255 |
| conformer_9 | -1234.397681 | 0.240560 | 0.180199 | -1234.431401 | -88.5 | -1234.190841 | -1234.248190 | 6.8 | -1228.934499 | -3.760013 | -1232.545021 |
| conformer_37 | -1234.398922 | 0.240563 | 0.181425 | -1234.431390 | -85.2 | -1234.190827 | -1234.246953 | 10.1 | -1228.933914 | -3.762458 | -1232.544404 |
| conformer_34 | -1234.389381 | 0.240578 | 0.178531 | -1234.431343 | -110.2 | -1234.190765 | -1234.249800 | 2.6 | -1228.928704 | -3.757322 | -1232.546444 |
| conformer_65 | -1234.389663 | 0.240416 | 0.178443 | -1234.431187 | -109.0 | -1234.190771 | -1234.249732 | 2.8 | -1228.926948 | -3.759830 | -1232.546846 |
| conformer_13 | -1234.397683 | 0.240690 | 0.179786 | -1234.431175 | -87.9 | -1234.190485 | -1234.248377 | 6.3 | -1228.938001 | -3.757573 | -1232.546269 |
| conformer_61 | -1234.388339 | 0.240418 | 0.177871 | -1234.431164 | -112.4 | -1234.190746 | -1234.250281 | 1.3 | -1228.927605 | -3.757279 | -1232.546827 |
| conformer_56 | -1234.394801 | 0.240674 | 0.180164 | -1234.431160 | -95.5 | -1234.190486 | -1234.247984 | 7.4 | -1228.933119 | -3.758982 | -1232.545283 |
| conformer_66 | -1234.389670 | 0.240396 | 0.177567 | -1234.430658 | -107.6 | -1234.190262 | -1234.250079 | 1.9 | -1228.930693 | -3.756010 | -1232.547113 |
| conformer_31 | -1234.397589 | 0.240645 | 0.179752 | -1234.430519 | -86.5 | -1234.189874 | -1234.247755 | 8.0 | -1228.938304 | -3.757384 | -1232.545854 |
| conformer_42 | -1234.391131 | 0.240501 | 0.177970 | -1234.430490 | -103.3 | -1234.189989 | -1234.249508 | 3.4 | -1228.932733 | -3.755671 | -1232.546781 |

| | | | | | | | | | | | |
|---|---|---|---|---|---|---|---|---|---|---|---|
| conformer_1 | -1234.400024 | 0.240958 | 0.180701 | -1234.430319 | -79.5 | -1234.189361 | -1234.246606 | 11.0 | -1228.939072 | -3.758349 | -1232.544003 |
| conformer_44 | -1234.396947 | 0.240459 | 0.179895 | -1234.430282 | -87.5 | -1234.189823 | -1234.247375 | 9.0 | -1228.934242 | -3.759872 | -1232.544542 |
| conformer_62 | -1234.389224 | 0.240489 | 0.178183 | -1234.430235 | -107.7 | -1234.189746 | -1234.249040 | 4.6 | -1228.929120 | -3.757045 | -1232.545981 |
| conformer_29 | -1234.394090 | 0.240960 | 0.181078 | -1234.430229 | -94.9 | -1234.189269 | -1234.246139 | 12.2 | -1228.930158 | -3.760868 | -1232.543076 |
| conformer_55 | -1234.395192 | 0.240634 | 0.179998 | -1234.430091 | -91.6 | -1234.189457 | -1234.247081 | 9.7 | -1228.934282 | -3.758618 | -1232.544790 |
| conformer_57 | -1234.391541 | 0.240526 | 0.178128 | -1234.429602 | -99.9 | -1234.189076 | -1234.248462 | 6.1 | -1228.933740 | -3.755242 | -1232.545903 |
| conformer_50 | -1234.387918 | 0.240627 | 0.179070 | -1234.429198 | -108.4 | -1234.188571 | -1234.247116 | 9.6 | -1228.924807 | -3.759239 | -1232.543244 |
| conformer_21 | -1234.393952 | 0.240749 | 0.180663 | -1234.429042 | -92.1 | -1234.188293 | -1234.245367 | 14.2 | -1228.929067 | -3.761629 | -1232.542110 |
| conformer_51 | -1234.394177 | 0.240589 | 0.178865 | -1234.429012 | -91.5 | -1234.188423 | -1234.247135 | 9.6 | -1228.935034 | -3.756814 | -1232.544806 |
| conformer_71 | -1234.395008 | 0.240851 | 0.181097 | -1234.428996 | -89.2 | -1234.188145 | -1234.244887 | 15.5 | -1228.930601 | -3.761597 | -1232.542078 |
| conformer_14 | -1234.395632 | 0.240846 | 0.180956 | -1234.428973 | -87.5 | -1234.188127 | -1234.245005 | 15.2 | -1228.931653 | -3.761296 | -1232.542323 |
| conformer_30 | -1234.391738 | 0.241008 | 0.180258 | -1234.428637 | -96.9 | -1234.187629 | -1234.245367 | 14.2 | -1228.930043 | -3.759102 | -1232.542774 |
| conformer_23 | -1234.392126 | 0.240875 | 0.180231 | -1234.427410 | -92.6 | -1234.186535 | -1234.244167 | 17.4 | -1228.930066 | -3.759766 | -1232.541874 |
| conformer_38 | -1234.390313 | 0.240997 | 0.180318 | -1234.426760 | -95.7 | -1234.185763 | -1234.243430 | 19.3 | -1228.928060 | -3.759286 | -1232.540464 |
| conformer_72 | -1234.388263 | 0.240587 | 0.179577 | -1234.426737 | -101.0 | -1234.186150 | -1234.244148 | 17.4 | -1228.926331 | -3.759573 | -1232.541789 |
| conformer_49 | -1234.387206 | 0.240804 | 0.179583 | -1234.425492 | -100.5 | -1234.184688 | -1234.242897 | 20.7 | -1228.924484 | -3.759718 | -1232.539892 |
| conformer_46 | -1234.389257 | 0.240809 | 0.179984 | -1234.425103 | -94.1 | -1234.184294 | -1234.242107 | 22.8 | -1228.926540 | -3.760052 | -1232.539443 |
| conformer_64 | -1234.388093 | 0.240664 | 0.179405 | -1234.425102 | -97.2 | -1234.184438 | -1234.242685 | 21.3 | -1228.926611 | -3.758857 | -1232.540061 |
| conformer_24 | -1234.391759 | 0.240766 | 0.179936 | -1234.424501 | -86.0 | -1234.183735 | -1234.241553 | 24.2 | -1228.930280 | -3.759226 | -1232.539300 |
| conformer_16 | -1234.392701 | 0.240808 | 0.180054 | -1234.423540 | -81.0 | -1234.182732 | -1234.240474 | 27.1 | -1228.931896 | -3.758932 | -1232.538602 |

[a]: Single-point calculation in aqueous phase with SMD model.

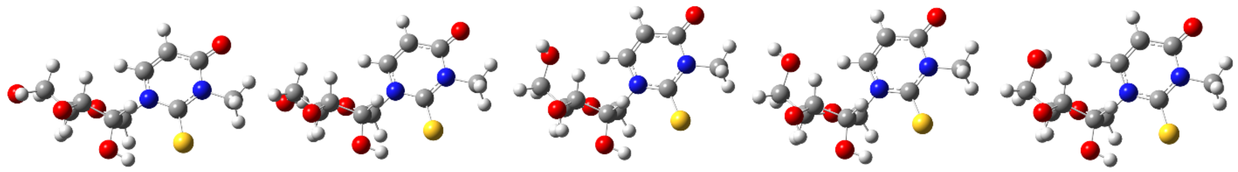

conformer_1 conformer_2 conformer_3 conformer_4 conformer_5

**Figure S36.** B3LYP-D3/def2-TZVPP optimized geometries of conformers for 2-thio-3-methyluridine (**1rb2th3mU**).

**Table S34.** Conformers of gas-phase optimized 2-thio-3-methyluridine (**1rb2th3mU**) at the B3LYP-D3/def2-TZVPP level of theory followed by aqueous phase single-point calculation. The columns display total energy without zero-point correction ($E_{Tot}$), thermal correction to enthalpy ($\delta H$), Gibbs free energy ($\delta G$), total energy without zero-point correction ($E_{Tot,W}$), Gibbs free energy ($G_{298,W}$) in water (W), total single-point energy ($E_{CBS}$) calculated at DLPNO-CCSD(T)/CBS level of theory, and their corresponding free energy $G_{CBS,W}$. $G_{298,W}$ and $G_{CBS,W}$ have been corrected to the standard state of 1 mol/L by addition of +7.908 kJ/mol. $\Delta G_{Solv}$ represents the Gibbs free energy of solvation. The data are arranged in the ascending numeric order of $E_{Tot,W}$. $\Delta G_{298,W}$ represents the respective energy difference to the lowest structure. Only conformers within the 24 kJ/mol (6 kcal/mol) energy window above the lowest in CREST are included in initial conformer sampling. Duplicates of the same structure are excluded. The overall optimum is marked bold.

| | **B3LYP-D3/def2-TZVPP** | | | **SMD($H_2O$)/B3LYP-D3/def2-TZVPP[a]** | | | | | **DLPNO-CCSD(T)/CBS** | | |
|---|---|---|---|---|---|---|---|---|---|---|---|
| **1rb2th3mU No.** | **$E_{Tot}$** (Hartree) | **$\delta H$** (Hartree) | **$\delta G$** (Hartree) | **$E_{Tot,W}$** (Hartree) | **$\Delta G_{Solv}$** (kJ/mol) | **$H_{298,W}$** (Hartree) | **$G_{298,W}$** (Hartree) | **$\Delta G_{298,W}$** (kJ/mol) | **$E_{CBS,HF}$** (Hartree) | **$E_{CBS,C}$** (Hartree) | **$G_{CBS,W}$** (Hartree) |
| conformer_3 | -1273.725581 | 0.270260 | 0.207133 | -1273.753045 | -72.1 | -1273.482785 | -1273.542900 | 1.4 | -1267.977915 | -3.972540 | -1271.767774 |
| **conformer_5** | **-1273.724437** | **0.270179** | **0.206255** | **-1273.752706** | **-74.2** | **-1273.482527** | **-1273.543439** | **0** | **-1267.976735** | **-3.972601** | **-1271.768337** |
| conformer_4 | -1273.726031 | 0.270231 | 0.207156 | -1273.752011 | -68.2 | -1273.481780 | -1273.541843 | 4.2 | -1267.979205 | -3.972188 | -1271.767206 |
| conformer_62 | -1273.718502 | 0.269950 | 0.205711 | -1273.751768 | -87.3 | -1273.481818 | -1273.543045 | 1.0 | -1267.969726 | -3.973795 | -1271.768064 |
| conformer_16 | -1273.722234 | 0.269710 | 0.206641 | -1273.751642 | -77.2 | -1273.481932 | -1273.541989 | 3.8 | -1267.971165 | -3.975730 | -1271.766651 |
| conformer_10 | -1273.724588 | 0.269769 | 0.207139 | -1273.751553 | -70.8 | -1273.481784 | -1273.541402 | 5.3 | -1267.973508 | -3.975274 | -1271.765597 |
| conformer_2 | -1273.723634 | 0.270142 | 0.206616 | -1273.751410 | -72.9 | -1273.481268 | -1273.541782 | 4.4 | -1267.977108 | -3.972288 | -1271.767543 |
| conformer_41 | -1273.715926 | 0.269935 | 0.206133 | -1273.751333 | -93.0 | -1273.481398 | -1273.542188 | 3.3 | -1267.966254 | -3.974536 | -1271.767051 |
| conformer_22 | -1273.722059 | 0.269614 | 0.205882 | -1273.751036 | -76.1 | -1273.481422 | -1273.542142 | 3.4 | -1267.971225 | -3.975471 | -1271.766780 |
| conformer_7 | -1273.722848 | 0.269752 | 0.207235 | -1273.750938 | -73.7 | -1273.481186 | -1273.540691 | 7.2 | -1267.971111 | -3.976704 | -1271.765658 |
| conformer_45 | -1273.714590 | 0.269861 | 0.204638 | -1273.750886 | -95.3 | -1273.481025 | -1273.543236 | 0.5 | -1267.966511 | -3.972902 | -1271.768059 |
| conformer_24 | -1273.715065 | 0.269912 | 0.205959 | -1273.750797 | -93.8 | -1273.480885 | -1273.541826 | 4.2 | -1267.965972 | -3.973013 | -1271.765747 |
| conformer_6 | -1273.725744 | 0.269875 | 0.207357 | -1273.750694 | -65.5 | -1273.480819 | -1273.540325 | 8.2 | -1267.975684 | -3.974777 | -1271.765043 |
| conformer_40 | -1273.713742 | 0.269910 | 0.205246 | -1273.750594 | -96.8 | -1273.480684 | -1273.542336 | 2.9 | -1267.965470 | -3.972832 | -1271.766896 |
| conformer_15 | -1273.721115 | 0.269933 | 0.205973 | -1273.750501 | -77.2 | -1273.480568 | -1273.541516 | 5.0 | -1267.975014 | -3.972042 | -1271.767456 |
| conformer_60 | -1273.715354 | 0.269799 | 0.205580 | -1273.750493 | -92.3 | -1273.480694 | -1273.541901 | 4.0 | -1267.968116 | -3.971736 | -1271.766399 |
| conformer_11 | -1273.720924 | 0.269708 | 0.206017 | -1273.750457 | -77.5 | -1273.480749 | -1273.541428 | 5.3 | -1267.971296 | -3.974438 | -1271.766238 |
| conformer_19 | -1273.716636 | 0.269993 | 0.206242 | -1273.750325 | -88.5 | -1273.480332 | -1273.541071 | 6.2 | -1267.968473 | -3.972610 | -1271.765518 |
| conformer_36 | -1273.721803 | 0.269651 | 0.207008 | -1273.750195 | -74.5 | -1273.480544 | -1273.540175 | 8.6 | -1267.970093 | -3.977304 | -1271.765770 |
| conformer_49 | -1273.714561 | 0.269854 | 0.204942 | -1273.750118 | -93.4 | -1273.480264 | -1273.542164 | 3.3 | -1267.968208 | -3.971394 | -1271.767205 |
| conformer_51 | -1273.717865 | 0.269842 | 0.206145 | -1273.750076 | -84.6 | -1273.480234 | -1273.540919 | 6.6 | -1267.969799 | -3.973550 | -1271.766404 |
| conformer_38 | -1273.717156 | 0.269856 | 0.205630 | -1273.749900 | -86.0 | -1273.480044 | -1273.541258 | 5.7 | -1267.970953 | -3.971102 | -1271.766157 |
| conformer_25 | -1273.720993 | 0.269892 | 0.205965 | -1273.749868 | -75.8 | -1273.479976 | -1273.540891 | 6.7 | -1267.975283 | -3.971840 | -1271.767022 |
| conformer_33 | -1273.712068 | 0.269796 | 0.204596 | -1273.749701 | -98.8 | -1273.479905 | -1273.542093 | 3.5 | -1267.964878 | -3.971772 | -1271.766675 |
| conformer_64 | -1273.711087 | 0.269692 | 0.204235 | -1273.749552 | -101.0 | -1273.479860 | -1273.542305 | 3.0 | -1267.963783 | -3.971771 | -1271.766772 |
| conformer_1 | -1273.723552 | 0.270184 | 0.206866 | -1273.749406 | -67.9 | -1273.479222 | -1273.539528 | 10.3 | -1267.976227 | -3.972818 | -1271.765021 |
| conformer_44 | -1273.720143 | 0.269621 | 0.205862 | -1273.749340 | -76.7 | -1273.479719 | -1273.540466 | 7.8 | -1267.970969 | -3.974281 | -1271.765572 |
| conformer_59 | -1273.718238 | 0.269820 | 0.206068 | -1273.749023 | -80.8 | -1273.479203 | -1273.539943 | 9.2 | -1267.970945 | -3.973226 | -1271.765877 |
| conformer_68 | -1273.712330 | 0.269653 | 0.203961 | -1273.748940 | -96.1 | -1273.479287 | -1273.541967 | 3.9 | -1267.966714 | -3.970561 | -1271.766912 |
| conformer_66 | -1273.711887 | 0.269731 | 0.204342 | -1273.748603 | -96.4 | -1273.478872 | -1273.541249 | 5.7 | -1267.965274 | -3.971476 | -1271.766112 |

| | | | | | | | | | | | |
|---|---|---|---|---|---|---|---|---|---|---|---|
| conformer_30 | -1273.716747 | 0.270136 | 0.207033 | -1273.748466 | -83.3 | -1273.478330 | -1273.538421 | 13.2 | -1267.966283 | -3.975601 | -1271.763559 |
| conformer_43 | -1273.717605 | 0.269762 | 0.204832 | -1273.748108 | -80.1 | -1273.478346 | -1273.540264 | 8.3 | -1267.972115 | -3.971197 | -1271.765971 |
| conformer_61 | -1273.714081 | 0.269733 | 0.204256 | -1273.747989 | -89.0 | -1273.478256 | -1273.540721 | 7.1 | -1267.969687 | -3.969787 | -1271.766114 |
| conformer_53 | -1273.710585 | 0.269841 | 0.204919 | -1273.747487 | -96.9 | -1273.477646 | -1273.539556 | 10.2 | -1267.960881 | -3.973893 | -1271.763746 |
| conformer_18 | -1273.718097 | 0.270027 | 0.206935 | -1273.747215 | -76.4 | -1273.477188 | -1273.537268 | 16.2 | -1267.967494 | -3.976174 | -1271.762839 |
| conformer_34 | -1273.714357 | 0.270216 | 0.206365 | -1273.746863 | -85.3 | -1273.476647 | -1273.537486 | 15.6 | -1267.966031 | -3.973887 | -1271.763046 |
| conformer_28 | -1273.714517 | 0.270023 | 0.206126 | -1273.745696 | -81.9 | -1273.475673 | -1273.536558 | 18.1 | -1267.965842 | -3.974638 | -1271.762521 |
| conformer_37 | -1273.712840 | 0.270203 | 0.206371 | -1273.744942 | -84.3 | -1273.474739 | -1273.535559 | 20.7 | -1267.964069 | -3.974145 | -1271.760934 |
| conformer_52 | -1273.711681 | 0.270053 | 0.206155 | -1273.743322 | -83.1 | -1273.473269 | -1273.534155 | 24.4 | -1267.962449 | -3.974790 | -1271.759713 |
| conformer_29 | -1273.714090 | 0.270022 | 0.206059 | -1273.742657 | -75.0 | -1273.472635 | -1273.533586 | 25.9 | -1267.966080 | -3.973976 | -1271.759552 |
| conformer_21 | -1273.714971 | 0.270057 | 0.206220 | -1273.741660 | -70.1 | -1273.471603 | -1273.532428 | 28.9 | -1267.967620 | -3.973696 | -1271.758773 |

[a]: Single-point calculation in aqueous phase with SMD model.

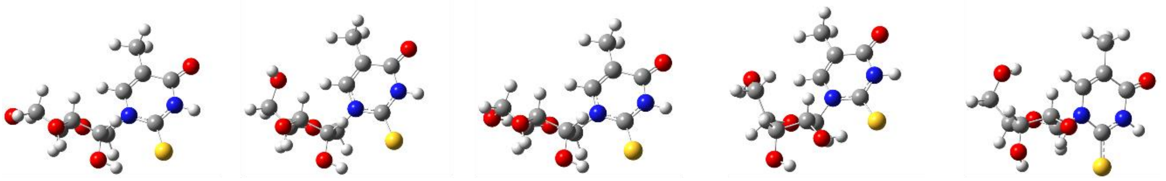

conformer_1 conformer_2 conformer_3 conformer_4 conformer_41

**Figure S37.** B3LYP-D3/def2-TZVPP optimized geometries of conformers for 2-thio-5-methyluridine (**1rb2th5mU**).

**Table S35.** Conformers of gas-phase optimized 2-thio-5-methyluridine (**1rb2th5mU**) at the B3LYP-D3/def2-TZVPP level of theory followed by aqueous phase single-point calculation. The columns display total energy without zero-point correction ($E_{Tot}$), thermal correction to enthalpy (δH), Gibbs free energy (δG), total energy without zero-point correction ($E_{Tot,W}$), Gibbs free energy ($G_{298,W}$) in water (W), total single-point energy ($E_{CBS}$) calculated at DLPNO-CCSD(T)/CBS level of theory, and their corresponding free energy $G_{CBS,W}$. $G_{298,W}$ and $G_{CBS,W}$ have been corrected to the standard state of 1 mol/L by addition of +7.908 kJ/mol. $\Delta G_{Solv}$ represents the Gibbs free energy of solvation. The data are arranged in the ascending numeric order of $E_{Tot,W}$. $\Delta G_{298,W}$ represents the respective energy difference to the lowest structure. Only conformers within the 24 kJ/mol (6 kcal/mol) energy window above the lowest in CREST are included in initial conformer sampling. Duplicates of the same structure are excluded. The overall optimum is marked bold.

| 1rb2th5mU | B3LYP-D3/def2-TZVPP | | | SMD(H2O)/B3LYP-D3/def2-TZVPP[a] | | | | | DLPNO-CCSD(T)/CBS | | |
|---|---|---|---|---|---|---|---|---|---|---|---|
| No. | $E_{Tot}$ (Hartree) | δH (Hartree) | δG (Hartree) | $E_{Tot,W}$ (Hartree) | $\Delta G_{Solv}$ (kJ/mol) | $H_{298,W}$ (Hartree) | $G_{298,W}$ (Hartree) | $\Delta G_{298,W}$ (kJ/mol) | $E_{CBS,HF}$ (Hartree) | $E_{CBS,C}$ (Hartree) | $G_{CBS,W}$ (Hartree) |
| conformer_2 | -1273.740402 | 0.270413 | 0.206841 | -1273.771098 | -80.6 | -1273.500685 | -1273.561245 | 2.9 | -1267.995933 | -3.968201 | -1271.784976 |
| conformer_5 | -1273.739161 | 0.270386 | 0.206142 | -1273.770981 | -83.5 | -1273.500595 | -1273.561827 | 1.4 | -1267.994619 | -3.968218 | -1271.785503 |
| conformer_6 | -1273.740728 | 0.270348 | 0.206607 | -1273.770027 | -76.9 | -1273.499679 | -1273.560408 | 5.1 | -1267.997156 | -3.967638 | -1271.784474 |
| conformer_54 | -1273.733579 | 0.270203 | 0.205214 | -1273.770004 | -95.6 | -1273.499801 | -1273.561778 | 1.5 | -1267.988031 | -3.969228 | -1271.785458 |
| **conformer_41** | **-1273.730180** | **0.270061** | **0.204504** | **-1273.769872** | **-104.2** | **-1273.499811** | **-1273.562356** | **0** | **-1267.985754** | **-3.968056** | **-1271.785986** |
| conformer_10 | -1273.739541 | 0.269888 | 0.206661 | -1273.769801 | -79.4 | -1273.499913 | -1273.560128 | 5.9 | -1267.991993 | -3.970612 | -1271.783191 |
| conformer_22 | -1273.730697 | 0.270160 | 0.205861 | -1273.769739 | -102.5 | -1273.499579 | -1273.560866 | 3.9 | -1267.985248 | -3.968247 | -1271.783663 |
| conformer_37 | -1273.729281 | 0.270066 | 0.204761 | -1273.769672 | -106.0 | -1273.499606 | -1273.561899 | 1.2 | -1267.984679 | -3.967929 | -1271.785225 |
| conformer_74 | -1273.734496 | 0.270140 | 0.206338 | -1273.769634 | -92.3 | -1273.499494 | -1273.560284 | 5.4 | -1267.989740 | -3.969037 | -1271.784565 |
| conformer_53 | -1273.730972 | 0.270087 | 0.205656 | -1273.769556 | -101.3 | -1273.499469 | -1273.560888 | 3.9 | -1267.987399 | -3.966889 | -1271.784204 |
| conformer_4 | -1273.738346 | 0.270035 | 0.207120 | -1273.769440 | -81.6 | -1273.499405 | -1273.559308 | 8.0 | -1267.989831 | -3.972227 | -1271.783019 |
| conformer_3 | -1273.738290 | 0.270266 | 0.206064 | -1273.769421 | -81.7 | -1273.499155 | -1273.560345 | 5.3 | -1267.995236 | -3.967417 | -1271.784707 |
| conformer_21 | -1273.737133 | 0.269744 | 0.205349 | -1273.769419 | -84.8 | -1273.499675 | -1273.561058 | 3.4 | -1267.989479 | -3.971015 | -1271.784420 |
| conformer_19 | -1273.732140 | 0.270217 | 0.206083 | -1273.769181 | -97.3 | -1273.498964 | -1273.560086 | 6.0 | -1267.987656 | -3.967749 | -1271.783351 |
| conformer_51 | -1273.730064 | 0.269952 | 0.204335 | -1273.769086 | -102.5 | -1273.499134 | -1273.561739 | 1.6 | -1267.987600 | -3.966439 | -1271.785714 |
| conformer_7 | -1273.740571 | 0.269961 | 0.206770 | -1273.768943 | -74.5 | -1273.498982 | -1273.559161 | 8.4 | -1267.993979 | -3.970114 | -1271.782683 |
| conformer_31 | -1273.737202 | 0.269960 | 0.207139 | -1273.768926 | -83.3 | -1273.498966 | -1273.558775 | 9.4 | -1267.988794 | -3.972608 | -1271.782974 |
| conformer_42 | -1273.732668 | 0.270109 | 0.205590 | -1273.768863 | -95.0 | -1273.498754 | -1273.560261 | 5.5 | -1267.990156 | -3.966277 | -1271.784025 |
| conformer_36 | -1273.727609 | 0.270011 | 0.204544 | -1273.768668 | -107.8 | -1273.498657 | -1273.561112 | 3.3 | -1267.984081 | -3.966878 | -1271.784462 |
| conformer_8 | -1273.735895 | 0.269752 | 0.205437 | -1273.768651 | -86.0 | -1273.498899 | -1273.560202 | 5.7 | -1267.989904 | -3.969647 | -1271.783858 |
| conformer_61 | -1273.726597 | 0.269950 | 0.204245 | -1273.768576 | -110.2 | -1273.498626 | -1273.561319 | 2.7 | -1267.982897 | -3.966946 | -1271.784565 |
| conformer_16 | -1273.735032 | 0.269658 | 0.205371 | -1273.768575 | -88.1 | -1273.498917 | -1273.560192 | 5.7 | -1267.988450 | -3.970107 | -1271.783717 |
| conformer_14 | -1273.735868 | 0.270087 | 0.205546 | -1273.768504 | -85.7 | -1273.498417 | -1273.559946 | 6.3 | -1267.993236 | -3.967256 | -1271.784569 |
| conformer_55 | -1273.732772 | 0.270213 | 0.206315 | -1273.768340 | -93.4 | -1273.498127 | -1273.559013 | 8.8 | -1267.987998 | -3.968993 | -1271.783231 |
| conformer_32 | -1273.735690 | 0.270033 | 0.205449 | -1273.767903 | -84.6 | -1273.497870 | -1273.559442 | 7.7 | -1267.993458 | -3.967045 | -1271.784255 |
| conformer_68 | -1273.727772 | 0.269865 | 0.203683 | -1273.767898 | -105.3 | -1273.498033 | -1273.561203 | 3.0 | -1267.986011 | -3.965561 | -1271.785003 |
| conformer_48 | -1273.729192 | 0.269905 | 0.203730 | -1273.767827 | -101.4 | -1273.497922 | -1273.561085 | 3.3 | -1267.987969 | -3.965244 | -1271.785106 |
| conformer_40 | -1273.735080 | 0.269632 | 0.204866 | -1273.767449 | -85.0 | -1273.497817 | -1273.559571 | 7.3 | -1267.989492 | -3.969545 | -1271.783528 |
| conformer_67 | -1273.727391 | 0.269941 | 0.204275 | -1273.767433 | -105.1 | -1273.497492 | -1273.560146 | 5.8 | -1267.984471 | -3.966565 | -1271.783792 |
| conformer_28 | -1273.732287 | 0.270265 | 0.206592 | -1273.767431 | -92.3 | -1273.497166 | -1273.557827 | 11.9 | -1267.985691 | -3.970436 | -1271.781667 |

| | | | | | | | | | | | |
|---|---|---|---|---|---|---|---|---|---|---|---|
| conformer_1 | -1273.738277 | 0.270456 | 0.206814 | -1273.767353 | -76.3 | -1273.496897 | -1273.557527 | 12.7 | -1267.994371 | -3.968102 | -1271.781723 |
| conformer_59 | -1273.733076 | 0.270136 | 0.206046 | -1273.767295 | -89.8 | -1273.497159 | -1273.558237 | 10.8 | -1267.989119 | -3.968572 | -1271.782853 |
| conformer_60 | -1273.729553 | 0.269933 | 0.203990 | -1273.766879 | -98.0 | -1273.496946 | -1273.559877 | 6.5 | -1267.988907 | -3.964864 | -1271.784095 |
| conformer_73 | -1273.729525 | 0.269993 | 0.206113 | -1273.766354 | -96.7 | -1273.496361 | -1273.557229 | 13.5 | -1267.980252 | -3.972486 | -1271.780443 |
| conformer_70 | -1273.733300 | 0.270268 | 0.206852 | -1273.766334 | -86.7 | -1273.496066 | -1273.556470 | 15.5 | -1267.986068 | -3.971225 | -1271.780464 |
| conformer_23 | -1273.732104 | 0.270076 | 0.206116 | -1273.766243 | -89.6 | -1273.496167 | -1273.557115 | 13.8 | -1267.984547 | -3.971198 | -1271.780756 |
| conformer_52 | -1273.732298 | 0.270042 | 0.204808 | -1273.766133 | -88.8 | -1273.496091 | -1273.558313 | 10.6 | -1267.990153 | -3.966587 | -1271.782754 |
| conformer_17 | -1273.733887 | 0.270271 | 0.206809 | -1273.766130 | -84.7 | -1273.495859 | -1273.556309 | 15.9 | -1267.987137 | -3.970903 | -1271.780462 |
| conformer_30 | -1273.729835 | 0.270349 | 0.205788 | -1273.765533 | -93.7 | -1273.495184 | -1273.556733 | 14.8 | -1267.985509 | -3.968610 | -1271.781017 |
| conformer_33 | -1273.730260 | 0.270246 | 0.205877 | -1273.764357 | -89.5 | -1273.494111 | -1273.555468 | 18.1 | -1267.985602 | -3.969293 | -1271.780103 |
| conformer_72 | -1273.726435 | 0.270075 | 0.205540 | -1273.763686 | -97.8 | -1273.493611 | -1273.555134 | 19.0 | -1267.981769 | -3.969132 | -1271.779600 |
| conformer_34 | -1273.728429 | 0.270369 | 0.206004 | -1273.763656 | -92.5 | -1273.493287 | -1273.554640 | 20.3 | -1267.983550 | -3.968819 | -1271.778580 |
| conformer_69 | -1273.725374 | 0.270203 | 0.205268 | -1273.762662 | -97.9 | -1273.492459 | -1273.554382 | 20.9 | -1267.979912 | -3.969256 | -1271.778177 |
| conformer_43 | -1273.727386 | 0.270197 | 0.205688 | -1273.762027 | -91.0 | -1273.491830 | -1273.553327 | 23.7 | -1267.982093 | -3.969528 | -1271.777562 |
| conformer_26 | -1273.729802 | 0.270179 | 0.205603 | -1273.761387 | -82.9 | -1273.491208 | -1273.552772 | 25.2 | -1267.985671 | -3.968728 | -1271.777369 |
| conformer_18 | -1273.730757 | 0.270211 | 0.205753 | -1273.760511 | -78.1 | -1273.490300 | -1273.551746 | 27.9 | -1267.987291 | -3.968422 | -1271.776702 |

[a]: Single-point calculation in aqueous phase with SMD model.

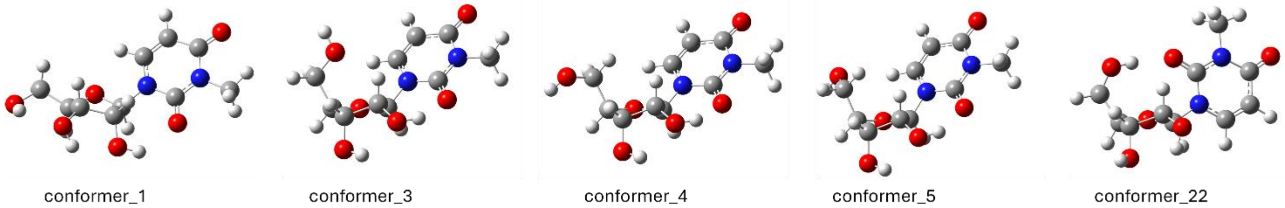

**Figure S38.** B3LYP-D3/def2-TZVPP optimized geometries of conformers for 3-methyluridine (**1rb3mU**).

**Table S36.** Conformers of gas-phase optimized 3-methyluridine (**1rb3mU**) at the B3LYP-D3/def2-TZVPP level of theory followed by aqueous phase single-point calculation. The columns display total energy without zero-point correction ($E_{Tot}$), thermal correction to enthalpy (δH), Gibbs free energy (δG), total energy without zero-point correction ($E_{Tot,W}$), Gibbs free energy ($G_{298,W}$) in water (W), total single-point energy ($E_{CBS}$) calculated at DLPNO-CCSD(T)/CBS level of theory, and their corresponding free energy $G_{CBS,W}$. $G_{298,W}$ and $G_{CBS,W}$ have been corrected to the standard state of 1 mol/L by addition of +7.908 kJ/mol. $\Delta G_{Solv}$ represents the Gibbs free energy of solvation. The data are arranged in the ascending numeric order of $E_{Tot,W}$. $\Delta G_{298,W}$ represents the respective energy difference to the lowest structure. Only conformers within the 24 kJ/mol (6 kcal/mol) energy window above the lowest in CREST are included in initial conformer sampling. Duplicates of the same structure are excluded. The overall optimum is marked bold.

| | B3LYP-D3/def2-TZVPP | | | SMD(H2O)/B3LYP-D3/def2-TZVPP[a] | | | | | DLPNO-CCSD(T)/CBS | | |
|---|---|---|---|---|---|---|---|---|---|---|---|
| 1rb3mU No. | $E_{Tot}$ (Hartree) | δH (Hartree) | δG (Hartree) | $E_{Tot,W}$ (Hartree) | $\Delta G_{Solv}$ (kJ/mol) | $H_{298,W}$ (Hartree) | $G_{298,W}$ (Hartree) | $\Delta G_{298,W}$ (kJ/mol) | $E_{CBS,HF}$ (Hartree) | $E_{CBS,C}$ (Hartree) | $G_{CBS,W}$ (Hartree) |
| **conformer_22** | **-950.780220** | **0.272354** | **0.209367** | **-950.812630** | **-85.1** | **-950.540276** | **-950.600251** | **0.0** | **-945.350498** | **-4.016546** | **-949.187075** |
| conformer_50 | -950.781235 | 0.272338 | 0.210715 | -950.812338 | -81.7 | -950.540000 | -950.598611 | 4.3 | -945.352141 | -4.016367 | -949.185884 |
| conformer_19 | -950.778483 | 0.272262 | 0.210464 | -950.812278 | -88.7 | -950.540016 | -950.598802 | 3.8 | -945.347459 | -4.017543 | -949.185322 |
| conformer_9 | -950.780576 | 0.272266 | 0.210433 | -950.812230 | -83.1 | -950.539964 | -950.598785 | 3.8 | -945.351397 | -4.016237 | -949.185843 |
| conformer_54 | -950.780772 | 0.272353 | 0.210147 | -950.811960 | -81.9 | -950.539607 | -950.598801 | 3.8 | -945.351781 | -4.016357 | -949.186167 |
| conformer_15 | -950.780078 | 0.272281 | 0.209865 | -950.811864 | -83.5 | -950.539583 | -950.598987 | 3.3 | -945.351005 | -4.016195 | -949.186108 |
| conformer_53 | -950.777960 | 0.272371 | 0.210130 | -950.811468 | -88.0 | -950.539097 | -950.598326 | 5.1 | -945.348698 | -4.016031 | -949.185096 |
| conformer_35 | -950.779029 | 0.272260 | 0.209215 | -950.810775 | -83.3 | -950.538515 | -950.598548 | 4.5 | -945.352207 | -4.013776 | -949.185502 |
| conformer_37 | -950.778886 | 0.272313 | 0.209257 | -950.810652 | -83.4 | -950.538339 | -950.598383 | 4.9 | -945.352087 | -4.013790 | -949.185375 |
| onformer_24 | -950.779953 | 0.272183 | 0.209820 | -950.810590 | -80.4 | -950.538407 | -950.597758 | 6.5 | -945.350698 | -4.016454 | -949.184957 |
| conformer_39 | -950.777824 | 0.272236 | 0.207621 | -950.810408 | -85.5 | -950.538172 | -950.599775 | 1.3 | -945.351120 | -4.013773 | -949.186844 |
| conformer_28 | -950.779653 | 0.272216 | 0.209647 | -950.810356 | -80.6 | -950.538140 | -950.597697 | 6.7 | -945.350515 | -4.016441 | -949.184999 |
| conformer_104 | -950.772219 | 0.272067 | 0.207707 | -950.810328 | -100.1 | -950.538261 | -950.599609 | 1.7 | -945.344628 | -4.014832 | -949.186850 |
| conformer_41 | -950.777674 | 0.272261 | 0.207103 | -950.810235 | -85.5 | -950.537974 | -950.600120 | 0.3 | -945.351078 | -4.013716 | -949.187241 |
| conformer_11 | -950.781362 | 0.272039 | 0.209350 | -950.810220 | -75.8 | -950.538181 | -950.597858 | 6.3 | -945.350480 | -4.017965 | -949.184940 |
| conformer_5 | -950.780193 | 0.272197 | 0.210169 | -950.810044 | -78.4 | -950.537847 | -950.596863 | 8.9 | -945.350865 | -4.016724 | -949.184258 |
| conformer_16 | -950.781032 | 0.272046 | 0.208730 | -950.809987 | -76.0 | -950.537941 | -950.598245 | 5.3 | -945.350157 | -4.018044 | -949.185414 |
| conformer_10 | -950.781038 | 0.272096 | 0.209549 | -950.809971 | -76.0 | -950.537875 | -950.597410 | 7.5 | -945.351172 | -4.016656 | -949.184200 |
| conformer_8 | -950.779942 | 0.272213 | 0.209758 | -950.809810 | -78.4 | -950.537597 | -950.597040 | 8.4 | -945.350677 | -4.016773 | -949.184548 |
| conformer_77 | -950.772125 | 0.272035 | 0.208714 | -950.809802 | -98.9 | -950.537767 | -950.598076 | 5.7 | -945.343466 | -4.014973 | -949.184390 |
| conformer_20 | -950.779541 | 0.272145 | 0.209224 | -950.809754 | -79.3 | -950.537609 | -950.597518 | 7.2 | -945.350001 | -4.016556 | -949.184535 |
| conformer_14 | -950.780779 | 0.272121 | 0.209091 | -950.809739 | -76.0 | -950.537618 | -950.597636 | 6.9 | -945.350989 | -4.016679 | -949.184525 |
| conformer_132 | -950.775073 | 0.272170 | 0.208609 | -950.809733 | -91.0 | -950.537563 | -950.598112 | 5.6 | -945.346657 | -4.015563 | -949.185260 |
| conformer_109 | -950.773614 | 0.272015 | 0.208513 | -950.809732 | -94.8 | -950.537717 | -950.598207 | 5.4 | -945.344236 | -4.015926 | -949.184755 |
| conformer_98 | -950.771141 | 0.272015 | 0.207703 | -950.809721 | -101.3 | -950.537706 | -950.599006 | 3.3 | -945.343916 | -4.014339 | -949.186120 |
| conformer_155 | -950.772416 | 0.271919 | 0.208269 | -950.809712 | -97.9 | -950.537793 | -950.598431 | 4.8 | -945.345708 | -4.013521 | -949.185245 |
| conformer_45 | -950.779426 | 0.272250 | 0.209156 | -950.809691 | -79.5 | -950.537441 | -950.597523 | 7.2 | -945.353492 | -4.013270 | -949.184859 |
| conformer_110 | -950.771725 | 0.272005 | 0.208242 | -950.809594 | -99.4 | -950.537589 | -950.598340 | 5.0 | -945.342876 | -4.015931 | -949.185422 |
| conformer_55 | -950.775290 | 0.272293 | 0.208498 | -950.809588 | -90.1 | -950.537295 | -950.598078 | 5.7 | -945.349043 | -4.013839 | -949.185671 |
| conformer_46 | -950.779278 | 0.272306 | 0.209210 | -950.809571 | -79.5 | -950.537265 | -950.597349 | 7.6 | -945.353368 | -4.013284 | -949.184723 |

| | | | | | | | | | | | |
|---|---|---|---|---|---|---|---|---|---|---|---|
| conformer_121 | -950.773442 | 0.272056 | 0.208535 | -950.809561 | -94.8 | -950.537505 | -950.598014 | 5.9 | -945.344095 | -4.015959 | -949.184625 |
| conformer_47 | -950.779330 | 0.272114 | 0.210052 | -950.809535 | -79.3 | -950.537421 | -950.596471 | 9.9 | -945.350552 | -4.016601 | -949.184294 |
| conformer_3 | -950.782480 | 0.272130 | 0.209600 | -950.809481 | -70.9 | -950.537351 | -950.596869 | 8.9 | -945.353383 | -4.016484 | -949.184256 |
| conformer_105 | -950.770839 | 0.271993 | 0.206988 | -950.809442 | -101.4 | -950.537449 | -950.599442 | 2.1 | -945.343672 | -4.014362 | -949.186637 |
| conformer_144 | -950.774771 | 0.272176 | 0.208279 | -950.809420 | -91.0 | -950.537244 | -950.598129 | 5.6 | -945.346404 | -4.015607 | -949.185369 |
| conformer_123 | -950.771557 | 0.272035 | 0.207979 | -950.809410 | -99.4 | -950.537375 | -950.598419 | 4.8 | -945.342721 | -4.015991 | -949.185574 |
| conformer_57 | -950.775010 | 0.272328 | 0.208383 | -950.809337 | -90.1 | -950.537009 | -950.597942 | 6.1 | -945.348808 | -4.013866 | -949.185605 |
| conformer_64 | -950.773752 | 0.272096 | 0.208988 | -950.809290 | -93.3 | -950.537194 | -950.597290 | 7.8 | -945.346014 | -4.014509 | -949.184061 |
| conformer_122 | -950.772049 | 0.272004 | 0.207901 | -950.809265 | -97.7 | -950.537261 | -950.598352 | 5.0 | -945.346729 | -4.012923 | -949.185956 |
| conformer_6 | -950.782201 | 0.272174 | 0.209297 | -950.809262 | -71.0 | -950.537088 | -950.596953 | 8.7 | -945.353171 | -4.016499 | -949.184422 |
| conformer_133 | -950.771673 | 0.272099 | 0.206980 | -950.809219 | -98.6 | -950.537120 | -950.599227 | 2.7 | -945.346269 | -4.012859 | -949.186682 |
| conformer_125 | -950.774324 | 0.271976 | 0.208415 | -950.809044 | -91.2 | -950.537068 | -950.597617 | 6.9 | -945.348588 | -4.012995 | -949.184876 |
| conformer_72 | -950.774917 | 0.272228 | 0.209925 | -950.809020 | -89.5 | -950.536792 | -950.596083 | 10.9 | -945.347064 | -4.015420 | -949.183650 |
| conformer_159 | -950.771344 | 0.272007 | 0.207698 | -950.809010 | -98.9 | -950.537003 | -950.598300 | 5.1 | -945.345271 | -4.013625 | -949.185853 |
| conformer_4 | -950.778851 | 0.272054 | 0.208851 | -950.808959 | -79.1 | -950.536905 | -950.597096 | 8.3 | -945.350750 | -4.015486 | -949.184482 |
| conformer_74 | -950.773970 | 0.272291 | 0.210127 | -950.808950 | -91.8 | -950.536659 | -950.595811 | 11.7 | -945.344027 | -4.017065 | -949.182933 |
| conformer_145 | -950.771403 | 0.272108 | 0.206711 | -950.808905 | -98.5 | -950.536797 | -950.599182 | 2.8 | -945.346033 | -4.012892 | -949.186704 |
| conformer_91 | -950.769429 | 0.271928 | 0.207491 | -950.808857 | -103.5 | -950.536929 | -950.598354 | 5.0 | -945.342785 | -4.013608 | -949.185317 |
| conformer_27 | -950.777974 | 0.271907 | 0.208432 | -950.808855 | -81.1 | -950.536948 | -950.597411 | 7.5 | -945.349446 | -4.015822 | -949.184705 |
| conformer_71 | -950.773100 | 0.272061 | 0.208137 | -950.808843 | -93.8 | -950.536782 | -950.597694 | 6.7 | -945.343367 | -4.016347 | -949.184307 |
| conformer_137 | -950.774014 | 0.271979 | 0.207930 | -950.808765 | -91.2 | -950.536786 | -950.597823 | 6.4 | -945.348334 | -4.013073 | -949.185216 |
| conformer_32 | -950.777670 | 0.271928 | 0.207993 | -950.808684 | -81.4 | -950.536756 | -950.597679 | 6.8 | -945.349193 | -4.015881 | -949.185082 |
| conformer_49 | -950.776299 | 0.272186 | 0.208753 | -950.808658 | -85.0 | -950.536472 | -950.596893 | 8.8 | -945.349836 | -4.014400 | -949.184831 |
| conformer_101 | -950.769135 | 0.271949 | 0.207049 | -950.808588 | -103.6 | -950.536639 | -950.598527 | 4.5 | -945.342532 | -4.013625 | -949.185549 |
| conformer_156 | -950.768367 | 0.271749 | 0.206206 | -950.808588 | -105.6 | -950.536839 | -950.599370 | 2.3 | -945.341799 | -4.013541 | -949.186343 |
| conformer_75 | -950.772812 | 0.272061 | 0.207738 | -950.808530 | -93.8 | -950.536469 | -950.597780 | 6.5 | -945.343160 | -4.016311 | -949.184439 |
| conformer_52 | -950.775981 | 0.272222 | 0.208605 | -950.808403 | -85.1 | -950.536181 | -950.596786 | 9.1 | -945.349561 | -4.014452 | -949.184818 |
| conformer_130 | -950.774025 | 0.272019 | 0.208437 | -950.808365 | -90.2 | -950.536346 | -950.596916 | 8.8 | -945.347073 | -4.014505 | -949.184468 |
| conformer_164 | -950.768075 | 0.271767 | 0.205894 | -950.808354 | -105.8 | -950.536587 | -950.599448 | 2.1 | -945.341497 | -4.013624 | -949.186494 |
| conformer_143 | -950.773837 | 0.272047 | 0.208356 | -950.808198 | -90.2 | -950.536151 | -950.596830 | 9.0 | -945.346899 | -4.014554 | -949.184447 |
| conformer_170 | -950.769043 | 0.271928 | 0.208139 | -950.808077 | -102.5 | -950.536149 | -950.596926 | 8.7 | -945.340323 | -4.015860 | -949.184067 |
| conformer_173 | -950.768874 | 0.271964 | 0.208083 | -950.808008 | -102.7 | -950.536044 | -950.596913 | 8.8 | -945.340197 | -4.015883 | -949.184118 |
| conformer_138 | -950.772625 | 0.271930 | 0.208020 | -950.808004 | -92.9 | -950.536074 | -950.596972 | 8.6 | -945.346366 | -4.014101 | -949.184814 |
| conformer_60 | -950.774225 | 0.272387 | 0.209242 | -950.807995 | -88.7 | -950.535608 | -950.595741 | 11.8 | -945.347461 | -4.014090 | -949.183067 |
| conformer_38 | -950.778251 | 0.271995 | 0.208757 | -950.807952 | -78.0 | -950.535957 | -950.596183 | 10.7 | -945.350672 | -4.015282 | -949.183887 |
| conformer_128 | -950.771123 | 0.271837 | 0.206771 | -950.807922 | -96.6 | -950.536085 | -950.598139 | 5.5 | -945.346907 | -4.011942 | -949.185866 |
| conformer_44 | -950.777948 | 0.272033 | 0.208398 | -950.807785 | -78.3 | -950.535752 | -950.596375 | 10.2 | -945.350408 | -4.015324 | -949.184158 |
| conformer_92 | -950.771257 | 0.272064 | 0.208444 | -950.807692 | -95.7 | -950.535628 | -950.596236 | 10.5 | -945.342148 | -4.015849 | -949.182976 |
| conformer_167 | -950.769289 | 0.271847 | 0.207199 | -950.807674 | -100.8 | -950.535827 | -950.597463 | 7.3 | -945.343239 | -4.013302 | -949.184715 |
| conformer_139 | -950.770817 | 0.271843 | 0.206188 | -950.807651 | -96.7 | -950.535808 | -950.598451 | 4.7 | -945.346640 | -4.011966 | -949.186240 |
| conformer_18 | -950.778612 | 0.272254 | 0.208585 | -950.807546 | -76.0 | -950.535292 | -950.595949 | 11.3 | -945.352331 | -4.014385 | -949.184053 |
| conformer_96 | -950.771098 | 0.272096 | 0.208371 | -950.807534 | -95.7 | -950.535438 | -950.596151 | 10.8 | -945.342014 | -4.015881 | -949.182948 |
| conformer_12 | -950.778829 | 0.272228 | 0.208793 | -950.807484 | -75.2 | -950.535256 | -950.595679 | 12.0 | -945.352446 | -4.014390 | -949.183685 |
| conformer_174 | -950.768991 | 0.271864 | 0.206736 | -950.807389 | -100.8 | -950.535525 | -950.597641 | 6.9 | -945.342988 | -4.013318 | -949.184956 |
| conformer_88 | -950.772519 | 0.271981 | 0.208306 | -950.807140 | -90.9 | -950.535159 | -950.595822 | 11.6 | -945.346385 | -4.013842 | -949.183530 |
| conformer_87 | -950.770467 | 0.272068 | 0.208723 | -950.807085 | -96.1 | -950.535017 | -950.595350 | 12.9 | -945.342040 | -4.015550 | -949.182473 |
| conformer_152 | -950.771534 | 0.271864 | 0.206924 | -950.806964 | -93.0 | -950.535100 | -950.597028 | 8.5 | -945.347966 | -4.011480 | -949.184941 |
| conformer_13 | -950.778482 | 0.272235 | 0.208732 | -950.806900 | -74.6 | -950.534665 | -950.595156 | 13.4 | -945.353274 | -4.013413 | -949.183362 |
| conformer_17 | -950.778293 | 0.272251 | 0.208375 | -950.806829 | -74.9 | -950.534578 | -950.595442 | 12.6 | -945.353170 | -4.013457 | -949.183776 |
| conformer_112 | -950.770999 | 0.272234 | 0.208664 | -950.806687 | -93.7 | -950.534453 | -950.595011 | 13.8 | -945.343295 | -4.014729 | -949.182037 |
| conformer_160 | -950.771226 | 0.271869 | 0.206330 | -950.806685 | -93.1 | -950.534816 | -950.597343 | 7.6 | -945.347699 | -4.011511 | -949.185327 |
| conformer_70 | -950.773873 | 0.272183 | 0.209089 | -950.806421 | -85.5 | -950.534238 | -950.594320 | 15.6 | -945.346826 | -4.014650 | -949.181923 |
| conformer_56 | -950.776053 | 0.272188 | 0.208949 | -950.806267 | -79.3 | -950.534079 | -950.594306 | 15.6 | -945.350023 | -4.013958 | -949.182233 |
| conformer_78 | -950.773527 | 0.272204 | 0.208841 | -950.806174 | -85.7 | -950.533970 | -950.594321 | 15.6 | -945.346548 | -4.014649 | -949.181991 |
| conformer_113 | -950.770420 | 0.272108 | 0.209094 | -950.806163 | -93.8 | -950.534055 | -950.594057 | 16.3 | -945.342244 | -4.015335 | -949.181216 |
| conformer_147 | -950.773396 | 0.272055 | 0.208504 | -950.806123 | -85.9 | -950.534068 | -950.594607 | 14.8 | -945.348025 | -4.013253 | -949.182488 |

| | | | | | | | | | | | |
|---|---|---|---|---|---|---|---|---|---|---|---|
| conformer_34 | -950.776229 | 0.272082 | 0.208500 | -950.806050 | -78.3 | -950.533968 | -950.594538 | 15.0 | -945.351340 | -4.013373 | -949.183021 |
| conformer_59 | -950.775726 | 0.272212 | 0.208712 | -950.805971 | -79.4 | -950.533759 | -950.594247 | 15.8 | -945.349771 | -4.013986 | -949.182278 |
| conformer_36 | -950.776019 | 0.272093 | 0.208163 | -950.805956 | -78.6 | -950.533863 | -950.594781 | 14.4 | -945.351232 | -4.013343 | -949.183337 |
| conformer_84 | -950.772986 | 0.272073 | 0.207912 | -950.805767 | -86.1 | -950.533694 | -950.594843 | 14.2 | -945.347671 | -4.013246 | -949.182774 |
| conformer_120 | -950.770926 | 0.271988 | 0.208060 | -950.805691 | -91.3 | -950.533703 | -950.594619 | 14.8 | -945.343417 | -4.014957 | -949.182066 |
| conformer_61 | -950.776171 | 0.272039 | 0.208406 | -950.805532 | -77.1 | -950.533493 | -950.594114 | 16.1 | -945.351738 | -4.013056 | -949.182737 |
| conformer_65 | -950.775962 | 0.272050 | 0.208060 | -950.805443 | -77.4 | -950.533393 | -950.594371 | 15.4 | -945.351623 | -4.013048 | -949.183080 |
| conformer_136 | -950.773497 | 0.271968 | 0.206676 | -950.805384 | -83.7 | -950.533416 | -950.595696 | 12.0 | -945.347868 | -4.013604 | -949.183671 |
| conformer_69 | -950.776772 | 0.272169 | 0.208789 | -950.805350 | -75.0 | -950.533181 | -950.593549 | 17.6 | -945.351480 | -4.013615 | -949.181873 |
| conformer_1 | -950.778232 | 0.272347 | 0.209364 | -950.805198 | -70.8 | -950.532851 | -950.592822 | 19.5 | -945.352361 | -4.013732 | -949.180683 |
| conformer_2 | -950.778064 | 0.272380 | 0.209231 | -950.805154 | -71.1 | -950.532774 | -950.592911 | 19.3 | -945.352250 | -4.013758 | -949.180856 |
| conformer_148 | -950.773198 | 0.271997 | 0.206929 | -950.805078 | -83.7 | -950.533081 | -950.595137 | 13.4 | -945.347622 | -4.013619 | -949.183180 |
| conformer_51 | -950.776444 | 0.272195 | 0.208523 | -950.805035 | -75.1 | -950.532840 | -950.593500 | 17.7 | -945.351228 | -4.013651 | -949.181935 |
| conformer_85 | -950.770265 | 0.272230 | 0.209680 | -950.804631 | -90.2 | -950.532401 | -950.591939 | 21.8 | -945.337106 | -4.019688 | -949.178468 |
| conformer_168 | -950.770054 | 0.272258 | 0.210095 | -950.804610 | -90.7 | -950.532352 | -950.591503 | 23.0 | -945.336221 | -4.020767 | -949.178438 |
| conformer_162 | -950.770913 | 0.272042 | 0.209111 | -950.803249 | -84.9 | -950.531207 | -950.591126 | 24.0 | -945.341806 | -4.017144 | -949.179163 |
| conformer_142 | -950.769238 | 0.272186 | 0.210119 | -950.801896 | -85.7 | -950.529710 | -950.588765 | 30.2 | -945.336819 | -4.019809 | -949.176154 |
| conformer_163 | -950.766194 | 0.272508 | 0.210514 | -950.797896 | -83.2 | -950.525388 | -950.584370 | 41.7 | -945.332101 | -4.021482 | -949.171759 |

[a]: Single-point calculation in aqueous phase with SMD model.

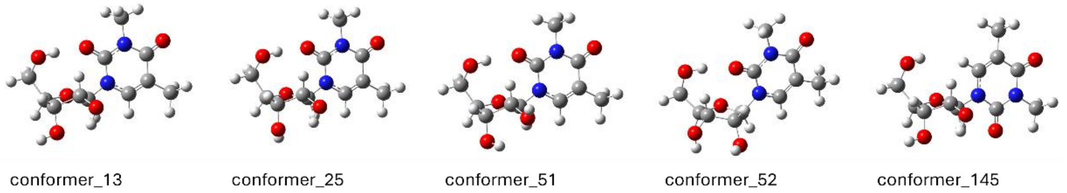

**Figure S39.** B3LYP-D3/def2-TZVPP optimized geometries of conformers for 3,5-dimethyluridine (**1rb3m5mU**).

**Table S37.** Conformers of gas-phase optimized 3,5-dimethyluridine (**1rb3m5mU**) at the B3LYP-D3/def2-TZVPP level of theory followed by aqueous phase single-point calculation. The columns display total energy without zero-point correction ($E_{Tot}$), thermal correction to enthalpy ($\delta H$), Gibbs free energy ($\delta G$), total energy without zero-point correction ($E_{Tot,W}$), Gibbs free energy ($G_{298,W}$) in water (W), total single-point energy ($E_{CBS}$) calculated at DLPNO-CCSD(T)/CBS level of theory, and their corresponding free energy $G_{CBS,W}$. $G_{298,W}$ and $G_{CBS,W}$ have been corrected to the standard state of 1 mol/L by addition of +7.908 kJ/mol. $\Delta G_{Solv}$ represents the Gibbs free energy of solvation. The data are arranged in the ascending numeric order of $E_{Tot,W}$. $\Delta G_{298,W}$ represents the respective energy difference to the lowest structure. Only conformers within the 24 kJ/mol (6 kcal/mol) energy window above the lowest in CREST are included in initial conformer sampling. Duplicates of the same structure are excluded. The overall optimum is marked bold.

| | B3LYP-D3/def2-TZVPP | | | SMD($H_2O$)/B3LYP-D3/def2-TZVPP[a] | | | | | DLPNO-CCSD(T)/CBS | | |
|---|---|---|---|---|---|---|---|---|---|---|---|
| 1rb3m5mU No. | $E_{Tot}$ (Hartree) | $\delta H$ (Hartree) | $\delta G$ (Hartree) | $E_{Tot,W}$ (Hartree) | $\Delta G_{Solv}$ (kJ/mol) | $H_{298,W}$ (Hartree) | $G_{298,W}$ (Hartree) | $\Delta G_{298,W}$ (kJ/mol) | $E_{CBS,HF}$ (Hartree) | $E_{CBS,C}$ (Hartree) | $G_{CBS,W}$ (Hartree) |
| conformer_25 | -990.1158949 | 0.3017420 | 0.2365280 | -990.1486787 | -86.1 | -989.846937 | -989.909139 | 5.3 | -984.4019192 | -4.2274616 | -988.4226245 |
| conformer_13 | -990.1179068 | 0.3017220 | 0.2364660 | -990.1486549 | -80.7 | -989.846933 | -989.909177 | 5.2 | -984.4057778 | -4.2261312 | -988.4231790 |
| conformer_51 | -990.1182672 | 0.3018710 | 0.2363140 | -990.1484638 | -79.3 | -989.846593 | -989.909138 | 5.3 | -984.4062832 | -4.2261979 | -988.4233517 |
| conformer_52 | -990.1152237 | 0.3016970 | 0.2357070 | -990.1475623 | -84.9 | -989.845865 | -989.908843 | 6.1 | -984.4030316 | -4.2259985 | -988.4226496 |
| conformer_39 | -990.1160334 | 0.3015970 | 0.2349720 | -990.1467957 | -80.8 | -989.845199 | -989.908812 | 6.1 | -984.4059058 | -4.2241344 | -988.4228184 |
| conformer_123 | -990.1095858 | 0.3014720 | 0.2335930 | -990.1467755 | -97.6 | -989.845304 | -989.910171 | 2.6 | -984.3989274 | -4.2248312 | -988.4243433 |
| conformer_41 | -990.1159741 | 0.3016150 | 0.2349310 | -990.1467420 | -80.8 | -989.845127 | -989.908799 | 6.2 | -984.4059028 | -4.2241579 | -988.4228856 |
| conformer_28 | -990.1172416 | 0.3015960 | 0.2357980 | -990.1466830 | -77.3 | -989.845087 | -989.907873 | 8.6 | -984.4049816 | -4.2264700 | -988.4220830 |
| conformer_38 | -990.1150151 | 0.3015990 | 0.2339850 | -990.1466659 | -83.1 | -989.845067 | -989.909669 | 3.9 | -984.4049346 | -4.2241006 | -988.4236890 |
| conformer_130 | -990.1093753 | 0.3014720 | 0.2332710 | -990.1466342 | -97.8 | -989.845162 | -989.910351 | 2.1 | -984.3987917 | -4.2248319 | -988.4245996 |
| conformer_40 | -990.1149613 | 0.3016370 | 0.2341000 | -990.1466013 | -83.1 | -989.844964 | -989.909489 | 4.4 | -984.4049346 | -4.2240877 | -988.4235503 |
| conformer_10 | -990.1186891 | 0.3014840 | 0.2355300 | -990.1465634 | -73.2 | -989.845079 | -989.908021 | 8.2 | -984.4044791 | -4.2282434 | -988.4220548 |
| conformer_14 | -990.1184593 | 0.3014800 | 0.2352730 | -990.1464979 | -73.6 | -989.845018 | -989.908213 | 7.7 | -984.4042703 | -4.2282836 | -988.4223075 |
| conformer_142 | -990.1123775 | 0.3015480 | 0.2343610 | -990.1462098 | -88.8 | -989.844662 | -989.908837 | 6.1 | -984.4007501 | -4.2257852 | -988.4229945 |
| conformer_3 | -990.1173702 | 0.3015110 | 0.2356790 | -990.1461395 | -75.5 | -989.844629 | -989.907449 | 9.7 | -984.4047943 | -4.2270083 | -988.4218809 |
| conformer_17 | -990.1180787 | 0.3014900 | 0.2351450 | -990.1461219 | -73.6 | -989.844632 | -989.907965 | 8.4 | -984.4050717 | -4.2268080 | -988.4217658 |
| conformer_80 | -990.1092338 | 0.3014560 | 0.2346260 | -990.1460398 | -96.6 | -989.844584 | -989.908402 | 7.2 | -984.3975292 | -4.2250309 | -988.4217281 |
| conformer_95 | -990.1085684 | 0.3013220 | 0.2333420 | -990.1460350 | -98.4 | -989.844713 | -989.909681 | 3.9 | -984.3978404 | -4.2247866 | -988.4237396 |
| conformer_5 | -990.1172358 | 0.3015510 | 0.2355630 | -990.1460336 | -75.6 | -989.844483 | -989.907459 | 9.7 | -984.4047343 | -4.2270464 | -988.4220035 |
| conformer_23 | -990.1179239 | 0.3015160 | 0.2350110 | -990.1460125 | -73.7 | -989.844496 | -989.907989 | 8.3 | -984.4050155 | -4.2267757 | -988.4218568 |
| conformer_136 | -990.1103926 | 0.3013020 | 0.2321380 | -990.1460070 | -93.5 | -989.844705 | -989.910857 | 0.8 | -984.4013646 | -4.2235721 | -988.4254012 |
| conformer_117 | -990.1083720 | 0.3013160 | 0.2327800 | -990.1458754 | -98.5 | -989.844559 | -989.910083 | 2.8 | -984.3977015 | -4.2247921 | -988.4242051 |
| conformer_182 | -990.1094456 | 0.3013230 | 0.2342180 | -990.1458719 | -95.6 | -989.844549 | -989.908642 | 6.6 | -984.3996873 | -4.2236108 | -988.4224944 |
| conformer_22 | -990.1168632 | 0.3014410 | 0.2350090 | -990.1458652 | -76.1 | -989.844424 | -989.907844 | 8.7 | -984.4042630 | -4.2267253 | -988.4219693 |
| conformer_27 | -990.1166969 | 0.3014600 | 0.2346430 | -990.1458487 | -76.5 | -989.844389 | -989.908194 | 7.8 | -984.4041603 | -4.2266446 | -988.4223017 |
| **conformer_145** | **-990.1101764** | **0.3012990** | **0.2316880** | **-990.1458480** | **-93.7** | **-989.844549** | **-989.911148** | **0** | **-984.4012130** | **-4.2235888** | **-988.4257734** |
| conformer_87 | -990.1090464 | 0.3014690 | 0.2344920 | -990.1458443 | -96.6 | -989.844375 | -989.908340 | 7.4 | -984.3974172 | -4.2250378 | -988.4217489 |
| conformer_47 | -990.1164089 | 0.3014200 | 0.2355590 | -990.1457434 | -77.0 | -989.844323 | -989.907172 | 10.4 | -984.4042178 | -4.2271230 | -988.4221043 |
| conformer_50 | -990.1162674 | 0.3014540 | 0.2355120 | -990.1456878 | -77.2 | -989.844234 | -989.907164 | 10.5 | -984.4041348 | -4.2271436 | -988.4221749 |
| conformer_191 | -990.1092564 | 0.3013360 | 0.2341060 | -990.1456658 | -95.6 | -989.844330 | -989.908548 | 6.8 | -984.3995643 | -4.2236178 | -988.4224735 |

| | | | | | | | | | | | |
|---|---|---|---|---|---|---|---|---|---|---|---|
| conformer_55 | -990.1125568 | 0.3017740 | 0.2346390 | -990.1456636 | -86.9 | -989.843890 | -989.908013 | 8.2 | -984.4033558 | -4.2236936 | -988.4225051 |
| conformer_132 | -990.1089246 | 0.3013890 | 0.2343700 | -990.1456531 | -96.4 | -989.844264 | -989.908271 | 7.6 | -984.3968549 | -4.2261175 | -988.4223188 |
| conformer_45 | -990.1162957 | 0.3015640 | 0.2348330 | -990.1456118 | -77.0 | -989.844048 | -989.907767 | 8.9 | -984.4071136 | -4.2235641 | -988.4221488 |
| conformer_149 | -990.1090190 | 0.3014470 | 0.2329130 | -990.1456074 | -96.1 | -989.844160 | -989.909682 | 3.8 | -984.4006692 | -4.2228002 | -988.4241328 |
| conformer_120 | -990.1106864 | 0.3014060 | 0.2341300 | -990.1455943 | -91.7 | -989.844188 | -989.908452 | 7.1 | -984.3980054 | -4.2262260 | -988.4219972 |
| conformer_133 | -990.1088575 | 0.3014090 | 0.2343300 | -990.1455670 | -96.4 | -989.844158 | -989.908225 | 7.7 | -984.3968297 | -4.2261207 | -988.4223179 |
| conformer_146 | -990.1162277 | 0.3015770 | 0.2347620 | -990.1455582 | -77.0 | -989.843981 | -989.907784 | 8.8 | -984.4071102 | -4.2235884 | -988.4222551 |
| conformer_57 | -990.1123648 | 0.3017910 | 0.2343140 | -990.1455240 | -87.1 | -989.843733 | -989.908198 | 7.7 | -984.4032709 | -4.2236629 | -988.4227671 |
| conformer_77 | -990.1107555 | 0.3015290 | 0.2348760 | -990.1455167 | -91.3 | -989.843988 | -989.907629 | 9.2 | -984.4000287 | -4.2244861 | -988.4213880 |
| conformer_125 | -990.1106054 | 0.3014160 | 0.2338630 | -990.1455083 | -91.6 | -989.844092 | -989.908633 | 6.6 | -984.3979927 | -4.2262295 | -988.4222502 |
| conformer_143 | -990.1093064 | 0.3012760 | 0.2332990 | -990.1455060 | -95.0 | -989.844230 | -989.909195 | 5.1 | -984.4009293 | -4.2230204 | -988.4238384 |
| conformer_155 | -990.1088564 | 0.3014530 | 0.2326860 | -990.1455012 | -96.2 | -989.844048 | -989.909803 | 3.5 | -984.4005602 | -4.2228306 | -988.4243376 |
| conformer_110 | -990.1067126 | 0.3013120 | 0.2331960 | -990.1453798 | -101.5 | -989.844068 | -989.909172 | 5.2 | -984.3970947 | -4.2235635 | -988.4231174 |
| conformer_148 | -990.1091041 | 0.3012710 | 0.2328100 | -990.1453311 | -95.1 | -989.844060 | -989.909509 | 4.3 | -984.4007946 | -4.2230373 | -988.4242370 |
| conformer_84 | -990.1105625 | 0.3015370 | 0.2346840 | -990.1453282 | -91.3 | -989.843791 | -989.907632 | 9.2 | -984.3999116 | -4.2244722 | -988.4214535 |
| conformer_9 | -990.1193799 | 0.3015290 | 0.2353540 | -990.1453250 | -68.1 | -989.843796 | -989.906959 | 11.0 | -984.4071552 | -4.2266634 | -988.4213977 |
| conformer_61 | -990.1106093 | 0.3015450 | 0.2341850 | -990.1453048 | -91.1 | -989.843760 | -989.908108 | 8.0 | -984.3976916 | -4.2264961 | -988.4216862 |
| conformer_11 | -990.1191970 | 0.3015560 | 0.2350900 | -990.1452720 | -68.5 | -989.843716 | -989.907170 | 10.4 | -984.4070792 | -4.2266407 | -988.4216929 |
| conformer_122 | -990.1065182 | 0.3013240 | 0.2329530 | -990.1452524 | -101.7 | -989.843928 | -989.909287 | 4.9 | -984.3969586 | -4.2235572 | -988.4232851 |
| conformer_163 | -990.1112460 | 0.3013890 | 0.2343220 | -990.1452482 | -89.3 | -989.843859 | -989.907914 | 8.5 | -984.4025136 | -4.2229855 | -988.4221673 |
| conformer_174 | -990.1056690 | 0.3012070 | 0.2326560 | -990.1452252 | -103.9 | -989.844018 | -989.909557 | 4.2 | -984.3960031 | -4.2235800 | -988.4234713 |
| conformer_63 | -990.1104299 | 0.3015550 | 0.2338000 | -990.1450944 | -91.0 | -989.843539 | -989.908282 | 7.5 | -984.3975995 | -4.2264366 | -988.4218886 |
| conformer_64 | -990.1104299 | 0.3015550 | 0.2338000 | -990.1450943 | -91.0 | -989.843539 | -989.908282 | 7.5 | -984.3975994 | -4.2264366 | -988.4218885 |
| conformer_186 | -990.1054781 | 0.3012150 | 0.2324170 | -990.1450929 | -104.0 | -989.843878 | -989.909664 | 3.9 | -984.3958668 | -4.2235609 | -988.4236136 |
| conformer_172 | -990.1085707 | 0.3015040 | 0.2340260 | -990.1450911 | -95.9 | -989.843587 | -989.908053 | 8.1 | -984.3995328 | -4.2234699 | -988.4224851 |
| conformer_169 | -990.1110503 | 0.3013970 | 0.2341520 | -990.1450391 | -89.2 | -989.843642 | -989.907875 | 8.6 | -984.4023845 | -4.2229981 | -988.4222073 |
| conformer_82 | -990.1120661 | 0.3016010 | 0.2357780 | -990.1450260 | -86.5 | -989.843425 | -989.906236 | 12.9 | -984.4013060 | -4.2254050 | -988.4208809 |
| conformer_72 | -990.1112096 | 0.3016650 | 0.2360120 | -990.1449817 | -88.7 | -989.843317 | -989.905958 | 13.6 | -984.3983492 | -4.2270822 | -988.4201795 |
| conformer_83 | -990.1112096 | 0.3016650 | 0.2360120 | -990.1449816 | -88.7 | -989.843317 | -989.905958 | 13.6 | -984.3983492 | -4.2270822 | -988.4201794 |
| conformer_4 | -990.1160204 | 0.3015050 | 0.2348450 | -990.1449292 | -75.9 | -989.843424 | -989.907072 | 10.7 | -984.4048498 | -4.2255001 | -988.4214017 |
| conformer_31 | -990.1151933 | 0.3013820 | 0.2346280 | -990.1448949 | -78.0 | -989.843513 | -989.907255 | 10.2 | -984.4035774 | -4.2258517 | -988.4214906 |
| conformer_7 | -990.1158327 | 0.3015460 | 0.2346650 | -990.1448855 | -76.3 | -989.843340 | -989.907209 | 10.3 | -984.4047260 | -4.2254919 | -988.4215937 |
| conformer_33 | -990.1150067 | 0.3014250 | 0.2344470 | -990.1448627 | -78.4 | -989.843438 | -989.907404 | 9.8 | -984.4034549 | -4.2258486 | -988.4217006 |
| conformer_48 | -990.1134961 | 0.3015550 | 0.2343340 | -990.1448564 | -82.3 | -989.843301 | -989.907510 | 9.6 | -984.4041130 | -4.2242709 | -988.4223981 |
| conformer_88 | -990.1109551 | 0.3016820 | 0.2357440 | -990.1447273 | -88.7 | -989.843045 | -989.905971 | 13.6 | -984.3981931 | -4.2270115 | -988.4202208 |
| conformer_53 | -990.1132734 | 0.3015790 | 0.2339190 | -990.1446742 | -82.4 | -989.843095 | -989.907743 | 8.9 | -984.4039533 | -4.2242679 | -988.4226910 |
| conformer_151 | -990.1082715 | 0.3012420 | 0.2325070 | -990.1444505 | -95.0 | -989.843209 | -989.908932 | 5.8 | -984.4010716 | -4.2218631 | -988.4235947 |
| conformer_150 | -990.1097811 | 0.3012930 | 0.2336740 | -990.1441840 | -90.3 | -989.842891 | -989.907498 | 9.6 | -984.4006386 | -4.2239770 | -988.4223325 |
| conformer_202 | -990.1065158 | 0.3012230 | 0.2328440 | -990.1441354 | -98.8 | -989.842912 | -989.908279 | 7.5 | -984.3975241 | -4.2232191 | -988.4225069 |
| conformer_60 | -990.1114614 | 0.3017660 | 0.2350450 | -990.1440696 | -85.6 | -989.842304 | -989.906013 | 13.5 | -984.4017724 | -4.2239622 | -988.4202858 |
| conformer_65 | -990.1112600 | 0.3017870 | 0.2349610 | -990.1439582 | -85.8 | -989.842171 | -989.905985 | 13.6 | -984.4016686 | -4.2239699 | -988.4203637 |
| conformer_43 | -990.1153390 | 0.3013990 | 0.2344630 | -990.1439154 | -75.0 | -989.842516 | -989.906440 | 12.4 | -984.4047124 | -4.2252810 | -988.4210948 |
| conformer_44 | -990.1151479 | 0.3014420 | 0.2343100 | -990.1438903 | -75.5 | -989.842448 | -989.906568 | 12.0 | -984.4045776 | -4.2252882 | -988.4212862 |
| conformer_91 | -990.1084223 | 0.3014880 | 0.2343950 | -990.1436203 | -92.4 | -989.842132 | -989.906213 | 13.0 | -984.3961155 | -4.2261409 | -988.4200473 |
| conformer_100 | -990.1083577 | 0.3015020 | 0.2342560 | -990.1435516 | -92.4 | -989.842050 | -989.906284 | 12.8 | -984.3960921 | -4.2261572 | -988.4201753 |
| conformer_185 | -990.1086448 | 0.3012650 | 0.2326930 | -990.1435412 | -91.6 | -989.842276 | -989.907836 | 8.7 | -984.4020539 | -4.2214327 | -988.4226779 |
| conformer_180 | -990.1061963 | 0.3013620 | 0.2333040 | -990.1435337 | -98.0 | -989.842172 | -989.907218 | 10.3 | -984.3939943 | -4.2263607 | -988.4213765 |
| conformer_192 | -990.1084413 | 0.3012730 | 0.2324110 | -990.1433859 | -91.7 | -989.842113 | -989.907963 | 8.4 | -984.4019080 | -4.2214542 | -988.4228838 |
| conformer_20 | -990.1158752 | 0.3016210 | 0.2342110 | -990.1432382 | -71.8 | -989.841617 | -989.906015 | 13.5 | -984.4065261 | -4.2245037 | -988.4211698 |
| conformer_15 | -990.1159845 | 0.3015900 | 0.2345190 | -990.1431671 | -71.4 | -989.841577 | -989.905636 | 14.5 | -984.4065271 | -4.2245668 | -988.4207455 |
| conformer_21 | -990.1155063 | 0.3016340 | 0.2346440 | -990.1429482 | -72.0 | -989.841314 | -989.905292 | 15.4 | -984.4073804 | -4.2234851 | -988.4206514 |
| conformer_16 | -990.1155819 | 0.3015860 | 0.2345950 | -990.1429294 | -71.8 | -989.841343 | -989.905322 | 15.3 | -984.4073591 | -4.2235220 | -988.4206216 |
| conformer_92 | -990.1076128 | 0.3014900 | 0.2347790 | -990.1428825 | -92.6 | -989.841392 | -989.905091 | 15.9 | -984.3962914 | -4.2254641 | -988.4192342 |
| conformer_128 | -990.1092989 | 0.3014100 | 0.2340060 | -990.1428561 | -88.1 | -989.841446 | -989.905838 | 13.9 | -984.4003440 | -4.2237256 | -988.4206088 |
| conformer_152 | -990.1103244 | 0.3013890 | 0.2337770 | -990.1427538 | -85.1 | -989.841365 | -989.905965 | 13.6 | -984.4009416 | -4.2239581 | -988.4205400 |
| conformer_121 | -990.1073199 | 0.3014720 | 0.2342210 | -990.1427051 | -92.9 | -989.841233 | -989.905472 | 14.9 | -984.3960919 | -4.2254293 | -988.4196734 |

| | | | | | | | | | | | |
|---|---|---|---|---|---|---|---|---|---|---|---|
| conformer_134 | -990.1081147 | 0.3015860 | 0.2341980 | -990.1426800 | -90.8 | -989.841094 | -989.905470 | 14.9 | -984.3975577 | -4.2246718 | -988.4195848 |
| conformer_158 | -990.1101305 | 0.3014050 | 0.2335710 | -990.1425844 | -85.2 | -989.841179 | -989.906001 | 13.5 | -984.4008084 | -4.2239800 | -988.4206593 |
| conformer_144 | -990.1079593 | 0.3016250 | 0.2342030 | -990.1425271 | -90.8 | -989.840902 | -989.905312 | 15.3 | -984.3974869 | -4.2246335 | -988.4194732 |
| conformer_70 | -990.1110861 | 0.3015880 | 0.2349470 | -990.1424620 | -82.4 | -989.840874 | -989.904503 | 17.4 | -984.4011234 | -4.2245364 | -988.4190767 |
| conformer_75 | -990.1108496 | 0.3016150 | 0.2347770 | -990.1423549 | -82.7 | -989.840740 | -989.904566 | 17.3 | -984.4009716 | -4.2245209 | -988.4192088 |
| conformer_56 | -990.1131769 | 0.3015670 | 0.2346880 | -990.1421973 | -76.2 | -989.840630 | -989.904497 | 17.5 | -984.4042630 | -4.2238169 | -988.4194003 |
| conformer_101 | -990.1076933 | 0.3014060 | 0.2346180 | -990.1421841 | -90.6 | -989.840778 | -989.904554 | 17.3 | -984.3966942 | -4.2252422 | -988.4187971 |
| conformer_37 | -990.1131581 | 0.3014170 | 0.2340920 | -990.1421000 | -76.0 | -989.840683 | -989.904996 | 16.2 | -984.4053886 | -4.2234061 | -988.4206326 |
| conformer_36 | -990.1132561 | 0.3013730 | 0.2341410 | -990.1420945 | -75.7 | -989.840721 | -989.904941 | 16.3 | -984.4053712 | -4.2234882 | -988.4205447 |
| conformer_59 | -990.1129542 | 0.3015840 | 0.2345550 | -990.1420593 | -76.4 | -989.840475 | -989.904492 | 17.5 | -984.4041354 | -4.2237902 | -988.4194638 |
| conformer_81 | -990.1104605 | 0.3014620 | 0.2344650 | -990.1420396 | -82.9 | -989.840578 | -989.904563 | 17.3 | -984.4021701 | -4.2231397 | -988.4194120 |
| conformer_89 | -990.1101544 | 0.3014560 | 0.2340330 | -990.1418499 | -83.2 | -989.840394 | -989.904805 | 16.7 | -984.4019401 | -4.2231476 | -988.4197382 |
| conformer_159 | -990.1106803 | 0.3013670 | 0.2336600 | -990.1416453 | -81.3 | -989.840278 | -989.904973 | 16.2 | -984.4021821 | -4.2235282 | -988.4200034 |
| conformer_137 | -990.1080656 | 0.3014060 | 0.2341710 | -990.1415539 | -87.9 | -989.840148 | -989.904371 | 17.8 | -984.3976162 | -4.2249143 | -988.4188359 |
| conformer_67 | -990.1130294 | 0.3013740 | 0.2340540 | -990.1415523 | -74.9 | -989.840178 | -989.904486 | 17.5 | -984.4057095 | -4.2231345 | -988.4203010 |
| conformer_167 | -990.1104846 | 0.3013820 | 0.2334570 | -990.1414736 | -81.4 | -989.840092 | -989.905005 | 16.1 | -984.4020464 | -4.2235382 | -988.4201045 |
| conformer_49 | -990.1138772 | 0.3015370 | 0.2343850 | -990.1411516 | -71.6 | -989.839615 | -989.903755 | 19.4 | -984.4056521 | -4.2235279 | -988.4190573 |
| conformer_85 | -990.1078214 | 0.3016560 | 0.2359160 | -990.1410160 | -87.2 | -989.839360 | -989.902088 | 23.8 | -984.3917390 | -4.2296109 | -988.4156165 |
| conformer_54 | -990.1136549 | 0.3015600 | 0.2342920 | -990.1409983 | -71.8 | -989.839438 | -989.903694 | 19.6 | -984.4055256 | -4.2235039 | -988.4190688 |
| conformer_2 | -990.1152727 | 0.3017790 | 0.2351800 | -990.1409531 | -67.4 | -989.839174 | -989.902761 | 22.0 | -984.4064470 | -4.2238155 | -988.4177508 |
| conformer_178 | -990.1075390 | 0.3015720 | 0.2356390 | -990.1409382 | -87.7 | -989.839366 | -989.902287 | 23.3 | -984.3906288 | -4.2308962 | -988.4162733 |
| conformer_1 | -990.1153402 | 0.3017450 | 0.2352530 | -990.1409187 | -67.2 | -989.839174 | -989.902654 | 22.3 | -984.4064351 | -4.2238398 | -988.4175884 |
| conformer_193 | -990.1079399 | 0.3014120 | 0.2347910 | -990.1393051 | -82.3 | -989.837893 | -989.901502 | 25.3 | -984.3959091 | -4.2271397 | -988.4166110 |
| conformer_161 | -990.1063032 | 0.3014210 | 0.2356170 | -990.1378944 | -82.9 | -989.836473 | -989.899265 | 31.2 | -984.3910168 | -4.2298125 | -988.4137915 |
| conformer_189 | -990.1027248 | 0.3015650 | 0.2341610 | -990.1373023 | -90.8 | -989.835737 | -989.900129 | 28.9 | -984.3880212 | -4.2292092 | -988.4146348 |

[a]: Single-point calculation in aqueous phase with SMD model.

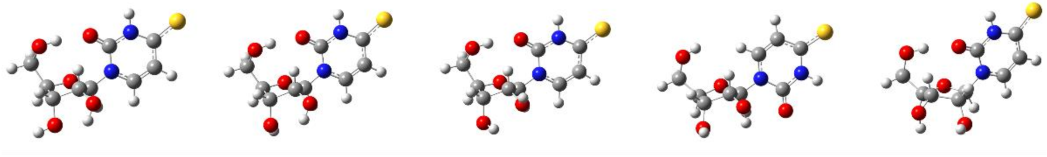

**Figure S40.** B3LYP-D3/def2-TZVPP optimized geometries of conformers for 4-thiouridine (**1rb4thU**).

**Table S38.** Conformers of gas-phase optimized 4-thiouridine (**1rb4thU**) at the B3LYP-D3/def2-TZVPP level of theory followed by aqueous phase single-point calculation. The columns display total energy without zero-point correction ($E_{Tot}$), thermal correction to enthalpy (δH), Gibbs free energy (δG), total energy without zero-point correction ($E_{Tot,W}$), Gibbs free energy ($G_{298,W}$) in water (W), total single-point energy ($E_{CBS}$) calculated at DLPNO-CCSD(T)/CBS level of theory, and their corresponding free energy $G_{CBS,W}$. $G_{298,W}$ and $G_{CBS,W}$ have been corrected to the standard state of 1 mol/L by addition of +7.908 kJ/mol. $\Delta G_{Solv}$ represents the Gibbs free energy of solvation. The data are arranged in the ascending numeric order of $E_{Tot,W}$. $\Delta G_{298,W}$ represents the respective energy difference to the lowest structure. Only conformers within the 24 kJ/mol (6 kcal/mol) energy window above the lowest in CREST are included in initial conformer sampling. Duplicates of the same structure are excluded. The overall optimum is marked bold.

| | B3LYP-D3/def2-TZVPP | | | SMD($H_2O$)/B3LYP-D3/def2-TZVPP[a] | | | | | DLPNO-CCSD(T)/CBS | | |
|---|---|---|---|---|---|---|---|---|---|---|---|
| 1rb4thU No. | $E_{Tot}$ (Hartree) | δH (Hartree) | δG (Hartree) | $E_{Tot,W}$ (Hartree) | $\Delta G_{Solv}$ (kJ/mol) | $H_{298,W}$ (Hartree) | $G_{298,W}$ (Hartree) | $\Delta G_{298,W}$ (kJ/mol) | $E_{CBS,HF}$ (Hartree) | $E_{CBS,C}$ (Hartree) | $G_{CBS,W}$ (Hartree) |
| conformer_18 | -1234.407324 | 0.241368 | 0.182161 | -1234.440501 | -87.1 | -1234.199133 | -1234.255328 | 5.6 | -1228.945926 | -3.756954 | -1232.550884 |
| conformer_5 | -1234.406673 | 0.241259 | 0.181818 | -1234.440438 | -88.6 | -1234.199179 | -1234.255608 | 4.9 | -1228.945225 | -3.756755 | -1232.550915 |
| conformer_8 | -1234.404403 | 0.241229 | 0.181767 | -1234.440388 | -94.5 | -1234.199159 | -1234.255609 | 4.9 | -1228.941051 | -3.758138 | -1232.550395 |
| conformer_38 | -1234.403474 | 0.241231 | 0.181190 | -1234.439258 | -93.9 | -1234.198027 | -1234.255056 | 6.3 | -1228.942012 | -3.756536 | -1232.550129 |
| **conformer_34** | **-1234.400951** | **0.241234** | **0.178759** | **-1234.439243** | **-100.5** | **-1234.198009** | **-1234.257472** | **0.0** | **-1228.942400** | **-3.754214** | **-1232.553135** |
| conformer_19 | -1234.405427 | 0.241211 | 0.180841 | -1234.439135 | -88.5 | -1234.197924 | -1234.255282 | 5.8 | -1228.946233 | -3.754522 | -1232.550610 |
| conformer_23 | -1234.404055 | 0.241182 | 0.179636 | -1234.438883 | -91.4 | -1234.197701 | -1234.256235 | 3.2 | -1228.944709 | -3.754638 | -1232.551527 |
| conformer_67 | -1234.398583 | 0.241043 | 0.179253 | -1234.438526 | -104.9 | -1234.197483 | -1234.256261 | 3.2 | -1228.938400 | -3.755683 | -1232.551761 |
| conformer_7 | -1234.407212 | 0.241043 | 0.181070 | -1234.438483 | -82.1 | -1234.197440 | -1234.254401 | 8.1 | -1228.944174 | -3.758469 | -1232.549832 |
| conformer_112 | -1234.398839 | 0.240877 | 0.179726 | -1234.438463 | -104.0 | -1234.197586 | -1234.255725 | 4.6 | -1228.939309 | -3.755377 | -1232.551572 |
| conformer_12 | -1234.405723 | 0.241200 | 0.181438 | -1234.438416 | -85.8 | -1234.197216 | -1234.253966 | 9.2 | -1228.944247 | -3.756909 | -1232.549399 |
| conformer_3 | -1234.406160 | 0.241057 | 0.181328 | -1234.438291 | -84.4 | -1234.197234 | -1234.253951 | 9.2 | -1228.944669 | -3.757300 | -1232.549759 |
| conformer_75 | -1234.401683 | 0.241164 | 0.180102 | -1234.438211 | -95.9 | -1234.197047 | -1234.255097 | 6.2 | -1228.940776 | -3.756290 | -1232.550480 |
| conformer_36 | -1234.398876 | 0.240994 | 0.180036 | -1234.438170 | -103.2 | -1234.197176 | -1234.255122 | 6.2 | -1228.937824 | -3.755739 | -1232.549808 |
| conformer_21 | -1234.405916 | 0.241188 | 0.180724 | -1234.438139 | -84.6 | -1234.196951 | -1234.254403 | 8.1 | -1228.947634 | -3.754004 | -1232.550125 |
| conformer_6 | -1234.407120 | 0.241018 | 0.180665 | -1234.438091 | -81.3 | -1234.197073 | -1234.254414 | 8.0 | -1228.945156 | -3.757224 | -1232.549675 |
| conformer_60 | -1234.397448 | 0.240897 | 0.179149 | -1234.438072 | -106.7 | -1234.197175 | -1234.255911 | 4.1 | -1228.937294 | -3.755478 | -1232.551236 |
| conformer_77 | -1234.397710 | 0.240885 | 0.179502 | -1234.438066 | -106.0 | -1234.197181 | -1234.255552 | 5.0 | -1228.936332 | -3.756681 | -1232.550855 |
| conformer_81 | -1234.399349 | 0.240885 | 0.179697 | -1234.438017 | -101.5 | -1234.197132 | -1234.255308 | 5.7 | -1228.940329 | -3.754179 | -1232.550467 |
| conformer_64 | -1234.400023 | 0.240965 | 0.180137 | -1234.437944 | -99.6 | -1234.196979 | -1234.254795 | 7.0 | -1228.938222 | -3.756671 | -1232.549665 |
| conformer_31 | -1234.400573 | 0.241098 | 0.180525 | -1234.437815 | -97.8 | -1234.196717 | -1234.254278 | 8.4 | -1228.940393 | -3.755216 | -1232.549314 |
| conformer_27 | -1234.405384 | 0.241001 | 0.181383 | -1234.437786 | -85.1 | -1234.196785 | -1234.253391 | 10.7 | -1228.944335 | -3.757326 | -1232.549668 |
| conformer_30 | -1234.401252 | 0.241218 | 0.180058 | -1234.437707 | -95.7 | -1234.196489 | -1234.254637 | 7.4 | -1228.942663 | -3.754432 | -1232.550479 |
| conformer_71 | -1234.398526 | 0.240931 | 0.179232 | -1234.437646 | -102.7 | -1234.196715 | -1234.255402 | 5.4 | -1228.940484 | -3.753892 | -1232.551252 |
| conformer_13 | -1234.405100 | 0.240949 | 0.180413 | -1234.437606 | -85.3 | -1234.196657 | -1234.254181 | 8.6 | -1228.943484 | -3.757050 | -1232.549615 |
| conformer_80 | -1234.398039 | 0.241027 | 0.178527 | -1234.437579 | -103.8 | -1234.196552 | -1234.256040 | 3.8 | -1228.940019 | -3.753729 | -1232.551749 |
| conformer_56 | -1234.401341 | 0.240964 | 0.179925 | -1234.437532 | -95.0 | -1234.196568 | -1234.254595 | 7.6 | -1228.943268 | -3.753641 | -1232.550163 |
| conformer_1 | -1234.408726 | 0.241126 | 0.181353 | -1234.437510 | -75.6 | -1234.196384 | -1234.253145 | 11.4 | -1228.947598 | -3.756959 | -1232.548976 |
| conformer_43 | -1234.399445 | 0.241108 | 0.179957 | -1234.437375 | -99.6 | -1234.196267 | -1234.254406 | 8.0 | -1228.937046 | -3.757184 | -1232.549191 |
| conformer_54 | -1234.395874 | 0.240864 | 0.178880 | -1234.437234 | -108.6 | -1234.196370 | -1234.255342 | 5.6 | -1228.936660 | -3.754429 | -1232.550558 |

| | | | | | | | | | | | |
|---|---|---|---|---|---|---|---|---|---|---|---|
| conformer_97 | -1234.397410 | 0.241015 | 0.179631 | -1234.437100 | -104.2 | -1234.196085 | -1234.254457 | 7.9 | -1228.938938 | -3.754279 | -1232.550264 |
| conformer_40 | -1234.400927 | 0.240983 | 0.180789 | -1234.437084 | -94.9 | -1234.196101 | -1234.253283 | 11.0 | -1228.941005 | -3.755822 | -1232.549183 |
| conformer_94 | -1234.394768 | 0.240683 | 0.178160 | -1234.437058 | -111.0 | -1234.196375 | -1234.255886 | 4.2 | -1228.935571 | -3.754468 | -1232.551158 |
| conformer_46 | -1234.399797 | 0.241042 | 0.181024 | -1234.436946 | -97.5 | -1234.195904 | -1234.252910 | 12.0 | -1228.937740 | -3.757498 | -1232.548351 |
| conformer_62 | -1234.400256 | 0.240903 | 0.179613 | -1234.436940 | -96.3 | -1234.196037 | -1234.254315 | 8.3 | -1228.939547 | -3.755973 | -1232.549578 |
| conformer_15 | -1234.403872 | 0.241045 | 0.180581 | -1234.436923 | -86.8 | -1234.195878 | -1234.253330 | 10.9 | -1228.943041 | -3.756339 | -1232.548838 |
| conformer_4 | -1234.404786 | 0.241142 | 0.180683 | -1234.436890 | -84.3 | -1234.195748 | -1234.253195 | 11.2 | -1228.944434 | -3.755977 | -1232.548820 |
| conformer_26 | -1234.402388 | 0.241074 | 0.179911 | -1234.436775 | -90.3 | -1234.195701 | -1234.253852 | 9.5 | -1228.943623 | -3.754967 | -1232.550054 |
| conformer_109 | -1234.396227 | 0.240670 | 0.177593 | -1234.436539 | -105.8 | -1234.195869 | -1234.255934 | 4.0 | -1228.939050 | -3.752898 | -1232.551655 |
| conformer_69 | -1234.397801 | 0.240805 | 0.178389 | -1234.436471 | -101.5 | -1234.195666 | -1234.255070 | 6.3 | -1228.940968 | -3.752743 | -1232.550981 |
| conformer_86 | -1234.398825 | 0.240860 | 0.179438 | -1234.436152 | -98.0 | -1234.195292 | -1234.253702 | 9.9 | -1228.940246 | -3.754703 | -1232.549827 |
| conformer_101 | -1234.395183 | 0.240886 | 0.179240 | -1234.436122 | -107.5 | -1234.195236 | -1234.253870 | 9.5 | -1228.933774 | -3.756833 | -1232.549295 |
| conformer_98 | -1234.395781 | 0.240764 | 0.178442 | -1234.436044 | -105.7 | -1234.195280 | -1234.254590 | 7.6 | -1228.937233 | -3.754057 | -1232.550099 |
| conformer_33 | -1234.400195 | 0.241266 | 0.180435 | -1234.436029 | -94.1 | -1234.194763 | -1234.252582 | 12.8 | -1228.940958 | -3.754846 | -1232.548191 |
| conformer_58 | -1234.397235 | 0.240961 | 0.179852 | -1234.435956 | -101.7 | -1234.194995 | -1234.253092 | 11.5 | -1228.935450 | -3.756824 | -1232.548130 |
| conformer_24 | -1234.404222 | 0.241019 | 0.180183 | -1234.435797 | -82.9 | -1234.194778 | -1234.252602 | 12.8 | -1228.944465 | -3.755727 | -1232.548572 |
| conformer_85 | -1234.398244 | 0.240798 | 0.178352 | -1234.435550 | -97.9 | -1234.194752 | -1234.254186 | 8.6 | -1228.942068 | -3.752291 | -1232.550301 |
| conformer_51 | -1234.399021 | 0.241021 | 0.180142 | -1234.435523 | -95.8 | -1234.194502 | -1234.252369 | 13.4 | -1228.940576 | -3.754325 | -1232.548249 |
| conformer_53 | -1234.396739 | 0.241057 | 0.180372 | -1234.435422 | -101.6 | -1234.194365 | -1234.252038 | 14.3 | -1228.935954 | -3.756087 | -1232.547341 |
| conformer_10 | -1234.404528 | 0.241137 | 0.180148 | -1234.435418 | -81.1 | -1234.194281 | -1234.252258 | 13.7 | -1228.945666 | -3.755150 | -1232.548545 |
| conformer_9 | -1234.404148 | 0.241115 | 0.180206 | -1234.435224 | -81.6 | -1234.194109 | -1234.252006 | 14.4 | -1228.946419 | -3.754168 | -1232.548445 |
| conformer_72 | -1234.399617 | 0.240885 | 0.178330 | -1234.434748 | -92.2 | -1234.193863 | -1234.253406 | 10.7 | -1228.940686 | -3.754775 | -1232.549250 |
| conformer_41 | -1234.399998 | 0.241151 | 0.180518 | -1234.434598 | -90.8 | -1234.193447 | -1234.251068 | 16.8 | -1228.940484 | -3.755326 | -1232.546879 |
| conformer_39 | -1234.399978 | 0.241090 | 0.180252 | -1234.434502 | -90.6 | -1234.193412 | -1234.251238 | 16.4 | -1228.942307 | -3.753718 | -1232.547285 |
| conformer_20 | -1234.402002 | 0.240964 | 0.179926 | -1234.434422 | -85.1 | -1234.193458 | -1234.251484 | 15.7 | -1228.944744 | -3.753970 | -1232.548197 |
| conformer_28 | -1234.402292 | 0.241102 | 0.180156 | -1234.434388 | -84.3 | -1234.193286 | -1234.251220 | 16.4 | -1228.943816 | -3.754640 | -1232.547384 |
| conformer_73 | -1234.396246 | 0.240977 | 0.180327 | -1234.434252 | -99.8 | -1234.193275 | -1234.250913 | 17.2 | -1228.935455 | -3.756198 | -1232.546320 |
| conformer_70 | -1234.397303 | 0.240937 | 0.179218 | -1234.433931 | -96.2 | -1234.192994 | -1234.251701 | 15.2 | -1228.937534 | -3.755409 | -1232.547342 |
| conformer_42 | -1234.402020 | 0.240931 | 0.179886 | -1234.433831 | -83.5 | -1234.192900 | -1234.250933 | 17.2 | -1228.945219 | -3.753673 | -1232.547805 |
| conformer_89 | -1234.400075 | 0.240877 | 0.178179 | -1234.433686 | -88.2 | -1234.192809 | -1234.252495 | 13.1 | -1228.942015 | -3.754376 | -1232.548810 |
| conformer_25 | -1234.403143 | 0.241180 | 0.180250 | -1234.433398 | -79.4 | -1234.192218 | -1234.250136 | 19.3 | -1228.945375 | -3.754267 | -1232.546636 |
| conformer_2 | -1234.403962 | 0.241328 | 0.181043 | -1234.433340 | -77.1 | -1234.192012 | -1234.249285 | 21.5 | -1228.945527 | -3.754499 | -1232.545349 |
| conformer_108 | -1234.393673 | 0.240884 | 0.177391 | -1234.433223 | -103.8 | -1234.192339 | -1234.252820 | 12.2 | -1228.935920 | -3.753508 | -1232.548574 |
| conformer_105 | -1234.396284 | 0.241138 | 0.181311 | -1234.433088 | -96.6 | -1234.191950 | -1234.248765 | 22.9 | -1228.930083 | -3.761360 | -1232.543925 |
| conformer_52 | -1234.396483 | 0.241159 | 0.181668 | -1234.432842 | -95.5 | -1234.191683 | -1234.248162 | 24.4 | -1228.930865 | -3.760429 | -1232.542973 |
| conformer_78 | -1234.398160 | 0.240988 | 0.180437 | -1234.431664 | -88.0 | -1234.190676 | -1234.248215 | 24.3 | -1228.936890 | -3.757643 | -1232.544589 |
| conformer_106 | -1234.396213 | 0.240873 | 0.179337 | -1234.430780 | -90.8 | -1234.189907 | -1234.248431 | 23.7 | -1228.934521 | -3.757718 | -1232.544457 |
| conformer_68 | -1234.396247 | 0.241042 | 0.181226 | -1234.430643 | -90.3 | -1234.189601 | -1234.246405 | 29.1 | -1228.931765 | -3.760230 | -1232.542153 |
| conformer_107 | -1234.392075 | 0.240995 | 0.179880 | -1234.430140 | -99.9 | -1234.189145 | -1234.247248 | 26.8 | -1228.928415 | -3.759610 | -1232.543199 |
| conformer_95 | -1234.391921 | 0.241476 | 0.181851 | -1234.425973 | -89.4 | -1234.184497 | -1234.241110 | 43.0 | -1228.926010 | -3.761630 | -1232.536829 |

[a]: Single-point calculation in aqueous phase with SMD model.

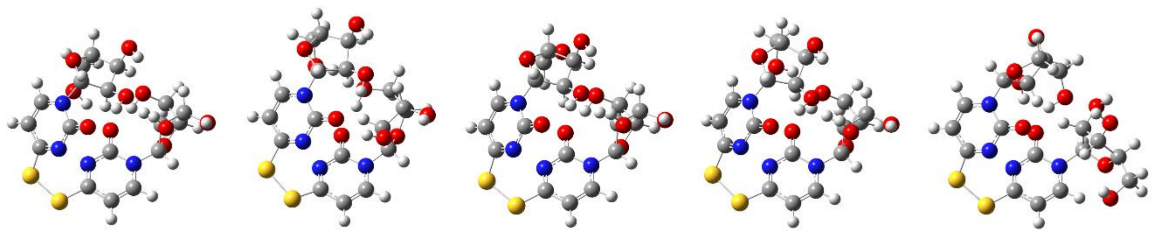

conformer_5 conformer_7 conformer_8 conformer_10 conformer_29

**Figure S41.** B3LYP-D3/def2-TZVPP optimized geometries of conformers for 4-thiouridine dimer (**1rb4thU_dimer**).

**Table S39.** Conformers of gas-phase optimized 4-thiouridine dimer (**1rb4thU_dimer**) at the B3LYP-D3/def2-TZVPP level of theory followed by aqueous phase single-point calculation. The columns display total energy without zero-point correction ($E_{Tot}$), thermal correction to enthalpy (δH), Gibbs free energy (δG), total energy without zero-point correction ($E_{Tot,W}$), Gibbs free energy ($G_{298,W}$) in water (W), total single-point energy ($E_{CBS}$) calculated at DLPNO-CCSD(T)/CBS level of theory, and their corresponding free energy $G_{CBS,W}$. $G_{298,W}$ and $G_{CBS,W}$ have been corrected to the standard state of 1 mol/L by addition of +7.908 kJ/mol. $\Delta G_{Solv}$ represents the Gibbs free energy of solvation. The data are arranged in the ascending numeric order of $E_{Tot,W}$. $\Delta G_{298,W}$ represents the respective energy difference to the lowest structure. Only conformers within the 24 kJ/mol (6 kcal/mol) energy window above the lowest in CREST are included in initial conformer sampling. Duplicates of the same structure are excluded. The overall optimum is marked bold.

| | B3LYP-D3/def2-TZVPP | | | SMD($H_2O$)/B3LYP-D3/def2-TZVPP[a] | | | | | DLPNO-CCSD(T)/CBS | | |
|---|---|---|---|---|---|---|---|---|---|---|---|
| 1rb4thU_dimer No. | $E_{Tot}$ (Hartree) | δH (Hartree) | δG (Hartree) | $E_{Tot,W}$ (Hartree) | $\Delta G_{Solv}$ (kJ/mol) | $H_{298,W}$ (Hartree) | $G_{298,W}$ (Hartree) | $\Delta G_{298,W}$ (kJ/mol) | $E_{CBS,HF}$ (Hartree) | $E_{CBS,C}$ (Hartree) | $G_{CBS,W}$ (Hartree) |
| conformer_5 | -2467.604289 | 0.459548 | 0.369779 | -2467.653235 | -128.5 | -2467.193687 | -2467.280444 | 7.5 | -2456.679038 | -7.519464 | -2463.874657 |
| conformer_7 | -2467.604109 | 0.459109 | 0.369192 | -2467.653225 | -129.0 | -2467.194116 | -2467.281021 | 6.0 | -2456.678848 | -7.519271 | -2463.875032 |
| **conformer_8** | **-2467.604035** | **0.459416** | **0.368826** | **-2467.655144** | **-134.2** | **-2467.195728** | **-2467.283306** | **0.0** | **-2456.681382** | **-7.517645** | **-2463.878299** |
| conformer_10 | -2467.602774 | 0.459049 | 0.367842 | -2467.651316 | -127.4 | -2467.192267 | -2467.280462 | 7.5 | -2456.681478 | -7.516348 | -2463.875515 |
| conformer_29 | -2467.602335 | 0.459231 | 0.367049 | -2467.652630 | -132.0 | -2467.193399 | -2467.282569 | 1.9 | -2456.680866 | -7.515477 | -2463.876577 |
| conformer_34 | -2467.601902 | 0.459217 | 0.367015 | -2467.651025 | -129.0 | -2467.191808 | -2467.280998 | 6.1 | -2456.681404 | -7.515108 | -2463.875608 |
| conformer_4 | -2467.601151 | 0.459290 | 0.367392 | -2467.651139 | -131.2 | -2467.191849 | -2467.280735 | 6.8 | -2456.681722 | -7.513477 | -2463.874783 |
| conformer_6 | -2467.601133 | 0.459380 | 0.367908 | -2467.652027 | -133.6 | -2467.192647 | -2467.281107 | 5.8 | -2456.680509 | -7.514130 | -2463.874612 |
| conformer_1 | -2467.601024 | 0.459300 | 0.367031 | -2467.652780 | -135.9 | -2467.193480 | -2467.282737 | 1.5 | -2456.680467 | -7.514495 | -2463.876675 |
| conformer_28 | -2467.601006 | 0.459326 | 0.368763 | -2467.651752 | -133.2 | -2467.192426 | -2467.279977 | 8.7 | -2456.676279 | -7.519115 | -2463.874366 |
| conformer_30 | -2467.599633 | 0.459310 | 0.366920 | -2467.651913 | -137.3 | -2467.192603 | -2467.281981 | 3.5 | -2456.679293 | -7.514699 | -2463.876340 |
| conformer_43 | -2467.599633 | 0.459310 | 0.366920 | -2467.651913 | -137.3 | -2467.192603 | -2467.281981 | 3.5 | -2456.679294 | -7.514699 | -2463.876340 |
| conformer_21 | -2467.599523 | 0.459372 | 0.367270 | -2467.649693 | -131.7 | -2467.190321 | -2467.279411 | 10.2 | -2456.678351 | -7.515376 | -2463.873614 |
| conformer_12 | -2467.599090 | 0.459028 | 0.367391 | -2467.647438 | -126.9 | -2467.188410 | -2467.277035 | 16.5 | -2456.676724 | -7.516623 | -2463.871292 |
| conformer_81 | -2467.599090 | 0.459028 | 0.367391 | -2467.647438 | -126.9 | -2467.188410 | -2467.277035 | 16.5 | -2456.676724 | -7.516623 | -2463.871292 |
| conformer_3 | -2467.598798 | 0.459072 | 0.365974 | -2467.651959 | -139.6 | -2467.192887 | -2467.282973 | 0.9 | -2456.680622 | -7.512432 | -2463.877230 |
| conformer_2 | -2467.598495 | 0.459180 | 0.366229 | -2467.649643 | -134.3 | -2467.190463 | -2467.280402 | 7.6 | -2456.679236 | -7.513240 | -2463.874383 |
| conformer_78 | -2467.597974 | 0.459285 | 0.368486 | -2467.653214 | -145.0 | -2467.193929 | -2467.281716 | 4.2 | -2456.675558 | -7.517490 | -2463.876788 |
| conformer_15 | -2467.597954 | 0.459418 | 0.367939 | -2467.648582 | -132.9 | -2467.189164 | -2467.277631 | 14.9 | -2456.674921 | -7.517632 | -2463.872229 |
| conformer_45 | -2467.597812 | 0.459591 | 0.368680 | -2467.650935 | -139.5 | -2467.191344 | -2467.279243 | 10.7 | -2456.676396 | -7.516958 | -2463.874784 |
| conformer_16 | -2467.597552 | 0.459325 | 0.368445 | -2467.646255 | -127.9 | -2467.186930 | -2467.274798 | 22.3 | -2456.671623 | -7.519597 | -2463.868467 |
| conformer_70 | -2467.597364 | 0.459004 | 0.365783 | -2467.651551 | -142.3 | -2467.192547 | -2467.282756 | 1.4 | -2456.677676 | -7.514057 | -2463.877125 |
| conformer_90 | -2467.597117 | 0.459417 | 0.369518 | -2467.651116 | -141.8 | -2467.191699 | -2467.278586 | 12.4 | -2456.671565 | -7.519509 | -2463.872542 |
| conformer_26 | -2467.596928 | 0.458913 | 0.367351 | -2467.649439 | -137.9 | -2467.190526 | -2467.279076 | 11.1 | -2456.677575 | -7.513638 | -2463.873361 |
| conformer_17 | -2467.596808 | 0.459358 | 0.367467 | -2467.648235 | -135.0 | -2467.188877 | -2467.277756 | 14.6 | -2456.676413 | -7.515628 | -2463.872989 |
| conformer_93 | -2467.596486 | 0.459590 | 0.368099 | -2467.649165 | -138.3 | -2467.189575 | -2467.278054 | 13.8 | -2456.673442 | -7.516830 | -2463.871839 |
| conformer_54 | -2467.596412 | 0.459251 | 0.367127 | -2467.648596 | -137.0 | -2467.189345 | -2467.278457 | 12.7 | -2456.676338 | -7.515158 | -2463.873539 |

| | | | | | | | | | | | |
|---|---|---|---|---|---|---|---|---|---|---|---|
| conformer_47 | -2467.595961 | 0.459399 | 0.366843 | -2467.645306 | -129.6 | -2467.185907 | -2467.275451 | 20.6 | -2456.680042 | -7.511171 | -2463.870704 |
| conformer_24 | -2467.595808 | 0.458562 | 0.365532 | -2467.650442 | -143.4 | -2467.191880 | -2467.281898 | 3.7 | -2456.679569 | -7.511062 | -2463.876720 |
| conformer_22 | -2467.595657 | 0.458875 | 0.365313 | -2467.651206 | -145.8 | -2467.192331 | -2467.282881 | 1.1 | -2456.677730 | -7.512270 | -2463.877224 |
| conformer_55 | -2467.595544 | 0.458845 | 0.365394 | -2467.649271 | -141.1 | -2467.190426 | -2467.280865 | 6.4 | -2456.681550 | -7.509550 | -2463.876420 |
| conformer_52 | -2467.595527 | 0.458897 | 0.365548 | -2467.650325 | -143.9 | -2467.191428 | -2467.281765 | 4.0 | -2456.680436 | -7.510244 | -2463.876918 |
| conformer_73 | -2467.595305 | 0.458830 | 0.365192 | -2467.650600 | -145.2 | -2467.191770 | -2467.282396 | 2.4 | -2456.677997 | -7.511987 | -2463.877075 |
| conformer_25 | -2467.594663 | 0.458609 | 0.364375 | -2467.641843 | -123.9 | -2467.183234 | -2467.274456 | 23.2 | -2456.681591 | -7.509830 | -2463.871215 |
| conformer_18 | -2467.594299 | 0.458602 | 0.364332 | -2467.642846 | -127.5 | -2467.184244 | -2467.275502 | 20.5 | -2456.680409 | -7.510239 | -2463.871851 |
| conformer_89 | -2467.594286 | 0.459476 | 0.367465 | -2467.648716 | -142.9 | -2467.189240 | -2467.278239 | 13.3 | -2456.672077 | -7.516367 | -2463.872396 |
| conformer_98 | -2467.594132 | 0.458933 | 0.364751 | -2467.650208 | -147.2 | -2467.191275 | -2467.282445 | 2.3 | -2456.685965 | -7.504814 | -2463.879092 |
| conformer_65 | -2467.594099 | 0.459300 | 0.366937 | -2467.648669 | -143.3 | -2467.189369 | -2467.278720 | 12.0 | -2456.671203 | -7.517738 | -2463.873562 |
| conformer_75 | -2467.593950 | 0.459576 | 0.367848 | -2467.646891 | -139.0 | -2467.187315 | -2467.276031 | 19.1 | -2456.671160 | -7.516717 | -2463.869958 |
| conformer_33 | -2467.593937 | 0.458811 | 0.364830 | -2467.640932 | -123.4 | -2467.182121 | -2467.273090 | 26.8 | -2456.675721 | -7.513967 | -2463.868842 |
| conformer_48 | -2467.593893 | 0.459021 | 0.365866 | -2467.650654 | -149.0 | -2467.191633 | -2467.281776 | 4.0 | -2456.678523 | -7.510557 | -2463.876963 |
| conformer_72 | -2467.593535 | 0.459156 | 0.366451 | -2467.647404 | -141.4 | -2467.188248 | -2467.277941 | 14.1 | -2456.677058 | -7.511559 | -2463.873024 |
| conformer_84 | -2467.593435 | 0.459208 | 0.367748 | -2467.645058 | -135.5 | -2467.185850 | -2467.274298 | 23.7 | -2456.669672 | -7.517576 | -2463.868111 |
| conformer_59 | -2467.593288 | 0.458739 | 0.365720 | -2467.649863 | -148.5 | -2467.191124 | -2467.281131 | 5.7 | -2456.676638 | -7.512485 | -2463.876965 |
| conformer_37 | -2467.592951 | 0.458882 | 0.365288 | -2467.647694 | -143.7 | -2467.188812 | -2467.279394 | 10.3 | -2456.679314 | -7.509046 | -2463.874803 |
| conformer_67 | -2467.592816 | 0.458664 | 0.364602 | -2467.649659 | -149.2 | -2467.190995 | -2467.282045 | 3.3 | -2456.679934 | -7.508525 | -2463.877688 |
| conformer_41 | -2467.592138 | 0.458500 | 0.363812 | -2467.642904 | -133.3 | -2467.184404 | -2467.276080 | 19.0 | -2456.678018 | -7.510411 | -2463.872371 |
| conformer_91 | -2467.592032 | 0.458716 | 0.364113 | -2467.648113 | -147.2 | -2467.189397 | -2467.280988 | 6.1 | -2456.674876 | -7.511619 | -2463.875451 |
| conformer_86 | -2467.591974 | 0.458875 | 0.368517 | -2467.648512 | -148.4 | -2467.189637 | -2467.276983 | 16.6 | -2456.664808 | -7.520857 | -2463.870674 |
| conformer_64 | -2467.591752 | 0.459152 | 0.366091 | -2467.644244 | -137.8 | -2467.185092 | -2467.275141 | 21.4 | -2456.671478 | -7.515540 | -2463.870407 |
| conformer_97 | -2467.591632 | 0.458514 | 0.365149 | -2467.646213 | -143.3 | -2467.187699 | -2467.278052 | 13.8 | -2456.677057 | -7.511101 | -2463.874579 |
| conformer_27 | -2467.591496 | 0.458434 | 0.363137 | -2467.642389 | -133.6 | -2467.183955 | -2467.276240 | 18.6 | -2456.679224 | -7.509046 | -2463.873012 |
| conformer_14 | -2467.591451 | 0.458579 | 0.363970 | -2467.640164 | -127.9 | -2467.181585 | -2467.273182 | 26.6 | -2456.678842 | -7.509270 | -2463.869842 |
| conformer_77 | -2467.591263 | 0.459088 | 0.366111 | -2467.640784 | -130.0 | -2467.181696 | -2467.271661 | 30.6 | -2456.672001 | -7.514317 | -2463.866715 |
| conformer_94 | -2467.591124 | 0.459147 | 0.367328 | -2467.642190 | -134.1 | -2467.183043 | -2467.271850 | 30.1 | -2456.668252 | -7.518722 | -2463.867701 |
| conformer_53 | -2467.590905 | 0.458465 | 0.363841 | -2467.643764 | -138.8 | -2467.185299 | -2467.276911 | 16.8 | -2456.675823 | -7.510834 | -2463.872663 |
| conformer_69 | -2467.590731 | 0.458961 | 0.367485 | -2467.643135 | -137.6 | -2467.184174 | -2467.272638 | 28.0 | -2456.667202 | -7.518115 | -2463.867224 |
| conformer_61 | -2467.590563 | 0.459390 | 0.365215 | -2467.644002 | -140.3 | -2467.184612 | -2467.275775 | 19.8 | -2456.677221 | -7.509997 | -2463.872430 |
| conformer_95 | -2467.590339 | 0.458711 | 0.366204 | -2467.649972 | -156.6 | -2467.191261 | -2467.280756 | 6.7 | -2456.672456 | -7.513255 | -2463.876128 |
| conformer_58 | -2467.590197 | 0.458466 | 0.364178 | -2467.644138 | -141.6 | -2467.185672 | -2467.276948 | 16.7 | -2456.671873 | -7.513695 | -2463.872319 |
| conformer_100 | -2467.590155 | 0.459303 | 0.366225 | -2467.645346 | -144.9 | -2467.186043 | -2467.276109 | 18.9 | -2456.675078 | -7.511276 | -2463.872309 |
| conformer_44 | -2467.589562 | 0.459425 | 0.365642 | -2467.642119 | -138.0 | -2467.182694 | -2467.273465 | 25.8 | -2456.676074 | -7.509935 | -2463.869912 |
| conformer_71 | -2467.589537 | 0.459346 | 0.366578 | -2467.642917 | -140.1 | -2467.183571 | -2467.273327 | 26.2 | -2456.667501 | -7.515425 | -2463.866716 |
| conformer_35 | -2467.589378 | 0.458303 | 0.362503 | -2467.641557 | -137.0 | -2467.183254 | -2467.276042 | 19.1 | -2456.677749 | -7.508465 | -2463.872878 |
| conformer_36 | -2467.589097 | 0.458850 | 0.366823 | -2467.642268 | -139.6 | -2467.183418 | -2467.272433 | 28.5 | -2456.666272 | -7.518139 | -2463.867747 |
| conformer_57 | -2467.588461 | 0.458195 | 0.363342 | -2467.639669 | -134.4 | -2467.181474 | -2467.273315 | 26.2 | -2456.672620 | -7.512545 | -2463.870020 |
| conformer_46 | -2467.588185 | 0.458330 | 0.362681 | -2467.642362 | -142.2 | -2467.184032 | -2467.276669 | 17.4 | -2456.673210 | -7.510860 | -2463.872554 |
| conformer_62 | -2467.587610 | 0.458555 | 0.363556 | -2467.640406 | -138.6 | -2467.181851 | -2467.273838 | 24.9 | -2456.673412 | -7.511179 | -2463.870818 |
| conformer_60 | -2467.587604 | 0.458786 | 0.368183 | -2467.643919 | -147.9 | -2467.185133 | -2467.272724 | 27.8 | -2456.658690 | -7.522818 | -2463.866627 |
| conformer_92 | -2467.587204 | 0.458193 | 0.362043 | -2467.642319 | -144.7 | -2467.184126 | -2467.277264 | 15.9 | -2456.671605 | -7.511386 | -2463.873050 |
| conformer_32 | -2467.587114 | 0.458611 | 0.362763 | -2467.638593 | -135.2 | -2467.179982 | -2467.272818 | 27.5 | -2456.675070 | -7.509085 | -2463.869859 |
| conformer_85 | -2467.586863 | 0.459079 | 0.365709 | -2467.639618 | -138.5 | -2467.180539 | -2467.270897 | 32.6 | -2456.666644 | -7.513807 | -2463.864487 |
| conformer_68 | -2467.586060 | 0.458282 | 0.363051 | -2467.638812 | -138.5 | -2467.180530 | -2467.272749 | 27.7 | -2456.671270 | -7.511763 | -2463.869721 |
| conformer_96 | -2467.584492 | 0.458628 | 0.362990 | -2467.638260 | -141.2 | -2467.179632 | -2467.272258 | 29.0 | -2456.672969 | -7.508044 | -2463.868780 |
| conformer_87 | -2467.583727 | 0.459059 | 0.365187 | -2467.640332 | -148.6 | -2467.181273 | -2467.272133 | 29.3 | -2456.666239 | -7.512750 | -2463.867395 |
| conformer_66 | -2467.583575 | 0.458681 | 0.363112 | -2467.635895 | -137.4 | -2467.177214 | -2467.269771 | 35.5 | -2456.673184 | -7.507705 | -2463.867086 |
| conformer_83 | -2467.582818 | 0.458967 | 0.366934 | -2467.638971 | -147.4 | -2467.180004 | -2467.269025 | 37.5 | -2456.655243 | -7.521340 | -2463.862790 |
| conformer_88 | -2467.581236 | 0.458646 | 0.361526 | -2467.635741 | -143.1 | -2467.177095 | -2467.271203 | 31.8 | -2456.671667 | -7.507644 | -2463.869278 |

[a]: Single-point calculation in aqueous phase with SMD model.

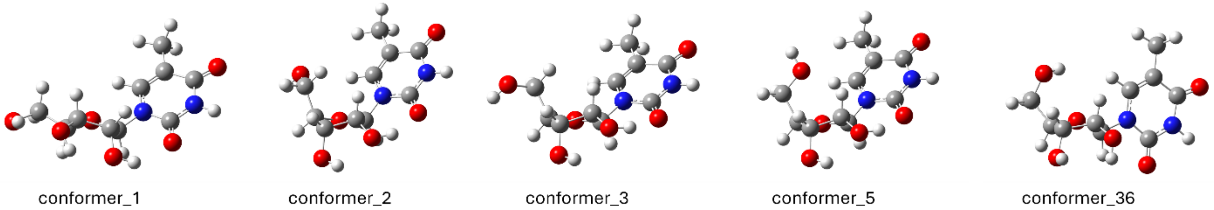

**Figure S42.** B3LYP-D3/def2-TZVPP optimized geometries of conformers for 5-methyluridine (**1rb5mU**).

**Table S40.** Conformers of gas-phase optimized 5-methyluridine (**1rb5mU**) at the B3LYP-D3/def2-TZVPP level of theory followed by aqueous phase single-point calculation. The columns display total energy without zero-point correction ($E_{Tot}$), thermal correction to enthalpy (δH), Gibbs free energy (δG), total energy without zero-point correction ($E_{Tot,W}$), Gibbs free energy ($G_{298,W}$) in water (W), total single-point energy ($E_{CBS}$) calculated at DLPNO-CCSD(T)/CBS level of theory, and their corresponding free energy $G_{CBS,W}$. $G_{298,W}$ and $G_{CBS,W}$ have been corrected to the standard state of 1 mol/L by addition of +7.908 kJ/mol. $\Delta G_{Solv}$ represents the Gibbs free energy of solvation. The data are arranged in the ascending numeric order of $E_{Tot,W}$. $\Delta G_{298,W}$ represents the respective energy difference to the lowest structure. Only conformers within the 24 kJ/mol (6 kcal/mol) energy window above the lowest in CREST are included in initial conformer sampling. Duplicates of the same structure are excluded. The overall optimum is marked bold.

| | B3LYP-D3/def2-TZVPP | | | SMD($H_2O$)/B3LYP-D3/def2-TZVPP[a] | | | | | DLPNO-CCSD(T)/CBS | | |
|---|---|---|---|---|---|---|---|---|---|---|---|
| 1rb5mU No. | $E_{Tot}$ (Hartree) | δH (Hartree) | δG (Hartree) | $E_{Tot,W}$ (Hartree) | $\Delta G_{Solv}$ (kJ/mol) | $H_{298,W}$ (Hartree) | $G_{298,W}$ (Hartree) | $\Delta G_{298,W}$ (kJ/mol) | $E_{CBS,HF}$ (Hartree) | $E_{CBS,C}$ (Hartree) | $G_{CBS,W}$ (Hartree) |
| conformer_18 | -950.793150 | 0.272517 | 0.210993 | -950.826731 | -88.2 | -950.554214 | -950.615738 | 4.9 | -945.365246 | -4.014791 | -949.199613 |
| conformer_6 | -950.792416 | 0.272403 | 0.210632 | -950.826572 | -89.7 | -950.554169 | -950.615940 | 4.4 | -945.364474 | -4.014643 | -949.199629 |
| conformer_11 | -950.790291 | 0.272357 | 0.210514 | -950.826560 | -95.2 | -950.554203 | -950.616046 | 4.1 | -945.360529 | -4.015951 | -949.199223 |
| **conformer_36** | **-950.786646** | **0.272385** | **0.207891** | **-950.825507** | **-102.0** | **-950.553122** | **-950.617616** | **0** | **-945.361490** | **-4.012247** | **-949.201695** |
| conformer_33 | -950.789342 | 0.272419 | 0.210172 | -950.825419 | -94.7 | -950.553000 | -950.615247 | 6.2 | -945.361485 | -4.014397 | -949.198775 |
| conformer_70 | -950.784326 | 0.272128 | 0.207981 | -950.825097 | -107.0 | -950.552969 | -950.617116 | 1.3 | -945.357585 | -4.013617 | -949.200980 |
| conformer_21 | -950.790541 | 0.272266 | 0.209355 | -950.824952 | -90.3 | -950.552686 | -950.615597 | 5.3 | -945.364367 | -4.012928 | -949.199339 |
| conformer_19 | -950.789543 | 0.272442 | 0.209082 | -950.824893 | -92.8 | -950.552451 | -950.615811 | 4.7 | -945.363066 | -4.013147 | -949.199470 |
| conformer_58 | -950.783303 | 0.272113 | 0.208185 | -950.824607 | -108.4 | -950.552494 | -950.616422 | 3.1 | -945.356286 | -4.013718 | -949.200111 |
| conformer_80 | -950.787197 | 0.272199 | 0.208671 | -950.824548 | -98.1 | -950.552349 | -950.615877 | 4.6 | -945.359479 | -4.014580 | -949.199727 |
| conformer_7 | -950.792900 | 0.272179 | 0.209762 | -950.824534 | -83.1 | -950.552355 | -950.614772 | 7.5 | -945.363006 | -4.016564 | -949.198430 |
| conformer_13 | -950.791459 | 0.272254 | 0.210015 | -950.824532 | -86.8 | -950.552278 | -950.614517 | 8.1 | -945.363602 | -4.014806 | -949.198453 |
| conformer_49 | -950.784175 | 0.272084 | 0.208802 | -950.824384 | -105.6 | -950.552300 | -950.615582 | 5.3 | -945.356381 | -4.013864 | -949.198640 |
| conformer_76 | -950.785242 | 0.272056 | 0.207153 | -950.824348 | -102.7 | -950.552292 | -950.617195 | 1.1 | -945.360086 | -4.012395 | -949.201421 |
| conformer_100 | -950.784437 | 0.271991 | 0.208524 | -950.824251 | -104.5 | -950.552260 | -950.615727 | 5.0 | -945.358609 | -4.012468 | -949.199355 |
| conformer_81 | -950.784085 | 0.272031 | 0.207849 | -950.824032 | -104.9 | -950.552001 | -950.616183 | 3.8 | -945.359462 | -4.011944 | -949.200491 |
| conformer_2 | -950.791712 | 0.272230 | 0.210292 | -950.823992 | -84.8 | -950.551762 | -950.613700 | 10.3 | -945.363126 | -4.015737 | -949.197839 |
| conformer_8 | -950.792444 | 0.272125 | 0.209491 | -950.823927 | -82.7 | -950.551802 | -950.614436 | 8.3 | -945.363432 | -4.015528 | -949.197940 |
| conformer_12 | -950.790987 | 0.272129 | 0.208964 | -950.823914 | -86.5 | -950.551785 | -950.614950 | 7.0 | -945.362322 | -4.015523 | -949.198796 |
| conformer_45 | -950.785679 | 0.272129 | 0.209000 | -950.823900 | -100.3 | -950.551771 | -950.614900 | 7.1 | -945.358837 | -4.013371 | -949.198417 |
| conformer_88 | -950.783785 | 0.272182 | 0.207413 | -950.823854 | -105.2 | -950.551672 | -950.616441 | 3.1 | -945.359384 | -4.011589 | -949.200617 |
| conformer_24 | -950.790810 | 0.272231 | 0.209110 | -950.823811 | -86.6 | -950.551580 | -950.614701 | 7.7 | -945.365579 | -4.012373 | -949.198831 |
| conformer_72 | -950.783211 | 0.272034 | 0.208100 | -950.823802 | -106.6 | -950.551768 | -950.615702 | 5.0 | -945.355261 | -4.014745 | -949.199484 |
| conformer_31 | -950.786918 | 0.272397 | 0.208845 | -950.823762 | -96.7 | -950.551365 | -950.614917 | 7.1 | -945.361941 | -4.012306 | -949.199234 |
| conformer_60 | -950.781512 | 0.272032 | 0.207805 | -950.823660 | -110.7 | -950.551628 | -950.615855 | 4.6 | -945.355828 | -4.012360 | -949.199519 |
| conformer_67 | -950.785232 | 0.272060 | 0.208248 | -950.823615 | -100.8 | -950.551555 | -950.615367 | 5.9 | -945.356456 | -4.015038 | -949.198617 |
| conformer_91 | -950.786224 | 0.271995 | 0.208376 | -950.823593 | -98.1 | -950.551598 | -950.615217 | 6.3 | -945.361412 | -4.011876 | -949.199269 |
| conformer_35 | -950.785310 | 0.272142 | 0.208352 | -950.823586 | -100.5 | -950.551444 | -950.615234 | 6.3 | -945.356221 | -4.015418 | -949.198551 |
| conformer_104 | -950.780465 | 0.271928 | 0.207231 | -950.823505 | -113.0 | -950.551577 | -950.616274 | 3.5 | -945.354744 | -4.012349 | -949.199889 |
| conformer_29 | -950.790763 | 0.272170 | 0.210297 | -950.823488 | -85.9 | -950.551318 | -950.613191 | 11.6 | -945.362676 | -4.015672 | -949.197764 |

| | | | | | | | | | | | |
|---|---|---|---|---|---|---|---|---|---|---|---|
| conformer_101 | -950.782986 | 0.272095 | 0.208000 | -950.823153 | -105.5 | -950.551058 | -950.615153 | 6.5 | -945.358085 | -4.012148 | -949.199389 |
| conformer_5 | -950.793760 | 0.272204 | 0.209830 | -950.823144 | -77.1 | -950.550940 | -950.613314 | 11.3 | -945.365625 | -4.015228 | -949.197394 |
| conformer_44 | -950.786554 | 0.272255 | 0.209930 | -950.822940 | -95.5 | -950.550685 | -950.613010 | 12.1 | -945.360087 | -4.013770 | -949.197301 |
| conformer_27 | -950.788017 | 0.272248 | 0.208694 | -950.822870 | -91.5 | -950.550622 | -950.614176 | 9.0 | -945.362843 | -4.012822 | -949.198811 |
| conformer_39 | -950.785599 | 0.272294 | 0.209981 | -950.822851 | -97.8 | -950.550557 | -950.612870 | 12.5 | -945.357055 | -4.015422 | -949.196737 |
| conformer_85 | -950.783177 | 0.271953 | 0.207057 | -950.822772 | -104.0 | -950.550819 | -950.615715 | 5.0 | -945.359837 | -4.010686 | -949.200050 |
| conformer_3 | -950.790295 | 0.272228 | 0.209297 | -950.822647 | -84.9 | -950.550419 | -950.613350 | 11.2 | -945.363271 | -4.014005 | -949.197319 |
| conformer_15 | -950.789471 | 0.272131 | 0.209201 | -950.822613 | -87.0 | -950.550482 | -950.613412 | 11.0 | -945.361981 | -4.014387 | -949.197297 |
| conformer_115 | -950.781321 | 0.271948 | 0.207468 | -950.822458 | -108.0 | -950.550510 | -950.614990 | 6.9 | -945.356265 | -4.012024 | -949.198945 |
| conformer_83 | -950.784358 | 0.271990 | 0.208105 | -950.822248 | -99.5 | -950.550258 | -950.614143 | 9.1 | -945.359331 | -4.012583 | -949.198686 |
| conformer_34 | -950.785937 | 0.272488 | 0.209414 | -950.822134 | -95.0 | -950.549646 | -950.612720 | 12.9 | -945.360262 | -4.012663 | -949.196696 |
| conformer_105 | -950.783553 | 0.271971 | 0.207201 | -950.821725 | -100.2 | -950.549754 | -950.614524 | 8.1 | -945.360869 | -4.010220 | -949.199049 |
| conformer_23 | -950.789614 | 0.272093 | 0.208660 | -950.821616 | -84.0 | -950.549523 | -950.612956 | 12.2 | -945.363131 | -4.013844 | -949.197304 |
| conformer_61 | -950.782713 | 0.272067 | 0.208514 | -950.821510 | -101.9 | -950.549443 | -950.612996 | 12.1 | -945.354207 | -4.015034 | -949.196512 |
| conformer_56 | -950.784263 | 0.272075 | 0.208609 | -950.821393 | -97.5 | -950.549318 | -950.612784 | 12.7 | -945.359341 | -4.012298 | -949.197148 |
| conformer_55 | -950.782181 | 0.272136 | 0.208907 | -950.821233 | -102.5 | -950.549097 | -950.612326 | 13.9 | -945.354959 | -4.014048 | -949.196141 |
| conformer_9 | -950.790128 | 0.272241 | 0.208908 | -950.821165 | -81.5 | -950.548924 | -950.612257 | 14.1 | -945.364670 | -4.013228 | -949.197015 |
| conformer_89 | -950.785168 | 0.272145 | 0.208181 | -950.821008 | -94.1 | -950.548863 | -950.612827 | 12.6 | -945.359685 | -4.012779 | -949.197111 |
| conformer_10 | -950.789780 | 0.272244 | 0.208902 | -950.820809 | -81.5 | -950.548565 | -950.611907 | 15.0 | -945.365498 | -4.012179 | -949.196792 |
| conformer_40 | -950.785660 | 0.272284 | 0.209220 | -950.820607 | -91.8 | -950.548323 | -950.611387 | 16.4 | -945.359731 | -4.013190 | -949.195636 |
| conformer_74 | -950.782495 | 0.272349 | 0.208813 | -950.820583 | -100.0 | -950.548234 | -950.611770 | 15.3 | -945.356001 | -4.013244 | -949.195508 |
| conformer_30 | -950.787740 | 0.272267 | 0.209092 | -950.820425 | -85.8 | -950.548158 | -950.611333 | 16.5 | -945.362855 | -4.012501 | -949.195937 |
| conformer_46 | -950.785193 | 0.272094 | 0.208560 | -950.820418 | -92.5 | -950.548324 | -950.611858 | 15.1 | -945.361032 | -4.011708 | -949.196393 |
| conformer_57 | -950.782121 | 0.272162 | 0.209067 | -950.820359 | -100.4 | -950.548197 | -950.611292 | 16.6 | -945.355042 | -4.013940 | -949.195141 |
| conformer_20 | -950.787572 | 0.272091 | 0.208614 | -950.819944 | -85.0 | -950.547853 | -950.611330 | 16.5 | -945.363606 | -4.012152 | -949.196504 |
| conformer_103 | -950.780656 | 0.272216 | 0.208601 | -950.819926 | -103.1 | -950.547710 | -950.611325 | 16.5 | -945.351498 | -4.016026 | -949.195181 |
| conformer_94 | -950.785525 | 0.272091 | 0.207959 | -950.819876 | -90.2 | -950.547785 | -950.611917 | 15.0 | -945.360936 | -4.012334 | -949.196650 |
| conformer_71 | -950.782704 | 0.272050 | 0.208262 | -950.819875 | -97.6 | -950.547825 | -950.611613 | 15.8 | -945.356382 | -4.013470 | -949.195749 |
| conformer_42 | -950.787456 | 0.272024 | 0.208451 | -950.819423 | -83.9 | -950.547399 | -950.610972 | 17.4 | -945.363956 | -4.011852 | -949.196312 |
| conformer_107 | -950.782077 | 0.272305 | 0.210158 | -950.819416 | -98.0 | -950.547111 | -950.609258 | 21.9 | -945.349646 | -4.019053 | -949.192869 |
| conformer_25 | -950.788503 | 0.272271 | 0.208980 | -950.819385 | -81.1 | -950.547114 | -950.610405 | 18.9 | -945.364329 | -4.012168 | -949.195387 |
| conformer_51 | -950.782350 | 0.272323 | 0.210406 | -950.818957 | -96.1 | -950.546634 | -950.608551 | 23.8 | -945.350556 | -4.018041 | -949.191787 |
| conformer_1 | -950.789517 | 0.272419 | 0.209666 | -950.818745 | -76.7 | -950.546326 | -950.609079 | 22.4 | -945.364588 | -4.012536 | -949.193673 |
| conformer_79 | -950.780744 | 0.272284 | 0.209069 | -950.818029 | -97.9 | -950.545745 | -950.608960 | 22.7 | -945.350158 | -4.017749 | -949.193111 |
| conformer_102 | -950.783022 | 0.272136 | 0.208906 | -950.817726 | -91.1 | -950.545590 | -950.608820 | 23.1 | -945.355630 | -4.015200 | -949.193616 |
| conformer_90 | -950.781234 | 0.272150 | 0.209861 | -950.816444 | -92.4 | -950.544294 | -950.606583 | 29.0 | -945.350338 | -4.018099 | -949.190774 |
| conformer_114 | -950.777453 | 0.272239 | 0.208645 | -950.815687 | -100.4 | -950.543448 | -950.607042 | 27.8 | -945.346927 | -4.017728 | -949.191232 |

[a]: Single-point calculation in aqueous phase with SMD model.

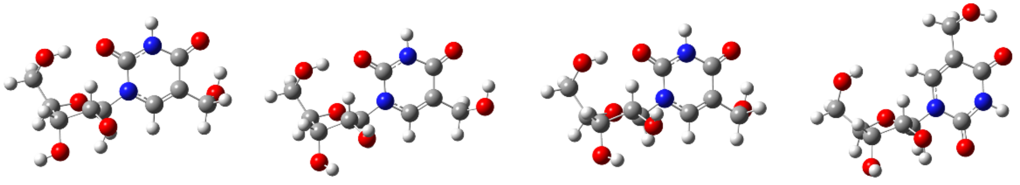

conformer_11 conformer_84 conformer_100 conformer_114

**Figure S43.** B3LYP-D3/def2-TZVPP optimized geometries of conformers for 5-hydroxymethyluridine (**1rb5hmU**).

**Table S41.** Conformers of gas-phase optimized 5-hydroxymethyluridine (**1rb5hmU**) at the B3LYP-D3/def2-TZVPP level of theory followed by aqueous phase single-point calculation. The columns display total energy without zero-point correction ($E_{Tot}$), thermal correction to enthalpy (δH), Gibbs free energy (δG), total energy without zero-point correction ($E_{Tot,W}$), Gibbs free energy ($G_{298,W}$) in water (W), total single-point energy ($E_{CBS}$) calculated at DLPNO-CCSD(T)/CBS level of theory, and their corresponding free energy $G_{CBS,W}$. $G_{298,W}$ and $G_{CBS,W}$ have been corrected to the standard state of 1 mol/L by addition of +7.908 kJ/mol. $\Delta G_{Solv}$ represents the Gibbs free energy of solvation. The data are arranged in the ascending numeric order of $E_{Tot,W}$. $\Delta G_{298,W}$ represents the respective energy difference to the lowest structure. Only conformers within the 24 kJ/mol (6 kcal/mol) energy window above the lowest in CREST are included in initial conformer sampling. Duplicates of the same structure are excluded. The overall optimum is marked bold.

| | B3LYP-D3/def2-TZVPP | | | SMD($H_2O$)/B3LYP-D3/def2-TZVPP[a] | | | | | DLPNO-CCSD(T)/CBS | | |
|---|---|---|---|---|---|---|---|---|---|---|---|
| 1rb5hmU No. | $E_{Tot}$ (Hartree) | δH (Hartree) | δG (Hartree) | $E_{Tot,W}$ (Hartree) | $\Delta G_{Solv}$ (kJ/mol) | $H_{298,W}$ (Hartree) | $G_{298,W}$ (Hartree) | $\Delta G_{298,W}$ (kJ/mol) | $E_{CBS,HF}$ (Hartree) | $E_{CBS,C}$ (Hartree) | $G_{CBS,W}$ (Hartree) |
| conformer_100 | -1026.042518 | 0.278984 | 0.215252 | -1026.081781 | -103.1 | -1025.802797 | -1025.863517 | 7.1 | -1020.256155 | -4.285523 | -1024.362676 |
| conformer_84 | -1026.042673 | 0.278974 | 0.215279 | -1026.081630 | -102.3 | -1025.802656 | -1025.863339 | 7.6 | -1020.256358 | -4.285478 | -1024.362501 |
| conformer_11 | -1026.042060 | 0.278914 | 0.215084 | -1026.081609 | -103.8 | -1025.802695 | -1025.863513 | 7.1 | -1020.255642 | -4.285373 | -1024.362469 |
| conformer_19 | -1026.039859 | 0.278869 | 0.214945 | -1026.081566 | -109.5 | -1025.802697 | -1025.863609 | 6.9 | -1020.251631 | -4.286675 | -1024.362056 |
| conformer_25 | -1026.039633 | 0.278859 | 0.214937 | -1026.081484 | -109.9 | -1025.802625 | -1025.863535 | 7.0 | -1020.251389 | -4.286634 | -1024.361925 |
| conformer_13 | -1026.041880 | 0.278870 | 0.214953 | -1026.081449 | -103.9 | -1025.802579 | -1025.863484 | 7.2 | -1020.255485 | -4.285347 | -1024.362437 |
| conformer_59 | -1026.039101 | 0.278771 | 0.213787 | -1026.080732 | -109.3 | -1025.801961 | -1025.863933 | 6.0 | -1020.252382 | -4.285252 | -1024.362467 |
| conformer_115 | -1026.036486 | 0.278866 | 0.212215 | -1026.080594 | -115.8 | -1025.801728 | -1025.865367 | 2.2 | -1020.252880 | -4.282948 | -1024.364709 |
| conformer_175 | -1026.036480 | 0.278657 | 0.214063 | -1026.080556 | -115.7 | -1025.801899 | -1025.863481 | 7.2 | -1020.250199 | -4.285373 | -1024.362573 |
| conformer_234 | -1026.034236 | 0.278671 | 0.214097 | -1026.080527 | -121.5 | -1025.801856 | -1025.863418 | 7.4 | -1020.246097 | -4.286693 | -1024.361972 |
| **conformer_114** | **-1026.036365** | **0.278841** | **0.211250** | **-1026.080481** | **-115.8** | **-1025.801640** | **-1025.866219** | **0** | **-1020.252731** | **-4.282912** | **-1024.365497** |
| conformer_128 | -1026.038758 | 0.278897 | 0.214451 | -1026.080479 | -109.5 | -1025.801582 | -1025.863016 | 8.4 | -1020.252408 | -4.285097 | -1024.361761 |
| conformer_112 | -1026.038874 | 0.278909 | 0.214359 | -1026.080472 | -109.2 | -1025.801563 | -1025.863101 | 8.2 | -1020.252587 | -4.285100 | -1024.361915 |
| conformer_43 | -1026.041195 | 0.278884 | 0.214192 | -1026.080441 | -103.0 | -1025.801557 | -1025.863237 | 7.8 | -1020.256117 | -4.284004 | -1024.362163 |
| conformer_56 | -1026.039319 | 0.278859 | 0.213055 | -1026.080233 | -107.4 | -1025.801374 | -1025.864166 | 5.4 | -1020.254258 | -4.283983 | -1024.363088 |
| conformer_203 | -1026.039928 | 0.278675 | 0.214273 | -1026.080150 | -105.6 | -1025.801475 | -1025.862865 | 8.8 | -1020.253827 | -4.285126 | -1024.361889 |
| conformer_159 | -1026.040110 | 0.278777 | 0.214555 | -1026.080149 | -105.1 | -1025.801372 | -1025.862582 | 9.5 | -1020.253971 | -4.285169 | -1024.361613 |
| conformer_54 | -1026.040549 | 0.278775 | 0.213761 | -1026.080079 | -103.8 | -1025.801304 | -1025.863306 | 7.6 | -1020.255756 | -4.283837 | -1024.362350 |
| conformer_44 | -1026.037577 | 0.278624 | 0.214135 | -1026.080053 | -111.5 | -1025.801429 | -1025.862906 | 8.7 | -1020.249576 | -4.286283 | -1024.361188 |
| conformer_89 | -1026.039559 | 0.278647 | 0.213450 | -1026.080039 | -106.3 | -1025.801392 | -1025.863577 | 6.9 | -1020.253823 | -4.284713 | -1024.362554 |
| conformer_53 | -1026.037306 | 0.278702 | 0.214349 | -1026.080021 | -112.1 | -1025.801319 | -1025.862660 | 9.3 | -1020.249249 | -4.286275 | -1024.360877 |
| conformer_41 | -1026.039398 | 0.278665 | 0.214217 | -1026.080005 | -106.6 | -1025.801340 | -1025.862776 | 9.0 | -1020.253215 | -4.284958 | -1024.361551 |
| conformer_36 | -1026.039544 | 0.278585 | 0.213938 | -1026.079996 | -106.2 | -1025.801411 | -1025.863046 | 8.3 | -1020.253422 | -4.284967 | -1024.361891 |
| conformer_49 | -1026.039663 | 0.278867 | 0.212859 | -1026.079986 | -105.9 | -1025.801119 | -1025.864115 | 5.5 | -1020.254688 | -4.283879 | -1024.363020 |
| conformer_188 | -1026.034292 | 0.278657 | 0.212196 | -1026.079950 | -119.9 | -1025.801293 | -1025.864742 | 3.9 | -1020.248973 | -4.284360 | -1024.363783 |
| conformer_201 | -1026.033868 | 0.278588 | 0.212148 | -1026.079820 | -120.6 | -1025.801232 | -1025.864660 | 4.1 | -1020.248672 | -4.284351 | -1024.363815 |
| conformer_181 | -1026.032990 | 0.278556 | 0.212342 | -1026.079686 | -122.6 | -1025.801130 | -1025.864332 | 5.0 | -1020.247411 | -4.284488 | -1024.363241 |
| conformer_111 | -1026.034537 | 0.278707 | 0.213477 | -1026.079599 | -118.3 | -1025.800892 | -1025.863110 | 8.2 | -1020.248037 | -4.284796 | -1024.361406 |
| conformer_27 | -1026.040980 | 0.278788 | 0.214386 | -1026.079568 | -101.3 | -1025.800780 | -1025.862170 | 10.6 | -1020.254650 | -4.285532 | -1024.361372 |
| conformer_157 | -1026.040885 | 0.278734 | 0.214170 | -1026.079547 | -101.5 | -1025.800813 | -1025.862365 | 10.1 | -1020.254586 | -4.285482 | -1024.361548 |

| | | | | | | | | | | | |
|---|---|---|---|---|---|---|---|---|---|---|---|
| conformer_187 | -1026.032982 | 0.278491 | 0.211823 | -1026.079541 | -122.2 | -1025.801050 | -1025.864706 | 4.0 | -1020.247832 | -4.284139 | -1024.363695 |
| conformer_82 | -1026.038566 | 0.278708 | 0.214138 | -1026.079514 | -107.5 | -1025.800806 | -1025.862364 | 10.1 | -1020.251240 | -4.285565 | -1024.360603 |
| conformer_126 | -1026.037412 | 0.278741 | 0.213182 | -1026.079452 | -110.4 | -1025.800711 | -1025.863258 | 7.8 | -1020.251056 | -4.285425 | -1024.362326 |
| conformer_207 | -1026.037414 | 0.278752 | 0.213072 | -1026.079423 | -110.3 | -1025.800671 | -1025.863339 | 7.6 | -1020.251014 | -4.285380 | -1024.362319 |
| conformer_17 | -1026.042333 | 0.278596 | 0.213779 | -1026.079415 | -97.4 | -1025.800819 | -1025.862624 | 9.4 | -1020.253900 | -4.287467 | -1024.361659 |
| conformer_81 | -1026.037965 | 0.278581 | 0.213964 | -1026.079357 | -108.7 | -1025.800776 | -1025.862381 | 10.1 | -1020.250878 | -4.285734 | -1024.361028 |
| conformer_51 | -1026.039602 | 0.278739 | 0.214105 | -1026.079349 | -104.4 | -1025.800610 | -1025.862232 | 10.5 | -1020.252508 | -4.285280 | -1024.360418 |
| conformer_219 | -1026.040483 | 0.278616 | 0.214021 | -1026.079338 | -102.0 | -1025.800722 | -1025.862305 | 10.3 | -1020.255299 | -4.284661 | -1024.361782 |
| conformer_3 | -1026.041581 | 0.278740 | 0.214750 | -1026.079316 | -99.1 | -1025.800576 | -1025.861554 | 12.2 | -1020.254478 | -4.286515 | -1024.360966 |
| conformer_153 | -1026.035961 | 0.278648 | 0.213567 | -1026.079306 | -113.8 | -1025.800658 | -1025.862727 | 9.2 | -1020.248272 | -4.286118 | -1024.361157 |
| conformer_192 | -1026.035345 | 0.278549 | 0.211352 | -1026.079290 | -115.4 | -1025.800741 | -1025.864926 | 3.4 | -1020.251668 | -4.283130 | -1024.364379 |
| conformer_52 | -1026.041369 | 0.278805 | 0.213793 | -1026.079245 | -99.4 | -1025.800440 | -1025.862440 | 9.9 | -1020.257278 | -4.283434 | -1024.361783 |
| conformer_58 | -1026.041193 | 0.278762 | 0.213665 | -1026.079192 | -99.8 | -1025.800430 | -1025.862515 | 9.7 | -1020.257228 | -4.283379 | -1024.361930 |
| conformer_113 | -1026.034403 | 0.278678 | 0.213391 | -1026.079182 | -117.6 | -1025.800504 | -1025.862779 | 9.0 | -1020.247929 | -4.284809 | -1024.361114 |
| conformer_200 | -1026.034049 | 0.278504 | 0.212112 | -1026.079179 | -118.5 | -1025.800675 | -1025.864055 | 5.7 | -1020.250748 | -4.282765 | -1024.363519 |
| conformer_4 | -1026.041542 | 0.278797 | 0.214977 | -1026.079168 | -98.8 | -1025.800371 | -1025.861179 | 13.2 | -1020.254446 | -4.286470 | -1024.360553 |
| conformer_189 | -1026.035236 | 0.278555 | 0.213009 | -1026.079168 | -115.3 | -1025.800613 | -1025.863147 | 8.1 | -1020.247783 | -4.286025 | -1024.361718 |
| conformer_96 | -1026.036095 | 0.278768 | 0.213808 | -1026.079149 | -113.0 | -1025.800381 | -1025.862329 | 10.2 | -1020.250419 | -4.284364 | -1024.361018 |
| conformer_216 | -1026.033237 | 0.278582 | 0.212911 | -1026.079134 | -120.5 | -1025.800552 | -1025.863211 | 7.9 | -1020.246400 | -4.285711 | -1024.362085 |
| conformer_29 | -1026.040452 | 0.278722 | 0.213985 | -1026.079098 | -101.5 | -1025.800376 | -1025.862101 | 10.8 | -1020.253372 | -4.286007 | -1024.361028 |
| conformer_30 | -1026.040469 | 0.278661 | 0.213446 | -1026.079068 | -101.3 | -1025.800407 | -1025.862610 | 9.5 | -1020.253572 | -4.285883 | -1024.361596 |
| conformer_241 | -1026.034823 | 0.278576 | 0.212956 | -1026.079062 | -116.1 | -1025.800486 | -1025.863094 | 8.2 | -1020.250269 | -4.283360 | -1024.361900 |
| conformer_259 | -1026.033452 | 0.278752 | 0.211965 | -1026.079047 | -119.7 | -1025.800295 | -1025.864070 | 5.6 | -1020.250369 | -4.282441 | -1024.363427 |
| conformer_238 | -1026.037845 | 0.278622 | 0.213389 | -1026.079046 | -108.2 | -1025.800424 | -1025.862645 | 9.4 | -1020.252712 | -4.284506 | -1024.362018 |
| conformer_243 | -1026.033057 | 0.278563 | 0.212975 | -1026.078997 | -120.6 | -1025.800434 | -1025.863010 | 8.4 | -1020.246350 | -4.285611 | -1024.361914 |
| conformer_211 | -1026.034027 | 0.278510 | 0.212143 | -1026.078981 | -118.0 | -1025.800471 | -1025.863826 | 6.3 | -1020.250949 | -4.282579 | -1024.363327 |
| conformer_221 | -1026.033739 | 0.278667 | 0.211514 | -1026.078967 | -118.7 | -1025.800300 | -1025.864441 | 4.7 | -1020.250735 | -4.282431 | -1024.363867 |
| conformer_93 | -1026.036296 | 0.278761 | 0.213813 | -1026.078964 | -112.0 | -1025.800203 | -1025.862139 | 10.7 | -1020.250649 | -4.284314 | -1024.360806 |
| conformer_62 | -1026.041465 | 0.278626 | 0.214491 | -1026.078958 | -98.4 | -1025.800332 | -1025.861455 | 12.5 | -1020.254603 | -4.286740 | -1024.361333 |
| conformer_12 | -1026.042336 | 0.278666 | 0.214021 | -1026.078953 | -96.1 | -1025.800287 | -1025.861920 | 11.3 | -1020.254862 | -4.286369 | -1024.360815 |
| conformer_78 | -1026.038353 | 0.278678 | 0.214116 | -1026.078931 | -106.5 | -1025.800253 | -1025.861803 | 11.6 | -1020.251090 | -4.285619 | -1024.360160 |
| conformer_205 | -1026.036424 | 0.278685 | 0.213788 | -1026.078907 | -111.5 | -1025.800222 | -1025.862107 | 10.8 | -1020.250295 | -4.284716 | -1024.360694 |
| conformer_173 | -1026.036864 | 0.278583 | 0.213063 | -1026.078865 | -110.3 | -1025.800282 | -1025.862790 | 9.0 | -1020.253173 | -4.282868 | -1024.361967 |
| conformer_97 | -1026.036647 | 0.278880 | 0.213238 | -1026.078861 | -110.8 | -1025.799981 | -1025.862611 | 9.5 | -1020.253088 | -4.283077 | -1024.362129 |
| conformer_15 | -1026.042308 | 0.278611 | 0.213858 | -1026.078819 | -95.9 | -1025.800208 | -1025.861949 | 11.2 | -1020.254778 | -4.286365 | -1024.360784 |
| conformer_101 | -1026.036486 | 0.278840 | 0.213028 | -1026.078780 | -111.0 | -1025.799940 | -1025.862740 | 9.1 | -1020.253057 | -4.283009 | -1024.362319 |
| conformer_165 | -1026.037024 | 0.278686 | 0.213402 | -1026.078749 | -109.5 | -1025.800063 | -1025.862335 | 10.2 | -1020.253335 | -4.282825 | -1024.361470 |
| conformer_1 | -1026.044570 | 0.278691 | 0.214482 | -1026.078686 | -89.6 | -1025.799995 | -1025.861192 | 13.2 | -1020.257324 | -4.286671 | -1024.360618 |
| conformer_167 | -1026.031341 | 0.278496 | 0.212132 | -1026.078641 | -124.2 | -1025.800145 | -1025.863497 | 7.1 | -1020.246994 | -4.283221 | -1024.362371 |
| conformer_171 | -1026.031367 | 0.278497 | 0.211996 | -1026.078596 | -124.0 | -1025.800099 | -1025.863588 | 6.9 | -1020.246995 | -4.283274 | -1024.362490 |
| conformer_10 | -1026.040657 | 0.278745 | 0.215692 | -1026.078579 | -99.6 | -1025.799834 | -1025.859875 | 16.7 | -1020.251356 | -4.288173 | -1024.358747 |
| conformer_279 | -1026.030363 | 0.278388 | 0.211458 | -1026.078533 | -126.5 | -1025.800145 | -1025.864063 | 5.7 | -1020.245989 | -4.283238 | -1024.362928 |
| conformer_120 | -1026.035320 | 0.278782 | 0.213346 | -1026.078528 | -113.4 | -1025.799746 | -1025.862170 | 10.6 | -1020.247564 | -4.286161 | -1024.360576 |
| conformer_88 | -1026.040456 | 0.278653 | 0.214702 | -1026.078522 | -99.9 | -1025.799869 | -1025.860808 | 14.2 | -1020.253858 | -4.286424 | -1024.360634 |
| conformer_286 | -1026.031615 | 0.278386 | 0.212430 | -1026.078493 | -123.1 | -1025.800107 | -1025.863051 | 8.3 | -1020.244847 | -4.284883 | -1024.361166 |
| conformer_125 | -1026.034927 | 0.278696 | 0.213066 | -1026.078484 | -114.4 | -1025.799788 | -1025.862406 | 10.0 | -1020.247178 | -4.286217 | -1024.360874 |
| conformer_298 | -1026.030202 | 0.278358 | 0.211327 | -1026.078416 | -126.6 | -1025.800058 | -1025.864077 | 5.6 | -1020.245877 | -4.283220 | -1024.362973 |
| conformer_2 | -1026.043798 | 0.278721 | 0.214233 | -1026.078322 | -90.6 | -1025.799601 | -1025.861077 | 13.5 | -1020.257089 | -4.286099 | -1024.360466 |
| conformer_196 | -1026.035004 | 0.278911 | 0.213989 | -1026.078311 | -113.7 | -1025.799400 | -1025.861310 | 12.9 | -1020.245546 | -4.287338 | -1024.359191 |
| conformer_285 | -1026.032882 | 0.278619 | 0.212595 | -1026.078306 | -119.3 | -1025.799687 | -1025.862699 | 9.2 | -1020.249446 | -4.282930 | -1024.362194 |
| conformer_99 | -1026.040091 | 0.278629 | 0.215381 | -1026.078229 | -100.1 | -1025.799600 | -1025.859836 | 16.8 | -1020.249717 | -4.289075 | -1024.358537 |
| conformer_185 | -1026.036051 | 0.278862 | 0.214641 | -1026.078203 | -110.7 | -1025.799341 | -1025.860550 | 14.9 | -1020.246331 | -4.288039 | -1024.358869 |
| conformer_184 | -1026.035679 | 0.278530 | 0.212698 | -1026.078120 | -111.4 | -1025.799590 | -1025.862410 | 10.0 | -1020.249235 | -4.285371 | -1024.361338 |
| conformer_139 | -1026.036142 | 0.278720 | 0.214235 | -1026.078111 | -110.2 | -1025.799391 | -1025.860864 | 14.1 | -1020.251148 | -4.284533 | -1024.360402 |
| conformer_317 | -1026.032509 | 0.278593 | 0.212420 | -1026.078078 | -119.6 | -1025.799485 | -1025.862646 | 9.4 | -1020.249148 | -4.282893 | -1024.362177 |
| conformer_174 | -1026.035801 | 0.278563 | 0.212979 | -1026.078071 | -111.0 | -1025.799508 | -1025.862080 | 10.9 | -1020.249265 | -4.285447 | -1024.360991 |

| | | | | | | | | | | | |
|---|---|---|---|---|---|---|---|---|---|---|---|
| conformer_7 | -1026.040057 | 0.278657 | 0.213251 | -1026.078034 | -99.7 | -1025.799377 | -1025.861771 | 11.7 | -1020.254500 | -4.284897 | -1024.361111 |
| conformer_130 | -1026.036240 | 0.278737 | 0.214190 | -1026.078013 | -109.7 | -1025.799276 | -1025.860811 | 14.2 | -1020.251301 | -4.284490 | -1024.360362 |
| conformer_68 | -1026.038416 | 0.278569 | 0.213696 | -1026.077998 | -103.9 | -1025.799429 | -1025.861290 | 12.9 | -1020.252297 | -4.285075 | -1024.360246 |
| conformer_136 | -1026.035001 | 0.278734 | 0.214247 | -1026.077996 | -112.9 | -1025.799262 | -1025.860737 | 14.4 | -1020.247974 | -4.286131 | -1024.359842 |
| conformer_206 | -1026.034128 | 0.278518 | 0.213704 | -1026.077922 | -115.0 | -1025.799404 | -1025.861206 | 13.2 | -1020.244491 | -4.287337 | -1024.358905 |
| conformer_85 | -1026.037724 | 0.278782 | 0.213384 | -1026.077902 | -105.5 | -1025.799120 | -1025.861506 | 12.4 | -1020.253941 | -4.283627 | -1024.361350 |
| conformer_138 | -1026.035160 | 0.278763 | 0.214351 | -1026.077900 | -112.2 | -1025.799137 | -1025.860537 | 14.9 | -1020.248208 | -4.286073 | -1024.359658 |
| conformer_163 | -1026.035160 | 0.278763 | 0.214350 | -1026.077900 | -112.2 | -1025.799137 | -1025.860538 | 14.9 | -1020.248208 | -4.286073 | -1024.359659 |
| conformer_61 | -1026.037091 | 0.278418 | 0.212424 | -1026.077894 | -107.1 | -1025.799476 | -1025.862458 | 9.9 | -1020.250206 | -4.286022 | -1024.361595 |
| conformer_33 | -1026.039154 | 0.278511 | 0.212789 | -1026.077877 | -101.7 | -1025.799366 | -1025.862076 | 10.9 | -1020.253264 | -4.285119 | -1024.361304 |
| conformer_118 | -1026.034464 | 0.278604 | 0.213236 | -1026.077852 | -113.9 | -1025.799248 | -1025.861604 | 12.1 | -1020.247270 | -4.285207 | -1024.359616 |
| conformer_9 | -1026.039878 | 0.278692 | 0.213687 | -1026.077851 | -99.7 | -1025.799159 | -1025.861152 | 13.3 | -1020.254266 | -4.284933 | -1024.360472 |
| conformer_37 | -1026.038884 | 0.278560 | 0.213355 | -1026.077812 | -102.2 | -1025.799252 | -1025.861445 | 12.5 | -1020.252833 | -4.285272 | -1024.360666 |
| conformer_284 | -1026.034675 | 0.278591 | 0.213650 | -1026.077802 | -113.2 | -1025.799211 | -1025.861140 | 13.3 | -1020.245045 | -4.287349 | -1024.358859 |
| conformer_208 | -1026.035920 | 0.278621 | 0.213089 | -1026.077782 | -109.9 | -1025.799161 | -1025.861681 | 11.9 | -1020.251113 | -4.284261 | -1024.361134 |
| conformer_86 | -1026.037588 | 0.278697 | 0.212769 | -1026.077731 | -105.4 | -1025.799034 | -1025.861950 | 11.2 | -1020.253928 | -4.283586 | -1024.361877 |
| conformer_283 | -1026.036041 | 0.278401 | 0.211958 | -1026.077728 | -109.5 | -1025.799327 | -1025.862758 | 9.1 | -1020.251208 | -4.283972 | -1024.361898 |
| conformer_227 | -1026.035020 | 0.278453 | 0.212620 | -1026.077674 | -112.0 | -1025.799221 | -1025.862042 | 11.0 | -1020.249991 | -4.283574 | -1024.360587 |
| conformer_71 | -1026.038236 | 0.278337 | 0.212450 | -1026.077629 | -103.4 | -1025.799292 | -1025.862167 | 10.6 | -1020.251658 | -4.285389 | -1024.360978 |
| conformer_106 | -1026.036364 | 0.278767 | 0.213843 | -1026.077619 | -108.3 | -1025.798852 | -1025.860764 | 14.3 | -1020.247979 | -4.286628 | -1024.359007 |
| conformer_79 | -1026.038064 | 0.278638 | 0.214374 | -1026.077612 | -103.8 | -1025.798974 | -1025.860226 | 15.7 | -1020.247545 | -4.288838 | -1024.358545 |
| conformer_75 | -1026.040565 | 0.278645 | 0.214905 | -1026.077593 | -97.2 | -1025.798948 | -1025.859676 | 17.2 | -1020.250614 | -4.289098 | -1024.358822 |
| conformer_28 | -1026.037991 | 0.278935 | 0.215414 | -1026.077579 | -103.9 | -1025.798644 | -1025.859153 | 18.6 | -1020.247319 | -4.289464 | -1024.357945 |
| conformer_300 | -1026.030728 | 0.278284 | 0.211660 | -1026.077560 | -123.0 | -1025.799276 | -1025.862888 | 8.7 | -1020.243687 | -4.285732 | -1024.361579 |
| conformer_182 | -1026.033832 | 0.278683 | 0.214452 | -1026.077539 | -114.8 | -1025.798856 | -1025.860075 | 16.1 | -1020.243306 | -4.288410 | -1024.357958 |
| conformer_45 | -1026.039227 | 0.278411 | 0.213357 | -1026.077490 | -100.5 | -1025.799079 | -1025.861121 | 13.4 | -1020.251497 | -4.286516 | -1024.359907 |
| conformer_6 | -1026.044312 | 0.278995 | 0.216416 | -1026.077489 | -87.1 | -1025.798494 | -1025.858061 | 21.4 | -1020.250878 | -4.291094 | -1024.355721 |
| conformer_305 | -1026.031323 | 0.278434 | 0.211894 | -1026.077479 | -121.2 | -1025.799045 | -1025.862573 | 9.6 | -1020.247572 | -4.282895 | -1024.361717 |
| conformer_233 | -1026.034212 | 0.278519 | 0.212703 | -1026.077299 | -113.1 | -1025.798780 | -1025.861584 | 12.2 | -1020.250613 | -4.283412 | -1024.361397 |
| conformer_293 | -1026.028998 | 0.278258 | 0.211244 | -1026.077291 | -126.8 | -1025.799033 | -1025.863035 | 8.4 | -1020.244711 | -4.283082 | -1024.361829 |
| conformer_193 | -1026.034049 | 0.278725 | 0.212602 | -1026.077271 | -113.5 | -1025.798546 | -1025.861657 | 12.0 | -1020.250778 | -4.282636 | -1024.361021 |
| conformer_50 | -1026.039713 | 0.278599 | 0.213162 | -1026.077261 | -98.6 | -1025.798662 | -1025.861087 | 13.5 | -1020.254562 | -4.284850 | -1024.360786 |
| conformer_116 | -1026.035465 | 0.278988 | 0.213915 | -1026.077244 | -109.7 | -1025.798256 | -1025.860317 | 15.5 | -1020.251243 | -4.283456 | -1024.359551 |
| conformer_18 | -1026.041824 | 0.278504 | 0.213592 | -1026.077225 | -92.9 | -1025.798721 | -1025.860621 | 14.7 | -1020.254601 | -4.286197 | -1024.359595 |
| conformer_110 | -1026.035690 | 0.278999 | 0.213870 | -1026.077214 | -109.0 | -1025.798215 | -1025.860332 | 15.5 | -1020.251466 | -4.283435 | -1024.359543 |
| conformer_140 | -1026.036657 | 0.278444 | 0.211718 | -1026.077212 | -106.5 | -1025.798768 | -1025.862482 | 9.8 | -1020.254213 | -4.282072 | -1024.362110 |
| conformer_119 | -1026.037980 | 0.278464 | 0.212535 | -1026.077196 | -103.0 | -1025.798732 | -1025.861649 | 12.0 | -1020.253086 | -4.283743 | -1024.360496 |
| conformer_303 | -1026.038620 | 0.278545 | 0.213215 | -1026.077177 | -101.2 | -1025.798632 | -1025.860950 | 13.8 | -1020.251632 | -4.285276 | -1024.359238 |
| conformer_197 | -1026.033903 | 0.278679 | 0.212525 | -1026.077160 | -113.6 | -1025.798481 | -1025.861623 | 12.1 | -1020.250618 | -4.282619 | -1024.360956 |
| conformer_179 | -1026.032734 | 0.278597 | 0.212974 | -1026.077154 | -116.6 | -1025.798557 | -1025.861168 | 13.3 | -1020.245726 | -4.285701 | -1024.359861 |
| conformer_215 | -1026.032431 | 0.278533 | 0.212852 | -1026.077150 | -117.4 | -1025.798617 | -1025.861286 | 13.0 | -1020.245400 | -4.285744 | -1024.359999 |
| conformer_204 | -1026.040778 | 0.278487 | 0.213354 | -1026.077146 | -95.5 | -1025.798659 | -1025.860780 | 14.3 | -1020.253785 | -4.285929 | -1024.359715 |
| conformer_122 | -1026.040696 | 0.278795 | 0.215858 | -1026.077118 | -95.6 | -1025.798323 | -1025.858248 | 20.9 | -1020.247947 | -4.291317 | -1024.356816 |
| conformer_149 | -1026.033722 | 0.278532 | 0.212893 | -1026.077083 | -113.8 | -1025.798551 | -1025.861178 | 13.2 | -1020.246668 | -4.285061 | -1024.359185 |
| conformer_306 | -1026.029017 | 0.278282 | 0.211312 | -1026.077054 | -126.1 | -1025.798772 | -1025.862730 | 9.2 | -1020.244710 | -4.283083 | -1024.361506 |
| conformer_270 | -1026.033880 | 0.278441 | 0.212195 | -1026.077052 | -113.3 | -1025.798611 | -1025.861845 | 11.5 | -1020.250381 | -4.283350 | -1024.361696 |
| conformer_266 | -1026.034389 | 0.278441 | 0.212421 | -1026.076948 | -111.7 | -1025.798507 | -1025.861515 | 12.4 | -1020.249492 | -4.283438 | -1024.360056 |
| conformer_152 | -1026.033795 | 0.278825 | 0.214549 | -1026.076926 | -113.2 | -1025.798101 | -1025.859365 | 18.0 | -1020.242503 | -4.288888 | -1024.356961 |
| conformer_247 | -1026.033777 | 0.278538 | 0.211803 | -1026.076909 | -113.2 | -1025.798371 | -1025.862094 | 10.8 | -1020.252353 | -4.281174 | -1024.361845 |
| conformer_220 | -1026.036509 | 0.278592 | 0.213013 | -1026.076805 | -105.8 | -1025.798213 | -1025.860780 | 14.3 | -1020.252431 | -4.283953 | -1024.360655 |
| conformer_76 | -1026.039238 | 0.278600 | 0.213391 | -1026.076793 | -98.6 | -1025.798193 | -1025.860390 | 15.3 | -1020.254161 | -4.284734 | -1024.360047 |
| conformer_262 | -1026.033679 | 0.278422 | 0.211431 | -1026.076782 | -113.2 | -1025.798360 | -1025.862339 | 10.2 | -1020.252334 | -4.281178 | -1024.362172 |
| conformer_91 | -1026.039142 | 0.278317 | 0.213095 | -1026.076752 | -98.7 | -1025.798435 | -1025.860645 | 14.6 | -1020.252367 | -4.285988 | -1024.359857 |
| conformer_32 | -1026.037494 | 0.278452 | 0.212919 | -1026.076653 | -102.8 | -1025.798201 | -1025.860722 | 14.4 | -1020.251967 | -4.284650 | -1024.359845 |
| conformer_77 | -1026.036560 | 0.278314 | 0.212575 | -1026.076646 | -105.2 | -1025.798332 | -1025.861059 | 13.5 | -1020.250608 | -4.284944 | -1024.360050 |
| conformer_22 | -1026.039805 | 0.278805 | 0.213525 | -1026.076639 | -96.7 | -1025.797834 | -1025.860102 | 16.1 | -1020.255870 | -4.283796 | -1024.359962 |

| | | | | | | | | | | | |
|---|---|---|---|---|---|---|---|---|---|---|---|
| conformer_64 | -1026.039744 | 0.278779 | 0.213178 | -1026.076625 | -96.8 | -1025.797846 | -1025.860435 | 15.2 | -1020.255842 | -4.283861 | -1024.360395 |
| conformer_170 | -1026.034135 | 0.278608 | 0.213145 | -1026.076577 | -111.4 | -1025.797969 | -1025.860420 | 15.2 | -1020.250582 | -4.283132 | -1024.359999 |
| conformer_35 | -1026.037117 | 0.278405 | 0.212605 | -1026.076519 | -103.5 | -1025.798114 | -1025.860902 | 14.0 | -1020.251426 | -4.284822 | -1024.360032 |
| conformer_249 | -1026.033795 | 0.278544 | 0.213489 | -1026.076497 | -112.1 | -1025.797953 | -1025.859996 | 16.3 | -1020.249023 | -4.284130 | -1024.359353 |
| conformer_160 | -1026.034220 | 0.278501 | 0.212820 | -1026.076477 | -110.9 | -1025.797976 | -1025.860645 | 14.6 | -1020.250750 | -4.283113 | -1024.360288 |
| conformer_269 | -1026.033647 | 0.278565 | 0.213544 | -1026.076450 | -112.4 | -1025.797885 | -1025.859894 | 16.6 | -1020.248859 | -4.284128 | -1024.359234 |
| conformer_272 | -1026.034561 | 0.278630 | 0.212271 | -1026.076439 | -110.0 | -1025.797809 | -1025.861156 | 13.3 | -1020.252213 | -4.282099 | -1024.360907 |
| conformer_261 | -1026.032913 | 0.278606 | 0.213734 | -1026.076438 | -114.3 | -1025.797832 | -1025.859692 | 17.1 | -1020.246087 | -4.285758 | -1024.358623 |
| conformer_291 | -1026.032023 | 0.278826 | 0.215199 | -1026.076431 | -116.6 | -1025.797605 | -1025.858220 | 21.0 | -1020.239779 | -4.290479 | -1024.356455 |
| conformer_178 | -1026.031903 | 0.278685 | 0.213506 | -1026.076421 | -116.9 | -1025.797736 | -1025.859903 | 16.6 | -1020.246080 | -4.284814 | -1024.358894 |
| conformer_83 | -1026.036055 | 0.278258 | 0.212268 | -1026.076385 | -105.9 | -1025.798127 | -1025.861105 | 13.4 | -1020.249948 | -4.285151 | -1024.360149 |
| conformer_280 | -1026.032666 | 0.278645 | 0.213808 | -1026.076337 | -114.7 | -1025.797692 | -1025.859517 | 17.6 | -1020.245811 | -4.285761 | -1024.358423 |
| conformer_164 | -1026.032041 | 0.278602 | 0.213237 | -1026.076301 | -116.2 | -1025.797699 | -1025.860052 | 16.2 | -1020.246311 | -4.284807 | -1024.359129 |
| conformer_109 | -1026.037013 | 0.278240 | 0.212686 | -1026.076274 | -103.1 | -1025.798034 | -1025.860576 | 14.8 | -1020.251059 | -4.285565 | -1024.360187 |
| conformer_172 | -1026.035003 | 0.278540 | 0.212371 | -1026.076271 | -108.3 | -1025.797731 | -1025.860888 | 14.0 | -1020.251466 | -4.283179 | -1024.360530 |
| conformer_20 | -1026.039453 | 0.278782 | 0.213259 | -1026.076258 | -96.6 | -1025.797476 | -1025.859987 | 16.4 | -1020.256696 | -4.282812 | -1024.360042 |
| conformer_166 | -1026.035043 | 0.278518 | 0.212304 | -1026.076215 | -108.1 | -1025.797697 | -1025.860899 | 14.0 | -1020.251517 | -4.283209 | -1024.360583 |
| conformer_264 | -1026.034292 | 0.278426 | 0.212366 | -1026.076106 | -109.8 | -1025.797680 | -1025.860728 | 14.4 | -1020.246921 | -4.285684 | -1024.359041 |
| conformer_21 | -1026.043533 | 0.278823 | 0.215263 | -1026.076094 | -85.5 | -1025.797271 | -1025.857819 | 22.1 | -1020.252049 | -4.289806 | -1024.356141 |
| conformer_26 | -1026.039330 | 0.278768 | 0.213377 | -1026.076074 | -96.5 | -1025.797306 | -1025.859685 | 17.2 | -1020.256590 | -4.282874 | -1024.359820 |
| conformer_232 | -1026.035112 | 0.278582 | 0.212241 | -1026.075891 | -107.1 | -1025.797309 | -1025.860638 | 14.7 | -1020.250955 | -4.283656 | -1024.360136 |
| conformer_281 | -1026.030383 | 0.278362 | 0.212087 | -1026.075847 | -119.4 | -1025.797485 | -1025.860748 | 14.4 | -1020.243259 | -4.285592 | -1024.359216 |
| conformer_47 | -1026.037804 | 0.278355 | 0.212562 | -1026.075818 | -99.8 | -1025.797463 | -1025.860244 | 15.7 | -1020.253102 | -4.284293 | -1024.359836 |
| conformer_107 | -1026.037139 | 0.278387 | 0.212728 | -1026.075813 | -101.5 | -1025.797426 | -1025.860073 | 16.1 | -1020.252088 | -4.284466 | -1024.359487 |
| conformer_102 | -1026.036873 | 0.278222 | 0.212277 | -1026.075767 | -102.1 | -1025.797545 | -1025.860478 | 15.1 | -1020.251703 | -4.284652 | -1024.359960 |
| conformer_134 | -1026.035363 | 0.278826 | 0.213724 | -1026.075724 | -106.0 | -1025.796898 | -1025.858988 | 19.0 | -1020.250867 | -4.283960 | -1024.358453 |
| conformer_214 | -1026.032298 | 0.278873 | 0.213290 | -1026.075714 | -114.0 | -1025.796841 | -1025.859412 | 17.9 | -1020.247248 | -4.284018 | -1024.358379 |
| conformer_146 | -1026.035163 | 0.278754 | 0.213550 | -1026.075624 | -106.2 | -1025.796870 | -1025.859062 | 18.8 | -1020.250709 | -4.283962 | -1024.358570 |
| conformer_137 | -1026.035055 | 0.278642 | 0.213157 | -1026.075595 | -106.4 | -1025.796953 | -1025.859426 | 17.8 | -1020.252271 | -4.282536 | -1024.359179 |
| conformer_218 | -1026.033067 | 0.278726 | 0.212989 | -1026.075593 | -111.7 | -1025.796867 | -1025.859592 | 17.4 | -1020.249178 | -4.282992 | -1024.358695 |
| conformer_304 | -1026.034666 | 0.278651 | 0.214716 | -1026.075551 | -107.3 | -1025.796900 | -1025.857823 | 22.0 | -1020.242872 | -4.289624 | -1024.355653 |
| conformer_295 | -1026.029085 | 0.278637 | 0.214861 | -1026.075532 | -121.9 | -1025.796895 | -1025.857659 | 22.5 | -1020.235614 | -4.292031 | -1024.356219 |
| conformer_95 | -1026.037485 | 0.278709 | 0.213197 | -1026.075488 | -99.8 | -1025.796779 | -1025.859279 | 18.2 | -1020.254088 | -4.283307 | -1024.359190 |
| conformer_94 | -1026.037555 | 0.278779 | 0.213496 | -1026.075468 | -99.5 | -1025.796689 | -1025.858960 | 19.1 | -1020.254096 | -4.283286 | -1024.358787 |
| conformer_246 | -1026.032818 | 0.278714 | 0.212860 | -1026.075465 | -112.0 | -1025.796751 | -1025.859593 | 17.4 | -1020.248901 | -4.282982 | -1024.358657 |
| conformer_131 | -1026.035119 | 0.278555 | 0.212891 | -1026.075440 | -105.9 | -1025.796885 | -1025.859537 | 17.5 | -1020.252411 | -4.282520 | -1024.359349 |
| conformer_199 | -1026.031767 | 0.278726 | 0.213699 | -1026.075413 | -114.6 | -1025.796687 | -1025.858702 | 19.7 | -1020.246130 | -4.284655 | -1024.357720 |
| conformer_240 | -1026.035546 | 0.278461 | 0.212114 | -1026.075413 | -104.7 | -1025.796952 | -1025.860287 | 15.6 | -1020.252901 | -4.282714 | -1024.360356 |
| conformer_48 | -1026.037517 | 0.278557 | 0.212507 | -1026.075368 | -99.4 | -1025.796811 | -1025.859849 | 16.7 | -1020.253763 | -4.283349 | -1024.359444 |
| conformer_57 | -1026.037298 | 0.278608 | 0.212894 | -1026.075355 | -99.9 | -1025.796747 | -1025.859449 | 17.8 | -1020.254925 | -4.282733 | -1024.359809 |
| conformer_67 | -1026.037060 | 0.278600 | 0.212961 | -1026.075144 | -100.0 | -1025.796544 | -1025.859171 | 18.5 | -1020.254672 | -4.282824 | -1024.359607 |
| conformer_212 | -1026.032421 | 0.278587 | 0.212756 | -1026.075057 | -111.9 | -1025.796470 | -1025.859289 | 18.2 | -1020.247520 | -4.284215 | -1024.358603 |
| conformer_226 | -1026.035650 | 0.278560 | 0.212222 | -1026.074959 | -103.2 | -1025.796399 | -1025.859725 | 17.0 | -1020.252348 | -4.283233 | -1024.359656 |
| conformer_169 | -1026.037264 | 0.278283 | 0.212259 | -1026.074914 | -98.8 | -1025.796631 | -1025.859643 | 17.3 | -1020.253003 | -4.284152 | -1024.359534 |
| conformer_202 | -1026.032535 | 0.278509 | 0.212515 | -1026.074873 | -111.2 | -1025.796364 | -1025.859346 | 18.0 | -1020.247739 | -4.284186 | -1024.358736 |
| conformer_132 | -1026.037374 | 0.278597 | 0.212945 | -1026.074845 | -98.4 | -1025.796248 | -1025.858888 | 19.2 | -1020.255424 | -4.282479 | -1024.359417 |
| conformer_74 | -1026.038213 | 0.278486 | 0.212264 | -1026.074842 | -96.2 | -1025.796356 | -1025.859566 | 17.5 | -1020.255327 | -4.282817 | -1024.359497 |
| conformer_301 | -1026.031574 | 0.278398 | 0.212398 | -1026.074814 | -113.5 | -1025.796416 | -1025.859404 | 17.9 | -1020.248343 | -4.282650 | -1024.358823 |
| conformer_311 | -1026.031438 | 0.278410 | 0.212411 | -1026.074791 | -113.8 | -1025.796381 | -1025.859368 | 18.0 | -1020.248167 | -4.282671 | -1024.358768 |
| conformer_294 | -1026.029599 | 0.278476 | 0.212739 | -1026.074731 | -118.5 | -1025.796255 | -1025.858980 | 19.0 | -1020.244076 | -4.284353 | -1024.357810 |
| conformer_308 | -1026.029374 | 0.278488 | 0.212789 | -1026.074676 | -118.9 | -1025.796188 | -1025.858875 | 19.3 | -1020.243792 | -4.284387 | -1024.357680 |
| conformer_299 | -1026.030344 | 0.278610 | 0.212952 | -1026.074575 | -116.1 | -1025.795965 | -1025.858611 | 20.0 | -1020.242259 | -4.287315 | -1024.357842 |
| conformer_133 | -1026.038254 | 0.278703 | 0.213088 | -1026.074487 | -95.1 | -1025.795784 | -1025.858387 | 20.6 | -1020.255606 | -4.282930 | -1024.358669 |
| conformer_80 | -1026.038303 | 0.278774 | 0.213326 | -1026.074453 | -94.9 | -1025.795679 | -1025.858115 | 21.3 | -1020.255546 | -4.282981 | -1024.358339 |
| conformer_314 | -1026.031661 | 0.278782 | 0.214290 | -1026.074443 | -112.3 | -1025.795661 | -1025.857141 | 23.8 | -1020.240749 | -4.289768 | -1024.355997 |
| conformer_5 | -1026.039275 | 0.278943 | 0.214052 | -1026.074422 | -92.3 | -1025.795479 | -1025.857358 | 23.3 | -1020.255803 | -4.283189 | -1024.357075 |

| | | | | | | | | | | | |
|---|---|---|---|---|---|---|---|---|---|---|---|
| conformer_63 | -1026.036625 | 0.278489 | 0.212234 | -1026.074358 | -99.1 | -1025.795869 | -1025.859112 | 18.7 | -1020.254109 | -4.282503 | -1024.359098 |
| conformer_73 | -1026.036534 | 0.278471 | 0.212285 | -1026.074275 | -99.1 | -1025.795804 | -1025.858978 | 19.0 | -1020.253994 | -4.282605 | -1024.359044 |
| conformer_277 | -1026.032553 | 0.278539 | 0.212835 | -1026.074073 | -109.0 | -1025.795534 | -1025.858226 | 21.0 | -1020.248364 | -4.283495 | -1024.357531 |
| conformer_8 | -1026.039043 | 0.278873 | 0.213700 | -1026.074026 | -91.8 | -1025.795153 | -1025.857314 | 23.4 | -1020.255625 | -4.283259 | -1024.357154 |
| conformer_288 | -1026.032530 | 0.278896 | 0.214929 | -1026.073859 | -108.5 | -1025.794963 | -1025.855918 | 27.0 | -1020.240480 | -4.290655 | -1024.354523 |
| conformer_123 | -1026.034385 | 0.278328 | 0.212009 | -1026.073827 | -103.6 | -1025.795499 | -1025.858806 | 19.5 | -1020.252170 | -4.282346 | -1024.358937 |
| conformer_190 | -1026.034793 | 0.278511 | 0.212623 | -1026.073826 | -102.5 | -1025.795315 | -1025.858191 | 21.1 | -1020.251605 | -4.282889 | -1024.357892 |
| conformer_191 | -1026.034781 | 0.278509 | 0.212505 | -1026.073811 | -102.5 | -1025.795302 | -1025.858294 | 20.8 | -1020.251590 | -4.282909 | -1024.358012 |
| conformer_168 | -1026.031638 | 0.278802 | 0.214822 | -1026.073793 | -110.7 | -1025.794991 | -1025.855959 | 26.9 | -1020.241026 | -4.289077 | -1024.354424 |
| conformer_257 | -1026.032474 | 0.278445 | 0.212421 | -1026.073781 | -108.5 | -1025.795336 | -1025.858348 | 20.7 | -1020.249987 | -4.282070 | -1024.357931 |
| conformer_267 | -1026.032388 | 0.278482 | 0.212570 | -1026.073771 | -108.7 | -1025.795289 | -1025.858189 | 21.1 | -1020.249854 | -4.282097 | -1024.357752 |
| conformer_254 | -1026.032715 | 0.278576 | 0.212745 | -1026.073673 | -107.5 | -1025.795097 | -1025.857916 | 21.8 | -1020.248503 | -4.283589 | -1024.357294 |
| conformer_148 | -1026.034330 | 0.278839 | 0.215383 | -1026.073594 | -103.1 | -1025.794755 | -1025.855199 | 28.9 | -1020.239194 | -4.292172 | -1024.352235 |
| conformer_87 | -1026.037233 | 0.278435 | 0.212126 | -1026.073507 | -95.2 | -1025.795072 | -1025.858369 | 20.6 | -1020.255629 | -4.282012 | -1024.358777 |
| conformer_162 | -1026.033776 | 0.278276 | 0.211886 | -1026.073505 | -104.3 | -1025.795229 | -1025.858607 | 20.0 | -1020.251468 | -4.282585 | -1024.358884 |
| conformer_268 | -1026.034495 | 0.278326 | 0.212061 | -1026.073301 | -101.9 | -1025.794975 | -1025.858228 | 21.0 | -1020.252738 | -4.282069 | -1024.358540 |
| conformer_315 | -1026.033843 | 0.278272 | 0.211878 | -1026.072985 | -102.8 | -1025.794713 | -1025.858095 | 21.3 | -1020.251993 | -4.282252 | -1024.358497 |
| conformer_156 | -1026.035520 | 0.278510 | 0.212409 | -1026.072891 | -98.1 | -1025.794381 | -1025.857470 | 23.0 | -1020.253057 | -4.282538 | -1024.357544 |
| conformer_154 | -1026.035526 | 0.278529 | 0.212512 | -1026.072874 | -98.1 | -1025.794345 | -1025.857350 | 23.3 | -1020.253084 | -4.282540 | -1024.357448 |
| conformer_255 | -1026.030117 | 0.278770 | 0.213268 | -1026.072837 | -112.2 | -1025.794067 | -1025.856557 | 25.4 | -1020.240979 | -4.288471 | -1024.355890 |
| conformer_34 | -1026.036384 | 0.278600 | 0.212684 | -1026.072698 | -95.3 | -1025.794098 | -1025.857002 | 24.2 | -1020.253163 | -4.282782 | -1024.356563 |
| conformer_225 | -1026.034355 | 0.278215 | 0.211627 | -1026.072686 | -100.6 | -1025.794471 | -1025.858047 | 21.5 | -1020.252971 | -4.281983 | -1024.358647 |
| conformer_252 | -1026.033109 | 0.278673 | 0.213720 | -1026.072653 | -103.8 | -1025.793980 | -1025.855921 | 27.0 | -1020.246884 | -4.286291 | -1024.355987 |
| conformer_42 | -1026.036053 | 0.278576 | 0.212786 | -1026.072564 | -95.9 | -1025.793988 | -1025.856766 | 24.8 | -1020.252779 | -4.282990 | -1024.356482 |
| conformer_222 | -1026.033364 | 0.278674 | 0.213751 | -1026.072284 | -102.2 | -1025.793610 | -1025.855521 | 28.1 | -1020.246485 | -4.286867 | -1024.355509 |
| conformer_239 | -1026.033601 | 0.278515 | 0.211856 | -1026.072071 | -101.0 | -1025.793556 | -1025.857203 | 23.7 | -1020.251704 | -4.282302 | -1024.357608 |
| conformer_245 | -1026.035908 | 0.278446 | 0.212206 | -1026.072047 | -94.9 | -1025.793601 | -1025.856829 | 24.7 | -1020.254398 | -4.282001 | -1024.357319 |
| conformer_217 | -1026.029705 | 0.278668 | 0.214584 | -1026.071881 | -110.7 | -1025.793213 | -1025.854285 | 31.3 | -1020.237670 | -4.290823 | -1024.353072 |
| conformer_60 | -1026.036684 | 0.278493 | 0.212385 | -1026.071804 | -92.2 | -1025.793311 | -1025.856407 | 25.8 | -1020.254368 | -4.282360 | -1024.356451 |
| conformer_253 | -1026.033430 | 0.278476 | 0.211509 | -1026.071701 | -100.5 | -1025.793225 | -1025.857180 | 23.7 | -1020.251422 | -4.282549 | -1024.357720 |
| conformer_198 | -1026.031326 | 0.278721 | 0.214368 | -1026.071551 | -105.6 | -1025.792830 | -1025.854171 | 31.6 | -1020.241843 | -4.288963 | -1024.353651 |
| conformer_282 | -1026.027490 | 0.279146 | 0.214830 | -1026.067167 | -104.2 | -1025.788021 | -1025.849325 | 44.4 | -1020.236427 | -4.290465 | -1024.348728 |
| conformer_289 | -1026.027464 | 0.279079 | 0.214888 | -1026.067075 | -104.0 | -1025.787996 | -1025.849175 | 44.7 | -1020.236486 | -4.290341 | -1024.348538 |

[a]: Single-point calculation in aqueous phase with SMD model.

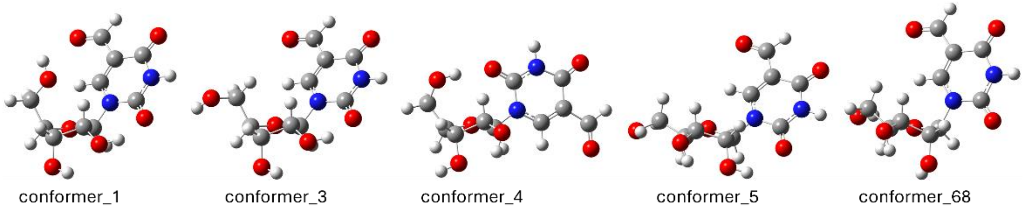

**Figure S44.** B3LYP-D3/def2-TZVPP optimized geometries of conformers for 5-formyluridine (**1rb5fU**).

**Table S42.** Conformers of gas-phase optimized 5-formyluridine (**1rb5fU**) at the B3LYP-D3/def2-TZVPP level of theory followed by aqueous phase single-point calculation. The columns display total energy without zero-point correction ($E_{Tot}$), Gibbs free energy (δG), total energy without zero-point correction ($E_{Tot,W}$), Gibbs free energy ($G_{298,W}$) in water (W), total single-point energy ($E_{CBS}$) calculated at DLPNO-CCSD(T)/CBS level of theory, and their corresponding free energy $G_{CBS}$. $G_{298,W}$ and $G_{CBS}$ have been corrected to the standard state of 1 mol/L by addition of +7.908 kJ/mol. $\Delta G_{Solv}$ represents the Gibbs free energy of solvation. The data are arranged in the ascending numeric order of $E_{Tot,W}$. $\Delta G_{298,W}$ represents the respective energy difference to the lowest structure. Only conformers within the 24 kJ/mol (6 kcal/mol) energy window above the lowest in CREST are included in initial conformer sampling. Duplicates of the same structure are excluded.

| 1rb5fU | B3LYP-D3/def2-TZVPP | | | SMD($H_2O$)/B3LYP-D3/def2-TZVPP[a] | | | | | DLPNO-CCSD(T)/CBS | | |
|---|---|---|---|---|---|---|---|---|---|---|---|
| No. | $E_{Tot}$ (Hartree) | δH (Hartree) | δG (Hartree) | $E_{Tot,W}$ (Hartree) | $\Delta G_{Solv}$ (kJ/mol) | $H_{298,W}$ (Hartree) | $G_{298,W}$ (Hartree) | $\Delta G_{298,W}$ (kJ/mol) | $E_{CBS,HF}$ (Hartree) | $E_{CBS,C}$ (Hartree) | $G_{CBS,W}$ (Hartree) |
| conformer_30 | -1024.828993 | 0.254407 | 0.191337 | -1024.866338 | -98.0 | -1024.611931 | -1024.671989 | 3.8 | -1019.089791 | -4.237052 | -1023.169839 |
| conformer_7 | -1024.826487 | 0.254368 | 0.191228 | -1024.866328 | -104.6 | -1024.611960 | -1024.672088 | 3.5 | -1019.085388 | -4.238130 | -1023.169120 |
| conformer_4 | -1024.828768 | 0.254343 | 0.191101 | -1024.866310 | -98.6 | -1024.611967 | -1024.672197 | 3.3 | -1019.089528 | -4.236767 | -1023.169724 |
| conformer_46 | -1024.825397 | 0.254397 | 0.190763 | -1024.865463 | -105.2 | -1024.611066 | -1024.671688 | 4.6 | -1019.086239 | -4.236565 | -1023.169094 |
| **conformer_68** | **-1024.820937** | **0.253952** | **0.188885** | **-1024.865333** | **-116.6** | **-1024.611381** | **-1024.673436** | **0** | **-1019.080861** | **-4.237349** | **-1023.170709** |
| conformer_9 | -1024.829692 | 0.254134 | 0.189643 | -1024.865231 | -93.3 | -1024.611097 | -1024.672576 | 2.3 | -1019.090698 | -4.235865 | -1023.169447 |
| conformer_69 | -1024.825848 | 0.254177 | 0.188678 | -1024.865115 | -103.1 | -1024.610938 | -1024.673425 | 0.1 | -1019.089602 | -4.234018 | -1023.171197 |
| conformer_1 | -1024.833143 | 0.254254 | 0.191000 | -1024.864881 | -83.3 | -1024.610627 | -1024.670869 | 6.7 | -1019.092450 | -4.237925 | -1023.168100 |
| conformer_14 | -1024.827435 | 0.253997 | 0.189656 | -1024.864751 | -98.0 | -1024.610754 | -1024.672083 | 3.6 | -1019.087875 | -4.236941 | -1023.169464 |
| conformer_53 | -1024.820333 | 0.253985 | 0.188402 | -1024.864682 | -116.4 | -1024.610697 | -1024.673268 | 0.4 | -1019.081926 | -4.235844 | -1023.170706 |
| conformer_12 | -1024.827466 | 0.254217 | 0.190489 | -1024.864503 | -97.2 | -1024.610286 | -1024.671002 | 6.4 | -1019.088318 | -4.236955 | -1023.168809 |
| conformer_10 | -1024.826574 | 0.254027 | 0.190524 | -1024.864319 | -99.1 | -1024.610292 | -1024.670783 | 7.0 | -1019.086708 | -4.237877 | -1023.168794 |
| conformer_52 | -1024.821683 | 0.254002 | 0.188457 | -1024.864299 | -111.9 | -1024.610297 | -1024.672830 | 1.6 | -1019.085314 | -4.234243 | -1023.170703 |
| conformer_16 | -1024.826835 | 0.254175 | 0.189977 | -1024.864228 | -98.2 | -1024.610053 | -1024.671239 | 5.8 | -1019.088425 | -4.235182 | -1023.168011 |
| conformer_6 | -1024.825863 | 0.254257 | 0.190395 | -1024.864222 | -100.7 | -1024.609965 | -1024.670815 | 6.9 | -1019.085196 | -4.236973 | -1023.167121 |
| conformer_62 | -1024.828080 | 0.254213 | 0.190996 | -1024.864200 | -94.8 | -1024.609987 | -1024.670192 | 8.5 | -1019.085589 | -4.238696 | -1023.166398 |
| conformer_20 | -1024.829193 | 0.254116 | 0.191902 | -1024.864019 | -91.4 | -1024.609903 | -1024.669105 | 11.4 | -1019.085950 | -4.240626 | -1023.166488 |
| conformer_3 | -1024.827047 | 0.254113 | 0.189880 | -1024.863935 | -96.9 | -1024.609822 | -1024.671043 | 6.3 | -1019.088510 | -4.236361 | -1023.168867 |
| conformer_28 | -1024.823387 | 0.254330 | 0.189490 | -1024.863929 | -106.4 | -1024.609599 | -1024.671427 | 5.3 | -1019.086738 | -4.234678 | -1023.169456 |
| conformer_15 | -1024.825685 | 0.253980 | 0.189579 | -1024.863856 | -100.2 | -1024.609876 | -1024.671265 | 5.7 | -1019.086706 | -4.236708 | -1023.168995 |
| conformer_49 | -1024.824518 | 0.254113 | 0.189454 | -1024.863727 | -102.9 | -1024.609614 | -1024.671261 | 5.7 | -1019.086319 | -4.235963 | -1023.169025 |
| conformer_34 | -1024.819030 | 0.253983 | 0.188357 | -1024.863653 | -117.2 | -1024.609670 | -1024.672284 | 3.0 | -1019.081285 | -4.235111 | -1023.169651 |
| conformer_74 | -1024.818008 | 0.253814 | 0.187393 | -1024.863580 | -119.6 | -1024.609766 | -1024.673175 | 0.7 | -1019.080281 | -4.235099 | -1023.170547 |
| conformer_26 | -1024.825555 | 0.254278 | 0.191495 | -1024.863473 | -99.6 | -1024.609195 | -1024.668966 | 11.7 | -1019.080731 | -4.240556 | -1023.164698 |
| conformer_36 | -1024.823260 | 0.254231 | 0.190492 | -1024.863332 | -105.2 | -1024.609101 | -1024.669828 | 9.5 | -1019.085320 | -4.235911 | -1023.167799 |
| conformer_56 | -1024.820122 | 0.254105 | 0.189489 | -1024.863243 | -113.2 | -1024.609138 | -1024.670742 | 7.1 | -1019.079666 | -4.237473 | -1023.167759 |
| conformer_45 | -1024.822006 | 0.254267 | 0.190590 | -1024.863180 | -108.1 | -1024.608913 | -1024.669578 | 10.1 | -1019.082052 | -4.237508 | -1023.167132 |
| conformer_18 | -1024.826822 | 0.254064 | 0.189756 | -1024.863078 | -95.2 | -1024.609014 | -1024.670310 | 8.2 | -1019.088666 | -4.236162 | -1023.168316 |
| conformer_42 | -1024.821234 | 0.253962 | 0.187787 | -1024.862951 | -109.5 | -1024.608989 | -1024.672152 | 3.4 | -1019.085964 | -4.233254 | -1023.170136 |
| conformer_23 | -1024.824424 | 0.254233 | 0.189533 | -1024.862941 | -101.1 | -1024.608708 | -1024.670396 | 8.0 | -1019.087551 | -4.235223 | -1023.168746 |
| conformer_39 | -1024.825675 | 0.254146 | 0.189657 | -1024.862823 | -97.5 | -1024.608677 | -1024.670154 | 8.6 | -1019.088065 | -4.235661 | -1023.168205 |

| | | | | | | | | | | | |
|---|---|---|---|---|---|---|---|---|---|---|---|
| conformer_63 | -1024.819597 | 0.253938 | 0.188191 | -1024.862711 | -113.2 | -1024.608773 | -1024.671508 | 5.1 | -1019.082286 | -4.234838 | -1023.169035 |
| conformer_37 | -1024.822138 | 0.253978 | 0.187976 | -1024.862348 | -105.6 | -1024.608370 | -1024.671360 | 5.5 | -1019.087327 | -4.232967 | -1023.169515 |
| conformer_35 | -1024.822358 | 0.254457 | 0.190060 | -1024.862182 | -104.6 | -1024.607725 | -1024.669110 | 11.4 | -1019.085077 | -4.235024 | -1023.166854 |
| conformer_11 | -1024.826201 | 0.254229 | 0.189510 | -1024.862060 | -94.1 | -1024.607831 | -1024.669538 | 10.2 | -1019.090569 | -4.234028 | -1023.167934 |
| conformer_43 | -1024.821638 | 0.254117 | 0.189484 | -1024.861684 | -105.1 | -1024.607567 | -1024.669188 | 11.2 | -1019.085104 | -4.234534 | -1023.167189 |
| conformer_51 | -1024.819312 | 0.254178 | 0.189786 | -1024.861628 | -111.1 | -1024.607450 | -1024.668830 | 12.1 | -1019.080504 | -4.236263 | -1023.166286 |
| conformer_24 | -1024.823343 | 0.254053 | 0.189112 | -1024.861155 | -99.3 | -1024.607102 | -1024.669031 | 11.6 | -1019.087994 | -4.234015 | -1023.167697 |
| conformer_61 | -1024.823941 | 0.254068 | 0.189197 | -1024.860753 | -96.7 | -1024.606685 | -1024.668544 | 12.8 | -1019.089077 | -4.233564 | -1023.167245 |
| conformer_47 | -1024.821973 | 0.254272 | 0.189939 | -1024.860742 | -101.8 | -1024.606470 | -1024.667791 | 14.8 | -1019.084392 | -4.235573 | -1023.165782 |
| conformer_32 | -1024.822519 | 0.254168 | 0.189555 | -1024.860658 | -100.1 | -1024.606490 | -1024.668091 | 14.0 | -1019.086739 | -4.233941 | -1023.166252 |
| conformer_22 | -1024.824534 | 0.254235 | 0.189637 | -1024.860416 | -94.2 | -1024.606181 | -1024.667767 | 14.9 | -1019.088017 | -4.234908 | -1023.166159 |
| conformer_5 | -1024.825670 | 0.254343 | 0.189957 | -1024.860340 | -91.0 | -1024.605997 | -1024.667371 | 15.9 | -1019.089252 | -4.234483 | -1023.165437 |
| conformer_70 | -1024.819795 | 0.254045 | 0.188611 | -1024.860109 | -105.8 | -1024.606064 | -1024.668486 | 13.0 | -1019.081988 | -4.235611 | -1023.166290 |
| conformer_21 | -1024.825285 | 0.254241 | 0.189640 | -1024.859519 | -89.9 | -1024.605278 | -1024.666867 | 17.2 | -1019.089523 | -4.234501 | -1023.165606 |
| conformer_55 | -1024.818525 | 0.254432 | 0.191634 | -1024.858711 | -105.5 | -1024.604279 | -1024.664065 | 24.6 | -1019.074759 | -4.240792 | -1023.161091 |
| conformer_72 | -1024.820239 | 0.254081 | 0.189577 | -1024.857657 | -98.2 | -1024.603576 | -1024.665068 | 22.0 | -1019.080689 | -4.238025 | -1023.163542 |

[a]: Single-point calculation in aqueous phase with SMD model.

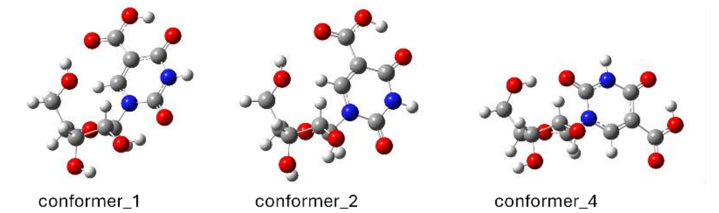

**Figure S45.** B3LYP-D3/def2-TZVPP optimized geometries of conformers for 5-carboxyluridine (**1rb5caU**).

**Table S43.** Conformers of gas-phase optimized 5-carboxyluridine (**1rb5caU**) at the B3LYP-D3/def2-TZVPP level of theory followed by aqueous phase single-point calculation. The columns display total energy without zero-point correction ($E_{Tot}$), Gibbs free energy ($\delta G$), total energy without zero-point correction ($E_{Tot,W}$), Gibbs free energy ($G_{298,W}$) in water (W), total single-point energy ($E_{CBS}$) calculated at DLPNO-CCSD(T)/CBS level of theory, and their corresponding free energy $G_{CBS}$. $G_{298,W}$ and $G_{CBS}$ have been corrected to the standard state of 1 mol/L by addition of +7.908 kJ/mol. $\Delta G_{Solv}$ represents the Gibbs free energy of solvation. The data are arranged in the ascending numeric order of $E_{Tot,W}$. $\Delta G_{298,W}$ represents the respective energy difference to the lowest structure. Only conformers within the 24 kJ/mol (6 kcal/mol) energy window above the lowest in CREST are included in initial conformer sampling. Duplicates of the same structure are excluded.

| | **B3LYP-D3/def2-TZVPP** | | | **SMD($H_2O$)/B3LYP-D3/def2-TZVPP[a]** | | | | | **DLPNO-CCSD(T)/CBS** | | |
|---|---|---|---|---|---|---|---|---|---|---|---|
| **1rb5caU No.** | **$E_{Tot}$** (Hartree) | **$\delta H$** (Hartree) | **$\delta G$** (Hartree) | **$E_{Tot,W}$** (Hartree) | **$\Delta G_{Solv}$** (kJ/mol) | **$H_{298,W}$** (Hartree) | **$G_{298,W}$** (Hartree) | **$\Delta G_{298,W}$** (kJ/mol) | **$E_{CBS,HF}$** (Hartree) | **$E_{CBS,C}$** (Hartree) | **$G_{CBS,W}$** (Hartree) |
| conformer_7 | -1100.118831 | 0.260925 | 0.196433 | -1100.161992 | -113.3 | -1099.901067 | -1099.962547 | 4.6 | -1094.016969 | -4.512051 | -1098.372736 |
| conformer_4 | -1100.121170 | 0.260887 | 0.196252 | -1100.161954 | -107.1 | -1099.901067 | -1099.962690 | 4.3 | -1094.021174 | -4.510702 | -1098.373395 |
| conformer_40 | -1100.121241 | 0.260959 | 0.196565 | -1100.161945 | -106.9 | -1099.900986 | -1099.962368 | 5.1 | -1094.021245 | -4.510974 | -1098.373347 |
| **conformer_19** | **-1100.118230** | **0.260687** | **0.194306** | **-1100.161629** | **-113.9** | **-1099.900942** | **-1099.964311** | **0** | **-1094.020716** | **-4.508367** | **-1098.375163** |
| conformer_76 | -1100.113566 | 0.260539 | 0.194353 | -1100.161302 | -125.3 | -1099.900763 | -1099.963937 | 1.0 | -1094.012637 | -4.511298 | -1098.374307 |
| conformer_70 | -1100.117636 | 0.260956 | 0.195932 | -1100.161161 | -114.3 | -1099.900205 | -1099.962217 | 5.5 | -1094.017679 | -4.510405 | -1098.372665 |
| conformer_10 | -1100.121148 | 0.260850 | 0.194894 | -1100.161031 | -104.7 | -1099.900181 | -1099.963125 | 3.1 | -1094.021345 | -4.509684 | -1098.373005 |
| conformer_1 | -1100.126246 | 0.260993 | 0.196498 | -1100.160804 | -90.7 | -1099.899811 | -1099.961294 | 7.9 | -1094.024057 | -4.512352 | -1098.371456 |
| conformer_75 | -1100.115003 | 0.260489 | 0.193584 | -1100.160704 | -120.0 | -1099.900215 | -1099.964108 | 0.5 | -1094.017873 | -4.508167 | -1098.375145 |
| conformer_12 | -1100.123016 | 0.260729 | 0.195206 | -1100.160642 | -98.8 | -1099.899913 | -1099.962424 | 5.0 | -1094.021552 | -4.511134 | -1098.372094 |
| conformer_57 | -1100.113095 | 0.260567 | 0.193703 | -1100.160463 | -124.4 | -1099.899896 | -1099.963748 | 1.5 | -1094.013902 | -4.509673 | -1098.374229 |
| conformer_14 | -1100.119681 | 0.260790 | 0.195667 | -1100.160167 | -106.3 | -1099.899377 | -1099.961488 | 7.4 | -1094.019731 | -4.510840 | -1098.372378 |
| conformer_53 | -1100.114639 | 0.260616 | 0.193952 | -1100.160146 | -119.5 | -1099.899530 | -1099.963182 | 3.0 | -1094.017454 | -4.508139 | -1098.374136 |
| conformer_17 | -1100.118575 | 0.260657 | 0.196009 | -1100.160015 | -108.8 | -1099.899358 | -1099.960994 | 8.7 | -1094.017692 | -4.511967 | -1098.372078 |
| conformer_26 | -1100.121923 | 0.260786 | 0.196563 | -1100.159925 | -99.8 | -1099.899139 | -1099.960350 | 10.4 | -1094.018106 | -4.513365 | -1098.369898 |
| conformer_16 | -1100.122343 | 0.260772 | 0.197507 | -1100.159919 | -98.7 | -1099.899147 | -1099.959400 | 12.9 | -1094.018084 | -4.514695 | -1098.369835 |
| conformer_6 | -1100.120667 | 0.260621 | 0.194911 | -1100.159818 | -102.8 | -1099.899197 | -1099.961895 | 6.3 | -1094.020928 | -4.509750 | -1098.371906 |
| conformer_5 | -1100.119252 | 0.260653 | 0.194838 | -1100.159790 | -106.4 | -1099.899137 | -1099.961940 | 6.2 | -1094.019879 | -4.510392 | -1098.372960 |
| conformer_2 | -1100.119665 | 0.260775 | 0.195490 | -1100.159704 | -105.1 | -1099.898929 | -1099.961202 | 8.2 | -1094.017368 | -4.511633 | -1098.370538 |
| conformer_37 | -1100.115750 | 0.260950 | 0.194946 | -1100.159682 | -115.3 | -1099.898732 | -1099.961724 | 6.8 | -1094.018248 | -4.508616 | -1098.372838 |
| conformer_48 | -1100.117551 | 0.260745 | 0.194984 | -1100.159645 | -110.5 | -1099.898900 | -1099.961649 | 7.0 | -1094.018393 | -4.510157 | -1098.372648 |
| conformer_36 | -1100.111921 | 0.260606 | 0.193666 | -1100.159583 | -125.1 | -1099.898977 | -1099.962905 | 3.7 | -1094.013204 | -4.509151 | -1098.373338 |
| conformer_20 | -1100.117750 | 0.260534 | 0.194723 | -1100.159502 | -109.6 | -1099.898968 | -1099.961767 | 6.7 | -1094.017974 | -4.510611 | -1098.372602 |
| conformer_66 | -1100.112673 | 0.260727 | 0.194938 | -1100.159275 | -122.4 | -1099.898548 | -1099.961325 | 7.8 | -1094.011285 | -4.511516 | -1098.371453 |
| conformer_81 | -1100.110927 | 0.260481 | 0.192788 | -1100.159271 | -126.9 | -1099.898790 | -1099.963471 | 2.2 | -1094.012286 | -4.509038 | -1098.373867 |
| conformer_18 | -1100.119346 | 0.260970 | 0.197426 | -1100.159200 | -104.6 | -1099.898230 | -1099.958762 | 14.6 | -1094.013293 | -4.515062 | -1098.367772 |
| conformer_41 | -1100.115713 | 0.260826 | 0.195792 | -1100.159034 | -113.7 | -1099.898208 | -1099.960230 | 10.7 | -1094.016959 | -4.509871 | -1098.371347 |
| conformer_22 | -1100.119147 | 0.260578 | 0.194687 | -1100.159032 | -104.7 | -1099.898454 | -1099.961333 | 7.8 | -1094.020027 | -4.510378 | -1098.372591 |
| conformer_60 | -1100.114372 | 0.260865 | 0.195928 | -1100.158911 | -116.9 | -1099.898046 | -1099.959971 | 11.4 | -1094.013620 | -4.511383 | -1098.370602 |
| conformer_38 | -1100.114264 | 0.260583 | 0.193329 | -1100.158901 | -117.2 | -1099.898318 | -1099.962560 | 4.6 | -1094.018048 | -4.507368 | -1098.373712 |
| conformer_30 | -1100.118929 | 0.260820 | 0.195317 | -1100.158764 | -104.6 | -1099.897944 | -1099.960435 | 10.2 | -1094.020318 | -4.509892 | -1098.371716 |

| | | | | | | | | | | | |
|---|---|---|---|---|---|---|---|---|---|---|---|
| conformer_31 | -1100.116751 | 0.260835 | 0.194904 | -1100.158685 | -110.1 | -1099.897850 | -1099.960769 | 9.3 | -1094.019012 | -4.509168 | -1098.372199 |
| conformer_52 | -1100.112733 | 0.260600 | 0.193713 | -1100.158676 | -120.6 | -1099.898076 | -1099.961951 | 6.2 | -1094.014315 | -4.509016 | -1098.372549 |
| conformer_33 | -1100.115352 | 0.260631 | 0.193563 | -1100.158366 | -112.9 | -1099.897735 | -1099.961791 | 6.6 | -1094.019579 | -4.506958 | -1098.372976 |
| conformer_15 | -1100.118379 | 0.260778 | 0.194815 | -1100.158135 | -104.4 | -1099.897357 | -1099.960308 | 10.5 | -1094.021870 | -4.507879 | -1098.371679 |
| conformer_84 | -1100.112706 | 0.260850 | 0.193465 | -1100.158023 | -119.0 | -1099.897173 | -1099.961546 | 7.3 | -1094.014394 | -4.508774 | -1098.372008 |
| conformer_50 | -1100.114723 | 0.261034 | 0.195367 | -1100.157918 | -113.4 | -1099.896884 | -1099.959539 | 12.5 | -1094.016605 | -4.508968 | -1098.370389 |
| conformer_83 | -1100.114427 | 0.260780 | 0.196870 | -1100.157820 | -113.9 | -1099.897040 | -1099.957938 | 16.7 | -1094.007310 | -4.517026 | -1098.367847 |
| conformer_54 | -1100.114284 | 0.260694 | 0.194780 | -1100.157427 | -113.3 | -1099.896733 | -1099.959635 | 12.3 | -1094.016974 | -4.508490 | -1098.370816 |
| conformer_61 | -1100.111884 | 0.260742 | 0.195051 | -1100.157378 | -119.4 | -1099.896636 | -1099.959315 | 13.1 | -1094.012292 | -4.510185 | -1098.369908 |
| conformer_44 | -1100.115360 | 0.260593 | 0.194323 | -1100.157254 | -110.0 | -1099.896661 | -1099.959919 | 11.5 | -1094.019106 | -4.507873 | -1098.371539 |
| conformer_69 | -1100.114275 | 0.260825 | 0.195204 | -1100.156435 | -110.7 | -1099.895610 | -1099.958219 | 16.0 | -1094.015865 | -4.509536 | -1098.369345 |
| conformer_39 | -1100.115129 | 0.260745 | 0.194872 | -1100.156401 | -108.4 | -1099.895656 | -1099.958517 | 15.2 | -1094.018566 | -4.507924 | -1098.369878 |
| conformer_55 | -1100.112390 | 0.260840 | 0.194217 | -1100.156382 | -115.5 | -1099.895542 | -1099.959153 | 13.5 | -1094.014583 | -4.508381 | -1098.369728 |
| conformer_9 | -1100.117724 | 0.260903 | 0.195196 | -1100.156314 | -101.3 | -1099.895411 | -1099.958106 | 16.3 | -1094.020465 | -4.508432 | -1098.369278 |
| conformer_27 | -1100.117012 | 0.260799 | 0.194876 | -1100.156118 | -102.7 | -1099.895319 | -1099.958230 | 16.0 | -1094.019681 | -4.508885 | -1098.369785 |
| conformer_46 | -1100.110546 | 0.260783 | 0.195046 | -1100.155889 | -119.0 | -1099.895106 | -1099.957831 | 17.0 | -1094.011400 | -4.509425 | -1098.368111 |
| conformer_32 | -1100.112730 | 0.260732 | 0.194808 | -1100.155852 | -113.2 | -1099.895120 | -1099.958032 | 16.5 | -1094.015487 | -4.508059 | -1098.368848 |
| conformer_23 | -1100.117743 | 0.260819 | 0.194947 | -1100.155217 | -98.4 | -1099.894398 | -1099.957258 | 18.5 | -1094.021166 | -4.508515 | -1098.369197 |
| conformer_43 | -1100.114145 | 0.260687 | 0.194847 | -1100.154872 | -106.9 | -1099.894185 | -1099.957013 | 19.2 | -1094.014910 | -4.508788 | -1098.366566 |
| conformer_58 | -1100.114314 | 0.260666 | 0.196394 | -1100.154505 | -105.5 | -1099.893839 | -1099.955099 | 24.2 | -1094.012701 | -4.512093 | -1098.365578 |
| conformer_56 | -1100.111161 | 0.260950 | 0.196722 | -1100.154497 | -113.8 | -1099.893547 | -1099.954763 | 25.1 | -1094.006432 | -4.514861 | -1098.364894 |
| conformer_71 | -1100.111519 | 0.260695 | 0.194465 | -1100.154012 | -111.6 | -1099.893317 | -1099.956535 | 20.4 | -1094.014247 | -4.508231 | -1098.367493 |
| conformer_11 | -1100.117883 | 0.260990 | 0.196801 | -1100.153865 | -94.5 | -1099.892875 | -1099.954052 | 26.9 | -1094.016381 | -4.511080 | -1098.363630 |
| conformer_67 | -1100.110452 | 0.260533 | 0.194586 | -1100.153702 | -113.6 | -1099.893169 | -1099.956104 | 21.5 | -1094.012328 | -4.509380 | -1098.367360 |
| conformer_78 | -1100.113209 | 0.260746 | 0.195584 | -1100.153561 | -105.9 | -1099.892815 | -1099.954965 | 24.5 | -1094.012342 | -4.512374 | -1098.366472 |
| conformer_24 | -1100.111051 | 0.260513 | 0.193276 | -1100.153464 | -111.4 | -1099.892951 | -1099.957176 | 18.7 | -1094.014457 | -4.507666 | -1098.368247 |
| conformer_45 | -1100.110166 | 0.260652 | 0.194543 | -1100.153392 | -113.5 | -1099.892740 | -1099.955837 | 22.2 | -1094.010277 | -4.509195 | -1098.365143 |
| conformer_77 | -1100.109784 | 0.260416 | 0.193307 | -1100.153217 | -114.0 | -1099.892801 | -1099.956898 | 19.5 | -1094.012753 | -4.507921 | -1098.367788 |
| conformer_62 | -1100.110228 | 0.260616 | 0.193243 | -1100.151673 | -108.8 | -1099.891057 | -1099.955418 | 23.3 | -1094.016560 | -4.505216 | -1098.366967 |
| conformer_35 | -1100.109623 | 0.260774 | 0.193806 | -1100.149767 | -105.4 | -1099.888993 | -1099.952949 | 29.8 | -1094.015187 | -4.505772 | -1098.364285 |

[a]: Single-point calculation in aqueous phase with SMD model.

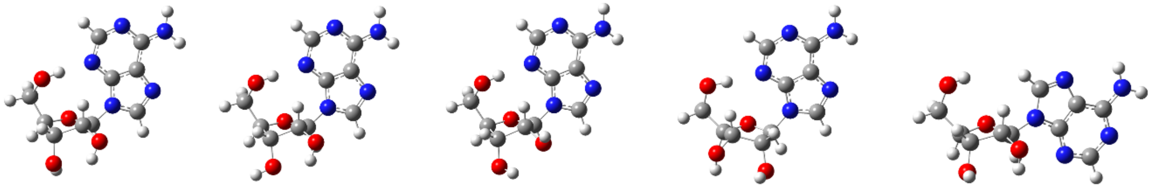

conformer_1 conformer_2 conformer_9 conformer_47 conformer_66

**Figure S46.** B3LYP-D3/def2-TZVPP optimized geometries of conformers for adenosine (**9rbA**).

**Table S44.** Conformers of gas-phase optimized adenosine (**9rbA**) at the B3LYP-D3/def2-TZVPP level of theory followed by aqueous phase single-point calculation. The columns display total energy without zero-point correction ($E_{Tot}$), Gibbs free energy (δG), total energy without zero-point correction ($E_{Tot,W}$), Gibbs free energy ($G_{298,W}$) in water (W), total single-point energy ($E_{CBS}$) calculated at DLPNO-CCSD(T)/CBS level of theory, and their corresponding free energy $G_{CBS}$. $G_{298,W}$ and $G_{CBS}$ have been corrected to the standard state of 1 mol/L by addition of +7.908 kJ/mol. $\Delta G_{Solv}$ represents the Gibbs free energy of solvation. The data are arranged in the ascending numeric order of $E_{Tot,W}$. $\Delta G_{298,W}$ represents the respective energy difference to the lowest structure. Only conformers within the 24 kJ/mol (6 kcal/mol) energy window above the lowest in CREST are included in initial conformer sampling. Duplicates of the same structure are excluded.

| | B3LYP-D3/def2-TZVPP | | | SMD(H2O)/B3LYP-D3/def2-TZVPP[a] | | | | | DLPNO-CCSD(T)/CBS | | |
|---|---|---|---|---|---|---|---|---|---|---|---|
| 9rbA No. | $E_{Tot}$ (Hartree) | δH (Hartree) | δG (Hartree) | $E_{Tot,W}$ (Hartree) | $\Delta G_{Solv}$ (kJ/mol) | $H_{298,W}$ (Hartree) | $G_{298,W}$ (Hartree) | $\Delta G_{298,W}$ (kJ/mol) | $E_{CBS,HF}$ (Hartree) | $E_{CBS,C}$ (Hartree) | $G_{CBS,W}$ (Hartree) |
| conformer_1 | -963.973397 | 0.269056 | 0.208337 | -964.012956 | -103.9 | -963.743900 | -963.801607 | 0.4 | -958.374775 | -4.133852 | -962.336836 |
| conformer_9 | -963.975529 | 0.269187 | 0.208672 | -964.012923 | -98.2 | -963.743736 | -963.801239 | 1.4 | -958.379083 | -4.132597 | -962.337390 |
| **conformer_2** | **-963.974987** | **0.269013** | **0.208045** | **-964.012821** | **-99.3** | **-963.743808** | **-963.801764** | **0.0** | **-958.378397** | **-4.132508** | **-962.337683** |
| conformer_47 | -963.969080 | 0.268992 | 0.206977 | -964.008546 | -103.6 | -963.739554 | -963.798557 | 8.4 | -958.372979 | -4.131756 | -962.334212 |
| conformer_66 | -963.963456 | 0.269045 | 0.204913 | -964.008210 | -117.5 | -963.739165 | -963.800285 | 3.9 | -958.373438 | -4.127194 | -962.337460 |
| conformer_42 | -963.965875 | 0.268796 | 0.207570 | -964.008069 | -110.8 | -963.739273 | -963.797487 | 11.2 | -958.367925 | -4.133306 | -962.332842 |
| conformer_5 | -963.971403 | 0.268927 | 0.207503 | -964.007910 | -95.8 | -963.738983 | -963.797395 | 11.5 | -958.375197 | -4.132450 | -962.333639 |
| conformer_39 | -963.967561 | 0.268747 | 0.206529 | -964.007842 | -105.8 | -963.739095 | -963.798301 | 9.1 | -958.371660 | -4.131759 | -962.334159 |
| conformer_82 | -963.961587 | 0.268833 | 0.205142 | -964.007828 | -121.4 | -963.738995 | -963.799674 | 5.5 | -958.369745 | -4.128618 | -962.336451 |
| conformer_71 | -963.964862 | 0.268921 | 0.206276 | -964.007599 | -112.2 | -963.738678 | -963.798311 | 9.1 | -958.372481 | -4.129303 | -962.335232 |
| conformer_11 | -963.966299 | 0.268966 | 0.207636 | -964.007506 | -108.2 | -963.738540 | -963.796858 | 12.9 | -958.369036 | -4.133337 | -962.332933 |
| conformer_43 | -963.965854 | 0.269070 | 0.206060 | -964.007377 | -109.0 | -963.738307 | -963.798305 | 9.1 | -958.374386 | -4.128373 | -962.335210 |
| conformer_4 | -963.973802 | 0.269023 | 0.207113 | -964.007309 | -88.0 | -963.738286 | -963.797184 | 12.0 | -958.378611 | -4.131815 | -962.333808 |
| conformer_25 | -963.966782 | 0.268897 | 0.207263 | -964.007266 | -106.3 | -963.738369 | -963.796991 | 12.5 | -958.371803 | -4.131753 | -962.333765 |
| conformer_79 | -963.960074 | 0.268671 | 0.204322 | -964.006937 | -123.0 | -963.738266 | -963.799603 | 5.7 | -958.367908 | -4.128819 | -962.336255 |
| conformer_55 | -963.962961 | 0.268992 | 0.205898 | -964.006684 | -114.8 | -963.737692 | -963.797774 | 10.5 | -958.368671 | -4.130406 | -962.333890 |
| conformer_7 | -963.971673 | 0.268943 | 0.206909 | -964.006260 | -90.8 | -963.737317 | -963.796339 | 14.2 | -958.377539 | -4.130968 | -962.333173 |
| conformer_17 | -963.970412 | 0.268831 | 0.206439 | -964.006073 | -93.6 | -963.737242 | -963.796622 | 13.5 | -958.375419 | -4.131326 | -962.332955 |
| conformer_81 | -963.958173 | 0.268700 | 0.204534 | -964.005875 | -125.2 | -963.737175 | -963.798329 | 9.0 | -958.367747 | -4.127103 | -962.335005 |
| conformer_75 | -963.965329 | 0.268906 | 0.206236 | -964.005804 | -106.3 | -963.736898 | -963.796556 | 13.7 | -958.374755 | -4.127910 | -962.333892 |
| conformer_6 | -963.970371 | 0.268924 | 0.206549 | -964.005556 | -92.4 | -963.736632 | -963.795995 | 15.1 | -958.376976 | -4.130087 | -962.332687 |
| conformer_8 | -963.969561 | 0.268813 | 0.206431 | -964.005553 | -94.5 | -963.736740 | -963.796110 | 14.8 | -958.375640 | -4.130492 | -962.332680 |
| conformer_84 | -963.961664 | 0.268916 | 0.206542 | -964.005503 | -115.1 | -963.736587 | -963.795949 | 15.3 | -958.369628 | -4.128584 | -962.332497 |
| conformer_16 | -963.968684 | 0.268800 | 0.206129 | -964.005451 | -96.5 | -963.736651 | -963.796310 | 14.3 | -958.375233 | -4.130131 | -962.332990 |
| conformer_19 | -963.965022 | 0.268996 | 0.205416 | -964.005345 | -105.9 | -963.736349 | -963.796917 | 12.7 | -958.374197 | -4.128007 | -962.334099 |
| conformer_63 | -963.965453 | 0.268880 | 0.205642 | -964.005171 | -104.3 | -963.736291 | -963.796517 | 13.8 | -958.374095 | -4.129061 | -962.334220 |
| conformer_10 | -963.971788 | 0.268870 | 0.205934 | -964.005170 | -87.6 | -963.736300 | -963.796224 | 14.5 | -958.378098 | -4.130774 | -962.333308 |
| conformer_60 | -963.960800 | 0.268742 | 0.205082 | -964.004838 | -115.6 | -963.736096 | -963.796744 | 13.2 | -958.370035 | -4.127961 | -962.333941 |
| conformer_86 | -963.965649 | 0.268847 | 0.205793 | -964.004776 | -102.7 | -963.735929 | -963.795971 | 15.2 | -958.375979 | -4.127347 | -962.333649 |
| conformer_73 | -963.968129 | 0.268731 | 0.205991 | -964.004727 | -96.1 | -963.735996 | -963.795724 | 15.9 | -958.375252 | -4.129998 | -962.332846 |

| | | | | | | | | | | | |
|---|---|---|---|---|---|---|---|---|---|---|---|
| conformer_89 | -963.965639 | 0.268820 | 0.205246 | -964.004706 | -102.6 | -963.735886 | -963.796448 | 14.0 | -958.375952 | -4.127270 | -962.334031 |
| conformer_65 | -963.959706 | 0.268652 | 0.206605 | -964.004657 | -118.0 | -963.736005 | -963.795040 | 17.7 | -958.365653 | -4.131527 | -962.332514 |
| conformer_31 | -963.969647 | 0.268874 | 0.206413 | -964.004424 | -91.3 | -963.735550 | -963.794999 | 17.8 | -958.376906 | -4.129752 | -962.332010 |
| conformer_12 | -963.966295 | 0.268913 | 0.205548 | -964.004412 | -100.1 | -963.735499 | -963.795852 | 15.5 | -958.374904 | -4.128835 | -962.333295 |
| conformer_76 | -963.961133 | 0.269011 | 0.206000 | -964.003961 | -112.4 | -963.734950 | -963.794949 | 17.9 | -958.367515 | -4.130577 | -962.331909 |
| conformer_28 | -963.964055 | 0.269098 | 0.206117 | -964.003871 | -104.5 | -963.734773 | -963.794742 | 18.4 | -958.372447 | -4.128533 | -962.331668 |
| conformer_53 | -963.962377 | 0.268675 | 0.205337 | -964.003847 | -108.9 | -963.735172 | -963.795498 | 16.5 | -958.371067 | -4.128750 | -962.332937 |
| conformer_67 | -963.963483 | 0.268822 | 0.205624 | -964.003831 | -105.9 | -963.735009 | -963.795195 | 17.2 | -958.372793 | -4.128292 | -962.332797 |
| conformer_36 | -963.963237 | 0.268786 | 0.205707 | -964.003798 | -106.5 | -963.735012 | -963.795079 | 17.6 | -958.372559 | -4.128182 | -962.332583 |
| conformer_32 | -963.962692 | 0.268877 | 0.206805 | -964.003764 | -107.8 | -963.734887 | -963.793947 | 20.5 | -958.365006 | -4.134334 | -962.330596 |
| conformer_15 | -963.961623 | 0.268854 | 0.206119 | -964.003636 | -110.3 | -963.734782 | -963.794505 | 19.1 | -958.368356 | -4.130068 | -962.331306 |
| conformer_90 | -963.955938 | 0.268796 | 0.204589 | -964.002933 | -123.4 | -963.734137 | -963.795332 | 16.9 | -958.361542 | -4.131246 | -962.332182 |
| conformer_30 | -963.964044 | 0.268842 | 0.205772 | -964.002803 | -101.8 | -963.733961 | -963.794019 | 20.3 | -958.374023 | -4.127682 | -962.331681 |
| conformer_26 | -963.960973 | 0.268754 | 0.204055 | -964.002777 | -109.8 | -963.734023 | -963.795710 | 15.9 | -958.367720 | -4.130042 | -962.332500 |
| conformer_64 | -963.960563 | 0.269015 | 0.208021 | -964.002718 | -110.7 | -963.733703 | -963.791685 | 26.5 | -958.361574 | -4.135778 | -962.328474 |
| conformer_21 | -963.964052 | 0.268916 | 0.205997 | -964.002489 | -100.9 | -963.733573 | -963.793480 | 21.7 | -958.371900 | -4.129257 | -962.330585 |
| conformer_27 | -963.961930 | 0.268833 | 0.205725 | -964.002397 | -106.2 | -963.733564 | -963.793660 | 21.3 | -958.369425 | -4.129655 | -962.330809 |
| conformer_14 | -963.961169 | 0.268819 | 0.205743 | -964.002374 | -108.2 | -963.733555 | -963.793619 | 21.4 | -958.367906 | -4.129804 | -962.330160 |
| conformer_72 | -963.960095 | 0.268957 | 0.205756 | -964.002343 | -110.9 | -963.733386 | -963.793575 | 21.5 | -958.366921 | -4.129883 | -962.330284 |
| conformer_41 | -963.963474 | 0.268846 | 0.205195 | -964.002328 | -102.0 | -963.733482 | -963.794121 | 20.1 | -958.372036 | -4.128571 | -962.331255 |
| conformer_22 | -963.966187 | 0.268874 | 0.204714 | -964.002229 | -94.6 | -963.733355 | -963.794503 | 19.1 | -958.374941 | -4.128642 | -962.331900 |
| conformer_58 | -963.958872 | 0.268904 | 0.205797 | -964.002085 | -113.5 | -963.733181 | -963.793276 | 22.3 | -958.364832 | -4.131331 | -962.330566 |
| conformer_20 | -963.961415 | 0.268879 | 0.205170 | -964.001841 | -106.1 | -963.732962 | -963.793659 | 21.3 | -958.365443 | -4.133138 | -962.330826 |
| conformer_77 | -963.960183 | 0.268932 | 0.204875 | -964.001310 | -108.0 | -963.732378 | -963.793423 | 21.9 | -958.367668 | -4.129121 | -962.330029 |
| conformer_18 | -963.966836 | 0.268910 | 0.204016 | -964.000825 | -89.2 | -963.731915 | -963.793797 | 20.9 | -958.376088 | -4.128525 | -962.331574 |
| conformer_57 | -963.960359 | 0.268785 | 0.206623 | -963.999923 | -103.9 | -963.731138 | -963.790288 | 30.1 | -958.363879 | -4.133977 | -962.327786 |
| conformer_38 | -963.958127 | 0.268820 | 0.205618 | -963.999898 | -109.7 | -963.731078 | -963.791268 | 27.6 | -958.361957 | -4.133540 | -962.328639 |

[a]: Single-point calculation in aqueous phase with SMD model.

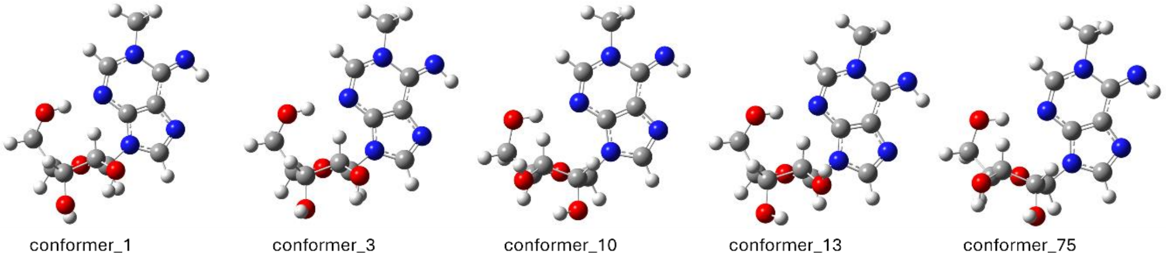

**Figure S47.** B3LYP-D3/def2-TZVPP optimized geometries of conformers for neutral $N^1$-methyladenosine (**9rb1mA**).

**Table S45.** Conformers of gas-phase optimized neutral $N^1$-methyladenosine (**9rb1mA**) at the B3LYP-D3/def2-TZVPP level of theory followed by aqueous phase single-point calculation. The columns display total energy without zero-point correction ($E_{Tot}$), Gibbs free energy (δG), total energy without zero-point correction ($E_{Tot,W}$), Gibbs free energy ($G_{298,W}$) in water (W), total single-point energy ($E_{CBS}$) calculated at DLPNO-CCSD(T)/CBS level of theory, and their corresponding free energy $G_{CBS}$. $G_{298,W}$ and $G_{CBS}$ have been corrected to the standard state of 1 mol/L by addition of +7.908 kJ/mol. $\Delta G_{Solv}$ represents the Gibbs free energy of solvation. The data are arranged in the ascending numeric order of $E_{Tot,W}$. $\Delta G_{298,W}$ represents the respective energy difference to the lowest structure. Only conformers within the 24 kJ/mol (6 kcal/mol) energy window above the lowest in CREST are included in initial conformer sampling. Duplicates of the same structure are excluded.

| 9rb1mA No. | B3LYP-D3/def2-TZVPP $E_{Tot}$ (Hartree) | δH (Hartree) | δG (Hartree) | SMD(H2O)/B3LYP-D3/def2-TZVPP[a] $E_{Tot,W}$ (Hartree) | $\Delta G_{Solv}$ (kJ/mol) | $H_{298,W}$ (Hartree) | $G_{298,W}$ (Hartree) | $\Delta G_{298,W}$ (kJ/mol) | DLPNO-CCSD(T)/CBS $E_{CBS,HF}$ (Hartree) | $E_{CBS,C}$ (Hartree) | $G_{CBS,W}$ (Hartree) |
|---|---|---|---|---|---|---|---|---|---|---|---|
| **conformer_3** | **-1003.280596** | **0.298837** | **0.235148** | **-1003.319621** | **-102.5** | **-1003.020784** | **-1003.081461** | **0.0** | **-997.396958** | **-4.346969** | **-1001.544792** |
| conformer_1 | -1003.279368 | 0.298730 | 0.235732 | -1003.319149 | -104.4 | -1003.020419 | -1003.080405 | 2.8 | -997.394232 | -4.347993 | -1001.543263 |
| conformer_13 | -1003.281526 | 0.298854 | 0.235869 | -1003.319026 | -98.5 | -1003.020172 | -1003.080145 | 3.5 | -997.398440 | -4.346864 | -1001.543923 |
| conformer_75 | -1003.275952 | 0.298800 | 0.234883 | -1003.314917 | -102.3 | -1003.016117 | -1003.077022 | 11.7 | -997.393233 | -4.345917 | -1001.540219 |
| conformer_10 | -1003.277817 | 0.298597 | 0.234782 | -1003.314293 | -95.8 | -1003.015696 | -1003.076499 | 13.0 | -997.394862 | -4.346766 | -1001.540310 |
| conformer_50 | -1003.274825 | 0.298455 | 0.234376 | -1003.314203 | -103.4 | -1003.015748 | -1003.076815 | 12.2 | -997.392288 | -4.346030 | -1001.540307 |
| conformer_91 | -1003.272666 | 0.298703 | 0.233806 | -1003.314014 | -108.6 | -1003.015311 | -1003.077196 | 11.2 | -997.394172 | -4.342804 | -1001.541507 |
| conformer_5 | -1003.279454 | 0.298743 | 0.234553 | -1003.313502 | -89.4 | -1003.014759 | -1003.075937 | 14.5 | -997.397303 | -4.346151 | -1001.539937 |
| conformer_20 | -1003.272497 | 0.298557 | 0.234612 | -1003.313396 | -107.4 | -1003.014839 | -1003.075772 | 14.9 | -997.388660 | -4.347588 | -1001.539524 |
| conformer_45 | -1003.273099 | 0.298495 | 0.234483 | -1003.313353 | -105.7 | -1003.014858 | -1003.075858 | 14.7 | -997.391552 | -4.346002 | -1001.540314 |
| conformer_16 | -1003.276972 | 0.298664 | 0.234173 | -1003.312286 | -92.7 | -1003.013622 | -1003.075101 | 16.7 | -997.396090 | -4.345280 | -1001.539500 |
| conformer_40 | -1003.271707 | 0.298646 | 0.232935 | -1003.312076 | -106.0 | -1003.013430 | -1003.076129 | 14.0 | -997.394081 | -4.342109 | -1001.540612 |
| conformer_66 | -1003.275657 | 0.298458 | 0.233793 | -1003.311838 | -95.0 | -1003.013380 | -1003.075033 | 16.9 | -997.393846 | -4.345687 | -1001.538909 |
| conformer_61 | -1003.273965 | 0.298544 | 0.233711 | -1003.311621 | -98.9 | -1003.013077 | -1003.074898 | 17.2 | -997.393661 | -4.344538 | -1001.539132 |
| conformer_12 | -1003.275851 | 0.298679 | 0.234184 | -1003.311554 | -93.7 | -1003.012875 | -1003.074358 | 18.7 | -997.395624 | -4.344429 | -1001.538559 |
| conformer_93 | -1003.271351 | 0.298569 | 0.232305 | -1003.311361 | -105.0 | -1003.012792 | -1003.076044 | 14.2 | -997.395273 | -4.340882 | -1001.540848 |
| conformer_28 | -1003.272424 | 0.298552 | 0.233152 | -1003.311085 | -101.5 | -1003.012533 | -1003.074921 | 17.2 | -997.394197 | -4.343057 | -1001.539752 |
| conformer_22 | -1003.277444 | 0.298555 | 0.233671 | -1003.311077 | -88.3 | -1003.012522 | -1003.074394 | 18.6 | -997.396820 | -4.345228 | -1001.538999 |
| conformer_99 | -1003.270851 | 0.298409 | 0.231909 | -1003.311044 | -105.5 | -1003.012635 | -1003.076123 | 14.0 | -997.393590 | -4.341996 | -1001.540858 |
| conformer_62 | -1003.270691 | 0.298747 | 0.233399 | -1003.310614 | -104.8 | -1003.011867 | -1003.074203 | 19.1 | -997.392187 | -4.342706 | -1001.538405 |
| conformer_63 | -1003.269943 | 0.298432 | 0.233019 | -1003.310535 | -106.6 | -1003.012103 | -1003.074504 | 18.3 | -997.392499 | -4.342226 | -1001.539286 |
| conformer_89 | -1003.275176 | 0.298608 | 0.233914 | -1003.310426 | -92.5 | -1003.011818 | -1003.073500 | 20.9 | -997.395596 | -4.344079 | -1001.538000 |
| conformer_34 | -1003.270005 | 0.298559 | 0.234073 | -1003.310252 | -105.7 | -1003.011693 | -1003.073167 | 21.8 | -997.385193 | -4.348759 | -1001.537114 |
| conformer_23 | -1003.268309 | 0.298530 | 0.233385 | -1003.310152 | -109.9 | -1003.011622 | -1003.073755 | 20.2 | -997.388162 | -4.344385 | -1001.537994 |
| conformer_56 | -1003.267311 | 0.298408 | 0.232525 | -1003.309847 | -111.7 | -1003.011439 | -1003.074310 | 18.8 | -997.387253 | -4.344175 | -1001.538428 |
| conformer_86 | -1003.270369 | 0.298489 | 0.233065 | -1003.309575 | -102.9 | -1003.011086 | -1003.073498 | 20.9 | -997.393637 | -4.341715 | -1001.538481 |
| conformer_48 | -1003.267111 | 0.298516 | 0.232920 | -1003.309366 | -110.9 | -1003.010850 | -1003.073434 | 21.1 | -997.387280 | -4.343813 | -1001.537415 |
| conformer_81 | -1003.270298 | 0.298529 | 0.232986 | -1003.309104 | -101.9 | -1003.010575 | -1003.073106 | 21.9 | -997.392004 | -4.342730 | -1001.537542 |
| conformer_41 | -1003.267962 | 0.298546 | 0.233526 | -1003.309102 | -108.0 | -1003.010556 | -1003.072564 | 23.4 | -997.387820 | -4.343982 | -1001.536404 |

| | | | | | | | | | | | |
|---|---|---|---|---|---|---|---|---|---|---|---|
| conformer_65 | -1003.268152 | 0.298452 | 0.232807 | -1003.308942 | -107.1 | -1003.010490 | -1003.073123 | 21.9 | -997.388871 | -4.343850 | -1001.537692 |
| conformer_33 | -1003.272835 | 0.298571 | 0.232691 | -1003.308933 | -94.8 | -1003.010362 | -1003.073230 | 21.6 | -997.394726 | -4.342948 | -1001.538068 |
| conformer_31 | -1003.268100 | 0.298557 | 0.233376 | -1003.308242 | -105.4 | -1003.009685 | -1003.071854 | 25.2 | -997.385187 | -4.347549 | -1001.536489 |
| conformer_54 | -1003.273232 | 0.298549 | 0.232621 | -1003.307869 | -90.9 | -1003.009320 | -1003.072236 | 24.2 | -997.395942 | -4.342567 | -1001.537513 |
| conformer_55 | -1003.265142 | 0.298571 | 0.233507 | -1003.306205 | -107.8 | -1003.007634 | -1003.069686 | 30.9 | -997.381719 | -4.348134 | -1001.534398 |
| conformer_32 | -1003.265842 | 0.298507 | 0.231867 | -1003.305542 | -104.2 | -1003.007035 | -1003.070663 | 28.4 | -997.383558 | -4.346801 | -1001.535180 |
| conformer_29 | -1003.265850 | 0.298562 | 0.232060 | -1003.305538 | -104.2 | -1003.006976 | -1003.070466 | 28.9 | -997.383597 | -4.346851 | -1001.535064 |

[a]: Single-point calculation in aqueous phase with SMD model.

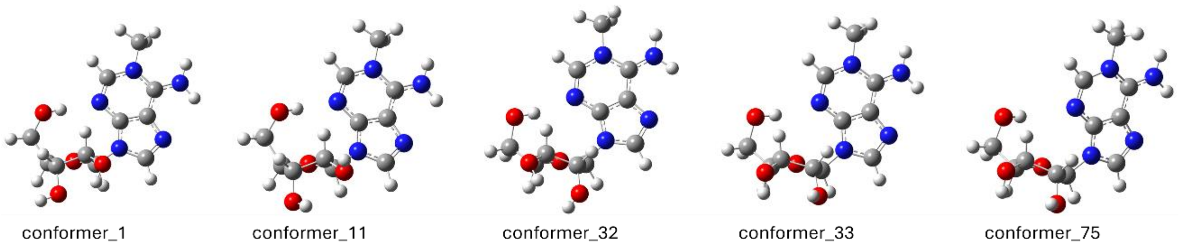

**Figure S48.** B3LYP-D3/def2-TZVPP optimized geometries of conformers for protonated *N*[1]-methyladenosine cation (**9rb1mA⁺**).

**Table S46.** Conformers of gas-phase optimized protonated *N*[1]-methyladenosine cation (**9rb1mA⁺**) at the B3LYP-D3/def2-TZVPP level of theory followed by aqueous phase single-point calculation. The columns display total energy without zero-point correction ($E_{Tot}$), Gibbs free energy (δG), total energy without zero-point correction ($E_{Tot,W}$), Gibbs free energy ($G_{298,W}$) in water (W), total single-point energy ($E_{CBS}$) calculated at DLPNO-CCSD(T)/CBS level of theory, and their corresponding free energy $G_{CBS}$. $G_{298,W}$ and $G_{CBS}$ have been corrected to the standard state of 1 mol/L by addition of +7.908 kJ/mol. $\Delta G_{Solv}$ represents the Gibbs free energy of solvation. The data are arranged in the ascending numeric order of $E_{Tot,W}$. $\Delta G_{298,W}$ represents the respective energy difference to the lowest structure. Only conformers within the 24 kJ/mol (6 kcal/mol) energy window above the lowest in CREST are included in initial conformer sampling. Duplicates of the same structure are excluded.

| 9rb1mA⁺ No. | B3LYP-D3/def2-TZVPP $E_{Tot}$ (Hartree) | δH (Hartree) | δG (Hartree) | SMD($H_2O$)/B3LYP-D3/def2-TZVPP[a] $E_{Tot,W}$ (Hartree) | $\Delta G_{Solv}$ (kJ/mol) | $H_{298,W}$ (Hartree) | $G_{298,W}$ (Hartree) | $\Delta G_{298,W}$ (kJ/mol) | DLPNO-CCSD(T)/CBS $E_{CBS,HF}$ (Hartree) | $E_{CBS,C}$ (Hartree) | $G_{CBS,W}$ (Hartree) |
|---|---|---|---|---|---|---|---|---|---|---|---|
| conformer_11 | -1003.678181 | 0.31198 | 0.247957 | -1003.789057 | -291.1 | -1003.477077 | -1003.538088 | 0.1 | -997.805187 | -4.331468 | -1001.996561 |
| **conformer_1** | **-1003.680379** | **0.311991** | **0.247877** | **-1003.789029** | **-285.3** | **-1003.477038** | **-1003.538140** | **0.0** | **-997.807155** | **-4.331489** | **-1001.996405** |
| conformer_33 | -1003.672423 | 0.311879 | 0.245769 | -1003.785382 | -296.6 | -1003.473503 | -1003.536601 | 4.0 | -997.799894 | -4.330582 | -1001.994655 |
| conformer_70 | -1003.672399 | 0.311768 | 0.245219 | -1003.784659 | -294.7 | -1003.472891 | -1003.536428 | 4.5 | -997.801787 | -4.329224 | -1001.995040 |
| conformer_32 | -1003.667636 | 0.31171 | 0.246847 | -1003.784616 | -307.1 | -1003.472906 | -1003.534757 | 8.9 | -997.792857 | -4.332287 | -1001.992264 |
| conformer_9 | -1003.67457 | 0.311705 | 0.246059 | -1003.784376 | -288.3 | -1003.472671 | -1003.535305 | 7.4 | -997.801661 | -4.331285 | -1001.993680 |
| conformer_22 | -1003.66626 | 0.311403 | 0.244988 | -1003.784317 | -310.0 | -1003.472914 | -1003.536317 | 4.8 | -997.795106 | -4.329198 | -1001.994360 |
| conformer_35 | -1003.669852 | 0.311729 | 0.245359 | -1003.784162 | -300.1 | -1003.472433 | -1003.535791 | 6.2 | -997.800958 | -4.327618 | -1001.994514 |
| conformer_45 | -1003.674366 | 0.311546 | 0.24483 | -1003.783664 | -287.0 | -1003.472118 | -1003.535822 | 6.1 | -997.803379 | -4.329796 | -1001.994630 |
| conformer_19 | -1003.673686 | 0.311646 | 0.245971 | -1003.783384 | -288.0 | -1003.471738 | -1003.534401 | 9.8 | -997.802923 | -4.329746 | -1001.993384 |
| conformer_23 | -1003.670434 | 0.311501 | 0.245113 | -1003.783254 | -296.2 | -1003.471753 | -1003.535129 | 7.9 | -997.801117 | -4.328370 | -1001.994182 |
| conformer_31 | -1003.674336 | 0.311607 | 0.245286 | -1003.782658 | -284.4 | -1003.471051 | -1003.534360 | 9.9 | -997.805844 | -4.327569 | -1001.993436 |
| conformer_12 | -1003.669700 | 0.311595 | 0.245940 | -1003.782593 | -296.4 | -1003.470998 | -1003.533641 | 11.8 | -997.798752 | -4.329606 | -1001.992299 |
| conformer_68 | -1003.672272 | 0.311876 | 0.246310 | -1003.782464 | -289.3 | -1003.470588 | -1003.533142 | 13.1 | -997.798445 | -4.332219 | -1001.991534 |
| conformer_18 | -1003.668908 | 0.311528 | 0.245105 | -1003.782441 | -298.1 | -1003.470913 | -1003.534324 | 10.0 | -997.799608 | -4.327560 | -1001.992584 |
| conformer_60 | -1003.669227 | 0.311826 | 0.245638 | -1003.782424 | -297.2 | -1003.470598 | -1003.533774 | 11.5 | -997.799628 | -4.328068 | -1001.992243 |
| conformer_21 | -1003.665958 | 0.311504 | 0.245401 | -1003.782393 | -305.7 | -1003.470889 | -1003.533980 | 10.9 | -997.794567 | -4.329136 | -1001.991725 |
| conformer_3 | -1003.677939 | 0.311605 | 0.246203 | -1003.782259 | -273.9 | -1003.470654 | -1003.533044 | 13.4 | -997.806421 | -4.330302 | -1001.991828 |
| conformer_57 | -1003.672360 | 0.311481 | 0.246306 | -1003.781864 | -287.5 | -1003.470383 | -1003.532546 | 14.7 | -997.800782 | -4.330113 | -1001.991081 |
| conformer_72 | -1003.672041 | 0.311596 | 0.245315 | -1003.781732 | -288.0 | -1003.470136 | -1003.533405 | 12.4 | -997.798365 | -4.332296 | -1001.992025 |
| conformer_14 | -1003.671340 | 0.311644 | 0.246080 | -1003.781663 | -289.7 | -1003.470019 | -1003.532571 | 14.6 | -997.800957 | -4.328973 | -1001.991161 |
| conformer_80 | -1003.674531 | 0.311602 | 0.245012 | -1003.781565 | -281.0 | -1003.469963 | -1003.533541 | 12.1 | -997.807040 | -4.326883 | -1001.992935 |
| conformer_36 | -1003.664800 | 0.311655 | 0.245628 | -1003.781234 | -305.7 | -1003.469579 | -1003.532594 | 14.6 | -997.793471 | -4.329294 | -1001.990560 |
| conformer_100 | -1003.663579 | 0.311499 | 0.245154 | -1003.780677 | -307.4 | -1003.469178 | -1003.532511 | 14.8 | -997.787498 | -4.334576 | -1001.991006 |
| conformer_51 | -1003.674376 | 0.311599 | 0.244980 | -1003.780402 | -278.4 | -1003.468803 | -1003.532410 | 15.0 | -997.805284 | -4.328437 | -1001.991755 |
| conformer_63 | -1003.670929 | 0.311557 | 0.245759 | -1003.780384 | -287.4 | -1003.468827 | -1003.531613 | 17.1 | -997.801155 | -4.328638 | -1001.990476 |
| conformer_28 | -1003.672432 | 0.311785 | 0.246918 | -1003.779734 | -281.7 | -1003.467949 | -1003.529804 | 21.9 | -997.797335 | -4.333715 | -1001.988421 |
| conformer_85 | -1003.674856 | 0.311633 | 0.245430 | -1003.779679 | -275.2 | -1003.468046 | -1003.531237 | 18.1 | -997.806591 | -4.327998 | -1001.990969 |

[a]: Single-point calculation in aqueous phase with SMD model.

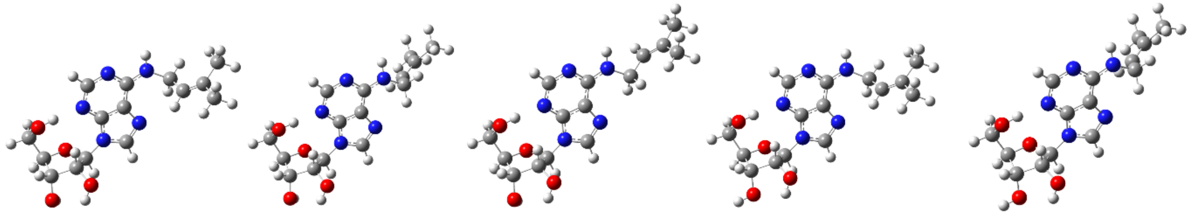

conformer_26 conformer_27 conformer_41 conformer_45 conformer_54

**Figure S49.** B3LYP-D3/def2-TZVPP optimized geometries of conformers for $N^6$-isoprenyladenosine (**9rb6iPA**).

**Table S47.** Conformers of gas-phase optimized $N^6$-isoprenyladenosine (**9rb6iPA**) at the B3LYP-D3/def2-TZVPP level of theory followed by aqueous phase single-point calculation. The columns display total energy without zero-point correction ($E_{Tot}$), Gibbs free energy (δG), total energy without zero-point correction ($E_{Tot,W}$), Gibbs free energy ($G_{298,W}$) in water (W), total single-point energy ($E_{CBS}$) calculated at DLPNO-CCSD(T)/CBS level of theory, and their corresponding free energy $G_{CBS}$. $G_{298,W}$ and $G_{CBS}$ have been corrected to the standard state of 1 mol/L by addition of +7.908 kJ/mol. $\Delta G_{Solv}$ represents the Gibbs free energy of solvation. The data are arranged in the ascending numeric order of $E_{Tot,W}$. $\Delta G_{298,W}$ represents the respective energy difference to the lowest structure. Only conformers within the 24 kJ/mol (6 kcal/mol) energy window above the lowest in CREST are included in initial conformer sampling. Duplicates of the same structure are excluded.

| 9rb6iPA No. | B3LYP-D3/def2-TZVPP $E_{Tot}$ (Hartree) | δH (Hartree) | δG (Hartree) | SMD(H2O)/B3LYP-D3/def2-TZVPP[a] $E_{Tot,W}$ (Hartree) | $\Delta G_{Solv}$ (kJ/mol) | $H_{298,W}$ (Hartree) | $G_{298,W}$ (Hartree) | $\Delta G_{298,W}$ (kJ/mol) | DLPNO-CCSD(T)/CBS $E_{CBS,HF}$ (Hartree) | $E_{CBS,C}$ (Hartree) | $G_{CBS,W}$ (Hartree) |
|---|---|---|---|---|---|---|---|---|---|---|---|
| conformer_27 | -1159.393204 | 0.393385 | 0.316412 | -1159.428896 | -93.7 | -1159.035511 | -1159.109472 | 3.2 | -1152.430254 | -5.133456 | -1157.279977 |
| conformer_26 | -1159.393273 | 0.393367 | 0.315985 | -1159.428820 | -93.3 | -1159.035453 | -1159.109823 | 2.3 | -1152.430356 | -5.133478 | -1157.280384 |
| conformer_45 | -1159.394815 | 0.393318 | 0.315781 | -1159.428741 | -89.1 | -1159.035423 | -1159.109948 | 2.0 | -1152.433905 | -5.132110 | -1157.281148 |
| conformer_54 | -1159.394779 | 0.393350 | 0.316227 | -1159.428730 | -89.1 | -1159.035380 | -1159.109491 | 3.2 | -1152.433839 | -5.132111 | -1157.280662 |
| **conformer_41** | **-1159.393048** | **0.393246** | **0.314900** | **-1159.428604** | **-93.4** | **-1159.035358** | **-1159.110692** | **0.0** | **-1152.431364** | **-5.132154** | **-1157.281163** |
| conformer_55 | -1159.394597 | 0.393225 | 0.314913 | -1159.428464 | -88.9 | -1159.035239 | -1159.110539 | 0.4 | -1152.434917 | -5.130805 | -1157.281665 |
| conformer_36 | -1159.393061 | 0.393360 | 0.315476 | -1159.428217 | -92.3 | -1159.034857 | -1159.109729 | 2.5 | -1152.430960 | -5.132339 | -1157.279967 |
| conformer_20 | -1159.394745 | 0.393450 | 0.315419 | -1159.427954 | -87.2 | -1159.034504 | -1159.109523 | 3.1 | -1152.431957 | -5.133783 | -1157.280518 |
| conformer_1 | -1159.393793 | 0.393367 | 0.316283 | -1159.427797 | -89.3 | -1159.034430 | -1159.108502 | 5.7 | -1152.429452 | -5.134811 | -1157.278973 |
| conformer_5 | -1159.395387 | 0.393398 | 0.316471 | -1159.427726 | -84.9 | -1159.034328 | -1159.108243 | 6.4 | -1152.433035 | -5.133499 | -1157.279389 |
| conformer_3 | -1159.393635 | 0.393404 | 0.316910 | -1159.427593 | -89.2 | -1159.034189 | -1159.107671 | 7.9 | -1152.429359 | -5.134823 | -1157.278217 |
| conformer_7 | -1159.395238 | 0.393393 | 0.316914 | -1159.427515 | -84.7 | -1159.034122 | -1159.107589 | 8.1 | -1152.432970 | -5.133467 | -1157.278788 |
| conformer_100 | -1159.393646 | 0.393342 | 0.315618 | -1159.425334 | -83.2 | -1159.031992 | -1159.106704 | 10.5 | -1152.433244 | -5.132190 | -1157.278493 |
| conformer_79 | -1159.391733 | 0.393355 | 0.316152 | -1159.422854 | -81.7 | -1159.029499 | -1159.103690 | 18.4 | -1152.429824 | -5.133500 | -1157.275280 |
| conformer_76 | -1159.391798 | 0.393329 | 0.316329 | -1159.422792 | -81.4 | -1159.029463 | -1159.103451 | 19.0 | -1152.429898 | -5.133430 | -1157.274982 |
| conformer_12 | -1159.394390 | 0.393342 | 0.315665 | -1159.422521 | -73.9 | -1159.029179 | -1159.103844 | 18.0 | -1152.433393 | -5.132797 | -1157.275644 |
| conformer_14 | -1159.394427 | 0.393396 | 0.315823 | -1159.422482 | -73.7 | -1159.029086 | -1159.103647 | 18.5 | -1152.433480 | -5.132811 | -1157.275512 |
| conformer_88 | -1159.392367 | 0.393289 | 0.315584 | -1159.421291 | -75.9 | -1159.028002 | -1159.102695 | 21.0 | -1152.432426 | -5.132056 | -1157.274809 |
| conformer_82 | -1159.392333 | 0.393271 | 0.315637 | -1159.421263 | -76.0 | -1159.027992 | -1159.102614 | 21.2 | -1152.432540 | -5.131879 | -1157.274700 |
| conformer_66 | -1159.390968 | 0.393189 | 0.314994 | -1159.420541 | -77.6 | -1159.027352 | -1159.102535 | 21.4 | -1152.431966 | -5.130887 | -1157.274420 |
| conformer_89 | -1159.390258 | 0.393167 | 0.315065 | -1159.420514 | -79.4 | -1159.027347 | -1159.102437 | 21.7 | -1152.430686 | -5.131297 | -1157.274162 |
| conformer_86 | -1159.390228 | 0.393124 | 0.315186 | -1159.420509 | -79.5 | -1159.027385 | -1159.102311 | 22.0 | -1152.430673 | -5.131424 | -1157.274180 |
| conformer_52 | -1159.391002 | 0.393261 | 0.315121 | -1159.420502 | -77.5 | -1159.027241 | -1159.102369 | 21.9 | -1152.431970 | -5.130943 | -1157.274280 |

[a]: Single-point calculation in aqueous phase with SMD model.

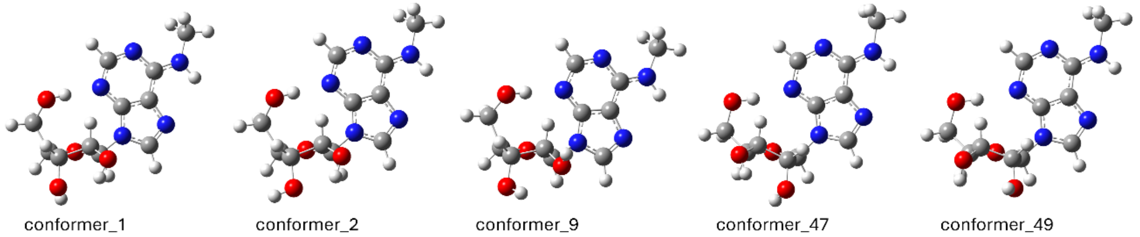

**Figure S50.** B3LYP-D3/def2-TZVPP optimized geometries of conformers for *N*6-methyladenosine (**9rb6mA**).

**Table S48.** Conformers of gas-phase optimized *N*6-methyladenosine (**9rb6mA**) at the B3LYP-D3/def2-TZVPP level of theory followed by aqueous phase single-point calculation. The columns display total energy without zero-point correction ($E_{Tot}$), Gibbs free energy ($\delta G$), total energy without zero-point correction ($E_{Tot,W}$), Gibbs free energy ($G_{298,W}$) in water (W), total single-point energy ($E_{CBS}$) calculated at DLPNO-CCSD(T)/CBS level of theory, and their corresponding free energy $G_{CBS}$. $G_{298,W}$ and $G_{CBS}$ have been corrected to the standard state of 1 mol/L by addition of +7.908 kJ/mol. $\Delta G_{Solv}$ represents the Gibbs free energy of solvation. The data are arranged in the ascending numeric order of $E_{Tot,W}$. $\Delta G_{298,W}$ represents the respective energy difference to the lowest structure. Only conformers within the 24 kJ/mol (6 kcal/mol) energy window above the lowest in CREST are included in initial conformer sampling. Duplicates of the same structure are excluded.

| 9rb6mA No. | B3LYP-D3/def2-TZVPP $E_{Tot}$ (Hartree) | $\delta H$ (Hartree) | $\delta G$ (Hartree) | SMD($H_2O$)/B3LYP-D3/def2-TZVPP[a] $E_{Tot,W}$ (Hartree) | $\Delta G_{Solv}$ (kJ/mol) | $H_{298,W}$ (Hartree) | $G_{298,W}$ (Hartree) | $\Delta G_{298,W}$ (kJ/mol) | DLPNO-CCSD(T)/CBS $E_{CBS,HF}$ (Hartree) | $E_{CBS,C}$ (Hartree) | $G_{CBS,W}$ (Hartree) |
|---|---|---|---|---|---|---|---|---|---|---|---|
| conformer_9 | -1003.299009 | 0.298998 | 0.234741 | -1003.334234 | -92.5 | -1003.035236 | -1003.096481 | 0.5 | -997.421106 | -4.339985 | -1001.558564 |
| conformer_1 | -1003.296919 | 0.298895 | 0.234601 | -1003.334227 | -98.0 | -1003.035332 | -1003.096614 | 0.2 | -997.416828 | -4.341272 | -1001.557795 |
| **conformer_2** | **-1003.298459** | **0.298857** | **0.234414** | **-1003.334104** | **-93.6** | **-1003.035247** | **-1003.096678** | **0.0** | **-997.420406** | **-4.339910** | **-1001.558534** |
| conformer_49 | -1003.292589 | 0.298872 | 0.233331 | -1003.329928 | -98.0 | -1003.031056 | -1003.093585 | 8.1 | -997.414999 | -4.339160 | -1001.555155 |
| conformer_47 | -1003.289417 | 0.298615 | 0.233494 | -1003.329376 | -104.9 | -1003.030761 | -1003.092870 | 10.0 | -997.410033 | -4.340722 | -1001.554209 |
| conformer_5 | -1003.294887 | 0.298788 | 0.233535 | -1003.329324 | -90.4 | -1003.030536 | -1003.092777 | 10.2 | -997.417250 | -4.339786 | -1001.554925 |
| conformer_73 | -1003.286750 | 0.298863 | 0.230886 | -1003.329255 | -111.6 | -1003.030392 | -1003.095357 | 3.5 | -997.415336 | -4.334409 | -1001.558353 |
| conformer_40 | -1003.291067 | 0.298574 | 0.232647 | -1003.329222 | -100.2 | -1003.030648 | -1003.093563 | 8.2 | -997.413739 | -4.339128 | -1001.555363 |
| conformer_87 | -1003.284879 | 0.298601 | 0.230733 | -1003.328931 | -115.7 | -1003.030330 | -1003.095186 | 3.9 | -997.411615 | -4.335880 | -1001.557801 |
| conformer_76 | -1003.288175 | 0.298761 | 0.232106 | -1003.328840 | -106.8 | -1003.030079 | -1003.093722 | 7.8 | -997.414331 | -4.336535 | -1001.556413 |
| conformer_12 | -1003.289787 | 0.298785 | 0.233453 | -1003.328671 | -102.1 | -1003.029886 | -1003.092206 | 11.7 | -997.411101 | -4.340759 | -1001.554279 |
| conformer_3 | -1003.297314 | 0.298854 | 0.233098 | -1003.328492 | -81.9 | -1003.029638 | -1003.092382 | 11.3 | -997.420696 | -4.339209 | -1001.554973 |
| conformer_26 | -1003.290234 | 0.298725 | 0.233230 | -1003.328473 | -100.4 | -1003.029748 | -1003.092231 | 11.7 | -997.413822 | -4.339140 | -1001.554960 |
| conformer_50 | -1003.289151 | 0.298821 | 0.232018 | -1003.328415 | -103.1 | -1003.029594 | -1003.093385 | 8.6 | -997.416310 | -4.335705 | -1001.556249 |
| conformer_83 | -1003.283433 | 0.298554 | 0.230775 | -1003.328092 | -117.3 | -1003.029538 | -1003.094305 | 6.2 | -997.409717 | -4.336232 | -1001.556821 |
| conformer_62 | -1003.286329 | 0.298839 | 0.232502 | -1003.327919 | -109.2 | -1003.029080 | -1003.092405 | 11.2 | -997.410630 | -4.337760 | -1001.554465 |
| conformer_7 | -1003.295206 | 0.298763 | 0.232949 | -1003.327392 | -84.5 | -1003.028629 | -1003.091431 | 13.8 | -997.419634 | -4.338371 | -1001.554230 |
| conformer_18 | -1003.293839 | 0.298685 | 0.232681 | -1003.327325 | -87.9 | -1003.028640 | -1003.091632 | 13.2 | -997.417408 | -4.338697 | -1001.553898 |
| conformer_82 | -1003.288574 | 0.298702 | 0.231567 | -1003.326974 | -100.8 | -1003.028272 | -1003.092395 | 11.2 | -997.416662 | -4.335036 | -1001.555519 |
| conformer_85 | -1003.281429 | 0.298483 | 0.230125 | -1003.326909 | -119.4 | -1003.028426 | -1003.093772 | 7.6 | -997.409641 | -4.334299 | -1001.556283 |
| conformer_6 | -1003.293844 | 0.298759 | 0.232657 | -1003.326751 | -86.4 | -1003.027992 | -1003.091082 | 14.7 | -997.419019 | -4.337476 | -1001.553732 |
| conformer_8 | -1003.293056 | 0.298637 | 0.232482 | -1003.326706 | -88.3 | -1003.028069 | -1003.091212 | 14.3 | -997.417688 | -4.337899 | -1001.553743 |
| conformer_16 | -1003.292146 | 0.298624 | 0.232283 | -1003.326649 | -90.6 | -1003.028025 | -1003.091354 | 14.0 | -997.417230 | -4.337565 | -1001.554004 |
| conformer_19 | -1003.288430 | 0.298859 | 0.231404 | -1003.326531 | -100.0 | -1003.027672 | -1003.092115 | 12.0 | -997.416221 | -4.335214 | -1001.555119 |
| conformer_11 | -1003.295172 | 0.298720 | 0.232207 | -1003.326370 | -81.9 | -1003.027650 | -1003.091151 | 14.5 | -997.420004 | -4.338244 | -1001.554227 |
| conformer_10 | -1003.295172 | 0.298720 | 0.232207 | -1003.326370 | -81.9 | -1003.027650 | -1003.091151 | 14.5 | -997.420004 | -4.338244 | -1001.554227 |
| conformer_66 | -1003.284178 | 0.298577 | 0.230889 | -1003.325981 | -109.8 | -1003.027404 | -1003.092080 | 12.1 | -997.411982 | -4.335180 | -1001.555064 |
| conformer_80 | -1003.291572 | 0.298550 | 0.232057 | -1003.325917 | -90.2 | -1003.027367 | -1003.090848 | 15.3 | -997.417226 | -4.337402 | -1001.553904 |
| conformer_74 | -1003.291572 | 0.298550 | 0.232057 | -1003.325917 | -90.2 | -1003.027367 | -1003.090848 | 15.3 | -997.417226 | -4.337402 | -1001.553904 |

| | | | | | | | | | | | |
|---|---|---|---|---|---|---|---|---|---|---|---|
| conformer_32 | -1003.293100 | 0.298707 | 0.232427 | -1003.325624 | -85.4 | -1003.026917 | -1003.090185 | 17.0 | -997.418921 | -4.337179 | -1001.553185 |
| conformer_33 | -1003.289684 | 0.298737 | 0.231542 | -1003.325439 | -93.9 | -1003.026702 | -1003.090885 | 15.2 | -997.416911 | -4.336143 | -1001.554255 |
| conformer_13 | -1003.289684 | 0.298737 | 0.231542 | -1003.325439 | -93.9 | -1003.026702 | -1003.090885 | 15.2 | -997.416911 | -4.336144 | -1001.554255 |
| conformer_30 | -1003.287445 | 0.298887 | 0.231866 | -1003.325096 | -98.9 | -1003.026209 | -1003.090218 | 17.0 | -997.414383 | -4.335800 | -1001.552956 |
| conformer_71 | -1003.286800 | 0.298619 | 0.231378 | -1003.325028 | -100.4 | -1003.026409 | -1003.090638 | 15.9 | -997.414763 | -4.335507 | -1001.554109 |
| conformer_35 | -1003.286108 | 0.298695 | 0.232490 | -1003.324982 | -102.1 | -1003.026287 | -1003.089480 | 18.9 | -997.407028 | -4.341512 | -1001.551912 |
| conformer_41 | -1003.286523 | 0.298554 | 0.231354 | -1003.324957 | -100.9 | -1003.026403 | -1003.090591 | 16.0 | -997.414513 | -4.335359 | -1001.553940 |
| conformer_59 | -1003.285741 | 0.298473 | 0.230997 | -1003.324892 | -102.8 | -1003.026419 | -1003.090883 | 15.2 | -997.413049 | -4.335975 | -1001.554167 |
| conformer_84 | -1003.284537 | 0.298770 | 0.231953 | -1003.324854 | -105.9 | -1003.026084 | -1003.089889 | 17.8 | -997.409421 | -4.337988 | -1001.552762 |
| conformer_78 | -1003.284537 | 0.298770 | 0.231953 | -1003.324854 | -105.9 | -1003.026084 | -1003.089889 | 17.8 | -997.409421 | -4.337988 | -1001.552762 |
| conformer_17 | -1003.284933 | 0.298615 | 0.231708 | -1003.324789 | -104.6 | -1003.026174 | -1003.090069 | 17.3 | -997.410290 | -4.337364 | -1001.552791 |
| conformer_60 | -1003.281874 | 0.298690 | 0.232013 | -1003.324137 | -111.0 | -1003.025447 | -1003.089112 | 19.9 | -997.406178 | -4.338389 | -1001.551805 |
| conformer_34 | -1003.287327 | 0.298620 | 0.231429 | -1003.323992 | -96.3 | -1003.025372 | -1003.089551 | 18.7 | -997.415976 | -4.334883 | -1001.553083 |
| conformer_69 | -1003.284100 | 0.298859 | 0.233855 | -1003.323880 | -104.4 | -1003.025021 | -1003.087013 | 25.4 | -997.403666 | -4.343220 | -1001.549799 |
| conformer_24 | -1003.284311 | 0.298534 | 0.230256 | -1003.323875 | -103.9 | -1003.025341 | -1003.090607 | 15.9 | -997.409629 | -4.337439 | -1001.553365 |
| conformer_22 | -1003.287447 | 0.298726 | 0.231903 | -1003.323731 | -95.3 | -1003.025005 | -1003.088816 | 20.6 | -997.413815 | -4.336626 | -1001.551810 |
| conformer_45 | -1003.286858 | 0.298632 | 0.230850 | -1003.323596 | -96.5 | -1003.024964 | -1003.089734 | 18.2 | -997.414020 | -4.335833 | -1001.552729 |
| conformer_46 | -1003.286858 | 0.298632 | 0.230850 | -1003.323596 | -96.5 | -1003.024964 | -1003.089734 | 18.2 | -997.414020 | -4.335833 | -1001.552729 |
| conformer_31 | -1003.285241 | 0.298589 | 0.231292 | -1003.323509 | -100.5 | -1003.024920 | -1003.089205 | 19.6 | -997.411342 | -4.336972 | -1001.552278 |
| conformer_27 | -1003.289558 | 0.298715 | 0.231484 | -1003.323485 | -89.1 | -1003.024770 | -1003.088989 | 20.2 | -997.416889 | -4.335954 | -1001.552274 |
| conformer_23 | -1003.289558 | 0.298715 | 0.231484 | -1003.323485 | -89.1 | -1003.024770 | -1003.088989 | 20.2 | -997.416889 | -4.335954 | -1001.552274 |
| conformer_75 | -1003.283438 | 0.298778 | 0.232063 | -1003.323351 | -104.8 | -1003.024573 | -1003.088276 | 22.1 | -997.409026 | -4.337061 | -1001.550925 |
| conformer_68 | -1003.282236 | 0.298728 | 0.231785 | -1003.323209 | -107.6 | -1003.024481 | -1003.088412 | 21.7 | -997.406832 | -4.338543 | -1001.551552 |
| conformer_63 | -1003.282236 | 0.298728 | 0.231785 | -1003.323209 | -107.6 | -1003.024481 | -1003.088412 | 21.7 | -997.406832 | -4.338543 | -1001.551552 |
| conformer_20 | -1003.284890 | 0.298711 | 0.231169 | -1003.323005 | -100.1 | -1003.024294 | -1003.088824 | 20.6 | -997.407389 | -4.340598 | -1001.551921 |
| conformer_79 | -1003.283548 | 0.298718 | 0.230836 | -1003.322354 | -101.9 | -1003.023636 | -1003.088506 | 21.5 | -997.409502 | -4.336534 | -1001.550994 |
| conformer_21 | -1003.290207 | 0.298751 | 0.231085 | -1003.321983 | -83.4 | -1003.023232 | -1003.087886 | 23.1 | -997.417963 | -4.335870 | -1001.551511 |
| conformer_44 | -1003.290207 | 0.298751 | 0.231085 | -1003.321983 | -83.4 | -1003.023232 | -1003.087886 | 23.1 | -997.417963 | -4.335870 | -1001.551511 |
| conformer_65 | -1003.290207 | 0.298751 | 0.231085 | -1003.321983 | -83.4 | -1003.023232 | -1003.087886 | 23.1 | -997.417962 | -4.335870 | -1001.551511 |
| conformer_15 | -1003.282730 | 0.298588 | 0.228227 | -1003.320263 | -98.5 | -1003.021675 | -1003.089024 | 20.1 | -997.406356 | -4.339428 | -1001.552078 |

[a]: Single-point calculation in aqueous phase with SMD model.

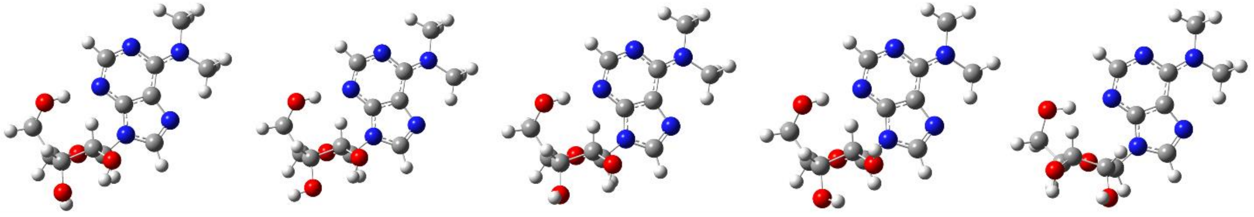

**Figure S51.** B3LYP-D3/def2-TZVPP optimized geometries of conformers for *N*[6],*N*[6]-dimethyladenosine (**9rb6dmA**).

**Table S49.** Conformers of gas-phase optimized *N*[6],*N*[6]-dimethyladenosine (**9rb6dmA**) at the B3LYP-D3/def2-TZVPP level of theory followed by aqueous phase single-point calculation. The columns display total energy without zero-point correction ($E_{Tot}$), Gibbs free energy (δG), total energy without zero-point correction ($E_{Tot,W}$), Gibbs free energy ($G_{298,W}$) in water (W), total single-point energy ($E_{CBS}$) calculated at DLPNO-CCSD(T)/CBS level of theory, and their corresponding free energy $G_{CBS}$. $G_{298,W}$ and $G_{CBS}$ have been corrected to the standard state of 1 mol/L by addition of +7.908 kJ/mol. $\Delta G_{Solv}$ represents the Gibbs free energy of solvation. The data are arranged in the ascending numeric order of $E_{Tot,W}$. $\Delta G_{298,W}$ represents the respective energy difference to the lowest structure. Only conformers within the 24 kJ/mol (6 kcal/mol) energy window above the lowest in CREST are included in initial conformer sampling. Duplicates of the same structure are excluded.

| 9rb6dmA | B3LYP-D3/def2-TZVPP | | | SMD($H_2O$)/B3LYP-D3/def2-TZVPP[a] | | | | | DLPNO-CCSD(T)/CBS | | |
|---|---|---|---|---|---|---|---|---|---|---|---|
| No. | $E_{Tot}$ (Hartree) | δH (Hartree) | δG (Hartree) | $E_{Tot,W}$ (Hartree) | $\Delta G_{Solv}$ (kJ/mol) | $H_{298,W}$ (Hartree) | $G_{298,W}$ (Hartree) | $\Delta G_{298,W}$ (kJ/mol) | $E_{CBS,HF}$ (Hartree) | $E_{CBS,C}$ (Hartree) | $G_{CBS,W}$ (Hartree) |
| **conformer_4** | **-1042.617869** | **0.328098** | **0.259116** | **-1042.648194** | **-79.6** | **-1042.320096** | **-1042.386066** | **0.0** | **-1036.453239** | **-4.552960** | **-1040.774396** |
| conformer_10 | -1042.618933 | 0.328163 | 0.259780 | -1042.647901 | -76.1 | -1042.319738 | -1042.385109 | 2.5 | -1036.455027 | -4.552667 | -1040.773870 |
| conformer_1 | -1042.616819 | 0.328027 | 0.259911 | -1042.647871 | -81.5 | -1042.319844 | -1042.384948 | 2.9 | -1036.450646 | -4.554066 | -1040.772842 |
| conformer_2 | -1042.618366 | 0.327993 | 0.259683 | -1042.647717 | -77.1 | -1042.319724 | -1042.385022 | 2.7 | -1036.454222 | -4.552666 | -1040.773544 |
| conformer_47 | -1042.612633 | 0.328067 | 0.258515 | -1042.643616 | -81.3 | -1042.315549 | -1042.382089 | 10.4 | -1036.449070 | -4.551881 | -1040.770406 |
| conformer_88 | -1042.606771 | 0.328016 | 0.256171 | -1042.643014 | -95.2 | -1042.314998 | -1042.383831 | 5.9 | -1036.449399 | -4.547165 | -1040.773624 |
| conformer_60 | -1042.609293 | 0.327801 | 0.258777 | -1042.642898 | -88.2 | -1042.315097 | -1042.381109 | 13.0 | -1036.443855 | -4.553482 | -1040.769153 |
| conformer_7 | -1042.614851 | 0.327986 | 0.258342 | -1042.642824 | -73.4 | -1042.314838 | -1042.381470 | 12.1 | -1036.451081 | -4.552613 | -1040.770314 |
| conformer_104 | -1042.604942 | 0.327760 | 0.255880 | -1042.642739 | -99.2 | -1042.314979 | -1042.383847 | 5.8 | -1036.445768 | -4.548535 | -1040.773207 |
| conformer_3 | -1042.617538 | 0.328019 | 0.258321 | -1042.642473 | -65.5 | -1042.314454 | -1042.381140 | 12.9 | -1036.454723 | -4.552057 | -1040.770381 |
| conformer_94 | -1042.608148 | 0.327869 | 0.257438 | -1042.642339 | -89.8 | -1042.314470 | -1042.381889 | 11.0 | -1036.448311 | -4.549277 | -1040.771330 |
| conformer_14 | -1042.609594 | 0.328001 | 0.259419 | -1042.642073 | -85.3 | -1042.314072 | -1042.379642 | 16.9 | -1036.444930 | -4.553451 | -1040.768429 |
| conformer_55 | -1042.609346 | 0.328037 | 0.257738 | -1042.641971 | -85.7 | -1042.313934 | -1042.381221 | 12.7 | -1036.450522 | -4.548375 | -1040.770772 |
| conformer_32 | -1042.610062 | 0.327942 | 0.259154 | -1042.641737 | -83.2 | -1042.313795 | -1042.379571 | 17.1 | -1036.447624 | -4.551889 | -1040.769022 |
| conformer_98 | -1042.603516 | 0.327797 | 0.256239 | -1042.641496 | -99.7 | -1042.313699 | -1042.382245 | 10.0 | -1036.443740 | -4.548912 | -1040.771380 |
| conformer_71 | -1042.606396 | 0.327956 | 0.257773 | -1042.641482 | -92.1 | -1042.313526 | -1042.380697 | 14.1 | -1036.444748 | -4.550447 | -1040.769497 |
| conformer_9 | -1042.615438 | 0.327914 | 0.258456 | -1042.641172 | -67.6 | -1042.313258 | -1042.379704 | 16.7 | -1036.453733 | -4.551165 | -1040.769164 |
| conformer_17 | -1042.614228 | 0.327897 | 0.258265 | -1042.641162 | -70.7 | -1042.313265 | -1042.379885 | 16.2 | -1036.451638 | -4.551509 | -1040.768804 |
| conformer_93 | -1042.608915 | 0.327862 | 0.257385 | -1042.640718 | -83.5 | -1042.312856 | -1042.380321 | 15.1 | -1036.450991 | -4.547856 | -1040.770252 |
| conformer_15 | -1042.612539 | 0.327817 | 0.258103 | -1042.640668 | -73.9 | -1042.312851 | -1042.379553 | 17.1 | -1036.451500 | -4.550362 | -1040.768875 |
| conformer_103 | -1042.601450 | 0.327626 | 0.255198 | -1042.640595 | -102.8 | -1042.312969 | -1042.382385 | 9.7 | -1036.443593 | -4.547078 | -1040.771606 |
| conformer_6 | -1042.614111 | 0.327938 | 0.258061 | -1042.640556 | -69.4 | -1042.312618 | -1042.379483 | 17.3 | -1036.453148 | -4.550297 | -1040.768817 |
| conformer_8 | -1042.613342 | 0.327798 | 0.257765 | -1042.640506 | -71.3 | -1042.312708 | -1042.379729 | 16.6 | -1036.451836 | -4.550683 | -1040.768905 |
| conformer_11 | -1042.615540 | 0.327950 | 0.257222 | -1042.640339 | -65.1 | -1042.312389 | -1042.380105 | 15.7 | -1036.454161 | -4.551036 | -1040.769763 |
| conformer_70 | -1042.608400 | 0.328045 | 0.257317 | -1042.640068 | -83.1 | -1042.312023 | -1042.379739 | 16.6 | -1036.450278 | -4.547912 | -1040.769529 |
| conformer_80 | -1042.611913 | 0.327727 | 0.257877 | -1042.639948 | -73.6 | -1042.312221 | -1042.379059 | 18.4 | -1036.451437 | -4.550220 | -1040.768803 |
| conformer_22 | -1042.608398 | 0.328046 | 0.257110 | -1042.639819 | -82.5 | -1042.311773 | -1042.379697 | 16.7 | -1036.450141 | -4.547973 | -1040.769413 |
| conformer_84 | -1042.608822 | 0.327928 | 0.257042 | -1042.639656 | -81.0 | -1042.311728 | -1042.379602 | 17.0 | -1036.449888 | -4.549185 | -1040.769853 |
| conformer_105 | -1042.609209 | 0.327831 | 0.257179 | -1042.639572 | -79.7 | -1042.311741 | -1042.379381 | 17.6 | -1036.452211 | -4.547267 | -1040.769650 |

| | | | | | | | | | | | |
|---|---|---|---|---|---|---|---|---|---|---|---|
| conformer_28 | -1042.613366 | 0.327889 | 0.257835 | -1042.639422 | -68.4 | -1042.311533 | -1042.378575 | 19.7 | -1036.453029 | -4.549955 | -1040.768193 |
| conformer_79 | -1042.604134 | 0.327758 | 0.256159 | -1042.639332 | -92.4 | -1042.311574 | -1042.380161 | 15.5 | -1036.445947 | -4.547866 | -1040.769840 |
| conformer_16 | -1042.609603 | 0.327907 | 0.256813 | -1042.639159 | -77.6 | -1042.311252 | -1042.379334 | 17.7 | -1036.450939 | -4.548761 | -1040.769430 |
| conformer_38 | -1042.609608 | 0.327925 | 0.256729 | -1042.638925 | -77.0 | -1042.311000 | -1042.379184 | 18.1 | -1036.450733 | -4.548876 | -1040.769185 |
| conformer_31 | -1042.607460 | 0.328096 | 0.257145 | -1042.638515 | -81.5 | -1042.310419 | -1042.378358 | 20.2 | -1036.448339 | -4.548528 | -1040.767765 |
| conformer_50 | -1042.606415 | 0.327781 | 0.257078 | -1042.638502 | -84.2 | -1042.310721 | -1042.378412 | 20.1 | -1036.448247 | -4.548172 | -1040.768417 |
| conformer_92 | -1042.604591 | 0.327943 | 0.257830 | -1042.638473 | -89.0 | -1042.310530 | -1042.377631 | 22.1 | -1036.443447 | -4.550654 | -1040.767141 |
| conformer_21 | -1042.604833 | 0.327879 | 0.257445 | -1042.638380 | -88.1 | -1042.310501 | -1042.377923 | 21.4 | -1036.444109 | -4.550061 | -1040.767261 |
| conformer_43 | -1042.605808 | 0.327857 | 0.258071 | -1042.638250 | -85.2 | -1042.310393 | -1042.377167 | 23.4 | -1036.440418 | -4.554466 | -1040.766243 |
| conformer_30 | -1042.604194 | 0.327793 | 0.255549 | -1042.637553 | -87.6 | -1042.309760 | -1042.378992 | 18.6 | -1036.443456 | -4.550032 | -1040.768286 |
| conformer_40 | -1042.607191 | 0.327840 | 0.257004 | -1042.637493 | -79.6 | -1042.309653 | -1042.377477 | 22.6 | -1036.449674 | -4.547680 | -1040.767639 |
| conformer_35 | -1042.605105 | 0.327841 | 0.256914 | -1042.637104 | -84.0 | -1042.309263 | -1042.377178 | 23.3 | -1036.445145 | -4.549596 | -1040.766814 |
| conformer_58 | -1042.606868 | 0.327877 | 0.256183 | -1042.637054 | -79.3 | -1042.309177 | -1042.377859 | 21.5 | -1036.447939 | -4.548525 | -1040.767455 |
| conformer_25 | -1042.607386 | 0.327928 | 0.257318 | -1042.637020 | -77.8 | -1042.309092 | -1042.376690 | 24.6 | -1036.447705 | -4.549312 | -1040.766321 |
| conformer_27 | -1042.609446 | 0.327922 | 0.256630 | -1042.636863 | -72.0 | -1042.308941 | -1042.377221 | 23.2 | -1036.450722 | -4.548639 | -1040.767135 |
| conformer_24 | -1042.604665 | 0.327923 | 0.257637 | -1042.636278 | -83.0 | -1042.308355 | -1042.375629 | 27.4 | -1036.440940 | -4.553444 | -1040.765347 |
| conformer_99 | -1042.603586 | 0.327862 | 0.255855 | -1042.635928 | -84.9 | -1042.308066 | -1042.377061 | 23.6 | -1036.443668 | -4.549033 | -1040.766175 |
| conformer_26 | -1042.610075 | 0.327960 | 0.256653 | -1042.635645 | -67.1 | -1042.307685 | -1042.375980 | 26.5 | -1036.451924 | -4.548437 | -1040.766266 |
| conformer_39 | -1042.610644 | 0.328017 | 0.257570 | -1042.634622 | -63.0 | -1042.306605 | -1042.374040 | 31.6 | -1036.450651 | -4.550008 | -1040.764056 |

[a]: Single-point calculation in aqueous phase with SMD model.

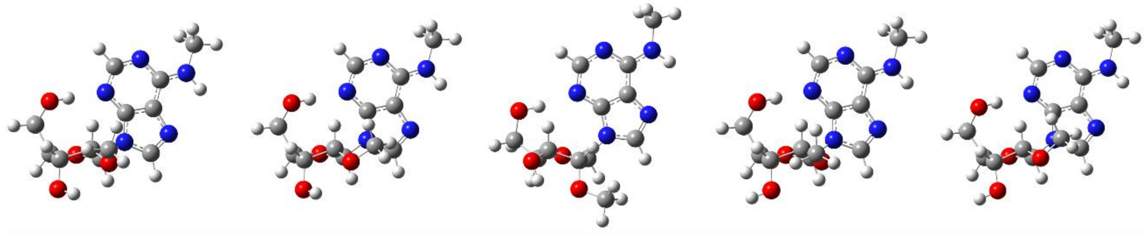

**Figure S52.** B3LYP-D3/def2-TZVPP optimized geometries of conformers for $N^6$-2’-*O*-dimethyladenosine (**9rbm6mA**).

**Table S50.** Conformers of gas-phase optimized $N^6$-2’-*O*-dimethyladenosine (**9rbm6mA**) at the B3LYP-D3/def2-TZVPP level of theory followed by aqueous phase single-point calculation. The columns display total energy without zero-point correction ($E_{Tot}$), Gibbs free energy (δG), total energy without zero-point correction ($E_{Tot,W}$), Gibbs free energy ($G_{298,W}$) in water (W), total single-point energy ($E_{CBS}$) calculated at DLPNO-CCSD(T)/CBS level of theory, and their corresponding free energy $G_{CBS}$. $G_{298,W}$ and $G_{CBS}$ have been corrected to the standard state of 1 mol/L by addition of +7.908 kJ/mol. $\Delta G_{Solv}$ represents the Gibbs free energy of solvation. The data are arranged in the ascending numeric order of $E_{Tot,W}$. $\Delta G_{298,W}$ represents the respective energy difference to the lowest structure. Only conformers within the 24 kJ/mol (6 kcal/mol) energy window above the lowest in CREST are included in initial conformer sampling. Duplicates of the same structure are excluded.

| 9rbm6dmA | B3LYP-D3/def2-TZVPP | | | SMD(H2O)/B3LYP-D3/def2-TZVPP[a] | | | | | DLPNO-CCSD(T)/CBS | | |
|---|---|---|---|---|---|---|---|---|---|---|---|
| No. | $E_{Tot}$ (Hartree) | δH (Hartree) | δG (Hartree) | $E_{Tot,W}$ (Hartree) | $\Delta G_{Solv}$ (kJ/mol) | $H_{298,W}$ (Hartree) | $G_{298,W}$ (Hartree) | $\Delta G_{298,W}$ (kJ/mol) | $E_{CBS,HF}$ (Hartree) | $E_{CBS,C}$ (Hartree) | $G_{CBS,W}$ (Hartree) |
| **conformer_1** | **-1042.617138** | **0.328299** | **0.259892** | **-1042.647847** | **-80.6** | **-1042.319548** | **-1042.384943** | **0.0** | **-1036.457903** | **-4.545632** | **-1040.771340** |
| conformer_2 | -1042.616405 | 0.328286 | 0.259523 | -1042.646501 | -79.0 | -1042.318215 | -1042.383966 | 2.6 | -1036.457141 | -4.545646 | -1040.770348 |
| conformer_31 | -1042.612190 | 0.328131 | 0.259650 | -1042.645632 | -87.8 | -1042.317501 | -1042.382970 | 5.2 | -1036.453990 | -4.544504 | -1040.769273 |
| conformer_76 | -1042.609321 | 0.328013 | 0.259099 | -1042.644563 | -92.5 | -1042.316550 | -1042.382452 | 6.5 | -1036.449987 | -4.545600 | -1040.768717 |
| conformer_4 | -1042.610141 | 0.328061 | 0.259008 | -1042.643045 | -86.4 | -1042.314984 | -1042.381025 | 10.3 | -1036.451643 | -4.544698 | -1040.767225 |
| conformer_50 | -1042.607941 | 0.328122 | 0.259115 | -1042.642289 | -90.2 | -1042.314167 | -1042.380162 | 12.6 | -1036.450359 | -4.544726 | -1040.767306 |
| conformer_8 | -1042.608544 | 0.328326 | 0.258036 | -1042.642166 | -88.3 | -1042.313840 | -1042.381118 | 10.0 | -1036.454111 | -4.541487 | -1040.768172 |
| conformer_52 | -1042.608371 | 0.328623 | 0.261109 | -1042.641693 | -87.5 | -1042.313070 | -1042.377572 | 19.4 | -1036.447028 | -4.546750 | -1040.762979 |
| conformer_59 | -1042.607404 | 0.328196 | 0.257278 | -1042.640383 | -86.6 | -1042.312187 | -1042.380093 | 12.7 | -1036.453538 | -4.541051 | -1040.767278 |
| conformer_96 | -1042.605111 | 0.327909 | 0.258579 | -1042.640264 | -92.3 | -1042.312355 | -1042.378673 | 16.5 | -1036.446809 | -4.544725 | -1040.765095 |
| conformer_15 | -1042.607116 | 0.328087 | 0.257381 | -1042.640093 | -86.6 | -1042.312006 | -1042.379700 | 13.8 | -1036.452805 | -4.542091 | -1040.767479 |
| conformer_18 | -1042.606355 | 0.328109 | 0.257402 | -1042.639541 | -87.1 | -1042.311432 | -1042.379127 | 15.3 | -1036.453484 | -4.540396 | -1040.766651 |
| conformer_13 | -1042.606899 | 0.328229 | 0.257051 | -1042.639489 | -85.6 | -1042.311260 | -1042.379426 | 14.5 | -1036.454189 | -4.540118 | -1040.766834 |
| conformer_70 | -1042.604403 | 0.327982 | 0.257269 | -1042.639361 | -91.8 | -1042.311379 | -1042.379080 | 15.4 | -1036.449596 | -4.542101 | -1040.766375 |
| conformer_64 | -1042.607779 | 0.328185 | 0.257213 | -1042.639271 | -82.7 | -1042.311086 | -1042.379046 | 15.5 | -1036.454731 | -4.540602 | -1040.766599 |
| conformer_77 | -1042.604417 | 0.328294 | 0.259765 | -1042.63917 | -91.2 | -1042.310876 | -1042.376393 | 22.5 | -1036.441591 | -4.547126 | -1040.760692 |
| conformer_78 | -1042.602563 | 0.327854 | 0.256827 | -1042.638833 | -95.2 | -1042.310979 | -1042.378994 | 15.6 | -1036.449685 | -4.540435 | -1040.766550 |
| conformer_27 | -1042.605047 | 0.328021 | 0.257077 | -1042.638811 | -88.6 | -1042.310790 | -1042.378722 | 16.3 | -1036.451379 | -4.541316 | -1040.766370 |
| conformer_16 | -1042.605495 | 0.328152 | 0.257267 | -1042.638733 | -87.3 | -1042.310581 | -1042.378454 | 17.0 | -1036.452393 | -4.540336 | -1040.765688 |
| conformer_88 | -1042.605053 | 0.327973 | 0.257945 | -1042.638697 | -88.3 | -1042.310724 | -1042.377740 | 18.9 | -1036.449255 | -4.542643 | -1040.764586 |
| conformer_30 | -1042.606940 | 0.328203 | 0.257812 | -1042.638661 | -83.3 | -1042.310458 | -1042.377837 | 18.7 | -1036.452383 | -4.542233 | -1040.765512 |
| conformer_67 | -1042.603563 | 0.327966 | 0.255931 | -1042.638544 | -91.8 | -1042.310578 | -1042.379601 | 14.0 | -1036.451133 | -4.540063 | -1040.767234 |
| conformer_44 | -1042.607684 | 0.328006 | 0.258466 | -1042.638406 | -80.7 | -1042.310400 | -1042.376928 | 21.0 | -1036.449509 | -4.544493 | -1040.763246 |
| conformer_65 | -1042.601668 | 0.327858 | 0.256942 | -1042.638352 | -96.3 | -1042.310494 | -1042.378398 | 17.2 | -1036.447450 | -4.541844 | -1040.766023 |
| conformer_66 | -1042.602614 | 0.327911 | 0.256731 | -1042.638114 | -93.2 | -1042.310203 | -1042.378371 | 17.3 | -1036.449222 | -4.540698 | -1040.765678 |
| conformer_92 | -1042.601305 | 0.327803 | 0.256590 | -1042.637879 | -96.0 | -1042.310076 | -1042.378277 | 17.5 | -1036.447378 | -4.541127 | -1040.765477 |
| conformer_9 | -1042.605838 | 0.328160 | 0.257918 | -1042.637846 | -84.0 | -1042.309686 | -1042.376916 | 21.1 | -1036.451948 | -4.541121 | -1040.764147 |
| conformer_33 | -1042.604798 | 0.328111 | 0.257108 | -1042.637663 | -86.3 | -1042.309552 | -1042.377543 | 19.4 | -1036.450956 | -4.541369 | -1040.765071 |
| conformer_62 | -1042.603442 | 0.327991 | 0.256582 | -1042.637662 | -89.8 | -1042.309671 | -1042.378068 | 18.1 | -1036.449213 | -4.541654 | -1040.765493 |

| | | | | | | | | | | | |
|---|---|---|---|---|---|---|---|---|---|---|---|
| conformer_63 | -1042.604468 | 0.327959 | 0.256717 | -1042.637621 | -87.0 | -1042.309662 | -1042.377892 | 18.5 | -1036.451403 | -4.541021 | -1040.765848 |
| conformer_5 | -1042.606449 | 0.328273 | 0.257271 | -1042.637506 | -81.5 | -1042.309233 | -1042.377223 | 20.3 | -1036.452902 | -4.540698 | -1040.764373 |
| conformer_29 | -1042.604753 | 0.328295 | 0.257781 | -1042.637052 | -84.8 | -1042.308757 | -1042.376259 | 22.8 | -1036.450564 | -4.540981 | -1040.763051 |
| conformer_85 | -1042.601565 | 0.327924 | 0.256879 | -1042.636694 | -92.2 | -1042.308770 | -1042.376803 | 21.4 | -1036.447283 | -4.541870 | -1040.764391 |
| conformer_84 | -1042.604236 | 0.328043 | 0.256771 | -1042.636453 | -84.6 | -1042.308410 | -1042.376670 | 21.7 | -1036.450943 | -4.541123 | -1040.764501 |

[a]: Single-point calculation in aqueous phase with SMD model.

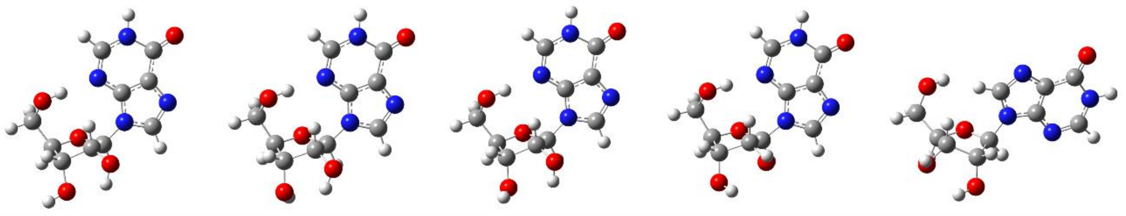

**Figure S53.** B3LYP-D3/def2-TZVPP optimized geometries of conformers for inosine (**9rbI**).

**Table S51.** Conformers of gas-phase optimized inosine (**9rbI**) at the B3LYP-D3/def2-TZVPP level of theory followed by aqueous phase single-point calculation. The columns display total energy without zero-point correction ($E_{Tot}$), Gibbs free energy (δG), total energy without zero-point correction ($E_{Tot,W}$), Gibbs free energy ($G_{298,W}$) in water (W), total single-point energy ($E_{CBS}$) calculated at DLPNO-CCSD(T)/CBS level of theory, and their corresponding free energy $G_{CBS}$. $G_{298,W}$ and $G_{CBS}$ have been corrected to the standard state of 1 mol/L by addition of +7.908 kJ/mol. $\Delta G_{Solv}$ represents the Gibbs free energy of solvation. The data are arranged in the ascending numeric order of $E_{Tot,W}$. $\Delta G_{298,W}$ represents the respective energy difference to the lowest structure. Only conformers within the 24 kJ/mol (6 kcal/mol) energy window above the lowest in CREST are included in initial conformer sampling. Duplicates of the same structure are excluded.

| | B3LYP-D3/def2-TZVPP | | | SMD(H2O)/B3LYP-D3/def2-TZVPP[a] | | | | | DLPNO-CCSD(T)/CBS | | |
|---|---|---|---|---|---|---|---|---|---|---|---|
| 9rbI No. | $E_{Tot}$ (Hartree) | δH (Hartree) | δG (Hartree) | $E_{Tot,W}$ (Hartree) | $\Delta G_{Solv}$ (kJ/mol) | $H_{298,W}$ (Hartree) | $G_{298,W}$ (Hartree) | $\Delta G_{298,W}$ (kJ/mol) | $E_{CBS,HF}$ (Hartree) | $E_{CBS,C}$ (Hartree) | $G_{CBS,W}$ (Hartree) |
| **conformer_3** | **-983.847915** | **0.256891** | **0.196751** | **-983.894365** | **-122.0** | **-983.637474** | **-983.694602** | **0.0** | **-978.212967** | **-4.160448** | **-982.220102** |
| conformer_2 | -983.846536 | 0.256772 | 0.197146 | -983.893930 | -124.4 | -983.637158 | -983.693772 | 2.2 | -978.210062 | -4.161528 | -982.218825 |
| conformer_7 | -983.848772 | 0.256886 | 0.197343 | -983.893808 | -118.2 | -983.636922 | -983.693453 | 3.0 | -978.214328 | -4.160374 | -982.219382 |
| conformer_1 | -983.848327 | 0.256737 | 0.196967 | -983.893740 | -119.2 | -983.637003 | -983.693761 | 2.2 | -978.213787 | -4.160229 | -982.219450 |
| conformer_54 | -983.840995 | 0.256633 | 0.195172 | -983.889866 | -128.3 | -983.633233 | -983.691682 | 7.7 | -978.210383 | -4.156998 | -982.218068 |
| conformer_79 | -983.837131 | 0.256552 | 0.193923 | -983.889849 | -138.4 | -983.633297 | -983.692914 | 4.4 | -978.206105 | -4.157072 | -982.218960 |
| conformer_51 | -983.843278 | 0.256775 | 0.196102 | -983.889806 | -122.2 | -983.633031 | -983.690692 | 10.3 | -978.209370 | -4.159420 | -982.216205 |
| conformer_45 | -983.840784 | 0.256721 | 0.195285 | -983.889573 | -128.1 | -983.632852 | -983.691276 | 8.7 | -978.210734 | -4.156484 | -982.217710 |
| conformer_80 | -983.838431 | 0.256439 | 0.193609 | -983.889534 | -134.2 | -983.633095 | -983.692913 | 4.4 | -978.209806 | -4.155318 | -982.219606 |
| conformer_63 | -983.835696 | 0.256393 | 0.193655 | -983.889249 | -140.6 | -983.632856 | -983.692582 | 5.3 | -978.204952 | -4.156874 | -982.218712 |
| conformer_82 | -983.839958 | 0.256546 | 0.196523 | -983.889095 | -129.0 | -983.632549 | -983.689560 | 13.2 | -978.204182 | -4.160930 | -982.214714 |
| conformer_5 | -983.845055 | 0.256660 | 0.196390 | -983.889076 | -115.6 | -983.632416 | -983.689674 | 12.9 | -978.210828 | -4.160214 | -982.215661 |
| conformer_32 | -983.842170 | 0.256505 | 0.195646 | -983.888989 | -122.9 | -983.632484 | -983.690331 | 11.2 | -978.208427 | -4.159516 | -982.216104 |
| conformer_56 | -983.838232 | 0.256702 | 0.195531 | -983.888870 | -132.9 | -983.632168 | -983.690327 | 11.2 | -978.205634 | -4.158328 | -982.216057 |
| conformer_128 | -983.839266 | 0.256402 | 0.193743 | -983.888675 | -129.7 | -983.632273 | -983.691920 | 7.0 | -978.211013 | -4.155225 | -982.218892 |
| conformer_109 | -983.837332 | 0.256422 | 0.193385 | -983.888617 | -134.6 | -983.632195 | -983.692220 | 6.3 | -978.208643 | -4.155550 | -982.219080 |
| conformer_10 | -983.839778 | 0.256656 | 0.196448 | -983.888594 | -128.2 | -983.631938 | -983.689134 | 14.4 | -978.204636 | -4.160954 | -982.214946 |
| conformer_4 | -983.846612 | 0.256660 | 0.195606 | -983.888544 | -110.1 | -983.631884 | -983.689926 | 12.3 | -978.212964 | -4.159812 | -982.216091 |
| conformer_88 | -983.836691 | 0.256337 | 0.193446 | -983.888493 | -136.0 | -983.632156 | -983.692035 | 6.7 | -978.209151 | -4.154368 | -982.218864 |
| conformer_105 | -983.837329 | 0.256361 | 0.193009 | -983.888337 | -133.9 | -983.631976 | -983.692316 | 6.0 | -978.208668 | -4.155723 | -982.219379 |
| conformer_117 | -983.839056 | 0.256573 | 0.194378 | -983.888322 | -129.3 | -983.631749 | -983.690932 | 9.6 | -978.210078 | -4.155294 | -982.217248 |
| conformer_19 | -983.840516 | 0.256559 | 0.196026 | -983.888273 | -125.4 | -983.631714 | -983.689235 | 14.1 | -978.207664 | -4.159354 | -982.215737 |
| conformer_90 | -983.839993 | 0.256487 | 0.194732 | -983.888090 | -126.3 | -983.631603 | -983.690346 | 11.2 | -978.210485 | -4.156181 | -982.217019 |
| conformer_65 | -983.833847 | 0.256365 | 0.193145 | -983.888064 | -142.3 | -983.631699 | -983.691907 | 7.1 | -978.204598 | -4.155345 | -982.218003 |
| conformer_151 | -983.835972 | 0.256235 | 0.193313 | -983.887968 | -136.5 | -983.631733 | -983.691643 | 7.8 | -978.206699 | -4.155473 | -982.217843 |
| conformer_78 | -983.835688 | 0.256482 | 0.194501 | -983.887936 | -137.2 | -983.631454 | -983.690423 | 11.0 | -978.204081 | -4.157209 | -982.216025 |
| conformer_125 | -983.832726 | 0.256210 | 0.192429 | -983.887882 | -144.8 | -983.631672 | -983.692441 | 5.7 | -978.203783 | -4.155007 | -982.218506 |
| conformer_112 | -983.838327 | 0.256382 | 0.194409 | -983.887795 | -129.9 | -983.631413 | -983.690374 | 11.1 | -978.209910 | -4.154970 | -982.216927 |
| conformer_57 | -983.837722 | 0.256599 | 0.195222 | -983.887782 | -131.4 | -983.631183 | -983.689548 | 13.3 | -978.206981 | -4.156710 | -982.215518 |
| conformer_24 | -983.839561 | 0.256671 | 0.194450 | -983.887432 | -125.7 | -983.630761 | -983.689970 | 12.2 | -978.210455 | -4.155835 | -982.216698 |

| | | | | | | | | | | | |
|---|---|---|---|---|---|---|---|---|---|---|---|
| conformer_135 | -983.839425 | 0.256436 | 0.194514 | -983.887380 | -125.9 | -983.630944 | -983.689854 | 12.5 | -978.210226 | -4.156149 | -982.216805 |
| conformer_101 | -983.835792 | 0.256329 | 0.193067 | -983.887316 | -135.3 | -983.630987 | -983.691237 | 8.8 | -978.209126 | -4.153548 | -982.218118 |
| conformer_11 | -983.844088 | 0.256624 | 0.195579 | -983.887295 | -113.4 | -983.630671 | -983.688704 | 15.5 | -978.211571 | -4.158909 | -982.215095 |
| conformer_49 | -983.839337 | 0.256554 | 0.193789 | -983.887226 | -125.7 | -983.630672 | -983.690425 | 11.0 | -978.211575 | -4.154733 | -982.217396 |
| conformer_139 | -983.836340 | 0.256231 | 0.192513 | -983.887226 | -133.6 | -983.630995 | -983.691701 | 7.6 | -978.208415 | -4.155102 | -982.218878 |
| conformer_87 | -983.834549 | 0.256620 | 0.193596 | -983.887124 | -138.0 | -983.630504 | -983.690516 | 10.7 | -978.202061 | -4.159382 | -982.217410 |
| conformer_137 | -983.839770 | 0.256360 | 0.194600 | -983.887123 | -124.3 | -983.630763 | -983.689511 | 13.4 | -978.208152 | -4.158628 | -982.216521 |
| conformer_97 | -983.840525 | 0.256450 | 0.194463 | -983.887120 | -122.3 | -983.630670 | -983.689645 | 13.0 | -978.211905 | -4.155665 | -982.216690 |
| conformer_40 | -983.843215 | 0.256549 | 0.195444 | -983.886969 | -114.9 | -983.630420 | -983.688513 | 16.0 | -978.209771 | -4.159428 | -982.214498 |
| conformer_75 | -983.836164 | 0.256518 | 0.194614 | -983.886916 | -133.2 | -983.630398 | -983.689290 | 13.9 | -978.204735 | -4.157894 | -982.215755 |
| conformer_118 | -983.833977 | 0.256329 | 0.193209 | -983.886880 | -138.9 | -983.630551 | -983.690659 | 10.4 | -978.205251 | -4.155049 | -982.216982 |
| conformer_124 | -983.836938 | 0.256441 | 0.194362 | -983.886818 | -131.0 | -983.630377 | -983.689444 | 13.5 | -978.206124 | -4.157165 | -982.215795 |
| conformer_39 | -983.841364 | 0.256545 | 0.195160 | -983.886705 | -119.0 | -983.630160 | -983.688533 | 15.9 | -978.209408 | -4.158328 | -982.214906 |
| conformer_8 | -983.845312 | 0.256628 | 0.195567 | -983.886513 | -108.2 | -983.629885 | -983.687934 | 17.5 | -978.212933 | -4.159000 | -982.214555 |
| conformer_6 | -983.843132 | 0.256604 | 0.195391 | -983.886491 | -113.8 | -983.629887 | -983.688088 | 17.1 | -978.211381 | -4.158100 | -982.214437 |
| conformer_41 | -983.836392 | 0.256455 | 0.194096 | -983.886442 | -131.4 | -983.629987 | -983.689334 | 13.8 | -978.205777 | -4.156955 | -982.215674 |
| conformer_26 | -983.842021 | 0.256457 | 0.195048 | -983.886441 | -116.6 | -983.629984 | -983.688381 | 16.3 | -978.209732 | -4.158500 | -982.214591 |
| conformer_111 | -983.836377 | 0.256360 | 0.193190 | -983.886423 | -131.4 | -983.630063 | -983.690221 | 11.5 | -978.210373 | -4.153062 | -982.217279 |
| conformer_18 | -983.840116 | 0.256555 | 0.194547 | -983.886412 | -121.5 | -983.629857 | -983.688853 | 15.1 | -978.210321 | -4.156756 | -982.215813 |
| conformer_59 | -983.838684 | 0.256403 | 0.193417 | -983.886280 | -125.0 | -983.629877 | -983.689851 | 12.5 | -978.209954 | -4.155672 | -982.216793 |
| conformer_74 | -983.838866 | 0.256574 | 0.194051 | -983.886150 | -124.1 | -983.629576 | -983.689087 | 14.5 | -978.210098 | -4.155736 | -982.216055 |
| conformer_129 | -983.836151 | 0.256377 | 0.193124 | -983.886140 | -131.2 | -983.629763 | -983.690004 | 12.1 | -978.208645 | -4.154606 | -982.217105 |
| conformer_142 | -983.841025 | 0.256470 | 0.195104 | -983.886105 | -118.4 | -983.629635 | -983.687989 | 17.4 | -978.209621 | -4.158167 | -982.214753 |
| conformer_31 | -983.838039 | 0.256467 | 0.194554 | -983.886000 | -125.9 | -983.629533 | -983.688434 | 16.2 | -978.209045 | -4.155923 | -982.215362 |
| conformer_44 | -983.838537 | 0.256786 | 0.195063 | -983.885900 | -124.4 | -983.629114 | -983.687825 | 17.8 | -978.208593 | -4.156396 | -982.214277 |
| conformer_94 | -983.836187 | 0.256294 | 0.193926 | -983.885857 | -130.4 | -983.629563 | -983.688919 | 14.9 | -978.206420 | -4.156685 | -982.215838 |
| conformer_12 | -983.836231 | 0.256545 | 0.195036 | -983.885829 | -130.2 | -983.629284 | -983.687781 | 17.9 | -978.204727 | -4.157676 | -982.213953 |
| conformer_15 | -983.837772 | 0.256570 | 0.195469 | -983.885609 | -125.6 | -983.629039 | -983.687128 | 19.6 | -978.201567 | -4.162332 | -982.213254 |
| conformer_34 | -983.835131 | 0.256447 | 0.194269 | -983.885584 | -132.5 | -983.629137 | -983.688303 | 16.5 | -978.203754 | -4.157581 | -982.214507 |
| conformer_89 | -983.837765 | 0.256399 | 0.193765 | -983.885415 | -125.1 | -983.629016 | -983.688638 | 15.7 | -978.209412 | -4.155504 | -982.215790 |
| conformer_52 | -983.842555 | 0.256558 | 0.195196 | -983.885395 | -112.5 | -983.628837 | -983.687187 | 19.5 | -978.211433 | -4.157785 | -982.213850 |
| conformer_64 | -983.833136 | 0.256497 | 0.194760 | -983.885329 | -137.0 | -983.628832 | -983.687557 | 18.5 | -978.200916 | -4.158398 | -982.213736 |
| conformer_29 | -983.835218 | 0.256607 | 0.194806 | -983.885203 | -131.2 | -983.628596 | -983.687385 | 18.9 | -978.203772 | -4.157643 | -982.213583 |
| conformer_106 | -983.831490 | 0.256553 | 0.194480 | -983.885138 | -140.9 | -983.628585 | -983.687646 | 18.3 | -978.198715 | -4.159350 | -982.214221 |
| conformer_66 | -983.839544 | 0.256575 | 0.194417 | -983.885011 | -119.4 | -983.628436 | -983.687582 | 18.4 | -978.211479 | -4.155522 | -982.215038 |
| conformer_43 | -983.838396 | 0.256472 | 0.194458 | -983.884955 | -122.2 | -983.628483 | -983.687485 | 18.7 | -978.210135 | -4.155394 | -982.214619 |
| conformer_38 | -983.838916 | 0.256697 | 0.193866 | -983.884753 | -120.3 | -983.628056 | -983.687875 | 17.7 | -978.210477 | -4.155098 | -982.214535 |
| conformer_93 | -983.833494 | 0.256488 | 0.194201 | -983.884715 | -134.5 | -983.628227 | -983.687502 | 18.6 | -978.201337 | -4.158932 | -982.214277 |
| conformer_42 | -983.836011 | 0.256477 | 0.194519 | -983.884572 | -127.5 | -983.628095 | -983.687041 | 19.9 | -978.205368 | -4.157169 | -982.213566 |
| conformer_73 | -983.835013 | 0.256820 | 0.197068 | -983.884518 | -130.0 | -983.627698 | -983.684438 | 26.7 | -978.198200 | -4.163432 | -982.211057 |
| conformer_47 | -983.838088 | 0.256516 | 0.194363 | -983.884460 | -121.8 | -983.627944 | -983.687085 | 19.7 | -978.208350 | -4.156376 | -982.213723 |
| conformer_27 | -983.835621 | 0.256514 | 0.194868 | -983.884428 | -128.1 | -983.627914 | -983.686548 | 21.1 | -978.204103 | -4.157632 | -982.212662 |
| conformer_16 | -983.840794 | 0.256559 | 0.194401 | -983.884408 | -114.5 | -983.627849 | -983.686995 | 20.0 | -978.211292 | -4.156522 | -982.214015 |
| conformer_146 | -983.834108 | 0.256483 | 0.194287 | -983.884318 | -131.8 | -983.627835 | -983.687019 | 19.9 | -978.202340 | -4.158818 | -982.214068 |
| conformer_83 | -983.834244 | 0.256392 | 0.194943 | -983.884057 | -130.8 | -983.627665 | -983.686102 | 22.3 | -978.198329 | -4.162125 | -982.212313 |
| conformer_153 | -983.834431 | 0.256482 | 0.194653 | -983.883902 | -129.9 | -983.627420 | -983.686237 | 22.0 | -978.200066 | -4.160519 | -982.212392 |
| conformer_25 | -983.835558 | 0.256623 | 0.194752 | -983.883579 | -126.1 | -983.626956 | -983.685815 | 23.1 | -978.201295 | -4.160963 | -982.212515 |
| conformer_23 | -983.841198 | 0.256556 | 0.194349 | -983.883340 | -110.6 | -983.626784 | -983.685979 | 22.6 | -978.212520 | -4.156171 | -982.213472 |
| conformer_86 | -983.834995 | 0.256618 | 0.193982 | -983.883339 | -126.9 | -983.626721 | -983.686345 | 21.7 | -978.204253 | -4.156815 | -982.212418 |
| conformer_144 | -983.833595 | 0.256265 | 0.195252 | -983.882232 | -127.7 | -983.625967 | -983.683968 | 27.9 | -978.199652 | -4.160898 | -982.210922 |
| conformer_30 | -983.832750 | 0.256621 | 0.195103 | -983.881704 | -128.5 | -983.625083 | -983.683589 | 28.9 | -978.197952 | -4.161645 | -982.210436 |
| conformer_21 | -983.833437 | 0.256659 | 0.193893 | -983.880951 | -124.7 | -983.624292 | -983.684046 | 27.7 | -978.199836 | -4.160213 | -982.210658 |
| conformer_141 | -983.831829 | 0.256451 | 0.194647 | -983.880851 | -128.7 | -983.624400 | -983.683192 | 30.0 | -978.197875 | -4.161300 | -982.210538 |
| conformer_9 | -983.839528 | 0.256712 | 0.195068 | -983.880693 | -108.1 | -983.623981 | -983.682613 | 31.5 | -978.209503 | -4.157206 | -982.209794 |

[a]: Single-point calculation in aqueous phase with SMD model.

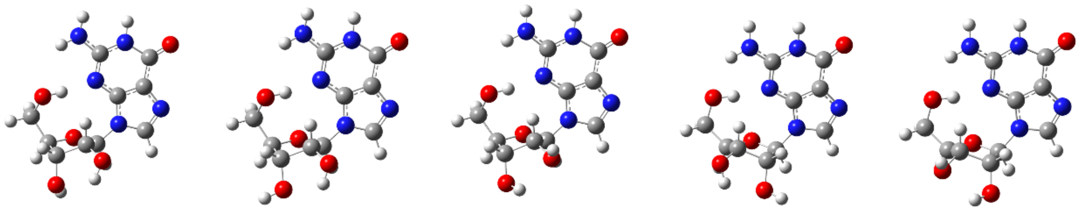

conformer_1 conformer_2 conformer_6 conformer_7 conformer_56

**Figure S54.** B3LYP-D3/def2-TZVPP optimized geometries of conformers for guanosine (**9rbG**).

**Table S52.** Conformers of gas-phase optimized guanosine (**9rbG**) at the B3LYP-D3/def2-TZVPP level of theory followed by aqueous phase single-point calculation. The columns display total energy without zero-point correction ($E_{Tot}$), Gibbs free energy ($\delta G$), total energy without zero-point correction ($E_{Tot,W}$), Gibbs free energy ($G_{298,W}$) in water (W), total single-point energy ($E_{CBS}$) calculated at DLPNO-CCSD(T)/CBS level of theory, and their corresponding free energy $G_{CBS}$. $G_{298,W}$ and $G_{CBS}$ have been corrected to the standard state of 1 mol/L by addition of +7.908 kJ/mol. $\Delta G_{Solv}$ represents the Gibbs free energy of solvation. The data are arranged in the ascending numeric order of $E_{Tot,W}$. $\Delta G_{298,W}$ represents the respective energy difference to the lowest structure. Only conformers within the 24 kJ/mol (6 kcal/mol) energy window above the lowest in CREST are included in initial conformer sampling. Duplicates of the same structure are excluded.

| | B3LYP-D3/def2-TZVPP | | | SMD(H2O)/B3LYP-D3/def2-TZVPP[a] | | | | | DLPNO-CCSD(T)/CBS | | |
|---|---|---|---|---|---|---|---|---|---|---|---|
| 9rbG No. | $E_{Tot}$ (Hartree) | $\delta H$ (Hartree) | $\delta G$ (Hartree) | $E_{Tot,W}$ (Hartree) | $\Delta G_{Solv}$ (kJ/mol) | $H_{298,W}$ (Hartree) | $G_{298,W}$ (Hartree) | $\Delta G_{298,W}$ (kJ/mol) | $E_{CBS,HF}$ (Hartree) | $E_{CBS,C}$ (Hartree) | $G_{CBS,W}$ (Hartree) |
| **conformer_1** | **-1039.245115** | **0.274774** | **0.212386** | **-1039.298128** | **-139.2** | **-1039.023354** | **-1039.082730** | **0.0** | **-1033.286208** | **-4.403786** | **-1037.527609** |
| conformer_6 | -1039.247527 | 0.274914 | 0.212436 | -1039.298004 | -132.5 | -1039.023090 | -1039.082556 | 0.5 | -1033.290704 | -4.402575 | -1037.528308 |
| conformer_2 | -1039.246687 | 0.274785 | 0.212320 | -1039.297877 | -134.4 | -1039.023092 | -1039.082545 | 0.5 | -1033.289699 | -4.402562 | -1037.528119 |
| conformer_56 | -1039.245030 | 0.274788 | 0.212249 | -1039.293354 | -126.9 | -1039.018566 | -1039.078093 | 12.2 | -1033.287754 | -4.403115 | -1037.523933 |
| conformer_7 | -1039.242935 | 0.274615 | 0.211732 | -1039.293179 | -131.9 | -1039.018564 | -1039.078435 | 11.3 | -1033.286025 | -4.402395 | -1037.523920 |
| conformer_8 | -1039.240656 | 0.274612 | 0.212168 | -1039.293045 | -137.5 | -1039.018433 | -1039.077865 | 12.8 | -1033.281775 | -4.403778 | -1037.522762 |
| conformer_88 | -1039.237474 | 0.274661 | 0.210067 | -1039.292959 | -145.7 | -1039.018298 | -1039.079880 | 7.5 | -1033.286565 | -4.397491 | -1037.526462 |
| conformer_78 | -1039.236948 | 0.274736 | 0.209998 | -1039.292697 | -146.4 | -1039.017961 | -1039.079687 | 8.0 | -1033.286786 | -4.396932 | -1037.526456 |
| conformer_86 | -1039.236725 | 0.274704 | 0.209942 | -1039.292596 | -146.7 | -1039.017892 | -1039.079642 | 8.1 | -1033.286557 | -4.396898 | -1037.526372 |
| conformer_11 | -1039.243030 | 0.274809 | 0.210974 | -1039.292536 | -130.0 | -1039.017727 | -1039.078550 | 11.0 | -1033.288640 | -4.400671 | -1037.524830 |
| conformer_9 | -1039.239383 | 0.274907 | 0.212632 | -1039.292254 | -138.8 | -1039.017347 | -1039.076610 | 16.1 | -1033.281133 | -4.403787 | -1037.522148 |
| conformer_13 | -1039.239949 | 0.274807 | 0.212320 | -1039.292184 | -137.1 | -1039.017377 | -1039.076852 | 15.4 | -1033.283885 | -4.402261 | -1037.523049 |
| conformer_90 | -1039.234622 | 0.274709 | 0.210461 | -1039.292134 | -151.0 | -1039.017425 | -1039.078661 | 10.7 | -1033.281750 | -4.398803 | -1037.524592 |
| conformer_38 | -1039.239999 | 0.274724 | 0.212037 | -1039.291618 | -135.5 | -1039.016894 | -1039.076569 | 16.2 | -1033.284372 | -4.402151 | -1037.523093 |
| conformer_18 | -1039.240276 | 0.274513 | 0.209525 | -1039.291198 | -133.7 | -1039.016685 | -1039.078661 | 10.7 | -1033.287463 | -4.399545 | -1037.525392 |
| conformer_43 | -1039.235865 | 0.274717 | 0.209364 | -1039.290809 | -144.3 | -1039.016092 | -1039.078433 | 11.3 | -1033.286348 | -4.396554 | -1037.525471 |
| conformer_47 | -1039.235788 | 0.274703 | 0.209325 | -1039.290749 | -144.3 | -1039.016046 | -1039.078412 | 11.3 | -1033.286230 | -4.396504 | -1037.525358 |
| conformer_68 | -1039.238983 | 0.274519 | 0.210151 | -1039.290601 | -135.5 | -1039.016082 | -1039.077438 | 13.9 | -1033.285294 | -4.400025 | -1037.523774 |
| conformer_99 | -1039.235282 | 0.274559 | 0.208978 | -1039.290581 | -145.2 | -1039.016022 | -1039.078591 | 10.9 | -1033.287236 | -4.395411 | -1037.525956 |
| conformer_100 | -1039.235118 | 0.274531 | 0.209018 | -1039.290493 | -145.4 | -1039.015962 | -1039.078463 | 11.2 | -1033.287048 | -4.395413 | -1037.525806 |
| conformer_15 | -1039.239283 | 0.274659 | 0.210115 | -1039.290486 | -134.4 | -1039.015827 | -1039.077359 | 14.1 | -1033.287159 | -4.398703 | -1037.523938 |
| conformer_57 | -1039.237290 | 0.274468 | 0.209334 | -1039.290236 | -139.0 | -1039.015768 | -1039.077890 | 12.7 | -1033.285144 | -4.398830 | -1037.524574 |
| conformer_58 | -1039.237287 | 0.274455 | 0.209397 | -1039.290070 | -138.6 | -1039.015615 | -1039.077661 | 13.3 | -1033.284992 | -4.398991 | -1037.524358 |
| conformer_23 | -1039.241042 | 0.274600 | 0.210086 | -1039.289913 | -128.3 | -1039.015313 | -1039.076815 | 15.5 | -1033.288483 | -4.399528 | -1037.523783 |
| conformer_72 | -1039.235332 | 0.274464 | 0.208613 | -1039.289508 | -142.2 | -1039.015044 | -1039.077883 | 12.7 | -1033.286111 | -4.396410 | -1037.525072 |
| conformer_48 | -1039.233115 | 0.274531 | 0.209289 | -1039.289500 | -148.0 | -1039.014969 | -1039.077199 | 14.5 | -1033.281898 | -4.397746 | -1037.523729 |
| conformer_62 | -1039.235240 | 0.274464 | 0.208814 | -1039.289487 | -142.4 | -1039.015023 | -1039.077661 | 13.3 | -1033.286008 | -4.396359 | -1037.524788 |
| conformer_49 | -1039.235456 | 0.274526 | 0.209014 | -1039.289461 | -141.8 | -1039.014935 | -1039.077435 | 13.9 | -1033.285079 | -4.397604 | -1037.524661 |
| conformer_65 | -1039.238697 | 0.274609 | 0.210056 | -1039.289385 | -133.1 | -1039.014776 | -1039.076317 | 16.8 | -1033.287192 | -4.398416 | -1037.523228 |
| conformer_50 | -1039.233864 | 0.274553 | 0.209695 | -1039.289321 | -145.6 | -1039.014768 | -1039.076614 | 16.1 | -1033.284176 | -4.396838 | -1037.523764 |

| | | | | | | | | | | | |
|---|---|---|---|---|---|---|---|---|---|---|---|
| conformer_81 | -1039.234660 | 0.274829 | 0.210007 | -1039.289185 | -143.2 | -1039.014356 | -1039.076166 | 17.2 | -1033.284138 | -4.397114 | -1037.522758 |
| conformer_71 | -1039.234794 | 0.274836 | 0.209888 | -1039.289156 | -142.7 | -1039.014320 | -1039.076256 | 17.0 | -1033.284302 | -4.397194 | -1037.522957 |
| conformer_64 | -1039.230886 | 0.274446 | 0.209137 | -1039.289113 | -152.9 | -1039.014667 | -1039.076964 | 15.1 | -1033.278844 | -4.398502 | -1037.523424 |
| conformer_28 | -1039.234747 | 0.274651 | 0.210768 | -1039.289092 | -142.7 | -1039.014441 | -1039.075312 | 19.5 | -1033.277951 | -4.402880 | -1037.521395 |
| conformer_21 | -1039.235045 | 0.274717 | 0.210948 | -1039.289055 | -141.8 | -1039.014338 | -1039.075095 | 20.0 | -1033.278265 | -4.402952 | -1037.521267 |
| conformer_45 | -1039.235819 | 0.274558 | 0.209424 | -1039.289051 | -139.8 | -1039.014493 | -1039.076615 | 16.1 | -1033.284844 | -4.398182 | -1037.523821 |
| conformer_66 | -1039.230772 | 0.274429 | 0.208961 | -1039.288993 | -152.9 | -1039.014564 | -1039.077020 | 15.0 | -1033.278785 | -4.398458 | -1037.523491 |
| conformer_16 | -1039.232436 | 0.274603 | 0.209722 | -1039.288817 | -148.0 | -1039.014214 | -1039.076083 | 17.5 | -1033.279872 | -4.398904 | -1037.522423 |
| conformer_80 | -1039.234280 | 0.274517 | 0.209447 | -1039.288689 | -142.9 | -1039.014172 | -1039.076230 | 17.1 | -1033.285175 | -4.396502 | -1037.523627 |
| conformer_97 | -1039.234321 | 0.274370 | 0.207880 | -1039.288251 | -141.6 | -1039.013881 | -1039.077359 | 14.1 | -1033.285388 | -4.396497 | -1037.524924 |
| conformer_46 | -1039.231857 | 0.274673 | 0.210073 | -1039.288023 | -147.5 | -1039.013350 | -1039.074938 | 20.5 | -1033.280294 | -4.397980 | -1037.521355 |
| conformer_84 | -1039.233807 | 0.274542 | 0.209370 | -1039.287949 | -142.2 | -1039.013407 | -1039.075567 | 18.8 | -1033.285167 | -4.396135 | -1037.523062 |
| conformer_44 | -1039.236770 | 0.274697 | 0.209827 | -1039.287798 | -134.0 | -1039.013101 | -1039.074959 | 20.4 | -1033.286642 | -4.397358 | -1037.522188 |
| conformer_73 | -1039.234673 | 0.274613 | 0.209476 | -1039.287796 | -139.5 | -1039.013183 | -1039.075308 | 19.5 | -1033.284377 | -4.397222 | -1037.522234 |
| conformer_52 | -1039.236888 | 0.274718 | 0.209983 | -1039.287777 | -133.6 | -1039.013059 | -1039.074782 | 20.9 | -1033.286707 | -4.397452 | -1037.522053 |
| conformer_61 | -1039.234528 | 0.274584 | 0.209474 | -1039.287758 | -139.8 | -1039.013174 | -1039.075272 | 19.6 | -1033.284219 | -4.397135 | -1037.522097 |
| conformer_35 | -1039.232195 | 0.274547 | 0.209614 | -1039.287677 | -145.7 | -1039.013130 | -1039.075051 | 20.2 | -1033.279976 | -4.398526 | -1037.521358 |
| conformer_31 | -1039.232201 | 0.274560 | 0.209782 | -1039.287629 | -145.5 | -1039.013069 | -1039.074835 | 20.7 | -1033.279942 | -4.398566 | -1037.521142 |
| conformer_59 | -1039.231706 | 0.274551 | 0.209227 | -1039.287291 | -145.9 | -1039.012740 | -1039.075052 | 20.2 | -1033.280302 | -4.398298 | -1037.521946 |
| conformer_51 | -1039.232595 | 0.274658 | 0.210043 | -1039.286908 | -142.6 | -1039.012250 | -1039.073853 | 23.3 | -1033.277588 | -4.401680 | -1037.520527 |
| conformer_26 | -1039.232634 | 0.274649 | 0.209967 | -1039.286774 | -142.1 | -1039.012125 | -1039.073795 | 23.5 | -1033.277784 | -4.401681 | -1037.520625 |
| conformer_63 | -1039.237212 | 0.274733 | 0.210033 | -1039.286655 | -129.8 | -1039.011922 | -1039.073610 | 23.9 | -1033.287939 | -4.397007 | -1037.521345 |
| conformer_83 | -1039.236898 | 0.274678 | 0.209815 | -1039.286652 | -130.6 | -1039.011974 | -1039.073825 | 23.4 | -1033.287705 | -4.396910 | -1037.521542 |
| conformer_33 | -1039.232586 | 0.274830 | 0.212172 | -1039.285888 | -139.9 | -1039.011058 | -1039.070704 | 31.6 | -1033.273603 | -4.404943 | -1037.516664 |
| conformer_67 | -1039.232041 | 0.274559 | 0.211381 | -1039.285437 | -140.2 | -1039.010878 | -1039.071044 | 30.7 | -1033.275553 | -4.402660 | -1037.517216 |
| conformer_30 | -1039.229876 | 0.274666 | 0.209920 | -1039.284892 | -144.4 | -1039.010226 | -1039.071960 | 28.3 | -1033.274338 | -4.402484 | -1037.518907 |
| conformer_41 | -1039.229515 | 0.274587 | 0.209628 | -1039.284862 | -145.3 | -1039.010275 | -1039.072222 | 27.6 | -1033.274182 | -4.402327 | -1037.519216 |
| conformer_19 | -1039.235585 | 0.274714 | 0.210038 | -1039.284443 | -128.3 | -1039.009729 | -1039.071393 | 29.8 | -1033.284932 | -4.398074 | -1037.518814 |
| conformer_85 | -1039.235565 | 0.274700 | 0.210029 | -1039.284433 | -128.3 | -1039.009733 | -1039.071392 | 29.8 | -1033.284728 | -4.398239 | -1037.518794 |
| conformer_93 | -1039.231850 | 0.274434 | 0.210821 | -1039.284144 | -137.3 | -1039.009710 | -1039.070311 | 32.6 | -1033.275626 | -4.402738 | -1037.516825 |
| conformer_91 | -1039.231734 | 0.274447 | 0.210280 | -1039.283341 | -135.5 | -1039.008894 | -1039.070049 | 33.3 | -1033.275823 | -4.402275 | -1037.516414 |
| conformer_24 | -1039.234561 | 0.274811 | 0.211209 | -1039.282816 | -126.7 | -1039.008005 | -1039.068595 | 37.1 | -1033.277824 | -4.402849 | -1037.514706 |
| conformer_34 | -1039.233360 | 0.274688 | 0.211212 | -1039.281286 | -125.8 | -1039.006598 | -1039.067062 | 41.1 | -1033.276552 | -4.402589 | -1037.512843 |
| conformer_20 | -1039.231869 | 0.274747 | 0.211618 | -1039.280897 | -128.7 | -1039.006150 | -1039.066267 | 43.2 | -1033.272975 | -4.404196 | -1037.511569 |

[a]: Single-point calculation in aqueous phase with SMD model.

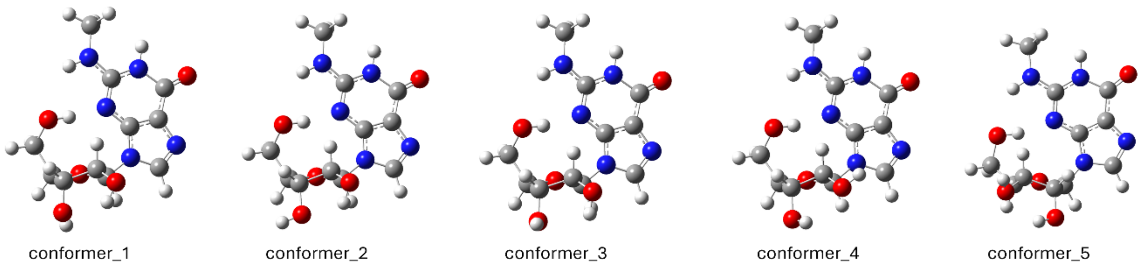

**Figure S55.** B3LYP-D3/def2-TZVPP optimized geometries of conformers for $N^2$-methylguanosine (**9rb2mG**).

**Table S53.** Conformers of gas-phase optimized $N^2$-methylguanosine (**9rb2mG**) at the B3LYP-D3/def2-TZVPP level of theory followed by aqueous phase single-point calculation. The columns display total energy without zero-point correction ($E_{Tot}$), Gibbs free energy (δG), total energy without zero-point correction ($E_{Tot,W}$), Gibbs free energy ($G_{298,W}$) in water (W), total single-point energy ($E_{CBS}$) calculated at DLPNO-CCSD(T)/CBS level of theory, and their corresponding free energy $G_{CBS}$. $G_{298,W}$ and $G_{CBS}$ have been corrected to the standard state of 1 mol/L by addition of +7.908 kJ/mol. $\Delta G_{Solv}$ represents the Gibbs free energy of solvation. The data are arranged in the ascending numeric order of $E_{Tot,W}$. $\Delta G_{298,W}$ represents the respective energy difference to the lowest structure. Only conformers within the 24 kJ/mol (6 kcal/mol) energy window above the lowest in CREST are included in initial conformer sampling. Duplicates of the same structure are excluded.

| | B3LYP-D3/def2-TZVPP | | | SMD(H2O)/B3LYP-D3/def2-TZVPP[a] | | | | | DLPNO-CCSD(T)/CBS | | |
|---|---|---|---|---|---|---|---|---|---|---|---|
| 9rb2mG No. | $E_{Tot}$ (Hartree) | δH (Hartree) | δG (Hartree) | $E_{Tot,W}$ (Hartree) | $\Delta G_{Solv}$ (kJ/mol) | $H_{298,W}$ (Hartree) | $G_{298,W}$ (Hartree) | $\Delta G_{298,W}$ (kJ/mol) | $E_{CBS,HF}$ (Hartree) | $E_{CBS,C}$ (Hartree) | $G_{CBS,W}$ (Hartree) |
| **conformer_3** | **-1078.570203** | **0.304570** | **0.237662** | **-1078.620814** | **-132.9** | **-1078.316244** | **-1078.380140** | **0.0** | **-1072.330371** | **-4.610180** | **-1076.750488** |
| conformer_1 | -1078.568949 | 0.304396 | 0.237629 | -1078.620166 | -134.5 | -1078.315770 | -1078.379525 | 1.6 | -1072.327605 | -4.611205 | -1076.749386 |
| conformer_2 | -1078.570415 | 0.304363 | 0.237453 | -1078.619853 | -129.8 | -1078.315490 | -1078.379388 | 2.0 | -1072.331042 | -4.609921 | -1076.749937 |
| conformer_4 | -1078.571083 | 0.304400 | 0.237369 | -1078.619195 | -126.3 | -1078.314795 | -1078.378814 | 3.5 | -1072.332540 | -4.609529 | -1076.749801 |
| conformer_5 | -1078.568535 | 0.304400 | 0.237916 | -1078.615392 | -123.0 | -1078.310992 | -1078.374464 | 14.9 | -1072.328989 | -4.610578 | -1076.745495 |
| conformer_6 | -1078.566685 | 0.304217 | 0.237160 | -1078.615271 | -127.6 | -1078.311054 | -1078.375099 | 13.2 | -1072.327527 | -4.609792 | -1076.745733 |
| conformer_7 | -1078.564460 | 0.304242 | 0.237650 | -1078.615044 | -132.8 | -1078.310802 | -1078.374382 | 15.1 | -1072.323282 | -4.611292 | -1076.744496 |
| conformer_80 | -1078.560555 | 0.304220 | 0.234711 | -1078.614681 | -142.1 | -1078.310461 | -1078.376958 | 8.4 | -1072.327617 | -4.604452 | -1076.748472 |
| conformer_27 | -1078.562262 | 0.304437 | 0.237474 | -1078.614571 | -137.3 | -1078.310134 | -1078.374085 | 15.9 | -1072.323212 | -4.609982 | -1076.745017 |
| conformer_9 | -1078.566461 | 0.304316 | 0.235746 | -1078.614435 | -126.0 | -1078.310119 | -1078.375677 | 11.7 | -1072.330010 | -4.607902 | -1076.747127 |
| conformer_69 | -1078.559936 | 0.304234 | 0.234920 | -1078.614342 | -142.8 | -1078.310108 | -1078.376410 | 9.8 | -1072.327777 | -4.604004 | -1076.748255 |
| conformer_8 | -1078.562928 | 0.304457 | 0.238172 | -1078.614135 | -134.4 | -1078.309678 | -1078.372951 | 18.9 | -1072.322330 | -4.611258 | -1076.743611 |
| conformer_13 | -1078.563348 | 0.304338 | 0.237695 | -1078.613906 | -132.7 | -1078.309568 | -1078.373199 | 18.2 | -1072.324933 | -4.609780 | -1076.744564 |
| conformer_79 | -1078.557755 | 0.304220 | 0.234871 | -1078.613725 | -147.0 | -1078.309505 | -1078.375842 | 11.3 | -1072.322930 | -4.605728 | -1076.746745 |
| conformer_31 | -1078.563546 | 0.304255 | 0.237417 | -1078.613238 | -130.5 | -1078.308983 | -1078.372809 | 19.2 | -1072.325531 | -4.609582 | -1076.744375 |
| conformer_16 | -1078.563931 | 0.304175 | 0.234611 | -1078.613065 | -129.0 | -1078.308890 | -1078.375442 | 12.3 | -1072.328857 | -4.606820 | -1076.747188 |
| conformer_36 | -1078.559063 | 0.304293 | 0.234486 | -1078.612506 | -140.3 | -1078.308213 | -1078.375008 | 13.5 | -1072.327382 | -4.603607 | -1076.746933 |
| conformer_55 | -1078.562375 | 0.304112 | 0.235406 | -1078.612393 | -131.3 | -1078.308281 | -1078.373975 | 16.2 | -1072.326759 | -4.607079 | -1076.745438 |
| conformer_14 | -1078.562713 | 0.304143 | 0.235099 | -1078.612113 | -129.7 | -1078.307970 | -1078.374002 | 16.1 | -1072.328622 | -4.605847 | -1076.745758 |
| conformer_48 | -1078.560740 | 0.304104 | 0.235172 | -1078.612070 | -134.8 | -1078.307966 | -1078.373886 | 16.4 | -1072.326483 | -4.606015 | -1076.745644 |
| conformer_24 | -1078.564400 | 0.304210 | 0.235243 | -1078.611893 | -124.7 | -1078.307683 | -1078.373638 | 17.1 | -1072.329759 | -4.606675 | -1076.745672 |
| conformer_45 | -1078.558582 | 0.304033 | 0.234035 | -1078.611289 | -138.4 | -1078.307256 | -1078.374242 | 15.5 | -1072.327304 | -4.603406 | -1076.746369 |
| conformer_50 | -1078.562085 | 0.304079 | 0.234776 | -1078.611012 | -128.5 | -1078.306933 | -1078.373224 | 18.2 | -1072.328614 | -4.605553 | -1076.745307 |
| conformer_38 | -1078.558839 | 0.304169 | 0.235292 | -1078.610905 | -136.7 | -1078.306736 | -1078.372601 | 19.8 | -1072.325548 | -4.605562 | -1076.744872 |
| conformer_51 | -1078.554018 | 0.304104 | 0.234742 | -1078.610803 | -149.1 | -1078.306699 | -1078.373049 | 18.6 | -1072.319658 | -4.605866 | -1076.744556 |
| conformer_56 | -1078.557934 | 0.304295 | 0.234763 | -1078.610759 | -138.7 | -1078.306464 | -1078.372984 | 18.8 | -1072.325349 | -4.604274 | -1076.744673 |
| conformer_43 | -1078.556941 | 0.304121 | 0.234844 | -1078.610754 | -141.3 | -1078.306633 | -1078.372898 | 19.0 | -1072.325045 | -4.604077 | -1076.745080 |
| conformer_19 | -1078.558291 | 0.304245 | 0.236124 | -1078.610737 | -137.7 | -1078.306492 | -1078.371601 | 22.4 | -1072.319375 | -4.610115 | -1076.742800 |
| conformer_15 | -1078.555794 | 0.304324 | 0.235568 | -1078.610484 | -143.6 | -1078.306160 | -1078.371904 | 21.6 | -1072.321019 | -4.606128 | -1076.743257 |

| | | | | | | | | | | | |
|---|---|---|---|---|---|---|---|---|---|---|---|
| conformer_35 | -1078.556260 | 0.304176 | 0.234333 | -1078.610359 | -142.0 | -1078.306183 | -1078.373014 | 18.7 | -1072.321619 | -4.605854 | -1076.744228 |
| conformer_63 | -1078.557421 | 0.304072 | 0.234621 | -1078.610332 | -138.9 | -1078.306260 | -1078.372699 | 19.5 | -1072.326061 | -4.603780 | -1076.745119 |
| conformer_70 | -1078.557813 | 0.304210 | 0.234018 | -1078.610028 | -137.1 | -1078.305818 | -1078.372998 | 18.8 | -1072.327225 | -4.602760 | -1076.745170 |
| conformer_42 | -1078.555002 | 0.304240 | 0.234818 | -1078.609576 | -143.3 | -1078.305336 | -1078.371746 | 22.0 | -1072.321307 | -4.605131 | -1076.743182 |
| conformer_81 | -1078.556617 | 0.304104 | 0.234820 | -1078.609503 | -138.9 | -1078.305399 | -1078.371671 | 22.2 | -1072.325765 | -4.603423 | -1076.744242 |
| conformer_49 | -1078.559952 | 0.304307 | 0.235515 | -1078.609498 | -130.1 | -1078.305191 | -1078.370971 | 24.1 | -1072.327677 | -4.604471 | -1076.743167 |
| conformer_41 | -1078.557907 | 0.304128 | 0.234383 | -1078.609424 | -135.3 | -1078.305296 | -1078.372029 | 21.3 | -1072.325531 | -4.604248 | -1076.743901 |
| conformer_29 | -1078.555576 | 0.304135 | 0.234795 | -1078.609248 | -140.9 | -1078.305113 | -1078.371441 | 22.8 | -1072.321296 | -4.605535 | -1076.742696 |
| conformer_22 | -1078.556078 | 0.304281 | 0.235101 | -1078.608688 | -138.1 | -1078.304407 | -1078.370575 | 25.1 | -1072.319039 | -4.608944 | -1076.742479 |
| conformer_61 | -1078.560144 | 0.304301 | 0.235492 | -1078.608312 | -126.5 | -1078.304011 | -1078.369808 | 27.1 | -1072.328806 | -4.604078 | -1076.742548 |
| conformer_30 | -1078.556065 | 0.304409 | 0.237709 | -1078.607426 | -134.8 | -1078.303017 | -1078.366705 | 35.3 | -1072.314684 | -4.612557 | -1076.737880 |
| conformer_47 | -1078.555683 | 0.304163 | 0.236827 | -1078.606833 | -134.3 | -1078.302670 | -1078.366994 | 34.5 | -1072.316647 | -4.610175 | -1076.738133 |
| conformer_18 | -1078.559110 | 0.304295 | 0.234620 | -1078.606445 | -124.3 | -1078.302150 | -1078.368813 | 29.7 | -1072.325942 | -4.605587 | -1076.741231 |
| conformer_76 | -1078.555408 | 0.304099 | 0.236422 | -1078.605598 | -131.8 | -1078.301499 | -1078.366164 | 36.7 | -1072.316638 | -4.610193 | -1076.737586 |
| conformer_21 | -1078.557937 | 0.304348 | 0.236630 | -1078.604732 | -122.9 | -1078.300384 | -1078.365090 | 39.5 | -1072.318777 | -4.610449 | -1076.736379 |
| conformer_26 | -1078.557099 | 0.304297 | 0.236904 | -1078.603188 | -121.0 | -1078.298891 | -1078.363272 | 44.3 | -1072.317796 | -4.610085 | -1076.734054 |
| conformer_17 | -1078.555665 | 0.304354 | 0.237284 | -1078.602870 | -123.9 | -1078.298516 | -1078.362574 | 46.1 | -1072.314318 | -4.611666 | -1076.732893 |

[a]: Single-point calculation in aqueous phase with SMD model.

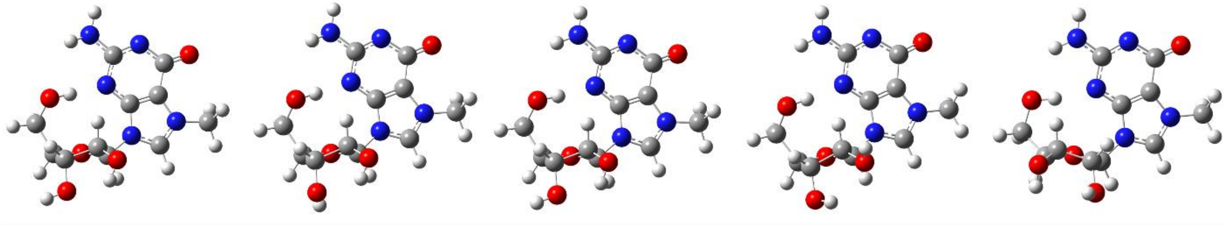

conformer_1 conformer_2 conformer_3 conformer_14 conformer_22

**Figure S56.** B3LYP-D3/def2-TZVPP optimized geometries of conformers for neutral $N^7$-methylguanosine (**9rb7mG**).

**Table S54.** Conformers of gas-phase optimized neutral $N^7$-methylguanosine (**9rb7mG**) at the B3LYP-D3/def2-TZVPP level of theory followed by aqueous phase single-point calculation. The columns display total energy without zero-point correction ($E_{Tot}$), Gibbs free energy (δG), total energy without zero-point correction ($E_{Tot,W}$), Gibbs free energy ($G_{298,W}$) in water (W), total single-point energy ($E_{CBS}$) calculated at DLPNO-CCSD(T)/CBS level of theory, and their corresponding free energy $G_{CBS}$. $G_{298,W}$ and $G_{CBS}$ have been corrected to the standard state of 1 mol/L by addition of +7.908 kJ/mol. $\Delta G_{Solv}$ represents the Gibbs free energy of solvation. The data are arranged in the ascending numeric order of $E_{Tot,W}$. $\Delta G_{298,W}$ represents the respective energy difference to the lowest structure. Only conformers within the 24 kJ/mol (6 kcal/mol) energy window above the lowest in CREST are included in initial conformer sampling. Duplicates of the same structure are excluded.

| | B3LYP-D3/def2-TZVPP | | | SMD(H2O)/B3LYP-D3/def2-TZVPP[a] | | | | | DLPNO-CCSD(T)/CBS | | |
|---|---|---|---|---|---|---|---|---|---|---|---|
| **9rb7mG No.** | **$E_{Tot}$ (Hartree)** | **δH (Hartree)** | **δG (Hartree)** | **$E_{Tot,W}$ (Hartree)** | **$\Delta G_{Solv}$ (kJ/mol)** | **$H_{298,W}$ (Hartree)** | **$G_{298,W}$ (Hartree)** | **$\Delta G_{298,W}$ (kJ/mol)** | **$E_{CBS,HF}$ (Hartree)** | **$E_{CBS,C}$ (Hartree)** | **$G_{CBS,W}$ (Hartree)** |
| conformer_14 | -1078.553451 | 0.304028 | 0.237119 | -1078.609466 | -147.1 | -1078.305438 | -1078.369335 | 2.0 | -1072.307898 | -4.615636 | -1076.739418 |
| **conformer_1** | **-1078.552889** | **0.303883** | **0.236283** | **-1078.609389** | **-148.3** | **-1078.305506** | **-1078.370094** | **0.0** | **-1072.307027** | **-4.615543** | **-1076.739775** |
| conformer_2 | -1078.550785 | 0.303874 | 0.236508 | -1078.609262 | -153.5 | -1078.305388 | -1078.369742 | 0.9 | -1072.302793 | -4.617061 | -1076.738813 |
| conformer_3 | -1078.552900 | 0.303928 | 0.237030 | -1078.609191 | -147.8 | -1078.305263 | -1078.369149 | 2.5 | -1072.306845 | -4.615733 | -1076.738827 |
| conformer_22 | -1078.547161 | 0.304108 | 0.236904 | -1078.604757 | -151.2 | -1078.300649 | -1078.364841 | 13.8 | -1072.300882 | -4.615759 | -1076.734320 |
| conformer_5 | -1078.551262 | 0.303839 | 0.235808 | -1078.604349 | -139.4 | -1078.300510 | -1078.365529 | 12.0 | -1072.308493 | -4.613804 | -1076.736564 |
| conformer_4 | -1078.549896 | 0.303919 | 0.236740 | -1078.604311 | -142.9 | -1078.300392 | -1078.364559 | 14.5 | -1072.303810 | -4.616296 | -1076.734770 |
| conformer_48 | -1078.545233 | 0.304020 | 0.236161 | -1078.603994 | -154.3 | -1078.299974 | -1078.364821 | 13.8 | -1072.300504 | -4.615444 | -1076.735535 |
| conformer_15 | -1078.544218 | 0.304019 | 0.235938 | -1078.603950 | -156.8 | -1078.299931 | -1078.365000 | 13.4 | -1072.297211 | -4.617062 | -1076.735055 |
| conformer_17 | -1078.545246 | 0.304023 | 0.237151 | -1078.603790 | -153.7 | -1078.299767 | -1078.363627 | 17.0 | -1072.300326 | -4.615571 | -1076.734277 |
| conformer_30 | -1078.544225 | 0.304022 | 0.236978 | -1078.603775 | -156.3 | -1078.299753 | -1078.363785 | 16.6 | -1072.297099 | -4.617195 | -1076.733855 |
| conformer_11 | -1078.550187 | 0.303792 | 0.235108 | -1078.603558 | -140.1 | -1078.299766 | -1078.365438 | 12.2 | -1072.308622 | -4.612656 | -1076.736529 |
| conformer_13 | -1078.549989 | 0.303743 | 0.235423 | -1078.603439 | -140.3 | -1078.299696 | -1078.365004 | 13.4 | -1072.308151 | -4.612771 | -1076.735937 |
| conformer_9 | -1078.549338 | 0.303829 | 0.235608 | -1078.603226 | -141.5 | -1078.299397 | -1078.364606 | 14.4 | -1072.307716 | -4.612866 | -1076.735850 |
| conformer_10 | -1078.548311 | 0.303821 | 0.235291 | -1078.602968 | -143.5 | -1078.299147 | -1078.364665 | 14.3 | -1072.307927 | -4.611693 | -1076.735973 |
| conformer_8 | -1078.548505 | 0.303883 | 0.235256 | -1078.602960 | -143.0 | -1078.299077 | -1078.364692 | 14.2 | -1072.308304 | -4.611642 | -1076.736133 |
| conformer_7 | -1078.548087 | 0.303797 | 0.235281 | -1078.602860 | -143.8 | -1078.299063 | -1078.364567 | 14.5 | -1072.307167 | -4.612015 | -1076.735662 |
| conformer_6 | -1078.548751 | 0.303868 | 0.235542 | -1078.602678 | -141.6 | -1078.298810 | -1078.364124 | 15.7 | -1072.308117 | -4.611794 | -1076.735285 |
| conformer_12 | -1078.551077 | 0.303840 | 0.235070 | -1078.602607 | -135.3 | -1078.298767 | -1078.364525 | 14.6 | -1072.310565 | -4.612238 | -1076.736251 |
| conformer_58 | -1078.544355 | 0.304052 | 0.235458 | -1078.602443 | -152.5 | -1078.298391 | -1078.363973 | 16.1 | -1072.307166 | -4.608843 | -1076.735627 |
| conformer_63 | -1078.542164 | 0.304161 | 0.235385 | -1078.602420 | -158.2 | -1078.298259 | -1078.364023 | 15.9 | -1072.303599 | -4.609928 | -1076.735386 |
| conformer_71 | -1078.544119 | 0.304003 | 0.235409 | -1078.602400 | -153.0 | -1078.298397 | -1078.363979 | 16.1 | -1072.306858 | -4.608874 | -1076.735592 |
| conformer_38 | -1078.547489 | 0.303754 | 0.235412 | -1078.602316 | -143.9 | -1078.298562 | -1078.363892 | 16.3 | -1072.307846 | -4.611386 | -1076.735635 |
| conformer_74 | -1078.541924 | 0.304086 | 0.235270 | -1078.602274 | -158.4 | -1078.298188 | -1078.363992 | 16.0 | -1072.303273 | -4.609958 | -1076.735299 |
| conformer_18 | -1078.548143 | 0.303804 | 0.235312 | -1078.601575 | -140.3 | -1078.297771 | -1078.363251 | 18.0 | -1072.308234 | -4.611434 | -1076.734776 |
| conformer_55 | -1078.539454 | 0.304031 | 0.234354 | -1078.599739 | -158.3 | -1078.295708 | -1078.362373 | 20.3 | -1072.300128 | -4.610531 | -1076.733579 |
| conformer_44 | -1078.540702 | 0.303889 | 0.233351 | -1078.599352 | -154.0 | -1078.295463 | -1078.362989 | 18.7 | -1072.301427 | -4.611010 | -1076.734724 |
| conformer_41 | -1078.541052 | 0.303945 | 0.234732 | -1078.599151 | -152.5 | -1078.295206 | -1078.361407 | 22.8 | -1072.301518 | -4.611297 | -1076.733169 |
| conformer_53 | -1078.538345 | 0.304010 | 0.233922 | -1078.598907 | -159.0 | -1078.294897 | -1078.361973 | 21.3 | -1072.299208 | -4.610254 | -1076.733090 |

| | | | | | | | | | | | |
|---|---|---|---|---|---|---|---|---|---|---|---|
| conformer_66 | -1078.537102 | 0.303635 | 0.232886 | -1078.598660 | -161.6 | -1078.295025 | -1078.362762 | 19.2 | -1072.298549 | -4.610467 | -1076.734676 |
| conformer_37 | -1078.538262 | 0.303684 | 0.234354 | -1078.598631 | -158.5 | -1078.294947 | -1078.361265 | 23.2 | -1072.299481 | -4.610467 | -1076.732952 |
| conformer_69 | -1078.537097 | 0.303634 | 0.232426 | -1078.598618 | -161.5 | -1078.294984 | -1078.363180 | 18.2 | -1072.298079 | -4.610740 | -1076.734903 |
| conformer_65 | -1078.535529 | 0.303825 | 0.234795 | -1078.598575 | -165.5 | -1078.294750 | -1078.360768 | 24.5 | -1072.295029 | -4.611398 | -1076.731665 |
| conformer_39 | -1078.538398 | 0.303736 | 0.234493 | -1078.598365 | -157.4 | -1078.294629 | -1078.360860 | 24.2 | -1072.299694 | -4.610434 | -1076.732590 |
| conformer_20 | -1078.536162 | 0.303753 | 0.234734 | -1078.598132 | -162.7 | -1078.294379 | -1078.360386 | 25.5 | -1072.294307 | -4.612807 | -1076.731338 |
| conformer_21 | -1078.536313 | 0.303786 | 0.234768 | -1078.598010 | -162.0 | -1078.294224 | -1078.360230 | 25.9 | -1072.294637 | -4.612623 | -1076.731177 |
| conformer_57 | -1078.535155 | 0.303636 | 0.233470 | -1078.597740 | -164.3 | -1078.294104 | -1078.361258 | 23.2 | -1072.293935 | -4.612360 | -1076.732399 |
| conformer_64 | -1078.538565 | 0.304002 | 0.234370 | -1078.597680 | -155.2 | -1078.293678 | -1078.360298 | 25.7 | -1072.299015 | -4.611064 | -1076.731812 |
| conformer_24 | -1078.539282 | 0.303924 | 0.234976 | -1078.597184 | -152.0 | -1078.293260 | -1078.359196 | 28.6 | -1072.301445 | -4.609885 | -1076.731245 |
| conformer_23 | -1078.537728 | 0.303928 | 0.235967 | -1078.596805 | -155.1 | -1078.292877 | -1078.357826 | 32.2 | -1072.291513 | -4.617416 | -1076.729027 |
| conformer_31 | -1078.537501 | 0.303839 | 0.235741 | -1078.596800 | -155.7 | -1078.292961 | -1078.358047 | 31.6 | -1072.291397 | -4.617246 | -1076.729188 |
| conformer_42 | -1078.541035 | 0.303922 | 0.233865 | -1078.596799 | -146.4 | -1078.292877 | -1078.359922 | 26.7 | -1072.301619 | -4.611069 | -1076.731575 |
| conformer_76 | -1078.535339 | 0.303835 | 0.234765 | -1078.596720 | -161.2 | -1078.292885 | -1078.358943 | 29.3 | -1072.290747 | -4.615813 | -1076.730164 |
| conformer_62 | -1078.541033 | 0.303921 | 0.234482 | -1078.596601 | -145.9 | -1078.292680 | -1078.359107 | 28.8 | -1072.301461 | -4.611224 | -1076.730758 |
| conformer_29 | -1078.540992 | 0.303876 | 0.234687 | -1078.596541 | -145.8 | -1078.292665 | -1078.358842 | 29.5 | -1072.301342 | -4.611238 | -1076.730431 |
| conformer_26 | -1078.536741 | 0.303836 | 0.234484 | -1078.596397 | -156.6 | -1078.292561 | -1078.358901 | 29.4 | -1072.296112 | -4.612163 | -1076.730436 |
| conformer_25 | -1078.542065 | 0.304046 | 0.234955 | -1078.595702 | -140.8 | -1078.291656 | -1078.357735 | 32.4 | -1072.303489 | -4.610728 | -1076.729888 |
| conformer_40 | -1078.533869 | 0.303801 | 0.233145 | -1078.594832 | -160.1 | -1078.291031 | -1078.358675 | 30.0 | -1072.290660 | -4.614429 | -1076.729895 |
| conformer_45 | -1078.533877 | 0.303801 | 0.233706 | -1078.594575 | -159.4 | -1078.290774 | -1078.357857 | 32.1 | -1072.290621 | -4.614464 | -1076.729064 |

[a]: Single-point calculation in aqueous phase with SMD model.

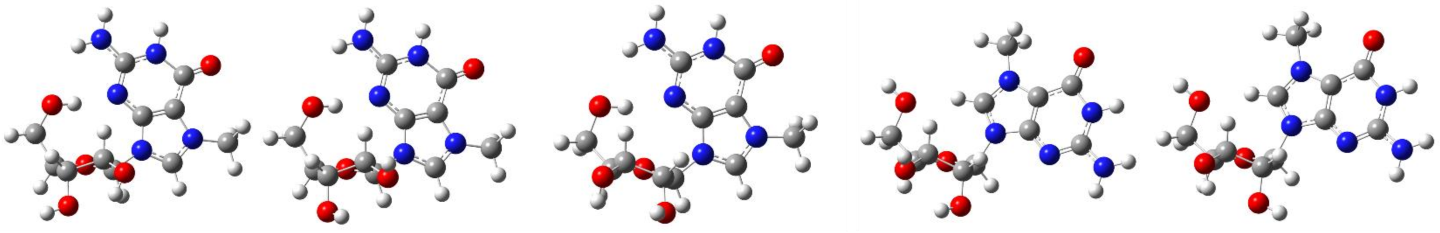

conformer_1 conformer_11 conformer_21 conformer_31 conformer_32

**Figure S57.** B3LYP-D3/def2-TZVPP optimized geometries of conformers for protonated $N^7$-methylguanosine cation (**9rb7mG⁺**).

**Table S55.** Conformers of gas-phase optimized protonated $N^7$-methylguanosine cation (**9rb7mG⁺**) at the B3LYP-D3/def2-TZVPP level of theory followed by aqueous phase single-point calculation. The columns display total energy without zero-point correction ($E_{Tot}$), Gibbs free energy (δG), total energy without zero-point correction ($E_{Tot,W}$), Gibbs free energy ($G_{298,W}$) in water (W), total single-point energy ($E_{CBS}$) calculated at DLPNO-CCSD(T)/CBS level of theory, and their corresponding free energy $G_{CBS}$. $G_{298,W}$ and $G_{CBS}$ have been corrected to the standard state of 1 mol/L by addition of +7.908 kJ/mol. $\Delta G_{Solv}$ represents the Gibbs free energy of solvation. The data are arranged in the ascending numeric order of $E_{Tot,W}$. $\Delta G_{298,W}$ represents the respective energy difference to the lowest structure. Only conformers within the 24 kJ/mol (6 kcal/mol) energy window above the lowest in CREST are included in initial conformer sampling. Duplicates of the same structure are excluded.

| | B3LYP-D3/def2-TZVPP | | | SMD(H₂O)/B3LYP-D3/def2-TZVPP[a] | | | | | DLPNO-CCSD(T)/CBS | | |
|---|---|---|---|---|---|---|---|---|---|---|---|
| **9rb7mG No.** | **$E_{Tot}$** (Hartree) | **δH** (Hartree) | **δG** (Hartree) | **$E_{Tot,W}$** (Hartree) | **$\Delta G_{Solv}$** (kJ/mol) | **$H_{298,W}$** (Hartree) | **$G_{298,W}$** (Hartree) | **$\Delta G_{298,W}$** (kJ/mol) | **$E_{CBS,HF}$** (Hartree) | **$E_{CBS,C}$** (Hartree) | **$G_{CBS,W}$** (Hartree) |
| conformer_11 | -1078.966164 | 0.317890 | 0.250949 | -1079.075998 | -288.4 | -1078.758108 | -1078.822037 | 0.8 | -1072.733128 | -4.598454 | -1077.187455 |
| **conformer_1** | **-1078.968371** | **0.317803** | **0.250546** | **-1079.075891** | **-282.3** | **-1078.758088** | **-1078.822333** | **0.0** | **-1072.734998** | **-4.598637** | **-1077.187598** |
| conformer_32 | -1078.967500 | 0.317272 | 0.247911 | -1079.071798 | -273.8 | -1078.754526 | -1078.820875 | 3.8 | -1072.741061 | -4.592994 | -1077.187431 |
| conformer_31 | -1078.968471 | 0.317259 | 0.247860 | -1079.071702 | -271.0 | -1078.754443 | -1078.820830 | 3.9 | -1072.741753 | -4.593254 | -1077.187366 |
| conformer_21 | -1078.960692 | 0.317758 | 0.250045 | -1079.071691 | -291.4 | -1078.753933 | -1078.818634 | 9.7 | -1072.727118 | -4.598687 | -1077.183747 |
| conformer_23 | -1078.964037 | 0.317291 | 0.248137 | -1079.071254 | -281.5 | -1078.753963 | -1078.820105 | 5.9 | -1072.735430 | -4.594398 | -1077.185895 |
| conformer_63 | -1078.951877 | 0.317043 | 0.247025 | -1079.070948 | -312.6 | -1078.753905 | -1078.820911 | 3.7 | -1072.723320 | -4.594101 | -1077.186454 |
| conformer_13 | -1078.962993 | 0.317604 | 0.249855 | -1079.070929 | -283.4 | -1078.753325 | -1078.818062 | 11.2 | -1072.729337 | -4.599183 | -1077.183589 |
| conformer_26 | -1078.964518 | 0.317316 | 0.247723 | -1079.070862 | -279.2 | -1078.753546 | -1078.820127 | 5.8 | -1072.735993 | -4.593919 | -1077.185521 |
| conformer_33 | -1078.968139 | 0.317331 | 0.247789 | -1079.070809 | -269.6 | -1078.753478 | -1078.820008 | 6.1 | -1072.741619 | -4.592602 | -1077.186090 |
| conformer_5 | -1078.967085 | 0.317455 | 0.248506 | -1079.070458 | -271.4 | -1078.753003 | -1078.818940 | 8.9 | -1072.738850 | -4.594882 | -1077.185587 |
| conformer_40 | -1078.956738 | 0.317296 | 0.247846 | -1079.070306 | -298.2 | -1078.753010 | -1078.819448 | 7.6 | -1072.728005 | -4.594962 | -1077.185678 |
| conformer_22 | -1078.955386 | 0.317496 | 0.250047 | -1079.070300 | -301.7 | -1078.752804 | -1078.817241 | 13.4 | -1072.719561 | -4.600399 | -1077.181815 |
| conformer_29 | -1078.962401 | 0.317573 | 0.250238 | -1079.070155 | -282.9 | -1078.752582 | -1078.816905 | 14.3 | -1072.729420 | -4.598684 | -1077.182608 |
| conformer_50 | -1078.959494 | 0.317456 | 0.250103 | -1079.069826 | -289.7 | -1078.752370 | -1078.816711 | 14.8 | -1072.726788 | -4.598565 | -1077.182571 |
| conformer_64 | -1078.956902 | 0.317216 | 0.247884 | -1079.069675 | -296.1 | -1078.752459 | -1078.818779 | 9.3 | -1072.728821 | -4.594233 | -1077.184931 |
| conformer_48 | -1078.959152 | 0.317363 | 0.248310 | -1079.069446 | -289.6 | -1078.752083 | -1078.818124 | 11.1 | -1072.731388 | -4.594009 | -1077.184369 |
| conformer_61 | -1078.955278 | 0.317272 | 0.247202 | -1079.068954 | -298.5 | -1078.751682 | -1078.818740 | 9.4 | -1072.728863 | -4.592457 | -1077.184782 |
| conformer_62 | -1078.952013 | 0.317238 | 0.247575 | -1079.068781 | -306.6 | -1078.751543 | -1078.818194 | 10.9 | -1072.723063 | -4.594225 | -1077.183470 |
| conformer_53 | -1078.959164 | 0.317336 | 0.247474 | -1079.067942 | -285.6 | -1078.750606 | -1078.817456 | 12.8 | -1072.732485 | -4.593218 | -1077.183995 |
| conformer_87 | -1078.948536 | 0.317302 | 0.247398 | -1079.066123 | -308.7 | -1078.748821 | -1078.815713 | 17.4 | -1072.719021 | -4.595025 | -1077.181223 |
| conformer_82 | -1078.959550 | 0.317340 | 0.247392 | -1079.065050 | -277.0 | -1078.747710 | -1078.814646 | 20.2 | -1072.733618 | -4.593145 | -1077.181858 |

[a]: Single-point calculation in aqueous phase with SMD model.

### S3 Results and Discussion

#### S3.1 Four Reaction Categories

The conformers with the lowest $G_{298,W}$ were taken for each species for the further calculation of thermodynamic driving force of natural decomposition in water. The energies of other basic molecules without conformers were taken directly for the calculation. Based on different types of decomposition, the reactions are divided into four categories: deglycosylation, deamination, deacetylation, desulfurization.

#### S3.1.1Deglycosylation

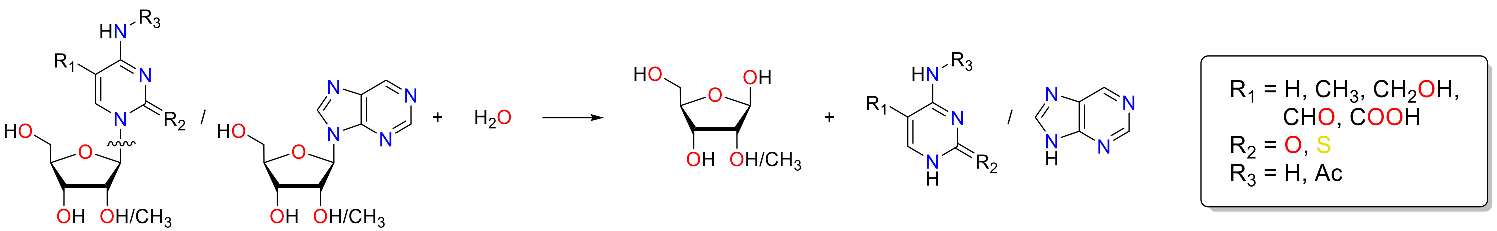

**Figure S58.** Deglycosylation reaction of modified cytidines and uridines.

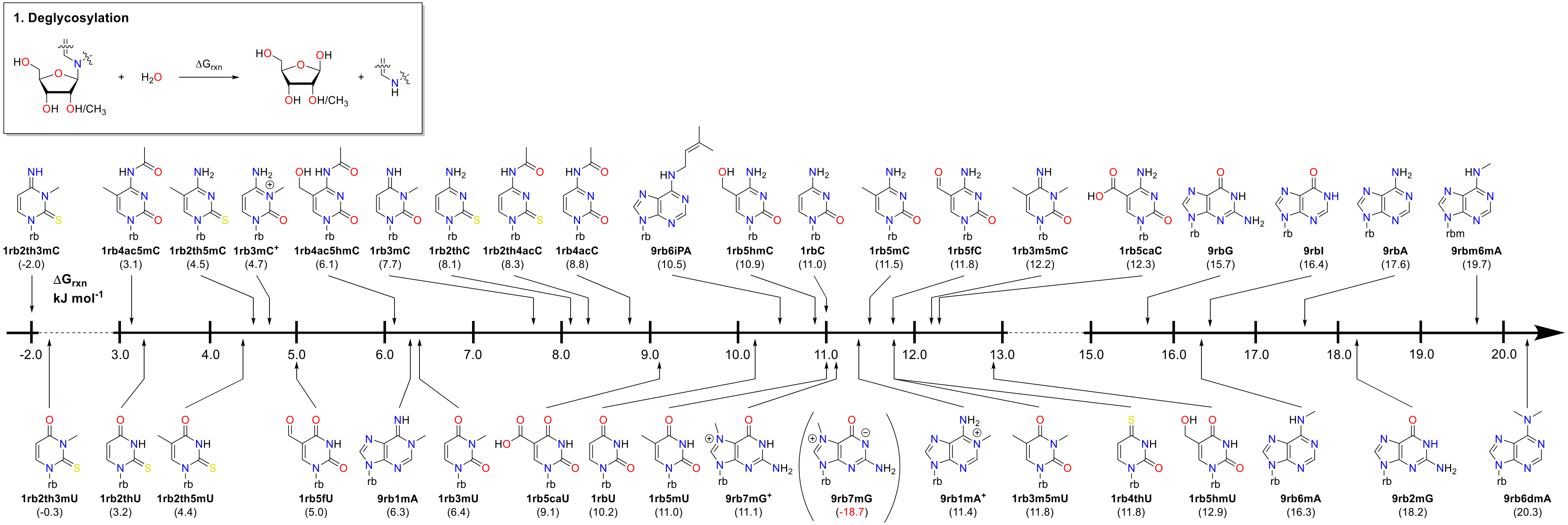

**Figure S59.** Reaction energy comparison for deglycosylation of different nucleosides calculated at DLPNO-CCSD(T)/CBS//SMD($H_2O$)/B3LYP-D3/def2-TZVPP level of theory.

**Table S56.** Energetics for the deglycosylation. $\Delta E_{Rxn,gas}$, $\Delta G_{Rxn,gas}$, and $\Delta G_{Rxn}$ represent the total energy difference in the gas phase and the Gibbs free difference in both gas and aqueous phase for the deglycosylation reaction.

| Reactant | Product | $\Delta E_{Rxn,gas}$ | $\Delta G_{Rxn,gas}$ | $\Delta G_{Rxn}$ |
|---|---|---|---|---|
| Nucleotide | Nucleobase | (kJ/mol) | (kJ/mol) | (kJ/mol) |
| **Depyrimidination** | | | | |
| 1rbC | C | +36.9 | +29.8 | +11.0 |
| 1rb2thC | 2thC | +30.5 | +25.9 | +8.1 |
| 1rb2th3mC | 2th3mC | -8.1 | -11.9 | -2.0 |
| 1rb2th4acC | 2th4acC | +29.6 | +24.2 | +8.3 |
| 1rb2th5mC | 2th5mC | +25.6 | +18.3 | +4.5 |
| 1rb3mC | 3mC | +21.5 | +12.7 | +7.7 |
| 1rb3mC$^+$ | 3mC$^+$ | +19.4 | +16.4 | +4.7 |
| 1rb3m5mC | 3m5mC | +5.7 | +5.3 | +12.2 |
| 1rb4acC | 4acC | +35.8 | +26.8 | +8.8 |
| 1rb4ac5mC | 4ac5mC | +67.4 | +51.3 | +3.1 |
| 1rb4ac5hmC | 4ac5hmC | +58.8 | +47.7 | +6.1 |
| 1rb5mC | 5mC | +38.4 | +31.1 | +11.5 |
| 1rb5hmC | 5hmC | +37.8 | +30.5 | +10.9 |
| 1rb5fC | 5fC | +15.7 | +14.8 | +11.8 |
| 1rb5caC | 5caC | +37.9 | +30.6 | +12.3 |
| 1rbU | U | +9.0 | +5.4 | +10.2 |
| 1rb2thU | 2thU | -4.4 | -6.3 | +3.2 |
| 1rb2th3mU | 2th3mU | +17.1 | +10.8 | -0.3 |
| 1rb2th5mU | 2th5mU | -3.4 | -5.8 | +4.4 |
| 1rb3mU | 3mU | +21.7 | +12.6 | +6.4 |
| 1rb3m5mU | 3m5mU | +5.2 | +6.1 | +11.8 |
| 1rb4thU | 4thU | +10.0 | +7.8 | +11.8 |
| 1rb5mU | 5mU | +9.8 | +6.3 | +11.0 |
| 1rb5hmU | 5hmU | +9.0 | +8.3 | +12.9 |
| 1rb5fU | 5fU | +21.2 | +15.5 | +5.0 |
| 1rb5caU | 5caU | +14.6 | +10.4 | +9.1 |
| **Depurination** | | | | |
| 9rbA | A | +40.9 | +29.4 | +17.6 |
| 9rb1mA | 1mA | +66.0 | +54.5 | +6.3 |
| 9rb1mA$^+$ | 1mA$^+$ | +39.8 | +30.6 | +11.4 |
| 9rb6mA | 6mA | +41.6 | +28.1 | +16.3 |
| 9rb6dmA | 6dmA | +39.8 | +28.7 | +20.3 |
| 9rbm6mA | 6mA | +43.4 | +31.1 | +19.7 |
| 9rb6iPA | 6iPA | +29.6 | +19.2 | +10.5 |
| 9rbG | G | +33.2 | +21.5 | +15.7 |
| 9rb2mG | 2mG | +40.2 | +26.8 | +18.2 |
| 9rb7mG | 7mG | -24.6 | -33.3 | -18.7 |
| 9rb7mG$^+$ | 7mG$^+$ | +53.1 | +41.6 | +11.1 |
| 9rbI | I | +32.9 | +22.8 | +16.4 |

#### S3.1.2Deamination

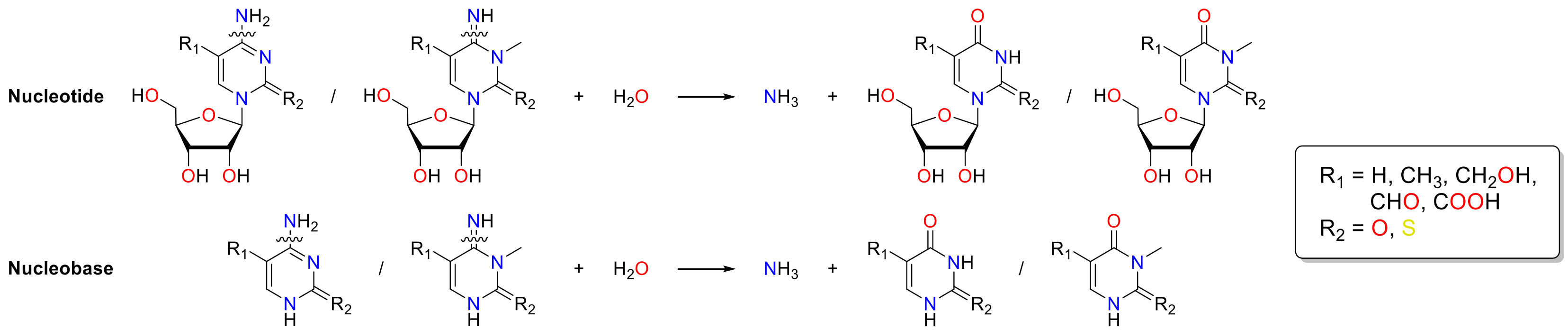

**Figure S60.** Deamination reaction of modified cytidines.

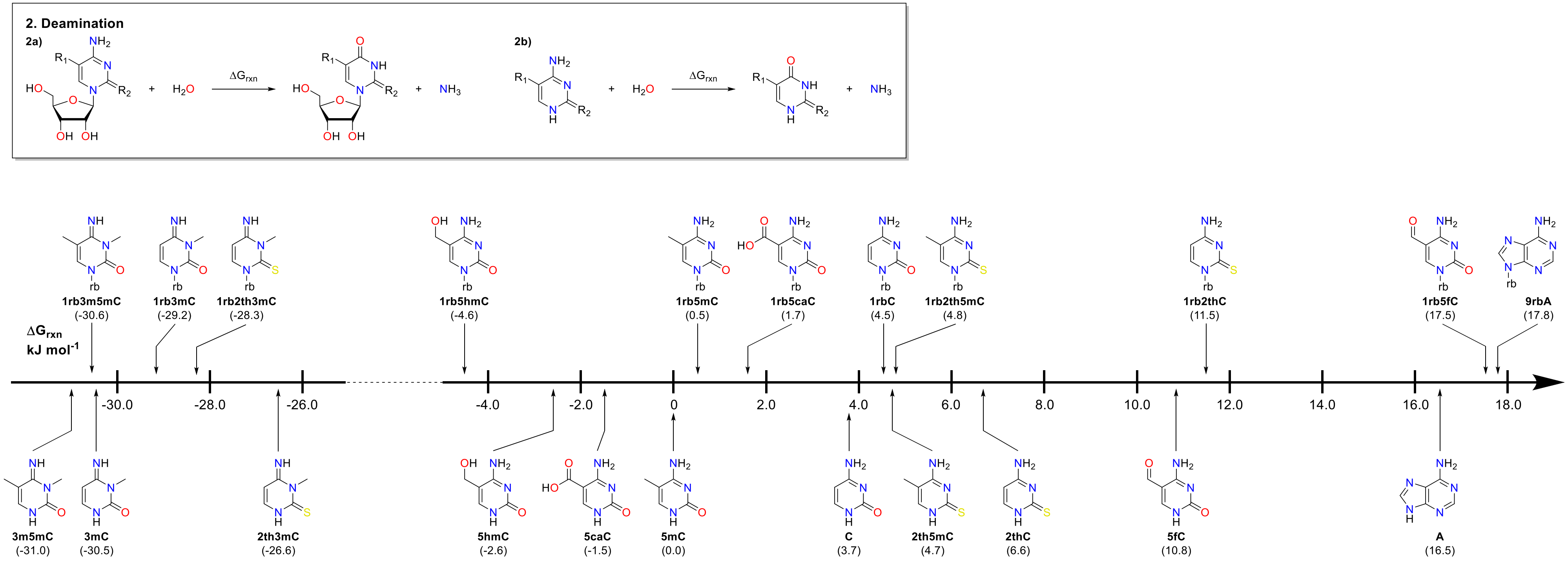

**Figure S61.** Reaction energy comparison for deamination of different nucleosides and their corresponding nucleobases calculated at DLPNO-CCSD(T)/CBS//SMD($H_2O$)/B3LYP-D3/def2-TZVPP level of theory.

**Table S57.** Energetics for the deamination. $\Delta E_{Rxn,gas}$, $\Delta G_{Rxn,gas}$, and $\Delta G_{Rxn}$ represent the total energy difference in the gas phase and the Gibbs free difference in both gas and aqueous phase for the deamination reaction.

| Nucleotide | | $\Delta E_{Rxn,gas}$ | $\Delta G_{Rxn,gas}$ | $\Delta G_{Rxn}$ |
|---|---|---|---|---|
| **Reactant** | **Product** | (kJ/mol) | (kJ/mol) | (kJ/mol) |
| **1rbC** | **1rbU** | -11.3 | -12.2 | +4.5 |
| **1rb2thC** | **1rb2thU** | -3.3 | -2.0 | +11.5 |
| **1rb2th5mC** | **1rb2th5mU** | -14 | -15.3 | +4.8 |
| **1rb5mC** | **1rb5mU** | -15.7 | -17.9 | +0.5 |
| **1rb5hmC** | **1rb5hmU** | -11.6 | -18.8 | -4.6 |
| **1rb5fC** | **1rb5fU** | -2.3 | -1.6 | +17.5 |
| **1rb5caC** | **1rb5caU** | -5.1 | -7.2 | +1.7 |
| **1rb2th3mC** | **1rb2th3mU** | -69.8 | -67.2 | -28.3 |
| **1rb3mC** | **1rb3mU** | -42.5 | -46.2 | -29.2 |
| **1rb3m5mC** | **1rb3m5mU** | -43.5 | -49.0 | -30.6 |
| **9rbA** | **9rbI** | +22.7 | +23.2 | +17.8 |
| **Nucleobase** | | $\Delta E_{Rxn,gas}$ | $\Delta G_{Rxn,gas}$ | $\Delta G_{Rxn}$ |
| **Reactant** | **Product** | (kJ/mol) | (kJ/mol) | (kJ/mol) |
| **C** | **U** | -39.2 | -36.6 | +3.7 |
| **2thC** | **2thU** | -38.2 | -34.2 | +6.6 |
| **2th5mC** | **2th5mU** | -43.0 | -39.4 | +4.7 |
| **5mC** | **5mU** | -44.2 | -42.8 | +0.1 |
| **5hmC** | **5hmU** | -40.4 | -41.1 | -2.6 |
| **5fC** | **5fU** | +3.2 | -0.8 | +10.8 |
| **5caC** | **5caU** | -28.4 | -27.3 | -1.5 |
| **2th3mC** | **2th3mU** | -44.6 | -44.5 | -26.6 |
| **3mC** | **3mU** | -42.3 | -46.4 | -30.5 |
| **3m5mC** | **3m5mU** | -44.0 | -48.2 | -31.0 |
| **A** | **I** | +14.6 | +16.6 | +16.5 |

##### S3.1.3Deacetylation

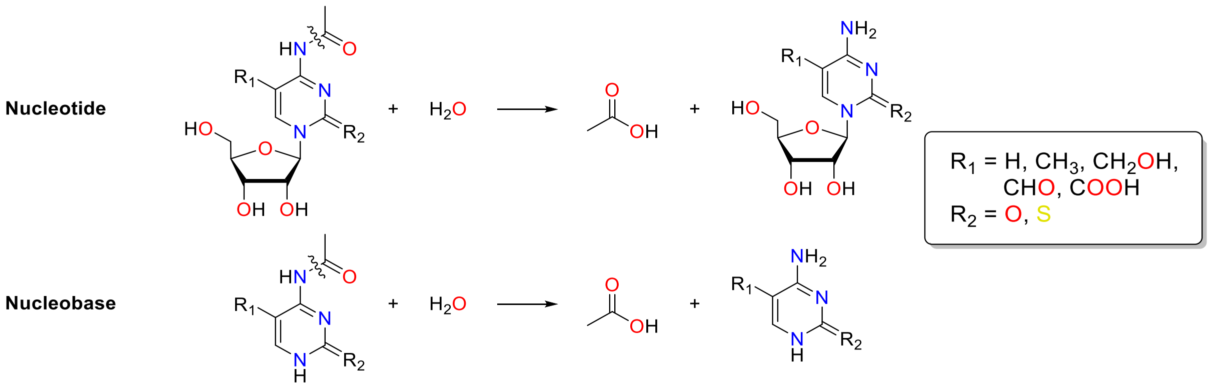

**Figure S62.** Deamination reaction of modified cytidine.

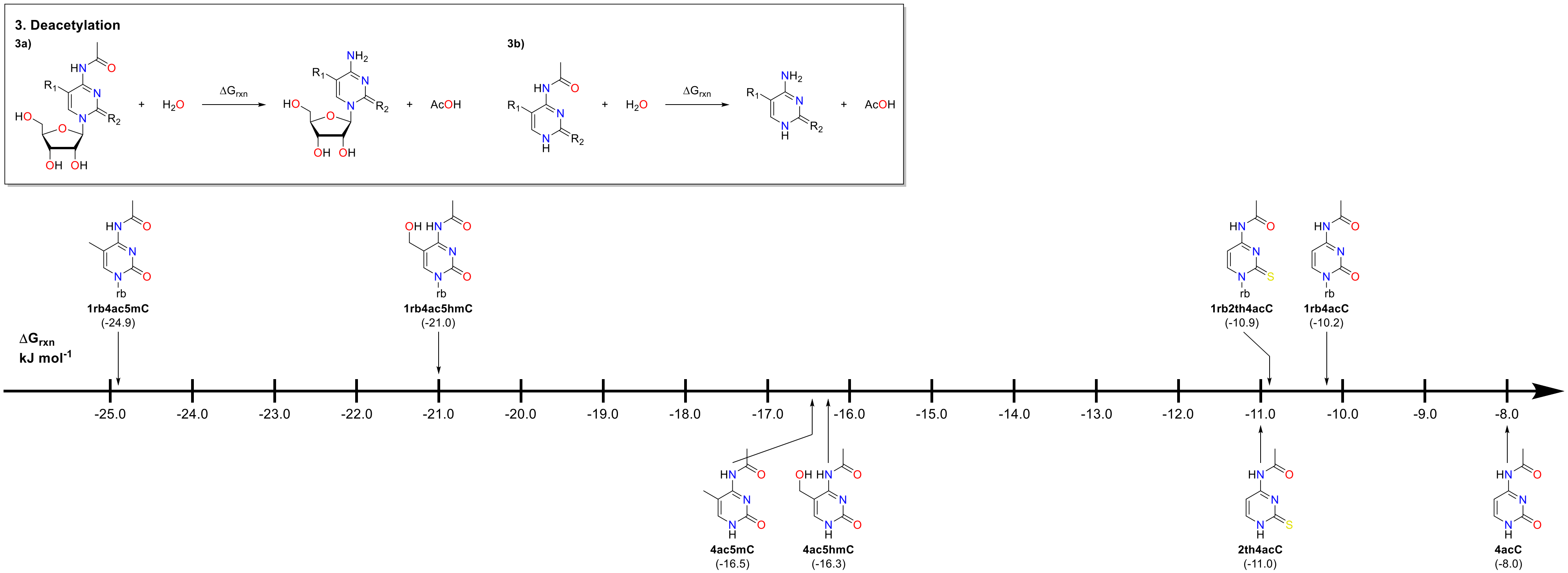

**Figure S63.** Reaction energy comparison for deacetylation of different nucleosides and their corresponding nucleobases calculated at DLPNO-CCSD(T)/CBS//SMD($H_2O$)/B3LYP-D3/def2-TZVPP level of theory.

**Table S58.** Energetics for the deacetylation. $\Delta E_{Rxn,gas}$, $\Delta G_{Rxn,gas}$, and $\Delta G_{Rxn}$ represent the total energy difference in the gas phase and the Gibbs free difference in both gas and aqueous phase for the deacetylation reaction.

| Nucleotide | | $\Delta E_{Rxn,gas}$ | $\Delta G_{Rxn,gas}$ | $\Delta G_{Rxn}$ |
|---|---|---|---|---|
| **Reactant** | **Product** | (kJ/mol) | (kJ/mol) | (kJ/mol) |
| **1rb2th4acC** | **1rb2thC** | -7.4 | -15.3 | -10.9 |
| **1rb4acC** | **1rbC** | -7.4 | -14.8 | -10.2 |
| **1rb4ac5mC** | **1rb5mC** | -18.7 | -27.6 | -24.9 |
| **1rb4ac5hmC** | **1rb5hmC** | -23.9 | -28.5 | -21.0 |
| **Nucleobase** | | $\Delta E_{Rxn,gas}$ | $\Delta G_{Rxn,gas}$ | $\Delta G_{Rxn}$ |
| **Reactant** | **Product** | (kJ/mol) | (kJ/mol) | (kJ/mol) |
| **2th4acC** | **2thC** | -6.5 | -13.6 | -11.0 |
| **4acC** | **C** | -6.2 | -11.7 | -8.0 |
| **4ac5mC** | **5mC** | -47.7 | -47.8 | -16.5 |
| **4ac5hmC** | **5hmC** | -44.9 | -45.7 | -16.3 |

#### S3.1.4Desulfurization

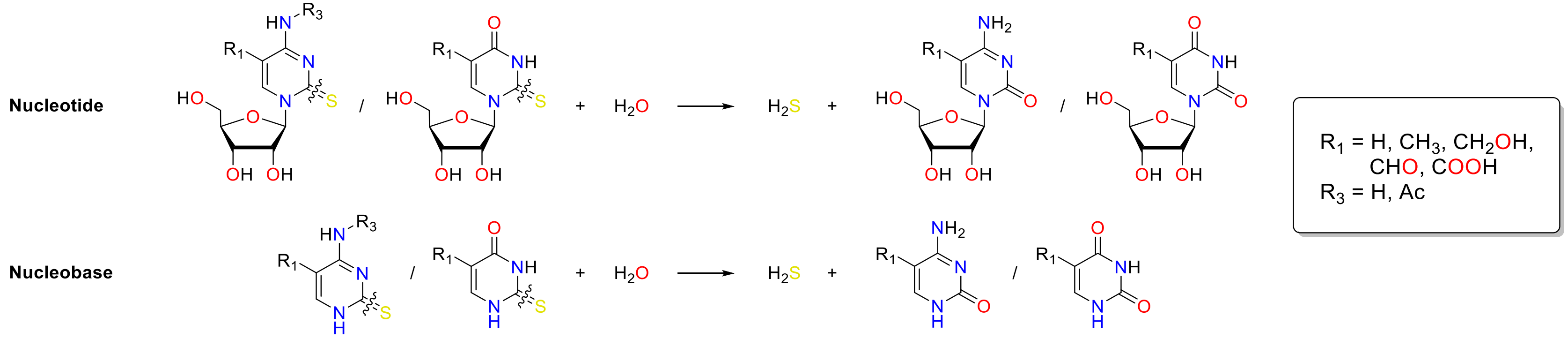

**Figure S64.** Desulfurization reaction of modified cytidines and uridines.

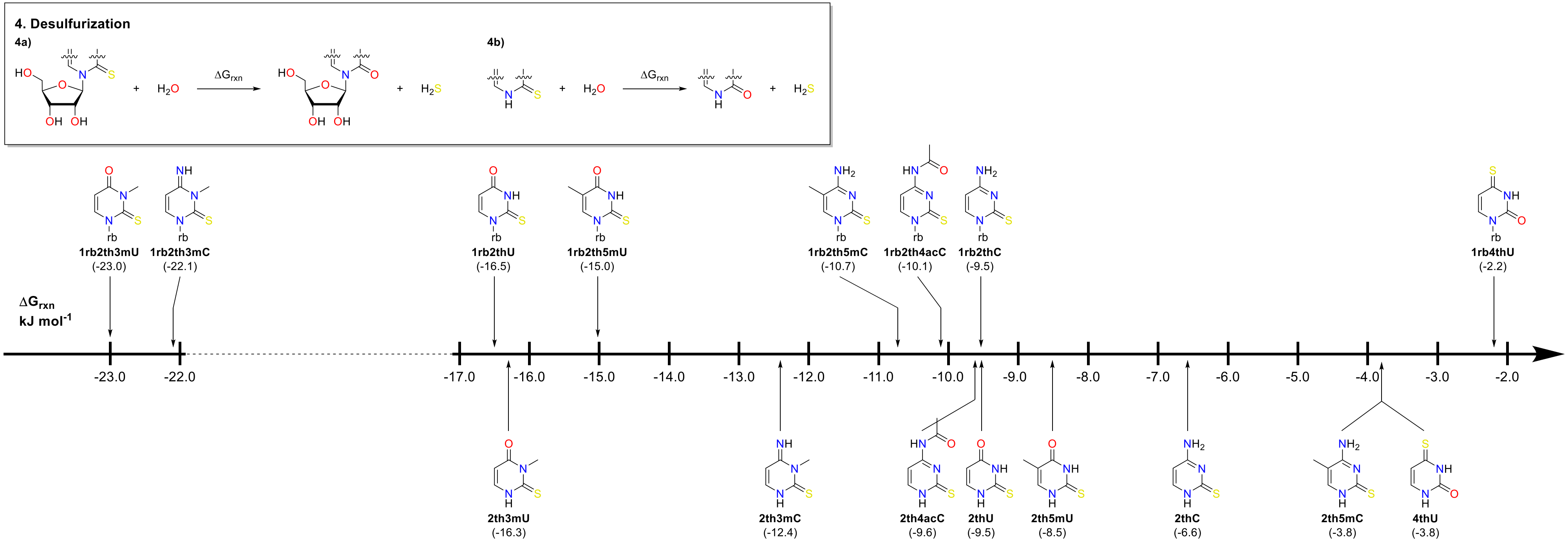

**Figure S65.** Reaction energy comparison for desulfurization of different nucleosides and their corresponding nucleobases calculated at DLPNO-CCSD(T)/CBS//SMD($H_2O$)/B3LYP-D3/def2-TZVPP level of theory.

**Table S59.** Energetics for the desulfurization. $\Delta E_{Rxn,gas}$, $\Delta G_{Rxn,gas}$, and $\Delta G_{Rxn}$ represent the total energy difference in the gas phase and the Gibbs free difference in both gas and aqueous phase for the desulfurization reaction.

| Nucleotide | | $\Delta E_{Rxn,gas}$ | $\Delta G_{Rxn,gas}$ | $\Delta G_{Rxn}$ |
|---|---|---|---|---|
| **Reactant** | **Product** | (kJ/mol) | (kJ/mol) | (kJ/mol) |
| **1rb2thC** | **1rbC** | -24.3 | -33.8 | -9.5 |
| **1rb2th3mC** | **1rb3mC** | -51.9 | -58.8 | -22.1 |
| **1rb2th4acC** | **1rb4acC** | -24.3 | -34.4 | -10.1 |
| **1rb2th5mC** | **1rb5mC** | -28.7 | -40.4 | -10.7 |
| **1rb2thU** | **1rbU** | -32.3 | -44.0 | -16.5 |
| **1rb2th3mU** | **1rb3mU** | -24.6 | -37.9 | -23.0 |
| **1rb2th5mU** | **1rb5mU** | -30.4 | -43.0 | -15.0 |
| **Nucleobase** | | $\Delta E_{Rxn,gas}$ | $\Delta G_{Rxn,gas}$ | $\Delta G_{Rxn}$ |
| **Reactant** | **Product** | (kJ/mol) | (kJ/mol) | (kJ/mol) |
| **2thC** | **C** | -17.9 | -29.9 | -6.6 |
| **2th3mC** | **3mC** | -22.3 | -34.3 | -12.4 |
| **2th4acC** | **4acC** | -18.2 | -31.8 | -9.6 |
| **2th5mC** | **5mC** | -16.0 | -27.6 | -3.8 |
| **2thU** | **U** | -18.9 | -32.4 | -9.5 |
| **2th3mU** | **3mu** | -20.0 | -36.1 | -16.3 |
| **2th5mU** | **5mU** | -17.2 | -30.9 | -8.5 |

#### S3.2 Corresponding Computational Results to Experimental Findings

##### S3.2.1Degradation of 4-thiouridine

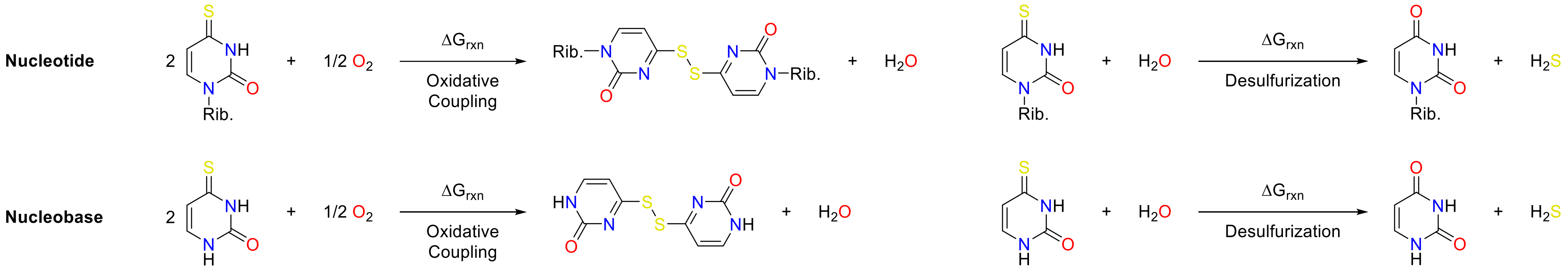

**Figure S66.** Reaction equation for the degradation of 4-thiouridine (**s⁴U**).

**Table S60.** Energetics for the degradation of 4-thiouridine (**s⁴U**). $\Delta E_{Rxn,gas}$, $\Delta G_{Rxn,gas}$, and $\Delta G_{Rxn}$ represent the total energy difference in the gas phase and the Gibbs free difference in both gas and aqueous phase for the deglycosylation reaction.

| **Nucleotide** | | | $\Delta E_{Rxn,gas}$ | $\Delta G_{Rxn,gas}$ | $\Delta G_{Rxn}$ |
|---|---|---|---|---|---|
| **Reactant** | **Product** | **Reaction Type** | (kJ/mol) | (kJ/mol) | (kJ/mol) |
| **s⁴U** | **s⁴U_dimer** | Oxidative Coupling | -215.0 | -154.4 | -121.6 |
| **s⁴U** | **U** | Desulfurization | -11.7 | -24.0 | -2.2 |
| **Nucleobase** | | | $\Delta E_{Rxn,gas}$ | $\Delta G_{Rxn,gas}$ | $\Delta G_{Rxn}$ |
| **Reactant** | **Product** | **Reaction Type** | (kJ/mol) | (kJ/mol) | (kJ/mol) |
| **s⁴U** | **s⁴U_dimer** | Oxidative Coupling | -108.0 | -90.9 | -133.4 |
| **s⁴U** | **U** | Desulfurization | -12.8 | -26.4 | -3.8 |

#### S3.2.2Degradation of 3-methylcytidine

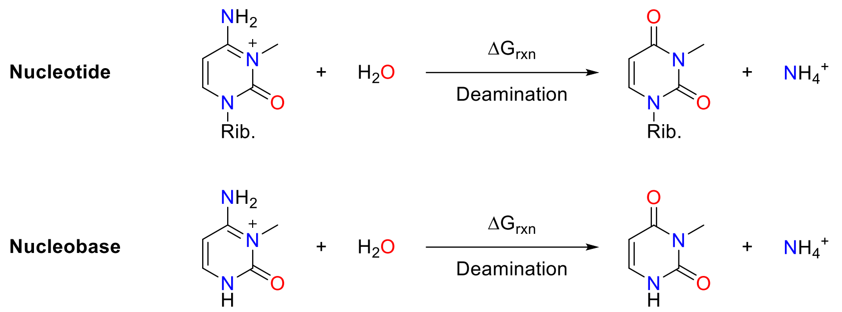

**Figure S67.** Reaction equation for the degradation of 3-methylcytidine (**m³C**).

**Table S61.** Energetics for the degradation of 3-methylcytidine (**m³C**). $\Delta E_{Rxn,gas}$, $\Delta G_{Rxn,gas}$, and $\Delta G_{Rxn}$ represent the total energy difference in the gas phase and the Gibbs free difference in both gas and aqueous phase for the deglycosylation reaction.

| Nucleotide | | | $\Delta E_{Rxn,gas}$ | $\Delta G_{Rxn,gas}$ | $\Delta G_{Rxn}$ |
|---|---|---|---|---|---|
| **Reactant** | **Product** | **Reaction Type** | (kJ/mol) | (kJ/mol) | (kJ/mol) |
| **m³C** | **m³U** | Deamination | 71.7 | 82.7 | -21.3 |
| **Nucleobase** | | | $\Delta E_{Rxn,gas}$ | $\Delta G_{Rxn,gas}$ | $\Delta G_{Rxn}$ |
| **Reactant** | **Product** | **Reaction Type** | (kJ/mol) | (kJ/mol) | (kJ/mol) |
| **m³C** | **m³U** | Deamination | 74.1 | 78.9 | -19.5 |

#### S3.2.3Degradation of $N^6$-isoprenyladenosine

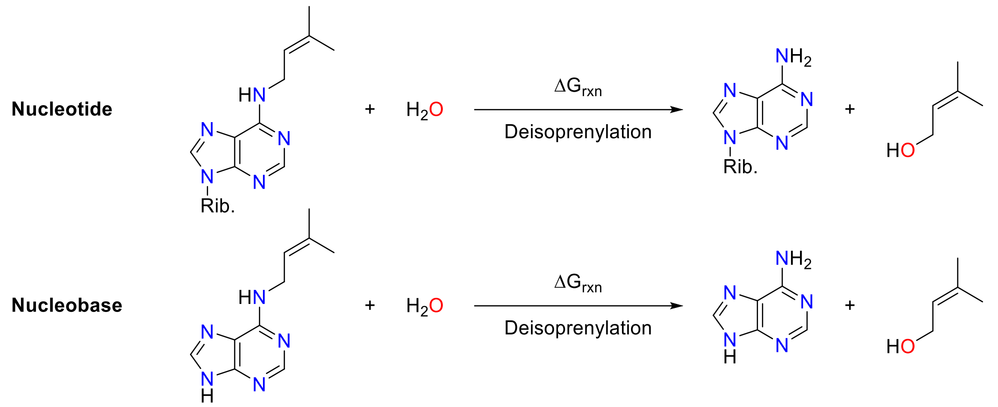

**Figure S68.** Reaction equation for the degradation of $N^6$-isoprenyladenosine (**i⁶A**).

**Table S62.** Energetics for the degradation of $N^6$-isoprenyladenosine (**i⁶A**). $\Delta E_{Rxn,gas}$, $\Delta G_{Rxn,gas}$, and $\Delta G_{Rxn}$ represent the total energy difference in the gas phase and the Gibbs free difference in both gas and aqueous phase for the deglycosylation reaction.

| **Nucleotide** | | | **$\Delta E_{Rxn,gas}$** | **$\Delta G_{Rxn,gas}$** | **$\Delta G_{Rxn}$** |
|---|---|---|---|---|---|
| **Reactant** | **Product** | **Reaction Type** | (kJ/mol) | (kJ/mol) | (kJ/mol) |
| **i⁶A** | **A** | Deisoprenylation | 19.5 | 16.0 | 24.5 |
| **Nucleobase** | | | **$\Delta E_{Rxn,gas}$** | **$\Delta G_{Rxn,gas}$** | **$\Delta G_{Rxn}$** |
| **Reactant** | **Product** | **Reaction Type** | (kJ/mol) | (kJ/mol) | (kJ/mol) |
| **i⁶A** | **A** | Deisoprenylation | 30.8 | 26.3 | 31.6 |

##### S3.2.4Degradation of *N*4-acetylcytidine

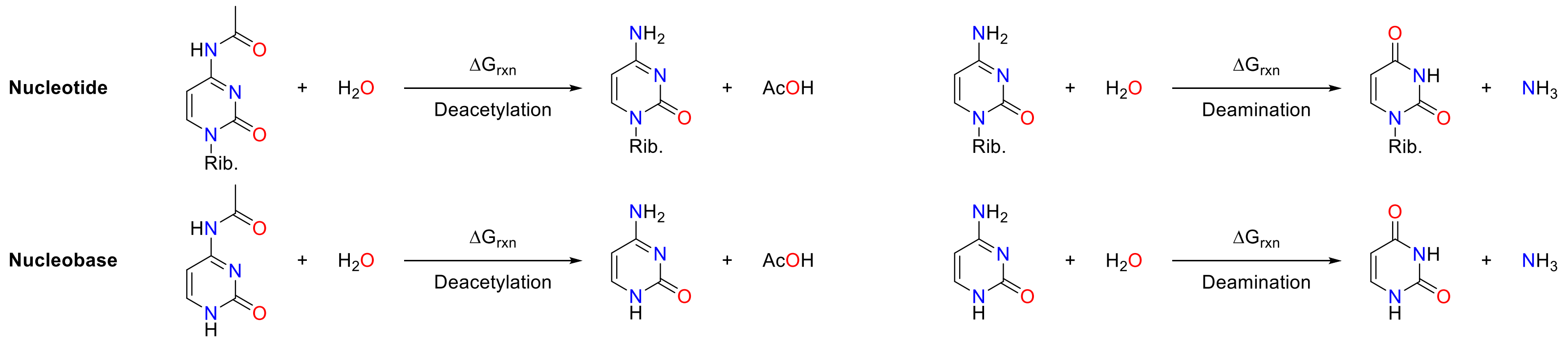

**Figure S69.** Reaction equation for the degradation of *N*4-acetylcytidine (**ac4C**).

**Table S63.** Energetics for the degradation of *N*4-acetylcytidine (**ac4C**). $\Delta E_{Rxn,gas}$, $\Delta G_{Rxn,gas}$, and $\Delta G_{Rxn}$ represent the total energy difference in the gas phase and the Gibbs free difference in both gas and aqueous phase for the deglycosylation reaction.

| Nucleotide | | | $\Delta E_{Rxn,gas}$ | $\Delta G_{Rxn,gas}$ | $\Delta G_{Rxn}$ |
|---|---|---|---|---|---|
| **Reactant** | **Product** | **Reaction Type** | (kJ/mol) | (kJ/mol) | (kJ/mol) |
| **ac4C** | **C** | Deacetylation | -7.4 | -14.8 | -10.2 |
| **C** | **U** | Deamination | -11.3 | -12.2 | 4.5 |
| **Nucleobase** | | | $\Delta E_{Rxn,gas}$ | $\Delta G_{Rxn,gas}$ | $\Delta G_{Rxn}$ |
| **Reactant** | **Product** | **Reaction Type** | (kJ/mol) | (kJ/mol) | (kJ/mol) |
| **ac4C** | **C** | Deacetylation | -6.2 | -11.7 | -8.0 |
| **C** | **U** | Deamination | -39.2 | -36.6 | 3.7 |
